# Variational inference in coupled models of amino acid substitution

**DOI:** 10.64898/2026.05.16.725674

**Authors:** Annabel Large, Ian Holmes

## Abstract

We investigate the use of Expectation–Maximization (em) and variational Bayes for inferring rates and interactions under models of molecular coevolution. We first review em theory for continuous-time Markov chains (ctmcs) and develop it for coevolutionary models, exploiting exchangeability and reversibility symmetries to constrain the parameter dimension. We fit several paired amino-acid coevolutionary models to structural alignments and compare the results to previous work. Our richest model trained on pooled coevolutionary data has explanatory power comparable to CherryML’s Q2 matrix (also trained on pooled data), with one quarter the parameters. However, we observe that aggregation of training data can lead to a form of Simpson’s Paradox: a mixture model, whose components are parameter-efficient continuous-time Bayes networks (ctbns), resolves signals that wash out when a single model tries to capture everything. These signals include both correlated and anticorrelated hydropathy and volume-packing in coevolving amino-acid pairs, as well as the anticorrelated acid/base compensation that was detected by CherryML’s Q2. We next present a closed-form evidence lower bound (ELBO) for ctbns using em statistics. Compared to the state of the art in variational modeling of ctbns, the Euler-Lagrange equations derived by Cohn et al (JMLR, 2010), our closed-form ELBO is competitive in accuracy, considerably more efficient, simpler, and more stable. We conclude by describing a covariant indel model: a Dirichlet process selecting coevolving sites within TKF92, yielding an Infinite Pair HMM over alignments and structures. This model slightly outperforms TKF92 on a structural-alignment benchmark. Code and data are at https://tkfdp.net/.

## 1. Introduction

Protein sequences evolve, to a first approximation, by indels and substitutions. The rates of these events are modulated by selection on the three-dimensional structure, and interacting sites are frequently observed to co-evolve. This selection produces both rate heterogeneity (hydrophobic cores, for instance, tolerate fewer mutations) and epistasis (residues in three-dimensional contact evolve co-ordinately). An integrated model of protein structure and phylogenetics should therefore treat indels, substitutions, rate heterogeneity, and epistasis as first-order phenomena.

In developing coevolutionary models for alignment and phylogenetics, several questions naturally arise. How should such models be constructed parametrically? How can they be fit to data? How can data likelihoods be computed efficiently, and how might the results be incorporated into phylogenetics and alignment algorithms?

Here, we address these questions using the tools of variational modeling. We build on a rich phylogenetics literature on studying non-independent models and their various approximations and bounds^1–12^ In particular, our work extends the application of Expectation Maximization to continuous-time Markov chains (CTMCs),^13–15^ the performance optimization of this technique via time discretization and composite likelihoods,^12^ the use of variational calculus to fit ELBOs to Continuous-Time Bayes Networks (CTBNs),^9^ the use of Dirichlet processes to assign parameters to substitution models,^16^ and the Thorne-Kishino-Felsenstein (TKF) model of indels^17,18^ and its generalizations.^19,20^

Our study is organized by symmetry. We start by noting that the single-site models of molecular evolution are often built on two null hypotheses: in particular, that evolutionary drift runs the same both ways (reversibility), and that sites in a protein are interchangeable (exchangeability). Neither of these null hypotheses have any claim to truth, but it is good practice to break them deliberately rather than accidentally.

We proceed from single sites (Section 2) to residue pairs (Section 3), fitting several models using EM algorithms we describe, and comparing results to prior work including CherryML.^12^ We present a closed-form ELBO for multi-site models, and (in evaluations) find it to be more stable than the Cohn et al. ODE-based ELBO^9^ (Section 5). Finally (Section 6), we describe how the various symmetries we have outlined can be used to add covariation to the TKF92 indel model.^18^

## 2. Single-site models

A single site evolves as a continuous-time Markov chain (CTMC) on the alphabet *A* with generator *Q*, off-diagonal rates *Q*_*xy*_ ≥ 0, *Q*_*xx*_ = − ∑_y≠*x*_ *Q*_*xy*_, and stationary *πQ* = 0. We keep the chain reversible, *π*_*x*_*Q*_*xy*_ = *π*_*y*_*Q*_*yx*_, so the symmetrised matrix 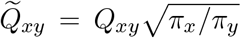 is real with spectral decomposition 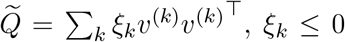. Conjugating back gives the branch transition matrix,

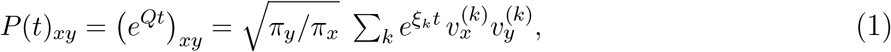

and the tree likelihood is the sum-product recursion of Felsenstein pruning. The generalised time-reversible (GTR) parameterisation writes *Q*_*xy*_ = *S*_*xy*_ *π*_*y*_ with *S* = *S*^⊤^ the exchangeabilities, so 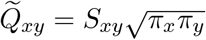 LG08 is one such *S* fixed across families.

### The E-step

The sufficient statistics of a branch trajectory X = (*X*(*u*))_0≤*u*≤*t*_ are the dwell times *W*_*x*_ and transition usages *U*_*xy*_. The complete-data log-likelihood is

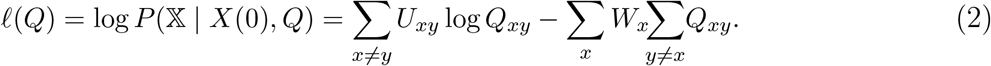

Their endpoint-conditioned expectations are the *bridge statistics*,^13–15^

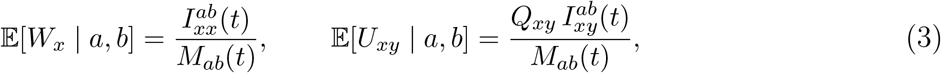

using the matrix exponential *M* (*t*) = *e*^*Qt*^, its time convolution 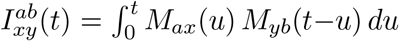, and the exponential divided-difference kernel

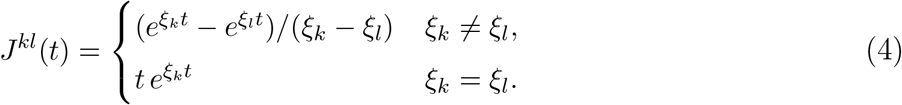

**The M-step** depends on the parameterization. For example, under GTR, letting *V*_*x*_ count the ancestral state (the event *X*(0) = *x*) and 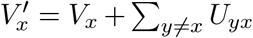, the complete-data log-likelihood is

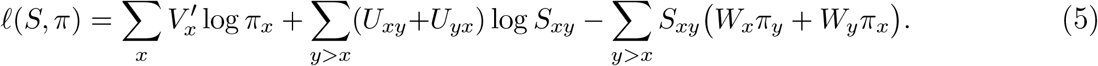

Fixing *π*, the exchangeabilities have the closed-form maximiser 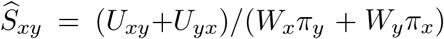 the joint equations for (*S, π*) are nonlinearly coupled with no closed form. There is no canonical optimization for this step, though several strategies may work. For example, one can fix *π* ∝ *V* ^′^ or iterate the two blocks, the *π*-block being strictly concave on the simplex at fixed *S*. Alternatively, a fixed-point *π* update is recovered by a secret-destination augmentation:^21^ attaching to each substitution attempt a hidden destination drawn from *π* makes the augmented counts Dirichlet-conjugate, with mode 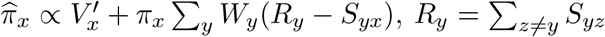, iterated to convergence. It is exact when *S* is off-diagonal-constant (the F81 case, where it reduces to 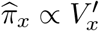) and biased when *S* varies across pairs as strongly as LG08.

## 3. Two-site models

### 3.1. The smallest coupled model: a pair of telegraph processes

The substitution models we consider are illustrated with a simple example. Consider two binary components *x, y* ∈ {0, 1} evolving on the state space 00, 01, 10, 11 (Figure 1). The stationary distribution has four probability parameters; normalization fixes one and exchangeability fixes another, leaving two parameters (corresponding to mean and covariance). As for flux parameters, reversibility leaves six (the Reversible model); exchangeability reduces this to four (Exchangeable). We then consider two different kinds of specialization: the asynchronous models, which forbid simultaneous flips, and the lumpable models, which must use them. The most general asynchronous coupled telegraph has two free rate parameters (Coupled); fixing the proposal (Metropolis) leaves only one rate to determine the time scale. Conversely, lumpability — the property that the chain stays Markovian if projected onto a single component — forces the chain to use simultaneous transitions to model correlations between the components. A non-exchangeably lumpable model is projectable onto the first component only (Half-lumpable); the exchangeable version is projectable onto both components (Lumpable). Comparing the asynchronous models (Coupled, Metropolis) with the synchronized models (Half-lumpable, Lumpable), the asynchronous ones are both favored by the data and significantly easier to train.

**Fig. 1.**
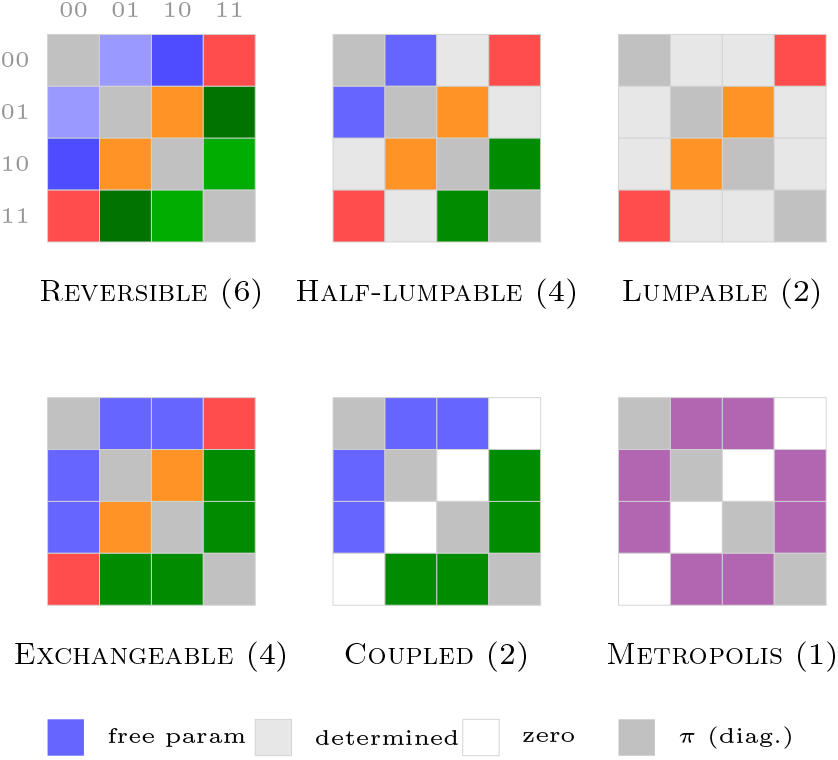
The pair parameterizations we explore in this paper, illustrated on the binary two-site chain. Each 4 × 4 block is the flux tensor over states 00, 01, 10, 11: the diagonal is the stationary *π* and the off-diagonal cells are the six symmetric edge fluxes — four single-site flips and the two simultaneous flips (correlated 00 ↔ 11, swap 01 ↔ 10). A distinct colour marks each free flux parameter (count in parentheses). Reversibility (symmetric flux) holds throughout.

### 3.2. The general pair chain

Two coupled sites evolve as a CTMC on ordered pairs *A*^2^, state *x* = (*i, j*), *y* = (*k, l*). Write the stationary distribution *π*_*ij*_ > 0 and the off-diagonal stationary fluxes *F*_*xy*_ = *π*_*x*_*Q*_*xy*_.

#### The Klein group on the flux tensor

Component exchangeability is *π*_*ij*_ = *π*_*ji*_; reversibility and exchangeability of the fluxes are *F*_*ij,kl*_ = *F*_*kl,ij*_ and *F*_*ij,kl*_ = *F*_*ji,lk*_. Let *R*(*i, j*; *k, l*) = (*k, l*; *i, j*) reverse the transition and *E*(*i, j*; *k, l*) = (*j, i*; *l, k*) swap the two components. The group *G* = {*e, R, E, RE*} ≅ *V*_4_, the Klein four-group, acts on directed off-diagonal transitions, and assigning one nonnegative variable *ϕ*_*o*_ per orbit *o* makes every reversibility and exchangeability equality automatic. Burnside’s lemma counts the orbits, |*O*| = (*A*^4^ + *A*^2^ − 2*A*)/4, so the reversible exchangeable pair generator has that many free flux parameters against the *A*^2^(*A*^2^−1) of an unconstrained one. The symmetric stationary has *A*(*A*+1)/2 − 1 free coordinates.

#### Lumpability versus single-component transitions

We now consider two different specializations of the pair chain. (1) A continuous-time Bayes network (CTBN)^9,22^ forbids simultaneous transitions at the two sites: every jump changes one component, so a compensatory double-substitution must pass through a transiently unfavourable intermediate. (2) A lumpable model, by contrast, is one which can be projected to a single component without losing the Markov property, implying

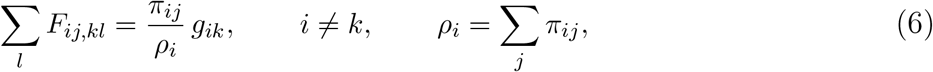

with *g*_*ik*_ = *g*_*ki*_ ≥ 0 the symmetric marginal edge fluxes and *A*_*ik*_ = *g*_*ik*_/*ρ*_*i*_ the marginal generator; exchangeability then gives lumpability of the second coordinate for free. Written compactly, Equation (6) is *Bϕ* = *D*(*π*)*g*, with *B* the sparse incidence map summing orbit fluxes *ϕ* over *l* and *D*(*π*) the diagonal ratio map *π*_*ij*_/*ρ*_*i*_. The two are generically incompatible: single-component transitions cannot also be lumpable under projection to either component. Biologically the CTBN excludes compensatory double-substitutions — the tension familiar from RNA base-pair and codon models, where the data favour simultaneous change.

#### The pair M-step

With transition usages *U*_*xy*_, dwell times *W*_*x*_, and ancestral counts giving 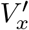 as in (5), the expected complete-data log-likelihood is 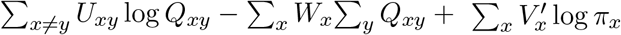, and how it maximises depends on the parameterisation. For the Metropolis family *Q*_*xy*_ = *S*_*xy*_*f* (*π*_*x*_, *π*_*y*_) the exchangeability enters log-linearly, so at fixed *π* it has one closed-form maximiser for every acceptance 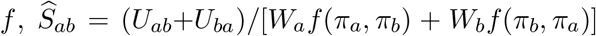 (pooled over both coordinates): the square root 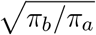, the GTR *π*_*b*_, Barker *π*_*b*_/(*π*_*a*_+*π*_*b*_), Hastings min(1, *π*_*b*_/*π*_*a*_). The stationary enters *f* nonlinearly in all of them (rationally, and non-smoothly for Hastings) with no closed form, so every Metropolis M-step is an expectation–conditional-maximisation alternation: the closed *S*-block above, then a numerical *π*-block (concave on the simplex for GTR, a damped step otherwise). The free reversible flux is the same in Klein-orbit coordinates, 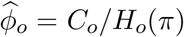 closed at fixed *π* (*C*_*o*_ = ∑_*o*_ *U*_*xy*_, *H*_*o*_(*π*) = ∑_*o*_ *W*_*x*_/*π*_*x*_), *π* numerical. Lumpability is the hard case: its constraint *Bϕ* = *D*(*π*)*g* (6) ties flux and stationary bilinearly, turning even the flux block into a constrained solve with multipliers *λ* (the rational Sinkhorn update *ϕ*_*o*_ = *C*_*o*_/[*H*_*o*_(*π*) + (*B*^⊤^*λ*)_*o*_], rational not exponential because the CTMC complete data is Poisson), and leaves *π* without closed form, so it needs a bilinear/ECM alternation of the constrained flux and *π* steps.

## 4. Comparing the pair couplings

We use the CherryML approach of fitting models to a fixed-size tensor of pairwise substitution counts *n*(*i, j*; *k, l*; *t*) where *t* is a binned time coordinate. The trained pair models are all reversible and exchangeable, so each fits from the coupled count tensor by expectation-maximisation with a common E-step and a model-specific M-step. The E-step is exact: for each time bin it applies the endpoint-conditioned bridge statistics of Equation (3) to the count slice, accumulating expected transition usages *U*_*xy*_ and dwell times *W*_*x*_ over the 400 pair states. The M-step is the model’s own flux update; the stationary *π* (which is symmetric for most models: *π*_*ij*_ = *π*_*ji*_, with *A*(*A*+1)/2 − 1 = 209 free coordinates) is fitted with the flux.

### Coupled amino acid models used in this study

Exchangeable, Coupled, and Lumpable are the twenty-state generalizations of the coupled telegraphs in Figure 1, as described in Section 3: the general chain over all Klein orbits, the single-transition chain, and the single-transition chain constrained to two-sided lumpability, Equation (6). The Metropolis family shares one single-site exchangeability *S* across both coordinates and writes each single-component rate as *Q*_*xy*_ = *S*_*xy*_ *f* (*π*_*x*_, *π*_*y*_) with a reversible acceptance kernel 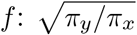, the symmetric kernel; *π*_*y*_, which recovers the single-site GTR form of Section 2 on the changing coordinate; *π*_*y*_/(*π*_*x*_+*π*_*y*_) (Barker); or min(1, *π*_*y*_/*π*_*x*_) (Hastings). All four are reversible with respect to *π*; each re-estimates the shared 190 off-diagonal exchangeabilities of *S* together with the 209 symmetric pair frequencies *π* (399 in all), the fewest of any coupling chain here.

We also fit a mixture of (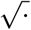 -symmetric) Metropolis kernels. Metropolis is the natural choice for a mixture component due to its economical parameter count. As well as the exchangeable models, we also tried a non-exchangeable variant for which we included each component twice in the mixture (once as itself, once transposed).

All models used a Γ + *I* rate model with four discrete Γ categories plus an invariant class, and a shared branch-length multiplier per family.

### Coupled count corpora

We assembled the coupled count tensor over structural contacts (C*β* distance below 8 Å, primary separation at least 7) from three sources at a common alphabet and time-bin grid. The Pfam-seed corpus draws contacts from experimentally-solved structures mapped by SIFTS; the AlphaFold corpus^23^ extends the same selective contact definition to 15,794 Pfam families whose seed sequences resolve to a confident AlphaFold model (per-residue pLDDT at least 70, low predicted aligned error), drawing its cherries from the shallow Pfam seed alignments; and the trRosetta corpus^24^ pairs the 15,051 structures of that training set with their deep coevolution alignments. Cherry divergences were set by a maximum-likelihood pairwise distance under LG08.

### Held-out comparison

Table 2 scores every model on the per-contact trRosetta corpus of Table 1. In general our symmetrically parameterized Exchangeable slightly outperforms CherryML’s asymmetric Q2 model, despite having a quarter of the parameters and being trained on the same data. Other single-component models (Coupled, the Metropolis variants) form a tight pack; Coupled leads, with its larger parameter count. The Lumpable and Half-lumpable models, forced to choose between dynamics and coupling on a pooled dataset dominated by dynamics, decline to choose correlation and perform poorly. The null models reinforce this: the null with dynamics but no coupling (LG08⊗LG08) significantly outperforms the null with coupling but renewal dynamics (F81).

**Table 1.** Coupled substitution count corpora. Contacts are C*β*<8 Å, separation ≥ 7; “double-sub cells” is the fraction of the 400 × 400 coupled transition matrix, restricted to entries where both residues change, with at least ten counts.

| Corpus | Families | Structures | Cherries | Pair counts | Double-sub cells |
| --- | --- | --- | --- | --- | --- |
| PFAM-seed | 470 | experimental | 47k | 0.63M | 0.9% |
| AlphaFold | 15,794 | predicted | 0.32M | 8.4M | 22% |
| trRosetta | 15,051 | experimental | 4.9M | 341M | 92% |

**Table 2.** Held-out coupled-substitution fits on the trRosetta structural alignment corpus (Section 4). The mutual information statistics are MI_stat_, measuring the coupling in the stationary (*π*), and MI_dyn_, measuring the dynamic coupling in the ancestor-descendant joint *π*_*x*_*P* (*t*)_*xy*_. MI statistics for the Metropolis-mixture rows are the mixture-weighted mean ± s.d. over the interaction classes (other rows carry one value). All models used Γ+I rate variation. The final two columns split the free-parameter count into flux (rate/exchangeability) and probability (the stationary *π*, plus mixture weights where applicable). The two null models are LG08 ⊗ LG08, which discards coupling in favor of dynamics, treating the two sites as independent; and F81-renewal, which preserves coupling but discards dynamics by making every transition resample the joint stationary.

| Model | train LL | val LL | $MI_{\text{stat}}$ | $MI_{\text{dyn}}$ | Flux | Prob. |
| --- | --- | --- | --- | --- | --- | --- |
| METROPOLIS mixture ($K=8$) | -2.5557 | -2.5316 | $0.089 \pm 0.070$ | $0.117 \pm 0.087$ | 190 | 1,679 |
| METROPOLIS mixture ($K=8$, transposed) | -2.5575 | -2.5332 | $0.046 \pm 0.028$ | $0.062 \pm 0.036$ | 190 | 1,599 |
| METROPOLIS mixture ($K=4$) | -2.5761 | -2.5515 | $0.044 \pm 0.019$ | $0.059 \pm 0.024$ | 190 | 839 |
| METROPOLIS mixture ($K=4$, transposed) | -2.5820 | -2.5573 | $0.039 \pm 0.005$ | $0.054 \pm 0.006$ | 190 | 799 |
| EXCHANGEABLE | -2.6169 | -2.5913 | 0.027 | 0.047 | 40,090 | 209 |
| CherryML Q2 (external) | -2.6170 | -2.5914 | 0.018 | 0.033 | 159,999 | — |
| COUPLED | -2.6176 | -2.5919 | 0.027 | 0.039 | 3,800 | 209 |
| METROPOLIS (Barker) | -2.6184 | -2.5928 | 0.029 | 0.036 | 190 | 209 |
| METROPOLIS ($\sqrt{\cdot}$) | -2.6185 | -2.5929 | 0.027 | 0.033 | 190 | 209 |
| METROPOLIS (Hastings) | -2.6192 | -2.5935 | 0.026 | 0.034 | 190 | 209 |
| HALF-LUMPABLE | -2.6248 | -2.5990 | 0.000 | 0.009 | 72,390 | 399 |
| LUMPABLE | -2.6258 | -2.6001 | 0.000 | 0.006 | 32,870 | 209 |
| METROPOLIS (GTR) | -2.6353 | -2.6096 | 0.011 | 0.026 | 190 | 209 |
| LG08$\otimes$LG08 null | -2.6456 | -2.6202 | 0.000 | 0.000 | 190 | 19 |
| F81 renewal from joint | -3.3955 | -3.3682 | 0.027 | 0.454 | — | 209 |

The results show that the coupling available to a single shared 400-state matrix is small: the mutual information of the stationary distribution is capped at ∼ 0.027 nats. This is significantly less than the *per-contact* mutual information, whose mean is 0.16 nats (median 0.13). The reason is that individual site-pairs carry mutual information that is washed out by pooling all sites. This is illustrated by the mixture fit (Figure 2), whose components show a wide range of coupling levels.

**Fig. 2.**
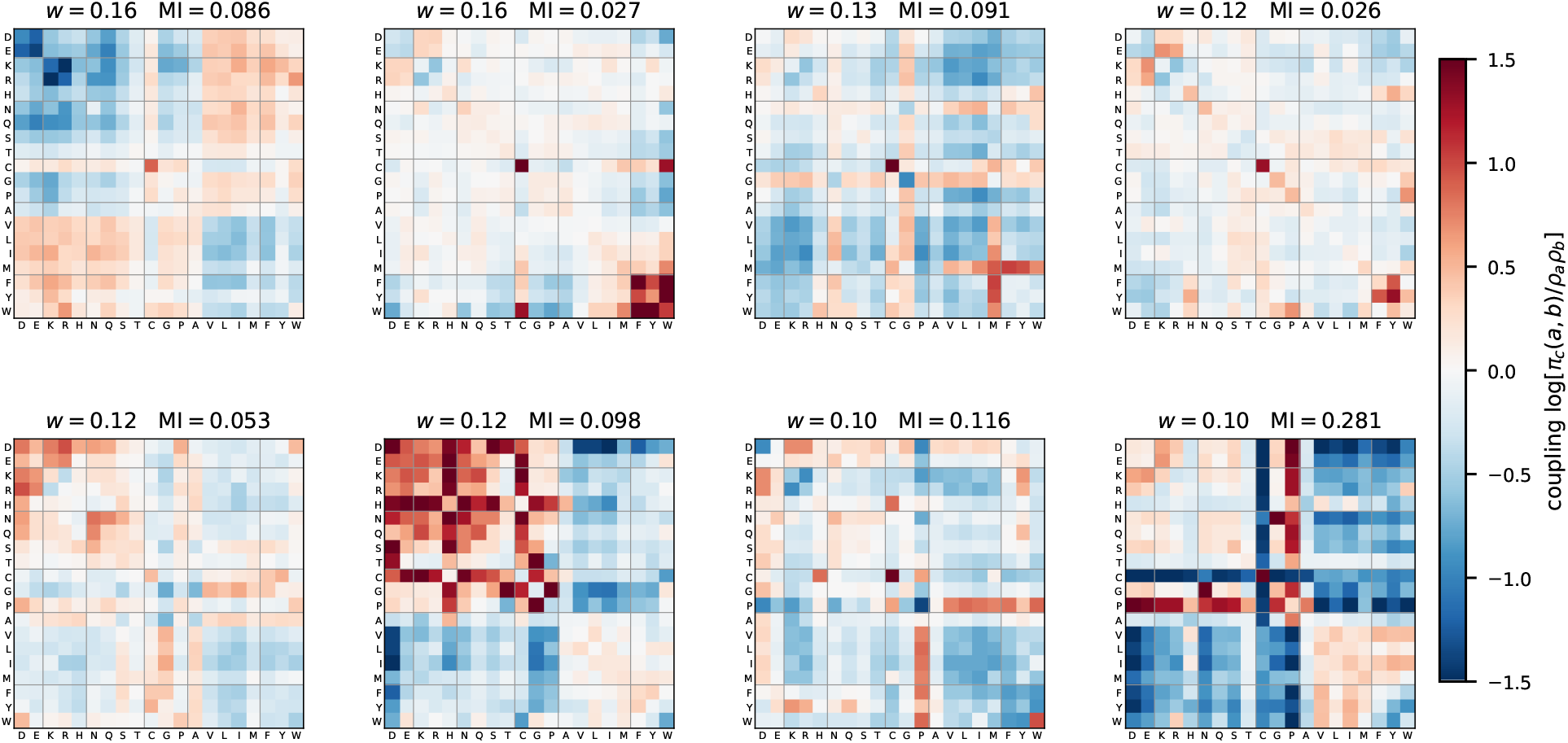
The eight Metropolis-mixture coupling classes (*K*=8). Each panel is one fitted interaction class’s coupling log[*π*_*c*_(*a, b*)/*ρ*_*a*_*ρ*_*b*_] over the 400 amino-acid pairs (red favoured, blue disfavoured; *ρ* the class marginal), with residues ordered by group (acidic | basic | polar | special | aliphatic | aromatic). Panels are sorted by mixture weight *w* and annotated with the stationary mutual information MI(*π*_*c*_). The classes carry coherent but *different* couplings—acid–base complementarity (the off-diagonal acidic/basic blocks) in some, aliphatic assortativity (the `AVLIM` block) in others—so pooling them into one shared chain averages opposite-sign couplings down to the 0.027-nat residual (Table 3); per-class fitting keeps each coupling intact.

We also tried dropping the exchangeability constraint, entering each asymmetric component as a *swap-pair* —its two orientations *π*_*c*_(*a, b*) and *π*_*c*_(*b, a*) tied to one parameter set, weighted equally—to keep the mixture exchangeable (the *transposed* rows of Table 2). At matched budget two symmetric components always beat one doubled-up asymmetric one, so we use symmetric throughout.

**Table 3.**
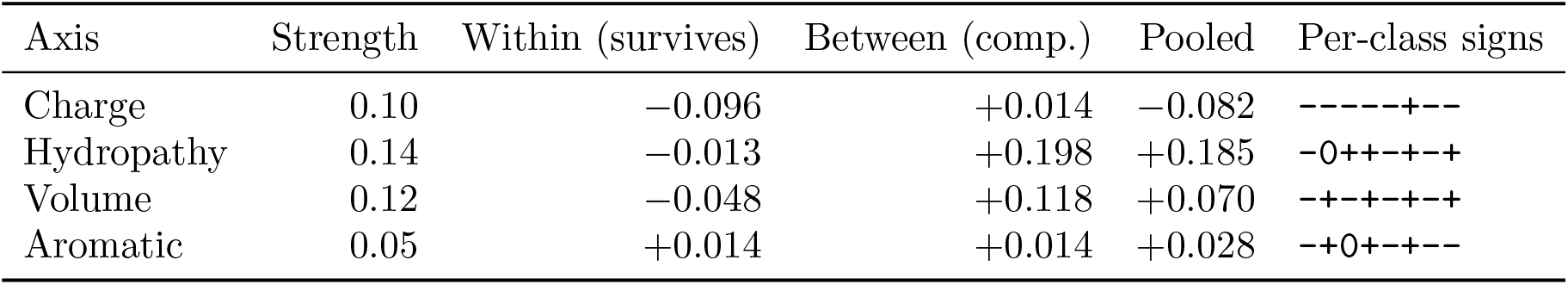
Biophysical coupling in trRosetta per-contact tensors. Each axis’s pooled pair correlation splits exactly as within + between; all values are standardised to the pooled marginal (unit property variance). “Strength” is the mean per-class |Cov_*c*_|; “signs” are the per-class within-contact signs. Charge is a weaker axis than hydropathy or volume, but it is the only one whose within-class coupling survives pooling—because it alone keeps a consistent sign (always complementary), while hydropathy and volume flip sign across classes and cancel.

To understand the washout, we project the fitted per-contact couplings onto four biophysical axes: charge, Kyte–Doolittle hydropathy, residue volume, and aromaticity (see Table 3). Hydropathy and volume are the strongest per-class couplings, but they flip sign across contact classes—assortative in a buried core (hydrophobe with hydrophobe, matched sizes), complementary at a solvent interface or a knob-into-hole packing—so their within-class coupling largely cancels on pooling. Charge is the weaker axis, but the only one with a consistent sign: it is complementary in nearly every class (opposite charges attract, the salt bridge), so its within-class coupling passes through pooling essentially intact. Aromatic stacking is likewise sign-consistent but too weak and narrow to matter.

## 5. Many components and variational inference

The pair models have *A*^2^ states, growing to *A*^*N*^ for *N* coupled sites; exact matrix exponentiation does not simplify for the general CTBN, and Felsenstein’s downward pass breaks the factorisation its upward pass keeps, so approximate inference is the operative regime beyond size two, with a long phylogenetics literature on context-dependent chains, ML/MCMC-EM estimators, and variational or truncation bounds.^1–8,10,11^ We take the variational law to be a product of independent single-site CTMC bridges;^9^ the coupling then contracts in closed form to the bridge statistics of (3), giving a bound linear in the number of contacts rather than exponential in cluster size.

### Path ELBO

Let a cluster of *L* sites carry the Potts target Π(*x*) ∝ *e*^−*E*(*x*)^, *x* ∈ *A*^*L*^, with *E*(*x*) = − ∑_*s*_ *h*_*s*_(*x*_*s*_) − ∑_*s*<*s*_′*H*_*ss*_′ (*x*_*s*_, *x*_*s*_′ ), and let the independent single-site GTR proposal *R*_*ab*_ = *S*_*ab*_*π*_*b*_ act site-wise. The coupling is carried by the *square-root* ^a^ Metropolis target rate for a move *x* → *x*^*s*:*c*^ (*a* = *x*_*s*_ → *c*),

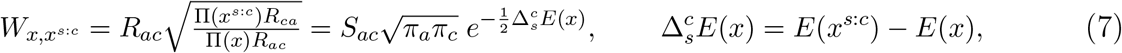

reversible w.r.t. Π for any *S, h, H*. On a branch of length *t* with *x*(0) = *a, x*(*t*) = *b*, using the product of independent *R*-chains as a variational proposal^b^, Girsanov’s formula gives the ELBO

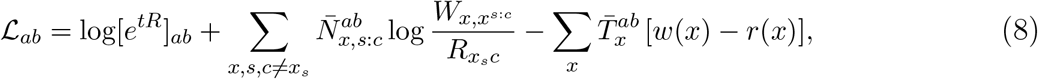

where 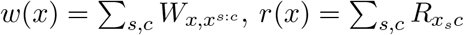, and 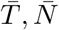 the dwell and usage bridge statistics (3) of the *A*^*L*^-state product chain. The partition function of Π cancels from every rate ratio in (7), so it never enters (8), surviving only in the root factor of the tree likelihood. The log-rate is *affine* in the energy,

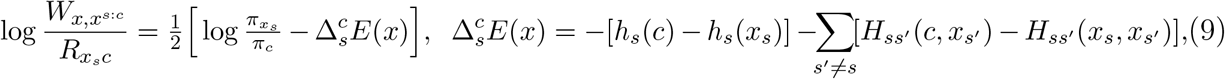

one site-*s* term plus one term per neighbour.

### Contraction to pairwise bridge statistics

The statistics 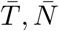 are joint over *A*^*L*^ and do *not* factorise, but (8) contracts them only against the one- and two-body factors of (9), and under the product bridge every such contraction collapses to a one-dimensional integral. For a factor *ϕ*_*C*_ on a site set *C*,

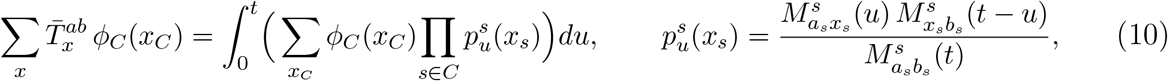

with *M* ^*s*^(*u*) = *e*^*uR*^ and 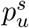 the site-*s* endpoint-conditioned occupancy. Proof: for the product chain 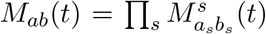 and 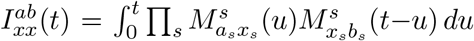, so 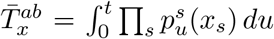 by (3); the *L*−|*C*| off-support sites sum out via 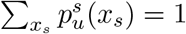 (Chapman–Kolmogorov), leaving (10). The usage 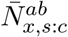 contracts identically with the site-*s* occupancy replaced by its jump density 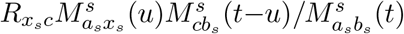. Each coupling *H*_*ss*_′ therefore enters through one two-site integral 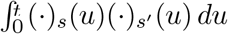 (a pairwise bridge statistic), so the ELBO costs *O m*(*L*+*E*)*A*^2^ for *m* quadrature nodes and *E* contacts.

The closed form is a faithful surrogate for the exact coupled matrix exponential (Figure 3): it is a certified lower bound, near-exact for *t* ≲ 1, and preserves the exact transition-probability ranking (global Spearman ≥ 0.998 across branch lengths and the Metropolis-mixture components) far better than the coupling-free product baseline, most clearly at long branches. The full Cohn mean field, by contrast, has an endpoint singularity in its rate integral and consequently fails to converge on up to 30–40% of pairs at short branches; where it does converge, it violates the bound on ≈ 41% of them; finally, its numerical ODE solver takes ∼ 100× longer to run. The closed form is the more dependable object.

**Fig. 3.**
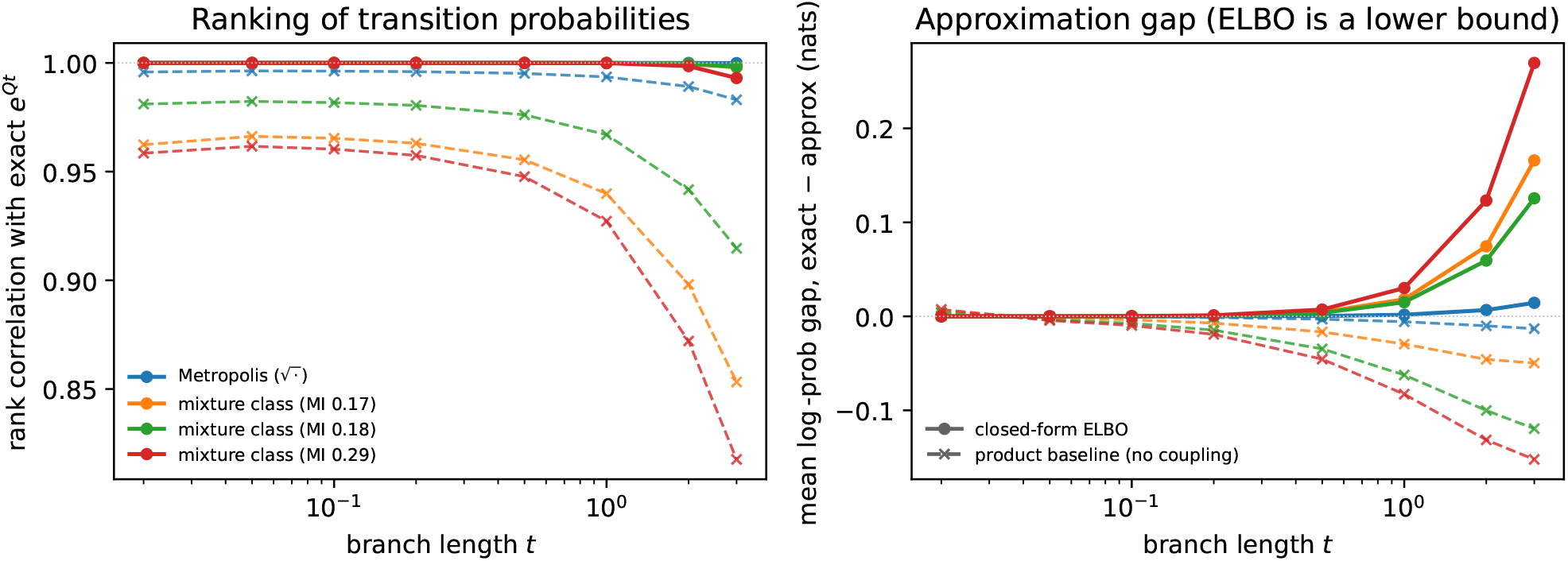
The closed-form ELBO tracks the exact matrix exponential. Path ELBO (8) versus the exact coupled transition matrix [*e*^*tQ*^] over branch length *t*, for the Metropolis chain and the three strongest-coupled Metropolis-mixture components (exact [*e*^*tQ*^] from the reversible eigen-decomposition). *Left:* global Spearman between the ELBO and exact log-transition entries (solid) stays ≈ 1 across *t*, well above the coupling-free product baseline [*e*^*tR*^] (dashed). *Right:* mean gap exact−approx; the ELBO (solid) is a lower bound near zero for *t* ≲ 1, growing only at long branches and strong coupling, while the product baseline (dashed) is not a bound.

Combining the closed-form ELBO above with the product-of-trees variational fit of Jojic et al.^7^ gives a method that jointly fits the Potts coupling and reconstructs ancestral sequences. We implemented this; it works, but on protein families it returns essentially the same reconstructions as independent-site ASR, concordant with prior reports.^25,26^ This matches our other findings in tone: amino-acid coupling is a minor effect in protein phylogenetics. The algorithm, the phylogenetic fixed-bridge Potts ELBO, is given in the supplementary appendix.

## 6. Combining coevolution with indels: the TKF–DP

The coupled substitution model supplies the substitution side of an indel model. Run TKF92 with indel parameters (*λ, µ, r*); deal each surviving residue, at birth, a key class *z*_*s*_ from the stick-breaking weights of a Dirichlet process with concentration *α*_*z*_, so on *L* sites the induced partition is Ewens^27,28^ and residues sharing a table are coupling partners, with coevolution mediated by a “shared field” common to each table. This is the TKF Dirichlet process (TKF– DP): TKF92 with site-independence relaxed via de Finetti.

### Stationarity implies marginal-consistency

For a coupled cluster *C* with joint stationary *π*_*C*_, the augmented substitution-plus-indel process is reversible at a deletion exactly when 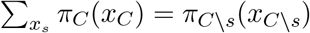 for every *s* ∈ *C*, so a residue joining a table draws from the conditional *π*_*C*_(*x*_*s*_ | *x*_*C*\*s*_) and its later deletion is exact marginalisation.

### Lumpability confers indifference to ghosts

In order to compute cluster likelihoods under the TKF–DP, we need to account for the possibility of transiently inserted lineages temporarily joining the cluster. This requires that the cluster likelihood is lumpable, surviving projection to the observed members. This is the motivation for having the coevolution be driven by a latent time-evolving variable associated with the site; the observed residues merely reflect the state of this variable. This further ensures marginal-consistency, as introduced in the previous paragraph. As a practical accommodation, if we limit clusters to two members, we can use any of the models from Section 4.

### Alignment: the infinite pair HMM

The generalization of the TKF92 Pair HMM to the TKF–DP is an *infinite Pair HMM* ^29^ whose match states track the Ewens partition. We sample the joint posterior over alignments and couplings by MCMC, precomputing partial Forward matrices for efficient resampling of alignments between coupling pins.

On the held-out Laminin EGF-like family (Pfam PF00053) the coupled sampler recovers the canonical disulfide pattern from sequence alone: the eight conserved cysteines generate 28 candidate Cys–Cys column pairs, and all 28 fall within the top 50 column-pair posteriors, with mean Cys–Cys posterior 0.087 against 0.004 background, about 22 times enriched (Figure 4). These are exactly the coevolving, indel-punctuated interactions that a fixed-graph alignment-conditioned Potts model is least equipped to recover, because the couplings and the gaps arrive together. Using the pairwise posteriors for multiple alignment yields improvements over plain TKF92 in sum-of-pairs (SP) and total-column (TC) accuracy (Table 4). The improvement is very slight, perhaps due to the weak coupling signal in pairwise alignments.

**Table 4.**
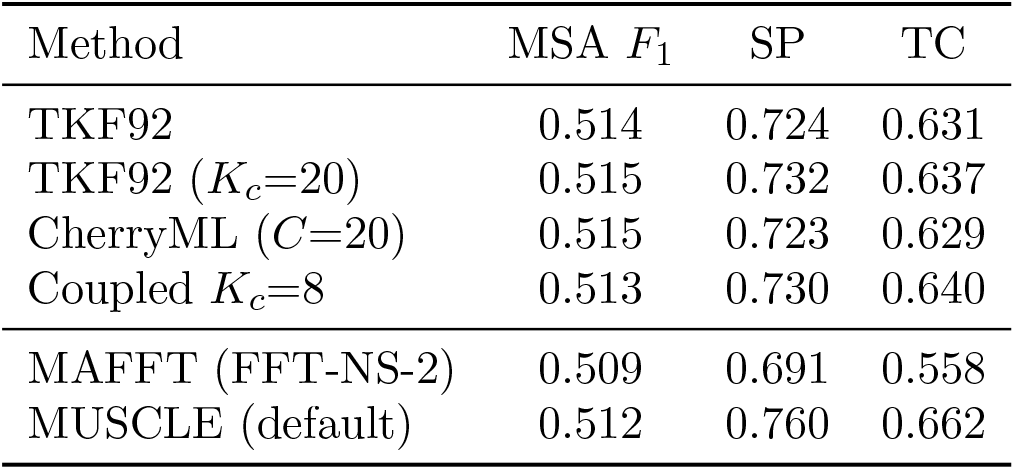
Pairwise-alignment accuracy on BAliBASE `bali3pdbm` (*L*<150 subset, 22 families / 187 sequence pairs). For the pair-HMM models (top block) the MSA is assembled by feeding pairwise match posteriors into FSA;^30^ *F*_1_, SP and TC are standard accuracy metrics. CherryML (*C*=20) is a 20-class site mixture fitted by CherryML.^12^ MAFFT^31^ and MUSCLE^32^ are shown for reference.

**Fig. 4.**
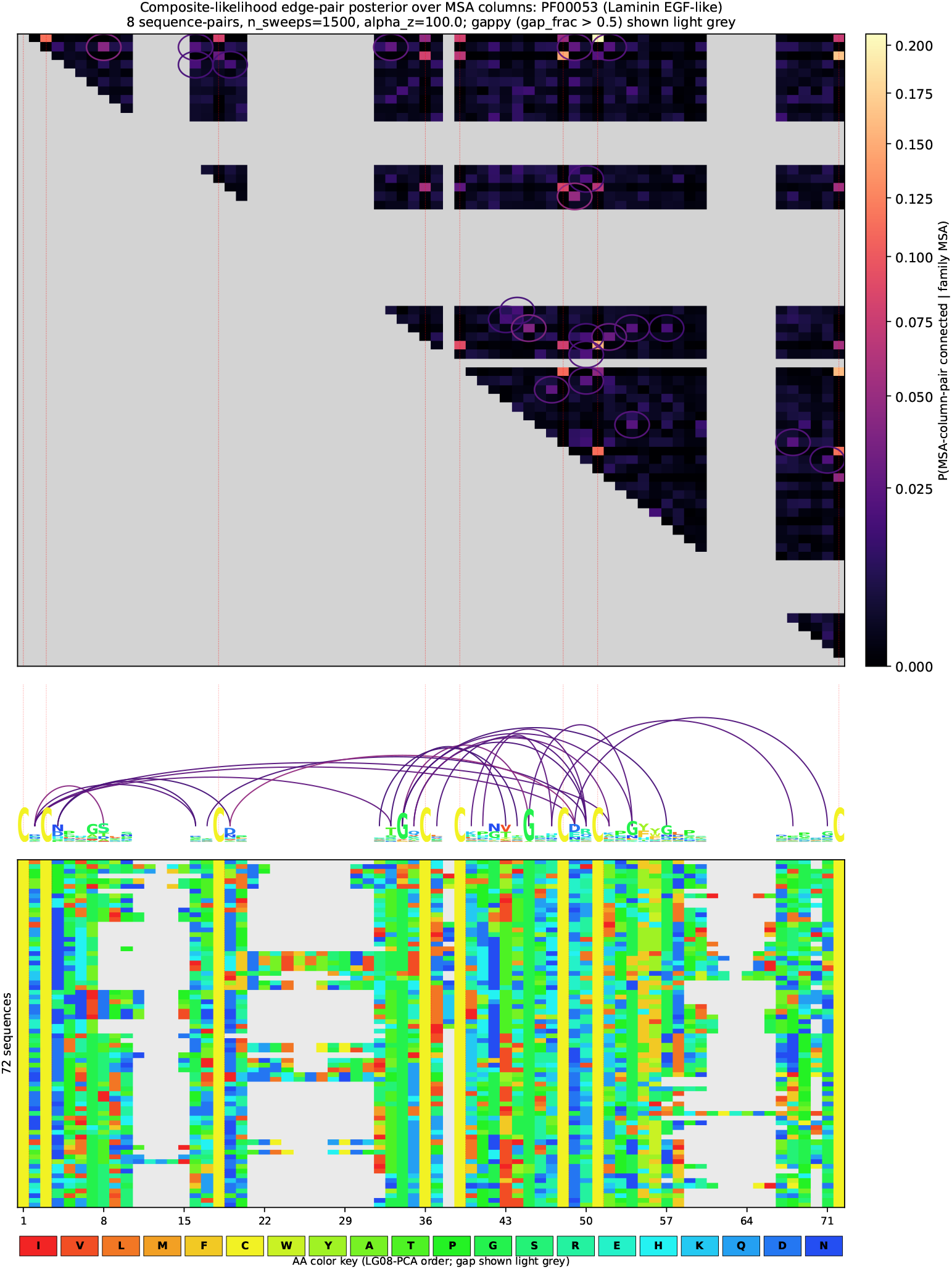
Coupled column-pair posterior on held-out PF00053. All 28 candidate Cys–Cys pairs fall in the top 50 posteriors; the non-Cys top arcs cluster on the cb-EGF Ca^2+^ pocket and conserved *β*-hairpin turns.

## 7. Discussion

Symmetry is a central feature of molecular evolution, and considering the invariants (reversibility, exchangeability) and the equivariants (lumpability) of a coupled substitution process informs the design of a coevolutionary model. Here, we have used it to extend the EM algorithm and the TKF92 model to coevolutionary substitution models, presenting a variational approximation for clusters too large to handle exactly.

We set aside the question of how to model locally-correlated rate heterogeneity, the kind needed for phylogenetic models that partition by domain or secondary structure (hydrophobic versus hydrophilic regions, say). Models introducing such local partitions have been developed elsewhere^20,33,34^ and are natural complements to the ones we present here.

## Acknowledgments

We thank Debora Marks, Yun Song, and Sebastian Prillo for helpful discussions. Selected mathematical derivations and exposition were generated by Anthropic and OpenAI products under the supervision of the human authors. The work was funded by NIH grants HG004483, GM080203, and HG013117. Code and data are at https://tkfdp.net/.

## Footnotes

a The ELBO (10) reduces to pairwise marginals whenever log(*W/R*) is a sum of one- and two-body potentials. This selects the *log-linear* (exponential-family) acceptances: the symmetric square root, the destination-weighted GTR (*f* = *π*_*y*_), and the Glauber form (*f* ∝ *e*^*h*+2*J*^ ).

b As described here, the proposal is fixed; it can be made variational by introducing midpoint-node distributions.

