## Supplemental Appendices A-E for "Variational inference in coupled models of amino acid substitution" (from another version of this preprint: version 1, 2026-05-19)

### Supplementary material for *Insertions, deletions, and exchangeable couplings: a Dirichlet process over TKF92 domains and sites*

Annabel Large and Ian Holmes

Department of Bioengineering, University of California, Berkeley
{annabel_large, ihh}@berkeley.edu

**Abstract**

This supplement collects the long technical derivations referenced by the main paper. It is organised into five appendices that mirror the conceptual ladder of the main text. Appendix A develops the finite-state CTMC and the linear birth–death–immigration process, derives the closed-form bridge-expectation and Fisher-score sufficient statistics for TKF91 and TKF92, resolves the L'Hôpital singularity at $\lambda = \mu$, and discusses the relationship between TKF92 and the latent-boundary-free General Geometric Indel (GGI) model. Appendix B collects the closed-form substitution M-steps for TKF91 / TKF92 / MixDom in their many GTR specialisations, the stochastic-variational Baum–Welch loop with its convergence theorem and linearised analysis at stationarity, Maraschino (the TKF92 cherry-distilled generalisation of CherryML) and its tree-level inference algorithms (FSA, BeamASR, VarAnc, svi-VarAnc), the mixture-of-trees variational ancestral presence/absence inference, and the structural bias of the BP cumulant under a column-factorised variational field. Appendix C develops the recursive TKF family: MixDom, the hierarchical-mixture-of-domains generalisation of TKF92, with its exact closed-form M-step via six-step chain restoration through a fully exploded null-state model; the order-1 Maraschino adjacency distillation; algebraic distillation of MixDom; the MixDom-specific SVI-BW convergence considerations; the tree-level VEM and ancestral-reconstruction algorithms; the generalised phylo-HMM; the labeled-MixDom Singlet and WFST; and the recursive-grammar-elaboration rules together with worked recursive examples (L-TKF, TKFST, TKFStack, TKF-Genome). Appendix D develops the TKF-DP generative model, the class-level path-measure variational likelihood with pairwise bridge expectations, the time-indexed gravestone-augmented pair SCFG, the SVI inference loop, and the pairwise alignment postprocessing landscape. Appendix E develops the infinite Pair HMM as the principled fixed point and the Gibbs+MH+replica-exchange MCMC sampler that draws from it.

B EM, composite likelihoods, and variational inference 37
B.1 Substitution M-Steps for Specific Models 37
B.1.1 JC69 (Jukes–Cantor) 37
B.1.2 K80 (Kimura 2-Parameter) 38
B.1.3 F81 (Felsenstein 1981) 38
B.1.4 HKY85 (Hasegawa–Kishino–Yano) 39
B.1.5 GTR (General Time-Reversible) 39
B.1.6 GY94 (Goldman–Yang Codon Model) 40
B.1.7 Summary and Practical Notes 41
B.1.8 Reversible Mixture with Per-Component GTR 42
B.1.9 Rate Rescaling: M-Step for a Global Scalar Multiplier 44
B.1.10 Joint Rate Rescaling and Equilibrium 45
B.1.11 Tied Equilibria across Class Blocks 48
B.1.12 Tied Equilibria with Per-Class Rate Rescaling 49
B.1.13 Stochastic Variational Baum-Welch 51
B.2 Stochastic Variational Baum–Welch Convergence 52
B.2.1 Setup 52
B.2.2 BDI sufficient statistics 52
B.2.3 Variance of Minibatch Estimates 52
B.2.4 Pseudocount representation and its advantages 53
B.2.5 SVB Convergence Rate 54
B.2.6 Practical Recommendations 55
B.2.7 Bias diagnostics: Hellinger, ESS, and Fisher readouts 56
B.3 Expected Statistics and Linearized Convergence 57
B.3.1 BDI expected statistics at stationarity 57
B.3.2 Per-pair variance of sufficient statistics 58
B.3.3 Relative error and per-pair coefficients of variation 59
B.4 Maraschino: Cherry-Counts for TKF92 60
B.4.1 Optimizing Maraschino: gradient methods vs. inner EM 61
B.4.2 EM-around-Maraschino: mixtures of TKF92 62
B.4.3 EM-around-CherryML: mixtures over site classes 62
B.4.4 Expected sufficient statistics as fast custom VJPs 64
B.5 Selected Inference Algorithms for TKF92 65
B.5.1 Fast Statistical Alignment (FSA) 65
B.5.2 Beam Search Ancestral Sequence Reconstruction (BeamASR) 65
B.5.3 Variational Ancestral Reconstruction (VarAnc) 67
B.5.4 Stochastic Variational VarAnc (svi-VarAnc) 69
B.6 Mixture-of-trees variational ancestral presence/absence 70
B.6.1 Setting and approximation 70
B.6.2 Restricted generative model 71
B.6.3 Singlet HMM at the root 71
B.6.4 Per-branch path log-likelihood 72
B.6.5 Tree-structured variational family 72
B.6.6 Expected log-likelihood under $q$ 73
B.6.7 ELBO 74
B.6.8 Stable computation: cumulant trick 75
B.6.9 Belief propagation for pairwise marginals 75
B.6.10 Special cases and scalability 76

#### Common notation

Several important symbols and their cross-appendix conventions are collected below.

| Symbol | Meaning | First defined |
|---|---|---|
| $\lambda, \mu$ | TKF insertion / deletion rates (per link) | §A |
| $r$ | TKF92 fragment-extension probability | §A |
| $\mathcal{A}$ | residue alphabet ($A = \lvert\mathcal{A}\rvert = 20$ for proteins) | §A |
| $\pi, Q, S$ | GTR stationary, asymmetric, exchangeability matrices | §A |
| $R$ | substitution rate matrix (generator) | §A |
| $n, f, c$ | domain, fragment, and substitution-class indices (MixDom) | §C |
| $\eta_s$ | per-site gamma rate multiplier | §D |
| $z_s, c_s, h_{cc'}$ | key, site-class, Potts-atom indices (TKF-DP) | §D |
| $\alpha_z, \alpha_c, \alpha_H$ | key/class/atom Dirichlet-process concentrations | §D |
| $\boldsymbol{Q}, \boldsymbol{H}$ | generator and Potts coupling tensor | §D |
| $\tau$ | TKF91 Pair HMM transition matrix | §A |
| $\upsilon$ | exploded 7×7 MixDom top-level transitions with null states | §C |
| $\mathcal{T}$ | effective 5×5 MixDom top-level matrix (nulls summed out) | §C |
| $\chi$ | final nested MixDom Pair HMM transition matrix | §C |
| $N_a, N_g$ | alive / gravestone counts (TKF-DP) | §D |
| $F_0, F_1, F_2$ | TKF92 Pair HMM 0/1/2-cell marginal partition functions | §E |
| $\pi_M$ | CRP partition of Match cells (infinite Pair HMM) | §E |

### A BDI and TKF foundations

This appendix builds the foundations on which the rest of the supplement rests. We develop the finite-state continuous-time Markov chain (CTMC) sufficient statistics; the linear birth–death–immigration (BDI) process and its endpoint-conditioned sufficient statistics via the Fisher score identity; the TKF91 model as a factorisation of these; its finite-state machine representations; the TKF92 generalisation; the explicit 6×6 WFST derivation by Pair-HMM-over-Singlet division; the L'Hôpital limits at $\lambda{=}\mu$ that close the 0/0 singularity in the score identity; and a discussion of how the latent-boundary TKF92 model relates to the latent-boundary-free General Geometric Indel (GGI) model via moment matching.

#### A.1 The TKF91 Model

##### A.1.1 Finite-State Continuous-Time Markov Chain (CTMC)

A finite-state continuous-time Markov chain (CTMC) on alphabet $\mathcal{A}$ has rate matrix (generator) $R$ with off-diagonal entries $R_{ij} \geq 0$ (instantaneous rate of substitution from $i$ to $j$) and diagonal entries $R_{ii} = -\sum_{j \neq i} R_{ij}$. The stationary distribution $\pi$ satisfies $\pi R = 0$ and $\sum_i \pi_i = 1$.

We restrict our analysis to time-reversible CTMCs, which satisfy detailed balance $\pi_i R_{ij} = \pi_j R_{ji}$ for all $i, j$. The symmetrized rate matrix $S_{ij} = R_{ij}\sqrt{\pi_i/\pi_j}$ is real symmetric, so it has a spectral decomposition $S = \sum_k \xi^{(k)} v^{(k)} v^{(k)\top}$ with real eigenvalues $\xi^{(k)} \leq 0$ and orthonormal eigenvectors $v^{(k)}$. Conjugating back, the transition matrix is

$$P(X(T) = j \mid X(0) = i, R) = \left(e^{RT}\right)_{ij} = \sum_k e^{\xi^{(k)}T} \sqrt{\frac{\pi_j}{\pi_i}}\, v_i^{(k)} v_j^{(k)}$$

##### A.1.2 Sufficient Statistics for Finite-State CTMCs

The sufficient statistics for a continuously-observed path $\mathbb{X} = \{X(t)\}_{0 \le t \le T}$ in the finite-state CTMC are the dwell times $W_i$ and transition counts $U_{ij}$. The complete-data log-likelihood is

$$\log P(\mathbb{X} \mid X(0), R) = \sum_{i \neq j} U_{ij} \log R_{ij} + \sum_i W_i R_{ii} = \sum_{i \neq j} U_{ij} \log R_{ij} - \sum_i W_i \sum_{j \neq i} R_{ij}. \tag{A.1}$$

This is an exponential-family form with natural parameters $\log R_{ij}$ and $-R_{ii}$, and sufficient statistics $U_{ij}$ and $W_i$.

The *a posteriori* expectations of these sufficient statistics for an endpoint-conditioned path in the finite-state CTMC are (19, 20, 25, 47)

$$\mathcal{C}_i^W(a,b,T) \equiv \mathbb{E}[W_i | a,b] = \frac{1}{M_{ab}(T)} I_{ii}^{ab}(T) \tag{A.2}$$

$$\mathcal{C}_{ij}^U(a,b,T) \equiv \mathbb{E}[U_{ij} | a,b] = \frac{R_{ij}}{M_{ab}(T)} I_{ij}^{ab}(T) \tag{A.3}$$

where $M(T) = e^{RT}$ is the matrix exponential and

$$I_{ij}^{ab}(T) = \int_0^T M_{ai}(t)\, M_{jb}(T-t)\, dt. \tag{A.4}$$

Writing $M_{ij}(t)$ in the eigenbasis of the symmetrized rate matrix $S_{ij} = R_{ij}\sqrt{\pi_i/\pi_j}$ (with eigenvalues $\xi^{(k)}$ and orthonormal eigenvectors $v^{(k)}$):

$$I_{ij}^{ab}(T) = \sqrt{\frac{\pi_i \pi_b}{\pi_a \pi_j}} \sum_k v_a^{(k)} v_i^{(k)} \sum_l v_j^{(l)} v_b^{(l)}\, J^{kl}(T) \tag{A.5}$$

where $J^{kl}(T) = T e^{\xi^{(k)} T}$ if $\xi^{(k)} = \xi^{(l)}$, and $(e^{\xi^{(k)} T} - e^{\xi^{(l)} T})/(\xi^{(k)} - \xi^{(l)})$ otherwise.

Using the GTR parameterization $R_{ij} = Q_{ij}\pi_j$, where $Q$ is a symmetric exchangeability matrix (so $S_{ij} = Q_{ij}\sqrt{\pi_i \pi_j}$), and including the prior $P(X(0) = i) = \pi_i$, the complete data log-likelihood is

$$\log P(\mathbb{X}, X(0) \mid Q, \pi) = \sum_i \left( V_i + \sum_{j \neq i} U_{ji} \right) \log \pi_i + \sum_{j > i} (U_{ij} + U_{ji}) \log Q_{ij} - \sum_{j > i} Q_{ij} \left( W_i \pi_j + W_j \pi_i \right)$$

where $V_i$ is the indicator/count for the initial state $X(0) = i$.

##### A.1.3 Linear Birth-Death-Immigration Process (BDI)

The linear birth-death-immigration (BDI) process (29) has per-capita birth rate $\lambda$, per-capita death rate $\mu$, and constant immigration rate $\nu$, so that the total event rates from state $k$ are $\lambda_k = k\lambda + \nu$ and $\mu_k = k\mu$.

The transition distribution for this process is well-studied in the general case. We confine our analysis to the specific case $\nu = \lambda, \mu > \lambda$ where the immigration rate is equal to the birth rate and the process is positive recurrent and has an equilibrium distribution, a zero-based geometric with parameter $\lambda/\mu$. This is the regime relevant to the TKF91 and related models. However, many of the formulas we derive will hold more generally, and the methods we use can be applied to other parameter regimes as well.

Let $Y_f$ denote the number of founder deaths and $Y_o$ the number of youngest orphans (i.e. the number of times a lineage goes extinct while leaving behind at least one surviving descendant). Then the finite-time transition probability is obtained by summing the joint probability

$$P(X(T) = j,\, Y_f = m,\, Y_o = n \mid X(0) = i) = \binom{i}{m}\binom{m}{n}\binom{j}{i-m+n} \\ \times\ \alpha^{i-m}(1-\alpha)^m\, \beta^{j-i+m-n}(1-\beta)^{i+1-m+n}\, \gamma^n(1-\gamma)^{m-n}$$

over all $m, n$ consistent with $i$ and $j$ ($0 \le m \le i$ and $0 \le n \le \min(m, j-i+m)$), where

$$\alpha(\lambda,\mu,T) = \exp(-\mu T) \tag{A.6}$$

$$\beta(\lambda,\mu,T) = \frac{\lambda \exp(-\lambda T) - \lambda\alpha}{\mu \exp(-\lambda T) - \lambda\alpha} \tag{A.7}$$

$$\tag{A.8}$$

$$\gamma(\lambda,\mu,T) = 1 - \frac{\mu\beta}{\lambda(1-\alpha)}. \tag{A.9}$$

##### A.1.4 Sufficient Statistics for Linear BDI Process

The sufficient statistics for an augmented continuously-observed path $\{X(t)\}_{0\le t\le T}$ in the linear BDI process wherein immigration events are labeled as such (i.e. we observe the full path and know which births are due to immigration and which are due to reproduction) are the individual-birth count $B_{\text{ind}}$, the immigration count $B_{\text{imm}}$ (with $B = B_{\text{ind}} + B_{\text{imm}}$ the total), the death count $D$, and the time-integrated population size $S = \int_0^T X(t)\,dt$. The complete-data log-likelihood is

$$\log P\big(\mathbb{X} \mid X(0), \lambda, \mu, \nu\big) = B_{\text{ind}} \log\lambda + B_{\text{imm}} \log\nu + D\log\mu - (\lambda+\mu)\,S - \nu T + c \tag{A.10}$$

where $c$ does not depend on $(\lambda, \mu, \nu)$. The natural parameters are $(\log\lambda, \log\mu, \log\nu, -(\lambda+\mu))$ and the sufficient statistics $(B_{\text{ind}}, D, B_{\text{imm}}, S)$. This is a curved exponential-family form because the natural parameters are not independent.

**Score function identity.** Let $P_{ij}(T;\theta)$ denote the observed likelihood, i.e. the transition probability $P(X(T) = j \mid X(0) = i;\, \theta)$, with $\theta = (\lambda, \mu, \nu)$. The *score function identity* states that the gradient of the observed log-likelihood equals the posterior expectation of the complete-data score:

$$\nabla_\theta \log P_{ij}(T;\theta) = \mathbb{E}[\nabla_\theta \log P(\mathbb{X} \mid X(0), \theta) \mid X(0) = i,\ X(T) = j;\ \theta]\,. \tag{A.11}$$

No approximation is involved; (A.11) holds for any parametric family and any choice of complete/incomplete data.

Specializing to $\nu = \lambda$ and writing $B = B_{\text{ind}} + B_{\text{imm}}$ for the total birth count, the complete-data log-likelihood (A.10) becomes $\log P = B\log\lambda + D\log\mu - (\lambda+\mu)\,S - \lambda T + c$ with two free parameters $(\lambda, \mu)$. Differentiating and applying (A.11):

$$\frac{\partial}{\partial\lambda}\log P_{ij}(T;\theta) = \frac{\mathbb{E}[B \mid i,j,T]}{\lambda} - \mathbb{E}[S \mid i,j,T] - T$$

$$\frac{\partial}{\partial\mu}\log P_{ij}(T;\theta) = \frac{\mathbb{E}[D \mid i,j,T]}{\mu} - \mathbb{E}[S \mid i,j,T]$$

Each equation involves two unknowns because the natural parameters $(\log\lambda, \log\mu, \lambda+\mu)$ are not independent. The conservation law $i + B - D = j$ provides the missing equation:

$$\mathbb{E}[B \mid i,j] - \mathbb{E}[D \mid i,j] = j - i \tag{A.12}$$

Solving this system (using $\lambda \neq \mu$):

$$\begin{aligned}
\mathbb{E}[S \mid i,j] &= \frac{j - i + \mu\frac{\partial}{\partial\mu}\log P_{ij} - \lambda\frac{\partial}{\partial\lambda}\log P_{ij} - \lambda T}{\lambda - \mu} \\
\mathbb{E}[B \mid i,j] &= \lambda\frac{\partial}{\partial\lambda}\log P_{ij} + \lambda\,\mathbb{E}[S \mid i,j] + \lambda T \\
\mathbb{E}[D \mid i,j] &= \mu\frac{\partial}{\partial\mu}\log P_{ij} + \mu\,\mathbb{E}[S \mid i,j]
\end{aligned}$$

Inclusion of an initial distribution $P(X(0) = L) = \kappa^L(1-\kappa)$ brings the complete-data log-likelihood into the form

$$\log P\big(\mathbb{X}, X(0) \mid \lambda, \mu\big) = B\log\lambda + D\log\mu + L\log\kappa + M\log(1-\kappa) - (\lambda+\mu)\,S - \lambda T + c$$

where $M = 1$ represents the number of BDI trajectories observed, introduced for later generalization.

##### A.1.5 The TKF91 Model: Linear BDI + Finite-State CTMC

Thorne, Kishino, and Felsenstein's 1991 model (TKF91) was introduced to describe the molecular evolution of a biological sequence (DNA, RNA, or protein) subject to insertions, deletions, and point substitutions (50).

The parameters $\theta = (\lambda, \mu, Q, \pi)$ consist of the insertion rate $\lambda$, deletion rate $\mu$, exchangeability matrix $Q$, and equilibrium distribution $\pi$.

The BDI part of the model, known as the *links model*, describes the evolution of an "immortal link" followed by zero or more "mortal links". Each link evolves according to an independent linear BDI process; births and immigrations correspond to insertions, which generate new mortal links, and deaths correspond to deletions.

In the TKF91 model, each mortal link is associated with an observed character that evolves according to an independent finite-state CTMC with rate matrix $Q$ and stationary distribution $\pi$. Inserted characters are drawn from the equilibrium distribution $\pi$.

In later TKF91-derived models, a more elaborate structure may be associated with each link; for example, in the TKF92 model, a fragment of characters of geometrically-distributed length (51).

To distinguish between these models we introduce the notation $\mathrm{Links}(M; \lambda, \mu)$ for the links model where an independent stationary stochastic process $x(t) \sim M$ is associated with every mortal link. This defines a stationary stochastic process over sequences $\mathcal{S}(t) \sim \mathrm{Links}(M; \lambda, \mu)$ where $\mathcal{S}$ is a sequence over the alphabet defined by the state space of $M$.

Let $\mathrm{Subst}(Q, \pi)$ denote the time-reversible continuous-time Markov chain with stationary distribution $\pi$ and generator $R = Q\mathrm{diag}(\pi)$, where $Q$ is the exchangeability matrix. Then the TKF91 model is just the links model with a point substitution process transpiring at each mortal link

$$\mathcal{S}(T) \sim \mathrm{Links}(\mathrm{Subst}(Q, \pi); \lambda, \mu)$$

##### A.1.6 Finite State Machines of the TKF91 Model

There are three distinct state machines associated with nodes and branches in a tree of TKF-related sequences.

**TKF91 Stationary HMM** The stationary distribution of the TKF91 model can be represented as a Hidden Markov Model (HMM) that emits $n \sim \mathrm{Geometric}(\kappa)$ characters drawn i.i.d. from $\pi$.

**TKF91 Transducer** The conditional distribution of a descendant sequence $\mathcal{S}(T)$ given ancestor sequence $\mathcal{S}(0)$ can be represented as a Weighted Finite-State Transducer (WFST) that consumes an ancestor sequence and emits a descendant sequence, with transition weights determined by the TKF91 parameters

$$\mathbb{T}(\lambda,\mu,T) = \left(\begin{array}{c|ccccc} & \mathtt{S} & \mathtt{M} & \mathtt{I} & \mathtt{D} & \mathtt{E} \\ \hline \mathtt{S} & 0 & (1-\beta)\alpha & \beta & (1-\beta)(1-\alpha) & 1-\beta \\ \mathtt{M} & 0 & (1-\beta)\alpha & \beta & (1-\beta)(1-\alpha) & 1-\beta \\ \mathtt{I} & 0 & (1-\beta)\alpha & \beta & (1-\beta)(1-\alpha) & 1-\beta \\ \mathtt{D} & 0 & (1-\gamma)\alpha & \gamma & (1-\gamma)(1-\alpha) & 1-\gamma \\ \mathtt{E} & 0 & 0 & 0 & 0 & 0 \end{array}\right)$$

with M aligning pairs of characters $(i,j)$ with probability $\exp(RT)_{ij}$, I emitting unaligned descendant characters $(\epsilon,i)$ with $i\sim\pi$, and D consuming unaligned ancestral characters $(i,\epsilon)$ without any additional weight penalty.

This transducer can be interpreted as a decomposition of each mortal-link BDI trajectory into founder survival/death, possible survival of the youngest orphan (which, by insertion order, is the leftmost surviving descendant when the founder dies), and a geometric tail of further offspring. This yields the parameters $\alpha,\beta,\gamma$ appearing in the WFST transition matrix and determines which transition counts correspond to birth and death events.

**TKF91 Pair HMM** The joint distribution of ancestor and descendant is given by the transducer composition product of the stationary HMM and the transition WFST. This product can itself be represented as a Pair HMM with states $\{\mathtt{S},\mathtt{M},\mathtt{I},\mathtt{D},\mathtt{E}\}$ that models the joint distribution, as opposed to the conditional distribution represented by the WFST. The transition matrix $\tau$ of the Pair HMM is similar to the transition matrix $\mathbb{T}$ of the transducer, with the differences that incoming transitions to M and D states acquire a factor of $\kappa$ for the extension of the ancestral sequence and transitions to E states acquire a factor of $1-\kappa$ for the termination of the ancestral sequence.

For concreteness, because we will refer to it frequently, the transition matrix of the TKF91 Pair HMM is

$$\tau(\lambda,\mu,T) = \left(\begin{array}{c|ccccc} & \mathtt{S} & \mathtt{M} & \mathtt{I} & \mathtt{D} & \mathtt{E} \\ \hline \mathtt{S} & 0 & (1-\beta)\alpha\kappa & \beta\kappa & (1-\beta)(1-\alpha)\kappa & (1-\beta)(1-\kappa) \\ \mathtt{M} & 0 & (1-\beta)\alpha\kappa & \beta\kappa & (1-\beta)(1-\alpha)\kappa & (1-\beta)(1-\kappa) \\ \mathtt{I} & 0 & (1-\beta)\alpha\kappa & \beta\kappa & (1-\beta)(1-\alpha)\kappa & (1-\beta)(1-\kappa) \\ \mathtt{D} & 0 & (1-\gamma)\alpha\kappa & \gamma\kappa & (1-\gamma)(1-\alpha)\kappa & (1-\gamma)(1-\kappa) \\ \mathtt{E} & 0 & 0 & 0 & 0 & 0 \end{array}\right)$$

The emission weights of the Pair HMM are similarly related to the input/output emission weights of the WFST, with M- and I-state emissions acquiring a factor of $\pi_i$ for the ancestral character $i$.

For the indel score identities associated with birth, death, and exposure statistics, it is convenient to work with the conditionally normalized TKF91 WFST representing $P(y|x,\theta)$, since its transition weights correspond directly to the finite-time BDI factors $\alpha,\beta,\gamma$. However, the M-step for $(\lambda,\mu)$ under the stationary TKF91 model must also include the prior contribution of the ancestral sequence length, which belongs to the joint distribution $P(x,y|\theta) = P(x|\theta)P(y|x,\theta)$. Accordingly, we use the conditional WFST to obtain posterior expectations of $B,D,S$ and then augment the M-step objective with the joint-model terms involving $L$ and $M$.

Appendix A.4 considers the limits and implications of these formulas in the biologically-relevant regime of long sequences, $\lambda\to\mu$.

##### A.1.7 Sufficient Statistics for the TKF91 Model

For observed ancestral and descendant sequences $x, y$ under TKF91 with parameters $\theta = (\lambda, \mu, Q, \pi)$, and a particular hidden Pair HMM/WFST path $z$, the $(\lambda, \mu)$-dependent part of the complete-data log-likelihood is linear in the transition and emission counts,

$$\log P\big(y, z \mid x, \theta\big) = \sum_{i,j} n_{ij}(z) \, \log \mathbb{T}_{ij}(\lambda, \mu, t) \; + \; \sum_{b} e^{\mathtt{I}}_{(\epsilon,b)}(z) \, \log \pi_b \; + \; \sum_{a,b} e^{\mathtt{M}}_{(a,b)}(z) \, \log \exp(Rt)_{ab}.$$

Let

$$\begin{aligned} \hat{n}_{ij} &= \mathbb{E}[n_{ij} \mid x, y; \theta] \\ \hat{e}^{\mathtt{I}}_{(\epsilon,b)} &= \mathbb{E}[e^{\mathtt{I}}_{(\epsilon,b)} \mid x, y; \theta] \\ \hat{e}^{\mathtt{M}}_{(a,b)} &= \mathbb{E}[e^{\mathtt{M}}_{(a,b)} \mid x, y; \theta] \end{aligned}$$

denote the posterior expectations of these counts returned by the Forward-Backward algorithm. By the score function identity,

$$\begin{aligned} \lambda \frac{\partial}{\partial \lambda} \log P\big(y \mid x, \theta\big) &= \sum_{i,j} \hat{n}_{ij} \, \mathcal{C}^{\lambda}_{ij}(\lambda, \mu, t) \\ \mu \frac{\partial}{\partial \mu} \log P\big(y \mid x, \theta\big) &= \sum_{i,j} \hat{n}_{ij} \, \mathcal{C}^{\mu}_{ij}(\lambda, \mu, t) \end{aligned}$$

where $\mathcal{C}^{\xi}_{ij}(\lambda, \mu, t) = \xi \frac{\partial}{\partial \xi} \log \mathbb{T}_{ij}(\lambda, \mu, t)$ for $\xi \in \{\lambda, \mu\}$. (See Sections A.4.2 and A.4.3 for explicit formulae.) The conservation law is likewise linear in the same counts:

$$\mathbb{E}[B \mid x, y] - \mathbb{E}[D \mid x, y] = \hat{n}_{\mathtt{SI}} + \hat{n}_{\mathtt{MI}} + \hat{n}_{\mathtt{II}} - \hat{n}_{\mathtt{SD}} - \hat{n}_{\mathtt{MD}} - \hat{n}_{\mathtt{DD}} - \hat{n}_{\mathtt{ID}} \qquad \text{(A.13)}$$

$$= \sum_{i,j} \hat{n}_{ij} \, \mathcal{C}^{\text{cons}}_{ij} \qquad \text{(A.14)}$$

where $\mathcal{C}^{\text{cons}}_{ij} = \delta(j = \mathtt{I}) - \delta(j = \mathtt{D})$ counts whether the WFST transition $i \to j$ corresponds to a birth, death, or neither. Substituting into the BDI score identities gives

$$\mathbb{E}[S \mid x, y] = \frac{\sum_{i,j} \hat{n}_{ij} \left( \mathcal{C}^{\text{cons}}_{ij} - \mathcal{C}^{\lambda}_{ij} + \mathcal{C}^{\mu}_{ij} \right) - \lambda T}{\lambda - \mu} \qquad \text{(A.15)}$$

$$\mathbb{E}[B \mid x, y] = \lambda \sum_{i,j} \hat{n}_{ij} \, \mathcal{C}^{\lambda}_{ij}(\lambda, \mu, t) + \lambda \, \mathbb{E}[S \mid x, y] + \lambda T \qquad \text{(A.16)}$$

$$\mathbb{E}[D \mid x, y] = \mu \sum_{i,j} \hat{n}_{ij} \, \mathcal{C}^{\mu}_{ij}(\lambda, \mu, t) + \mu \, \mathbb{E}[S \mid x, y] \qquad \text{(A.17)}$$

Thus the posterior expectations of the BDI sufficient statistics $\mathcal{C}^B(\mathbf{n}, T) = \mathbb{E}[B|x, y]$, $\mathcal{C}^D(\mathbf{n}, T) = \mathbb{E}[D|x, y]$, and $\mathcal{C}^S(\mathbf{n}, T) = \mathbb{E}[S|x, y]$ are affine functions of the Forward-Backward expected transition counts $\hat{n}_{ij}$.

The TKF91 CTMC sufficient statistics take the form

$$\begin{aligned} \mathbb{E}[W_i \mid x, y] &= \sum_{a,b} \hat{e}^{\mathtt{M}}_{(a,b)} \, \mathcal{C}^{W}_{i}(a, b, T) \; + \mathcal{G}^{W}_{i}(x, y) \\ \mathbb{E}[U_{ij} \mid x, y] &= \sum_{a,b} \hat{e}^{\mathtt{M}}_{(a,b)} \, \mathcal{C}^{U}_{ij}(a, b, T) \; + \mathcal{G}^{U}_{ij}(x, y) \end{aligned}$$

where the first term is the contribution from observed endpoint-matched lineages, obtained from CTMC bridge expectations conditional on $a \to b$ over time $T$. The correction terms $\mathcal{G}_i^W, \mathcal{G}_{ij}^U$ account for unobserved partial CTMC trajectories on lineages that are deleted before time $T$, inserted after time 0, or born and deleted within $(0, T)$. Exact evaluation of these terms would require genealogical information about the latent BDI history, including birth and death times and lineage lifetimes. In the reduced Pair-HMM EM algorithm used here, we optimize the endpoint alignment likelihood rather than the complete-data likelihood of the full birth-death-substitution process. Consequently, the reduced substitution update for $Q$ retains only matched-lineage bridge contributions. For the equilibrium distribution $\pi$, we additionally retain observable endpoint composition counts from insertion and deletion states, while omitting genealogical correction terms from nonterminal birth and death times and from fully transient hidden lineages.

Considering now the joint distribution $P(x, y|\theta)$ and including the sufficient statistics for the stationary composition and length distribution of ancestral sequences, as well as the compositional statistics for inserted sequences that are already included in the conditional case, the expectations are

$$\begin{aligned}
\mathbb{E}[L \mid x, y] &= \sum_{i \in \{\mathtt{S},\mathtt{M},\mathtt{I},\mathtt{D}\}} (\hat{n}_{i\mathtt{M}} + \hat{n}_{i\mathtt{D}}) = |x| \\
\mathbb{E}[M \mid x, y] &= 1 \\
\mathbb{E}[V_i \mid x, y] &= \sum_b \hat{e}^{\mathtt{M}}_{(i,b)} + \hat{e}^{\mathtt{I}}_{(\epsilon,i)} + \hat{e}^{\mathtt{D}}_{(i,\epsilon)} + \mathcal{G}_i^V(x, y)
\end{aligned}$$

where the three observable terms count, respectively, ancestral residues at match positions (drawn from $\pi$ at time 0 and surviving until $t$), descendant residues at insert positions (drawn from $\pi$ at the time of insertion), and ancestral residues at delete positions (drawn from $\pi$ at time 0 and deleted before $t$); $\mathcal{G}_i^V$ is a similar genealogical correction term, the omission of which again corresponds to assuming (for the purposes of the CTMC parameter update) that insertion and deletion events are synchronized with the trajectory endpoints, and that no transient characters are born and die within the trajectory. Thus $V_i$ counts every observable equilibrium $\pi$-draw of character $i$ in the joint pair HMM, regardless of whether the residue subsequently underwent a match-position substitution, was deleted, or never had an ancestor (insertion). The conditional $P(y|x, t)$ analogue drops the two ancestor-derived terms (matches and deletions), since the ancestral sequence is given.

##### A.1.8 Baum-Welch Algorithm for TKF91

Given a collection of ancestral and descendant sequences $\{x^{(n)}, y^{(n)}\}$, the Baum-Welch algorithm iterates between an E-step and an M-step.

**E-step.** For each of the sequence pairs, run the Forward-Backward algorithm on the TKF91 WFST yielding the expected transition counts $\hat{n}_{ab}$ for $a, b \in \{\mathtt{S},\mathtt{M},\mathtt{I},\mathtt{D},\mathtt{E}\}$ and the expected emission counts $\hat{e}^{\mathtt{M}}_{(a,b)}, \hat{e}^{\mathtt{I}}_{(\epsilon,b)}, \hat{e}^{\mathtt{D}}_{(a,\epsilon)}$. Use these to accumulate expectations for the sufficient statistics $S, B, D, W, U, L, M, V$ as described in the previous section. In this section we will simply refer to the accumulated expectations for these statistics using the variable names for the statistics themselves. Thus $S \equiv \mathbb{E}[S \mid \{x^{(n)}, y^{(n)}\}]$ and so forth.

**M-step.** The goal is to maximize $\ell(\theta) = \ell_1(\lambda, \mu) + \ell_2(Q, \pi)$ where

$$\ell_1(\lambda,\mu) = (B+L)\log\lambda + (D-L-M)\log\mu + M\log(\mu-\lambda) - (\lambda+\mu)\,S - \lambda T \quad (A.18)$$
$$\ell_2(Q,\pi) = \sum_i V_i' \log\pi_i + \sum_{j>i}(U_{ij}+U_{ji})\log Q_{ij} - \sum_{j>i} Q_{ij}\,(W_i\pi_j + W_j\pi_i) \quad (A.19)$$

using the reversible parameterization $\kappa = \lambda/\mu$ and $R_{ij} = Q_{ij}\pi_j$ and letting $V_i' = V_i + \sum_{j\neq i} U_{ji}$ for notational convenience. Differentiating and setting to zero yields equations the MLEs must satisfy. For $\lambda, \mu$ these are simultaneous quadratics

$$\partial_\lambda \ell_1 = \frac{B+L}{\lambda} - \frac{M}{\mu-\lambda} - S - T = 0 \quad (A.20)$$
$$\partial_\mu \ell_1 = \frac{D-L-M}{\mu} + \frac{M}{\mu-\lambda} - S = 0 \quad (A.21)$$

which reduce to a single quadratic in $\kappa$. Setting $a = B + L$ and $b = D - L - M$

$$\kappa^2(S+T)b - \kappa\,(S(a+b+2M) + T(b+M)) + Sa = 0. \quad (A.22)$$

For the coefficients $A_\kappa = (S+T)b$, $B_\kappa = -\,(S(a+b+2M) + T(b+M))$, and $C_\kappa = Sa$, the root $\kappa \in (0,1)$ will be the smaller of the two roots

$$\hat\kappa(B,D,L,M,S,T) = \frac{-B_\kappa - \sqrt{B_\kappa^2 - 4A_\kappa C_\kappa}}{2A_\kappa} \quad (A.23)$$
$$\hat\mu(B,D,L,M,S,T) = \frac{1}{S}\left(b + \frac{M}{1-\hat\kappa}\right) \quad (A.24)$$
$$\hat\lambda(B,D,L,M,S,T) = \hat\kappa\hat\mu \quad (A.25)$$

For $Q, \pi$ the MLE equations are

$$\partial_{Q_{ij}} \ell_2 = \frac{U_{ij}+U_{ji}}{Q_{ij}} - (W_i\pi_j + W_j\pi_i) = 0 \quad (A.26)$$
$$\partial_{\pi_i} \ell_2 = \frac{V_i'}{\pi_i} - \sum_{j\neq i} Q_{ij}W_j + \eta = 0 \quad (A.27)$$

where in the second equation $\eta$ is a Lagrange multiplier for the constraint $\sum_i \pi_i = 1$. These equations are nonlinearly coupled and do not, in general, have a closed-form solution, nor even a closed-form iterative solution. There are several practical alternatives:

1. Set $\pi$ from insertions and ancestral composition, $\pi_i = V_i/\sum_j V_j$, and then solve for $Q$ in closed form
$$Q_{ij} = \frac{U_{ij}+U_{ji}}{W_i\pi_j + W_j\pi_i}.$$

2. Fixing $\pi$, solve for $Q$ using the above closed form; then solve for $\eta$ from $\sum_i V_i'/(\sum_{j\neq i} Q_{ij}W_j - \eta) = 1$, and set $\pi_i \leftarrow V_i'/(\sum_{j\neq i} Q_{ij}W_j - \eta)$, iterating until convergence. For fixed $Q$, the optimization over $\pi$ is strictly concave on the simplex since $V_i' > 0$ for all $i$ and the Hessian is negative definite.

3. Eliminate $Q$ to yield a self-consistency map for $\pi$. Banach fixed-point guarantees are non-obvious for this approach.

4. Use the unconstrained MLE for $R_{ij} = Q_{ij}\pi_j$ and then project or reparameterize it into reversible GTR form. If the underlying process is reversible, this can provide a useful approximation or initialization, though finite-sample estimates need not satisfy detailed balance exactly.

##### A.1.9 Extending TKF91 to Phylogenetic Trees

Hein (18) showed how to compute a multi-sequence Forward likelihood for TKF91 on a binary tree. Holmes and Bruno (24) used a similar approach to compute posterior marginals for Gibbs sampling of ancestral sequences and alignments. Lunter *et al* showed how to calculate alignment likelihoods efficiently (36). Suchard and Redelings (42) extended the MCMC sampling approach to jointly sample over MSAs and trees. Westesson et al. (52) showed how this approach can be seen as a generalization of Felsenstein's pruning algorithm for substitution-only models, where the WFST plays the role of a matrix exponential defined on whole sequences, with the WFST's Forward algorithm computing the entries of this matrix exponential. The linear algebraic interpretation was further remarked upon by Bouchard-Côté (**?** ).

#### A.2 The TKF92 Model

In sequence-alignment terms, TKF92 is the classical fragment-based extension of TKF91 and induces gap behavior analogous to an affine gap penalty (51). Instead of each mortal link getting a single character from $\mathcal{A}$, each link is associated with a fragment consisting of $K \geq 1$ characters from $\mathcal{A}$, where $K \sim \text{Geometric}(r)$. We write this as

$$\mathcal{S}'(T) \sim \text{TKF92}(\text{Subst}(Q, \pi); \lambda, \mu, r)$$

where $\mathcal{S}' \in (\mathcal{A}^+)^*$ is a sequence of multi-character fragments. Specifically

$$\text{TKF92}(\text{Subst}(Q, \pi); \lambda, \mu, r) \equiv \text{Links}(\text{Frag}(\text{Subst}(Q, \pi); r); \lambda, \mu)$$

where $\text{Frag}(M, r)$ denotes a tuple of $K$ i.i.d. random variables, each governed by $M$, and $K$ is geometrically-distributed with parameter $r$

$$x \sim \text{Frag}(M, r) \;\Leftrightarrow\; K \sim \text{Geometric}(r),\; x = (x_1 \ldots x_K),\; x_k \sim M$$

The TKF92 model thus has the same BDI process as TKF91, but with a more complex substitution process that emits fragments instead of single characters.

##### A.2.1 Latent Information in TKF92

The mean length of an insertion in TKF92 is $1/(1-r)$, i.e. the expected fragment length. For deletions, things are slightly more complicated. Superficially, it looks like deletions are also fragments so they should have the same mean length. However, if we do not have knowledge of the fragment boundaries, but are conditioning just on the current sequence length, we have to weight this by the posterior probability that the subsequence being deleted does in fact constitute a single fragment. For a subsequence of length $k$, this posterior probability is proportional to $\left(\frac{r}{r+(1-r)\frac{\lambda}{\mu}}\right)^{k-1}$ (indels at the end of the sequence have a slightly different correction factor since they are more likely to constitute a fragment, but the $k$-dependence is the same). The rate of starting an insertion or deletion *at a particular site* is subject to a similar correction factor.

This illustrates how the derivation of the WFST from the Pair HMM of such models becomes more complicated as the models carry greater amounts of latent information in their state space.

##### A.2.2 Singlet HMM, Pair HMM, and WFST Representations

The TKF92 Singlet HMM, Pair HMM, and WFST have the same state spaces $\{\mathtt{S},\mathtt{M},\mathtt{I},\mathtt{D},\mathtt{E}\}$ as TKF91, but with transition probabilities modified to account for fragment extension. Within a link's fragment, each character beyond the first extends the fragment with probability $r$, so the fragment terminates with probability $1-r$.

**Singlet HMM (stationary distribution).** The total sequence length in TKF92 is a geometric sum of geometric fragment lengths. Since $N \sim \text{Geometric}(\kappa)$ links and each fragment has length $K \sim \text{Geometric}(r)$, the total length $L = \sum_{i=1}^{N} K_i$ is distributed as $P(L = 0) = 1 - \kappa$ and $P(L = \ell \mid L \geq 1) = (1-p)\,p^{\ell-1}$ where $p = r + \kappa(1-r)$ is the effective continuation probability. The HMM emits no characters with probability $1-\kappa$, emits a first character with probability $\kappa$, and then continues with probability $p$ after each emitted character. This is equivalent to a zero-inflated geometric distribution with parameters $\kappa$ and $p$.

**Pair HMM (finite-time joint distribution).** Each emitting state has a fragment extension self-loop with probability $r$. On fragment termination (probability $1-r$), the TKF91 link-level transitions apply. The Pair HMM transition matrix is

$$\tau'(\lambda,\mu,T,r) = \left(\begin{array}{c|ccccc} & \mathtt{S} & \mathtt{M} & \mathtt{I} & \mathtt{D} & \mathtt{E} \\ \hline \mathtt{S} & 0 & \tau_{\mathtt{SM}} & \tau_{\mathtt{SI}} & \tau_{\mathtt{SD}} & \tau_{\mathtt{SE}} \\ \mathtt{M} & 0 & r+(1-r)\tau_{\mathtt{MM}} & (1-r)\tau_{\mathtt{MI}} & (1-r)\tau_{\mathtt{MD}} & (1-r)\tau_{\mathtt{ME}} \\ \mathtt{I} & 0 & (1-r)\tau_{\mathtt{IM}} & r+(1-r)\tau_{\mathtt{II}} & (1-r)\tau_{\mathtt{ID}} & (1-r)\tau_{\mathtt{IE}} \\ \mathtt{D} & 0 & (1-r)\tau_{\mathtt{DM}} & (1-r)\tau_{\mathtt{DI}} & r+(1-r)\tau_{\mathtt{DD}} & (1-r)\tau_{\mathtt{DE}} \\ \mathtt{E} & 0 & 0 & 0 & 0 & 0 \end{array}\right)$$

where $\tau_{ab}(\lambda,\mu,T)$ are the TKF91 Pair HMM transition probabilities. Each row sums to 1: for $a \in \{\mathtt{M},\mathtt{I},\mathtt{D}\}$, $r + (1-r)\sum_b \tau_{ab} = r + (1-r) = 1$.

**WFST (finite-time conditional distribution).** Forming a WFST for TKF92 is complicated by the latent information associated with fragment boundaries. If those boundaries are known, one can construct a WFST that respects them exactly. If they are not known, one can still construct a plausible WFST that imputes them on the fly. This trick, however, ceases to be straightforward for more highly decorated descendants of TKF91. Alternatively, one can expose the fragment boundary information in the transducer alphabet, at the cost of reifying indivisibility of fragments over time. The choice between these approaches is non-obvious and presumably application-dependent. The explicit TKF92 WFST construction, obtained by dividing the Pair HMM by the Singlet HMM, is given in Appendix A.3. That construction requires a 6×6 state space $(\mathtt{S},\mathtt{M},\mathtt{I}_0,\mathtt{I}_1,\mathtt{D},\mathtt{E})$ rather than the 5×5 form one might naively borrow from TKF91, because the singlet's outgoing-emission weight differs between the start state ($\kappa$ for the first ancestor character) and any subsequent emit state ($p = r + (1-r)\kappa$), so the insert state must be split into a pre-immortal-link variant ($\mathtt{I}_0$) and a post-immortal-link variant ($\mathtt{I}_1$). The resulting WFST is conditionally normalised in the global sense: for any input ancestor sequence $x$, the total weight of all paths from $\mathtt{S}$ to $\mathtt{E}$ that read $x$ exactly equals 1, summing over all alignments and descendant outputs. Equivalently, Singlet $\circ\, \mathbb{T}'' = \tau''$ is row-stochastic on the 6-state space. Local single-step row sums and per-state $\varepsilon$-closure normalisation hold for the $\mathtt{S},\mathtt{M},\mathtt{I}_0,\mathtt{D}$ rows but not for the $\mathtt{I}_1$ row, which is improper in isolation but accumulates the correct $\kappa/p$ factor whenever it is reached from a proper source state along an $\mathtt{S}$-rooted path (Appendix A.3.7). The same trade-off recurs for MixDom WFSTs in Section C.1.5, and is discussed more carefully in Appendix A.3.7.

These issues are not unique to TKF92, but are a general feature of models with latent information that is not directly observable in the sequence data. They are dealt with at greater length in the section on MixDom WFSTs (Section C.1.5), where the same issues arise in a more complex setting. To state the general principle: as TKF91-derived models acquire additional latent decoration, that decoration must either be promoted to explicit symbols in the interface alphabet of the transducer construction, or integrated out, in which case the resulting transducer is only approximate.

##### A.2.3 Baum-Welch Algorithm for TKF92

The Baum-Welch algorithm for TKF92 is similar to that for TKF91, but we must now resolve the Pair HMM's self-looping transition counts onto their separate components (fragment extension vs. new link), and accumulate fragment-level sufficient statistics in addition to the substitution-level sufficient statistics.

**E-step.** Run Forward-Backward on the TKF92 Pair HMM. This yields expected transition counts $\hat{n}'_{ab}$. For $a \in \{\mathtt{M}, \mathtt{I}, \mathtt{D}\}$, the self-loop count $\hat{n}'_{aa}$ combines fragment extension (probability $r$) and new-link-same-state (probability $(1-r)\tau_{aa}$). The expected number of fragment extensions from state $a$ is $\hat{n}'_{aa} \cdot r/(r + (1-r)\tau_{aa})$ and the expected number of new-link self-transitions is the remainder. Thus, we accumulate new expectations for the additional fragment-level sufficient statistics, $F$ the number of fragment extensions and $E$ the number of fragment ends, and then recover the TKF91 link-level counts by removing the fragment extensions

$$
\begin{aligned}
F_a &= \frac{\hat{n}'_{aa} \cdot r}{r + (1-r)\tau_{aa}}, \ a \in \{\mathtt{M}, \mathtt{I}, \mathtt{D}\} \\
F &= \sum_{a \in \{\mathtt{M}, \mathtt{I}, \mathtt{D}\}} F_a \\
E &= \sum_{a \in \{\mathtt{M}, \mathtt{I}, \mathtt{D}\}} \sum_b \hat{n}'_{ab} - F \\
\hat{n}_{ab} &= \hat{n}'_{ab} - \delta_{ab} F_a
\end{aligned}
$$

We then proceed to accumulate $S, B, D, W, U, L, M, V$ as for TKF91. Note that $L$ counts *links*, which are now fragments rather than simply residues: $L = \hat{n}_\kappa = \sum_i (\hat{n}_{i\mathtt{M}} + \hat{n}_{i\mathtt{D}})$ where $\hat{n}$ is the resolved (TKF91-level) count matrix. The character-level statistics $W, U, V$ are accumulated per character exactly as in TKF91, because fragment extension affects the indel process estimates but not the substitution process estimates.

**M-step.** This is identical to TKF91, except we also set $r \leftarrow F/(F+E)$.

##### A.2.4 Maraschino: Distilled Cherries

The TKF92 Baum-Welch algorithm, as with other instances of the Expectation Maximization algorithm, is most useful if there is hidden information that we need to marginalize during parameter estimation; e.g. an unknown alignment that needs to be summed over, or (as in Section C.1.1) latent site or fragment classes that are not directly observed.

For many applications, we can take the alignment data (and potentially site classes) as a given, and then a more efficient estimation strategy becomes available. This is the CherryML approach: a composite likelihood consisting of a product of alignments likelihoods for nearest-neighbor sibling pairs, a.k.a. "cherries" (40). The CherryML developers observe that, given cherry branch lengths, the alignment substitution counts are sufficient to estimate substitution rates and can be performed efficiently using autograd, independently of data size except for a fast preprocessing step We observe that this can be generalized to TKF92 by including the alignment transition statistics i.e. the sixteen transition counts $n_{\mathtt{X}\to\mathtt{Y}}$ where $\mathtt{X} \in \{\mathtt{S}, \mathtt{M}, \mathtt{I}\,del\}$ and $\mathtt{Y} \in \{\mathtt{M}, \mathtt{I}, \mathtt{D}, \mathtt{E}\}$ states. The CherryML approach can then be applied: 1) discretize times, yielding a fixed-shape tensor of summary counts; 2) write down the observed data likelihood in terms of the summary counts; 3) maximize using autograd and standard gradient-based optimizers.

We call this algorithm *Maraschino*, as it involves distilling cherries to state machines.

##### A.3 TKF92 WFST by Singlet Division

This appendix derives the TKF92 conditional WFST $\mathbb{T}'(\lambda, \mu, T, r)$ representing $P(\text{descendant} \mid \text{ancestor}, \theta)$ by dividing the TKF92 Pair HMM by the TKF92 Singlet HMM.

The TKF91 WFST (Section A.1.6) is a 5×5 matrix obtained from the TKF91 Pair HMM $\tau$ by stripping $\kappa$ from ancestor-consuming columns ($\mathtt{M}, \mathtt{D}$) and $1-\kappa$ from the end column ($\mathtt{E}$). The TKF92 WFST cannot be expressed as a 5×5 matrix in the same state space. As we show in Section A.3.2, the singlet's outgoing-emission weight differs between its $\mathtt{S}$ state and its emit state, so the two cases must be carried as distinct insert states (*Ins0* and *Ins1*) in the WFST. The natural object is therefore 6×6, with state space $\{\mathtt{S}, \mathtt{M}, \mathtt{I}_0, \mathtt{I}_1, \mathtt{D}, \mathtt{E}\}$. A 5×5 presentation that lumps $\mathtt{I}_0$ and $\mathtt{I}_1$ into a single $\mathtt{I}$ state and uses the emit-state divisor $p$ throughout is incorrect for paths that traverse $\mathtt{S} \to \mathtt{I} \to \cdots \to \{\mathtt{M}, \mathtt{D}\}$; this is discussed in Section A.3.6.

###### A.3.1 TKF92 Singlet HMM Transitions

The TKF92 singlet at stationarity emits sequences whose length $L$ follows a zero-inflated geometric distribution (Section A.2.2). The singlet HMM has two transition contexts:

- *From* $\mathtt{S}$ *(before any character):* $P(\text{emit first character}) = \kappa, \quad P(\text{end}) = 1 - \kappa$.
- *From an emitting state (after character* $i \geq 1$*):* $P(\text{emit next character}) = p \equiv r + (1-r)\kappa$, $P(\text{end}) = (1-r)(1-\kappa)$.

The effective continuation probability $p$ decomposes as: fragment extension with probability $r$, or fragment termination $(1-r)$ followed by a new nonempty link ($\kappa$). Each emitted character is drawn i.i.d. from $\pi$.

The key consequence for the WFST construction below is that $\kappa \neq p$ whenever $r > 0$.

###### A.3.2 Why an Ins0 / Ins1 Split is Necessary

Transducer composition Singlet ∘ WFST = Pair HMM couples each ancestor-consuming Pair HMM transition to one of the two singlet emission events:

- the $\mathtt{S} \to$ emit event (weight $\kappa$), used *exactly once*, for the first ancestor character; or
- the emit $\to$ emit event (weight $p$), used for every subsequent ancestor character.

The Pair HMM's $\mathtt{I}$ state corresponds to "the descendant is in the middle of a descendant-only fragment." This state is reached either *before* the first ancestor character has been consumed (entered from $\mathtt{S}$, possibly via $\mathtt{I} \to \mathtt{I}$ self-loops) or *after* $\geq 1$ ancestor characters have been consumed (entered from $\mathtt{M}, \mathtt{D}, \mathtt{I}$). In the first case, the next ancestor-consuming transition out of $\mathtt{I}$ corresponds to the $\mathtt{S} \to$ emit event in the singlet (weight $\kappa$); in the second case, it corresponds to emit $\to$ emit (weight $p$). The two divisors differ unless $r = 0$.

The 5×5 Pair HMM $\tau'(\lambda, \mu, T, r)$ of Section A.2.2 marginalises over the "has any ancestor character been consumed yet?" bit and is correct as a generator of the joint distribution. But because that bit controls which singlet divisor applies, the WFST construction must expose it. We therefore split the Pair HMM $\mathtt{I}$ state into

- $\mathtt{I}_0$ — reachable only from $\mathtt{S}$ (via $\mathtt{S} \to \mathtt{I}_0$) and from itself (via $\mathtt{I}_0 \to \mathtt{I}_0$): the singlet has not yet emitted.
- $\mathtt{I}_1$ — reachable from $\mathtt{M}, \mathtt{D}, \mathtt{I}_1$: the singlet has emitted at least one character.

The 6×6 Pair HMM in this state space is row-stochastic (verified below), and the 5×5 Pair HMM is recovered by lumping $\mathtt{I}_0$ and $\mathtt{I}_1$ into a single $\mathtt{I}$ row. The lumping preserves the joint distribution because the outgoing transitions from $\mathtt{I}_0$ and $\mathtt{I}_1$ have identical Pair HMM structure once weighted by their singlet contributions; what fails to lump is the WFST.

##### A.3.3 The 6×6 TKF92 Pair HMM

Writing $\alpha, \beta, \gamma, \kappa$ for the TKF link-level parameters (as in Section A.1.6), and $r = r$ for the fragment-extension probability:

$$\tau''(\lambda, \mu, T, r) = \left( \begin{array}{c|cccccc} & \mathtt{S} & \mathtt{M} & \mathtt{I}_0 & \mathtt{I}_1 & \mathtt{D} & \mathtt{E} \\ \hline \mathtt{S} & 0 & (1-\beta)\kappa\alpha & \beta & 0 & (1-\beta)\kappa(1-\alpha) & (1-\beta)(1-\kappa\text{[illegible]} \\ \mathtt{M} & 0 & r + (1-r)(1-\beta)\kappa\alpha & 0 & (1-r)\beta & (1-r)(1-\beta)\kappa(1-\alpha) & (1-r)(1-\beta)(1-\text{[illegible]} \\ \mathtt{I}_0 & 0 & (1-r)(1-\beta)\kappa\alpha & r + (1-r)\beta & 0 & (1-r)(1-\beta)\kappa(1-\alpha) & (1-r)(1-\beta)(1-\text{[illegible]} \\ \mathtt{I}_1 & 0 & (1-r)(1-\beta)\kappa\alpha & 0 & r + (1-r)\beta & (1-r)(1-\beta)\kappa(1-\alpha) & (1-r)(1-\beta)(1-\text{[illegible]} \\ \mathtt{D} & 0 & (1-r)(1-\gamma)\kappa\alpha & 0 & (1-r)\gamma & r + (1-r)(1-\gamma)\kappa(1-\alpha) & (1-r)(1-\gamma)(1-\text{[illegible]} \\ \mathtt{E} & 0 & 0 & 0 & 0 & 0 & 0 \end{array} \right. \tag{A.28}$$

The structural zeros are: $\mathtt{S}$ cannot reach $\mathtt{I}_1$ in one step (no ancestor has yet been emitted), and $\mathtt{M}, \mathtt{D}, \mathtt{I}_1$ cannot reach $\mathtt{I}_0$ in one step (an ancestor has already been emitted). The $\mathtt{I}_0$ and $\mathtt{I}_1$ rows differ only in their self-loop column ($\mathtt{I}_0 \to \mathtt{I}_0$ vs $\mathtt{I}_1 \to \mathtt{I}_1$); their entries to $\mathtt{M}, \mathtt{D}, \mathtt{E}$ are identical.

**Row-stochasticity.** For each non-$\mathtt{E}$ source row, the row sum equals

$$r + (1-r)\Big[\beta + (1-\beta)\kappa\alpha + (1-\beta)\kappa(1-\alpha) + (1-\beta)(1-\kappa)\Big] = r + (1-r)\big[\beta + (1-\beta)\big] = 1$$

(replacing $\beta$ with $\gamma$ for the $\mathtt{D}$ row), confirming that $\tau''$ is a valid Markov transition matrix.

**Recovery of the 5×5 Pair HMM.** Lumping $\mathtt{I}_0$ and $\mathtt{I}_1$ into a single $\mathtt{I}$ state preserves all four non-$\mathtt{I}$-column entries (which are identical between the two rows by construction). In the self-loop column, $\mathtt{I}_0 \to \mathtt{I}_0$ and $\mathtt{I}_1 \to \mathtt{I}_1$ both equal $r + (1-r)\beta$, so the lumped self-loop is also $r + (1-r)\beta$. The result is the 5×5 Pair HMM of Section A.2.2, recovered exactly.

##### A.3.4 The 6×6 TKF92 WFST

Each Pair HMM transition consumes either one ancestor character ($\to \mathtt{M}$ or $\to \mathtt{D}$), zero ancestor characters ($\to \mathtt{I}_0$ or $\to \mathtt{I}_1$), or signals termination ($\to \mathtt{E}$). The WFST $\mathbb{T}''$ is obtained by dividing each Pair HMM entry by the corresponding singlet weight:

- Ancestor-consuming entries from $\mathtt{S}, \mathtt{I}_0$ (the next ancestor character is the *first*): divide by $\kappa$.

- Ancestor-consuming entries from $\mathtt{M}, \mathtt{I}_1, \mathtt{D}$ (the next ancestor character is a *subsequent* one): divide by $p = r + (1-r)\kappa$.
- $\mathtt{E}$ entries from $\mathtt{S}, \mathtt{I}_0$ (the ancestor sequence has length zero): divide by $1-\kappa$.
- $\mathtt{E}$ entries from $\mathtt{M}, \mathtt{I}_1, \mathtt{D}$ (the ancestor sequence has length $\geq 1$): divide by $(1-r)(1-\kappa)$.
- $\mathtt{I}_0$ and $\mathtt{I}_1$ entries (no ancestor consumed): divide by 1.

The emission factor $\pi_a$ is also stripped from the emission weights, exactly as in TKF91, so that M-state emission becomes $\exp(RT)_{ab}$ and D-state emission becomes 1; we focus on the transition matrix below.

$$\mathbb{T}''(\lambda, \mu, T, r) = \left(\begin{array}{c|cccccc} & \mathtt{S} & \mathtt{M} & \mathtt{I}_0 & \mathtt{I}_1 & \mathtt{D} & \mathtt{E} \\ \hline \mathtt{S} & 0 & (1-\beta)\alpha & \beta & 0 & (1-\beta)(1-\alpha) & 1-\beta \\ \mathtt{M} & 0 & \dfrac{r + (1-r)(1-\beta)\kappa\alpha}{p} & 0 & (1-r)\beta & \dfrac{(1-r)(1-\beta)\kappa(1-\alpha)}{p} & 1-\beta \\ \mathtt{I}_0 & 0 & (1-r)(1-\beta)\alpha & r + (1-r)\beta & 0 & (1-r)(1-\beta)(1-\alpha) & (1-r)(1-\beta) \\ \mathtt{I}_1 & 0 & \dfrac{(1-r)(1-\beta)\kappa\alpha}{p} & 0 & r + (1-r)\beta & \dfrac{(1-r)(1-\beta)\kappa(1-\alpha)}{p} & 1-\beta \\ \mathtt{D} & 0 & \dfrac{(1-r)(1-\gamma)\kappa\alpha}{p} & 0 & (1-r)\gamma & \dfrac{r + (1-r)(1-\gamma)\kappa(1-\alpha)}{p} & 1-\gamma \\ \mathtt{E} & 0 & 0 & 0 & 0 & 0 & 0 \end{array}\right) \tag{A.29}$$

The $\mathtt{I}_1$ row uses the $p$ divisor on its M and D exits and $1-\beta$ on its E exit, matching the post-immortal-link regime. The $\mathtt{I}_0$ row uses $\kappa$ on its M, D exits (absorbed into the numerator: $(1-r)(1-\beta)\kappa\alpha/\kappa = (1-r)(1-\beta)\alpha$, etc.) and $1-\kappa$ on its E exit (giving $(1-r)(1-\beta)$). These are exactly the TKF91 WFST S/M/I-row outputs scaled by $(1-r)$, reflecting the leading-extension fragment's chance of terminating before any ancestor is consumed.

##### A.3.5 Structure and Verification

**Sanity check 1: S row.** The S row inherits the TKF91 form because the first character of the immortal-link descendant fragment has no extension self-loop. Multiplying by the singlet emission weights from S ($\kappa$ for M, D, 1 for $\mathtt{I}_0$, $1-\kappa$ for E) recovers the Pair HMM S row of (A.28).

**Sanity check 2: $\mathtt{I}_0$ row.** Multiplying $\mathbb{T}''$ entries by the appropriate singlet weights ("from S" weights: $\kappa$ for M/D, 1 for $\mathtt{I}_0$, $1-\kappa$ for E):

$$\begin{aligned} \kappa \cdot (1-r)(1-\beta)\alpha = (1-r)(1-\beta)\kappa\alpha &= \tau''_{\mathtt{I}_0\mathtt{M}}, \\ 1 \cdot [r + (1-r)\beta] = r + (1-r)\beta &= \tau''_{\mathtt{I}_0\mathtt{I}_0}, \\ \kappa \cdot (1-r)(1-\beta)(1-\alpha) = (1-r)(1-\beta)\kappa(1-\alpha) &= \tau''_{\mathtt{I}_0\mathtt{D}}, \\ (1-\kappa) \cdot (1-r)(1-\beta) = (1-r)(1-\beta)(1-\kappa) &= \tau''_{\mathtt{I}_0\mathtt{E}}. \end{aligned}$$

All four entries match the Pair HMM, so Singlet $\circ\, \mathbb{T}'' = \tau''$ on the $\mathtt{I}_0$ row.

**Sanity check 3: $\texttt{M}$ row.** Multiplying by emit-state weights ($p$ for $\texttt{M}/\texttt{D}$, 1 for $\texttt{I}_1$, $(1-r)(1-\kappa)$ for $\texttt{E}$):

$$\begin{aligned}
p \cdot \frac{r + (1-r)(1-\beta)\kappa\alpha}{p} &= r + (1-r)(1-\beta)\kappa\alpha \;=\; \tau''_{\texttt{MM}}, \\
1 \cdot (1-r)\beta &= (1-r)\beta \;=\; \tau''_{\texttt{MI}_1}, \\
p \cdot \frac{(1-r)(1-\beta)\kappa(1-\alpha)}{p} &= (1-r)(1-\beta)\kappa(1-\alpha) \;=\; \tau''_{\texttt{MD}}, \\
(1-r)(1-\kappa) \cdot (1-\beta) &= (1-r)(1-\beta)(1-\kappa) \;=\; \tau''_{\texttt{ME}}.
\end{aligned}$$

The $\texttt{I}_1$ and $\texttt{D}$ rows verify identically (with $\gamma$ for $\beta$ in the $\texttt{D}$ row).

**TKF91 limit ($r = 0$).** At $r = 0$, $p = \kappa$ and $r = 0$. The $\texttt{I}_0$ and $\texttt{I}_1$ rows become:

$$\begin{aligned}
\mathbb{T}''_{\texttt{I}_0 * \,|\, r=0} &= \big(0,\ (1-\beta)\alpha,\ \beta,\ 0,\ (1-\beta)(1-\alpha),\ 1-\beta\big), \\
\mathbb{T}''_{\texttt{I}_1 * \,|\, r=0} &= \big(0,\ (1-\beta)\alpha,\ 0,\ \beta,\ (1-\beta)(1-\alpha),\ 1-\beta\big).
\end{aligned}$$

With the $\texttt{I}_0$ and $\texttt{I}_1$ self-loops both equal to $\beta$ and all other columns identical, the two rows reduce to the same TKF91-Ins row when $\texttt{I}_0, \texttt{I}_1$ are merged. The TKF91 WFST (Section A.1.6) is recovered.

**Latent fragment-boundary posterior.** The $\texttt{M}$ row's $\texttt{M} \to \texttt{M}$ entry decomposes as

$$\mathbb{T}''_{\texttt{MM}} = \underbrace{\frac{r}{p}}_{\substack{\text{same}\\\text{frag}}} \cdot 1 \;+\; \underbrace{\frac{(1-r)\kappa}{p}}_{\substack{\text{new}\\\text{frag}}} \cdot (1-\beta)\alpha,$$

exactly as in the original construction; under same-fragment continuation the descendant certainly stays in $\texttt{M}$ (the link persists within the fragment), and under new-fragment start the descendant must survive $(1-\beta)$ and align $(\alpha)$. The fragment-boundary posterior $r/p + (1-r)\kappa/p = 1$ holds by definition of $p$. The $\texttt{I}_0$ row has *no* same-fragment ancestor-consuming term — since the singlet has not yet emitted, the fragment-boundary posterior is degenerate at "new fragment" — which is why the $\texttt{I}_0 \to \texttt{M}$ entry has the simple $(1-r)(1-\beta)\alpha$ form rather than a $1/p$-divided form.

##### A.3.6 Comparison with the 5×5 Form

A 5×5 presentation that replaces the two rows $\texttt{I}_0, \texttt{I}_1$ of (A.29) by a single $\texttt{I}$ row equal to the $\texttt{I}_1$ row is correct only at $r = 0$. For $r > 0$, that 5×5 form misweights paths $\texttt{S} \to \texttt{I} \to \cdots \to \{\texttt{M}, \texttt{D}, \texttt{E}\}$ with one or more leading inserts before the first ancestor consumption: the divisor on the leading-insert exit should be $\kappa$ (or $1-\kappa$ for $\texttt{E}$) rather than $p$ (or $(1-r)(1-\kappa)$). For a single-leading-insert path $\texttt{S} \to \texttt{I} \to \texttt{M} \to \texttt{E}$, the multiplicative discrepancy is $(1-r)\kappa/p < 1$ when $r > 0$, so the 5×5 WFST *underweights* such paths.

For benchmarks dominated by alignments without leading inserts (typical when the ancestor's $N$-terminus is well-conserved), the numerical effect of the discrepancy is small per pair. However, the 5×5 form does not represent the true conditional distribution $P(\text{descendant} \mid \text{ancestor}, \theta)$, and the 6×6 WFST in (A.29) is the construction we recommend.

##### A.3.7 Normalization Structure

The WFST is conditionally normalised in the global sense: for every input sequence $x \in \Sigma^*$,

$$\sum_{(y,\,\text{alignment})} \big[\text{WFST path-weight}\big] \;=\; 1,$$

where the sum is over all paths from $\mathtt{S}$ to $\mathtt{E}$ that read $x$ exactly. This is a direct consequence of Singlet $\circ\, \mathbb{T}'' = \tau''$ (verified row by row in Section A.3.5) combined with the fact that $\tau''$ is row-stochastic and the Singlet HMM defines a proper distribution on $x$.

The WFST is *not* stochastic in the local single-step sense: its rows do not sum to 1, and the divisor structure (ancestor-consuming columns divide by $\kappa$ from $\mathtt{S}, \mathtt{I}_0$ and by $p$ from $\mathtt{M}, \mathtt{I}_1, \mathtt{D}$; the $\mathtt{E}$ column divides by $1{-}\kappa$ from $\mathtt{S}, \mathtt{I}_0$ and by $(1{-}r)(1{-}\kappa)$ from $\mathtt{M}, \mathtt{I}_1, \mathtt{D}$; insert columns divide by 1) means a fixed source row's outgoing weights are not constrained to sum to 1.

**$\varepsilon$-closure conditional normalisation.** A more useful local property is the following. Define $w_X^L$ as the total WFST weight of paths starting at state $X$ that consume the next non-$\varepsilon$ input symbol $L$ (a character or end-of-input sentinel \$), where each path is a (possibly empty) geometric chain of insert self-loops followed by a single input-consuming exit. For our 6×6 WFST:

| $X$ | $w_X^c$ | $w_X^\$$ |
|---|---|---|
| $\mathtt{S}$ | 1 | 1 |
| $\mathtt{M}$ | 1 | 1 |
| $\mathtt{I}_0$ | 1 | 1 |
| $\mathtt{I}_1$ | $\kappa/p$ | $1/(1-r)$ |
| $\mathtt{D}$ | 1 | 1 |

(A.30)

The $\mathtt{S}, \mathtt{M}, \mathtt{I}_0, \mathtt{D}$ rows are conditionally normalised: each sums to exactly 1. The $\mathtt{I}_1$ row alone is not, because its destination divisors mix the $r$ (extension) and $1{-}r$ (exit) weights asymmetrically into the single-step edges.

This local non-normalisation does *not* affect the WFST's correctness as a representation of $P(\text{descendant} \mid \text{ancestor}, \theta)$: every WFST path begins at $\mathtt{S}$, and from $\mathtt{S}$ the chain-through-$\mathtt{I}_1$ contribution accumulates exactly the right $\kappa/p$ factor at the appropriate step to bring the total back to 1. Concretely, starting from $\mathtt{M}$ the $\varepsilon$-closure weight to consume the next character is

$$w_{\mathtt{M}}^c = \big(\mathbb{T}''_{\mathtt{MM}} + \mathbb{T}''_{\mathtt{MD}}\big) + \mathbb{T}''_{\mathtt{MI}_1} \cdot \frac{\mathbb{T}''_{\mathtt{I}_1\mathtt{M}} + \mathbb{T}''_{\mathtt{I}_1\mathtt{D}}}{1 - \mathbb{T}''_{\mathtt{I}_1\mathtt{I}_1}} = \frac{r + (1{-}r)(1{-}\beta)\kappa}{p} + (1{-}r)\beta \cdot \frac{\kappa}{p} = \frac{r + (1{-}r)\kappa}{p} = 1,$$

so the $\kappa/p$ factor that the $\mathtt{I}_1$ row alone undershoots is exactly recovered when $\mathtt{I}_1$ is reached from a "proper" source state. The $\mathtt{I}_1$ row is improper only if the WFST were started mid-stream from $\mathtt{I}_1$ in isolation, which never happens in normal use.

**Per-input single-step row sums.** The simpler invariant $\sum_b \mathbb{T}''_{Xb} = 1$ does *not* hold for any of the body rows. This is a state-encoding artefact of folding the Bernoulli-$r$ extension-vs-exit decision into the same outgoing edges that carry the destination-singlet factor. An alternative WFST that exposes the extension event as a silent "decision" state would be locally row-stochastic at the cost of more states. The compact 6×6 form trades local stochasticity for a smaller graph; the global property Singlet $\circ\, \mathbb{T}'' = \tau''$ (with $\tau''$ row-stochastic) is preserved either way.

##### A.3.8 Testable Invariants

The following invariants hold for any valid parameters $0 < \lambda, \mu$, $T > 0$, $0 \leq r < 1$ and can be verified numerically from the WFST tensor code.

1. **Fragment-boundary posterior:** $r/p + (1-r)\kappa/p = 1$.
2. **Pair HMM reconstruction:** For each state $a$ in $\{\mathtt{S}, \mathtt{M}, \mathtt{I}_0, \mathtt{I}_1, \mathtt{D}\}$, reconstruct the Pair HMM row by multiplying WFST entries by their singlet factors (per the rules above), and verify $\sum_b \tau''_{ab} = 1$.
3. **Mixture decomposition:** $\mathbb{T}''_{\mathtt{MM}} = r/p + [(1-r)\kappa/p](1-\beta)\alpha$ (and analogously for $\mathtt{D} \to \mathtt{D}$ with $\gamma$). Both mixture components are non-negative.
4. **TKF91 limit:** At $r = 0$: $p = \kappa$, the $\mathtt{I}_0$ and $\mathtt{I}_1$ rows become identical (and equal the TKF91 $\mathtt{I}$ row), and the 6×6 WFST reduces to the TKF91 5×5 WFST.
5. **$\mathtt{E}$-column dichotomy:** $\mathbb{T}''_{X\mathtt{E}} = 1-\beta$ for $X \in \{\mathtt{S}, \mathtt{M}, \mathtt{I}_1\}$ and $1-\gamma$ for $X = \mathtt{D}$, but $\mathbb{T}''_{\mathtt{I}_0\mathtt{E}} = (1-r)(1-\beta)$ scales linearly in $1-r$ because the $\mathtt{I}_0$ exit weight is divided by the smaller $\mathtt{S} \to \mathtt{E}$ singlet weight $1-\kappa$ rather than $(1-r)(1-\kappa)$.
6. **$\mathtt{I}_1$ column scaling:** $\mathbb{T}''_{\mathtt{MI}_1} = (1-r)\beta$ and $\mathbb{T}''_{\mathtt{DI}_1} = (1-r)\gamma$ scale linearly in $(1-r)$, reflecting that inserts can only begin at fragment boundaries.
7. **Structural zeros:** $\mathbb{T}''_{\mathtt{S}\,\mathtt{I}_1} = \mathbb{T}''_{\mathtt{M}\,\mathtt{I}_0} = \mathbb{T}''_{\mathtt{I}_1\,\mathtt{I}_0} = \mathbb{T}''_{\mathtt{D}\,\mathtt{I}_0} = 0$, reflecting that $\mathtt{I}_0$ is reachable only before any ancestor has been consumed and $\mathtt{I}_1$ only after.

#### A.4 Equal-Rate Limits for TKF Parameters

For biologically realistic equilibrium lengths under TKF91/TKF92, one is typically driven into the near-critical regime $\lambda \approx \mu$ ($\kappa \approx 1$). Under TKF91, the expected equilibrium sequence length is $\kappa/(1-\kappa)$ characters. Under TKF92 with fragment extension probability $r$, it is $\kappa/((1-r)(1-\kappa))$ characters (over $\kappa/(1-\kappa)$ fragments). The average protein in UniProtKB/Swiss-Prot is approximately 360 amino acids long (49). Even under TKF92 with a mean indel length of $1/(1-r) \approx 3$ residues (matching the empirically observed predominance of 1–5 residue indels in protein evolution (1, 7)), the required $\kappa$ is $120/121 \approx 0.992$. Under TKF91, $\kappa = 360/361 \approx 0.997$. For DNA sequences (a typical prokaryotic ORF or eukaryotic intron-free coding sequence is ~1000 bp (53); intron-bearing eukaryotic genes and syntenic genomic regions can be much longer), $\kappa > 0.999$. Since $1 - \kappa = (\mu - \lambda)/\mu$, these biological sequence lengths force $\lambda$ very close to $\mu$.

The TKF91 transition probabilities $\alpha, \beta, \gamma$ and the BDI score/sufficient statistics all contain factors of $(\mu - \lambda)$ in denominators. When $\lambda \to \mu$ ($\kappa \to 1$), these become 0/0 indeterminate forms. This section derives the L'Hôpital limits, providing numerically stable expressions in the equal-rate regime.

Let $\varepsilon = \mu - \lambda \to 0$ and $s = \mu T$. We parameterize $\lambda = \mu - \varepsilon$ and take $\varepsilon \to 0^+$.

Throughout, we write $\Phi = 1-\alpha$ for the complement of the survival probability ($\Phi$ is well-defined at any $(\lambda, \mu)$). In the equal-rate regime, where $s = \mu T$ and $\alpha = e^{-s}$, we additionally define:

$$\phi = \Phi/s = (1 - e^{-s})/s$$

Note $\phi(0) = 1$ and $\phi(s) \to 0$ as $s \to \infty$.

##### A.4.1 TKF91 Transition Parameters

**$\alpha$ (survival probability).** $\alpha = e^{-\mu T}$ has no dependence on $\lambda$; no singularity.

**$\beta$ (insertion continuation probability).**

$$\beta = \frac{\lambda(e^{-\lambda T} - e^{-\mu T})}{\mu\, e^{-\lambda T} - \lambda\, e^{-\mu T}} \tag{A.31}$$

Write $e^{-\lambda T} = \alpha\, e^{\varepsilon T}$ and let $u = \varepsilon T$. Then:

$$\begin{aligned} \text{Numerator} &= (\mu - \varepsilon)\, \alpha\, (e^u - 1) \\ \text{Denominator} &= \alpha\big(\mu\, e^u - \mu + \varepsilon\big) \end{aligned}$$

Cancelling $\alpha$ and expanding $e^u = 1 + u + u^2/2 + \cdots$ with $u = \varepsilon T$:

$$\begin{aligned} \text{Numer} &= (\mu - \varepsilon)(\varepsilon T + \varepsilon^2 T^2/2 + \cdots) = \varepsilon\big(\mu T + O(\varepsilon)\big) \\ \text{Denom} &= \mu\varepsilon T + \varepsilon(1 + O(\varepsilon)) = \varepsilon\big(\mu T + 1 + O(\varepsilon)\big) \end{aligned}$$

After cancelling $\varepsilon$:

$$\boxed{\beta \xrightarrow{\lambda\to\mu} \frac{s}{1+s}} \tag{A.32}$$

**$\gamma$ (orphan insertion probability).** Using $\gamma = 1 - \mu\beta/(\lambda(1-\alpha))$:

$$\boxed{\gamma \xrightarrow{\lambda\to\mu} 1 - \frac{1}{(1+s)\,\phi} = \frac{(1+s)\phi - 1}{(1+s)\phi}} \tag{A.33}$$

**Complementary parameters.**

$$1 - \beta \;\to\; \frac{1}{1+s} \tag{A.34}$$

$$1 - \gamma \;\to\; \frac{1}{(1+s)\,\phi} \tag{A.35}$$

##### A.4.2 Score Derivatives: General Case

Define $\eta = e^{-\lambda T}$ and $\delta = \mu\eta - \lambda\alpha$ (the denominator of $\beta$). The score of the observed log-likelihood decomposes into 8 log-parameter groups. Below we list the derivatives of each log-factor with respect to $\lambda$ and $\mu$, from which the log-elasticities follow by multiplication:

$$\frac{\partial \log \xi}{\partial \log \lambda} = \lambda\, \frac{\partial \log \xi}{\partial \lambda}, \qquad \frac{\partial \log \xi}{\partial \log \mu} = \mu\, \frac{\partial \log \xi}{\partial \mu}, \qquad \frac{\partial \log \xi}{\partial \log(\lambda{+}\mu)} = \frac{\partial \log \xi}{\partial \log \lambda} + \frac{\partial \log \xi}{\partial \log \mu}$$

The third identity holds because differentiating with respect to $\log(\lambda{+}\mu)$ at fixed ratio $\kappa = \lambda/\mu$ is equivalent to scaling both rates by a common factor $c$ and differentiating with respect to $\log c$, which gives $\lambda\, \partial_\lambda + \mu\, \partial_\mu$.

$\log \alpha$ **and** $\log(1{-}\alpha)$ Since $\alpha = e^{-\mu T}$ depends only on $\mu$:

$$\frac{\partial \log \alpha}{\partial \lambda} = 0, \qquad \frac{\partial \log \alpha}{\partial \mu} = -T \tag{A.36}$$

$$\frac{\partial \log(1{-}\alpha)}{\partial \lambda} = 0, \qquad \frac{\partial \log(1{-}\alpha)}{\partial \mu} = \frac{T\alpha}{1-\alpha} = \frac{T\alpha}{\Phi} \tag{A.37}$$

$\log\beta$ **and** $\log(1-\beta)$ From $\log\beta = \log\lambda + \log(\eta-\alpha) - \log\delta$:

$$\frac{\partial\log\beta}{\partial\lambda} = \frac{1}{\lambda} - \frac{T\eta}{\eta-\alpha} + \frac{T\mu\eta+\alpha}{\delta} \tag{A.38}$$

$$\frac{\partial\log\beta}{\partial\mu} = \frac{T\alpha}{\eta-\alpha} - \frac{\eta+T\lambda\alpha}{\delta} \tag{A.39}$$

Both $(\eta-\alpha)\to 0$ and $\delta\to 0$ as $\lambda\to\mu$, so each term individually diverges; the combination is finite (Section A.4.3).

For $\log(1-\beta)$, using the chain rule $\partial\log(1-\beta) = -\frac{\beta}{1-\beta}\,\partial\log\beta$:

$$\frac{\partial\log(1-\beta)}{\partial\lambda} = -\frac{\beta}{1-\beta}\cdot\frac{\partial\log\beta}{\partial\lambda} \tag{A.40}$$

$$\frac{\partial\log(1-\beta)}{\partial\mu} = -\frac{\beta}{1-\beta}\cdot\frac{\partial\log\beta}{\partial\mu} \tag{A.41}$$

$\log(1-\gamma)$ **and** $\log\gamma$ From $1-\gamma = \mu\beta/(\lambda(1-\alpha))$, so $\log(1-\gamma) = \log\mu + \log\beta - \log\lambda - \log(1-\alpha)$:

$$\frac{\partial\log(1-\gamma)}{\partial\lambda} = \frac{\partial\log\beta}{\partial\lambda} - \frac{1}{\lambda} \tag{A.42}$$

$$\frac{\partial\log(1-\gamma)}{\partial\mu} = \frac{1}{\mu} + \frac{\partial\log\beta}{\partial\mu} - \frac{T\alpha}{1-\alpha} \tag{A.43}$$

For $\log\gamma$:

$$\frac{\partial\log\gamma}{\partial\lambda} = -\frac{1-\gamma}{\gamma}\cdot\frac{\partial\log(1-\gamma)}{\partial\lambda} \tag{A.44}$$

$$\frac{\partial\log\gamma}{\partial\mu} = -\frac{1-\gamma}{\gamma}\cdot\frac{\partial\log(1-\gamma)}{\partial\mu} \tag{A.45}$$

##### Summary Table: General Case

| | $\log\alpha$ | $\log(1-\alpha)$ | $\log\beta$ | $\log(1-\beta)$ | $\log\gamma$ | $\log(1-\gamma)$ |
|---|---|---|---|---|---|---|
| $\frac{\partial}{\partial\log\lambda}$ | $0$ | $0$ | $1-\frac{s_\lambda\eta}{\eta-\alpha}+\frac{s_\lambda\mu\eta/\lambda+\lambda\alpha/\lambda}{\delta/\lambda}$ | $-\frac{\beta}{1-\beta}\cdot[\cdot]^\lambda_\beta$ | $-\frac{1-\gamma}{\gamma}\cdot[\cdot]^\lambda_{1-\gamma}$ | $[\cdot]^\lambda_\beta-1$ |
| $\frac{\partial}{\partial\log\mu}$ | $-s$ | $\frac{s\alpha}{\Phi}$ | $\frac{s\alpha}{\eta-\alpha}-\frac{\mu\eta+s\lambda\alpha}{\delta}$ | $-\frac{\beta}{1-\beta}\cdot[\cdot]^\mu_\beta$ | $-\frac{1-\gamma}{\gamma}\cdot[\cdot]^\mu_{1-\gamma}$ | $1+[\cdot]^\mu_\beta-\frac{s\alpha}{\Phi}$ |
| $\frac{\partial}{\partial\log(\lambda+\mu)}$ | (sum of above two rows) | | | | | |

where $s_\lambda = \lambda T$, $s=\mu T$, and $[\cdot]^\theta_\xi$ denotes the $\partial\log\xi/\partial\log\theta$ entry.

The table is dense because each $\log\beta$ and $\log\gamma$ entry involves the singular terms $(\eta-\alpha)^{-1}$ and $\delta^{-1}$. These singularities cancel in each entry, as we show next.

##### A.4.3 Score Derivatives: L'Hôpital Limits

We systematically expand each derivative to first order in $\varepsilon = \mu - \lambda$ and extract the finite limit. With $u = \varepsilon T$, the key expansions are:

$$\eta - \alpha = \alpha(e^u - 1) = \alpha\bigl(u + u^2/2 + \cdots\bigr) \tag{A.46}$$

$$\delta = \mu\eta - \lambda\alpha = \alpha\varepsilon(1 + s + O(\varepsilon)) \tag{A.47}$$

and the Laurent expansion $1/(e^u - 1) = 1/u - 1/2 + u/12 - \cdots$, giving:

$$\frac{T\eta}{\eta - \alpha} = \frac{Te^u}{e^u - 1} = \frac{1}{\varepsilon} + \frac{T}{2} + O(\varepsilon) \tag{A.48}$$

$$\frac{T\alpha}{\eta - \alpha} = \frac{T}{e^u - 1} = \frac{1}{\varepsilon} - \frac{T}{2} + O(\varepsilon) \tag{A.49}$$

$$\frac{T\mu\eta + \alpha}{\delta} = \frac{1}{\varepsilon} + \frac{sT}{2(1+s)} + O(\varepsilon) \tag{A.50}$$

$$\frac{\eta + T\lambda\alpha}{\delta} = \frac{1}{\varepsilon} - \frac{sT}{2(1+s)} + O(\varepsilon) \tag{A.51}$$

Derivation of (A.50): the numerator is $T\mu\eta + \alpha = \alpha(se^u + 1) = \alpha\bigl((1+s) + s\varepsilon T + O(\varepsilon^2)\bigr)$ and the denominator is $\delta = \alpha\varepsilon\bigl((1+s) + s\varepsilon T/2 + O(\varepsilon^2)\bigr)$ from (A.47). So:

$$\frac{T\mu\eta + \alpha}{\delta} = \frac{(1+s) + s\varepsilon T + \cdots}{\varepsilon\bigl((1+s) + s\varepsilon T/2 + \cdots\bigr)} = \frac{1}{\varepsilon} \cdot \frac{1 + \frac{s\varepsilon T}{1+s} + \cdots}{1 + \frac{s\varepsilon T}{2(1+s)} + \cdots}$$

The $O(\varepsilon)$ coefficients differ: $sT/(1+s)$ in the numerator vs. $sT/(2(1+s))$ in the denominator. Their difference gives the $O(1)$ correction: $1/\varepsilon + sT/(2(1+s)) + O(\varepsilon)$. Similarly for (A.51).

**$\partial \log\beta/\partial\lambda$ and $\partial \log\beta/\partial\mu$** Using (A.38) with (A.48) and (A.50):

$$\begin{aligned}\frac{\partial \log\beta}{\partial\lambda} &= \frac{1}{\lambda} - \left(\frac{1}{\varepsilon} + \frac{T}{2}\right) + \left(\frac{1}{\varepsilon} + \frac{sT}{2(1+s)}\right) + O(\varepsilon)\\ &= \frac{1}{\mu} - \frac{T}{2} + \frac{sT}{2(1+s)} = \frac{1}{\mu} - \frac{T}{2(1+s)}\end{aligned}$$

Using (A.39) with (A.49) and (A.51):

$$\begin{aligned}\frac{\partial \log\beta}{\partial\mu} &= \left(\frac{1}{\varepsilon} - \frac{T}{2}\right) - \left(\frac{1}{\varepsilon} - \frac{sT}{2(1+s)}\right) + O(\varepsilon)\\ &= -\frac{T}{2} + \frac{sT}{2(1+s)} = -\frac{T}{2(1+s)}\end{aligned}$$

Converting to log-elasticities:

$$\boxed{\frac{\partial \log\beta}{\partial \log\lambda} \to \frac{s+2}{2(1+s)}, \qquad \frac{\partial \log\beta}{\partial \log\mu} \to -\frac{s}{2(1+s)}} \tag{A.52}$$

**Check.** As a consistency check, we verify that the sum of the two elasticities (which gives the response to uniformly scaling both rates at fixed $\kappa = \lambda/\mu$) matches the direct derivative of the limit formula. The sum is $(s+2)/(2(1+s)) + (-s)/(2(1+s)) = 1/(1+s)$. At the limit, $\beta = s/(1+s)$ where $s = \mu T$, so $d\log\beta/d\log s = 1/(1+s)$. ✓

$\partial \log(1{-}\beta)/\partial \log \lambda$ **and** $\partial \log(1{-}\beta)/\partial \log \mu$  Using $\beta/(1-\beta) \to s$ at equal rates:

$$\boxed{\frac{\partial \log(1{-}\beta)}{\partial \log \lambda} \to -\frac{s(s+2)}{2(1+s)}, \qquad \frac{\partial \log(1{-}\beta)}{\partial \log \mu} \to \frac{s^2}{2(1+s)}} \tag{A.53}$$

**Check.** Summing the two elasticities (uniform rate scaling): $(-s^2{-}2s{+}s^2)/(2(1{+}s)) = -s/(1{+}s)$. Direct: $d \log(1{-}\beta)/d \log s = -s/(1{+}s)$. ✓

$\partial \log(1{-}\gamma)/\partial \log \lambda$ **and** $\partial \log(1{-}\gamma)/\partial \log \mu$  From (A.42): $\partial \log(1{-}\gamma)/\partial\lambda = \partial \log \beta/\partial\lambda - 1/\lambda \to 1/\mu - T/(2(1{+}s)) - 1/\mu = -T/(2(1{+}s))$.

From (A.43): $\partial \log(1{-}\gamma)/\partial\mu = 1/\mu + \partial \log \beta/\partial\mu - T\alpha/\Phi \to 1/\mu - T/(2(1{+}s)) - T\alpha/\Phi$.

Converting:

$$\boxed{\frac{\partial \log(1{-}\gamma)}{\partial \log \lambda} \to -\frac{s}{2(1+s)}, \qquad \frac{\partial \log(1{-}\gamma)}{\partial \log \mu} \to \frac{s+2}{2(1+s)} - \frac{s\alpha}{\Phi}} \tag{A.54}$$

**Check.** Summing the two elasticities (uniform rate scaling): $-s/(2(1{+}s)) + (s{+}2)/(2(1{+}s)) - s\alpha/\Phi = 1/(1{+}s) - s\alpha/\Phi$. Direct: at the limit, $1-\gamma = s/((1{+}s)\Phi)$, so $\log(1{-}\gamma) = \log s - \log(1{+}s) - \log\Phi$. Under $s \to cs$: $d/d\log c|_{c=1} = 1 - s/(1{+}s) - s\alpha/\Phi = 1/(1{+}s) - s\alpha/\Phi$. ✓

$\partial \log\gamma/\partial \log \lambda$ **and** $\partial \log\gamma/\partial \log \mu$  With $R = (1{-}\gamma)/\gamma$ at equal rates. Since $1-\gamma = 1/((1{+}s)\phi)$:

$$R = \frac{1}{(1+s)\phi - 1}$$

$$\boxed{\frac{\partial \log \gamma}{\partial \log \lambda} \to \frac{Rs}{2(1+s)}, \qquad \frac{\partial \log \gamma}{\partial \log \mu} \to -R\Big(\frac{s+2}{2(1+s)} - \frac{s\alpha}{\Phi}\Big)} \tag{A.55}$$

**Summary Table: L'Hôpital Limits**  At $\lambda = \mu$, with $s = \mu T$, $\alpha = e^{-s}$, $\Phi = 1-\alpha$, $\phi = \Phi/s$, $R = 1/((1{+}s)\phi - 1)$:

| | $\log\alpha$ | $\log(1{-}\alpha)$ | $\log\beta$ | $\log(1{-}\beta)$ | $\log\gamma$ | $\log(1{-}\gamma)$ |
|---|---|---|---|---|---|---|
| $\frac{\partial}{\partial \log \lambda}$ | 0 | 0 | $\frac{s+2}{2(1+s)}$ | $-\frac{s(s+2)}{2(1+s)}$ | $\frac{Rs}{2(1+s)}$ | $-\frac{s}{2(1+s)}$ |
| $\frac{\partial}{\partial \log \mu}$ | $-s$ | $\frac{s\alpha}{\Phi}$ | $-\frac{s}{2(1+s)}$ | $\frac{s^2}{2(1+s)}$ | $-R\Big(\frac{s+2}{2(1+s)} - \frac{s\alpha}{\Phi}\Big)$ | $\frac{s+2}{2(1+s)} - \frac{s\alpha}{\Phi}$ |
| $\frac{\partial}{\partial \log(\lambda+\mu)}$ | $-s$ | $\frac{s\alpha}{\Phi}$ | $\frac{1}{1+s}$ | $-\frac{s}{1+s}$ | $R\Big(\frac{s}{2(1+s)} - \frac{s+2}{2(1+s)} + \frac{s\alpha}{\Phi}\Big)$ | $\frac{1}{1+s} - \frac{s\alpha}{\Phi}$ |

All entries are finite. The third row equals the sum of the first two (as it must by construction). The $\log\alpha$ and $\log(1{-}\alpha)$ columns are trivially continuous (no $\lambda$ dependence). The $\log\beta$ and $\log(1{-}\beta)$ entries have a particularly clean form involving only $s/(1{+}s)$. The $\log\gamma$ entries additionally involve $\alpha/\Phi = e^{-s}/(1-e^{-s}) = 1/(e^s - 1)$ and the ratio $R = 1/((1{+}s)\phi - 1)$.

##### A.4.4 BDI Sufficient Statistics

The EM-relevant quantities are the posterior expectations $\mathbb{E}[B|i,j,T]$, $\mathbb{E}[D|i,j,T]$, $\mathbb{E}[S|i,j,T]$, recovered from the score via the conservation law $i+B-D=j$:

$$\mathbb{E}[S] = \frac{j-i+\mu\,\partial_\mu \log P_{ij} - \lambda\,\partial_\lambda \log P_{ij} - \lambda T}{\lambda-\mu} \tag{A.56}$$

$$\mathbb{E}[B] = \lambda\,\partial_\lambda \log P_{ij} + \lambda\,\mathbb{E}[S] + \lambda T \tag{A.57}$$

$$\mathbb{E}[D] = \mu\,\partial_\mu \log P_{ij} + \mu\,\mathbb{E}[S] \tag{A.58}$$

The score $\partial_\lambda \log P_{ij}$ itself is computed from the transition count groups:

$$\partial_\lambda \log P_{ij} = \sum_{\xi\in\{\alpha,1-\alpha,\beta,\ldots\}} n_{\log\xi}\cdot\frac{\partial\log\xi}{\partial\lambda}$$

Each derivative $\partial\log\xi/\partial\lambda$ has a finite L'Hôpital limit (Section A.4.3), so the score itself has a finite limit. The denominator $\lambda-\mu$ in (A.56) is what creates the 0/0 for $\mathbb{E}[S]$.

**The $\mathbb{E}[S]$ singularity.** The numerator of (A.56) also vanishes at $\lambda=\mu$, since the complete-data log-likelihood for $\nu=\lambda$ is $\ell = B\log\lambda + D\log\mu - (\lambda+\mu)S - \lambda T + c$, and at $\lambda=\mu$ the numerator becomes $j-i+\mathbb{E}[D]-\mathbb{E}[B]=0$ by conservation.

To resolve the 0/0, write $f(\lambda) = j-i+\mu\,\partial_\mu\log P - \lambda\,\partial_\lambda \log P - \lambda T$ and $g(\lambda)=\lambda-\mu$. By L'Hôpital ($g'=1$):

$$\lim_{\lambda\to\mu}\mathbb{E}[S] = f'(\mu) = \mu\,\partial^2_{\lambda\mu}\log P - \partial_\lambda\log P - \mu\,\partial^2_{\lambda^2}\log P - T \tag{A.59}$$

where all derivatives are evaluated at $\lambda=\mu$.

The first and second derivatives of $\log P$ that appear in (A.59) can be expressed in terms of the transition count groups and the L'Hôpital-limited score derivatives from Section A.4.3, together with their first-order corrections in $\varepsilon$. Specifically, the second derivatives of $\log P$ require the $O(\varepsilon)$ terms in the Laurent expansions (A.48)–(A.51).

**Alternative: direct gradient on $\mu$.** At equal rates, the BDI has a single parameter $\mu$. The gradient is:

$$\left.\frac{\partial\log P}{\partial\mu}\right|_{\lambda=\nu=\mu} = \sum_\xi n_{\log\xi}\left(\frac{\partial\log\xi}{\partial\lambda}+\frac{\partial\log\xi}{\partial\mu}\right) = \frac{\mathbb{E}[B+D]}{\mu} - 2\mathbb{E}[S] - T$$

This is well-defined using the L'Hôpital limits, and gives the combined quantity $\mathbb{E}[B+D]-\mu(2\mathbb{E}[S]+T)$ without needing to separate $\mathbb{E}[B]$, $\mathbb{E}[D]$, $\mathbb{E}[S]$ individually. The M-step (A.60) sets this to zero: $\hat{\mu} = (\mathbb{E}[B]+\mathbb{E}[D])/(2\mathbb{E}[S]+T)$.

To *break out* of equal rates (determine whether $\lambda$ should differ from $\mu$), we need the individual $\partial\log P/\partial\lambda$ and $\partial\log P/\partial\mu$, which the table in Section A.4.3 provides, or we need to use additional information from the ancestral length distribution.

##### A.4.5 Direct BDI Rate EM at Equal Rates

If one ignores the stationary root-length prior and considers only the conditional BDI bridge model with $\lambda=\nu=\mu$, the complete-data log-likelihood (A.10) reduces to:

$$\ell = (B+D)\log\mu - 2\mu S - \mu T + c$$

This has a single free parameter $\mu$. Setting $\partial\ell/\partial\mu = 0$:

$$\boxed{\hat{\mu} = \frac{B+D}{2S+T}} \tag{A.60}$$

where $B$, $D$, $S$ are understood as posterior expectations.

**Implication.** At equal rates, the M-step collapses to estimating a single rate from the total event count and total sojourn time.

##### A.4.6 $\log\kappa$ and $\log(1-\kappa)$ Derivatives

The $\kappa = \lambda/\mu$ and $(1-\kappa)$ groups are not among the six $(\alpha, \beta, \gamma)$ factors but appear in the transition matrix. At $\kappa = 1$:

$$\frac{\partial \log\kappa}{\partial\lambda} = \frac{1}{\lambda} \to \frac{1}{\mu}, \qquad \frac{\partial \log\kappa}{\partial\mu} = -\frac{1}{\mu}$$

No singularity. However, $\log(1-\kappa)$ diverges:

$$\frac{\partial \log(1-\kappa)}{\partial\lambda} = \frac{-\kappa}{(1-\kappa)\lambda} \to -\infty$$

Unlike the conditional WFST factors $\alpha, \beta, \gamma$, the stationary-length factors $\kappa$ and $1-\kappa$ do not admit a regular equal-rate limit in the full joint TKF model. As $\kappa \to 1$, the equilibrium length distribution ceases to be normalizable, and the $\log(1-\kappa)$ contribution diverges. Accordingly, the L'Hôpital limits derived above apply to the conditional bridge/WFST quantities, whereas the full stationary joint model must either remain in the subcritical regime $\kappa < 1$ or be reparameterized with a nonstationary root-length model.

##### A.4.7 Joint vs Conditional Likelihoods

The TKF91 transition matrix includes factors of $\kappa = \lambda/\mu$ and $(1-\kappa)$ arising from the geometric($\kappa$) prior on ancestor length: $\Pr(|\text{anc}| = i) = \kappa^i(1-\kappa)$. This prior is part of the *joint* likelihood $P(x, y)$ but can be factored out with respect to the ancestor-length prior to yield an *ancestor length-conditioned* likelihood $P_{\text{cond}}(x, y \mid |x| = i) = P(x, y)/[\kappa^i(1-\kappa)]$.

**The four regimes.** Of the four combinations of $\kappa < 1$ vs $\kappa = 1$ and joint vs ancestor length-conditioned, only one is degenerate:

| | $\kappa < 1$ | $\kappa = 1$ |
|---|---|---|
| Joint $P(x, y)$ | well-defined | **degenerate** ($\to 0$) |
| Ancestor length-conditioned $P_{\text{cond}}$ | well-defined | well-defined |

At $\kappa = 1$, the geometric prior places vanishing mass on finite sequences, so $P(x, y) \to 0$ and $\log P \to -\infty$. The ancestor length-conditioned WFST removes the $\kappa$ and $(1-\kappa)$ factors corresponding to the stationary length distribution from the transition matrix, leaving only $(\alpha, \beta, \gamma)$-dependent terms. These have finite L'Hôpital limits (Sections A.4.3–A.4.4), so $P_{\text{cond}}$ is finite and well-defined even at $\kappa = 1$.

**BDI sufficient statistics.** The BDI sufficient statistics $\mathbb{E}[B]$, $\mathbb{E}[D]$, $\mathbb{E}[S]$ are most naturally recovered from the ancestor length-conditioned WFST, whose score depends only on the six TKF bridge groups $(\alpha, 1-\alpha, \beta, 1-\beta, \gamma, 1-\gamma)$. In the joint model, additional contributions from the ancestral length prior appear through the $\kappa$ and $1-\kappa$ factors. These must be included in the M-step when optimizing the full stationary TKF likelihood, but they are conceptually distinct from the bridge statistics themselves.

Equations (A.15)-A.17 give the correct expectations for the BDI sufficient statistics in terms of the counts $\hat{n}_{ij}$ regardless of whether those counts were computed using the joint Pair HMM or the conditionally-normalized transducer, as long as the WFST transition matrix $\mathbb{T}$ is used for the counts $\mathcal{C}_{ij}^{\xi} = \xi \partial_\xi \log \mathbb{T}_{ij}$ (e.g. using the tables in Section A.4.2 or Section A.4.3). The M-step for the joint likelihood includes additional terms from the $\kappa$ and $1-\kappa$ factors, leading to a coupled system of equations that can be solved in closed form (Section A.1.8).

**Closed-form M-steps.** Both formulations yield closed-form M-steps:

- *Ancestor-length conditioned:* pure BDI exponential family. For the conditional model away from the $\kappa = 1$ singularity of the joint prior, setting $\partial Q_{\text{cond}}/\partial\lambda = 0$ gives

$$\hat{\lambda} = \frac{\max(0,\ \mathbb{E}[B] + \alpha_\lambda - 1)}{\mathbb{E}[S] + T + \beta_\lambda}, \qquad \hat{\mu} = \frac{\max(0,\ \mathbb{E}[D] + \alpha_\mu - 1)}{\mathbb{E}[S] + \beta_\mu} \tag{A.61}$$

  where $(\alpha, \beta)$ are Gamma prior parameters.

- *Joint ($\kappa < 1$):* includes $L \log\kappa + M \log(1-\kappa)$. Since, for a single sequence-pair, $L = i$ (ancestor length, exact) and $M = 1$ (one End transition, exact), with corresponding accumulated values across multiple sequence-pairs, the optimality conditions $\partial Q/\partial\lambda = 0$ and $\partial Q/\partial\mu = 0$ form a coupled system in $(\lambda, \mu)$. These can be reduced to a single quadratic equation in $\kappa$ with a closed-form solution, which can be substituted back to get $\hat{\lambda}$ and $\hat{\mu}$, as described in Section A.1.8.

In both cases, the M-step maximizes the correct $Q$-function, so EM monotonicity is guaranteed for the corresponding tracked objective ($\log P_{\text{cond}}$ for the ancestor length-conditioned model, $\log P$ for the full joint model).

##### A.4.8 Irreversible Models and the $\lambda > \mu$ Regime

**The $\kappa > 1$ regime.** When $\lambda > \mu$, the ratio $\kappa = \lambda/\mu > 1$ and the geometric($\kappa$) prior on ancestor length is not normalizable: $\sum_{i=0}^{\infty} \kappa^i (1-\kappa)$ diverges. Therefore the joint likelihood $P(x, y)$ does not exist. However, the ancestor length-conditioned likelihood $P_{\text{cond}}(x, y \mid |x|)$ is well-defined for all $\kappa > 0$. The transition parameters $\alpha$, $\beta$, $\gamma$ are continuous functions of $(\lambda, \mu, T)$ with no singularity at $\lambda = \mu$ or $\lambda > \mu$. The BDI sufficient statistics $\mathbb{E}[B]$, $\mathbb{E}[D]$, $\mathbb{E}[S]$ remain well-defined, and the conditioned M-step (A.61) applies unchanged: it is unconstrained in $\lambda$ and $\mu$ and permits $\hat{\lambda} > \hat{\mu}$ if the data support it.

**Irreversible substitution models.** For completeness, our implementations support dropping the reversibility assumption for the substitution CTMC. The integrals described in Section A.1.2 and in (25) exploit the symmetric factorization of $R$ using $\pi$-weighted similarity, which yields real eigenvalues. In the irreversible case, $R$ is a general rate matrix with no detailed-balance constraint.

The eigendecomposition $R = V \operatorname{diag}(d) V^{-1}$ may have complex eigenvalues. Assuming $R$ is diagonalizable, the spectral formulae for the integrals of Section A.1.2 carry over, with $V^{-1}$ replacing $V^\top$ and complex arithmetic throughout:

$$I_{ij}^{ab}(T) = \sum_{k,l} V_{ak} V_{ki}^{-1} \frac{e^{d_k T} - e^{d_l T}}{d_k - d_l} V_{jl} V_{lb}^{-1}$$

with the convention $(e^{d_k T} - e^{d_l T})/(d_k - d_l) = T e^{d_k T}$ when $d_k = d_l$. The result is real (complex eigenvalues come in conjugate pairs).

**Fixed-point iteration for $\pi$.** In the reversible case, the equilibrium distribution $\pi$ is constrained to be the stationary distribution of $R$ by by detailed balance, and may either be fixed a priori or updated jointly with the reversible rate parameters during the M-step. In the irreversible case, $\pi$ must be recomputed as the left eigenvector of $R$ (with eigenvalue 0) after each M-step update of $R$ from the Holmes–Rubin sufficient statistics. This gives an ECM (Expectation-Conditional-Maximization) scheme (37): the E-step uses the current $(R, \pi)$; the M-step updates $R$, then $\pi$ is recomputed from the updated $R$. Standard monotonicity results apply providing each conditional maximization step is well-defined.

**Differentiability.** Standard autograd libraries provide general eigendecomposition and matrix-exponential primitives (with Fréchet derivatives). The eigendecomposition route is convenient for EM-style closed-form updates, while gradient-based optimization is often more naturally based on the matrix exponential. We do not rely on differentiating through eigenvectors here.

##### A.4.9 Summary of Limits

| **Quantity** | **Formula** | **Limit as $\lambda \to \mu$** |
|---|---|---|
| $\alpha$ | $e^{-\mu T}$ | $e^{-s}$ (continuous) |
| $\beta$ | $\frac{\lambda(\eta-\alpha)}{\mu\eta-\lambda\alpha}$ | $\frac{s}{1+s}$ |
| $1-\beta$ | | $\frac{1}{1+s}$ |
| $\gamma$ | $1 - \frac{\mu\beta}{\lambda(1-\alpha)}$ | $1 - \frac{1}{(1+s)\phi}$ |
| $\kappa$ | $\lambda/\mu$ | 1 |
| $\partial_{\log\lambda} \log\beta$ | (A.38) | $\frac{s+2}{2(1+s)}$ |
| $\partial_{\log\mu} \log\beta$ | (A.39) | $-\frac{s}{2(1+s)}$ |
| $\hat{\mu}_{\text{MLE}}$ | | $\frac{\mathbb{E}[B]+\mathbb{E}[D]}{2\mathbb{E}[S]+T}$ |

where $s = \mu T$ throughout.

#### A.5 TKF91 Score Function

We derive the explicit partial derivatives needed to compute the observed-data score from Pair HMM transition counts, as described in Section A.1.8.

##### A.5.1 Derivatives of TKF parameters

Define $\eta = e^{-\lambda T}$ (so that $\alpha = e^{-\mu T}$ and $\eta = e^{-\lambda T}$) and $\Delta = \mu\eta - \lambda\alpha$ (the denominator of $\beta$).

$\alpha = e^{-\mu T}$.

$$\frac{\partial \log \alpha}{\partial \lambda} = 0, \qquad \frac{\partial \log \alpha}{\partial \mu} = -T$$

$\beta = \lambda(\eta - \alpha)/\Delta$. Using $\log \beta = \log \lambda + \log(\eta - \alpha) - \log \Delta$:

$$\begin{aligned} \frac{\partial \log \beta}{\partial \lambda} &= \frac{1}{\lambda} - \frac{T\eta}{\eta - \alpha} + \frac{T\mu\eta + \alpha}{\Delta} \\ \frac{\partial \log \beta}{\partial \mu} &= \frac{T\alpha}{\eta - \alpha} - \frac{\eta + T\lambda\alpha}{\Delta} \end{aligned}$$

$\gamma = 1 - \mu\beta/(\lambda(1 - \alpha))$. Using $\log(1 - \gamma) = \log \mu + \log \beta - \log \lambda - \log(1 - \alpha)$:

$$\begin{aligned} \frac{\partial \log(1-\gamma)}{\partial \lambda} &= \frac{\partial \log \beta}{\partial \lambda} - \frac{1}{\lambda} \\ \frac{\partial \log(1-\gamma)}{\partial \mu} &= \frac{1}{\mu} + \frac{\partial \log \beta}{\partial \mu} + \frac{T\alpha}{1-\alpha} \end{aligned}$$

and $\frac{\partial \log \gamma}{\partial \theta} = -\frac{1-\gamma}{\gamma} \frac{\partial \log(1-\gamma)}{\partial \theta}$ for $\theta \in \{\lambda, \mu\}$.

$\kappa = \lambda/\mu$.

$$\frac{\partial \log \kappa}{\partial \lambda} = \frac{1}{\lambda}, \qquad \frac{\partial \log \kappa}{\partial \mu} = -\frac{1}{\mu}$$

**Complementary parameters.** For any parameter $\xi$: $\frac{\partial \log(1-\xi)}{\partial \theta} = -\frac{\xi}{1-\xi} \frac{\partial \log \xi}{\partial \theta}$.

##### A.5.2 Observed-data score

Table A.1: Correspondence between transition counts, complete-data log-likelihood terms, and lineage fates.

| Log-likelihood term | Coefficient | Interpretation |
|---|---|---|
| $\log \alpha$ | $n_{SM} + n_{MM} + n_{DM} + n_{IM}$ | Founder survival (match) |
| $\log(1 - \alpha)$ | $n_{SD} + n_{MD} + n_{DD} + n_{ID}$ | Founder death (delete) |
| $\log(1 - \gamma)$ | $n_{DM} + n_{DD} + n_{DE}$ | Lineage extinction (no `D` $\to$ `I`) |
| $\log \gamma$ | $n_{DI}$ | Youngest orphan (`D` $\to$ `I`) |
| $\log \beta$ | $n_{SI} + n_{MI} + n_{II}$ | Further offspring (insertions continue) |
| $\log(1 - \beta)$ | $n_{IM} + n_{ID} + n_{IE}$ | No further offspring (insertions end) |

Using the transition count decomposition from Table A.1, the Pair HMM log-likelihood under the *conditional* model $P(y|x)$ is

$$\ell_{\text{cond}} = n_{\log \alpha} \log \alpha + n_{\log(1-\alpha)} \log(1-\alpha) + n_{\log \beta} \log \beta + n_{\log(1-\beta)} \log(1-\beta) + n_{\log \gamma} \log \gamma + n_{\log(1-\gamma)} \log(1-\gamma)$$

where we have suppressed the $\kappa$ and $(1 - \kappa)$ terms since they cancel in the conditional likelihood (they appear symmetrically in the transition probabilities and the ancestral sequence prior).

The score with respect to $\lambda$ is

$$\frac{\partial \ell_{\text{cond}}}{\partial \lambda} = \sum_{\xi} n_{\xi} \frac{\partial \xi}{\partial \lambda}$$

where $\xi$ ranges over $\{\log \alpha, \log(1-\alpha), \log \beta, \log(1-\beta), \log \gamma, \log(1-\gamma)\}$, and similarly for $\mu$.

##### A.5.3 Joint likelihood correction

For the *joint* model $P(x, y)$, the log-likelihood includes the ancestral sequence prior

$$\ell_{\text{joint}} = \ell_{\text{cond}} + |x| \log \kappa + \log(1 - \kappa) + \sum_i n_i \log \pi_i$$

where $|x|$ is the ancestor length, $n_i$ counts character $i$ in the ancestor, and the $\kappa/1-\kappa$ terms come from the geometric length distribution. The additional score contributions are

$$\frac{\partial}{\partial \lambda}\left[|x| \log \kappa + \log(1-\kappa)\right] = \frac{|x|}{\lambda} - \frac{1}{\mu - \lambda}, \qquad \frac{\partial}{\partial \mu}\left[|x| \log \kappa + \log(1-\kappa)\right] = -\frac{|x|}{\mu} + \frac{1}{\mu - \lambda}$$

##### A.5.4 Recovering sufficient statistics and M-step

The gradients $\partial \ell / \partial \lambda$ and $\partial \ell / \partial \mu$ relate to the BDI sufficient statistics via

$$\frac{\partial \ell}{\partial \lambda} = \frac{\mathbb{E}[B]}{\lambda} - \mathbb{E}[S] - T, \qquad \frac{\partial \ell}{\partial \mu} = \frac{\mathbb{E}[D]}{\mu} - \mathbb{E}[S]$$

Combined with the conservation law (A.12), the closed-form M-step updates are $\lambda_{\text{new}} = \mathbb{E}[B]/\mathbb{E}[S]$ and $\mu_{\text{new}} = \mathbb{E}[D]/\mathbb{E}[S]$.

Alternatively, gradient ascent can be performed directly using the score, without needing to recover the individual sufficient statistics.

#### A.6 General BDI Sufficient Statistics

We derive the endpoint-conditioned sufficient statistics for the general linear BDI process with birth rate $\lambda$, death rate $\mu$, and immigration rate $\nu$ (not necessarily equal to $\lambda$), conditioned on $X(0) = i$ and $X(T) = j$.

##### A.6.1 Complete-data log-likelihood

The complete-data log-likelihood for a path is

$$\log P(\mathbb{X}) = B_{\text{ind}} \log \lambda + B_{\text{imm}} \log \nu + D \log \mu - (\lambda + \mu) S - \nu T + c$$

with sufficient statistics $B_{\text{ind}}$ (individual births), $B_{\text{imm}}$ (immigrations), $D$ (deaths), and $S = \int_0^T X(t)\, dt$ (time-integrated population).

##### A.6.2 Score equations

Differentiating with respect to $(\lambda, \mu, \nu)$:

$$\begin{aligned}
\frac{\partial}{\partial \lambda} \log P_{ij} &= \frac{\mathbb{E}[B_{\text{ind}}]}{\lambda} - \mathbb{E}[S] \\
\frac{\partial}{\partial \mu} \log P_{ij} &= \frac{\mathbb{E}[D]}{\mu} - \mathbb{E}[S] \\
\frac{\partial}{\partial \nu} \log P_{ij} &= \frac{\mathbb{E}[B_{\text{imm}}]}{\nu} - T
\end{aligned}$$

##### A.6.3 Conservation law

Since population changes only by births and deaths, $i + B_{\text{ind}} + B_{\text{imm}} - D = j$, giving

$$\mathbb{E}[B_{\text{ind}}] + \mathbb{E}[B_{\text{imm}}] - \mathbb{E}[D] = j - i \tag{A.62}$$

##### A.6.4 Closed-form solution

Solving the system of four equations (three score equations plus conservation) in four unknowns, using $\lambda \neq \mu$:

$$\begin{aligned}
\mathbb{E}[B_{\text{imm}}] &= \nu \frac{\partial}{\partial \nu} \log P_{ij} + \nu T \\
\mathbb{E}[S] &= \frac{j - i + \mu \frac{\partial}{\partial \mu} \log P_{ij} - \lambda \frac{\partial}{\partial \lambda} \log P_{ij} - \nu \frac{\partial}{\partial \nu} \log P_{ij} - \nu T}{\lambda - \mu} \\
\mathbb{E}[B_{\text{ind}}] &= \lambda \frac{\partial}{\partial \lambda} \log P_{ij} + \lambda \, \mathbb{E}[S] \\
\mathbb{E}[D] &= \mu \frac{\partial}{\partial \mu} \log P_{ij} + \mu \, \mathbb{E}[S]
\end{aligned}$$

##### A.6.5 Transition probability for general BDI

For the general linear BDI process with per-capita birth rate $\lambda$, per-capita death rate $\mu$, and constant immigration rate $\nu$ (where $\nu$ is a free parameter, not necessarily 0 or $\lambda$), the transition probability $P_{ij}(T) = P(X(T) = j \mid X(0) = i)$ is known in closed form (29).

Using the $\alpha$ and $\beta$ from Section A.1.3, define $\rho = \nu/\lambda$ (the immigration-to-birth ratio). The transition probability factorizes as:

$$P_{ij}(T) = \underbrace{P_{ij}^{(0)}(T)}_{\text{no-immigration}} \cdot \underbrace{R_{ij}(T)}_{\text{immigration factor}} \tag{A.63}$$

The no-immigration factor is

$$P_{ij}^{(0)}(T) = \sum_{k=0}^{\min(i,j)} \binom{i}{k} \binom{j-1}{k-1} \alpha^k (1-\alpha)^{i-k} (1-\gamma)^{i-k} \gamma^{j-k} (1-\beta)^{k+1} \beta^{j-k-1} \cdot \delta_{j>0} \tag{A.64}$$

(with the $k = 0$ term understood as $(1-\alpha)^i (1-\gamma)^i$ when $j = 0$, and the convention $\binom{-1}{-1} = 1$).

For the immigration factor, when $\nu \neq \lambda$ (i.e. $\rho \neq 1$):

$$R_{ij}(T) = (1-\beta)^\rho \cdot {}_2F_1\left(-j, \rho; 1; \frac{\beta}{1-\beta} \cdot \frac{1}{\beta/(1-\beta)}\right) \tag{A.65}$$

More explicitly, the full $P_{ij}(T)$ with immigration can be written using the negative binomial form (29):

$$P_{ij}(T) = \sum_{k=0}^{\min(i,j)} \binom{i}{k} \binom{j-1}{k-1} \alpha^k (1-\alpha-\gamma+\alpha\gamma)^{i-k} \gamma^{j-k} (1-\beta)^{k+1} \beta^{j-k-1} \cdot (1-\beta)^\rho \sum_{m=0}^{j-k} \binom{\rho+m-1}{m} \beta^m \tag{A.66}$$

where the inner sum is a truncated negative binomial in $\beta$ with parameter $\rho$. When $\rho = 1$ (i.e. $\nu = \lambda$), the immigration factor simplifies to a factor of $(1-\beta)$, recovering the TKF91 transition probabilities. When $\rho = 0$ (no immigration), $R = 1$ and we recover $P_{ij}^{(0)}$.

##### A.6.6 Score derivatives for $\nu$

The score with respect to $\nu$ is

$$\frac{\partial}{\partial \nu} \log P_{ij} = \frac{1}{P_{ij}} \frac{\partial P_{ij}}{\partial \nu}$$

Since $\nu$ enters through $\rho = \nu/\lambda$ and the factor $(1-\beta)^\rho$ times the negative binomial sum, we have

$$\frac{\partial}{\partial \nu} \log P_{ij} = \frac{1}{\lambda} \left( \log(1-\beta) + \psi(\rho) \cdot [\text{correction from the sum}] \right) \tag{A.67}$$

where $\psi$ is the digamma function (arising from $\partial/\partial\rho \binom{\rho+m-1}{m}$). The full expression is:

$$\frac{\partial}{\partial \nu} \log P_{ij} = \frac{1}{\lambda} \frac{\sum_k (\cdots) \sum_m \binom{\rho+m-1}{m} \beta^m \left[ \log(1-\beta) + \sum_{\ell=0}^{m-1} \frac{1}{\rho+\ell} \right]}{\sum_k (\cdots) \sum_m \binom{\rho+m-1}{m} \beta^m} \tag{A.68}$$

where $(\cdots)$ abbreviates the terms from the outer sum in (A.66).

##### A.6.7 Chain rule

In practice, automatic differentiation of (A.66) is the preferred approach for computing $\partial \log P_{ij}/\partial\lambda$, $\partial \log P_{ij}/\partial\mu$, and $\partial \log P_{ij}/\partial\nu$, since the nested sums and special functions make manual differentiation unwieldy.

For completeness, the chain rule decomposes as:

$$\frac{\partial \log P_{ij}}{\partial \lambda} = \frac{\partial \log P_{ij}}{\partial \alpha}\frac{\partial \alpha}{\partial \lambda} + \frac{\partial \log P_{ij}}{\partial \beta}\frac{\partial \beta}{\partial \lambda} + \frac{\partial \log P_{ij}}{\partial \gamma}\frac{\partial \gamma}{\partial \lambda} + \frac{\partial \log P_{ij}}{\partial \rho}\frac{\partial \rho}{\partial \lambda}$$

with the $\alpha$, $\beta$, $\gamma$ derivatives given in Appendix A.5, and the additional term

$$\frac{\partial \rho}{\partial \lambda} = -\frac{\nu}{\lambda^2}, \qquad \frac{\partial \rho}{\partial \nu} = \frac{1}{\lambda}$$

The derivative $\partial \log P_{ij}/\partial\rho$ is the quantity in (A.68) times $\lambda$. The $\mu$ derivative has no $\rho$ term since $\rho$ does not depend on $\mu$.

The formulas above hold for any $(i, j, \lambda, \mu, \nu)$ with $\lambda \neq \mu$ and $P_{ij}(T) > 0$. In the TKF91 regime ($\nu = \lambda$, $i \in \{0, 1\}$), they specialize to the results of the main text.

#### A.7 The General Geometric model

Ideally, the underlying evolutionary model we'd use for indels would be the General Geometric Indel (GGI) model (11, 23). The GGI model is, in some sense, the maximum-entropy indel model for a given expected rate and length of indels. Unfortunately, the probability distributions of aligned ancestor-descendant sequences at finite times under this model are hard to solve. It turns out that the pairwise alignment distributions under the GGI model are well approximated by the TKF92 model (51), so we use that model for inference. In this section we discuss the relationship between the models and, in particular, how we can map parameters between them in a principled way.

The GGI model has parameters $(\lambda_0, \mu_0, x, y)$. In this model, the instantaneous rate of insertion of $k$ residues at any available site is $\lambda_0 x^{k-1}(1-x)$ (with the residues themselves sampled from a stationary distribution), while the instantaneous rate of deletion of a stretch of exactly $k$ contiguous residues is $\mu_0 y^{k-1}(1-y)$. For reversibility (or, indeed, the existence of an equilibrium) we require that $\lambda_0 y(1-x) = \mu_0 x(1-y)$. The mean insertion and deletion lengths are $1/(1-x)$ and $1/(1-y)$.

The rate at which any given residue is deleted, counting deletions that start at or before that residue (in an infinitely long sequence), is $\mu_0/(1-y)$. The probability that a sequence has length $n$ at equilibrium is $(x/y)^n(1-x/y)$, so the expected sequence length at equilibrium is $\ell_{\mathrm{GGI}} = \frac{x/y}{1-x/y} = \frac{1}{y/x-1}$.

The TKF92 model has parameters $(\lambda, \mu, r)$. In this model, $k$-residue fragments (where $k > 0$ and $P(k) = r^{k-1}(1-r)$) are inserted and deleted at rates $\lambda, \mu$. The rate at which any given residue is deleted is just the rate at which its fragment is deleted, $\mu$. The mean number of fragments at equilibrium is $\frac{\lambda/\mu}{1-\lambda/\mu} = \frac{1}{\mu/\lambda-1}$ and the mean fragment length is $1/(1-r)$, so the mean sequence length at equilibrium is $\ell_{\mathrm{TKF}} = \frac{1}{(\mu/\lambda-1)(1-r)}$. The precise distribution of sequence lengths at equilibrium is a zero-altered geometric distribution

$$P(n) = \begin{cases} 1 - \frac{\lambda}{\mu} & \text{if } n = 0 \\ \frac{\lambda}{\mu}\left(r + (1-r)\frac{\lambda}{\mu}\right)^{n-1}(1-r)\left(1-\frac{\lambda}{\mu}\right) & \text{if } n > 0 \end{cases}$$

The mean length of an insertion in TKF92 is $1/(1-r)$, i.e. the expected fragment length. For deletions, things are slightly more complicated. Superficially, it looks like deletions are also fragments so they should have the same mean length. However, if we do not have knowledge of the fragment boundaries, but are conditioning just on the current sequence length (as in GGI), we have to weight this by the posterior probability that the subsequence being deleted does in fact constitute a single fragment. For a subsequence of length $k$, this posterior probability is proportional to $\left(\frac{r}{r+(1-r)\frac{\lambda}{\mu}}\right)^{k-1}$ (indels at the end of the sequence have a slightly different correction factor since they are more likely to constitute a fragment, but the $k$-dependence is the same). The rate of starting an insertion or deletion *at a particular site* is subject to a similar correction factor.

Given the equilibrium constraint on GGI, each model has three free parameters. In order to find a correspondence, it is useful to match moments (expectations). In view of the above discussion, the most natural expectations to equate are (in the form "GGI expectation = TKF92 expectation")

- The mean deletion rate of a given residue: $\mu_0/(1-y) = \mu$
- The mean length at equilibrium: $\frac{1}{y/x-1} = \frac{1}{(\mu/\lambda-1)(1-r)}$
- The mean length of an insertion event: $1/(1-x) = 1/(1-r)$ so $x = r$.

These equations, together with the reversibility constraint $\lambda_0 y(1-x) = \mu_0 x(1-y)$, fully specify a one-to-one mapping between GGI and TKF92. In what follows, we will simply use the TKF92 parameters $(\lambda, \mu, r)$ and likelihoods (including the zero-adjusted equilibrium length distribution), but with the understanding that we are approximating GGI. In particular, when we jointly sample the pairwise alignments around any given node in the phylogenetic tree, we will *not* require the fragment boundaries to coincide (as would be the case in TKF92). Technically speaking, any indel model parameter priors should be on the GGI parameters rather than the TKF92 ones, but since we are using uninformative priors this may be an acceptable compromise.

One way in which the TKF92 model, where the process state is augmented with latent fragment boundaries, deviates from the sequence-only GGI is that when we use the TKF92 Pair HMM (with the fragment boundaries marginalized out) as an approximation to $P(X_t|X_0)$, we cannot expect it perfectly to obey the Chapman-Kolmogorov equation $P(X_{t+u} = x''|X_0 = x) = \sum_{x'} P(X_t = x'|X_0 = x)P(X_{t+u} = x''|X_t = x')$. This has consequences for any MCMC moves that split branches using a Pair HMM (which models exactly the Chapman-Kolmogorov equation), since it means that those moves will be miscalibrated in probability, but the approximation should be mild. In fact, it seems possible that no five-state ($\{\mathtt{S}, \mathtt{M}, \mathtt{I}, \mathtt{D}, \mathtt{E}\}$) Pair HMM exactly obeys the Chapman-Kolmogorov equation for a GGI model, except in the special case where the model reduces to the TKF91 model. The best approximations known are obtained by matching moments of the underlying counting process and thereby deriving ODEs for the transition probabilities (23). However, the TKF92 model is a close runner-up (and actually appears to fit real data better).

### B EM, composite likelihoods, and variational inference

Building on Appendix A, this appendix develops the M-step and variational-inference machinery for TKF91 / TKF92 / MixDom at full generality. First, closed-form GTR substitution M-steps in their many specialisations (JC69, K80, F81, HKY85, GTR, GY94, plus rate-rescaling, tied-equilibria, and mixture-of-GTR variants); then the stochastic variational Baum–Welch (SVI-BW) loop with its convergence theorem (ELBO derivation, natural-gradient updates, minibatch variance bounds), the linearised analysis at stationarity, Maraschino as the cherry-count distillation of TKF92, the tree-level inference algorithms it composes with (FSA, BeamASR, VarAnc, svi-VarAnc), the mixture-of-trees variational ancestral presence/absence inference, and the structural-bias analysis of the BP cumulant under a column-factorised variational field.

#### B.1 Substitution M-Steps for Specific Models

This appendix specializes the general GTR substitution M-step (Section A.1.8) to specific DNA and codon substitution models. In each case, we state the rate matrix $R$ in terms of the model parameters, derive the closed-form MLE (or MAP) update given the Holmes–Rubin sufficient statistics $(W_i, U_{ij}, V_i)$, and note the spectral structure when it simplifies the endpoint-conditioned integrals (A.5).

Throughout, we work with the complete-data log-likelihood (A.1)

$$\ell_2 = \sum_i V'_i \log \pi_i + \sum_{j>i} (U_{ij} + U_{ji}) \log Q_{ij} - \sum_{j>i} Q_{ij}(W_i \pi_j + W_j \pi_i)$$

where $V'_i = V_i + \sum_{j \neq i} U_{ji}$, under the GTR parameterization $R_{ij} = Q_{ij}\pi_j$ with symmetric exchangeability $Q_{ij} = Q_{ji}$.

The indel M-step (quadratic (A.22) for $\kappa$, with L'Hôpital limits in Section A.4) is orthogonal and unchanged across all substitution models.

For DNA models ($|\mathcal{A}| = 4$), the Holmes–Rubin integral $I^{ab}_{ij}(T)$ (A.4) involves only $4 \times 4 = 16$ matrix entries per endpoint pair, and the $J^{kl}(T)$ matrix (A.5) is $4 \times 4$, making the endpoint-conditioned expectations cheap to compute.

##### B.1.1 JC69 (Jukes–Cantor)

**Rate matrix.** All off-diagonal rates are equal: $R_{ij} = \rho/3$ for $i \neq j$ (total rate out of any state is $\rho$). Equivalently, $\pi_i = 1/4$ for all $i$ and $Q_{ij} = 4\rho/3$ for all $i \neq j$.

The single free parameter is the overall rate $\rho$.

**M-step.** By detailed balance, $Q_{ij} = (U_{ij} + U_{ji})/(W_i \pi_j + W_j \pi_i)$ is the GTR MLE. Under JC69, all exchangeabilities are forced equal, so we pool:

$$\hat{Q} = \frac{\sum_{j>i}(U_{ij} + U_{ji})}{\sum_{j>i}(W_i \pi_j + W_j \pi_i)} = \frac{U_\bullet}{3W_\bullet/4} = \frac{4U_\bullet}{3W_\bullet} \tag{B.1}$$

where $U_\bullet = \sum_{i \neq j} U_{ij}$ is the total substitution count and $W_\bullet = \sum_i W_i$ is the total dwell time. (The denominator uses $\sum_{j>i}(W_i/4 + W_j/4) = 3W_\bullet/4$ since each of the 4 states appears in 3 of the $\binom{4}{2} = 6$ pairs.) The overall rate is then $\hat{\rho} = 3\hat{Q}/4 = U_\bullet/W_\bullet$, which is simply the total substitution count divided by total dwell time.

The equilibrium frequencies are fixed: $\hat{\pi}_i = 1/4$.

**Spectral structure.** $S = (\rho/3)J - (4\rho/3)\, I$ where $J$ is the $4 \times 4$ all-ones matrix. Eigenvalues: 0 (once, eigenvector $\mathbf{1}/2$) and $-4\rho/3$ (triply degenerate). Thus $J^{kl}(T)$ has only two distinct diagonal values ($T$ and $Te^{-4\rho T/3}$) and the off-diagonal $J^{01}$ term $(1 - e^{-4\rho T/3})/(4\rho/3)$.

##### B.1.2 K80 (Kimura 2-Parameter)

**Rate matrix.** Equal equilibrium frequencies $\pi_i = 1/4$. Transitions (purine↔purine, pyrimidine↔pyrimidine) occur at rate $\kappa_s$ and transversions at rate 1 (or vice versa, depending on convention). Labeling the states A, G, C, T and writing $i \sim j$ for a transition pair:

$$R_{ij} = \begin{cases} \kappa_s/4 & \text{if } i \sim j \text{ (transition)} \\ 1/4 & \text{if } i \not\sim j \text{ (transversion)} \end{cases}$$

with total rate out of any state $(\kappa_s + 2)/4$. The single free parameter beyond rate scaling is $\kappa_s$ (the transition/transversion rate ratio).

**M-step.** Partition the substitution counts into transitions $U_s = \sum_{i \sim j} U_{ij}$ and transversions $U_v = \sum_{i \not\sim j} U_{ij} = U_\bullet - U_s$. Similarly partition the dwell-weighted opportunities. Under K80, the exchangeability for transition pairs is $\kappa_s$ and for transversion pairs is 1 (up to a common scale). The MLE for each group, from the pooled GTR formula:

$$\hat{\kappa}_s = \frac{U_s/N_s}{U_v/N_v} \tag{B.2}$$

where $N_s = \sum_{i \sim j,\, i<j}(W_i \pi_j + W_j \pi_i) = W_\bullet/4$ is the opportunity for transitions (2 pairs: A–G and C–T, each contributing $(W_i + W_j)/4$, summing to $W_\bullet/4$) and $N_v = \sum_{i \not\sim j,\, i<j}(W_i \pi_j + W_j \pi_i) = W_\bullet/2$ is the opportunity for transversions (4 pairs, each state appearing in 2 transversion pairs, summing to $2W_\bullet/4 = W_\bullet/2$). Simplifying:

$$\hat{\kappa}_s = \frac{2\, U_s}{U_v} \tag{B.3}$$

Given $\hat{\kappa}_s$, the overall scale $\hat{\sigma}$ satisfies $\hat{\sigma}(\hat{\kappa}_s N_s + N_v) = U_\bullet$, giving overall rate $\hat{\rho} = \hat{\sigma}(\hat{\kappa}_s + 2)/4 = U_\bullet/W_\bullet$.

**Spectral structure.** Three distinct eigenvalues: 0, $-(\kappa_s + 1)/2$ (doubly degenerate, within-purine and within-pyrimidine contrasts), and $-1$ (purine-vs-pyrimidine contrast).

##### B.1.3 F81 (Felsenstein 1981)

**Rate matrix.** Equal exchangeabilities, variable equilibrium: $R_{ij} = \rho\, \pi_j$ for $i \neq j$ (equivalently, $Q_{ij} = \rho$ for all $i \neq j$).

**M-step.** Since $Q_{ij} = \rho$ is constant across all pairs, the exchangeability MLE pools all pairs:

$$\hat{\rho} = \frac{U_\bullet}{\sum_{j>i}(W_i\pi_j + W_j\pi_i)} = \frac{U_\bullet}{\sum_i W_i(1-\pi_i)} \tag{B.4}$$

using the identity $\sum_{j>i}(W_i\pi_j + W_j\pi_i) = \sum_i W_i \sum_{j\neq i}\pi_j = \sum_i W_i(1-\pi_i)$.

For equilibrium frequencies:

$$\hat{\pi}_i = \frac{V_i'}{\sum_j V_j'} \tag{B.5}$$

This is the standard empirical-frequency estimator (exact MLE when $Q$ is held fixed; approximate when $Q$ and $\pi$ are jointly optimized, as discussed in Section A.1.8).

In practice, iterate: fix $\hat{\pi}$, solve for $\hat{\rho}$ (B.4), update $\hat{\pi}$ (B.5), repeat. The decoupling is exact after one iteration when $W_i \propto \pi_i$ (which holds approximately for long dwell times).

**Spectral structure.** $S = \rho(\sqrt{\pi}\sqrt{\pi}^\top - I)$ where $\sqrt{\pi}$ is the vector with entries $\sqrt{\pi_i}$. Eigenvalues: 0 (once) and $-\rho$ ($|\mathcal{A}|-1$ times, degenerate). The $J^{kl}$ matrix has the same simple structure as JC69.

##### B.1.4 HKY85 (Hasegawa–Kishino–Yano)

**Rate matrix.** Variable equilibrium frequencies $\pi$ and a transition/transversion ratio $\kappa_s$:

$$R_{ij} = \begin{cases} \kappa_s\,\pi_j & \text{if } i \sim j \text{ (transition)} \\ \pi_j & \text{if } i \not\sim j \text{ (transversion)} \end{cases}$$

Equivalently, $Q_{ij} = \kappa_s$ for transition pairs and $Q_{ij} = 1$ for transversion pairs. This is K80 + F81: the $\kappa_s$ structure of K80 with the variable-$\pi$ structure of F81.

**M-step.** Pool exchangeabilities within each class:

$$\hat{\kappa}_s = \frac{U_s \,/\, N_s}{U_v \,/\, N_v} \tag{B.6}$$

where now (unlike K80) the opportunities depend on $\pi$:

$$N_s = \sum_{i\sim j,\, i<j} (W_i\pi_j + W_j\pi_i) \tag{B.7}$$

$$N_v = \sum_{i\not\sim j,\, i<j} (W_i\pi_j + W_j\pi_i) \tag{B.8}$$

With $U_s$ and $U_v$ as in K80 (summed over directed transition and transversion pairs respectively).

For $\pi$, use $\hat{\pi}_i = V_i' / \sum_j V_j'$ as in F81, then iterate with $\kappa_s$ if desired.

**Spectral structure.** Let $\pi_R = \pi_A + \pi_G$ and $\pi_Y = \pi_C + \pi_T$. The eigenvalues are 0, $-(\pi_R + \kappa_s\pi_Y)$ (within-pyrimidine contrast), $-(\pi_Y + \kappa_s\pi_R)$ (within-purine contrast), and $-1$ (purine-vs-pyrimidine contrast). Generically four distinct values; when $\pi_R = \pi_Y = 1/2$ (i.e. K80), the two within-class eigenvalues merge to $-(\kappa_s + 1)/2$.

##### B.1.5 GTR (General Time-Reversible)

For completeness, we restate the general case from Section A.1.8.

**Rate matrix.** $R_{ij} = Q_{ij}\,\pi_j$ with $Q_{ij} = Q_{ji} \geq 0$ for $i \neq j$. For DNA, $Q$ has 6 free parameters (upper triangle) and $\pi$ has 3 free parameters (4 frequencies summing to 1).

**M-step.** Fixing $\pi$:

$$\hat{Q}_{ij} = \frac{U_{ij} + U_{ji}}{W_i\pi_j + W_j\pi_i} \tag{B.9}$$

Fixing $Q$, the $\pi$ update requires solving a nonlinear system (Section A.1.8). In practice, set $\hat{\pi}_i \propto V_i'$ and iterate with (B.9).

**Spectral structure.** $S_{ij} = Q_{ij}\sqrt{\pi_i\pi_j}$ is real symmetric with eigenvalues $0 = \xi^{(0)} > \xi^{(1)} \geq \cdots \geq \xi^{(|\mathcal{A}|-1)}$ (generically all distinct for DNA). No further simplification beyond the general formulas (A.5).

##### B.1.6 GY94 (Goldman–Yang Codon Model)

**Rate matrix.** The state space is the 61 sense codons ($|\mathcal{A}| = 61$, excluding stop codons). The instantaneous rate from codon $i$ to codon $j$ is:

$$R_{ij} = \begin{cases} 0 & \text{if } i \text{ and } j \text{ differ at more than one position} \\ \pi_j & \text{if synonymous transversion} \\ \kappa_s\,\pi_j & \text{if synonymous transition} \\ \omega\,\pi_j & \text{if nonsynonymous transversion} \\ \omega\,\kappa_s\,\pi_j & \text{if nonsynonymous transition} \end{cases}$$

where $\omega$ is the nonsynonymous/synonymous rate ratio (dN/dS), $\kappa_s$ is the transition/transversion ratio, and $\pi_j$ is the equilibrium frequency of codon $j$. This is a reversible model with exchangeability

$$Q_{ij} = \begin{cases} 0 & \text{(multi-nucleotide change)} \\ 1 & \text{(synonymous transversion)} \\ \kappa_s & \text{(synonymous transition)} \\ \omega & \text{(nonsynonymous transversion)} \\ \omega\kappa_s & \text{(nonsynonymous transition)} \end{cases}$$

**M-step.** Partition the substitution counts into four classes indexed by (synonymous/nonsynonymous) $\times$ (transition/transversion):

$U_{S,s}$ = total synonymous transition counts    $U_{S,v}$ = total synonymous transversion counts
$U_{N,s}$ = total nonsynonymous transition counts    $U_{N,v}$ = total nonsynonymous transversion counts

and the corresponding dwell-weighted opportunities $N_{S,s}, N_{S,v}, N_{N,s}, N_{N,v}$ defined analogously to (B.7)–(B.8) but summing over single-nucleotide codon pairs in each class.

The exchangeabilities are $Q = 1$ (synonymous transversion), $\kappa_s$ (synonymous transition), $\omega$ (nonsynonymous transversion), $\omega\kappa_s$ (nonsynonymous transition). The complete-data log-likelihood for $(\omega, \kappa_s)$, with $\pi$ fixed, is:

$$\begin{aligned} \ell_2(\omega, \kappa_s) = {} & U_{S,v}\log 1 + U_{S,s}\log\kappa_s + U_{N,v}\log\omega + U_{N,s}\log(\omega\kappa_s) \\ & - N_{S,v} - \kappa_s N_{S,s} - \omega N_{N,v} - \omega\kappa_s N_{N,s} \end{aligned}$$

Setting $\partial\ell_2/\partial\kappa_s = 0$ and $\partial\ell_2/\partial\omega = 0$:

$$\frac{U_{S,s} + U_{N,s}}{\kappa_s} = N_{S,s} + \omega\, N_{N,s} \tag{B.10}$$

$$\frac{U_{N,v} + U_{N,s}}{\omega} = N_{N,v} + \kappa_s\, N_{N,s} \tag{B.11}$$

These are two equations in two unknowns. Substituting (B.11) into (B.10) to eliminate $\omega$:

$$\omega = \frac{U_{N,v} + U_{N,s}}{N_{N,v} + \kappa_s N_{N,s}}$$

$$\frac{U_{S,s} + U_{N,s}}{\kappa_s} = N_{S,s} + \frac{(U_{N,v} + U_{N,s})\, N_{N,s}}{N_{N,v} + \kappa_s N_{N,s}}$$

Multiplying both sides by $\kappa_s(N_{N,v} + \kappa_s N_{N,s})$ and rearranging:

$$N_{S,s}\, N_{N,s}\, \kappa_s^2 + \big(N_{S,s}\, N_{N,v} + (U_{N,v} - U_{S,s})\, N_{N,s}\big)\, \kappa_s - (U_{S,s} + U_{N,s})\, N_{N,v} = 0 \tag{B.12}$$

The positive root gives $\hat\kappa_s$, and then

$$\hat\omega = \frac{U_{N,v} + U_{N,s}}{N_{N,v} + \hat\kappa_s\, N_{N,s}} \tag{B.13}$$

The codon equilibrium frequencies $\pi$ are estimated as $\hat\pi_j \propto V'_j$.

**Spectral structure.** For 61 sense codons, the symmetrized rate matrix is $61 \times 61$ with at most 61 distinct eigenvalues. No closed-form eigendecomposition is known in general for arbitrary $\pi$. When $\pi$ factors as $\pi_{abc} = \pi^{(1)}_a \pi^{(2)}_b \pi^{(3)}_c$ (the F3x4 model), the symmetrized matrix is a sum of Kronecker products and its eigenvectors factor as tensor products of $4 \times 4$ matrices, reducing the eigendecomposition to three $4 \times 4$ problems. In practice, for the full GY94, numerical eigendecomposition of the $61 \times 61$ symmetric matrix is inexpensive ($O(61^3) \approx 2 \times 10^5$ flops) and need only be performed once per M-step.

##### B.1.7 Summary and Practical Notes

| Model | Free params | $\pi$ | M-step for exchangeability |
|---|---|---|---|
| JC69 | 1 ($\rho$) | fixed 1/4 | $\hat\rho = U_\bullet / W_\bullet$ |
| K80 | 1 ($\kappa_s$) | fixed 1/4 | $\hat\kappa_s = 2U_s/U_v$ |
| F81 | 4 ($\rho, \pi$) | $V'_i / \sum V'_j$ | $\hat\rho = U_\bullet / \sum_i W_i(1 - \pi_i)$ |
| HKY85 | 5 ($\kappa_s, \pi$) | $V'_i / \sum V'_j$ | $\hat\kappa_s = (U_s/N_s)/(U_v/N_v)$ |
| GTR | 9 ($Q, \pi$) | $V'_i / \sum V'_j$ | $\hat Q_{ij} = (U_{ij} + U_{ji})/(W_i\pi_j + W_j\pi_i)$ |
| GY94 | 62 ($\omega, \kappa_s, \pi$) | $V'_j / \sum V'_k$ | quadratic in $\kappa_s$ (B.12) |

For all models above, the indel M-step is identical: solve the $\kappa$ quadratic (A.22) for $(\lambda, \mu)$. The substitution and indel parameter blocks decouple completely in the M-step (they share no sufficient statistics).

**MAP estimates.** To obtain MAP estimates, augment the sufficient statistics before applying the MLE formulas: $U_{ij} \to U_{ij} + \alpha_Q - 1$, $W_i \to W_i + \beta_Q$, $V_i \to V_i + \alpha_\pi - 1$, as in Section C.1.4. The independent $\text{Gamma}(\alpha_Q, \beta_Q)$ priors on each $Q_{ij}$ together with a Dirichlet on $\pi$ are conjugate to the *irreversible* CTMC complete-data likelihood (independent $R_{ij}$) but not to the reversible GTR parameterization—the symmetric coupling $Q_{ij} = Q_{ji}$ together with the requirement that the joint complete-data likelihood include the stationary draw $X(0) \sim \pi$ means the proper conjugate prior is the cycle-corrected edge-flow density of Diaconis and Rolles (12), with a Kirchhoff matrix-tree determinant accounting for the cycle structure of the state graph. See the discussion in Section C.1.4 for the form of the Diaconis–Rolles prior and its modification to incorporate the initial-state contribution.The simpler Gamma × Dirichlet pseudocounts used here are non-conjugate regularizers and remain perfectly valid for MAP, with the closed-form augmented M-step above; we adopt them throughout for simplicity.

**Degenerate eigenvalues.** Models with symmetry (JC69, K80, F81) have degenerate eigenvalues in $S$, which means the $J^{kl}(T)$ matrix has repeated diagonal entries. This reduces the number of distinct $J$ values that must be computed, but requires care to use the $\xi^{(k)} = \xi^{(l)}$ branch ($J^{kl} = T\, e^{\xi^{(k)}T}$) rather than the difference quotient, which is a removable 0/0 singularity.

##### B.1.8 Reversible Mixture with Per-Component GTR

We now consider a $C$-component reversible mixture in which each component is a full GTR model with its own exchangeability matrix $Q^{(c)}$ and equilibrium distribution $\pi^{(c)}$:

$$R_{ij}^{(c)} = Q_{ij}^{(c)}\, \pi_j^{(c)} \quad (i \neq j), \qquad c \in \{1, \ldots, C\}. \tag{B.14}$$

Each site is assigned (latently) to one component, and conditional on $c$, the site evolves under the reversible CTMC with rate matrix $R^{(c)}$. This is the parameterisation used by the per-(domain, fragment-type) site-class emissions of MixDom (Section C.1.4), where $c$ is the site-class drawn independently at each position from $u_{nf}$.Because each component is an independent GTR model, the overall rate scale is absorbed into $Q^{(c)}$ itself and no separate rate multiplier is needed; this makes the parameterisation a strict generalisation of per-domain GTR (the per-domain GTR case is recovered when $u_{nf,c} = \mathbf{1}[c = n]$) and of the discrete-gamma rate-across-sites model (54).

**E-step counts.** The component assignment is latent. From the standard E-step, each branch produces an endpoint pair $(a, b)$ with marginal posterior weight $w_{ab}$; conditional on $(a, b)$ at a site with prior class distribution $P(c \mid \text{site}) = u_c$, the component posterior is

$$P(c \mid a, b, T, \text{site}) = \frac{u_c\, \pi_a^{(c)}\, M_{ab}^{(c)}(T)}{\sum_{c'} u_{c'}\, \pi_a^{(c')}\, M_{ab}^{(c')}(T)}, \tag{B.15}$$

with $M^{(c)}(T) = \exp(R^{(c)}T)$. Per-component Holmes–Rubin sufficient statistics are then accumulated by weighting the standard expressions (A.3)–(A.2) by (B.15):

$$V_i^{(c)} = \textstyle\sum_{\text{starts}} w_a\, P(c \mid a, \cdot)\, \mathbf{1}[X(0) = i] \tag{B.16}$$

$$U_{ij}^{(c)} = \textstyle\sum_{\text{branches}} w_{ab}\, P(c \mid a, b, T, \cdot)\, \mathcal{C}_{ij}^{U}\big(a, b, T;\, R^{(c)}\big) \tag{B.17}$$

$$W_i^{(c)} = \textstyle\sum_{\text{branches}} w_{ab}\, P(c \mid a, b, T, \cdot)\, \mathcal{C}_{i}^{W}\big(a, b, T;\, R^{(c)}\big) \tag{B.18}$$

and we write $V_i'^{(c)} = V_i^{(c)} + \sum_{j \neq i} U_{ji}^{(c)}$ as before. Writing $U_\bullet^{(c)} = \sum_{i \neq j} U_{ij}^{(c)}$ for the total per-component substitution count.

**Complete-data log-likelihood.** Because each class is an independent GTR model under this parameterisation, the complete-data log-likelihood decomposes over classes:

$$\ell_2(\{Q^{(c)}\}, \{\pi^{(c)}\}) = \sum_c \ell_2^{(c)}, \tag{B.19}$$

$$\begin{aligned} \ell_2^{(c)} = \sum_i V_i'^{(c)} \log \pi_i^{(c)} + \sum_{j>i} \big(U_{ij}^{(c)} + U_{ji}^{(c)}\big) \log Q_{ij}^{(c)} \\ - \sum_{j>i} Q_{ij}^{(c)} \big(W_i^{(c)} \pi_j^{(c)} + W_j^{(c)} \pi_i^{(c)}\big). \end{aligned} \tag{B.20}$$

**Identifiability.** No gauge to fix: the rate scale of class $c$ lives inside $Q^{(c)}$, and the $\{Q^{(c)}\}$ are uncoupled in both (B.14) and (B.20). Every class can be updated independently.

**Per-class GTR M-step.** Since $\ell_2$ decomposes class by class, the update for class $c$ is exactly the GTR M-step of Section A.1.8 applied to that class's sufficient statistics. Differentiating (B.20) w.r.t. $Q_{ij}^{(c)}$:

$$\frac{\partial \ell_2^{(c)}}{\partial Q_{ij}^{(c)}} = \frac{U_{ij}^{(c)} + U_{ji}^{(c)}}{Q_{ij}^{(c)}} - \big(W_i^{(c)} \pi_j^{(c)} + W_j^{(c)} \pi_i^{(c)}\big) = 0,$$

giving the closed-form update

$$\hat{Q}_{ij}^{(c)} = \frac{U_{ij}^{(c)} + U_{ji}^{(c)}}{W_i^{(c)} \pi_j^{(c)} + W_j^{(c)} \pi_i^{(c)}}. \tag{B.21}$$

The equilibrium update uses the empirical-frequency approximation

$$\hat{\pi}_i^{(c)} \propto V_i'^{(c)}, \tag{B.22}$$

which is exact when $Q_{ij}^{(c)} W_j^{(c)}$ is independent of $i$ (F81 limit) and approximate otherwise, identical to Section A.1.8. No alternation or projection is needed; one pass through each class completes the M-step.

**Connection to standard models.** With $C = 1$, (B.21)–(B.22) reduce exactly to the GTR M-step (B.9). With $u_{nf,c} = \mathbf{1}[c = n]$ so each domain uses its own dedicated class, the model is the per-domain GTR ($Q^{(n)}$) special case; the per-class M-step here reduces to the per-domain M-step of Section A.1.8 applied once per domain. With $\pi^{(c)} \equiv \pi$ and $Q^{(c)} = \gamma_c Q$ for some shared $Q$, the model is the discrete-gamma rate model (54); a constrained M-step that enforces $Q^{(c)} \propto Q$ recovers the shared-exchangeability case as a special case. Free-rate variants (46) are the shared-$Q$ / free-$\gamma_c$ special case.

**MAP estimates.** Per-class regularising priors apply independently to each class. A Dirichlet pseudocount $\alpha_i^{(c)}$ is added to each $V_i^{(c)}$, and a Gamma($a_S, b_S$) prior on each $Q_{ij}^{(c)}$ (e.g. calibrated to LG via $a_S = b_S\, Q_{ij}^{\mathrm{LG}}$) adds $a_S - 1$ to the numerator and $b_S$ to the denominator of (B.21). As above, this independent-Gamma × Dirichlet structure is conjugate only in the irreversible parameterization; it is a non-conjugate regularizer for the reversible GTR mixture component, with the proper conjugate being a per-class Diaconis–Rolles prior (12). We use the simpler form here for the same reasons as the single-component case (Section C.1.4). The MAP objective

$$\mathcal{L} = \ell_2 + \sum_{c,i} (\alpha_i^{(c)} - 1) \log \pi_i^{(c)} + \sum_{c,j>i} \big[(a_S - 1) \log Q_{ij}^{(c)} - b_S Q_{ij}^{(c)}\big] \tag{B.23}$$

decomposes across classes and has a *unique* joint maximum at

$$\hat{Q}_{ij}^{(c)} = \frac{U_{ij}^{(c)} + U_{ji}^{(c)} + a_S - 1}{W_i^{(c)} \pi_j^{(c)} + W_j^{(c)} \pi_i^{(c)} + b_S}, \tag{B.24}$$

with the $\pi^{(c)}$ update obtained by the same bisection as Section A.1.8. Each class is a standalone GTR M-step; no cross-class coupling.

**Hyperparameter sizing under SVI.** Under SVI-BW (B.48) the EMA sufficient statistics are scaled to the full-dataset size $N$, so prior pseudocounts must live on the same scale to retain their relative weight. Parameterising by *effective sample size*, set $\alpha_i^{(c)} = N_\pi \, \pi_i^{\mathrm{LG}}$ and $b_S = N_S$ with $a_S = N_S \, Q_{ij}^{\mathrm{LG}}$. Defaults $N_\pi, N_S \sim 10^2$–$10^3$ (data-independent) give priors that vanish in the data limit but identify the posterior in small batches.

##### B.1.9 Rate Rescaling: M-Step for a Global Scalar Multiplier

In some training scenarios the relative pattern of substitution rates is known (e.g. from a previously-estimated GTR matrix or from an external published matrix such as LG) and only the overall *rate* needs to adapt to the current data. Concretely, we hold both the exchangeability matrix $Q$ *and* the equilibrium distribution $\pi$ fixed, and seek a single scalar multiplier $\sigma > 0$ such that the rescaled rate matrix

$$R_{ij}^{\mathrm{new}} = \sigma \, Q_{ij} \, \pi_j \quad (i \neq j) \tag{B.25}$$

maximises the complete-data log-likelihood $\ell_2$. Equivalently, the new exchangeability is $Q_{ij}^{\mathrm{new}} = \sigma \, Q_{ij}$, so all relative exchangeabilities are preserved and only the overall tempo of the model changes.

**Closed-form $\sigma$.** Substituting $Q_{ij} \to \sigma Q_{ij}$ into the GTR complete-data log-likelihood,

$$\ell_2(\sigma) = \text{const} + \log \sigma \sum_{j>i} (U_{ij} + U_{ji}) - \sigma \sum_{j>i} Q_{ij} \big(W_i \pi_j + W_j \pi_i\big).$$

The terms not involving $\sigma$ (the $V_i' \log \pi_i$ part and $\sum_{j>i} (U_{ij} + U_{ji}) \log Q_{ij}$) are constants under rescaling. Differentiating and setting to zero yields the unique maximiser

$$\hat{\sigma} = \frac{U_\bullet}{D}, \qquad U_\bullet = \sum_{i \neq j} U_{ij}, \qquad D = \sum_{j>i} Q_{ij} \big(W_i \pi_j + W_j \pi_i\big), \tag{B.26}$$

where $U_\bullet$ is the total expected substitution count and $D$ is the total dwell-weighted opportunity for substitution under the fixed shape $(Q, \pi)$. The objective is strictly concave in $\sigma$ on $(0, \infty)$ (the Hessian $-U_\bullet / \sigma^2 < 0$), so (B.26) is the global maximiser.

**Mixture form (per-class).** Applied class by class to the reversible mixture of Section B.1.8, each class $c$ has its own scalar $\sigma_c$ updated independently from its own per-class statistics:

$$\hat{\sigma}_c = \frac{U_\bullet^{(c)}}{D^{(c)}}, \qquad D^{(c)} = \sum_{j>i} Q_{ij}^{(c)} \big(W_i^{(c)} \pi_j^{(c)} + W_j^{(c)} \pi_i^{(c)}\big). \tag{B.27}$$

The shapes $Q^{(c)}$ and $\pi^{(c)}$ are held at their current values and only the per-class rate scale $\sigma_c$ moves; the class-decomposition (B.19) ensures classes do not couple. A common-rate variant (a single shared $\hat{\sigma}$ across all classes) is recovered by pooling: $\hat{\sigma} = \sum_c U_\bullet^{(c)} / \sum_c D^{(c)}$. The per-class form (B.27) attains a strictly higher $\ell_2$ unless all $\sigma_c$ already coincide.

**Aggregation of counts.** The only sufficient statistics required are $U_\bullet^{(c)} = \sum_{i \neq j} U_{ij}^{(c)}$ (a scalar per class) and the dwell–frequency contraction $D^{(c)}$, which is itself a scalar inner product of the fixed quantity $Q_{ij}^{(c)}(W_i^{(c)} \pi_j^{(c)} + W_j^{(c)} \pi_i^{(c)})$ against the per-pair count. Both are linear in the underlying per-pair $(W, U)$ statistics, so under SVI-BW they accumulate via the same EMA used for the standard M-step (no special handling).

**MAP estimate.** With a Gamma$(a_\sigma, b_\sigma)$ prior on $\sigma$, the augmented M-step is

$$\hat{\sigma} = \frac{U_\bullet + a_\sigma - 1}{D + b_\sigma}. \tag{B.28}$$

The Gamma prior on the global rate is conjugate (the model is linear in $\sigma$), in contrast to the non-conjugate prior on the full-shape M-step. In the rate-rescaling regime the standard $(N_\pi, N_S)$ pseudocounts on $(Q, \pi)$ are inactive (those parameters are frozen); $(a_\sigma, b_\sigma)$ centred at $\sigma = 1$ (e.g. $a_\sigma = b_\sigma = N_\sigma$ for some effective sample size $N_\sigma$) regularises the rate towards the current frozen scale.

**When to use.** Rate rescaling is a useful intermediate regime between fully-frozen substitution parameters (no M-step) and the full-shape mixture M-step of Section B.1.8. It is particularly natural when warm-starting from a published matrix and one wishes to absorb a calibration drift in the time units of the data *without* re-estimating the relative chemistry. In an alternating schedule (e.g. alternate $\sigma_c$ updates with tied-$\pi$ (B.42) updates) the rescaling step contributes a single guaranteed $\ell_2$ ascent per class per iteration, and the alternation does not destabilise the joint objective because each step is an exact maximisation in its own coordinate.

##### B.1.10 Joint Rate Rescaling and Equilibrium

A natural extension of the rate-rescaling regime (Section B.1.9) frees the equilibrium distribution $\pi$ jointly with the scale $\sigma$ while keeping the exchangeability shape $Q$ fixed. This matches the substitution degrees of freedom of a Maraschino-style fit with `--freeze-class-S-shape` (per-class $\sigma_c$ and $\pi^{(c)}$ free, $Q^{(c)}$ shape locked at LG), and is the natural SVI–BW counterpart when warm-starting from such a fit: the previous `rescaling-rates` mode artificially froze $\pi^{(c)}$ at the warm-start values, dropping $A - 1$ degrees of freedom per class.

**Joint complete-data log-likelihood.** With $R_{ij} = \sigma\, Q_{ij}\, \pi_j$ and $Q$ frozen, the complete-data log-likelihood as a function of $(\sigma, \pi)$ is

$$\ell_2(\sigma, \pi) = \sum_i V_i' \log \pi_i + U_\bullet \log \sigma - \sigma \sum_{j>i} Q_{ij}\big(W_i \pi_j + W_j \pi_i\big) + \text{const}, \tag{B.29}$$

where the constant collects the frozen $\sum_{j>i}(U_{ij} + U_{ji}) \log Q_{ij}$ term. Using $Q_{ij} = Q_{ji}$ and $Q_{ii} = 0$, the dwell penalty contracts as

$$\sum_{j>i} Q_{ij}\big(W_i \pi_j + W_j \pi_i\big) = \sum_b \pi_b\, r_b, \qquad r_b := \sum_{a \neq b} Q_{ab} W_a = (QW)_b,$$

so $r_b$ depends only on the frozen quantities $(Q, W)$ and is constant under variation of $(\sigma, \pi)$. Equation (B.29) simplifies to

$$\ell_2(\sigma, \pi) = \sum_b V_b' \log \pi_b + U_\bullet \log \sigma - \sigma\, \mathbf{r}^\top \pi + \text{const}. \tag{B.30}$$

**Stationary conditions.** Lagrangian for $\sum_b \pi_b = 1$ with multiplier $\lambda$:

$$\mathcal{L}(\sigma, \pi, \lambda) = \ell_2(\sigma, \pi) - \lambda\Big(\sum_b \pi_b - 1\Big).$$

The first-order conditions are

$$\frac{\partial \mathcal{L}}{\partial \sigma} = \frac{U_\bullet}{\sigma} - \mathbf{r}^\top \pi = 0 \implies \sigma = \frac{U_\bullet}{\mathbf{r}^\top \pi}, \tag{B.31}$$

$$\frac{\partial \mathcal{L}}{\partial \pi_b} = \frac{V_b'}{\pi_b} - \sigma r_b - \lambda = 0 \implies \pi_b = \frac{V_b'}{\lambda + \sigma r_b}. \tag{B.32}$$

**The Lagrange multiplier is fixed.** A clean structural property obtains by multiplying (B.32) through by $\pi_b$ and summing:

$$\sum_b V_b' = \lambda \sum_b \pi_b + \sigma\, \mathbf{r}^\top \pi = \lambda + \sigma\, \mathbf{r}^\top \pi.$$

The $\sigma$-FOC (B.31) sets $\sigma\, \mathbf{r}^\top \pi = U_\bullet$, so

$$N \;=\; \lambda + U_\bullet, \qquad N := \sum_b V_b' = V_\bullet + U_\bullet,$$

which gives

$$\boxed{\lambda \;=\; V_\bullet,} \tag{B.33}$$

*independent of* $\sigma$: the multiplier is fully determined by the boundary count $V_\bullet = \sum_i V_i$.

**Closed-form $\pi(\sigma)$ and 1-D root in $\sigma$.** Substituting (B.33) into (B.32),

$$\hat{\pi}_b(\sigma) = \frac{V_b'}{V_\bullet + \sigma\, r_b}. \tag{B.34}$$

Note $\sum_b \hat{\pi}_b(\sigma) = 1$ at $\sigma$ satisfying the $\sigma$-FOC, automatically. Plugging back into the $\sigma$-FOC (B.31), $\sigma$ satisfies the single one-dimensional equation

$$g(\sigma) := \sum_b \frac{\sigma\, r_b\, V_b'}{V_\bullet + \sigma\, r_b} \;-\; U_\bullet \;=\; 0, \qquad \sigma > 0. \tag{B.35}$$

**Existence and uniqueness.** $g$ is well-defined and smooth for $\sigma > 0$ (denominators are positive when $V_\bullet > 0$ or $r_b > 0$). Boundary behaviour:

$$g(0) = -U_\bullet \le 0, \qquad g(\sigma) \xrightarrow[\sigma\to\infty]{} \sum_b V'_b - U_\bullet = V_\bullet \ge 0,$$

and the derivative is strictly positive,

$$g'(\sigma) = \sum_b \frac{r_b V'_b V_\bullet}{(V_\bullet + \sigma r_b)^2} > 0 \quad \text{whenever some } r_b V'_b > 0.$$

Hence $g$ is strictly increasing and crosses zero exactly once at a unique $\hat\sigma > 0$ (provided $V_\bullet > 0$ and $U_\bullet > 0$; otherwise the boundary $\hat\sigma = 0$ or $\hat\sigma \to \infty$ is the maximiser and the regime is degenerate). The joint stationary point $(\hat\sigma, \hat\pi)$ is thus globally unique on the simplex.

**Numerical solution: 1-D Newton.** $g$ is monotone with a closed-form derivative, so a few Newton steps on $\log\sigma$ (or bisection in $\sigma$) starting from the warm $\sigma_{\text{old}}$ converges to machine precision in $\le 10$ iterations. Once $\hat\sigma$ is found, $\hat\pi$ follows from (B.34).

**Equivalent: coordinate ascent.** Both individual updates (B.31) and (B.32) are exact closed-form maximisations of $\ell_2$ in their own coordinate (for fixed value of the other), so alternating

$$\begin{aligned} &\sigma \leftarrow U_\bullet \,/\, (\mathbf{r}^\top \pi), \\ &\pi_b \leftarrow V'_b \,/\, (V_\bullet + \sigma\, r_b) \quad \text{(automatically normalised once } \lambda = V_\bullet \text{ holds)} \end{aligned}$$

gives a monotone $\ell_2$ ascent and converges to the unique fixed point identified above. In code, $\sim 5$ alternation rounds typically suffice; the 1-D Newton on (B.35) is slightly faster and gives the same answer.

**Mixture form (per-class).** Applied class by class to the reversible mixture of Section B.1.8, each class $c$ has its own pair $(\sigma_c, \pi^{(c)})$ updated independently from its own per-class sufficient statistics $(W^{(c)}, U^{(c)}, V^{(c)})$ and its own fixed $Q^{(c)}$ shape:

$$\hat\sigma_c \text{ solves } \sum_b \frac{\hat\sigma_c\, r_b^{(c)}\, V_b'^{(c)}}{V_\bullet^{(c)} + \hat\sigma_c\, r_b^{(c)}} = U_\bullet^{(c)}, \qquad \hat\pi_b^{(c)} = \frac{V_b'^{(c)}}{V_\bullet^{(c)} + \hat\sigma_c\, r_b^{(c)}}, \tag{B.36}$$

with $r_b^{(c)} = \sum_{a\ne b} Q_{ab}^{(c)} W_a^{(c)}$. Classes do not couple, by the same class-decomposition (B.19) that justifies per-class M-steps in the standard mixture regime.

**Aggregation of counts.** The required sufficient statistics per class are $(U_\bullet^{(c)}, V_\bullet^{(c)}, V'^{(c)}, r^{(c)})$. All are linear contractions of the underlying per-pair $(W, U, V)$ statistics against the fixed quantities $(Q^{(c)}, V_\bullet^{(c)} = \text{count of } V \text{ entries})$, so under SVI–BW they accumulate via the same EMA used for the standard M-step (no special handling).

**Reduction to pure rescaling.** Setting $\pi = \pi_{\text{old}}$ in (B.31) recovers the closed form $\hat\sigma = U_\bullet / D$ of (B.26). The joint mode strictly dominates pure rescaling whenever $\hat\pi \ne \pi_{\text{old}}$ at the joint optimum (i.e. whenever the Maraschino-fit warm $\pi$ is not exactly the M-step optimum given the current $W, U, V$); in practice the gain is largest immediately after a warm start that fixed $\pi$ away from its joint optimum.

**When to use.** Use the joint $(\sigma, \pi)$ rescaling when warm-starting from a fit that already learnt per-class $(\sigma_c, \pi^{(c)})$ but the SVI–BW M-step should keep updating both with new evidence. Use the pure $\sigma$-only rescaling when the warm $\pi^{(c)}$ should be treated as a strong prior or when one wishes to attribute any substitution-rate drift purely to the time scale.

##### B.1.11 Tied Equilibria across Class Blocks

A second restricted regime ties the equilibrium distribution $\pi^{(c)}$ across blocks of site classes while leaving exchangeabilities $Q^{(c)}$ free. Partition the $C$ classes $\{1, \ldots, C\}$ into $B = C/N_{\text{tied}}$ disjoint blocks $\mathcal{B}_1, \ldots, \mathcal{B}_B$ each of size $N_{\text{tied}}$ (we require $N_{\text{tied}} \mid C$). Within each block $b$ all classes share a common equilibrium distribution

$$\pi^{(c)} \equiv \tilde{\pi}^{(b)} \quad \text{for all } c \in \mathcal{B}_b. \tag{B.37}$$

**Block-pooled complete-data log-likelihood.** Substituting (B.37) into (B.19), the log-likelihood decomposes across blocks rather than across individual classes:

$$\ell_2 = \sum_{b=1}^{B} \ell_2^{(b)}, \tag{B.38}$$

$$\begin{aligned} \ell_2^{(b)} = \sum_i \tilde{V}_i'^{(b)} \log \tilde{\pi}_i^{(b)} \; + \; \sum_{c \in \mathcal{B}_b} \sum_{j>i} \big(U_{ij}^{(c)} + U_{ji}^{(c)}\big) \log Q_{ij}^{(c)} \\ - \sum_{c \in \mathcal{B}_b} \sum_{j>i} Q_{ij}^{(c)} \left(W_i^{(c)} \tilde{\pi}_j^{(b)} + W_j^{(c)} \tilde{\pi}_i^{(b)}\right), \end{aligned} \tag{B.39}$$

with the pooled equilibrium-character count

$$\tilde{V}_i'^{(b)} = \sum_{c \in \mathcal{B}_b} V_i'^{(c)}. \tag{B.40}$$

The exchangeability parts $Q^{(c)}$ remain class-specific; only the equilibrium term pools.

**Block M-step.** Within block $b$, alternate between the per-class $Q^{(c)}$ update (holding $\tilde{\pi}^{(b)}$ fixed) and the pooled $\tilde{\pi}^{(b)}$ update (holding all $Q^{(c)}, c \in \mathcal{B}_b$ fixed). The $Q$-step uses the standard mixture formula (B.21) with the shared $\tilde{\pi}^{(b)}$:

$$\hat{Q}_{ij}^{(c)} = \frac{U_{ij}^{(c)} + U_{ji}^{(c)}}{W_i^{(c)} \tilde{\pi}_j^{(b)} + W_j^{(c)} \tilde{\pi}_i^{(b)}} \quad (c \in \mathcal{B}_b). \tag{B.41}$$

The $\tilde{\pi}^{(b)}$-step is the same Lagrange-multiplier solve as the unconstrained mixture M-step, but with pooled coefficients. Differentiating (B.39):

$$\tilde{\pi}_i^{(b)} = \frac{\tilde{V}_i'^{(b)}}{\tilde{c}_i^{(b)} - \eta}, \qquad \tilde{c}_i^{(b)} = \sum_{c \in \mathcal{B}_b} \sum_{j \neq i} Q_{ij}^{(c)} W_j^{(c)}, \tag{B.42}$$

with $\eta$ found by 1D bisection from $\sum_i \tilde{V}_i'^{(b)} / (\tilde{c}_i^{(b)} - \eta) = 1$, exactly as in Section A.1.8. Each coordinate update is an exact maximiser within its coordinate; the joint objective on $(Q^{(\cdot)}, \tilde{\pi}^{(b)})$ is strictly concave on $\mathbb{R}_+^{|\mathcal{B}_b| \cdot \binom{|\mathcal{A}|}{2}} \times \Delta^{|\mathcal{A}|-1}$ (the latter the simplex), so the coordinate ascent converges to the unique block MLE.

**Connection to standard models.** With $N_{\mathrm{tied}} = 1$ (one class per block) the tying is trivial and (B.42) reduces to the per-class Lagrange solve of Section B.1.8. With $N_{\mathrm{tied}} = C$ (one block containing all classes) all classes share a single $\tilde{\pi}$ — the discrete-rate-classes parameterisation of (54) when the $Q^{(c)}$ are also proportional to a shared shape. For intermediate $N_{\mathrm{tied}}$ the tying captures shared equilibrium chemistry across classes that nonetheless differ in their exchangeability (and hence in their substitution *pattern* and *rate*), without forcing every class to maintain its own equilibrium estimate.

**Pseudocounts.** The LG-informed Dirichlet pseudocount on $\tilde{\pi}^{(b)}$ adds $N_\pi \, \pi_i^{\mathrm{LG}}$ to $\tilde{V}_i'^{(b)}$ once per block (not once per class within the block); the per-class Gamma$(a_S, b_S)$ pseudocount on $Q_{ij}^{(c)}$ enters the class-specific update (B.41) unchanged.

**Combination with rate rescaling.** Tied-$\pi$ and rate rescaling can also be *combined* within a single M-step: each class has its own scalar $\sigma_c$ but classes within a block share the equilibrium $\tilde{\pi}^{(b)}$. The coordinate-ascent factorisation does dispatch in that joint regime; see Section B.1.12. The frozen-$\pi$ mode (which fixes $\pi^{(c)}$ at its initialisation, with no M-step update) remains an alternative restriction. Alternation *across* M-steps (rate rescaling on odd iterations, tied-$\pi$ on even iterations) is valid in addition because each step is an unconditional ascent on the joint $\ell_2$.

##### B.1.12 Tied Equilibria with Per-Class Rate Rescaling

The two restrictions of Sections B.1.9 and B.1.11 can be applied jointly to the same mixture M-step: each class $c$ in block $\mathcal{B}_b$ has its own *scalar* rate multiplier $\sigma_c$, with the exchangeability shape $Q^{(c)}$ frozen, while all classes within the block share a common equilibrium $\tilde{\pi}^{(b)}$. This matches the substitution degrees of freedom of a Maraschino-style fit with `--freeze-class-S-shape` under a block-tied `class_pi` constraint and is strictly more constrained than the joint $(\sigma_c, \pi^{(c)})$ M-step of Section B.1.10.

**Block-pooled complete-data log-likelihood.** With $R_{ij}^{(c)} = \sigma_c \, Q_{ij}^{(c)} \, \tilde{\pi}_j^{(b)}$ for $c \in \mathcal{B}_b$ (with $Q^{(c)}$ frozen), the per-block log-likelihood is

$$\ell_2^{(b)}(\{\sigma_c\}_{c\in\mathcal{B}_b}, \tilde{\pi}^{(b)}) = \sum_{b}^{\prime} \tilde{V}_{b'}'^{(b)} \log \tilde{\pi}_{b'}^{(b)} + \sum_{c\in\mathcal{B}_b} \Big[ U_\bullet^{(c)} \log \sigma_c - \sigma_c \, \mathbf{r}^{(c)\top} \tilde{\pi}^{(b)} \Big] + \text{const}, \tag{B.43}$$

where the constant collects $\sum_c \sum_{j>i} (U_{ij}^{(c)} + U_{ji}^{(c)}) \log Q_{ij}^{(c)}$ (frozen), $\tilde{V}_{b'}'^{(b)} = \sum_{c\in\mathcal{B}_b} V_{b'}'^{(c)}$ is the block-pooled boundary count, and $r_{b'}^{(c)} = \sum_{a\neq b'} Q_{a,b'}^{(c)} W_a^{(c)}$ depends only on the frozen $(Q^{(c)}, W^{(c)})$.

**Stationary conditions.** Lagrangian for $\sum_{b'} \tilde{\pi}_{b'}^{(b)} = 1$ with multiplier $\lambda^{(b)}$:

$$\frac{\partial}{\partial \sigma_c} = \frac{U_\bullet^{(c)}}{\sigma_c} - \mathbf{r}^{(c)\top} \tilde{\pi}^{(b)} = 0 \implies \sigma_c = \frac{U_\bullet^{(c)}}{\mathbf{r}^{(c)\top} \tilde{\pi}^{(b)}}, \tag{B.44}$$

$$\frac{\partial}{\partial \tilde{\pi}_{b'}^{(b)}} = \frac{\tilde{V}_{b'}'^{(b)}}{\tilde{\pi}_{b'}^{(b)}} - \sum_{c\in\mathcal{B}_b} \sigma_c \, r_{b'}^{(c)} - \lambda^{(b)} = 0 \implies \tilde{\pi}_{b'}^{(b)} = \frac{\tilde{V}_{b'}'^{(b)}}{\lambda^{(b)} + \sum_{c\in\mathcal{B}_b} \sigma_c r_{b'}^{(c)}}. \tag{B.45}$$

**Block Lagrange multiplier is fixed.** Multiplying (B.45) by $\tilde{\pi}^{(b)}_{b'}$ and summing over $b'$,

$$\sum_{b'} \tilde{V}'^{(b)}_{b'} = \lambda^{(b)} + \sum_{c \in \mathcal{B}_b} \sigma_c \, \mathbf{r}^{(c)\top} \tilde{\pi}^{(b)}.$$

Substituting (B.44) into the right-hand side collapses the $\sigma_c$-dependent term to $\sum_c U^{(c)}_\bullet$, giving

$$\boxed{\lambda^{(b)} \;=\; \tilde{V}^{(b)}_\bullet,} \qquad \tilde{V}^{(b)}_\bullet := \sum_{c \in \mathcal{B}_b} V^{(c)}_\bullet = \sum_{c \in \mathcal{B}_b} \sum_i V^{(c)}_i, \tag{B.46}$$

generalising the single-class result (B.33). The multiplier is fully determined by the block-pooled boundary count.

**Closed-form coupled updates.** Substituting (B.46) into (B.45),

$$\hat{\tilde{\pi}}^{(b)}_{b'} = \frac{\tilde{V}'^{(b)}_{b'}}{\tilde{V}^{(b)}_\bullet + \sum_{c \in \mathcal{B}_b} \sigma_c \, r^{(c)}_{b'}}. \tag{B.47}$$

Equation (B.47) together with (B.44) forms a closed nonlinear system in the $|\mathcal{B}_b| + A$ unknowns $(\{\sigma_c\}, \tilde{\pi}^{(b)})$ per block. Two practical solvers, both monotone in $\ell_2^{(b)}$:

1. *Coordinate ascent.* Alternate $\sigma_c \leftarrow U^{(c)}_\bullet / (\mathbf{r}^{(c)\top} \tilde{\pi}^{(b)})$ for each $c \in \mathcal{B}_b$ and $\tilde{\pi}^{(b)} \leftarrow \tilde{V}'^{(b)} \,/\, (\tilde{V}^{(b)}_\bullet + \sum_c \sigma_c r^{(c)})$ (componentwise), with a final renormalisation safety guard. Each update is the exact closed-form maximiser in its own coordinate; convergence to the unique block-optimum follows in $\sim$ 5–20 rounds.

2. *Substitute and root-find.* Eliminate $\sigma_c$ from (B.47) via (B.44); the result is a vector fixed-point equation in $\tilde{\pi}^{(b)} \in \Delta^{A-1}$ alone. When $|\mathcal{B}_b|$ is small (typical: 2–5), the resulting low-dimensional system is amenable to a damped Newton iteration on the $A-1$ free $\tilde{\pi}$ components; in practice coordinate ascent is faster and simpler.

Both solvers reach the same fixed point because $\ell_2^{(b)}$ is strictly concave in each block of variables given the others (the log terms ensure positivity through the FOCs).

**Mixture-wide M-step.** The blocks decouple in the same way the classes do under Section B.1.11: the joint $\ell_2 = \sum_b \ell_2^{(b)}$ separates over $b$, so each block can be solved independently.

**Aggregation of counts.** Per block, the required sufficient statistics are the per-class $(U^{(c)}_\bullet, V'^{(c)}, r^{(c)})$ for $c \in \mathcal{B}_b$ plus the block totals $\tilde{V}'^{(b)}, \tilde{V}^{(b)}_\bullet$. All are linear in $(W, U, V)$ and accumulate via the same EMA used by the standard M-step.

**Reduction to special cases.** Setting $|\mathcal{B}_b| = 1$ (each class its own block) recovers the joint $(\sigma_c, \pi^{(c)})$ M-step of Section B.1.10; setting all $\sigma_c = 1$ (no rate rescaling within block) recovers a degenerate version of the tied-$\pi$ M-step with frozen exchangeabilities; freezing $\tilde{\pi}^{(b)}$ at its current value recovers the rate-only rescaling of Section B.1.9. Strict dominance holds in both directions: this regime improves on Section B.1.9 by adding the $\pi$ degree of freedom and improves on Section B.1.11 (with $Q^{(c)}$ frozen at LG) by adding per-class $\sigma_c$.

**When to use.** Use this regime when the warm-start fit (e.g. Maraschino with `--freeze-class-S-shape`) ties classes into blocks with shared equilibrium chemistry but distinct rates, and the SVI–BW M-step should preserve that tying while continuing to update both $(\sigma_c, \tilde{\pi}^{(b)})$ from new evidence. The block tying is a strong inductive bias when the number of classes per block is small relative to $A$ and the per-class equilibrium counts $V'^{(c)}$ would otherwise be too sparse to estimate $\pi^{(c)}$ reliably.

##### B.1.13 Stochastic Variational Baum-Welch

For large datasets it is impractical to run the E-step over all training pairs every iteration. A natural stochastic extension is to sample a minibatch of pairs, run the E-step on those, and update parameters. Naively replacing the full sufficient statistics with each batch's statistics discards information from previous batches and results in noisy, slowly converging parameter estimates.

The framework of stochastic variational inference (21) provides a principled resolution in the conditionally conjugate case. Each M-step parameter group in TKF91 (and TKF92 and MixDom) has an exponential-family complete-data log-likelihood, regular for the multinomial groups (mixture weights, fragment-type transitions, site-class distributions) and curved for the BDI rates and the reversible-CTMC (GTR) submodel (Sections A.1.3, A.1.2). For the multinomial groups we use Dirichlet priors that are strictly conjugate; for the BDI and GTR groups we use independent Gamma $\times$ Dirichlet pseudocounts that are non-conjugate regularisers in the reversible / curved-family parameterisation (Section C.1.4). For the conjugate groups, Hoffman et al.'s natural-gradient argument applies in full and the update (B.48) below is exact natural-gradient ascent on the variational ELBO. For the non-conjugate groups, the natural-gradient interpretation no longer holds verbatim, but the same exponential-moving-average update remains a sound stochastic-MAP-EM scheme: each M-step is affine in the augmented sufficient statistics, so an EMA of statistics commutes with the M-step and converges to the MAP fixed point of full-batch EM under the standard Robbins–Monro step-size conditions. In practice both groups are updated by the same rule:

$$\bar{T}^{(t+1)} = (1-\rho_t)\,\bar{T}^{(t)} + \rho_t\left(\bar{T}_0 + \frac{N}{|B_t|}\,T_{B_t}\right), \tag{B.48}$$

where $\bar{T}^{(t)}$ denotes the running sufficient statistics at iteration $t$ (transition counts $\hat{n}_{ij}$, Holmes–Rubin dwell times $W_i$ and jump counts $U_{ij}$, character counts $V_i$, and BDI expectations $\hat{B}, \hat{D}, \hat{S}$), $\bar{T}_0$ the prior pseudocounts, $T_{B_t}$ the batch sufficient statistics from the E-step on minibatch $B_t$, $N$ the total number of training pairs, $|B_t|$ the batch size, and $\rho_t = (t+\tau)^{-\kappa}$ the Robbins–Monro step size with delay $\tau \geq 0$ and forgetting rate $\kappa \in (0.5, 1]$.

The M-step—quadratic solve for indel rates (A.22), Dirichlet MAP for mixture weights, Beta MAP for extension rates, and the iterative coordinate ascent for $(Q, \pi)$—is then applied to $\bar{T}^{(t+1)}$ exactly as in full-batch EM. Each M-step maximizes a (penalised) Q-function that is linear in the sufficient statistics, so averaging the statistics before solving is equivalent to finding the MAP of the averaged Q-function. For the conjugate parameter groups this maximizer is also the natural parameter of the updated variational posterior; for the non-conjugate groups it is the MAP of the regularised Q-function and not a variational posterior, but the iteration is identical.

This retains the closed-form exactness of every M-step while enabling online processing of arbitrarily large pair collections, with each training pair visited only once. The two hyperparameters $\tau$ and $\kappa$ have robust defaults ($\tau = 10$, $\kappa = 0.7$) that work across a range of dataset sizes without tuning (21).

**Equivalent formulations of the prior pseudocounts.** Equation (B.48) places the prior pseudocounts $\bar{T}_0$ inside the EMA update. An equivalent implementation accumulates only the data contribution in $\bar{T}$ and adds $\bar{T}_0$ at M-step time:

$$\tilde{T}^{(t+1)} = (1-\rho_t)\,\tilde{T}^{(t)} + \rho_t\,\tfrac{N}{|B_t|}\,T_{B_t}, \qquad \theta^{(t+1)} = \text{M-step}\big(\tilde{T}^{(t+1)} + \bar{T}_0\big).$$

For every M-step considered in this paper (Dirichlet MAP, Gamma-augmented counts, pooled GTR formula), the M-step operator is affine in its sufficient statistics, so $\mathrm{M}(\bar{T}^{(t)}) = \mathrm{M}(\tilde{T}^{(t)} + \bar{T}_0)$ exactly at every iteration; the two prescriptions yield numerically identical estimates. This affineness is what we actually need for the EMA-of-pseudocounts formulation, and it holds whether or not the prior is strictly conjugate to the complete-data likelihood (Section C.1.4 discusses the distinction). The post-hoc form is slightly more convenient for implementation because each M-step keeps the prior encapsulated in its closed-form update; see the code comment at the SVI loop for details.

#### B.2 Stochastic Variational Baum–Welch Convergence

##### B.2.1 Setup

We consider the idealized infinite-data setting: each minibatch of $B$ sequence pairs is drawn i.i.d. from the true generative model, so there is no model misspecification. Stochastic Variational Baum–Welch (SVB) alternates:

1. **E-step**: Run forward–backward on a minibatch of $B$ pairs, accumulate expected sufficient statistics $\hat{T}_B$.
2. **M-step**: Update parameters $\theta_{k+1} = M(\hat{T}_B)$ using the closed-form M-step.

At convergence, the true parameters $\theta^*$ are a fixed point of $\theta \mapsto M\big(\mathbb{E}_\theta[T]\big)$.

##### B.2.2 BDI sufficient statistics

For a single sequence pair $(x, y)$ with ancestor length $i$, descendant length $j$, and evolutionary time $t$, the BDI sufficient statistics are $T = (\mathbb{E}[B \mid x, y],\ \mathbb{E}[D \mid x, y],\ \mathbb{E}[S \mid x, y])$, recovered via the score function identity. The M-step is (Section A.1.8):

$$\hat{\lambda} = \frac{\bar{B}}{\bar{S} + t}, \qquad \hat{\mu} = \frac{\bar{D}}{\bar{S}}, \tag{B.49}$$

where $\bar{B}, \bar{D}, \bar{S}$ denote averages (or sums) over the minibatch.

##### B.2.3 Variance of Minibatch Estimates

**Per-pair variance** Let $\sigma_B^2, \sigma_D^2, \sigma_S^2$ denote the variances of $\mathbb{E}[B \mid x, y]$, $\mathbb{E}[D \mid x, y]$, $\mathbb{E}[S \mid x, y]$ under the data-generating distribution (randomness over $(x, y)$ pairs drawn from the model). For a minibatch of $B$ independent pairs, the sample means satisfy

$$\mathrm{Var}[\bar{B}] = \frac{\sigma_B^2}{B}, \qquad \mathrm{Var}[\bar{D}] = \frac{\sigma_D^2}{B}, \qquad \mathrm{Var}[\bar{S}] = \frac{\sigma_S^2}{B}.$$

**Parameter variance via the delta method** By the delta method applied to (B.49), the relative error of $\hat{\lambda}$ after one M-step on a minibatch of size $B$ is

$$\mathrm{Var}[\hat{\lambda}]/\lambda^2 \;\approx\; \frac{1}{B}\left[\frac{\sigma_B^2}{\mathbb{E}[B]^2} + \frac{\sigma_S^2}{(\mathbb{E}[S]+t)^2} - \frac{2\,\mathrm{Cov}[B,S]}{\mathbb{E}[B](\mathbb{E}[S]+t)}\right] \equiv \frac{v_\lambda}{B}, \tag{B.50}$$

and similarly for $\hat{\mu}$:

$$\mathrm{Var}[\hat{\mu}]/\mu^2 \;\approx\; \frac{1}{B}\left[\frac{\sigma_D^2}{\mathbb{E}[D]^2} + \frac{\sigma_S^2}{\mathbb{E}[S]^2} - \frac{2\,\mathrm{Cov}[D,S]}{\mathbb{E}[D]\,\mathbb{E}[S]}\right] \equiv \frac{v_\mu}{B}. \tag{B.51}$$

The constants $v_\lambda, v_\mu$ depend on $(\lambda, \mu, t)$ and the sequence length distribution. They are the diagonal entries of the inverse Fisher information matrix (per observation), expressed in the $(\lambda, \mu)$ parameterization.

**Scaling of per-pair variance with sequence length** The dominant source of randomness is the ancestor/descendant length pair $(i, j)$. At stationarity with $\kappa = \lambda/\mu < 1$:

- $\mathbb{E}[i] = \kappa/(1-\kappa), \quad \mathrm{Var}[i] = \kappa/(1-\kappa)^2$.
- $\mathbb{E}[B \mid i] \approx \lambda\,\mathbb{E}[S \mid i], \quad \mathbb{E}[S \mid i] \approx i/(\mu-\lambda)$ for $t \gg 1/(\mu-\lambda)$.
- The relative variance $v_\lambda$ is $O(1)$ in the number of links (it does *not* decrease as sequences get longer, because the M-step aggregates over the pair, not per-link).

Thus the per-pair Fisher information is $O(1)$ in sequence length — longer sequences do not help much for indel parameter estimation (in contrast to substitution parameters, where Fisher information grows linearly with alignment length).

**Proposition B.1** (Per-pair Fisher information, long-time limit)**.** *For $\delta = \mu - \lambda > 0$ and $t \to \infty$:*

$$\mathbb{E}[B] \to \frac{\kappa}{1-\kappa}, \qquad \mathbb{E}[D] \to \frac{\kappa}{1-\kappa}, \qquad \mathbb{E}[S] \to \frac{2\kappa}{(1-\kappa)\delta},$$

*and the coefficient of variation of $\hat{\lambda}$ from a single pair is $O(1)$ (bounded away from zero).*

This reflects the fact that, at stationarity, each pair provides a single "observation" of the BDI process endpoint distribution $P(j \mid i)$ — the information content is fundamentally per-pair, not per-residue.

##### B.2.4 Pseudocount representation and its advantages

The SVB iterates can be represented equivalently in two ways: as an EMA of data-only sufficient statistics $s_k$ with priors $\alpha$ added inside each M-step, or as an EMA directly in *posterior pseudocount* space $\tilde{\alpha}_k = s_k + \alpha$. In the latter view, the update rule is

$$\tilde{\alpha}_k \;=\; (1-\eta_k)\,\tilde{\alpha}_{k-1} + \eta_k\big(\alpha + (N/|B|)\,s_{\mathrm{batch}_k}\big), \tag{B.52}$$

matching (B.48) with the prior $T_0 \equiv \alpha$ absorbed into the EMA target. By induction, $\tilde{\alpha}_k = s_k + \alpha$ at every step given a matched initial state, so the two formulations produce identical parameter iterates.

For the Dirichlet-multinomial groups, $\tilde{\alpha}$ is the natural parameter of the conjugate posterior; for the BDI / GTR groups it is the augmented sufficient-statistic vector at which the regularised Q-function is maximised (the underlying priors there are non-conjugate regularisers; see Section C.1.4). In either case, the closed-form M-step maximises

$$J(\theta \mid \tilde{\alpha}) \;=\; \tilde{\alpha}\cdot\log\theta - Z(\theta), \tag{B.53}$$

where $Z$ is the log-partition function of the family. For a Dirichlet-multinomial component (transitions, mixture weights, class pis, classdist), $J = \sum_i (\tilde{\alpha}_i - 1) \log \theta_i$ up to a constant, with unique maximizer $\theta_i \propto \tilde{\alpha}_i - 1$. The BDI rates $(\lambda, \mu)$ follow the same pattern but the joint $J$ couples them through the geometric fragment length prior and the M-step is obtained from the quadratic in $\kappa = \lambda/\mu$ of (A.22). The SVB iteration thus reduces to updating one pytree of pseudocounts per step, with M-step parameters computed on demand.

**Variance via a single Jacobian.** Let $f\colon \tilde{\alpha} \mapsto \theta$ denote the M-step map. The delta method gives

$$\mathrm{Var}[\theta_k] \;\approx\; J_f(\tilde{\alpha}_k)\,\mathrm{Var}[\tilde{\alpha}_k]\,J_f(\tilde{\alpha}_k)^\top. \tag{B.54}$$

Equations (B.50)–(B.51) are the diagonal entries of (B.54) in the $(\lambda, \mu)$ parameterization; in pseudocount form, the same delta method is one derivative of $f$ away regardless of parameter group.

**Polyak–Ruppert averaging is built in.** Expanding (B.52),

$$\tilde{\alpha}_K \;=\; \sum_{j=1}^{K} w_{j,K}\big(\alpha + (N/|B|)\, s_{\mathrm{batch}_j}\big), \qquad w_{j,K} \;=\; \eta_j \prod_{i>j}^{K} (1 - \eta_i).$$

The pseudocount state is itself a weighted average of batch-level pseudocounts, and the M-step applied to this average is the Rao–Blackwell improvement over averaging parameter iterates — which matters when $f$ is nonlinear, as for the BDI rate M-step (§C.1.4, eq. (A.22)). An effective sample size

$$\mathrm{ESS}_K \;=\; \frac{\big(\sum_j w_{j,K}\big)^2}{\sum_j w_{j,K}^2}$$

quantifies how "warmed up" the EMA is and is strictly less than $K$ whenever $\eta_k < 1$. For a parameter group receiving a non-negligible contribution in only a fraction $\varepsilon$ of batches (a rare domain, a rarely-visited fragment state, a rarely-active class), the per-group effective sample size is $\varepsilon\,\mathrm{ESS}_K$, not $\mathrm{ESS}_K$ — exposing the rare-parameter bottleneck of §B.3 as a direct quantity rather than an approximation.

**Complete-data Fisher information.** For a Dirichlet-multinomial component, the Hessian of $J$ at its maximum is diagonal in natural-parameter coordinates with $[I_{\mathrm{comp}}(\tilde{\alpha})]_{ii} = 1/\tilde{\alpha}_i$. Rare categories (small $\tilde{\alpha}_i$) therefore have large per-component variance, and the bottleneck for each parameter group can be read off directly as $\tilde{\alpha}_i/\mathrm{ESS}$ — a diagnostic in natural-parameter space that replaces the CV-based convergence recommendation of item 6 in §B.2.

**Practical consequence.** An implementation that carries $\tilde{\alpha}_k$ as its primary state — rather than separately tracking $s_k$ and re-adding $\alpha$ in each M-step — makes (B.54), the Polyak–Ruppert weights, and the per-group Fisher diagnostics all computable from the same pytree, with no additional bookkeeping.

##### B.2.5 SVB Convergence Rate

**Stochastic approximation framework** SVB is a stochastic fixed-point iteration. By the results of Cappé & Moulines (2009), under regularity conditions satisfied by exponential families, the parameter iterates $\theta_k$ satisfy

$$\sqrt{K}(\bar{\theta}_K - \theta^*) \xrightarrow{d} \mathcal{N}(0, \Sigma) \tag{B.55}$$

where $\bar{\theta}_K = \frac{1}{K}\sum_{k=1}^{K} \theta_k$ is the Polyak–Ruppert average and $\Sigma$ depends on the learning rate schedule.

**Direct M-step (no averaging).** With direct M-step updates (step size 1), the iterate $\theta_K$ has variance

$$\mathrm{Var}[\theta_K] \approx \frac{\Sigma_{\text{per-pair}}}{B} \tag{B.56}$$

after convergence of the transient, where $\Sigma_{\text{per-pair}}$ is the asymptotic covariance of the M-step estimator applied to a single pair. This does *not* decrease with $K$ — the iterates fluctuate around $\theta^*$ with amplitude $O(1/\sqrt{B})$.

**Polyak–Ruppert averaging.** Averaging the iterates:

$$\mathrm{Var}[\bar{\theta}_K] \approx \frac{\Sigma_{\text{per-pair}}}{BK}. \tag{B.57}$$

This is optimal: the total number of pairs processed is $N = BK$, and we achieve variance $\Sigma/N$ regardless of how $N$ is split between $B$ and $K$.

**Exponential moving average (EMA).** In practice, one often uses $\theta_{k+1} = (1-\eta)\theta_k + \eta\, M(\hat{T}_B)$ with step size $\eta \in (0, 1]$. The asymptotic variance is

$$\mathrm{Var}[\theta_\infty] \approx \frac{\eta}{2-\eta} \cdot \frac{\Sigma_{\text{per-pair}}}{B} \approx \frac{\eta\, \Sigma_{\text{per-pair}}}{2B} \quad (\eta \ll 1).$$

This trades convergence speed for lower asymptotic variance.

**Convergence transient** The EM rate matrix governs the transient. For exponential families, the EM convergence rate is $\rho = 1 - I_{\text{obs}}/I_{\text{comp}}$, where $I_{\text{obs}}$ and $I_{\text{comp}}$ are the observed and complete-data Fisher information matrices. For BDI, the fraction of missing information is typically moderate ($\rho \approx$ 0.3–0.7 for typical protein evolution times $t \sim$ 0.5–2), so the transient dies out in $O(10)$ iterations.

##### B.2.6 Practical Recommendations

**Minibatch size** For target relative error $\varepsilon$ on indel parameters after $K$ steps with Polyak–Ruppert averaging:

$$BK \geq \frac{v_\theta}{\varepsilon^2}, \tag{B.58}$$

where $v_\theta \sim O(1)$ is the per-pair relative variance from (B.50)–(B.51).

| Target $\varepsilon$ | $BK$ needed | $B = 50, K =?$ | $B = 200, K =?$ |
|---|---|---|---|
| 0.10 | $\sim 100\, v_\theta$ | 2 | 1 |
| 0.05 | $\sim 400\, v_\theta$ | 8 | 2 |
| 0.01 | $\sim 10{,}000\, v_\theta$ | 200 | 50 |

**Rule of thumb.** Take $v_\theta \approx 5$ as a conservative estimate for typical protein BDI parameters ($\kappa \approx 0.9$, $t \approx 1$). Then:

- $B \geq 50$ ensures that each M-step estimate has $\lesssim 30\%$ relative error on indel parameters.
- $B \geq 200$ brings single-step error below $\sim 15\%$.
- For rare domains ($w_d \approx 0.05$), multiply $B$ by $1/w_d = 20$.
- After $K$ steps with averaging, total error scales as $1/\sqrt{BK}$.

**Natural gradient** The BDI complete-data statistics form a curved exponential family with natural parameters $(\log\lambda, \log\mu, -(\lambda+\mu))$ and sufficient statistics $(B, D, S)$. The natural gradient replaces the M-step update $\Delta\theta = -I_F^{-1}\nabla_\theta Q(\theta)$ where $I_F$ is the Fisher information matrix of the complete-data family.

For the BDI M-step, the closed-form solution (B.49) already implicitly incorporates the complete-data Fisher information structure. Stochastic natural gradient (SNatGrad) would precondition the *stochastic* update direction using $I_F^{-1}$, but since $\hat\lambda = \bar B/(\bar S + t)$ is already an unbiased estimator of $\lambda$ (to first order), the benefit of explicit preconditioning is marginal.

**Recommendation**: Natural gradient is not worth the implementation complexity for SVB on TKF-family models. The closed-form M-step already provides the natural parameterization. Use Polyak–Ruppert averaging of the M-step outputs instead.

**Summary of recommendations**

1. **Minibatch size**: $B \geq 50$ for general parameters; $B \geq 200$ or use averaging if rare domains ($w_d < 0.1$) are present.
2. **Averaging**: Use Polyak–Ruppert averaging of parameter iterates. Total error $\propto 1/\sqrt{BK}$, optimal regardless of $B/K$ split.
3. **Step size**: With direct M-step ($\eta = 1$), iterates oscillate with amplitude $O(1/\sqrt{B})$. EMA with $\eta = 2/(k+2)$ (Polyak schedule) achieves $O(1/\sqrt{BK})$.
4. **Decoupled updates**: Substitution parameters converge $\sim \bar L$ times faster than indel parameters. Consider accumulating indel statistics over $\sim \bar L/B$ minibatches before updating.
5. **Natural gradient**: Not recommended; the closed-form M-step is already natural.
6. **Convergence diagnostic**: Monitor the running coefficient of variation of $\hat\lambda_d, \hat\mu_d$ across minibatches. Convergence when CV $< \varepsilon$.

##### B.2.7 Bias diagnostics: Hellinger, ESS, and Fisher readouts

The convergence diagnostics above answer "have the parameter iterates stabilised?" but are silent on a distinct question that arises in practice: when validation log-likelihood *regresses* during training, is the regression within the noise floor predicted by the minibatch variance, or is it evidence of a structural problem (class-assignment drift, over-concentration of a mixture component, over-fitting of the training likelihood)? The pseudocount view of §B.2.4 makes several readouts available that are cheap to compute from saved checkpoints and sharply discriminate among these hypotheses.

**Hellinger distance on mixture distributions.** For any probability vector $p$ (e.g. a row of `classdist[d, f, :]`, a fragment-transition row $r^{(d)}_{f,\cdot}$, or a domain-weight vector), the Hellinger distance

$$H(p,q) \;=\; \tfrac{1}{\sqrt{2}}\,\|\sqrt{p}-\sqrt{q}\|_2 \;\in\; [0,1]$$

is the natural metric: bounded, symmetric (unlike KL), and *locally linear in probability-mass movement.* A transfer of $\delta$ mass from one category to another produces $H \approx \sqrt{\delta}/\sqrt{2}$, so a 1% reassignment registers as $H \approx 0.07$ and a clean class-swap gives $H = 1$. The drift hypothesis — that a mixture component is discretely reassigned between iterations $k$ and $k{+}1$ and that this reassignment accounts for the val-LL regression — predicts one or more iter-to-iter Hellinger values well above the

EM-concentration noise floor, where the noise floor is empirically $\sim 0.005$ on training runs seeded from an informed mixture profile (e.g. IQ-TREE's C10). A "sharp" drift signature appropriate to a $\sim 10\%$ per-pair val-LL regression would be $H > 0.15$ at the responsible iter; a flat trajectory at $H \leq 0.02$ throughout the regression window falsifies the discrete-drift hypothesis and redirects investigation to continuous alternatives (over-fitting of the training likelihood, slow limit cycles, or systematic mis-specification).

**Effective sample size per parameter group.** As derived in §B.2.4, the EMA effective sample size $\mathrm{ESS}_K = (\sum_j w_{j,K})^2 / \sum_j w_{j,K}^2$ is a first-class quantity in pseudocount space, strictly smaller than $K$ whenever $\eta_k < 1$. For a parameter group receiving a non-negligible batch contribution in only a fraction $\varepsilon$ of minibatches, the *group-specific* effective sample size is $\varepsilon\,\mathrm{ESS}_K$, turning the rare-parameter bottleneck of §B.3 from a theoretical concern into a direct quantity that can be logged per iter.

**Fisher readout of remaining uncertainty.** Combining the two quantities, the per-component posterior uncertainty for a Dirichlet-multinomial parameter group is proportional to $\tilde{\alpha}_i/\mathrm{ESS}$: rare categories with small posterior pseudocount and small per-group ESS are exactly those whose point estimates are least certain. This replaces the CV-of-$\hat{\theta}$ heuristic of item 6 above with a reading in natural-parameter space and lets one rank parameter groups by residual uncertainty without running a second pass of DP.

**Suggested dashboard.** At minimum, log per iter: (i) $\eta_k$ and $\mathrm{ESS}_k$; (ii) for each mixture group (`classdist`, `ext` rows, `dom_weights`), the max Hellinger distance to the previous iter and the entropy of the current iterate; (iii) $\|\Delta\tilde{\alpha}_k\|_1/\|\tilde{\alpha}_k\|_1$ per parameter group as an $\tilde{\alpha}$-space convergence diagnostic; (iv) the Hungarian-matched distance of the current class_pis (or analogous mixture-profile tensor) to the initial informed seed, as a "drift-from-prior" signature independent of relabelling. All are computable from saved parameters alone; no DP is required. Taken together, these give a rapid read-out on whether a val-LL regression reflects minibatch noise, a discrete drift event, or systematic over-fitting — and thus whether the response should be a larger minibatch, a frozen-component diagnostic, or a stronger prior.

#### B.3 Expected Statistics and Linearized Convergence

The preceding sections characterized SVB convergence in terms of abstract per-pair variances $\sigma_B^2, \sigma_D^2, \sigma_S^2$. We now derive these quantities in closed form at stationarity, giving concrete convergence estimates for the BDI parameter group of any TKF-family model. The corresponding specialisation to the MixDom hierarchy of top-level, per-domain, and intra-fragment chains is collected in Appendix C.6.

##### B.3.1 BDI expected statistics at stationarity

**Lemma B.2** (Expected BDI sufficient statistics)**.** *Consider a single BDI process with insertion rate $\lambda$, deletion rate $\mu$, $\kappa = \lambda/\mu < 1$, $\delta = \mu - \lambda > 0$, and define $\alpha = e^{-\mu T}$, $\Phi = 1 - \alpha$. For an ancestor–descendant pair $(i, j)$ with ancestor length $i$ drawn from the stationary distribution $i \sim$* $\mathrm{Geom}(\kappa)$

*(i.e. $P(i) = (1-\kappa)\kappa^i$ for $i = 0, 1, 2, \ldots$), the expected BDI sufficient statistics are:*

$$\mathbb{E}_\pi[i] = L \;=\; \frac{\kappa}{1-\kappa} \tag{B.59}$$

$$\mathbb{E}_\pi[S] = L\,T \tag{B.60}$$

$$\mathbb{E}_\pi[B] = \lambda\,(L+1)\,T \;=\; \frac{\lambda T}{1-\kappa} \tag{B.61}$$

$$\mathbb{E}_\pi[D] = \mu\,L\,T \;=\; \frac{\mu\,\kappa\,T}{1-\kappa} \;=\; \mathbb{E}_\pi[B] \tag{B.62}$$

*with $M = 1$ (one trajectory endpoint per pair) and $T = T$ (total observation time for a single process).*

*The key intermediate result is: for a BDI process starting from $N(0) = i$ mortal links, the expected number at time $s$ is*

$$\mathbb{E}[N(s) \mid N(0) = i] = i\,e^{-\delta s} + L\,(1 - e^{-\delta s}), \tag{B.63}$$

*so the time-integrated expected count is*

$$\mathbb{E}[S \mid i] = \int_0^T \mathbb{E}[N(s) \mid i]\,ds = (i - L)\,\frac{1 - e^{-\delta T}}{\delta} + L\,T. \tag{B.64}$$

*At stationarity, $\mathbb{E}_\pi[i] = L$, so $\mathbb{E}_\pi[S] = L\,T$. The birth and death expectations follow from $\mathbb{E}[B \mid i] = \lambda\,\mathbb{E}[S \mid i] + \lambda T$ (the $\lambda T$ term is the immortal link's contribution) and $\mathbb{E}[D \mid i] = \mu\,\mathbb{E}[S \mid i]$. The conservation law $\mathbb{E}_\pi[B] = \mathbb{E}_\pi[D]$ confirms that stationarity is maintained.*

*Proof.* Equation (B.63) is the standard result for a linear birth-death process with immigration at rate $\lambda$ and per-capita death rate $\mu$. The $i$ initial links each survive for $\mathrm{Exp}(\mu)$ time (giving the $i\,e^{-\delta s}$ decay), while the immigration generates new links at rate $\lambda$ from the immortal link plus the mortal links themselves; the mean converges to the stationary value $L = \kappa/(1-\kappa)$ as $s \to \infty$.

Integrating (B.63):

$$\begin{aligned}
\mathbb{E}[S \mid i] &= \int_0^T \left[i\,e^{-\delta s} + L\,(1 - e^{-\delta s})\right] ds \\
&= i\,\frac{1 - e^{-\delta T}}{\delta} + L\Big(T - \frac{1 - e^{-\delta T}}{\delta}\Big) \\
&= (i - L)\,\frac{1 - e^{-\delta T}}{\delta} + L\,T.
\end{aligned}$$

Taking $\mathbb{E}_\pi[i] = L$, the $(i - L)$ term vanishes, giving $\mathbb{E}_\pi[S] = L\,T$.

For births: $\mathbb{E}_\pi[B] = \lambda\,\mathbb{E}_\pi[S] + \lambda T = \lambda\,(L+1)\,T = \lambda T/(1-\kappa)$.

For deaths: $\mathbb{E}_\pi[D] = \mu\,\mathbb{E}_\pi[S] = \mu\,L\,T = \mu\,\kappa\,T/(1-\kappa) = \lambda T/(1-\kappa) = \mathbb{E}_\pi[B]$. □

##### B.3.2 Per-pair variance of sufficient statistics

The randomness in $(B, D, S)$ across pairs arises from the randomness in the ancestor and descendant lengths $(i, j)$. At stationarity, $i \sim \mathrm{Geom}(\kappa)$ with $\mathrm{Var}[i] = \kappa/(1-\kappa)^2$.

Since $\mathbb{E}[S \mid i]$ is linear in $i$ (eq. (B.64)), the variance of $\mathbb{E}[S \mid i]$ across pairs is:

$$\mathrm{Var}_\pi[\mathbb{E}[S \mid i]] = \left(\frac{1 - e^{-\delta T}}{\delta}\right)^2 \cdot \frac{\kappa}{(1-\kappa)^2} \;\equiv\; \sigma^2_{S,\mathrm{outer}}. \tag{B.65}$$

Similarly, since $\mathbb{E}[B \mid i] = \lambda\, \mathbb{E}[S \mid i] + \lambda\, T$:

$$\mathrm{Var}_\pi[\mathbb{E}[B \mid i]] = \lambda^2\, \sigma^2_{S,\text{outer}}, \tag{B.66}$$

and $\mathbb{E}[D \mid i] = \mu\, \mathbb{E}[S \mid i]$:

$$\mathrm{Var}_\pi[\mathbb{E}[D \mid i]] = \mu^2\, \sigma^2_{S,\text{outer}}. \tag{B.67}$$

By the law of total variance, $\mathrm{Var}_\pi[S] = \mathrm{Var}_\pi[\mathbb{E}[S \mid i]] + \mathbb{E}_\pi[\mathrm{Var}[S \mid i]]$. The inner variance $\mathrm{Var}[S \mid i]$ captures the randomness of the BDI trajectory given a fixed starting state $i$. This is harder to compute in closed form, but for the purpose of convergence estimates, the outer variance (B.65) provides a *lower bound* on $\mathrm{Var}[S]$ and is the dominant term when $\kappa$ is near 1 (long sequences), since both sources of variance scale as $O(\kappa/(1-\kappa)^2)$.

The covariance structure is also dominated by the outer term:

$$\mathrm{Cov}_\pi[\mathbb{E}[B \mid i],\, \mathbb{E}[S \mid i]] = \lambda\, \sigma^2_{S,\text{outer}}. \tag{B.68}$$

##### B.3.3 Relative error and per-pair coefficients of variation

Substituting into the delta-method formulas (B.50)–(B.51) with $\mathbb{E}[B] = \lambda\, T/(1-\kappa)$, $\mathbb{E}[S] = L\, T$, and the outer-variance approximation:

Define

$$\psi = \frac{1 - e^{-\delta T}}{\delta\, T} \;\in\; (0, 1)$$

(the ratio of the "regression to mean" time scale to the total time). Then:

$$\sigma^2_{S,\text{outer}} = \psi^2\, T^2\, \frac{\kappa}{(1-\kappa)^2} \tag{B.69}$$

$$\frac{\sigma^2_{S,\text{outer}}}{\mathbb{E}_\pi[S]^2} = \frac{\psi^2}{\kappa\,/(1-\kappa)^2 \cdot T^2} \cdot \frac{\kappa}{(1-\kappa)^2} \;=\; \frac{\psi^2\,(1-\kappa)^2}{\kappa} \;\cdot\; \frac{1}{1}$$

The relative variance of $\hat{\lambda}$ from a single pair becomes (using the outer-variance approximation in (B.50), and noting that the $B$–$S$ covariance term partially cancels the $B$ and $S$ terms):

$$v_\lambda \;\approx\; \frac{\psi^2\,(1-\kappa)^2}{\kappa} \left[1 + \frac{\lambda^2}{(\lambda + 1/T)^2} - \frac{2\lambda}{\lambda + 1/T}\right] \;=\; \frac{\psi^2\,(1-\kappa)^2}{\kappa\,(1+\lambda T)^2} \tag{B.70}$$

This uses $\mathbb{E}[S] + T = (L+1)\, T = T/(1-\kappa)$, so $\lambda/(\lambda + 1/T) = \lambda T/(1+\lambda T) = \kappa\delta T/(\delta + \delta/(\kappa \cdots))$—more directly, substitute into (B.50):

$$\begin{aligned}
v_\lambda &= \frac{\lambda^2 \sigma^2_{S,\text{outer}}}{\mathbb{E}[B]^2} + \frac{\sigma^2_{S,\text{outer}}}{(\mathbb{E}[S]+T)^2} - \frac{2\lambda \sigma^2_{S,\text{outer}}}{\mathbb{E}[B](\mathbb{E}[S]+T)} \\
&= \sigma^2_{S,\text{outer}} \left(\frac{\lambda^2(1-\kappa)^2}{\lambda^2 T^2} + \frac{(1-\kappa)^2}{T^2} - \frac{2\lambda(1-\kappa)^2}{\lambda T^2}\right) \\
&= \frac{\psi^2 \kappa}{(1-\kappa)^2} \cdot \frac{(1-\kappa)^2}{T^2} \cdot \big(1 + 1 - 2\big) \\
&= 0.
\end{aligned}$$

The outer-variance contribution to $v_\lambda$ vanishes exactly! This is because $\mathbb{E}[B \mid i] = \lambda\, (\mathbb{E}[S \mid i] + T)$, so the ratio $\mathbb{E}[B \mid i]/(\mathbb{E}[S \mid i] + T) = \lambda$ is constant in $i$—the delta-method perturbation is zero.

This means the **dominant source of variance in $\hat{\lambda}$ is the inner (trajectory) variance**, not the length distribution. The per-pair coefficient of variation of $\hat{\lambda}$ is determined by the fluctuations in $(B, D, S)$ *given* the endpoint pair $(i, j)$, which depends on the stochastic path structure of the BDI process.

**Proposition B.3** (Per-pair relative variance from inner fluctuations)**.** *For $\hat{\lambda} = \bar{B}/(\bar{S}+T)$ estimated from a single pair with ancestor length $i$ drawn from stationarity:*

$$v_\lambda \;\approx\; \frac{\mathbb{E}_\pi[\mathrm{Var}[B \mid i,j]]}{\mathbb{E}_\pi[B]^2} + \frac{\mathbb{E}_\pi[\mathrm{Var}[S \mid i,j]]}{(\mathbb{E}_\pi[S]+T)^2} - \frac{2\,\mathbb{E}_\pi[\mathrm{Cov}[B,S \mid i,j]]}{\mathbb{E}_\pi[B]\,(\mathbb{E}_\pi[S]+T)}. \tag{B.71}$$

*The inner variances* $\mathrm{Var}[B \mid i,j]$ *and* $\mathrm{Var}[S \mid i,j]$ *are the posterior variances of the BDI trajectory statistics given the observed endpoints. These are computable from the observed Fisher information (Hessian of the log-likelihood), but do not simplify to elementary closed forms.*

*However, we can bound the total per-pair relative variance using the observed Fisher information. For a single BDI process observed at one timepoint, the complete-data Fisher information for $\lambda$ is $I_{comp}(\lambda) = \mathbb{E}[S+T]/\lambda^2 = T/(\lambda^2(1-\kappa))$, and the observed (incomplete-data) information satisfies $I_{obs} \le I_{comp}$. The per-pair relative variance is $v_\lambda \ge 1/(\lambda^2\, I_{obs}) \ge 1/(\lambda^2\, I_{comp})$, giving:*

$$v_\lambda \;\ge\; \frac{1-\kappa}{\lambda T} \;=\; \frac{1}{\mathbb{E}_\pi[B]}. \tag{B.72}$$

*The per-pair relative variance is at least $1/\mathbb{E}_\pi[B]$. When $\mathbb{E}_\pi[B]$ is small (few births per pair), the relative error is large.*

**Practical approximation.** For moderate times ($\delta T \sim 1$), the missing-information fraction is $\rho \approx 0.3$–$0.7$, so $I_{\mathrm{obs}} \approx (1-\rho)\, I_{\mathrm{comp}}$ and $v_\lambda \approx (1-\kappa)/(\lambda T\,(1-\rho))$. Taking $\rho \approx 0.5$ as a rough estimate:

$$v_\lambda \;\approx\; \frac{2}{\mathbb{E}_\pi[B]} \;=\; \frac{2(1-\kappa)}{\lambda T}. \tag{B.73}$$

Similarly:

$$v_\mu \;\approx\; \frac{2}{\mathbb{E}_\pi[D]} \;=\; \frac{2(1-\kappa)}{\mu\,\kappa\, T}. \tag{B.74}$$

#### B.4 Maraschino: Cherry-Counts for TKF92

We earlier introduced Maraschino as a TKF92 generalization of CherryML in Section A.2.4. We now develop it in full.

The CherryML composite likelihood replaces the joint phylogenetic likelihood with a product of pairwise sibling (“cherry”) likelihoods (40), exploiting the fact that for a fixed alignment, the alignment-substitution counts are sufficient statistics for the substitution rate parameters. This reduces the training data—which can be many gigabytes of alignments—to a fixed-shape counts tensor whose size depends only on the alphabet $\mathcal{A}$ and the number of discretized cherry-time bins, and is independent of the number of training alignments. The resulting compact tensor fits in GPU memory and may be optimized by standard autograd-based gradient methods.

**Maraschino** extends CherryML to TKF92 by additionally including the alignment *transition* counts as sufficient statistics for the indel rates and fragment-extension probability. Concretely, for each pair of consecutive (non-empty) alignment columns we record the source column type ($\mathtt{X} \in \{\mathtt{S},\mathtt{M},\mathtt{I},\mathtt{D}\}$), the destination column type ($\mathtt{Y} \in \{\mathtt{M},\mathtt{I},\mathtt{D},\mathtt{E}\}$), and the ancestor/descendant characters in each column. We refer to the resulting tensor of counts—binned by discretized pairwise divergence time and aggregated over training cherries—as the *cherry-count tensor*, and the procedure for training TKF92 from it as Maraschino.

The Maraschino composite log-likelihood for TKF92 is the sum, over divergence-time bins $b$ and adjacency types $\texttt{X} \to \texttt{Y}$, of the cherry-count multiplied by the model log-probability of that adjacency at that bin's representative time:

$$\mathcal{L}_{\text{cherry}}^{\text{TKF92}}(\lambda, \mu, r, Q, \pi) = \sum_{b=1}^{n_\tau} \Bigg[ \sum_{\texttt{X},\texttt{Y}} n_{\texttt{X}\to\texttt{Y}}^{(b)} \log \tau'_{\texttt{X}\to\texttt{Y}}(\lambda, \mu, \bar{T}_b, r) + \sum_{a,b} n_{a,b}^{\texttt{M},(b)} \log \left[\pi_a \exp(R\bar{T}_b)_{ab}\right] + \sum_b n_b^{\texttt{I},(b)} \log \pi_b + \sum_a n_a^{\texttt{D},(b)} \log \pi_a \Bigg] \quad \text{(B.75)}$$

where $n_{\texttt{X}\to\texttt{Y}}^{(b)}$ are the binned transition counts, $n_{a,b}^{\texttt{M},(b)}, n_b^{\texttt{I},(b)}, n_a^{\texttt{D},(b)}$ are the binned per-state emission counts, $\tau'$ is the TKF92 Pair HMM transition matrix from Section A.2.2, and $\bar{T}_b$ is the representative time of bin $b$ (typically the geometric mean of the bin endpoints). The first term scores indel/fragment behaviour; the remaining three score the substitution model. Boundary contributions ($\texttt{S} \to \cdot$, $\cdot \to \texttt{E}$, $\texttt{S} \to \texttt{E}$ for empty alignments) are scored against the corresponding $\texttt{S}$-row and $\texttt{E}$-column entries of $\tau'$.

**Why this is a sufficient-statistic compression.** For fixed alignment and fixed discretized cherry times, the entire collection of training cherries enters equation (B.75) only through the binned counts. In particular, two cherry collections with identical counts yield identical $\mathcal{L}_{\text{cherry}}^{\text{TKF92}}$, identical gradients, and identical maximum-likelihood estimates. The counts tensor is therefore a sufficient statistic for the TKF92 parameters under the cherry-count composite likelihood.

**Sharded count accumulation.** For a corpus such as Pfam-Full, cherries are extracted from each MSA in parallel and contribute additively to the global counts tensor. A simple map-reduce over Pfam clans accumulates the counts in constant memory per worker; the global tensor is the elementwise sum of per-shard tensors and is independent of the order of accumulation.

##### B.4.1 Optimizing Maraschino: gradient methods vs. inner EM

The cherry-count log-likelihood $\mathcal{L}_{\text{cherry}}^{\text{TKF92}}$ is a smooth function of the unconstrained parameters $(\log \lambda, \log \mu, \log r, \log Q, \log \pi)$ (with the simplex and exchangeability constraints handled by softmax / log-Cholesky reparameterisation). Two distinct optimizers apply.

**Maraschino-as-autograd.** The objective is differentiable, so Adam, L-BFGS, and similar optimizers may be applied directly via autograd. Each step requires one matrix exponential per cherry-time bin, and a forward+backward pass through equation (B.75); both steps are highly parallel on GPU.

**Maraschino-around-EM.** Alternatively, the full-batch EM iterates of Section A.2.3 apply, with the cherry-count tensor playing the role of the E-step output. Concretely:

1. For each bin $b$, run a one-step expected-count update on the TKF92 Pair HMM at parameters $\theta^{(t)}$ and time $\bar{T}_b$, using the cherry-count tensor as the input "observed alignment." This replaces the usual Forward–Backward E-step with a closed-form linear pass that resolves the $\texttt{M}, \texttt{I}, \texttt{D}$ self-loop counts into fragment-extension counts and link-level counts (Section A.2.3).

2. Accumulate the BDI sufficient statistics $(B, D, L, M, S, T)$ via the score identity (Section A.1.2), and the Holmes–Rubin substitution statistics $(W, U, V)$ via the matched-pair contribution.
3. Apply the closed-form M-step (equations (A.22)–(A.25) for $\lambda, \mu$; $r \leftarrow F/(F + E)$ for fragment extension; Section A.1.8 item 1 for $Q, \pi$).

Because every M-step in Section A.2.3 is closed-form and linear in the sufficient statistics, the inner EM iterates are exact ascent on $\mathcal{L}^{\mathrm{TKF92}}_{\mathrm{cherry}}$, with no gradient-step or learning-rate hyperparameter. In practice, EM converges in ∼10–30 iterations on protein cherry-count tensors and is competitive with Adam on the same objective.

**Practical comparison.** Adam handles arbitrary regularizers and is straightforward to combine with stochastic minibatching over time-bins or alphabet slices. Inner EM does not require a learning rate, has monotonic convergence guarantees, and produces the natural-gradient direction "for free." When the cherry-count tensor fits in memory (it does, even for Pfam-Full at $|\mathcal{A}| = 20$ and $n_\tau = 20$), inner-EM Maraschino is the recommended default.

##### B.4.2 EM-around-Maraschino: mixtures of TKF92

A mixture of $K$ TKF92 components, each with its own indel rates, fragment-extension probability, and substitution model, captures heterogeneity within and across protein families. We fit such a mixture by an outer EM loop with Maraschino as the inner optimizer of each component:

**Algorithm 1** EM-around-Maraschino for a mixture of $K$ TKF92 components.

1: Initialize component parameters $\theta_1^{(0)}, \ldots, \theta_K^{(0)}$ and mixture weights $\pi_1^{(0)}, \ldots, \pi_K^{(0)}$.
2: **for** $t = 0, 1, \ldots$ **do**
3: **E-step.** For each cherry $c$ in the training set, compute the responsibility $\gamma_{c,k}^{(t)} \propto \pi_k^{(t)} P(c \mid \theta_k^{(t)})$ (the per-component cherry-count likelihood normalised across components). Each $P(c \mid \theta_k)$ is a single forward pass through the cherry-count likelihood at the cherry's counts subtensor.
4: **Component-weighted count accumulation.** For each component $k$, accumulate the responsibility-weighted cherry-count tensor $C_k^{(t+1)} = \sum_c \gamma_{c,k}^{(t)} C_c$.
5: **Component M-step (Maraschino).** For each component $k$, run Maraschino (Section B.4.1) on $C_k^{(t+1)}$ to obtain $\theta_k^{(t+1)}$.
6: **Mixture-weight M-step.** $\pi_k^{(t+1)} \propto \sum_c \gamma_{c,k}^{(t)}$.
7: **end for**

The outer EM is monotone in the mixture log-likelihood $\sum_c \log \sum_k \pi_k P(c \mid \theta_k)$ because both the responsibilities (E-step) and the per-component Maraschino solves (M-step) are exact ascent on their respective Q-functions.

The decisive feature is that the outer M-step does *not* require re-extracting cherries or recomputing alignments per iteration: the per-component cherry-count tensor $C_k^{(t+1)}$ is a responsibility-weighted average of the same per-cherry tensors that were extracted once and for all at the start of training.

##### B.4.3 EM-around-CherryML: mixtures over site classes

A complementary class of mixture model places the latent variable on *MSA columns* rather than on whole cherries: each MSA column $c$ draws an unobserved *site class* $z_c \sim \mathrm{Categorical}(\pi_1, \ldots, \pi_C)$ from a shared per-class substitution model $\mathrm{Subst}(Q^{(k)}, \pi^{(k)})$, and every cherry in the family that has a residue pair at column $c$ evolves its match emission under the same class. Because no indel rates vary across mixture components, this is strictly a substitution-only mixture, and the corresponding fitter is CherryML rather than Maraschino: the cherry-count tensor of Section B.4 contributes only the matched substitution observations $(a, b, t_p)$ and the indel-free composite likelihood reduces to that of (40).

**Column-shared responsibilities.** The defining observation is that within a single MSA, all cherries that have a match at column $c$ inherit the *same* latent class $z_c$. This couples cherries across the same column: the column-level posterior

$$\gamma_c(k) \;\propto\; \pi_k \prod_{p\,:\,\mathtt{M}\text{ at }c} P\big(a_{p,c}, b_{p,c} \mid k, t_p\big), \qquad P(a, b \mid k, t) = \pi_a^{(k)} \exp(R^{(k)} t)_{ab},$$

multiplies the per-cherry match-emission likelihoods over all cherries $p$ that contribute a residue pair to column $c$. Cherries that have a gap at $c$ contribute no factor for that column. The resulting composite log-likelihood

$$\mathcal{L}_{\text{site-mix}} \;=\; \sum_m \sum_{c\,\in\,\text{cols}(m)} \log \sum_{k=1}^{C} \pi_k \prod_{p\,:\,\mathtt{M}\text{ at }c} \pi^{(k)}_{a_{p,c}} \exp(R^{(k)} t_p)_{a_{p,c}\, b_{p,c}} \tag{B.76}$$

is a sum, over MSAs $m$ and over columns $c$ within each MSA, of a log-mixture in which cherries share the column's class. This is in direct contrast to EM-around-Maraschino (Section B.4.2), whose responsibilities $\gamma^{(t)}_{c,k}$ are per *cherry*: there each cherry is assigned to a single component independently of the columns it spans.

**Outer / inner EM.** We fit (B.76) by alternating an MSA-wide column-responsibility update with a per-class Holmes–Rubin M-step. At iteration $t$, the column responsibilities are

$$\gamma^{(t)}_{m,c}(k) \;\propto\; \pi^{(t)}_k \prod_{p\,:\,\mathtt{M}\text{ at }c} \pi^{(t,k)}_{a_{p,c}} \exp\!\big(R^{(t,k)}\, t_p\big)_{a_{p,c}\, b_{p,c}}, \tag{B.77}$$

normalised across $k$ for each $(m, c)$, and the responsibility-weighted Holmes–Rubin sufficient statistics for class $k$ are

$$\begin{aligned}
\hat{W}_i^{(t+1,k)} &= \sum_{m,c} \gamma^{(t)}_{m,c}(k) \sum_{p\,:\,\mathtt{M}\text{ at }c} \mathcal{C}^W_i\big(a_{p,c}, b_{p,c}, t_p\big),\\
\hat{U}_{ij}^{(t+1,k)} &= \sum_{m,c} \gamma^{(t)}_{m,c}(k) \sum_{p\,:\,\mathtt{M}\text{ at }c} \mathcal{C}^U_{ij}\big(a_{p,c}, b_{p,c}, t_p\big),\\
\hat{V}_i^{(t+1,k)} &= \sum_{m,c} \gamma^{(t)}_{m,c}(k) \sum_{p\,:\,\mathtt{M}\text{ at }c} \delta\big(a_{p,c} = i\big),
\end{aligned} \tag{B.78}$$

where $\mathcal{C}^W_i, \mathcal{C}^U_{ij}$ are the standard Holmes–Rubin endpoint-conditioned bridge expectations (Section A.1.2, eqs. (A.2)–(A.3)) at the class-$k$ rate matrix $R^{(t,k)}$. The full outer iteration is then Algorithm 2.
The outer iteration is monotone in $\mathcal{L}_{\text{site-mix}}$: the column responsibilities are the exact posterior over $z_c$ and the per-class GTR M-step is exact ascent on its (responsibility-weighted) Q-function. The inner GTR EM is itself iterative because the GTR M-step couples $(Q^{(k)}, \pi^{(k)})$ nonlinearly (Section A.1.8, item 2 of the M-step list); a few inner sweeps per outer iteration suffice in practice.

**Algorithm 2** EM-around-CherryML for $C$ site classes.

1: Initialize per-class GTR parameters $\theta_k^{(0)}$ and class weights $\pi_k^{(0)}$.
2: **for** $t = 0, 1, \ldots$ **do**
3: **Outer E-step.** Compute column responsibilities $\gamma_{m,c}^{(t)}(k)$ via (B.77).
4: **Sufficient-statistic accumulation.** Compute $(\hat{W}^{(t+1,k)}, \hat{U}^{(t+1,k)}, \hat{V}^{(t+1,k)})$ via (B.78).
5: **Per-class GTR inner EM.** For each class $k$, run the closed-form GTR coordinate-ascent M-step of Section A.1.8 on $(\hat{W}^{(t+1,k)}, \hat{U}^{(t+1,k)}, \hat{V}^{(t+1,k)})$ to obtain $\theta_k^{(t+1)}$. Because the responsibility weights are fixed, this step is a standard CherryML rate fit on a single GTR model with weighted observations.
6: **Class-weight update.** $\pi_k^{(t+1)} \propto \sum_{m,c} \gamma_{m,c}^{(t)}(k)$.
7: **end for**

**Why CherryML, not Maraschino.** No indel rates appear in (B.76): the model assumes the alignment is given and that all alignment columns share a single TKF92 indel process whose parameters are held fixed (or estimated elsewhere, e.g. by Maraschino on the same cherries). Mixing *only* the substitution model means the $(\lambda, \mu, r)$-related row-normalisation and fragment-extension responsibility issues of TKF92 do not arise; the relevant sufficient statistics are exactly those of CherryML. The only structural change relative to plain CherryML is the per-MSA, per-column responsibility weighting in (B.78), which couples cherries that share columns of the same MSA.

**Relation to per-class GTR mixtures elsewhere.** The per-class GTR M-step of equation (B.78) is exactly the reversible-mixture update of Appendix B.1.8; the only difference between this appendix and Algorithm 2 is the provenance of the responsibilities (here: cross-cherry column sharing within an MSA; in the appendix: any user-supplied per-site posterior). Site-mixture parameters fitted by EM-around-CherryML can therefore be plugged into mixture-aware downstream methods — including Maraschino with a fixed indel-process backbone — without further re-derivation.

##### B.4.4 Expected sufficient statistics as fast custom VJPs

The Maraschino objective (equation (B.75)) is concretely a sum of log-probabilities, each of which has a closed-form gradient via the score identity and the BDI/Holmes–Rubin sufficient statistics. Specifically, the $(\lambda, \mu)$-gradient of $\log \tau'_{\mathtt{X}\to\mathtt{Y}}(\lambda, \mu, t, r)$ is given exactly by the score-derivative tables of Sections A.4.2–A.4.3, and the $(Q, \pi)$-gradient of $\log[\pi_a \exp(Rt)_{ab}]$ is given by the Holmes–Rubin expected dwell times and jump counts (equations (A.2)–(A.3)).

For training pipelines built around stochastic-gradient optimizers such as Adam, this observation enables a substantial speedup. Rather than relying on autograd to back-propagate through a matrix exponential and an eigendecomposition, we register a *custom vector-Jacobian product (VJP)* that returns the score-derivative-and-bridge-expectation closed forms directly:

- Forward pass: compute and cache the matrix exponential $P(t) = e^{Rt}$ and the BDI link-level transition factors $(\alpha, \beta, \gamma)$ at each cherry-time bin.
- Backward pass: contract the upstream cotangent with the expected sufficient statistics (Holmes–Rubin and BDI score expressions) instead of with the autograd-computed Jacobian.

The forward and backward costs are both $O(|\mathcal{A}|^2)$ per bin per emission state for the substitution part and $O(1)$ for the indel part, matching the cost of evaluating the closed-form M-step itself.

This pattern generalises beyond Maraschino. Any model whose M-step is closed-form linear in expected sufficient statistics admits a corresponding fast custom VJP that exposes those expectations as the gradient of the observed log-likelihood (this is, again, the score identity (A.11)).

Concretely, this enables Adam and other gradient-based optimizers to train TKF91, TKF92, and mixture-of-TKF92 models at a per-step cost matching exact EM, while retaining Adam's flexibility to incorporate arbitrary differentiable regularizers (e.g., weight decay on the BDI rates, KL-divergence priors on the GTR exchangeability), to interleave updates with non-EM-amenable parameters (e.g., the per-component mixture weights of Section B.4.2 when fit jointly with neural-network embeddings), and to work with the mini-batch SVI-BW pseudocount stream of Section B.1.13.

#### B.5 Selected Inference Algorithms for TKF92

This section describes four inference algorithms that operate on TKF91 / TKF92 models and on phylogenetic trees: the *Fast Statistical Alignment* (FSA) framework with a Newton-step time-optimization on the per-cherry expected log-likelihood (Section C.2.1); the *Beam Search Ancestral Reconstruction* (BeamASR) algorithm (Section C.2.2); the *Variational Ancestral Reconstruction* (VarAnc) algorithm that operates on a fixed MSA with known gap structure (Section B.5.3); and its stochastic-variational training extension *svi-VarAnc* (Section B.5.4).

##### B.5.1 Fast Statistical Alignment (FSA)

Given a set of sequences and a phylogenetic tree with branch-specific TKF91 or TKF92 pair HMMs, we construct a multiple sequence alignment using the *sequence annealing* approach of (6), to which we refer the reader for a full description of the algorithm.

Briefly, the method proceeds as follows. For each pair of sequences $(x, y)$ in a selected subset (either all $\binom{N}{2}$ pairs or an $O(N \log N)$ Erdős–Rényi sample), we compute pairwise residue alignment posteriors $P(x_i \sim y_j)$ by running the Forward-Backward algorithm on the pair HMM at an optimized evolutionary time $\hat{\tau}$.

**Per-pair time optimization (NR step).** The time $\hat{\tau}$ is found by Newton–Raphson optimization of the expected log-likelihood (the "NR step"):

$$\hat{\tau} = \underset{\tau}{\operatorname{argmax}}\ \mathbb{E}_{P(\pi|x,y,\tau_0)}\big[\log P(x, y, \pi \mid \tau)\big] \tag{B.79}$$

where $\pi$ ranges over alignment paths and $\tau_0$ is an initial estimate. This expectation is computed from Forward–Backward expected counts at $\tau_0$, and—because the TKF92 expected log-likelihood is quadratic-like in $\tau$ in a neighbourhood of the maximum, with analytic derivatives via the BDI score identities of Section A.1.3—the Newton iteration typically converges in 3–5 steps. This time-maximisation differs from the original FSA approach of (6), which attempts to optimize all model parameters via unregularized EM for every pair, and consequently must terminate the EM recursion early to avoid instability. Restricting the optimization to the per-pair time $\tau$, while keeping the model parameters fixed at the population estimate, is both cheaper and more stable.

**Sequence annealing.** The pairwise posteriors are then assembled into a multiple alignment by the greedy sequence annealing procedure of (6), which iteratively merges alignment columns to maximize a sum-of-pairs posterior objective.

##### B.5.2 Beam Search Ancestral Sequence Reconstruction (BeamASR)

We now describe an alternative progressive reconstruction method that finds the maximum-likelihood ancestral sequence at each internal node by beam search, without materializing the full composite automaton.

At each internal node $v$ with children $l, r$ and observed descendant sequences $c_l, c_r$, we seek

$$\hat{a}_v = \underset{a}{\operatorname{argmax}} \Big[ \log P(a, c_l \mid B_l) + \log P(a, c_r \mid B_r) - \log P(a \mid R) \Big] \tag{B.80}$$

where $P(a, c_k \mid B_k)$ is the pair HMM forward probability on branch $k$ and $P(a \mid R)$ is the singlet probability under the root generator, subtracted to avoid double-counting the prior on $a$.

**Incremental forward profiles** Since the branches are conditionally independent given the ancestor, we can evaluate (C.9) by maintaining *incremental forward profiles*: for each branch $k$, a 1D forward table $F_k[i, q]$ giving the log-probability that descendant positions $1, \ldots, i$ have been emitted and the branch machine is in state $q$, given ancestor positions $1, \ldots, j$ processed so far.

Each ancestor character extends both profiles independently in $O(L_k)$ time per branch, where $L_k = |c_k|$.

**Beam search** The ancestor sequence $\hat{a}_v$ is built left-to-right by beam search. At each position $j$, the beam maintains $B$ candidate partial ancestors. For each candidate and each alphabet character $\sigma$:

1. Extend both branch profiles by one ancestor character $\sigma$, comprising a *match/delete phase* (the ancestor emits $\sigma$, descendant positions advance via M or D transitions) and an *insertion phase* (descendant-only insertions following the ancestor emission).
2. Update the singlet forward score for $\sigma$.
3. Score the extension: $\Delta(j, \sigma) = \Delta F_l + \Delta F_r - \Delta_{\text{singlet}}$.

The top $B$ extensions (by cumulative score) are retained. Total cost per node is $O(K \cdot B \cdot A \cdot (L_l + L_r))$ where $K = |\hat{a}_v|$ and $A$ is the alphabet size.

**Insertion phase via associative scan** The insertion recurrence within each branch profile has the form

$$x_{i+1} = \operatorname{logsumexp}\big(A_{II}\, x_i,\ b_i\big) + e_i \tag{B.81}$$

where $A_{II}$ is the I-to-I log-transition submatrix, $b_i$ collects transitions into insertion states from M and D, and $e_i$ is the emission score. This is a log-semiring affine recurrence, parallelizable via an associative scan with operator

$$(A_1, b_1) \oplus (A_2, b_2) = (A_2 \otimes A_1,\ \operatorname{logsumexp}(A_2\, b_1,\ b_2))$$

where $\otimes$ denotes log-semiring matrix multiplication. This reduces the insertion phase from $O(L)$ sequential depth to $O(\log L)$.

**Supported model types** The beam search interface is generic over the pair HMM used on each branch. For the present (TKF) paper we specialize to:

1. **TKF91** — order-0, 5-state pair HMM.
2. **TKF92** — order-0, 5-state pair HMM with fragment self-loops (the standard model of Section A.2.2).

The same beam-search framework extends without algorithmic change to hierarchical TKF extensions in which each branch carries a larger latent-state pair HMM (30); we do not pursue that direction here.

##### B.5.3 Variational Ancestral Reconstruction (VarAnc)

Given an MSA with known gap structure—i.e., for each internal node $v$ and MSA column $c$, we know whether $v$ is present (ungapped) or absent (gapped)—but unknown ancestral residue identities, we optimize a variational lower bound (ELBO) on the marginal likelihood. Our approach is a "product of trees" structured variational approximation (28).

**Model structure** The joint distribution over observed leaf sequences $\mathbf{y}$ and ancestral sequences $\mathbf{a}$ factorises over tree edges. With the alignment structure fixed, the log-likelihood decomposes into contributions from consecutive alignment-column pairs along each edge $e = (u, v)$:

$$\log P(\mathbf{y}, \mathbf{a}) = \sum_{e \in \text{edges}} \sum_{k=1}^{K_e} \log w_e\big(\text{type}_{k-1},\ \text{type}_k,\ \text{ctx}_{k-1},\ \text{ctx}_k\big) \tag{B.82}$$

where $w_e$ is the TKF92 pair-HMM transition weight on edge $e$ (depending on the previous and current column types and context characters, plus the branch length). The first and last columns contribute start (S) and end (E) transition factors.

The key computational challenge is that even the order-0 TKF92 transitions couple adjacent alignment columns through the fragment-extension self-loops: the pair-HMM weight at column $c$ depends on the ancestral and descendant characters at both column $c$ and its predecessor. This makes the full graphical model a tree (phylogeny) $\times$ chain (sequence) grid with undirected cycles, and exact inference is intractable.

**Product-of-trees variational distribution** Following (28), we factorize the variational posterior over MSA columns while retaining the full tree joint at each column:

$$q(\mathbf{a}) = \prod_{c=1}^{L} q_c\big(\mathbf{h}_c\big) \tag{B.83}$$

where $\mathbf{h}_c = \{a_{v,c} : v \in \text{ungapped}(c)\}$ denotes the set of all ancestral characters at MSA column $c$, and each $q_c$ is a tree-structured distribution over the characters at the internal nodes that are ungapped at column $c$. Leaf characters are observed (clamped).

This approximation decouples columns in $q$ but preserves the full parent-child correlation structure within each column. In particular, the pairwise joint marginal $q_c^{(u,v)}(a_u, a_v)$ on each tree edge is available exactly from the Felsenstein peeling/unpeeling pass used to represent $q_c$ (Section B.5.3).

**Free energy and ELBO** The evidence lower bound is

$$\mathcal{L}(q) = \mathbb{E}_q\big[\log P(\mathbf{y}, \mathbf{a})\big] + H(q) \ \geq\ \log P(\mathbf{y}) \tag{B.84}$$

where $H(q) = -\sum_c \sum_{\mathbf{h}_c} q_c(\mathbf{h}_c) \log q_c(\mathbf{h}_c)$ is the entropy of the product-of-trees distribution. Since each $q_c$ is tree-structured, its entropy decomposes as

$$H(q_c) = \sum_{v \in \text{ungapped}(c)} H\big(q_c^{(v)}\big) - \sum_{e=(u,v)} \text{MI}_c(u, v) \tag{B.85}$$

where $H(q_c^{(v)})$ is the marginal entropy at node $v$ and $\text{MI}_c(u, v)$ is the mutual information between parent $u$ and child $v$ at column $c$, both computable from the Felsenstein marginals.

**Potentials from neighboring columns** The inter-column coupling enters through the TKF92 pair-HMM transitions. For each edge $e = (u, v)$ and MSA column $c$ where $e$ has an event of type $\tau_c$, let $c^-$ denote the predecessor column (the previous column where $e$ had an event) and $c^+$ the successor column. The *potential* at column $c$ for edge $e$ receives two contributions:

**As-child term (from $c^-$).** The transition from column $c^-$ to $c$ on edge $e$ depends on the characters at both columns. Using the pairwise marginal from $q_{c^-}$:

$$\log \phi_e^{\text{child}}(a_{u,c}, a_{v,c}) = \sum_{a',b'} q_{c^-}^{(u,v)}(a', b') \, \log w_e(\tau_{c^-}, \tau_c, a', b', a_{u,c}, a_{v,c}) \tag{B.86}$$

where the sum over $(a', b')$ uses the *joint* pairwise marginal $q_{c^-}^{(u,v)}(a', b')$—not the product of independent marginals. This is the key advantage over mean-field: the within-tree parent-child correlation at the predecessor column is preserved exactly.

**As-parent term (from $c^+$).** Symmetrically, column $c$ acts as the predecessor for column $c^+$:

$$\log \phi_e^{\text{parent}}(a_{u,c}, a_{v,c}) = \sum_{a'',b''} q_{c^+}^{(u,v)}(a'', b'') \, \log w_e(\tau_c, \tau_{c^+}, a_{u,c}, a_{v,c}, a'', b'') \tag{B.87}$$

For insert transitions (only the descendant is present at $c$), the potential reduces to a per-node function $\psi_v(a_{v,c})$. For delete transitions (only the ancestor is present), it becomes $\psi_u(a_{u,c})$. For match transitions, it contributes a per-edge potential $\phi_e(a_{u,c}, a_{v,c})$. The start and end transitions contribute analogous per-node or per-edge terms.

**Felsenstein coordinate ascent** Each coordinate ascent step updates $q_c$ for a single MSA column $c$, holding all other columns fixed. We accumulate, for each edge $e$ and node $v$ in the tree at column $c$:

- Per-edge log-potentials $\log \phi_e(a_u, a_v) = \log \phi_e^{\text{child}} + \log \phi_e^{\text{parent}}$ (for match transitions where both endpoints are present).
- Per-node log-potentials $\log \psi_v(a_v)$ (from insert/delete transitions on incident edges, plus the root prior $\log \pi(a)$ at the root node).

The optimal $q_c$, given the potentials, is the Gibbs distribution on the tree at column $c$:

$$q_c(\mathbf{h}_c) \propto \prod_v \psi_v(a_v) \prod_{e=(u,v)} \phi_e(a_u, a_v) \prod_{\ell \in \text{leaves}} \delta(a_\ell = y_\ell) \tag{B.88}$$

Since this is a tree-structured MRF, the normalising constant and all node and edge marginals can be computed *exactly* by Felsenstein peeling (postorder) and unpeeling (preorder) in $O(|E| \cdot |\mathcal{A}|^2)$ time.

**Peeling (postorder).** For each node $v$ in postorder, compute the conditional likelihood:

$$\text{CL}_v(a) = \psi_v(a) \prod_{\text{children } c} \left[ \sum_{a_c} \phi_{(v,c)}(a, a_c) \, \text{CL}_c(a_c) \right] \tag{B.89}$$

with $\text{CL}_\ell(a) = \delta(a = y_\ell)$ for observed leaves. The log-partition function is $\log Z_c = \log \sum_a \pi(a) \, \text{CL}_{\text{root}}(a)$.

**Unpeeling (preorder).** Propagate top-down to obtain the posterior marginal at each node:

$$q_c^{(v)}(a) \propto \mathrm{CL}_v(a) \cdot \mathrm{msg}_{\mathrm{parent}\to v}(a) \tag{B.90}$$

and the pairwise marginal on each edge:

$$q_c^{(u,v)}(a_u, a_v) \propto \mathrm{msg}_{\mathrm{above}\, u}(a_u) \cdot \phi_{(u,v)}(a_u, a_v) \cdot \mathrm{CL}_v(a_v) \tag{B.91}$$

where the "message from above $u$" combines the top-down message to $u$ with $u$'s conditional likelihood excluding child $v$.

**Sweep.** One iteration sweeps through all MSA columns $c = 1, \ldots, L$: for each column, recompute the potentials from the current neighbor marginals, run peeling/unpeeling, and store the updated node and edge marginals.

**Properties** The product-of-trees approximation enjoys the same monotonic convergence guarantee as mean-field coordinate ascent (each column update minimises the free energy in its coordinate), with the additional guarantee that the ELBO is at least as tight as the fully-factored mean-field bound. This follows because the product-of-trees family *contains* the mean-field family as a special case (where each $q_c$ is itself fully factored).

**Computational cost.** The dominant cost per sweep is the potential computation: $O(|E| \cdot L \cdot |\mathcal{A}|^4)$ for Match→Match transitions (contracting the $(|\mathcal{A}|, |\mathcal{A}|)$ pairwise marginal against the $(|\mathcal{A}|, |\mathcal{A}|, |\mathcal{A}|, |\mathcal{A}|)$ pair-HMM tensor). The Felsenstein passes add $O(|E| \cdot L \cdot |\mathcal{A}|^2)$, which is subdominant.

##### B.5.4 Stochastic Variational VarAnc (svi-VarAnc)

The VarAnc algorithm of Section B.5.3 computes a variational posterior over ancestral residues for a *fixed* TKF92 model $\theta$ on a *single* fixed-MSA family. We now extend it to a training algorithm that updates $\theta$ on a corpus of tree-structured MSAs. The construction parallels SVI-BW (Section B.1.13) but uses an MSA-conditioned VarAnc E-step in place of the per-pair Pair-HMM Forward–Backward.

**Outer EM.** The outer iteration is a stochastic-EM loop over MSAs. At each iteration:

1. Sample a minibatch of MSAs (with their phylogenetic trees, obtained e.g. by FastTree).
2. For each MSA, run a few sweeps of the VarAnc coordinate-ascent algorithm of Section B.5.3 to refine $q$. This is the *family-level E-step*.
3. Read out, from the converged $q$ and the pairwise edge marginals $q_c^{(u,v)}$, the expected sufficient statistics $(B, D, L, M, S, T, F, E, W, U, V)$ for the TKF92 model:
   - Indel/fragment statistics $(B, D, L, M, S, T, F, E)$ are read column-by-column from the per-edge expected event types (match/insert/delete) implied by the gap structure, weighted by the pairwise edge marginals. These are the same TKF92 sufficient statistics defined in Section A.2.3.
   - Substitution statistics $(W, U, V)$ are accumulated by contracting the pairwise edge marginal $q_c^{(u,v)}(a_u, a_v)$ against the Holmes–Rubin bridge expectations $\mathcal{C}_i^W(a_u, a_v, t_e)$ and $\mathcal{C}_{ij}^U(a_u, a_v, t_e)$ at each match column.
4. Apply the SVI exponential moving average update (equation (B.48)) to the running sufficient statistics, mixing the minibatch contribution with the prior pseudocounts and the previous EMA estimate.
5. Apply the closed-form M-steps of Section A.2.3 (quadratic for $\lambda, \mu$, ratio for $r$, GTR coordinate ascent for $Q, \pi$).

**Comparison to SVI-BW.** The two algorithms differ in their E-step and converge to different fixed points:

- SVI-BW (Section B.1.13) treats each pair as conditionally independent given the population indel/substitution parameters, and sums over alignments with a 2D Pair-HMM Forward–Backward. It uses no tree information.
- svi-VarAnc takes a tree-structured MSA as input (alignment fixed, gap structure fixed, ancestral residues marginalised) and shares information across all branches in the tree at each MSA column.

For pairwise data (a tree of two leaves), the two algorithms coincide. For deep trees with many siblings, svi-VarAnc tightens the indel-rate posterior considerably by coupling sibling pairs through the tree. ELBO monitoring, mini-batch construction, and warm-start initialisation from cherry-trained TKF92 parameters follow the SVI-BW recipe of Section B.1.13 verbatim, replacing the Pair-HMM Forward–Backward E-step with the family-level VarAnc sweep of Section B.5.3.

#### B.6 Mixture-of-trees variational ancestral presence/absence

This appendix derives a variational lower bound on the indel log-likelihood of an MSA under the TKF92 conditional WFST $\mathbb{T}'(\lambda, \mu, T, r)$ of Section A.3 when applied independently on each branch of a phylogenetic tree, combined with a TKF stationary HMM at the root. The bound is parameterised by a tree-structured graphical model on per-column ancestral presence/absence indicators with three irreversible states {NotYetInserted, Present, Deleted}. It is a proper lower bound on the full TKF92 marginal log-likelihood $\log p(\text{MSA} \mid \mathbf{t}, \theta)$, with gap split into a standard $q$-dependent variational KL term and a $q$-independent "restriction" term arising because our latent space cannot represent ancestor residues that get inserted and then fully deleted on every leaf-bound lineage before reaching any leaf (Section B.6.2). For ancestral-presence inference at the observed columns the restriction term is irrelevant, since it cancels from the q-optimal posterior; for parameter learning it does not, and a tighter bound would require explicit modelling of those ghost-column histories. Special-case parameter settings recover an irreversible 3-state analogue of Felsenstein gap-CTMC reconstruction and Fitch parsimony.

##### B.6.1 Setting and approximation

Fix a rooted phylogenetic tree $\mathbf{t}$ with internal-node set $\mathcal{I}$, leaf set $\mathcal{L}$, and a branch length $d_e > 0$ on each edge $e$. We are given a multiple-sequence alignment with $L$ columns whose rows are the leaf sequences of $\mathbf{t}$; let $X_n^v \in \{0, 1\}$ denote the observed presence indicator (1 if leaf $v \in \mathcal{L}$ has a residue in column $n$). The task is to infer per-internal-node presence indicators $\{X_n^v\}_{v \in \mathcal{I}, 1 \le n \le L}$.

We replace the full TKF92 generative model on the tree with the following *per-branch conditional approximation*: each branch $v \to w$ is taken to evolve its row according to the TKF92 conditional WFST $\mathbb{T}'$ acting independently on the whole parent sequence, with no shared latent fragment structure across branches. A residue inserted by the WFST on one branch may therefore be partially deleted, character by character, on a later branch (unlike in the strict TKF92 process, where fragments are indivisible). The WFST $\mathbb{T}'$ is itself a closed-form approximation to the finite-time transition probabilities of the General Geometric Indel (GGI) model — the maximum-entropy CTMC with given expected per-site indel rates and expected inserted-fragment length — and has been shown empirically to fit GGI Gillespie simulations (23) and pairwise-alignment data (30) well. Using it as the per-branch transition law of the tree-level model in this appendix is therefore a small additional approximation on top of the GGI approximation itself.

##### B.6.2 Restricted generative model

The model joint over leaf and internal-node presence patterns at the $L$ MSA columns is

$$p(\text{MSA},\, \{X^v\}_{v\in\mathcal{I}} \mid \mathbf{t}, \theta) = p_{\text{singlet}}\big(X^{\text{root}} \mid \theta\big) \prod_{(v\to w)\in\mathbf{t}} P\big(X^w \mid X^v, d_{vw}, \theta\big), \tag{B.92}$$

where $p_{\text{singlet}}$ is the TKF stationary distribution (Section B.6.3, below) and each $P(X^w \mid X^v, t)$ is the per-branch presence-pattern conditional defined by the WFST path log-likelihood (Section B.6.4). Marginalising (B.92) over internal patterns supported on the $L$ observed MSA columns gives a quantity we denote $\tilde{p}(\text{MSA} \mid \mathbf{t}, \theta)$. The full TKF92 marginal $p(\text{MSA} \mid \mathbf{t}, \theta)$ also sums over evolutionary histories with ancestral residues that get inserted and then fully deleted on every leaf-bound lineage before reaching any leaf — *ghost columns* that leave no MSA trace and so are not representable as values of $X_n^v$ at observed columns $1, \ldots, L$. Since each ghost-column history adds a non-negative probability to $p$ that is absent from $\tilde{p}$,

$$\tilde{p}(\text{MSA} \mid \mathbf{t}, \theta) \;\le\; p(\text{MSA} \mid \mathbf{t}, \theta). \tag{B.93}$$

The gap $\log p - \log \tilde{p} \ge 0$ depends on the model parameters and the tree but not on the variational distribution $q$ introduced below. For the ancestral-presence reconstruction task — fix $\theta$, optimise $q$ to recover internal-node marginals at the observed columns — this gap is a $q$-independent constant and the restricted-model posterior at observed columns coincides with the full-model posterior conditioned on the same support, so all inferences below proceed unchanged.

##### B.6.3 Singlet HMM at the root

The TKF92 stationary HMM (Section A.2.2) emits a sequence of length $L_{\text{root}}$ with

$$p_{\text{singlet}}(L_{\text{root}}) = \begin{cases} 1-\kappa & L_{\text{root}} = 0, \\ \kappa\, p^{L_{\text{root}}-1}\,(1-r)(1-\kappa) & L_{\text{root}} \ge 1, \end{cases} \tag{B.94}$$

where $\kappa = \lambda/\mu$ and $p = r + (1-r)\kappa$. Identifying $L_{\text{root}}$ with the number of MSA columns at which the root is in state $xP$ ($L_{\text{root}} = \sum_n \delta(X_n^{\text{root}} = 1)$) gives the root prior in the joint (B.92). The two-cases form arises because the TKF92 singlet HMM has distinct “from $\mathtt{S}$” (probability $\kappa$ to emit, $1-\kappa$ to end) and “from emitting state” (probability $p$ to extend, $(1-r)(1-\kappa)$ to end) transition contexts.

For TKF91 ($r = 0$): $p = \kappa$ and the two cases collapse to the single form $p_{\text{singlet}}(L_{\text{root}}) = (1-\kappa)\kappa^{L_{\text{root}}}$.

**Note on a tempting shortcut.** One might hope to avoid the singlet by interpreting the root as the output of a pseudo-branch with $T \to \infty$ and an all-absent pseudo-parent. For TKF91 this works — the $t \to \infty$ limit of $\mathbb{T}$ with an empty input does recover the singlet sequence-length distribution. For TKF92 it does *not*: the WFST's $(1-r)$ factors are tied to the fragment-extension structure of the Pair HMM and the limiting output mass for length $L \ge 1$ is $\kappa\, p^{L-1}(1-\kappa)$ rather than the correct $\kappa\, p^{L-1}(1-r)(1-\kappa)$ — i.e. the WFST does not normalise on an empty input under TKF92. We therefore include the singlet term explicitly.

##### B.6.4 Per-branch path log-likelihood

Consider a single branch $v \to w$ of length $d$. Reading the parent and child rows of the MSA column by column gives, for each $n \in \{1, \ldots, L\}$, an ordered pair $(X_n^v, X_n^w) \in \{0,1\}^2$. Map each pair to a WFST column-state $S_n \in \{\mathtt{M}, \mathtt{I}, \mathtt{D}, \mathsf{Ig}\}$ via

$$S_n = \begin{cases} \mathtt{M} & \text{if } (X_n^v, X_n^w) = (1,1) \\ \mathtt{I} & \text{if } (X_n^v, X_n^w) = (0,1) \\ \mathtt{D} & \text{if } (X_n^v, X_n^w) = (1,0) \\ \mathsf{Ig} & \text{if } (X_n^v, X_n^w) = (0,0), \end{cases} \tag{B.95}$$

where the symbol $\mathsf{Ig}$ ("Ignore") marks columns absent in both parent and child and so contributing no transition to the WFST path. Add boundary sentinels $S_0 = \mathtt{S}$ and $S_{L+1} = \mathtt{E}$. Stripping out columns with $S_n = \mathsf{Ig}$ collapses the sequence $(S_0, S_1, \ldots, S_L, S_{L+1})$ to a unique path through the WFST whose nontrivial transitions are between consecutive non-Ignore columns. The branch log-likelihood is the sum of the corresponding entries of $\log \mathbb{T}'(\lambda, \mu, d, r)$ from equation (A.29). Equivalently, summing over each column's incoming transition,

$$\log P(X^w \mid X^v, d, \theta) = \sum_{N=1}^{L+1} \delta(S_N \neq \mathsf{Ig}) \sum_{M=0}^{N-1} \delta(S_M \neq \mathsf{Ig}) \log \mathbb{T}'_{S_M S_N} \prod_{K=M+1}^{N-1} \delta(S_K = \mathsf{Ig}), \tag{B.96}$$

where $\theta = (\lambda, \mu, r)$, $\delta(\cdot)$ is a Kronecker indicator, and the empty product (when $M + 1 > N - 1$) equals 1. For each non-Ignore column $N$ the inner sum picks out the unique $M < N$ that is also non-Ignore with all columns strictly between $M$ and $N$ in state $\mathsf{Ig}$; that pair contributes $\log \mathbb{T}'_{S_M S_N}$ and all other $M$ contribute zero.

##### B.6.5 Tree-structured variational family

We approximate the joint posterior $P(\{X_n^v\}_{v \in \mathcal{I}} \mid \{X_n^v\}_{v \in \mathcal{L}})$ by a product over MSA columns

$$q(\{X_n^v\}_{v \in \mathcal{I}, 1 \leq n \leq L}) = \prod_{n=1}^{L} q_n(\{X_n^v\}_{v \in \mathcal{I}}), \tag{B.97}$$

where each per-column distribution $q_n$ is a directed graphical model on the tree with node values in

$$\mathcal{Z} = \{\textsc{NotYetInserted},\ \textsc{Present},\ \textsc{Deleted}\} \equiv \{xN, xP, xD\}. \tag{B.98}$$

The variable at the root is drawn from a free distribution $q_n^{\text{root}} \in \Delta^{\mathcal{Z}}$; along each edge $v \to w$ the variable $Z_n^w \in \mathcal{Z}$ is drawn conditionally on $Z_n^v$ from a free distribution $q_n^{v \to w}(\cdot \mid \cdot)$. The 9-entry matrix $q_n^{v \to w}(z' \mid z)$ has 4 entries pinned to zero to enforce irreversibility:

$$q_n^{v \to w}(xD \mid xN) = q_n^{v \to w}(xN \mid xP) = q_n^{v \to w}(xN \mid xD) = q_n^{v \to w}(xP \mid xD) = 0. \tag{B.99}$$

The two unconstrained rows ($xN$ and $xP$ parents) each have two surviving outcomes, contributing one free parameter per row; the $xD$ row is deterministic ($q_n^{v \to w}(xD \mid xD) = 1$). This leaves two free parameters per (edge, column). Leaf-node states $Z_n^v$ for $v \in \mathcal{L}$ are clamped: if $X_n^v = 1$, then $Z_n^v = xP$; if $X_n^v = 0$, then $Z_n^v \in \{xN, xD\}$ with the variational mass split between the two by $q_n$.

The presence indicator at internal node $v$ is recovered as

$$X_n^v = \delta(Z_n^v = xP), \tag{B.100}$$

so that any realisation drawn from $q_n$ has the property that the nodes with $X_n^v = 1$ form a connected subtree of $\mathbf{t}$ (or the empty set, which by hypothesis cannot arise once the leaf clamps are respected, since every column of the supplied MSA contains at least one Present leaf).

**Why this support.** Under the strict TKF92 process, each MSA column corresponds to a single insertion event in the history of the tree; the residue descends, and the set of Present nodes is necessarily a connected subtree containing the insertion point. A factorised mean-field $q$ with independent Bernoulli marginals at each $(v, n)$ would assign positive mass to disconnected Present-sets, which (i) are unreachable under the model, and (ii) would force the substitution likelihood to factor as a product of independent Felsenstein recursions on the two pieces — corresponding to two separate insertions co-located in one MSA column, a different generative process. The 3-state irreversible $q$ rules out both pathologies by construction.

##### B.6.6 Expected log-likelihood under $q$

For a parent-child pair $(v, w)$ on the tree, define the per-column state probability under $q_n$:

$$P_q(S_n = \mathtt{M}) = q_n(Z_n^v = xP,\ Z_n^w = xP), \tag{B.101}$$
$$P_q(S_n = \mathtt{D}) = q_n(Z_n^v = xP,\ Z_n^w = xD), \tag{B.102}$$
$$P_q(S_n = \mathtt{I}) = q_n(Z_n^v = xN,\ Z_n^w = xP), \tag{B.103}$$
$$P_q(S_n = \mathsf{lg}) = q_n(Z_n^v = xN,\ Z_n^w = xN) + q_n(Z_n^v = xD,\ Z_n^w = xD). \tag{B.104}$$

The four other $(z, z')$ joint marginals are zero by equation (B.99). Each pairwise marginal $q_n(Z_n^v, Z_n^w)$ is computed by belief propagation (Section B.6.9).

The product factorisation (B.97) makes the per-branch expected log-likelihood factor across columns:

$$\mathbb{E}_q\big[\log P(X^w \mid X^v, d, \theta)\big] = \sum_{s,s'} \log \mathbb{T}'_{ss'}\, W_{ss'}^{(v\to w)}, \tag{B.105}$$

where $s$ ranges over $\{\mathtt{S},\mathtt{M},\mathtt{I},\mathtt{D},\mathtt{E}\} \setminus \{\mathtt{E}\}$ and $s'$ ranges over $\{\mathtt{S},\mathtt{M},\mathtt{I},\mathtt{D},\mathtt{E}\} \setminus \{\mathtt{S}\}$ (so $\mathtt{S}$ enters only as a source and $\mathtt{E}$ only as a destination, both via the deterministic boundary sentinels), and

$$W_{ss'}^{(v\to w)} = \sum_{N=1}^{L+1} \sum_{M=0}^{N-1} P_q(S_M = s)\, P_q(S_N = s') \prod_{K=M+1}^{N-1} P_q(S_K = \mathsf{lg}) \tag{B.106}$$

is the expected number of times the path uses transition $s \to s'$ on this branch. Boundary terms have $P_q(S_0 = \mathtt{S}) = 1$ and $P_q(S_{L+1} = \mathtt{E}) = 1$ deterministically.

**Root-prior contribution.** Under the per-column factorisation (B.97), the expected singlet log-likelihood at the root (equation (B.94)) reduces to

$$\mathbb{E}_q\big[\log p_{\text{singlet}}(X^{\text{root}})\big] = \log p \cdot \sum_{n=1}^{L} q_n(X_n^{\text{root}} = 1) + \big(1 - P_q(L_{\text{root}}{=}0)\big)\, c_1 + P_q(L_{\text{root}}{=}0)\, \log(1-\kappa), \tag{B.107}$$

with $c_1 = \log\big[\kappa(1-r)(1-\kappa)/p\big]$ and $P_q(L_{\text{root}}{=}0) = \prod_{n=1}^{L} q_n(X_n^{\text{root}} = 0)$ the variational mass on an empty root sequence. For TKF91 ($r = 0$) the second and third terms collapse to a single $q$-independent constant $\log(1-\kappa)$ and (B.107) reduces to the textbook form $\log(1-\kappa) + \log\kappa \cdot \sum_n q_n(X_n^{\text{root}} = 1)$. For TKF92 the $L_{\text{root}}{=}0$ special case must be retained; note that $P_q(L_{\text{root}}{=}0)$ is small but generically nonzero, since some columns may have positive variational mass on root being $xN$ (the column was inserted on a strict subtree below the root).

##### B.6.7 ELBO

For any product-of-trees $q$ that gives joint marginals $q_n(Z_n^v, Z_n^w)$ at every (edge, column),

$$\log p(\text{MSA} \mid \mathbf{t}, \theta) \;\geq\; \log \tilde{p}(\text{MSA} \mid \mathbf{t}, \theta) \;\geq\; \mathcal{L}[q] \;=\; \mathbb{E}_q\big[\log p_{\text{singlet}}(X^{\text{root}})\big] + \sum_{(v \to w) \in \mathbf{t}} \mathbb{E}_q\big[\log P(X^w \mid X^v, d_{vw}, \theta)\big] + \text{[illegible]} \tag{B.108}$$

The first two terms are given by (B.107) and (B.105) respectively; the third is the entropy of the leaf-conditioned variational distribution. The leftmost inequality is (B.93); the right inequality is the standard Jensen bound on the restricted joint (B.92).

**Entropy decomposition.** The variational $q$ is parameterised through a directed graphical model on the full tree (root + internal + leaf nodes) with edge conditionals $q_n^{v \to w}(\cdot \mid \cdot)$. Writing $q_{\text{joint}}$ for this joint and conditioning on the observed leaf states gives $q(\{X^v\}_{v \in \mathcal{I}} \mid \text{MSA}) = q_{\text{joint}}(\text{internals}, \text{MSA}) \,/\, Z_q$, where $Z_q = \sum_{\text{internals}} q_{\text{joint}}$ is the per-column belief-propagation partition function summed over columns. The leaf-conditioned entropy then decomposes as

$$H\big[q(\text{internals} \mid \text{MSA})\big] = -\,\mathbb{E}_q\big[\log q_{\text{joint}}\big] + \log Z_q, \tag{B.109}$$

where the first term factors over the directed graphical model:

$$-\mathbb{E}_q\big[\log q_{\text{joint}}\big] = -\sum_{n=1}^{L}\sum_{z \in \mathcal{Z}} q_n^{\text{root}}(z) \log q_n^{\text{root}}(z) - \sum_{(v \to w) \in \mathbf{t}} \sum_{n=1}^{L} \sum_{z, z' \in \mathcal{Z}} q_n(Z_n^v{=}z, Z_n^w{=}z') \log q_n^{v \to w}(z' \mid z). \tag{B.110}$$

The first term is the per-column root entropy. Each per-edge term is a *cross-entropy* between the BP-derived joint marginal $q_n(Z_n^v, Z_n^w)$ and the prior edge conditional $q_n^{v \to w}$; it is *not* the parent-marginal-weighted entropy $\mathbb{E}_{q_n(Z_n^v)}\big[H(q_n^{v \to w}(\cdot \mid Z_n^v))\big]$, which only equals (B.110) when the leaves impose no evidence on the edge — generically a different quantity once the BP-conditioning shifts pair marginals away from the prior factorisation. The $\log Z_q$ term in (B.109) comes out of the up-pass partition function and ensures the entropy is on the conditioned distribution rather than the joint.

**Bound interpretation.** Equation (B.108) is a proper lower bound on the full TKF92 marginal log-likelihood $\log p(\text{MSA})$, with the gap split into two non-negative pieces:

$$\log p(\text{MSA}) - \mathcal{L}[q] \;=\; \underbrace{\big(\log p(\text{MSA}) - \log \tilde{p}(\text{MSA})\big)}_{\text{restriction gap}} \;+\; \underbrace{\mathrm{KL}\big[q(\text{internals} \mid \text{MSA}) \,\|\, \tilde{p}(\text{internals} \mid \text{MSA})\big]}_{\text{variational gap}}. \tag{B.111}$$

The variational gap is the standard Jensen slack on the restricted joint (B.92) and depends on $q$; the restriction gap is the contribution of ghost-column histories that our latent space cannot represent (Section B.6.2) and depends on the model parameters $\theta$ and the tree but not on $q$. For ancestral-presence reconstruction (fix $\theta$, optimise $q$) the restriction gap is a constant additive offset, so maximising (B.108) is equivalent to minimising the KL divergence from $q$ to the true model posterior on the support of observed-column internal-node configurations. For parameter learning over $\theta$ this equivalence breaks: the restriction gap shifts with $\theta$, so the bound is not tight up to a parameter-independent constant. A tighter bound would require augmenting $q$ to model ghost columns explicitly — for instance via per-(branch, gap-region) latent counts of inserted-then-fully-deleted residues — at the cost of additional machinery beyond the per-MSA-column factorisation used here. Equivalently and intuitively, one recovers the full marginal in the limit of inserting infinitely many additional "empty" columns into the alignment between (and around) every observed column, with all leaves clamped to absent at those columns and the variational $q$ free over their internal states; the present formulation simply truncates this padding to zero.

##### B.6.8 Stable computation: cumulant trick

The expected transition count $W_{ss'}^{(v \to w)}$ in (B.106) appears at first sight to require $O(L^2)$ work per (branch, state-pair). A standard prefix-sum reduces this to $O(L)$.

For brevity drop the branch superscript and write $P_n^{\mathsf{Ig}} = P_q(S_n = \mathsf{Ig})$, $P_n^s = P_q(S_n = s)$ for $s \in \{\mathtt{S}, \mathtt{M}, \mathtt{D}, \mathtt{I}, \mathtt{E}\}$ (with $P_0^{\mathtt{S}} = 1$, $P_{L+1}^{\mathtt{E}} = 1$, all other boundary values zero). Define the running log-Ignore mass

$$C_N \;=\; \sum_{k=1}^{N} \log P_k^{\mathsf{Ig}}, \qquad C_0 = 0. \tag{B.112}$$

Then $\prod_{K=M+1}^{N-1} P_K^{\mathsf{Ig}} = \exp(C_{N-1} - C_M)$, and (B.106) factorises as

$$W_{ss'} \;=\; \sum_{N=1}^{L+1} P_N^{s'} \, e^{C_{N-1}} \;\cdot\; \underbrace{\sum_{M=0}^{N-1} P_M^{s} \, e^{-C_M}}_{\text{prefix sum in } M}. \tag{B.113}$$

Both inner factors are non-negative; the difference $C_{N-1} - C_M$ is non-positive (it is a sum of log-probabilities), so the geometric factor lies in $(0, 1]$ and (B.113) is numerically benign in float64 for any practical $L$. For very long alignments where individual exponents could underflow, one $\epsilon$-floors $P_n^{\mathsf{Ig}}$ before computing $C_n$; the floor is harmless because the product weights paths that traverse many Ignore columns, which contribute negligibly to the expectation.

The total work for the per-branch expected log-likelihood is $O(L \cdot |S|^2)$ with $|S| \leq 5$ states, yielding $O(BL)$ overall for $B$ branches. This is fully vectorisable across columns and branches.

**Why the inner sum is in probability space.** Equation (B.96) for a fixed deterministic realisation of the indicators is purely a sum of $\log \mathbb{T}'$ entries: no probabilities, no products, no exponentials. The probability-space arithmetic in (B.113) arises only when we marginalise over the variational distribution: each summand is a probability mass (product of independent column factors) times a real-valued $\log \mathbb{T}'$ entry, and the sum collapses these weighted log-transitions into the expected transition count.

##### B.6.9 Belief propagation for pairwise marginals

The pairwise marginals $q_n(Z_n^v, Z_n^w)$ feeding equations (B.101) are computed by the standard Felsenstein up-down algorithm on $\mathbf{t}$. For each column $n$:

1. *Up pass:* for every node $v$ and every state $z \in \mathcal{Z}$, accumulate $\beta_v^n(z) = \prod_{w:\text{child of } v} \sum_{z'} q_n^{v \to w}(z' \mid z)\, \beta_w^n(z')$, with leaf base case $\beta_v^n(z) = \delta(\text{leaf clamp at } v, n \text{ is consistent with } z)$.
2. *Down pass:* for every internal $v$ and state $z$, accumulate the root-conditioned posterior $\alpha_v^n(z)$ in standard fashion.
3. *Marginals:* for every edge $v \to w$ and every state pair, $q_n(Z_n^v = z, Z_n^w = z') = \alpha_v^n(z)\, q_n^{v \to w}(z' \mid z)\, \beta_w^n(z') / \mathcal{Z}_n$, where $\mathcal{Z}_n$ is the per-column partition function from the up pass at the root.

The up and down passes are $O(|\mathcal{I}| \cdot |\mathcal{Z}|^2)$ per column and trivially vectorise across columns (same tree topology, different leaf clamps and different per-column edge conditionals). For typical protein alignments ($L \approx 360$ amino acids on average in UniProtKB/Swiss-Prot (49), with the long tail extending to a few thousand) this is unproblematic; for DNA MSAs (gene-scale regions in the few-kbp range (53), longer for syntenic blocks) the per-column independence keeps cost linear in $L$.

##### B.6.10 Special cases and scalability

**Irreversible Felsenstein-3.** Tie the per-(edge, column) variational conditionals $q_n^{v\to w}(\cdot \mid \cdot)$ across columns to a single per-edge transition matrix $\bar{q}^{v\to w}(\cdot \mid \cdot)$ — equivalently, parameterise each $\bar{q}^{v\to w}$ as the transition matrix of an irreversible 3-state CTMC over $(xN, xP, xD)$ with rates fitted by maximum likelihood on the leaf data. The variational marginals $q_n$ are then exactly the Felsenstein posteriors of this CTMC at column $n$, and the ELBO reduces to the log-likelihood of the leaf clamps under that CTMC plus the (now column-independent) expected log of $\mathbb{T}'$.

**Fitch parsimony.** The zero-temperature limit of the irreversible Felsenstein-3 posterior — assigning each internal node deterministically to its most-likely state under the CTMC — recovers the Fitch parsimony labelling of presence/absence under uniform priors and unit transition costs.

**Scalability of stochastic ELBO optimisation.** The free-parameter count scales as $2 \cdot |\mathcal{E}| \cdot L$ (plus a per-column root parameter) where $|\mathcal{E}|$ is the number of tree edges and $L$ is the number of MSA columns. The variational $q$ factorises over columns, but the ELBO is non-trivially coupled across columns through the prefix-of-Ignore factor in equation (B.113), which weights each $(M, N)$ transition contribution by the chain of intervening Ignore probabilities. Match-rich columns (where most lineages are clamped to $xP$) have $P_n^{\mathsf{Ig}} \approx 0$ and so act as anchors that break the chain: contributions from $(M, N)$ pairs that straddle such an anchor column are damped to zero, decoupling the gradient on either side. For both protein-scale and DNA-scale alignments with non-trivial column coverage, ELBO optimisation should therefore scale linearly with total residue count without running into long-range coupling pathologies.

#### B.7 Theory: structural bias of the BP cumulant under column-factorised $q$

We give a precise derivation of the structural bias displayed empirically in Section **??**. The bias is not a finite-data noise artifact — it is a property of the column-factorised variational family used by SVI-VarAnc, and persists at any $q$ whose per-column marginals match the truth.

**Setup.** On a single branch $e$ of length $t_e$, the per-branch WFST is a chain HMM over the 5 states $\{S, M, I, D, E\}$ with chain transition matrix $\mathbf{T}_e[s, s']$ given by the standard TKF91 / TKF92 closed forms (`tkf92_wfst_T`, eq. A.29). The branch's contribution to the data log-likelihood is $\sum_{n=1}^{L+1} \log T_e[\zeta_{n-1}, \zeta_n]$ where $\boldsymbol{\zeta} = (\zeta_0, \ldots, \zeta_{L+1})$ is the chain trajectory across the $L+2$ columns (including boundaries). Conventionally $\zeta_n \in \{S, M, I, D, E, \mathrm{Ig}\}$ where Ig denotes a non-emitting "insertion-gap" column where the branch is in NYI/NYI or $D/D$ joint state and contributes nothing to the chain transitions; chain transitions are indexed by consecutive non-Ig columns.

The variational $q$ factorises across columns: $q(\boldsymbol{\zeta}) = \prod_{n=1}^{L} q_n(\zeta_n)$ with each $q_n$ a tree-structured distribution over edge presence/absence carrying the leaf clamps of column $n$. Define the per-column WFST-state probabilities at branch $e$:

$$P_n^{(e)}(s) = q_n(\text{branch state } e \text{ is } s), \quad s \in \{M, I, D\}, \qquad P_n^{(e)}(\mathrm{Ig}) = q_n(\mathrm{NYI}, \mathrm{NYI}) + q_n(D, D). \tag{B.114}$$

**The cumulant formula and what it computes exactly.** Section A.2.3 introduces the BP cumulant $\mathbf{W}_e \in \mathbb{R}^{5\times 5}$ as the sufficient statistic the M-step ingests:

$$W_e[s,s'] = \sum_{1\le M<N\le L+1} P_M^{(e)}(s)\cdot P_N^{(e)}(s')\cdot \prod_{k=M+1}^{N-1} P_k^{(e)}(\mathrm{Ig}). \tag{B.115}$$

Under the factorised $q$, $W_e[s,s']$ is the *exact* expected count of WFST chain transitions $s \to s'$ on branch $e$:

$$\begin{aligned}\mathbb{E}_q\big[\#\{\text{chain transitions } s\to s'\}\big] &= \sum_{M<N} q\big(\zeta_M = s, \zeta_N = s', \zeta_{M+1} = \cdots = \zeta_{N-1} = \mathrm{Ig}\big)\\ &= \sum_{M<N} P_M(s)\, P_N(s') \prod_{k=M+1}^{N-1} P_k(\mathrm{Ig}) \quad = \quad W_e[s,s'],\end{aligned}$$

where the second equality uses the column-factorisation of $q$. The sum over $(M,N)$ counts each chain transition once because of the explicit Ig-padding constraint: any single trajectory realisation $\boldsymbol{\zeta}$ with non-Ig columns at indices $n_1 < n_2 < \cdots < n_K$ contributes weight 1 to exactly the adjacent-pair transitions $(n_0,n_1),(n_1,n_2),\ldots,(n_K,n_{K+1})$ (with $n_0 = 0, n_{K+1} = L+1$); for any other $(M,N)$ pair, at least one intervening column is non-Ig and the indicator product is zero. This $O(L)$ prefix-sum implementation agrees with the $O(L^2)$ direct sum to machine precision ($6.7\times 10^{-16}$).

**The structural bias.** The factorisation assumption is *not* satisfied by the truth posterior $p^*(\boldsymbol{\zeta} \mid \text{data})$, even for TKF91 ($r = 0$). Although TKF91's generative model is per-position i.i.d., the per-branch WFST chain has nonzero transition probabilities $T_e[D,D]$, $T_e[I,I]$, $T_e[I,D]$ and so on (closed-form expressions in (A.29) reduce, at $r = 0$, to the standard TKF91 Pair HMM weights). These chain memories propagate from data conditioning into a non-factorised joint posterior:

$$p^*\big(\zeta_M = s, \zeta_N = s', \zeta_{M+1..N-1} = \mathrm{Ig} \mid \text{data}\big) \;\ne\; \prod_n p^*\big(\zeta_n \mid \text{data}\big), \tag{B.116}$$

even though the per-column truth marginal $p^*(\zeta_n \mid \text{data})$ is exactly recoverable by per-column Felsenstein BP. The bias of the cumulant against the truth-posterior chain expectation is the integrated difference between the factorised-q product of marginals and the truth-posterior joint:

$$\boxed{W_e[s,s'] - \mathbb{E}_{p^*}\big[\#\{s\to s'\} \bigm| \text{data}\big] = \sum_{M<N}\Big[p_M^*(s)\, p_N^*(s') \prod_{k=M+1}^{N-1} p_k^*(\mathrm{Ig}) \;-\; p^*\big(\zeta_M = s, \zeta_N = s', \zeta_{M+1..N-1} = \mathrm{Ig} \bigm| \text{data}\big) \text{[illegible]}} \tag{B.117}$$

The bracketed term is the connected $(N - M + 1)$-way correlation of the truth posterior over the columns $\{M, M+1, \ldots, N\}$; it vanishes only when the truth posterior factorises across these columns. For TKF91, $T_e[D,D] > 0$ and $T_e[I,I] > 0$ at $r = 0$ alone suffice to make the right-hand side of (B.117) nonzero on the $(s,s') = (I,I)$ and $(D,D)$ entries: an opened gap is positively correlated with continued gap by the chain transition kernel itself, even before any explicit fragment-extension parameter is invoked.

**Entropy reward exacerbates the bias on rare entries.** The ELBO objective decomposes as $\mathrm{ELBO} = \mathbb{E}_q[\log p(z, \text{data}; \theta)] - \mathbb{E}_q[\log q]$ with the entropy term $-\mathbb{E}_q[\log q]$ rewarding diffuse $q$. At ELBO stationarity, $q$'s per-column marginals balance the data-fit term against the entropy term;

for tied or weakly-fitted logits, the resulting $q$ is slightly more diffuse than the truth posterior on individual rare states (i.e., $P_n^{(e)}(I)$ above the per-column truth marginal $p_n^*(I)$). Through the cumulant (B.115), even a small per-column inflation $P_n(s) = p_n^*(s) + \epsilon$ contributes a leading linear-in-$\epsilon$ bias on $W[s, s']$ with positive coefficient $\partial W / \partial P_n(s) = \sum_{N>n} P_N^{(e)}(s') \prod_{k=n+1}^{N-1} P_k^{(e)}(\text{Ig}) > 0$, summed over $L$ source columns; the cumulant amplifies per-column diffuseness via its $O(L^2)$ pair structure.

**Why the EM converges to a fixed point.** The EM map $\theta \mapsto \text{MLE}(\mathbf{W}(\theta))$ is self-consistent at the biased fixed point $\theta_\infty$ when the cumulant bias on $\theta$ saturates. Each entry of the cumulant is bounded uniformly in $\theta$: $W_e[s, s'] \leq L + 1$ on a single branch (each $(M, N)$ pair contributes at most 1, and the geometric Ig-product is summable), giving a $\theta$-independent upper bound on the aggregated sufficient-statistic count. The exposure denominator $T_{\text{eff}} = \sum_e t_e$ in the BDI rate-MLE (`tkf92_vbem.py:223`) is itself $\theta$-independent — it depends only on the tree branch lengths, not on the inferred rates — and is bounded below by the smallest tree branch. Composing these, the rate-MLE $\hat{\lambda}(\theta) \propto \sum_e W[\cdot, I]/T_{\text{eff}}$ (with the precise BDI quadratic-MLE form in `tkf92_vbem.py:m_step_indel_quadratic`) is a continuous, bounded self-map on a compact rate interval $[0, \bar{\lambda}]$ with $\bar{\lambda} \leq (L+1)/\min_e t_e$ (continuity follows because $W(\theta)$ depends continuously on $\theta$ via the per-edge TKF92 transition matrix (A.29), and the M-step quadratic root is continuous in its coefficients). Brouwer's fixed-point theorem applies: a fixed point $\theta_\infty$ exists. Consistent with this boundedness argument, the EM trajectory of Section **??** reaches a finite biased fixed point and does not diverge.

**Implications for parameter inference.** The structural bias is a property of any inference scheme that uses the cumulant tensor $\mathbf{W}$ as a sufficient statistic under a column-factorised variational family. It does not affect: (i) the ELBO value itself (which is a coherent $\log p(\text{data})$ lower bound regardless of what the cumulant looks like), nor (ii) ancestral indel-presence reconstruction (Section **??**: VarAnc and Fitch parsimony are statistically indistinguishable on the unified Pfam test splits, $|\Delta\text{F1}| < 0.006$ across short / hard / xhard), nor (iii) downstream MSA-conditioned rate estimation via exact-EM (Maraschino, Section **??**). It does affect any rate estimate computed from the variational cumulant, including the SVI-VarAnc refinement of cherry-trained parameters (Section **??**), where parameters drift toward $\theta_\infty$ rather than toward truth. We report both initial cherry-trained and $\theta_\infty$ estimates without correction; the structural bias is the principal known limitation of variational training in this paper.

### C Recursive TKF

MixDom adds two levels of nested mixture structure on top of TKF92: a top-level TKF91 process governing domain births and deaths, and per-domain TKF92 processes governing fragments within. Each fragment also carries a substitution-class index. The exact Baum–Welch M-step proceeds via a six-step chain-restoration identity through a fully exploded null-state model. The same nesting pattern — a parent BDI / TKF process emitting child fragments that themselves carry latent class / domain / structure indices — supports a family of recursive TKF models that this appendix collects: MixDom, its order-1 Maraschino adjacency distillation, the algebraic full-Woodbury distillation, the MixDom-specific SVI-BW convergence theory, the tree-level VEM and ancestral-reconstruction algorithms, the generalised phylo-HMM, the labeled-MixDom Singlet and WFST, the formal recursive-grammar-elaboration rules, and four worked recursive examples (L-TKF, TKFST, TKFStack, TKF-Genome).

#### C.1 The TKF-Mixed Domain Model (MixDom)

This model was preliminarily described and empirically evaluated in (30).

##### C.1.1 The MixDom Model

The Mixture of TKF92 Domains (MixDom) is a multiply-nested hierarchical mixture model. At the top level is a TKF91-like links process where each link is associated with a domain of random type $n \sim \text{Categorical}(v_1, \ldots, v_{\mathcal{N}})$. Each top-level link emits its own domain sequence, with model parameters determined by domain type.

**Three nesting levels.** MixDom is generated as three nested processes:

1. **Top-level TKF91 over domains**: a sequence of top-level links, each of domain type $n \sim \text{Categorical}(v_1, \ldots, v_{\mathcal{N}})$, evolving under its own per-domain TKF92 indel process.
2. **Per-domain TKF92 over fragments**: within each domain of type $n$, a TKF92 process generates a sequence of nested links. Each such nested link is called a *fragment*. Different fragments are statistically independent.
3. **Intra-fragment Markov chain on fragment-types**: a single fragment consists of a sequence of fragment-type characters drawn from a per-domain Markov chain with $\mathcal{F}+2$ states (start, end, and $\mathcal{F}$ emitting states $f \in \{1, \ldots, \mathcal{F}\}$). The initial fragment-type is drawn from $w_{nf}$. From type $f$, the chain advances within the current fragment to type $g$ with probability $r_{fg}^{(n)}$, or terminates the fragment with probability $\rho_f^{(n)} = 1 - \sum_g r_{fg}^{(n)}$.

The chain restarts at Start for each fresh fragment; different fragments are independent Markov realisations.

Each emitted site within a fragment of domain $n$ and fragment-type $f$ independently draws a site class $c \sim \text{Categorical}(u_{nf1}, \ldots, u_{nfC})$, with $C$ the number of site classes. The site is then governed by the substitution process $\text{Subst}(Q^{(c)}, \pi^{(c)})$.

We denote the per-domain process (TKF92 fragments with an intra-fragment fragment-type chain and per-(domain, fragment-type) site-class mixture) by $\text{HMM}(\{Q^{(c)}, \pi^{(c)}\}_c; r^{(n)}, u_n)$. The full MixDom model is then

$$\begin{aligned} M_{\text{dom}} &= \text{Links}(\text{HMM}(\{Q^{(c)}, \pi^{(c)}\}_c; r^{(n)}, u_n); \lambda_n, \mu_n) \\ \text{MixDom} &= \text{Links}(\text{Mix}_{n \sim v_n}(M_{\text{dom}}); \lambda_0, \mu_0) \end{aligned}$$

where Mix(...) denotes a mixture model, with weights $p$, over parameters $\theta$ for model $M$

$$x \sim \text{Mix}_{k \sim p}(M(\theta_k); p) \;\Leftrightarrow\; k \sim \text{Categorical}(p),\; x \sim M(\theta_k)$$

*Remark* C.1 (Relationship to TKF92). When $\mathcal{F} = 1$ and $C = 1$, each fragment's intra-fragment chain has a single emitting state, so the fragment length is $\text{Geometric}(r_{11}^{(n)})$ and all sites share the single substitution model $\text{Subst}(Q, \pi)$, recovering TKF92. In general, the $\mathcal{F} \times \mathcal{F}$ transition matrix $r^{(n)}$ allows *intra-fragment correlations* between the fragment-types of adjacent positions within a single fragment (e.g., fragment-type 1 positions tend to follow fragment-type 1 positions when $r_{11}^{(n)}$ is the dominant row entry), and the per-(fragment-type) class distributions $u_{nfc}$ allow the resulting substitution patterns to vary with fragment-type. *Different fragments remain statistically independent under this scheme*; Markov correlations are strictly *within* a fragment, carried by the fragment-type chain.

##### C.1.2 Singlet HMM for MixDom

The Singlet HMM generates sequences from the stationary distribution. Each domain may be empty with probability $\varphi_n \equiv 1-\kappa_n$, so the probability that a top-level link generates a zero-length domain sequence is $z_0 = \sum_{n\in\mathcal{N}} v_n \varphi_n$. This leads to null cycles (a link is entered but immediately terminates). Eliminating these—by the Schur complement procedure described in the next section—yields a collapsed Singlet HMM with state space $\Psi^{(\mathrm{eqm})} = \{\mathtt{S},\mathtt{E}\} \cup \{\mathtt{I}_{nf} : n \in \mathcal{N}, f \in \mathcal{F}\}$ ($\mathcal{N}\mathcal{F}+2$ states). Each emitting state $\mathtt{I}_{nf}$ emits a character from $\sum_c u_{nfc}\pi^{(c)}$. The transition matrix $\mathcal{T}^{(\mathrm{eqm})}$ for this collapsed Singlet HMM has entries

$$\mathcal{T}^{(\mathrm{eqm})}_{\mathtt{I}_{lf},\mathtt{I}_{mg}} = \frac{\rho^{(l)}_f \varphi_l \cdot \kappa_0 \cdot v_m \kappa_m w_{mg}}{1-\kappa_0 z_0} + \delta(l{=}m)\,\rho^{(l)}_f \kappa_l w_{lg} + \delta(l{=}m)\,r^{(l)}_{fg}$$

where $\rho^{(n)}_f = 1 - \sum_g r^{(n)}_{fg}$ is the fragment termination probability, and the three terms represent (respectively) inter-domain transitions via the null-corrected top-level geometric process, same-domain new-fragment transitions, and same-domain intra-fragment Markov transitions on fragment-types. The $\mathtt{S}$ row is $\mathcal{T}^{(\mathrm{eqm})}_{\mathtt{S},\mathtt{I}_{mg}} = \kappa_0 v_m \kappa_m w_{mg}/(1-\kappa_0 z_0)$ and $\mathcal{T}^{(\mathrm{eqm})}_{\mathtt{S},\mathtt{E}} = (1-\kappa_0)/(1-\kappa_0 z_0)$; the $\mathtt{E}$ column is $\mathcal{T}^{(\mathrm{eqm})}_{\mathtt{I}_{lf},\mathtt{E}} = \rho^{(l)}_f \varphi_l (1-\kappa_0)/(1-\kappa_0 z_0)$.

##### C.1.3 Pair HMM for MixDom

In the joint TKF92 Pair HMM, the start→end weight is $\tau(\lambda,\mu,T)_{\mathtt{SE}}$. In MixDom, the probability that a $\mathtt{M}$ state emits no sequence is obtained by summing this TKF92 null output probability over domain types, $z_T = \sum_{n\in\mathcal{N}} v_n \tau^{(n)}_{\mathtt{SE}}$ where $\tau^{(n)} \equiv \tau(\lambda_n,\mu_n,T)$. The Singlet HMM's null probability $z_0$ also appears in the Pair HMM, governing the $\mathtt{I}$ and $\mathtt{D}$ state null outputs (which involve only one sequence).

Start with the Pair HMM and split the $\mathtt{M}$, $\mathtt{I}$, and $\mathtt{D}$ states into non-emitting and emitting states. Let $\mathtt{A}$, $\mathtt{B}$, $\mathtt{C}$ denote the separated empty-match, empty-insert, and empty-delete states, respectively. The $8\times 8$ transition matrix for this null-separated joint pair HMM is

$$\upsilon = \left(\begin{array}{c|cccccccc} & \mathtt{S} & \mathtt{M} & \mathtt{I} & \mathtt{D} & \mathtt{E} & \mathtt{A} & \mathtt{B} & \mathtt{C} \\ \hline \mathtt{S} & 0 & (1-z_T)\tau_{\mathtt{SM}} & (1-z_0)\tau_{\mathtt{SI}} & (1-z_0)\tau_{\mathtt{SD}} & \tau_{\mathtt{SE}} & z_T\tau_{\mathtt{SM}} & z_0\tau_{\mathtt{SI}} & z_0\tau_{\mathtt{SD}} \\ \mathtt{M} & 0 & (1-z_T)\tau_{\mathtt{MM}} & (1-z_0)\tau_{\mathtt{MI}} & (1-z_0)\tau_{\mathtt{MD}} & \tau_{\mathtt{ME}} & z_T\tau_{\mathtt{MM}} & z_0\tau_{\mathtt{MI}} & z_0\tau_{\mathtt{MD}} \\ \mathtt{I} & 0 & (1-z_T)\tau_{\mathtt{IM}} & (1-z_0)\tau_{\mathtt{II}} & (1-z_0)\tau_{\mathtt{ID}} & \tau_{\mathtt{IE}} & z_T\tau_{\mathtt{IM}} & z_0\tau_{\mathtt{II}} & z_0\tau_{\mathtt{ID}} \\ \mathtt{D} & 0 & (1-z_T)\tau_{\mathtt{DM}} & (1-z_0)\tau_{\mathtt{DI}} & (1-z_0)\tau_{\mathtt{DD}} & \tau_{\mathtt{DE}} & z_T\tau_{\mathtt{DM}} & z_0\tau_{\mathtt{DI}} & z_0\tau_{\mathtt{DD}} \\ \mathtt{E} & 0 & 0 & 0 & 0 & 0 & 0 & 0 & 0 \\ \mathtt{A} & 0 & (1-z_T)\tau_{\mathtt{MM}} & (1-z_0)\tau_{\mathtt{MI}} & (1-z_0)\tau_{\mathtt{MD}} & \tau_{\mathtt{ME}} & z_T\tau_{\mathtt{MM}} & z_0\tau_{\mathtt{MI}} & z_0\tau_{\mathtt{MD}} \\ \mathtt{B} & 0 & (1-z_T)\tau_{\mathtt{IM}} & (1-z_0)\tau_{\mathtt{II}} & (1-z_0)\tau_{\mathtt{ID}} & \tau_{\mathtt{IE}} & z_T\tau_{\mathtt{IM}} & z_0\tau_{\mathtt{II}} & z_0\tau_{\mathtt{ID}} \\ \mathtt{C} & 0 & (1-z_T)\tau_{\mathtt{DM}} & (1-z_0)\tau_{\mathtt{DI}} & (1-z_0)\tau_{\mathtt{DD}} & \tau_{\mathtt{DE}} & z_T\tau_{\mathtt{DM}} & z_0\tau_{\mathtt{DI}} & z_0\tau_{\mathtt{DD}} \end{array}\right) \tag{C.1}$$

with $\tau \equiv \tau(\lambda_0,\mu_0,T)$; or, elementwise,

$$\upsilon_{ij} = \begin{cases} 0 & \text{if } j = \mathtt{S} \\ (1-z_T)\tau_{i\mathtt{M}} & \text{if } j = \mathtt{M} \\ (1-z_0)\tau_{ij} & \text{if } j \in \{\mathtt{I},\mathtt{D}\} \\ \tau_{i\mathtt{E}} & \text{if } j = \mathtt{E} \\ z_T\tau_{i\mathtt{M}} & \text{if } j = \mathtt{A} \\ z_0\tau_{i\mathtt{I}} & \text{if } j = \mathtt{B} \\ z_0\tau_{i\mathtt{D}} & \text{if } j = \mathtt{C} \end{cases} \tag{C.2}$$

Let $\mathcal{U}_{\Phi_1,\Phi_2}$ be the matrix formed by zeroing all rows except $i \in \Phi_1$ and all columns except $j \in \Phi_2$ of $\upsilon$. Consider the matrix of transitions between the empty states $\mathtt{A}, \mathtt{B}, \mathtt{C}$

$$\mathcal{U}_{\mathtt{ABC},\mathtt{ABC}} = \left( \begin{array}{c|cccc} & \dots & \mathtt{A} & \mathtt{B} & \mathtt{C} \\ \hline \dots & \dots & \dots & \dots & \dots \\ \mathtt{A} & \dots & z_T \tau_{\mathtt{MM}} & z_0 \tau_{\mathtt{MI}} & z_0 \tau_{\mathtt{MD}} \\ \mathtt{B} & \dots & z_T \tau_{\mathtt{IM}} & z_0 \tau_{\mathtt{II}} & z_0 \tau_{\mathtt{ID}} \\ \mathtt{C} & \dots & z_T \tau_{\mathtt{DM}} & z_0 \tau_{\mathtt{DI}} & z_0 \tau_{\mathtt{DD}} \end{array} \right)$$

where the unshown rows and columns (corresponding to transitions to, from, or between $\mathtt{S}, \mathtt{M}, \mathtt{I}, \mathtt{D}, \mathtt{E}$) have zero entries. Summing over paths of all (including zero) lengths via $\mathtt{A}, \mathtt{B}, \mathtt{C}$ (i.e. performing the Schur complement) yields

$$\sum_{k=0}^{\infty} \mathcal{U}^k_{\mathtt{ABC},\mathtt{ABC}} = (I - \mathcal{U}_{\mathtt{ABC},\mathtt{ABC}})^{-1} = \mathcal{Z}$$

which has a relatively simple closed form (reducing to $3\times3$ matrix inversion). The effective nonempty $5 \times 5$ transition matrix (with $\mathtt{A}, \mathtt{B}, \mathtt{C}$ summed out) is

$$\mathcal{T} \equiv \mathcal{T}(\theta, T) = \mathcal{U}_{\mathtt{SMIDE},\mathtt{SMIDE}} \; + \; \mathcal{U}_{\mathtt{SMIDE},\mathtt{ABC}} \cdot \mathcal{Z} \cdot \mathcal{U}_{\mathtt{ABC},\mathtt{SMIDE}}$$

where $\theta = (\lambda_0, \mu_0, \{\upsilon_n\}, \{\lambda_n, \mu_n\}, \{w_{nf}\}, \{r^{(n)}_{fg}\}, \{u_{nfc}\}, \{Q^{(c)}, \pi^{(c)}\})$.

Following null state elimination, the collapsed Pair HMM has $5\mathcal{N}\mathcal{F} + 2$ states, namely

$$\Psi = \{\mathtt{SS}, \mathtt{EE}\} \cup \{\mathtt{UX}_{lf} : \mathtt{UX} \in \{\mathtt{MM}, \mathtt{MI}, \mathtt{MD}, \mathtt{II}, \mathtt{DD}\}, 1 \leq l \leq \mathcal{N}, 1 \leq f \leq \mathcal{F}\}$$

where $l$ is a domain index and $f$ is a fragment index. For notational convenience, we treat the distinguished states $\mathtt{SS}$ and $\mathtt{EE}$ as carrying sentinel indices $(l, f) = (0, 0)$, which are unused by other states as true domain and fragment indices are 1-based. Thus expressions written in terms of a generic state $\mathtt{UX}_{lf}$ are understood to include the cases $\mathtt{SS}$ and $\mathtt{EE}$ by setting $(l, f) = (0, 0)$, except where formulae explicitly require $1 \leq l \leq \mathcal{N}$ or $1 \leq f \leq \mathcal{F}$.

Loosely speaking, and noting the above caveat, every transition $\mathtt{UX}_{lf} \to \mathtt{VY}_{mg}$ involves a potential domain exit transition from $\mathtt{X} \to \mathtt{E}$ in the nested model (weight $p_{\text{out}}$), an inter-domain transition $\mathtt{U} \to \mathtt{V}$ in the top-level model that factors in paths through empty domains (weight $\mathcal{T}_{\mathtt{UV}}$), and a potential domain re-entry transition from $\mathtt{S} \to \mathtt{Y}$ in the nested model (weight $p_{\text{in}}$), which must include a factor of $(1 - z_T)^{-1}$ (for top-level Match states) or $(1 - z_0)^{-1}$ (for top-level Insert/Delete states) to account for domain entry being conditional on the domain being nonempty (these factors will precisely cancel out the factors in the $\mathtt{M}$, $\mathtt{I}$, and $\mathtt{D}$ columns of $\upsilon$, which have been included here solely to preserve row-normalization of $\upsilon$ and $\chi$). If the domain type and top-level state are the same for source and destination state ($\mathtt{U} = \mathtt{V}$ and $l = m$), then an additional intra-domain transition which extends the current domain, starting a new fragment, is folded in (weight $p_{\text{SameDom}}$). If the domain is the same ($l = m$), then an additional intra-fragment fragment-type transition from $f$ to $g$ is folded in (weight $p_{\text{SameFrag}}$), governed by the $\mathcal{F} \times \mathcal{F}$ transition matrix $r^{(l)}_{fg}$ of the Markov chain within the current fragment.

The transition matrix $\chi(\theta, T)$ for this collapsed Pair HMM has entries

| Source $i$ $(\mathtt{UX}_{lf})$ | Destination $j$ $(\mathtt{VY}_{mg})$ | $\chi_{ij} = p_{\text{out}} \times \mathcal{T}_{\mathtt{UV}} \times p_{\text{in}} + \delta(\mathtt{U}=\mathtt{V})\delta(l=m)(p_{\text{SameDom}} + \delta(\mathtt{X}=\mathtt{Y})p_{\text{SameFrag}})$ $p_{\text{out}}(\mathtt{U},\mathtt{X},l,f)$ | $\mathcal{T}_{\mathtt{UV}}$ | $p_{\text{in}}(\mathtt{V},\mathtt{Y},m,g)$ | $p_{\text{SameDom}}(\mathtt{X},\mathtt{Y},l,f,g)$ | $p_{\text{SameFrag}}(l,f,g)$ |
|---|---|---|---|---|---|---|
| $\mathtt{SS}$ | $\mathtt{MY}_{mg}$ | $1$ | $\mathcal{T}_{\mathtt{SM}}$ | $(1-z_T)^{-1} v_m \tau^{(m)}_{\mathtt{SY}} w_{mg}$ | $0$ | $0$ |
| | $\mathtt{II}_{mg}$ | $1$ | $\mathcal{T}_{\mathtt{SI}}$ | $(1-z_0)^{-1} v_m \kappa_m w_{mg}$ | $0$ | $0$ |
| | $\mathtt{DD}_{mg}$ | $1$ | $\mathcal{T}_{\mathtt{SD}}$ | $(1-z_0)^{-1} v_m \kappa_m w_{mg}$ | $0$ | $0$ |
| | $\mathtt{EE}$ | $1$ | $\mathcal{T}_{\mathtt{SE}}$ | $1$ | $0$ | $0$ |
| $\mathtt{MX}_{lf}$ | $\mathtt{MY}_{mg}$ | $\rho^{(l)}_f \tau^{(l)}_{\mathtt{XE}}$ | $\mathcal{T}_{\mathtt{MM}}$ | $(1-z_T)^{-1} v_m \tau^{(m)}_{\mathtt{SY}} w_{mg}$ | $\rho^{(l)}_f \tau^{(l)}_{\mathtt{XY}} w_{lg}$ | $r^{(l)}_{fg}$ |
| | $\mathtt{II}_{mg}$ | $\rho^{(l)}_f \tau^{(l)}_{\mathtt{XE}}$ | $\mathcal{T}_{\mathtt{MI}}$ | $(1-z_0)^{-1} v_m \kappa_m w_{mg}$ | $0$ | $0$ |
| | $\mathtt{DD}_{mg}$ | $\rho^{(l)}_f \tau^{(l)}_{\mathtt{XE}}$ | $\mathcal{T}_{\mathtt{MD}}$ | $(1-z_0)^{-1} v_m \kappa_m w_{mg}$ | $0$ | $0$ |
| | $\mathtt{EE}$ | $\rho^{(l)}_f \tau^{(l)}_{\mathtt{XE}}$ | $\mathcal{T}_{\mathtt{ME}}$ | $1$ | $0$ | $0$ |
| $\mathtt{II}_{lf}$ | $\mathtt{MY}_{mg}$ | $\rho^{(l)}_f \varphi_l$ | $\mathcal{T}_{\mathtt{IM}}$ | $(1-z_T)^{-1} v_m \tau^{(m)}_{\mathtt{SY}} w_{mg}$ | $0$ | $0$ |
| | $\mathtt{II}_{mg}$ | $\rho^{(l)}_f \varphi_l$ | $\mathcal{T}_{\mathtt{II}}$ | $(1-z_0)^{-1} v_m \kappa_m w_{mg}$ | $\rho^{(l)}_f \kappa_l w_{lg}$ | $r^{(l)}_{fg}$ |
| | $\mathtt{DD}_{mg}$ | $\rho^{(l)}_f \varphi_l$ | $\mathcal{T}_{\mathtt{ID}}$ | $(1-z_0)^{-1} v_m \kappa_m w_{mg}$ | $0$ | $0$ |
| | $\mathtt{EE}$ | $\rho^{(l)}_f \varphi_l$ | $\mathcal{T}_{\mathtt{IE}}$ | $1$ | $0$ | $0$ |
| $\mathtt{DD}_{lf}$ | $\mathtt{MY}_{mg}$ | $\rho^{(l)}_f \varphi_l$ | $\mathcal{T}_{\mathtt{DM}}$ | $(1-z_T)^{-1} v_m \tau^{(m)}_{\mathtt{SY}} w_{mg}$ | $0$ | $0$ |
| | $\mathtt{II}_{mg}$ | $\rho^{(l)}_f \varphi_l$ | $\mathcal{T}_{\mathtt{DI}}$ | $(1-z_0)^{-1} v_m \kappa_m w_{mg}$ | $0$ | $0$ |
| | $\mathtt{DD}_{mg}$ | $\rho^{(l)}_f \varphi_l$ | $\mathcal{T}_{\mathtt{DD}}$ | $(1-z_0)^{-1} v_m \kappa_m w_{mg}$ | $\rho^{(l)}_f \kappa_l w_{lg}$ | $r^{(l)}_{fg}$ |
| | $\mathtt{EE}$ | $\rho^{(l)}_f \varphi_l$ | $\mathcal{T}_{\mathtt{DE}}$ | $1$ | $0$ | $0$ |

, (C.3)

where $\tau^{(n)} \equiv \tau(\lambda_n, \mu_n, T)$, $\rho^{(n)}_f \equiv 1 - \sum_g r^{(n)}_{fg}$ is the fragment termination probability for fragment state $f$ in domain $n$, and (as before) $\varphi_l \equiv 1 - \kappa_l$.

The substitution parameters $(Q^{(c)}, \pi^{(c)})$ do not appear in the transition matrix, but in the emission probabilities of the various states. The probability of emitting token $(a, b)$ from state $\mathtt{MM}_{nf}$ is

$$P(a, b \mid \mathtt{MM}_{nf}, T) = \sum_{c=1}^{C} u_{nfc}\, \pi^{(c)}_a \exp(R^{(c)} T)_{a,b} \tag{C.4}$$

where $R^{(c)} = Q^{(c)} \cdot \text{diag}(\pi^{(c)})$ is the rate matrix for site class $c$, and $u_{nfc}$ is the probability that fragment state $f$ in domain $n$ generates site class $c$. The probability of emitting ancestral token $a$ from states $\{\mathtt{MD}_{nf}, \mathtt{DD}_{nf}\}$, or descendant token $a$ from states $\{\mathtt{MI}_{nf}, \mathtt{II}_{nf}\}$, is $\sum_c u_{nfc}\, \pi^{(c)}_a$.

##### C.1.4 Baum-Welch Algorithm for MixDom Pair HMM

In order to map HMM transition counts back to the BDI sufficient statistics, we need to correct for the null transition elimination performed in the previous section, resolve the transition counts in the nested Pair HMM onto the separate components of the model, and then apply the formulas from earlier sections.

**E-step.** Run Forward-Backward on the collapsed $(5\mathcal{N}\mathcal{F} + 2)$-state Pair HMM (Section C.1.1). This yields expected transition counts $\hat{n}''_{ij}$ for all state pairs $i, j$ in the collapsed Pair HMM, and expected emission counts $\hat{e}^{i''}_{(a,b)}$ for each state $i$ and input-output token pair $(a, b)$. The sufficient statistics for the mixture component selectors $\{v_n, w_{nf}\}$ can be recovered directly at this stage, and the emission counts accumulated onto the appropriate $W, U, V$ statistics for the per-domain CTMCs.

Conceptually speaking, we next resolve the collapsed-HMM transition counts $\hat{n}''_{ij}$ to TKF91 Pair HMM-like transition counts $\hat{n}^{(0)}_{ij}$ for transitions in the top-level inter-domain model; TKF92 Pair HMM-like transition counts $\hat{n}^{\mathtt{M}(l)'}_{\mathtt{XY}}$ for the nested intra-domain match, insert, and delete states $\mathtt{MX}_{lf}$ of each domain-match submodel (for $\mathtt{X}, \mathtt{Y} \in \{\mathtt{M}, \mathtt{I}, \mathtt{D}\}$); and singlet-style transition counts for the within-domain fragment Markov chain $\hat{n}^{\mathtt{II}(l)'}_0$, $\hat{n}^{\mathtt{II}(l)'}_1$, $\hat{n}^{\mathtt{DD}(l)'}_0$, $\hat{n}^{\mathtt{DD}(l)'}_1$ for the domain-insert and domain-delete submodels (where the "continuation" is now governed by the $\mathcal{F} \times \mathcal{F}$ Markov chain $r^{(l)}$ rather than a scalar geometric parameter).

The intra-domain transition counts arise solely from the $p_{\mathrm{SameDom}}$-weighted transitions, and can be isolated proportionally, while the inter-domain counts must correct for the Schur complement null cycle elimination. The zero-adjustment length counts $\hat{n}^{\mathtt{II}(l)'}_{\kappa}$, $\hat{n}^{\mathtt{II}(l)'}_{\varphi}$, $\hat{n}^{\mathtt{DD}(l)'}_{\kappa}$, $\hat{n}^{\mathtt{DD}(l)'}_{\varphi}$ accounting for the number of times a domain was empty *vs* nonempty must also receive contributions from the Schur complement correction.

The intra-domain counts can then be resolved to TKF91-like counts $\hat{n}^{(l)}_{ab}$ and Intra-fragment fragment-type transition counts $\hat{n}^{(n)}_{fg}$ (the expected number of transitions from fragment-type $f$ to fragment-type $g$ within a single fragment of domain $n$), together with fragment termination counts $\hat{n}^{(n)}_{\rho,f}$. The singlet-style counts for each domain $\hat{n}^{\mathtt{II}(l)'}_0$, $\hat{n}^{\mathtt{II}(l)'}_1$, $\hat{n}^{\mathtt{DD}(l)'}_0$, $\hat{n}^{\mathtt{DD}(l)'}_1$ can be resolved via a similar procedure to $\hat{n}_\kappa$, $\hat{n}_\varphi$, which are directly link sequence extension/termination counts $L^{(l)}, M^{(l)}$, to be accumulated onto the running totals for those counts. Finally, all TKF91-like transition counts for the top-level and nested models are accumulated onto the BDI sufficient statistics $S, B, D, L, M$ as in Section A.1.8.

We now consider the null count restorations in more detail. In practice, we will use a notational shortcut (that is, nevertheless, correct and reliable, and entirely equivalent to the procedure we just outlined) to greatly simplify many of these calculations, bypassing much of this conceptual tower of piecewise count restorations. In doing this we again exploit a form of the score function identity—in this case, that the expected transition usage is the derivative of the log-likelihood with respect to the log-transition weight.

**Resolving transition counts.** Following the notation of Section C.1.1, let $(\mathcal{U}_{\Phi_1,\Phi_2})_{ij} = \upsilon_{ij}\delta(i \in \Phi_1, j \in \Phi_2)$ be the masked transition matrix so that $\mathcal{Z} = (I - \mathcal{U}_{\mathtt{ABC},\mathtt{ABC}})^{-1}$ represents sums over null paths and $\mathcal{T} = \mathcal{U}_{\mathtt{SMIDE},\mathtt{SMIDE}} + \mathcal{U}_{\mathtt{SMIDE},\mathtt{ABC}} \cdot \mathcal{Z} \cdot \mathcal{U}_{\mathtt{ABC},\mathtt{SMIDE}}$ the null-eliminated transition matrix.

Considering paths in $\upsilon$ that begin and end in $i, j \in \{\mathtt{S}, \mathtt{M}, \mathtt{I}, \mathtt{D}, \mathtt{E}\}$, whose intermediate states (if any) are in $\{\mathtt{A}, \mathtt{B}, \mathtt{C}\}$, the expected transition usages and state occupancies are

$$\begin{aligned} \mathbb{E}[n_{ab}|i \to j] &= \frac{(I + \mathcal{U}_{\mathtt{SMIDE},\mathtt{ABC}} \cdot \mathcal{Z})_{ia} \mathcal{T}_{ab} (\mathcal{Z} \cdot \mathcal{U}_{\mathtt{ABC},\mathtt{SMIDE}})_{bj}}{\mathcal{T}_{ij}} \\ &= \frac{\partial \log \mathcal{T}_{ij}}{\partial \log \upsilon_{ab}} \end{aligned}$$

The expected transition usage is the score function identity, appearing in a new form: the $n_{ab}$ are sufficient statistics for the $\upsilon$ path log-likelihood. We can use this identity, with the chain rule, for many of the counts we seek:

$$\hat{n}^{(0)}_{\texttt{UV}} = \sum_{i,j} \hat{n}''_{ij} \sum_{k,l} \frac{\partial \log \chi_{ij}}{\partial \log \mathcal{T}_{kl}} \sum_{a,b} \frac{\partial \log \mathcal{T}_{kl}}{\partial \log \upsilon_{ab}} \frac{\partial \log \upsilon_{ab}}{\partial \log \tau^{(0)}_{\texttt{UV}}} \tag{C.5}$$

$$\hat{n}^{(l)}_{\texttt{UV}} = \sum_{i,j} \hat{n}''_{ij} \frac{\partial \log \chi_{ij}}{\partial \log \tau^{(l)}_{\texttt{UV}}} \tag{C.6}$$

$$\hat{n}_{\vartheta} = \sum_{i,j} \hat{n}''_{ij} \left( \frac{\partial \log \chi_{ij}}{\partial \log \vartheta} + \sum_{k,l} \frac{\partial \log \chi_{ij}}{\partial \log \mathcal{T}_{kl}} \sum_{a,b} \frac{\partial \log \mathcal{T}_{kl}}{\partial \log \upsilon_{ab}} \frac{\partial \log \upsilon_{ab}}{\partial \log \vartheta} \right) \tag{C.7}$$

for $\vartheta \in \{v_n, \kappa_n, \varphi_n, w_{nf}, r^{(n)}_{fg}, \rho^{(n)}_f\}$. A few notes:

1. In these expressions we have written $\tau^{(0)}_{\texttt{UV}}$ for the inter-domain TKF91 transition probabilities that just appear as $\tau_{\texttt{UV}}$ in Equation (C.1).

2. The term $\frac{\partial \log \mathcal{T}_{ij}}{\partial \log \upsilon_{ab}}$ is used to highlight use of the chain rule, but the actual calculation of this term can be done via the matrix formula for $\mathbb{E}[n_{ab}|i \to j]$ above.

3. We must be careful to treat $(\kappa_n, \varphi_n)$ as independent free parameters of $\chi_{ij}$ for the purpose of the partial derivatives in Equation (C.7), and similarly for $(r^{(n)}_{fg}, \rho^{(n)}_f)$; even though they are deterministically related by $\kappa_n + \varphi_n = 1$ and $\rho^{(n)}_f = 1 - \sum_g r^{(n)}_{fg}$, we should not differentiate through those constraints when calculating the derivatives of $\upsilon_{ab}$ with respect to $\kappa_n$, $\varphi_n$, $r^{(n)}_{fg}$, and $\rho^{(n)}_f$.

4. We are also treating $\upsilon_{ab}$ and $\vartheta$ as independent free parameters of $\chi_{ij}$ for the purpose of Equation (C.7), which is why we expand the derivative into two terms: one term involving $\frac{\partial \log \chi_{ij}}{\partial \log \vartheta}$ which captures the direct dependence of $\chi_{ij}$ on $\vartheta$ (e.g. via $v_m$ or $w_{mg}$ in the transition formulas above), and another term which captures the indirect dependence of $\chi_{ij}$ on $\vartheta$ via $\upsilon_{ab}$.

5. Similar points apply to the roles of $\tau^{(l)}_{\texttt{UV}}$ in Equation (C.6), and of $\tau^{(0)}_{\texttt{UV}}$ in Equation (C.5): these must be treated as free parameters and not differentiated through.

6. In contrast, however, we must differentiate through the $z_0$ and $z_T$ terms in $\upsilon$ when calculating the derivatives of $\upsilon_{ab}$ with respect to $v_n$, $\varphi_n$, and $\tau^{(n)}_{\texttt{SE}}$.

7. Similarly, we must differentiate through $p_{\text{out}}$, $p_{\text{in}}$, $p_{\text{SameDom}}$, and $p_{\text{SameFrag}}$ when calculating the derivatives of $\mathcal{T}_{ij}$. These terms do not represent free parameters.

Finally, we use these counts to accumulate onto the BDI sufficient statistics for the top-level inter-domain TKF91 model and nested intra-domain TKF92 models as described in Section A.1.8. The fragment correction for the intra-domain counts has effectively already been performed by the way we resolved the intra-domain counts from the collapsed Pair HMM counts, which account for the probability of fragment continuation *vs* new fragments via the $p_{\text{SameDom}}$-weighted terms which were differentiated through by the score function identity. We also need to accumulate the expected counts for the mixture component selectors $\{v_n, w_{nf}\}$, which are $\hat{n}_{\vartheta}$ for $\vartheta \in \{v_n, w_{nf}\}$ as calculated above.

**Accumulating sufficient statistics.** Initialize all sufficient statistics to zero. Then, for each training pair $(x, y)$ with evolutionary time $T$, accumulate as follows.

*Top-level BDI (inter-domain TKF91).* From the top-level count matrix $\hat{n}^{(0)}_{ij}$, compute BDI expectations using (A.15)–(A.17) with parameters $(\lambda_0, \mu_0, T)$, and accumulate:

$$\begin{aligned}
B^{(0)} &\mathrel{+}= \mathcal{C}^B(\hat{\mathbf{n}}^{(0)}, T) \\
D^{(0)} &\mathrel{+}= \mathcal{C}^D(\hat{\mathbf{n}}^{(0)}, T) \\
S^{(0)} &\mathrel{+}= \mathcal{C}^S(\hat{\mathbf{n}}^{(0)}, T) \\
L^{(0)} &\mathrel{+}= \hat{n}^{(0)}_{\kappa} \quad \text{(expected ancestral \& inserted domain count)} \\
M^{(0)} &\mathrel{+}= 1 \quad \text{(expected top-level terminations)} \\
T^{(0)} &\mathrel{+}= T \quad \text{(top-level BDI observation time)}
\end{aligned}$$

Note that $L^{(0)}$ counts *domains* (top-level links), not residues. It is not directly observed, but is recovered from the M and D column sums of $\hat{n}^{(0)}_{ij}$: $\hat{n}^{(0)}_{\kappa} = \sum_i (\hat{n}^{(0)}_{i\mathtt{M}} + \hat{n}^{(0)}_{i\mathtt{D}})$. The time statistic $T^{(0)}$ is weighted by $\hat{n}^{(0)}_{\varphi}$, since each independent BDI process contributes one trajectory of duration $T$.

*Per-domain BDI (intra-domain TKF92, domain $l$).* The intra-domain counts $\hat{n}^{(l)}_{ij}$ have already had the TKF92 fragment correction applied (fragment continuation *vs* new fragments was resolved by differentiating through the $p_{\text{SameFrag}}$ terms). Thus:

$$\begin{aligned}
B^{(l)} &\mathrel{+}= \mathcal{C}^B(\hat{\mathbf{n}}^{(l)}, T) \\
D^{(l)} &\mathrel{+}= \mathcal{C}^D(\hat{\mathbf{n}}^{(l)}, T) \\
S^{(l)} &\mathrel{+}= \mathcal{C}^S(\hat{\mathbf{n}}^{(l)}, T) \\
L^{(l)} &\mathrel{+}= \hat{n}^{(l)}_{\kappa} \quad \text{(expected ancestral \& inserted fragment count)} \\
M^{(l)} &\mathrel{+}= \hat{n}^{(l)}_{\varphi} \quad \text{(expected domain terminations)} \\
T^{(l)} &\mathrel{+}= T \cdot \hat{n}^{(l)}_{\varphi} \quad \text{(expected domain-level BDI time)} \\
F^{(n)}_{fg} &\mathrel{+}= \hat{n}^{(n)}_{r,fg} \quad \text{(fragment } f \to g \text{ transitions)} \\
E^{(n)}_{f} &\mathrel{+}= \hat{n}^{(n)}_{\rho,f} \quad \text{(fragment } f \text{ terminations)}
\end{aligned}$$

Here $L^{(l)}$ counts fragments (links within the domain), not residues. Each domain-$l$ entry runs an independent fragment-level BDI process for time $T$, so $T^{(l)}$ is weighted by the expected number of entries. The fragment-type transition counts $F^{(n)}_{fg}$ form an $\mathcal{F} \times \mathcal{F}$ matrix per domain, recording the expected number of intra-fragment Markov transitions between fragment-types within a fragment of domain $n$.

*Mixture selectors.*

$$\begin{aligned}
N_{v_l} &\mathrel{+}= \hat{n}_{v_l} \\
N_{w_{lf}} &\mathrel{+}= \hat{n}_{w_{lf}}
\end{aligned}$$

*CTMC substitution (per site class).* For each match emission at state $\mathtt{MM}_{nf}$ with observed pair $(a, b)$, the posterior probability of site class $c$ given the emission is $\gamma_c \propto u_{nfc}\, \pi^{(c)}_a \exp(R^{(c)}T)_{a,b}$. Using this posterior weight, accumulate endpoint-conditioned CTMC expectations (A.2)–(A.3) on the $|\mathcal{A}|$-state chain with rate matrix $R^{(c)}$:

$$W_a^{(c)} \mathrel{+}= \text{(dwell in state } a\text{, weighted by } \gamma_c \text{ and HMM posterior)}$$
$$U_{a,a'}^{(c)} \mathrel{+}= \text{(transition } a \to a'\text{, weighted by } \gamma_c\text{)}$$
$$V_a^{(c)} \mathrel{+}= \text{(composition count for } a \text{ at match-position ancestors, insertions, and deletions, weighted by } \gamma_c\text{)}$$

*Site class assignment counts.* For each emission at state $\texttt{MM}_{nf}$ (or the corresponding insert/delete states), accumulate the posterior site class assignment:

$$N_{u_{nfc}} \mathrel{+}= \gamma_c$$

*Remark* C.2 (Genealogical correction terms in nested models). As noted in Section A.1.8, the CTMC sufficient statistics $W, U, V$ omit genealogical correction terms from transient and partially-observed lineages. An analogous omission applies at each nesting level of the MixDom model: transient domain insertions and deletions (domains that are born and die between times 0 and $T$ without being directly observed) contribute to the top-level BDI sufficient statistics $B$, $D$, $S$ via the null count restoration, but their internal fragment-level processes (intra-fragment fragment-type Markov chain and BDI) are not modeled. Similarly, domains that are inserted after time 0 or deleted before time $T$ do not accumulate fragment-level statistics for the period of their non-existence. This is consistent with the principle that the M-step optimizes only the complete-data log-likelihood for structures whose existence is certified by the HMM state path.

**M-step.** All M-step updates use MAP estimates whose priors contribute additive pseudocounts to the sufficient statistics, so the maximizer formulas are the same as the MLE formulas applied to prior-augmented statistics. For the multinomial parameter groups (mixture weights, fragment-type transitions, site-class distributions) the priors below are the standard Dirichlet conjugates; for the BDI rates and the reversible GTR submodel the priors below are non-conjugate regularizers and the true conjugate priors are different (see "Comments on conjugacy" below for what the proper conjugate priors look like and why we use the simpler ones here).

*Priors.*

- Gamma$(\alpha_\lambda, \beta_{\lambda\mu})$ on $\lambda$ and Gamma$(\alpha_\mu, \beta_{\lambda\mu})$ on $\mu$, sharing the rate parameter $\beta_{\lambda\mu}$. Augmented statistics: $B \to B + \alpha_\lambda - 1$, $D \to D + \alpha_\mu - 1$, $S \to S + \beta_{\lambda\mu}$.
- Gamma$(\alpha_Q, \beta_Q)$ on each $Q_{ij}$ (shared $\beta_Q$ per row): $U_{ij} \to U_{ij} + \alpha_Q - 1$, $W_i \to W_i + \beta_Q$.
- Dirichlet$(\alpha_\pi)$ on $\pi$: $V_i \to V_i + \alpha_\pi - 1$.
- Dirichlet$(\alpha_v)$ on domain weights: $N_{v_n} \to N_{v_n} + \alpha_v - 1$.
- Dirichlet$(\alpha_w)$ on fragment weights: $N_{w_{nf}} \to N_{w_{nf}} + \alpha_w - 1$.
- Dirichlet$(\alpha_r)$ on each row of the fragment transition matrix (including the termination probability): $F_{fg}^{(n)} \to F_{fg}^{(n)} + \alpha_r - 1$, $E_f^{(n)} \to E_f^{(n)} + \alpha_r - 1$.
- Dirichlet$(\alpha_u)$ on site class distributions: $N_{u_{nfc}} \to N_{u_{nfc}} + \alpha_u - 1$.

**Comments on conjugacy.** The Dirichlet priors on $v$, $w$, $u$, and the fragment-type transition matrix are conjugate to the corresponding multinomial likelihoods in the standard way. The two remaining cases warrant comment:

*Reversible CTMC.* Treated as an irreversible CTMC (independent off-diagonal rates $R_{ij}$), the complete-data likelihood (A.1) is a regular exponential family and a product of independent Gammas on $R_{ij}$ together with a Dirichlet on the initial distribution is conjugate. For the *reversible* parameterization $R_{ij} = Q_{ij}\pi_j$ with symmetric exchangeabilities, the conjugate prior is *not* a product of independent Gammas on $Q_{ij}$ and a Dirichlet on $\pi$: detailed balance couples the two factors, and the conjugate prior is the cycle-corrected edge-flow density of Diaconis and Rolles (12), which arises as the de Finetti mixing measure for an edge-reinforced random walk on the state graph (10, 43). Reparameterizing in terms of an undirected edge weight $x_e = \pi_i Q_{ij} \pi_j$ on each edge $e = \{i, j\}$ (with loops $x_{ii} = \pi_i^2 R_{ii}^{\text{loop}}$ in the Grassmann-uniformized variant) and normalising so that $\sum_e x_e = 1$, the prior takes the form

$$\phi_{v_0,a}(x) \ \propto \ \Big(\prod_e x_e^{a_e - 1/2}\Big)\, x_{v_0}^{a_{v_0}/2} \prod_{v \neq v_0} x_v^{-(a_v+1)/2} \sqrt{\det A(x)},$$

where $x_v = \sum_{e \ni v} x_e$ is the total flow at vertex $v$, $a_v = \sum_{e \ni v} a_e$ is the corresponding pseudocount, and $A(x)$ is a matrix indexed by a basis of cycles in the state graph whose determinant equals $\sum_T \prod_{e \notin T} x_e^{-1}$ (sum over spanning trees $T$), by Kirchhoff's matrix-tree theorem. For the complete graph with loops (i.e. GTR), $\binom{|\mathcal{A}|-1}{2}$ independent cycles contribute and the joint prior cannot be factored across edges. The Diaconis–Rolles prior conditions on the initial state $v_0$; including a stationary observation of $X(0) = v_0$ multiplies the prior by $\pi(v_0) \propto \sqrt{x_{v_0}}$ in the edge-flow chart, which shifts the $v_0$-exponent and stays within the same family with adjusted hyperparameters. The independent Gamma $\times$ Dirichlet priors used here are non-conjugate regularizers in the reversible parameterization and are perfectly valid for MAP, but the closed-form posterior is in the Diaconis–Rolles family rather than back in Gamma $\times$ Dirichlet.

*TKF91 BDI rates.* The joint complete-data log-likelihood $\ell_1(\lambda, \mu)$ from (A.18), which includes the prior probability of the ancestral sequence length, is a *curved* exponential family in $(\lambda, \mu)$ because the $\log(\mu - \lambda)$ term couples the natural parameters. A product of independent Gammas on $\lambda$ and $\mu$ is therefore not conjugate; it is conjugate to the complete-data likelihood of the underlying linear birth–death process *conditioned on* the initial sequence length, but not to the joint likelihood that includes a stationary $L$-prior. The proper conjugate prior, derived in the queueing-theory literature by Armero and Bayarri (3) and applied to the linear-growth BDI by Conti (9), is most naturally written in terms of $\kappa = \lambda/\mu \in (0, 1)$ and $\mu$:

$$\phi(\kappa, \mu) \ \propto \ \kappa^{a-1}(1-\kappa)^{b-1}\, \mu^{c-1} \exp\big(-\mu\,(\tau_1\,\kappa + \tau_2)\big),$$

with five hyperparameters $(a, b, c, \tau_1, \tau_2)$ updated by $a \to a + B + L$, $b \to b + M$, $c \to c + B + D$, $\tau_1 \to \tau_1 + S + T$, $\tau_2 \to \tau_2 + S$. Marginalising $\mu$ yields a Gauss-hypergeometric distribution on $\kappa$ (in the Johnson–Kotz–Balakrishnan family); the normaliser is a ${}_2F_1$ value and the posterior is tractable by one-dimensional quadrature or Gibbs sampling. Equivalently, the $M \log(\mu - \lambda) + L \log \lambda$ contribution from incorporating the stationary initial-length prior amounts to an extra $\text{Beta}(L+1, M+1)$-like factor in $\kappa$ on top of the dynamics-only Gamma evidence. As above, our independent Gamma priors on $\lambda$ and $\mu$ are non-conjugate regularizers in this parameterisation. In the long-time stationary regime where $B \approx D$, the $M \log(\mu - \lambda)$ and $L \log \lambda - (L + M) \log \mu$ stationary contributions are doing the work of identifying $\kappa$ separately from the overall rate scale; ignoring them entirely (i.e. Gamma EM with no $L, M$ counts) silently underuses the data when only one long observation is available.

We use simple Gamma$\times$Dirichlet pseudocounts throughout because they are easy to set, behave well as MAP regularizers, and keep the M-step closed-form (the augmented sufficient statistics still have a unique maximiser via the same quadratic in $\kappa$ and the same pooled GTR formula). A fully Bayesian treatment would substitute the Diaconis–Rolles and Armero–Bayarri/Conti priors above; we leave that to future work.

*Indel rates.* For the top-level rates $(\lambda_0, \mu_0)$ and each per-domain rate pair $(\lambda_n, \mu_n)$, solve the quadratic (A.22) with augmented $(B, D, L, M, S, T)$ and extract $\kappa, \mu, \lambda$ via (A.23)–(A.25).

*Fragment transition matrix.* Row-normalize the augmented fragment-type transition counts per domain: $r^{(n)}_{fg} \leftarrow F^{(n)}_{fg} / (E^{(n)}_f + \sum_{g'} F^{(n)}_{fg'})$ (with augmented $F$, $E$).

*Mixture weights.* Normalize augmented counts: $v_n \propto N_{v_n}$, $w_{nf} \propto N_{w_{nf}}$.

*Site class distributions.* $u_{nfc} \propto N_{u_{nfc}}$.

*CTMC parameters.* The sufficient statistics are projected onto parameter groups.

*Per-class equilibrium:* $\pi^{(c)}_a \propto V^{(c)}_a + \alpha_\pi - 1$. (This is the standard empirical-frequency estimator; the exact EM M-step couples $\pi$ with $Q$ through dwell-time statistics, but the approximation is standard practice for GTR models.)

*Per-class exchangeability:* The rate is $Q^{(c)}_{a,a'} \cdot \pi^{(c)}_{a'}$. The exchangeability is estimated from Holmes–Rubin transition and dwell counts:

$$Q^{(c)}_{a,a'} = \frac{U^{(c)}_{a,a'} + U^{(c)}_{a',a}}{W^{(c)}_a \cdot \pi^{(c)}_{a'} + W^{(c)}_{a'} \cdot \pi^{(c)}_a}$$

##### C.1.5 WFSTs for MixDom

As with TKF92 (Section A.2.2), constructing a WFST for MixDom is complicated by latent information—in this case, the domain type and fragment type associated with each position. We here outline two approaches to this issue.

The first approach is to preserve the latent information by promoting it to the transducer's input/output alphabet: each character is decorated with its domain and fragment labels, yielding a *Labeled-MixDom WFST* whose state space is comparable to the Pair HMM but whose alphabet is enlarged. This approach is exact but produces larger machines.

The second approach integrates out the latent variables and approximates the result using compact *order-1* machines whose transitions depend only on the most recently emitted characters. These are smaller and more efficient for tree-based inference, at the cost of approximating the full latent structure via local context.

#### C.2 Selected Inference Algorithms for MixDom

##### C.2.1 Fast Statistical Alignment (FSA)

Given a set of sequences and a phylogenetic tree with branch-specific pair HMMs (TKF92, MixDom, or distilled order-1 transducers), we construct a multiple sequence alignment using the *sequence annealing* approach of (6), to which we refer the reader for a full description of the algorithm.

Briefly, the method proceeds as follows. For each pair of sequences $(x, y)$ in a selected subset (either all $\binom{N}{2}$ pairs or an $O(N \log N)$ Erdős–Rényi sample), we compute pairwise residue alignment posteriors $P(x_i \sim y_j)$ by running the Forward-Backward algorithm on the pair HMM at an optimized evolutionary time $\hat{\tau}$. The time $\hat{\tau}$ is found by Newton–Raphson optimization of the expected log-likelihood (the "NR step"):

$$\hat{\tau} = \underset{\tau}{\operatorname{argmax}} \; \mathbb{E}_{P(\pi|x,y,\tau_0)} \big[ \log P(x, y, \pi \mid \tau) \big] \tag{C.8}$$

where $\pi$ ranges over alignment paths and $\tau_0$ is an initial estimate. This expectation is computed from Forward-Backward expected counts at $\tau_0$, and typically converges in 3–5 Newton steps. (This time-maximization differs slightly from the approach of (6) which attempts to optimize all model parameters via unregularized EM for every pair, and consequently must terminate the EM recursion early to avoid instability.)

The pairwise posteriors are then assembled into a multiple alignment by the greedy sequence annealing procedure of (6), which iteratively merges alignment columns to maximize a sum-of-pairs posterior objective.

##### C.2.2 Beam Search Ancestral Sequence Reconstruction (BeamASR)

We now describe an alternative progressive reconstruction method that finds the maximum-likelihood ancestral sequence at each internal node by beam search, without materializing the full composite automaton.

At each internal node $v$ with children $l, r$ and observed descendant sequences $c_l, c_r$, we seek

$$\hat{a}_v = \underset{a}{\operatorname{argmax}}\big[\log P(a, c_l \mid B_l) + \log P(a, c_r \mid B_r) - \log P(a \mid R)\big] \tag{C.9}$$

where $P(a, c_k \mid B_k)$ is the pair HMM forward probability on branch $k$ and $P(a \mid R)$ is the singlet probability under the root generator, subtracted to avoid double-counting the prior on $a$.

**Incremental forward profiles** Since the branches are conditionally independent given the ancestor, we can evaluate (C.9) by maintaining *incremental forward profiles*: for each branch $k$, a 1D forward table $F_k[i, q]$ giving the log-probability that descendant positions $1, \ldots, i$ have been emitted and the branch machine is in state $q$, given ancestor positions $1, \ldots, j$ processed so far.

Each ancestor character extends both profiles independently in $O(L_k)$ time per branch, where $L_k = |c_k|$.

**Beam search** The ancestor sequence $\hat{a}_v$ is built left-to-right by beam search. At each position $j$, the beam maintains $B$ candidate partial ancestors. For each candidate and each alphabet character $\sigma$:

1. Extend both branch profiles by one ancestor character $\sigma$, comprising a *match/delete phase* (the ancestor emits $\sigma$, descendant positions advance via M or D transitions) and an *insertion phase* (descendant-only insertions following the ancestor emission).
2. Update the singlet forward score for $\sigma$.
3. Score the extension: $\Delta(j, \sigma) = \Delta F_l + \Delta F_r - \Delta_{\text{singlet}}$.

The top $B$ extensions (by cumulative score) are retained. Total cost per node is $O(K \cdot B \cdot A \cdot (L_l + L_r))$ where $K = |\hat{a}_v|$ and $A$ is the alphabet size.

**Insertion phase via associative scan** The insertion recurrence within each branch profile has the form

$$x_{i+1} = \operatorname{logsumexp}\big(A_{II}\, x_i,\ b_i\big) + e_i \tag{C.10}$$

where $A_{II}$ is the I-to-I log-transition submatrix, $b_i$ collects transitions into insertion states from M and D, and $e_i$ is the emission score. This is a log-semiring affine recurrence, parallelizable via an associative scan with operator

$$(A_1, b_1) \oplus (A_2, b_2) = (A_2 \otimes A_1,\ \operatorname{logsumexp}(A_2\, b_1,\ b_2))$$

where $\otimes$ denotes log-semiring matrix multiplication. This reduces the insertion phase from $O(L)$ sequential depth to $O(\log L)$.

**Supported model types** The beam search interface is generic over the pair HMM used on each branch:

1. **TKF92** — order-0, 5-state pair HMM (the standard model).

2. **MixDom** — the full latent-state pair HMM (Section C.1.1) with $2+5NK$ states; latent-state correlations are marginalized in the forward pass without distillation.

**Potentials from neighboring columns** The inter-column coupling enters through the order-1 WFST transitions. For each edge $e = (u, v)$ and MSA column $c$ where $e$ has an event of type $\tau_c$, let $c^-$ denote the predecessor column (the previous column where $e$ had an event) and $c^+$ the successor column. The *potential* at column $c$ for edge $e$ receives two contributions:

**As-child term (from $c^-$).** The transition from column $c^-$ to $c$ on edge $e$ depends on the characters at both columns. Using the pairwise marginal from $q_{c^-}$:

$$\log \phi_e^{\text{child}}(a_{u,c}, a_{v,c}) = \sum_{a',b'} q_{c^-}^{(u,v)}(a', b') \ \log w_e(\tau_{c^-}, \tau_c, a', b', a_{u,c}, a_{v,c}) \tag{C.11}$$

where the sum over $(a', b')$ uses the *joint* pairwise marginal $q_{c^-}^{(u,v)}(a', b')$—not the product of independent marginals. This is the key advantage over mean-field: the within-tree parent-child correlation at the predecessor column is preserved exactly.

**As-parent term (from $c^+$).** Symmetrically, column $c$ acts as the predecessor for column $c^+$:

$$\log \phi_e^{\text{parent}}(a_{u,c}, a_{v,c}) = \sum_{a'',b''} q_{c^+}^{(u,v)}(a'', b'') \ \log w_e(\tau_c, \tau_{c^+}, a_{u,c}, a_{v,c}, a'', b'') \tag{C.12}$$

For insert transitions (only the descendant is present at $c$), the potential reduces to a per-node function $\psi_v(a_{v,c})$. For delete transitions (only the ancestor is present), it becomes $\psi_u(a_{u,c})$. For match transitions, it contributes a per-edge potential $\phi_e(a_{u,c}, a_{v,c})$. The start and end transitions contribute analogous per-node or per-edge terms.

**Felsenstein coordinate ascent** Each coordinate ascent step updates $q_c$ for a single MSA column $c$, holding all other columns fixed. We accumulate, for each edge $e$ and node $v$ in the tree at column $c$:

- Per-edge log-potentials $\log \phi_e(a_u, a_v) = \log \phi_e^{\text{child}} + \log \phi_e^{\text{parent}}$ (for match transitions where both endpoints are present).

- Per-node log-potentials $\log \psi_v(a_v)$ (from insert/delete transitions on incident edges, plus the root prior $\log \pi(a)$ at the root node).

The optimal $q_c$, given the potentials, is the Gibbs distribution on the tree at column $c$:

$$q_c(\mathbf{h}_c) \propto \prod_v \psi_v(a_v) \prod_{e=(u,v)} \phi_e(a_u, a_v) \prod_{\ell \in \text{leaves}} \delta(a_\ell = y_\ell) \tag{C.13}$$

Since this is a tree-structured MRF, the normalizing constant and all node and edge marginals can be computed *exactly* by Felsenstein peeling (postorder) and unpeeling (preorder) in $O(|E| \cdot |\mathcal{A}|^2)$ time.

**Peeling (postorder).** For each node $v$ in postorder, compute the conditional likelihood:

$$\mathrm{CL}_v(a) = \psi_v(a) \prod_{\text{children } c} \Big[\sum_{a_c} \phi_{(v,c)}(a, a_c)\, \mathrm{CL}_c(a_c)\Big] \tag{C.14}$$

with $\mathrm{CL}_\ell(a) = \delta(a = y_\ell)$ for observed leaves. The log-partition function is $\log Z_c = \log \sum_a \pi(a)\, \mathrm{CL}_{\mathrm{root}}(a)$.

**Unpeeling (preorder).** Propagate top-down to obtain the posterior marginal at each node:

$$q_c^{(v)}(a) \propto \mathrm{CL}_v(a) \cdot \mathrm{msg}_{\mathrm{parent}\to v}(a) \tag{C.15}$$

and the pairwise marginal on each edge:

$$q_c^{(u,v)}(a_u, a_v) \propto \mathrm{msg}_{\mathrm{above}\, u}(a_u) \cdot \phi_{(u,v)}(a_u, a_v) \cdot \mathrm{CL}_v(a_v) \tag{C.16}$$

where the "message from above $u$" combines the top-down message to $u$ with $u$'s conditional likelihood excluding child $v$.

**Sweep.** One iteration sweeps through all MSA columns $c = 1, \ldots, L$: for each column, recompute the potentials from the current neighbor marginals, run peeling/unpeeling, and store the updated node and edge marginals. The sweep order is left-to-right; the "as-child" potentials use the just-updated predecessor marginals, while the "as-parent" potentials use stale successor marginals from the previous iteration.

**Properties** The product-of-trees approximation enjoys the same monotonic convergence guarantee as mean-field coordinate ascent (each column update minimizes the free energy in its coordinate), with the additional guarantee that the ELBO is at least as tight as the fully-factored mean-field bound. This follows because the product-of-trees family *contains* the mean-field family as a special case (where each $q_c$ is itself fully factored).

**Computational cost.** The dominant cost per sweep is the potential computation: $O(|E| \cdot L \cdot |\mathcal{A}|^4)$ for Match→Match transitions (contracting the $(|\mathcal{A}|, |\mathcal{A}|)$ pairwise marginal against the $(|\mathcal{A}|, |\mathcal{A}|, |\mathcal{A}|, |\mathcal{A}|)$ WFST tensor). The Felsenstein passes add $O(|E| \cdot L \cdot |\mathcal{A}|^2)$, which is subdominant.

##### C.2.3 Phylogenetic Hidden Markov Model (PhyloHMM)

If the top-level indel rates in MixDom are low, and the ancestral presence/absence fully specified by the MSA, the phylogenetic likelihood calculation and ancestral reconstruction problems admit systematic approximation by a generalized Phylo-HMM, yielding $O(L^2)$-complexity versions of the Forward and Forward-Backward algorithms. This approach is described in Section C.9.

##### C.2.4 Phylogenetic composition

The order-1 HMM (Section C.4.5) and transducers (Section C.4.6) can be composed on a phylogenetic tree to yield a single composite machine whose state encodes the joint configuration of all branch machines.

Given a rooted binary tree with $n$ leaves:

- Number nodes $0, \ldots, 2n-2$ in *preorder* (root = 0).

- Place the order-1 Singlet HMM on a notional branch above the root (node 0).
- Place an order-1 Pair Transducer on each real branch.
- Each node $v$ carries a *tag* $\in \mathcal{A} \cup \{\varepsilon\}$, initially $\varepsilon$.

Each branch machine is either the root HMM or a branch transducer. All machines are converted to waiting-machine form (they already are, by construction above).

**Composition rules**

**State constraint.** A node's machine may advance (take a transition) only if all higher-numbered nodes' machines are in waiting states.

**Tagging.** When an internal node $v$'s transducer takes a transition with output symbol $b \neq \varepsilon$, node $v$ is tagged with $b$. If $v$'s tag is non-$\varepsilon$, the next move must feed $v$'s tag as input to both child branch transducers (forcing them out of their waiting states). This clears $v$'s tag (but the children's transitions may tag downstream nodes).

**Priority.** If multiple nodes are tagged simultaneously, the lowest-numbered (closest to root) tagged node is cleared first.

**Cascading.** This system of tags allows upward propagation from observed leaf emissions (MSA columns) via Felsenstein-style pruning: working from leaves (known emissions) upward, each internal node's ancestral character is a latent variable marginalized by the DP.

**Composite state space** A *configuration* of the composed machine is a tuple

$$\sigma = (q_0, q_1, \ldots, q_{2n-2},\ \tau_0, \tau_1, \ldots, \tau_{2n-2})$$

where $q_v$ is node $v$'s machine state and $\tau_v \in \mathcal{A} \cup \{\varepsilon\}$ is its tag. The start configuration has all machines in `S` and all tags $\varepsilon$. The end configuration has all machines in `E` and all tags $\varepsilon$.

**Practical caveat.** While this composition defines a valid single machine whose language is the set of MSAs weighted by the full phylogenetic likelihood, the composite state space is $O(|Q|^{2n-1} \cdot |\mathcal{A}|^{2n-1})$ where $|Q|$ is the number of states per branch machine—i.e. geometric in the number of taxa. Explicitly constructing the composed machine is therefore impractical for all but the smallest trees. The beam algorithms that follow (Sections C.2.5–C.2.6) avoid this by enumerating only the configurations reachable within a pruned beam, so that the effective state space remains manageable. The composition formalism is nonetheless useful as a *specification*: it defines the target distribution from which the beam search samples or whose expected counts the Forward-Backward algorithm estimates.

##### C.2.5 Beam Backward algorithm (BeamMSA)

Given a multiple sequence alignment (MSA) with $L$ columns and the composite machine from Section C.2.4, we compute the alignment likelihood using a *beam Backward* algorithm, working from the end configuration backward.

**Columns and the emission constraint.** Each MSA column $\ell = 1, \ldots, L$ specifies, for each leaf $v$, either a character $y_v^\ell \in \mathcal{A}$ or a gap. A configuration $\sigma$ is *compatible* with column $\ell$ if the set of leaf emissions implied by $\sigma$'s tags matches the column.

**Backward recurrence.** Let $B(\sigma)$ denote the Backward variable: the total probability of generating MSA columns $\ell, \ell+1, \ldots, L$ and reaching the end state, given that the composite machine is currently in configuration $\sigma$ just before column $\ell$.

$$B(\sigma_{\mathtt{E}}) = 1 \qquad \text{(end configuration)} \tag{C.17}$$

$$B(\sigma) = \sum_{\sigma'} T(\sigma, \sigma')\, B(\sigma') \qquad \text{(all other configurations)} \tag{C.18}$$

where $T(\sigma, \sigma')$ is the composite transition weight (product of individual machine transitions, subject to the composition rules above) and the sum is over all successor configurations $\sigma'$.

The alignment likelihood is $B(\sigma_{\mathtt{S}})$.

**Beam pruning.** Maintain a beam $\mathcal{B}_\ell$ of at most $W$ configurations per column, ranked by $B(\sigma)$. When expanding $\mathcal{B}_\ell$ from $\mathcal{B}_{\ell+1}$, discard any $\sigma$ whose $B(\sigma)$ falls below $B_{\max}/\Delta$ where $B_{\max}$ is the current maximum and $\Delta$ is the beam width ratio. If the beam collapses (no configurations remain), backtrack.

**Epsilon closures within a column.** A *column-emitting move* is any transition whose output cascades down the tree to produce a new MSA column: specifically, an insertion (a transition that outputs a character without consuming input), which then tags the node and cascades to its descendants. Between column-emitting moves, machines may make silent transitions (tag propagation, waiting-state transitions, etc.). These form an $\varepsilon$-closure that must be computed at each step. For each configuration in the beam, enumerate all reachable configurations via silent transitions (respecting the priority ordering), accumulating weights multiplicatively along each path. In practice, null cycles (if any) can either be ignored (assuming the model's null-state topology is acyclic) or handled by allowing a configurable number of extra exploratory steps in the beam search.

**Forward traceback** After the Backward pass reaches $\sigma_{\mathtt{S}}$, a stochastic Forward traceback samples a path from $\sigma_{\mathtt{S}}$ to $\sigma_{\mathtt{E}}$:

$$P(\sigma'|\sigma) = \frac{T(\sigma, \sigma')\, B(\sigma')}{B(\sigma)} \tag{C.19}$$

At each step, sample the next configuration proportional to (C.19). This yields a sampled alignment (including ancestral sequences at internal nodes).

**Beam Forward-Backward** Alternatively, after the Backward beam pass:

1. Prune dead-end configurations from each $\mathcal{B}_\ell$ (those with no predecessor in $\mathcal{B}_{\ell-1}$).
2. Run a Forward pass over the pruned beam:

$$F(\sigma_{\mathtt{S}}) = 1 \tag{C.20}$$

$$F(\sigma') = \sum_{\sigma \in \mathcal{B}} T(\sigma, \sigma')\, F(\sigma) \tag{C.21}$$

3. Posterior marginals for any feature $\phi$:

$$P(\phi|\text{MSA}) = \frac{1}{B(\sigma_{\mathtt{S}})} \sum_{\sigma \to \sigma'} F(\sigma)\, T(\sigma, \sigma')\, B(\sigma')\, [\phi(\sigma, \sigma')] \tag{C.22}$$

**Beam Viterbi** Replace $\sum$ with max in the Backward recurrence (C.18) and store argmax pointers:

$$B^V(\sigma_{\mathtt{E}}) = 1 \tag{C.23}$$

$$B^V(\sigma) = \max_{\sigma'} T(\sigma, \sigma')\, B^V(\sigma') \tag{C.24}$$

The optimal alignment is recovered by Forward traceback following the argmax pointers.

##### C.2.6 Progressive alignment via profile construction (ProgRec)

We now describe a progressive multiple sequence alignment algorithm using the order-1 machines from Sections C.4.5–C.4.6, with model parameters from (30). This follows the transducer-composition approach of (52), adapted here for Mealy machines (I/O on transitions rather than states).

The antecedents of this approach are the full multidimensional alignment algorithm of (18) which computes the Forward algorithm for TKF91 on a binary tree. This may be seen as unifying the tree-based Viterbi multiple alignment approach of (44) with the statistical phylogenetics of (16). The approach described here also maintains a partial order graph of intermediate alignments (33), which essentially is the approach used by (35).

**Recognizers and Profiles** Let the phylogenetic tree have $n$ leaves with observed sequences $\{y_v : v \in \text{leaves}\}$, nodes numbered in preorder. Let $R$ denote the order-1 Singlet HMM (root generator, Section C.4.5) and $B_v$ the order-1 Pair Transducer on the branch to node $v$ (Section C.4.6).

**Recognizers.** For each leaf $v$, the *exact-match recognizer* $\mathcal{R}(y_v)$ is a transducer with empty output alphabet that accepts only $y_v$: it has $|y_v| + 1$ states (positions $0, \ldots, |y_v|$), with a single input-consuming transition $i \to i + 1$ labeled by $y_v[i + 1]$ at each position. All states are waiting states except the start.

**Profiles.** A *profile* $E_v$ at node $v$ is a recognizer (empty output alphabet) that accepts a set of plausible ancestral sequences at $v$, weighted by their approximate posterior probability given the descendants of $v$. For leaves, $E_v = \mathcal{R}(y_v)$.

**Progressive reconstruction** Working from the leaves toward the root, for each internal node $v$ with children $l, r$:

**Step 1: Compose branch and profile.** For each child $c \in \{l, r\}$, form the composition $B_c \circ E_c$, which is a transducer mapping the sequence at $v$ to the constrained sequences at child $c$. Since $E_c$ has empty output, $B_c \circ E_c$ is a recognizer (it reads a candidate parent sequence and recognizes it with weight proportional to the probability of generating the descendant data at $c$).

**Step 2: Intersect siblings.** Form the intersection

$$H_v = (B_l \circ E_l) \cap (B_r \circ E_r)$$

This recognizer reads a candidate sequence at $v$ and scores it by the joint probability of both children's descendant data, given that parent sequence. In the Mealy-machine intersection, the composite state is $(q_l, e_l, q_r, e_r)$ where $q_c \in B_c$ and $e_c \in E_c$. Both sides must agree on the same input symbol when both are ready (waiting); when one side is not waiting, it advances silently while the other stays put.

**Step 3: Compose with root prior.** Form the generator

$$M_v = R \circ H_v$$

This has empty input and empty output: it is a weighted automaton over the empty string, whose total weight $Z = \sum_\pi w(\pi)$ over all paths $\pi$ is the marginal likelihood of the descendant data below $v$ (under the stationary prior $R$ at $v$).

**Step 4: Sample paths and construct profile.** Sample $K$ paths from $P(\pi|M_v) = w(\pi)/Z$ using a Forward pass followed by stochastic traceback (C.19). For each sampled path $\pi$, extract the $H_v$-component states visited. The profile $E_v$ is the sub-recognizer of $H_v$ containing exactly those states visited by at least $\tau \geq 2$ of the $K$ sampled paths. This bounds $|E_v| = O(KL)$ where $L = \max_v |y_v|$.

**Mealy-machine composition and intersection** For completeness, we state the composition and intersection rules for Mealy machines in waiting-machine normal form ("ready" states = waiting states with input-consuming transitions only; "unready" states = non-waiting, silent transitions only).

**Composition.** Given $T = (\Omega_X, \Omega_Y, \ldots)$ and $U = (\Omega_Y, \Omega_Z, \ldots)$ in Mealy normal form, $T \circ U$ has states $\subseteq \mathcal{S}_T \times \mathcal{S}_U$ with transition weight:

$$w''((t,u), \omega_x, \omega_z, (t',u')) = \begin{cases} \delta_{tt'}\, \delta_{\omega_x \varepsilon}\, w'(u, \varepsilon, \omega_z, u') & \text{if } u \text{ unready} \\ \delta_{uu'}\, \delta_{\omega_z \varepsilon}\, w(t, \omega_x, \varepsilon, t') + \sum_{\omega_y} w(t, \omega_x, \omega_y, t')\, w'(u, \omega_y, \omega_z, u') & \text{if } u \text{ ready} \end{cases}$$

**Intersection.** Given $T = (\Omega_X, \Omega_T, \ldots)$ and $U = (\Omega_X, \Omega_U, \ldots)$ in Mealy normal form, $T \cap U$ has states $\subseteq \mathcal{S}_T \times \mathcal{S}_U$ with output alphabet $\Omega_T \times \Omega_U$ and transition weight:

$$w''((t,u), \omega_x, (\omega_y, \omega_z), (t',u')) = \begin{cases} \delta_{tt'}\, \delta_{\omega_x \varepsilon}\, \delta_{\omega_y \varepsilon}\, w'(u, \varepsilon, \omega_z, u') & \text{if } u \text{ unready} \\ \delta_{uu'}\, \delta_{\omega_x \varepsilon}\, \delta_{\omega_z \varepsilon}\, w(t, \varepsilon, \omega_y, t') & \text{if } t \text{ unready, } u \text{ ready} \\ w(t, \omega_x, \omega_y, t')\, w'(u, \omega_x, \omega_z, u') & \text{if both ready} \end{cases}$$

**Forward recursion for $M_v$** The generator $M_v = R \circ H_v$ has states $m = (\rho, q_l, e_l, q_r, e_r)$ where $\rho \in R$, $(q_l, e_l) \in B_l \circ E_l$, and $(q_r, e_r) \in B_r \circ E_r$. The Forward variable $Z(m)$ satisfies:

$$Z(\mathtt{S}_M) = 1 \tag{C.25}$$

$$Z(m') = \sum_{m:(m,\varepsilon,\varepsilon,m') \in \mathcal{T}} w(m, \varepsilon, \varepsilon, m')\, Z(m) \tag{C.26}$$

where $\mathcal{T}$ is the transition set of $M_v$. The total likelihood is $Z(\mathtt{E}_M)$.

The fill order iterates over $e_l$ and $e_r$ in topological order (corresponding to positions in the child profiles), with an inner loop over $Q_v = R \circ (B_l \cap B_r)$ states (the "comparison kernel" Pair HMM). This has time complexity $O(|B|^2\, |E_l|\, |E_r|)$ per internal node, where $|B|$ is the branch transducer state count.

**Profile extraction** Given the Forward table $Z$, sample paths $\pi^{(1)}, \ldots, \pi^{(K)}$ from $M_v$ using the stochastic traceback (C.19). For each path $\pi^{(k)}$, let $\mathcal{H}^{(k)} = \{(q_l, e_l, q_r, e_r) : (\rho, q_l, e_l, q_r, e_r) \in \pi^{(k)}\}$ be the $H_v$-states visited.

The profile $E_v$ is the sub-automaton of $H_v$ induced by the states

$$\mathcal{S}_{E_v} = \{h \in H_v : |\{k : h \in \mathcal{H}^{(k)}\}| \geq \tau\}$$

with the same transition weights as $H_v$, restricted to $\mathcal{S}_{E_v}$. Adding appropriate wait states places $E_v$ in Mealy normal form.

**Bubble merging.** Paths through $E_v$ that traverse the same sequence of wait states but differ only in latent-state assignments can be merged by collapsing bubbles (identifying states with identical incoming and outgoing wait-state connectivity). This further compresses the profile without changing the recognized language.

**MSA extraction** A sampled path through $M_1$ (the root) determines, at each position, which $H_1$-state is visited, and therefore which states of $E_l$, $E_r$ are aligned. Recursing into the child profiles yields a full column assignment for all leaves.

Specifically, each emitting transition in the sampled path implies:

- a character at the current node (from the root generator or branch match),
- advancement of the left profile, right profile, or both,
- and therefore a column in the MSA (with gaps for non-advancing sides).

When bubble merging has been applied, the canonical path (chosen during merging) is used to resolve any ambiguity in the sub-alignment of the clade below the merged bubble.

**Viterbi Progressive Reconstruction** An alternative to the sampling-based profile construction of Step 4 above is to use Viterbi decoding at each internal node. Instead of sampling $K$ paths from $M_v$ and building a multi-path profile, the Viterbi variant computes a single maximum-likelihood path through $M_v$ and uses the resulting ancestral sequence directly as the reconstructed sequence at node $v$. This gives a deterministic progressive reconstruction that avoids the $O(KL)$ profile size but sacrifices the ability to represent uncertainty in the ancestral sequence.

Table C.1: Exploded MixDom Pair HMM state space.

| Category | States | Count |
|---|---|---|
| **Start/End** | `S`, `E` | 2 |
| **Domain-level** (top-level TKF91 states) | `MatDom`, `InsDom`, `DelDom`, `MatDomEnd`, `InsDomEnd`, `DelDomEnd` | 6 |
| **Domain type selection** (one per domain type $k$) | `MatDomType`$[k]$, `InsDomType`$[k]$, `DelDomType`$[k]$ | $3\mathcal{N}$ |
| **Fragment-level** (inner TKF states within `MatDomType`$[k]$) | `MatFrag`$[k]$, `InsFrag`$[k]$, `DelFrag`$[k]$ | $3\mathcal{N}$ |
| **Fragment-level** (single looping state within `InsDomType`$[k]$, `DelDomType`$[k]$) | `IFrag`$[k]$, `DFrag`$[k]$ | $2\mathcal{N}$ |
| **Fragment type selection** (one per fragment type $f$) | `MatFragType`$[k,f]$, `InsFragType`$[k,f]$, `DelFragType`$[k,f]$, `IFragType`$[k,f]$, `DFragType`$[k,f]$ | $5\mathcal{N}\mathcal{F}$ |
| **Emit states** (the only emitting states) | `MatEmit`$[k,f]$, `InsEmit`$[k,f]$, `DelEmit`$[k,f]$, `IEmit`$[k,f]$, `DEmit`$[k,f]$ | $5\mathcal{N}\mathcal{F}$ |
| **Fragment end** (fragment termination) | `MatFragEnd`$[k,f]$, `InsFragEnd`$[k,f]$, `DelFragEnd`$[k,f]$, `IFragEnd`$[k,f]$, `DFragEnd`$[k,f]$ | $5\mathcal{N}\mathcal{F}$ |
| **Total** | | $8+8\mathcal{N}+15\mathcal{N}\mathcal{F}$ |

#### C.3 Exploded MixDom Pair HMM

##### C.3.1 State Space

The exploded MixDom Pair HMM makes every structural decision explicit as a separate state transition. Let $\mathcal{N}$ denote the number of domain types, $\mathcal{F}$ the number of fragment types per domain. Parameters are indexed by domain type $k$ and fragment type $f$.

The states are shown in Table C.1 (emitting states marked with $\star$).

The emitting states correspond to the compound states of the collapsed model: `MatEmit`$[k,f]$ = `MM`$_{kf}$, `InsEmit`$[k,f]$ = `MI`$_{kf}$, `DelEmit`$[k,f]$ = `MD`$_{kf}$, `IEmit`$[k,f]$ = `II`$_{kf}$, `DEmit`$[k,f]$ = `DD`$_{kf}$.

##### C.3.2 Transition Weights

All transitions are between non-emitting states, or from non-emitting to emitting, or from emitting to non-emitting (Mealy machine: emissions occur on the transitions into emit states).

We use BDI parameters for two TKF91 processes:

- Top-level (domain sequence): $\alpha_0, \beta_0, \gamma_0, \kappa_0$ from $(\lambda_0, \mu_0, T)$
- Per-domain $k$ (fragment sequence): $\alpha_k, \beta_k, \gamma_k, \kappa_k$ from $(\lambda_k, \mu_k, T)$

**Top-level transitions** These implement the TKF91 Pair HMM structure with `MatDom/InsDom/DelDom/E` (for incoming connections) and `S/MatDomEnd/InsDomEnd/DelDomEnd` (for outgoing connections) playing the roles of S/M/I/D/E:

$$\texttt{S} \to \texttt{MatDom}: \quad \tau_{\texttt{SM}}(\lambda_0, \mu_0, T) \tag{C.27}$$
$$\texttt{S} \to \texttt{InsDom}: \quad \tau_{\texttt{SI}}(\lambda_0, \mu_0, T) \tag{C.28}$$
$$\texttt{S} \to \texttt{DelDom}: \quad \tau_{\texttt{SD}}(\lambda_0, \mu_0, T) \tag{C.29}$$
$$\texttt{S} \to \texttt{E}: \quad \tau_{\texttt{SE}}(\lambda_0, \mu_0, T) \tag{C.30}$$
$$\texttt{MatDomEnd} \to \texttt{MatDom}: \quad \tau_{\texttt{MM}}(\lambda_0, \mu_0, T) \tag{C.31}$$
$$\texttt{MatDomEnd} \to \texttt{InsDom}: \quad \tau_{\texttt{MI}}(\lambda_0, \mu_0, T) \tag{C.32}$$
$$\texttt{MatDomEnd} \to \texttt{DelDom}: \quad \tau_{\texttt{MD}}(\lambda_0, \mu_0, T) \tag{C.33}$$
$$\texttt{MatDomEnd} \to \texttt{E}: \quad \tau_{\texttt{ME}}(\lambda_0, \mu_0, T) \tag{C.34}$$
$$\texttt{InsDomEnd} \to \texttt{MatDom}: \quad \tau_{\texttt{IM}}(\lambda_0, \mu_0, T) \tag{C.35}$$
$$\texttt{InsDomEnd} \to \texttt{InsDom}: \quad \tau_{\texttt{II}}(\lambda_0, \mu_0, T) \tag{C.36}$$
$$\texttt{InsDomEnd} \to \texttt{DelDom}: \quad \tau_{\texttt{ID}}(\lambda_0, \mu_0, T) \tag{C.37}$$
$$\texttt{InsDomEnd} \to \texttt{E}: \quad \tau_{\texttt{IE}}(\lambda_0, \mu_0, T) \tag{C.38}$$
$$\texttt{DelDomEnd} \to \texttt{MatDom}: \quad \tau_{\texttt{DM}}(\lambda_0, \mu_0, T) \tag{C.39}$$
$$\texttt{DelDomEnd} \to \texttt{InsDom}: \quad \tau_{\texttt{DI}}(\lambda_0, \mu_0, T) \tag{C.40}$$
$$\texttt{DelDomEnd} \to \texttt{DelDom}: \quad \tau_{\texttt{DD}}(\lambda_0, \mu_0, T) \tag{C.41}$$
$$\texttt{DelDomEnd} \to \texttt{E}: \quad \tau_{\texttt{DE}}(\lambda_0, \mu_0, T) \tag{C.42}$$

**Domain type selection**

$$\texttt{MatDom} \to \texttt{MatDomType}[k]: \quad v_k \quad \text{(domain weight)} \tag{C.43}$$
$$\texttt{InsDom} \to \texttt{InsDomType}[k]: \quad v_k \tag{C.44}$$
$$\texttt{DelDom} \to \texttt{DelDomType}[k]: \quad v_k \tag{C.45}$$

**Domain-to-fragment entry (M-type domains)** Within MatDomType[$k$], the fragment-level TKF91 begins. Again, this follows the TKF91 Pair HMM structure, now with MatFrag[$k$]/InsFrag[$k$]/DelFrag[$k$]Ma (for incoming connections) and MatDomType[$k$]/MatFragEnd[$k$, /]InsFragEnd[$k$, /]DelFragEnd[$k$,] states (for outgoing connections) playing the roles of $\texttt{S/M/I/D/E}$:

$$\texttt{MatDomType}[k] \to \texttt{MatFrag}[k]: \quad \tau_{\texttt{SM}}(\lambda_k, \mu_k, T) \tag{C.46}$$
$$\texttt{MatDomType}[k] \to \texttt{InsFrag}[k]: \quad \tau_{\texttt{SI}}(\lambda_k, \mu_k, T) \tag{C.47}$$
$$\texttt{MatDomType}[k] \to \texttt{DelFrag}[k]: \quad \tau_{\texttt{SD}}(\lambda_k, \mu_k, T) \tag{C.48}$$
$$\texttt{MatDomType}[k] \to \texttt{MatDomEnd}: \quad \tau_{\texttt{SE}}(\lambda_k, \mu_k, T) \tag{C.49}$$
$$\texttt{MatFragEnd}[k,] \to \texttt{MatFrag}[k]: \quad \tau_{\texttt{MM}}(\lambda_k, \mu_k, T) \tag{C.50}$$
$$\texttt{MatFragEnd}[k,] \to \texttt{InsFrag}[k]: \quad \tau_{\texttt{MI}}(\lambda_k, \mu_k, T) \tag{C.51}$$
$$\texttt{MatFragEnd}[k,] \to \texttt{DelFrag}[k]: \quad \tau_{\texttt{MD}}(\lambda_k, \mu_k, T) \tag{C.52}$$
$$\texttt{MatFragEnd}[k,] \to \texttt{MatDomEnd}: \quad \tau_{\texttt{ME}}(\lambda_k, \mu_k, T) \tag{C.53}$$
$$\texttt{InsFragEnd}[k,] \to \texttt{MatFrag}[k]: \quad \tau_{\texttt{IM}}(\lambda_k, \mu_k, T) \tag{C.54}$$
$$\texttt{InsFragEnd}[k,] \to \texttt{InsFrag}[k]: \quad \tau_{\texttt{II}}(\lambda_k, \mu_k, T) \tag{C.55}$$
$$\texttt{InsFragEnd}[k,] \to \texttt{DelFrag}[k]: \quad \tau_{\texttt{ID}}(\lambda_k, \mu_k, T) \tag{C.56}$$
$$\texttt{InsFragEnd}[k,] \to \texttt{MatDomEnd}: \quad \tau_{\texttt{IE}}(\lambda_k, \mu_k, T) \tag{C.57}$$
$$\texttt{DelFragEnd}[k,] \to \texttt{MatFrag}[k]: \quad \tau_{\texttt{DM}}(\lambda_k, \mu_k, T) \tag{C.58}$$
$$\texttt{DelFragEnd}[k,] \to \texttt{InsFrag}[k]: \quad \tau_{\texttt{DI}}(\lambda_k, \mu_k, T) \tag{C.59}$$
$$\texttt{DelFragEnd}[k,] \to \texttt{DelFrag}[k]: \quad \tau_{\texttt{DD}}(\lambda_k, \mu_k, T) \tag{C.60}$$
$$\texttt{DelFragEnd}[k,] \to \texttt{MatDomEnd}: \quad \tau_{\texttt{DE}}(\lambda_k, \mu_k, T) \tag{C.61}$$

The $\texttt{MatDomType}[k] \to \texttt{MatDomEnd}$ transition is the "phantom" null path: the domain is entered but the inner model immediately terminates with no fragments emitted. This leads to null cycles that must be eliminated by Schur complement.

**Domain-to-fragment entry (I/D-type domains)** $\texttt{InsDomType}[k]$ and $\texttt{DelDomType}[k]$ have a single looping fragment state:

$$\texttt{InsDomType}[k] \to \texttt{IFrag}[k]: \quad \kappa_k \tag{C.62}$$
$$\texttt{InsDomType}[k] \to \texttt{InsDomEnd}: \quad 1 - \kappa_k \tag{C.63}$$
$$\texttt{DelDomType}[k] \to \texttt{DFrag}[k]: \quad \kappa_k \tag{C.64}$$
$$\texttt{DelDomType}[k] \to \texttt{DelDomEnd}: \quad 1 - \kappa_k \tag{C.65}$$

Again, the $\to$ $\texttt{InsDomEnd/DelDomEnd}$ transitions are null (empty domain).

**Fragment type selection**

$$\texttt{MatFrag}[k] \to \texttt{MatFragType}[k, f]: \quad w_{kf} \quad \text{(fragment weight)} \tag{C.66}$$
$$\texttt{InsFrag}[k] \to \texttt{InsFragType}[k, f]: \quad w_{kf} \tag{C.67}$$
$$\texttt{DelFrag}[k] \to \texttt{DelFragType}[k, f]: \quad w_{kf} \tag{C.68}$$
$$\texttt{IFrag}[k] \to \texttt{IFragType}[k, f]: \quad w_{kf} \tag{C.69}$$
$$\texttt{DFrag}[k] \to \texttt{DFragType}[k, f]: \quad w_{kf} \tag{C.70}$$

**Fragment emission** These are the only transitions with emissions (the exploded HMM is represented as a Mealy machine, so emissions occur on transitions; the collapsed HMM treats emissions as state-based). The emission probability is summed over site classes $c \in \{1, \dots, C\}$, weighted by the per-fragment class distribution $u_{kfc}$:

$$\texttt{MatFragType}[k, f] \xrightarrow{(a,b)} \texttt{MatEmit}[k, f] : \quad \sum_{c=1}^{C} u_{kfc}\, \pi_a^{(c)} \exp(R^{(c)}T)_{ab} \quad \text{(align } (a, b)) \tag{C.71}$$

$$\texttt{InsFragType}[k, f] \xrightarrow{(\epsilon,b)} \texttt{InsEmit}[k, f] : \quad \sum_{c=1}^{C} u_{kfc}\, \pi_b^{(c)} \quad \text{(insert } b) \tag{C.72}$$

$$\texttt{DelFragType}[k, f] \xrightarrow{(a,\epsilon)} \texttt{DelEmit}[k, f] : \quad \sum_{c=1}^{C} u_{kfc}\, \pi_a^{(c)} \quad \text{(delete } a) \tag{C.73}$$

$$\texttt{IFragType}[k, f] \xrightarrow{(\epsilon,b)} \texttt{IEmit}[k, f] : \quad \sum_{c=1}^{C} u_{kfc}\, \pi_b^{(c)} \quad \text{(insert } b) \tag{C.74}$$

$$\texttt{DFragType}[k, f] \xrightarrow{(a,\epsilon)} \texttt{DEmit}[k, f] : \quad \sum_{c=1}^{C} u_{kfc}\, \pi_a^{(c)} \quad \text{(delete } a) \tag{C.75}$$

**Intra-fragment fragment-type transition vs. fragment termination** Within each fragment of domain $k$ the fragment-type process is a Markov chain on $\mathcal{F}+2$ states (start, end, and $\mathcal{F}$ fragment-type states): from the current emit state with fragment-type $f$, the chain either advances within the fragment to fragment-type $g$ with probability $r_{fg}^{(k)}$, or terminates the fragment (transition to the end state) with probability $\rho_f^{(k)} = 1 - \sum_g r_{fg}^{(k)}$. Different fragments are statistically independent realisations of this chain. The transition from each emit state goes to any fragment-type's type-selection state within the current fragment (not just the same type), or to the fragment end:

$$\texttt{MatEmit}[k, f] \to \texttt{MatFragType}[k, g] : \quad r_{fg}^{(k)} \quad \text{(intra-fragment type transition } f \to g) \tag{C.76}$$

$$\texttt{MatEmit}[k, f] \to \texttt{MatFragEnd}[k, f] : \quad \rho_f^{(k)} \quad \text{(fragment termination)} \tag{C.77}$$

$$\texttt{InsEmit}[k, f] \to \texttt{InsFragType}[k, g] : \quad r_{fg}^{(k)} \tag{C.78}$$

$$\texttt{InsEmit}[k, f] \to \texttt{InsFragEnd}[k, f] : \quad \rho_f^{(k)} \tag{C.79}$$

$$\texttt{DelEmit}[k, f] \to \texttt{DelFragType}[k, g] : \quad r_{fg}^{(k)} \tag{C.80}$$

$$\texttt{DelEmit}[k, f] \to \texttt{DelFragEnd}[k, f] : \quad \rho_f^{(k)} \tag{C.81}$$

$$\texttt{IEmit}[k, f] \to \texttt{IFragType}[k, g] : \quad r_{fg}^{(k)} \tag{C.82}$$

$$\texttt{IEmit}[k, f] \to \texttt{IFragEnd}[k, f] : \quad \rho_f^{(k)} \tag{C.83}$$

$$\texttt{DEmit}[k, f] \to \texttt{DFragType}[k, g] : \quad r_{fg}^{(k)} \tag{C.84}$$

$$\texttt{DEmit}[k, f] \to \texttt{DFragEnd}[k, f] : \quad \rho_f^{(k)} \tag{C.85}$$

where $g$ ranges over all $\mathcal{F}$ fragment types. For $\mathcal{F} = 1$, this reduces to a scalar self-extension with $r_{11}^{(k)} = r_{k1}$ and $\rho_1^{(k)} = 1 - r_{k1}$.

##### C.3.3 Null State Classification

Every state except `S`, `E`, and the five emit states `MatEmit`$[k, f]$, `InsEmit`$[k, f]$, `DelEmit`$[k, f]$, `IEmit`$[k, f]$, `DEmit`$[k, f]$ is a **null state** (non-emitting). The collapsed model retains only `S`, `E`, and the $5\mathcal{N}\mathcal{F}$ emit states.

##### C.3.4 Null Elimination

The null states are eliminated by the standard HMM null closure:

$$\chi_{\text{emit,emit}} = T_{\text{emit,emit}} + T_{\text{emit,null}}(I - T_{\text{null,null}})^{-1}T_{\text{null,emit}}$$

Since there are no direct emit→emit transitions in the exploded model (every path between emit states passes through at least one null state), $T_{\text{emit,emit}} = 0$ and:

$$\chi = T_{\text{emit,null}}(I - T_{\text{null,null}})^{-1}T_{\text{null,emit}}$$

This gives the collapsed $\chi$ matrix with states $\{\mathtt{S}, \mathtt{E}, \mathtt{MM}_{kf}, \mathtt{MI}_{kf}, \mathtt{MD}_{kf}, \mathtt{II}_{kf}, \mathtt{DD}_{kf}\}$, matching the collapsed MixDom Pair HMM in Section C.1.1.

Each state in the $(5\mathcal{N}\mathcal{F}+2)$-state collapsed HMM corresponds to an uneliminated state in the $(15\mathcal{N}\mathcal{F}+8\mathcal{N}+8)$-state exploded HMM: either `S` (`SS`), `E` (`EE`), or one of the emit states $\mathtt{MatEmit}[l,f]$ ($\mathtt{MM}_{lf}$), $\mathtt{InsEmit}[l,f]$ ($\mathtt{MI}_{lf}$), $\mathtt{DelEmit}[l,f]$ ($\mathtt{MD}_{lf}$), $\mathtt{IEmit}[l,f]$ ($\mathtt{II}_{lf}$), $\mathtt{DEmit}[l,f]$ ($\mathtt{DD}_{lf}$). The transition weight from $\mathtt{UX}_{lf}$ to $\mathtt{VY}_{mg}$ in the collapsed Pair HMM has the form $p_{\text{out}}(\mathtt{U},\mathtt{X},l,f) \times \mathcal{T}_{\mathtt{UV}}(\mathtt{U},\mathtt{V}) \times p_{\text{in}}(\mathtt{V},\mathtt{Y},m,g) + \delta_{\mathtt{UV}}\delta(l=m)(p_{\text{SameDom}}(\mathtt{U},\mathtt{X},l,f,g) + \delta_{\mathtt{XY}}p_{\text{SameFrag}}(l,f,g))$ corresponding to the following path segments

- $p_{\text{out}}(\mathtt{U},\mathtt{X},l,f)$ represents transitions from the emit state to the end state of the domain, e.g. $\tau_{\mathtt{ME}}(\mathtt{M},\mathtt{I},l,f)$ represents $\mathtt{InsEmit}[l,f] \to \mathtt{InsFragEnd}[l,f] \to \mathtt{MatDomEnd}$;

- $\mathcal{T}_{\mathtt{UV}}(\mathtt{U},\mathtt{V})$ represents the sum over all paths from the domain end state (or `S`), through zero or more empty domains, to the next (nonempty) domain start (or `E`), e.g. $\mathcal{T}_{\mathtt{MD}}(\mathtt{M},\mathtt{D})$ represents paths like $\mathtt{MatDomEnd} \to (\ldots\mathtt{InsDom} \to \mathtt{InsDomType}[\mathcal{N}'] \to \mathtt{InsDomEnd}\ldots)^* \to \mathtt{DelDom}$. This is where null cycle elimination happens;

- $p_{\text{in}}(\mathtt{V},\mathtt{Y},m,g)$ represents paths from the domain start state to an emit state inside the domain, e.g. $\tau_{\mathtt{SM}}(\mathtt{M},\mathtt{I},m,g)$ represents $\mathtt{MatDom} \to \mathtt{MatDomType}[m] \to \mathtt{InsFrag}[m] \to \mathtt{InsFragType}[m,g]$;

- $p_{\text{SameDom}}(\mathtt{X},\mathtt{Y},l,f,g)$ represents paths from the emit state to another emit state of similar profile but (potentially) different fragment type within the same domain, e.g. $p_{\text{SameDom}}(\mathtt{M},\mathtt{I},l,f,g)$ represents $\mathtt{InsEmit}[l,f] \to \mathtt{MatDomEnd} \to \mathtt{MatDom} \to \mathtt{MatDomType}[l] \to \mathtt{InsFrag}[l] \to \mathtt{InsFragType}[l,g]$;

- $p_{\text{SameFrag}}(l,f,g) = r^{(l)}_{fg}$ represents intra-fragment Markov transitions between fragment-types (extending the current fragment by one position), e.g. $\mathtt{MatEmit}[l,f] \to \mathtt{MatFragType}[l,g]$ with weight $r^{(l)}_{fg}$. Different fragments are independent; the Markov structure is strictly *within* a single fragment, allowing the fragment-type to change at each position (not just self-loop).

##### C.3.5 Exact Count Restoration

Given Forward-Backward expected transition counts $\hat{n}_\chi(i,j)$ on the collapsed model, we recover the expected counts on the exploded model using the null closure inverse.

Define:

$$C = (I - T_{\text{null,null}})^{-1} \quad \text{(null closure)} \tag{C.86}$$

$$C_{ab} = \text{expected visits to null state } b\text{, starting from null state } a \tag{C.87}$$

For each collapsed transition $\hat{n}_\chi(s,s')$ from emit state $s$ to emit state $s'$, the path in the exploded model is:

$$s \xrightarrow{1} \text{FragEnd}(s) \xrightarrow{\text{null chain}} \text{FragType}(s') \xrightarrow{1} s'$$

The null chain from FragEnd($s$) to FragType($s'$) passes through a sequence of null states. The expected count for each null transition ($a \to b$) along this chain is:

$$\hat{n}_{\text{exploded}}(a, b) = \sum_{s,s'} \hat{n}_\chi(s, s') \cdot \frac{T_{\text{emit},a'} \cdot C_{a',a} \cdot T_{a,b} \cdot C_{b,b'} \cdot T_{b',\text{emit}'}}{\chi(s, s')} \tag{C.88}$$

where $a'$ is the first null state entered from $s$, and $b'$ is the last null state before reaching $s'$.

More explicitly, each collapsed transition decomposes into contributions to the following parameter groups.

**Intra-fragment type transitions vs. new-fragment transitions** For a transition $\hat{n}_\chi(s, s')$ where $s = \text{Emit}^X_{kf}$ and $s' = \text{Emit}^Y_{kg}$ with the same domain $k$, the path splits into:

- **Intra-fragment fragment-type transition**: $s \to \text{FragType}_{kg} \to s'$, with weight $r^{(k)}_{fg}$. This applies to all transitions where $X = Y$ (same TKF state type) and allows $f \neq g$. It extends the current fragment by one position without generating a new TKF92 link.
- **New fragment via domain loop**: $s \to \text{FragEnd}_{kf} \to \ldots \to \text{Frag}_k \to \text{FragType}_{kg} \to s'$, with weight $\rho^{(k)}_f \cdot \tau_k[X, Y] \cdot w_{kg}$ (for M-type) or $\rho^{(k)}_f \cdot \kappa_k \cdot w_{kg}$ (for I/D-type). This terminates the current fragment and initiates an independent fresh fragment via the TKF92 process.

The expected intra-fragment fragment-type transition count from $f$ to $g$ is:

$$\hat{n}_{\text{ext}}(k, f, g) = \hat{n}_\chi(s, s') \cdot \frac{r^{(k)}_{fg}}{r^{(k)}_{fg} + \rho^{(k)}_f \cdot p_{\text{new},g}} \tag{C.89}$$

where $p_{\text{new},g}$ is the new-fragment-to-type-$g$ probability. These counts form an $\mathcal{F} \times \mathcal{F}$ matrix per domain, and the M-step row-normalizes $(\hat{n}_{\text{ext}}(k, f, g), \hat{n}_\rho(k, f))$ to obtain the updated $r^{(k)}_{fg}$.

**Intra-domain TKF transitions** Each new-fragment transition (after fragment termination) contributes one TKF transition at the domain level: $\tau_k[X, Y]$ for M-type domains, or $\kappa_k$ / $(1 - \kappa_k)$ for I/D-type domains.

These counts go into the domain-$k$ TKF91 count matrix (for M-type) or the $\kappa_k$ / $(1 - \kappa_k)$ accumulators (for I/D-type).

**Domain entry/exit and phantom counts** Inter-domain transitions pass through `MatDomEnd`/`InsDomEnd`/`DelD`[illegible] (exit from source domain) and `MatDom`/`InsDom`/`DelDom` then `MatDomType`[$k$] (entry to destination domain).

Within the entry, the path `MatDomType`[$k$] $\to$ `MatFrag`[$k$] uses $\tau_k[\texttt{S}, \cdot]$. The phantom path `MatDomType`[$k$] $\to$ `MatDomEnd` has probability $\tau_k[\texttt{S}, \texttt{E}] = (1 - \beta_k)(1 - \kappa_k)$ and contributes a phantom birth-death event to domain $k$'s BDI statistics.

Similarly for I/D-type entries: `InsDomType`[$k$] $\to$ `InsDomEnd` with probability $(1 - \kappa_k)$ is a phantom I/D-type domain.

**Top-level TKF transitions** Each inter-domain transition contributes one TKF transition at the top level: $\tau_0[U, V]$ where $U \in \{\texttt{S}, \texttt{M}, \texttt{I}, \texttt{D}\}$ is the domain-end type and $V$ is the domain-start type.

The null domain paths (empty domains via `MatDomType`[$k$] $\to$ `MatDomEnd`) contribute additional phantom top-level transitions via the null closure $(I - T_{\text{null,null}})^{-1}$.

**Domain and fragment weight counts** Each domain entry contributes one count to $v_k$ (domain weight). Each fragment-type selection contributes one count to $w_{kf}$ (fragment weight).

##### C.3.6 Parameter Group Decomposition

Each transition in the exploded model involves exactly one of the following parameter factors:

| Parameter | Factor | Where it appears |
|---|---|---|
| $\alpha_0$ | $(1-\beta_0)\kappa_0\alpha_0$ | Top-level $\to$ `MatDom` |
| $1-\alpha_0$ | $(1-\beta_0)\kappa_0(1-\alpha_0)$ | Top-level $\to$ `DelDom` |
| $\beta_0$ | $\beta_0$ | Top-level $\to$ `InsDom` |
| $1-\beta_0$ | $(1-\beta_0)$ | Top-level $\to$ `MatDom`, `DelDom`, `E` |
| $\gamma_0$ | $\gamma_0$ | `DelDomEnd` $\to$ `InsDom` |
| $1-\gamma_0$ | $(1-\gamma_0)$ | `DelDomEnd` $\to$ `MatDom`, `DelDom`, `E` |
| $\kappa_0$ | $\kappa_0$ | Top-level $\to$ `MatDom`, `DelDom` |
| $1-\kappa_0$ | $(1-\kappa_0)$ | Top-level $\to$ `E` |
| $\alpha_k$ | $(1-\beta_k)\kappa_k\alpha_k$ | Domain-$k$ $\to$ MatFrag |
| $\beta_k$ | $\beta_k$ | Domain-$k$ $\to$ InsFrag |
| $\gamma_k$ | $\gamma_k$ | Domain-$k$ DelFragEnd $\to$ InsFrag |
| $\kappa_k$ | $\kappa_k$ | Domain-$k$ $\to$ MatFrag/DelFrag, I/D-type continuation |
| $1-\kappa_k$ | $(1-\kappa_k)$ | Domain-$k$ $\to$ DomEnd |
| $r_{fg}^{(k)}$ | $r_{fg}^{(k)}$ | Intra-fragment fragment-type transition $f \to g$ |
| $\rho_f^{(k)}$ | $1-\sum_g r_{fg}^{(k)}$ | Fragment termination |
| $v_k$ | $v_k$ | Domain type selection |
| $w_{kf}$ | $w_{kf}$ | Fragment type selection |

Because each exploded transition involves a product of these factors, and each factor's log depends on at most one natural parameter ($\lambda_k$ or $\mu_k$), the Q-function on the exploded model decomposes into independent BDI score terms. The null-state count restoration maps collapsed counts exactly onto exploded counts, allowing the M-step to decompose into the same parameter-group updates used in the component TKF91/TKF92 models, together with standard mixture-weight updates.

#### C.4 Order-1 Maraschino: Distilled Adjacency Frequencies

"Cherries" are pairwise training examples chosen, in place of full phylogenetically-annotated multiple sequence alignments, as a composite likelihood approximation to the full phylogenetic likelihood (40).

"Maraschino Cherries" are order-1 counts tensors that summarize the adjacency statistics of such pairwise alignments. MixDom's Maraschino Cherries generalize CherryML to include context-dependent substitution and indel patterns (40).

The Maraschino pipeline has two phases. First (Section C.4.1), pairwise alignments are reduced to fixed-shape *cherry-count* tensors that aggregate adjacency statistics binned by divergence time. Second (Section C.4.2), the parameters of the MixDom Pair HMM (Section C.1.1) are estimated by maximizing the cherry-count log-likelihood: a composite likelihood that scores each adjacency in each time bin under the latent-marginalized collapsed Pair HMM transition matrix $\chi(\theta, T)$ derived in Section C.1.1. After fitting, the MixDom model is then distilled to compact order-1 machines (an HMM and a WFST) suitable for use in tree algorithms (Sections C.4.5 and C.4.6).

##### C.4.1 Cherry-count summary statistics

The input to the Maraschino fitter is a precomputed tensor of pairwise adjacency counts. For each multiple sequence alignment, sibling pairs are extracted, gapped columns dropped, and the resulting pairwise alignment classified into *adjacency contexts*: for every pair of consecutive non-empty alignment columns, we record the source column type (`S`, `M`, `I`, `D`), the destination column type (`M`, `I`, `D`, `E`), and the ancestor/descendant characters in each.

The pairwise $p$-distance of each cherry is converted to an estimated divergence time $T$, and $T$ is discretized into $n_\tau$ geometric bins $\{T_1, \ldots, T_{n_\tau}\}$ with representative bin centres $\bar{T}_b$. Counts are accumulated per bin into the following tensors over the amino-acid alphabet $\mathcal{A}$ ($|\mathcal{A}| = 20$), with extended vocabulary $\mathcal{A} \cup \{\mathtt{S}, \mathtt{E}\}$ for boundary positions:

| Tensor | Shape | Meaning |
|---|---|---|
| $B$ | $n_\tau \times (\|\mathcal{A}\| + 2)^2$ | Singlet bigrams (incl. `S`/`E`) |
| $C^{\mathtt{MM}}$ | $n_\tau \times \|\mathcal{A}\|^4$ | Match→Match: $(a, b, a', b')$ |
| $C^{\mathtt{MI}}$ | $n_\tau \times \|\mathcal{A}\|^3$ | Match→Insert: $(a, b, b')$ |
| $C^{\mathtt{MD}}$ | $n_\tau \times \|\mathcal{A}\|^3$ | Match→Delete: $(a, b, a')$ |
| $C^{\mathtt{IM}}$ | $n_\tau \times \|\mathcal{A}\|^3$ | Insert→Match: $(b, a', b')$ |
| $C^{\mathtt{II}}$ | $n_\tau \times \|\mathcal{A}\|^2$ | Insert→Insert: $(b, b')$ |
| $C^{\mathtt{ID}}$ | $n_\tau \times \|\mathcal{A}\|^2$ | Insert→Delete: $(b, a')$ |
| $C^{\mathtt{DM}}$ | $n_\tau \times \|\mathcal{A}\|^3$ | Delete→Match: $(a, a', b')$ |
| $C^{\mathtt{DD}}$ | $n_\tau \times \|\mathcal{A}\|^2$ | Delete→Delete: $(a, a')$ |
| $C^{\mathtt{DI}}$ | $n_\tau \times \|\mathcal{A}\|^2$ | Delete→Insert: $(a, b')$ |
| $C^{\mathtt{SM}}$ | $n_\tau \times \|\mathcal{A}\|^2$ | Start→Match: $(a', b')$ |
| $C^{\mathtt{SI}}$ | $n_\tau \times \|\mathcal{A}\|$ | Start→Insert: $(b')$ |
| $C^{\mathtt{SD}}$ | $n_\tau \times \|\mathcal{A}\|$ | Start→Delete: $(a')$ |
| $C^{\mathtt{ME}}$ | $n_\tau \times \|\mathcal{A}\|^2$ | Match→End: $(a, b)$ |
| $C^{\mathtt{IE}}$ | $n_\tau \times \|\mathcal{A}\|$ | Insert→End: $(b)$ |
| $C^{\mathtt{DE}}$ | $n_\tau \times \|\mathcal{A}\|$ | Delete→End: $(a)$ |
| $C^{\mathtt{SE}}$ | $n_\tau$ | Start→End (empty alignment) |

The post-Insert and post-Delete tensors carry only the adjacent emitted character ($b$ for inserts, $a$ for deletes) as context, not the previous match's full $(a, b)$ context: the model will marginalise its context-rich frequencies down to this reduced context when computing the likelihood. The Match-to-Match tensor $C^{\mathtt{MM}}$ is the largest and dominates the parameter budget. Boundary tensors record alignments that begin or end in a particular adjacency type.

##### C.4.2 Cherry-count likelihood for the MixDom Pair HMM

The cherry-count tensors of Section C.4.1 are scored against the collapsed MixDom Pair HMM $\chi(\theta, T)$ defined by equation (C.3) of Section C.1.1. The free parameters are

$$\theta = \big(\lambda_0, \mu_0,\ \{\lambda_n, \mu_n\}_{n=1}^{\mathcal{N}},\ \{\upsilon_n\}_{n=1}^{\mathcal{N}},\ \{w_{nf}\}_{n,f},\ \{r_{fg}^{(n)}\}_{n,f,g},\ \{u_{nfc}\}_{n,f,c},\ \{\pi^{(c)}, Q^{(c)}\}_{c=1}^{C}\big),$$

parameterised in unconstrained space (log-rates, log-Dirichlet weights, log-exchangeabilities) so that gradient methods are unconstrained. Note in particular:

- $w_{nf}$ is the per-domain Dirichlet over fragment entry types (initial state of the intra-fragment Markov chain).

- $r^{(n)}_{fg}$ is a per-domain $\mathcal{F} \times \mathcal{F}$ row-stochastic matrix (with row sums $\leq 1$) giving the intra-fragment Markov transition probability from fragment-type $f$ to fragment-type $g$ within the same fragment; the residual mass $\rho^{(n)}_f = 1 - \sum_g r^{(n)}_{fg}$ is the fragment-termination probability. The TKF92 scalar self-extension is the special case $\mathcal{F} = 1$, with $r^{(n)}_{11}$ playing the role of the TKF92 extension probability (the M-step closed forms for both reduce to a single binary count split per row of $r^{(n)}$).
- $u_{nfc}$ is a per-(domain, fragment-type) Dirichlet over $C$ *static* site classes (drawn independently per emitted site, with no chain-time class switching).
- Each site class $c$ has its own reversible substitution model $\text{Subst}(Q^{(c)}, \pi^{(c)})$ with rate matrix $R^{(c)} = Q^{(c)} \,\text{diag}(\pi^{(c)})$. All rate variation across sites is captured by these per-class GTR matrices; there is no separate Yang-style discretized-gamma rate-multiplier mechanism.

**Pair adjacency frequencies.** For each $T$-bin centre $\bar{T}_b$, the collapsed $(5\mathcal{N}\mathcal{F}+2)$-state Pair HMM transition matrix $\chi^{(b)} \equiv \chi(\theta, \bar{T}_b)$ is constructed via the closed form of equation (C.3). The marginal stationary distribution $\pi^{\text{stat}}_b$ over emitting states is obtained as the left null vector of $I - \chi^{(b)}_{\Omega\Omega}$, where $\Omega$ denotes the set of $5\mathcal{N}\mathcal{F}$ emitting states.

For source machine state $u \in \{\mathtt{S}, \mathtt{M}, \mathtt{I}, \mathtt{D}\}$, destination machine state $v \in \{\mathtt{M}, \mathtt{I}, \mathtt{D}, \mathtt{E}\}$, and characters $(a, b, a', b')$ in their respective slots, the *model-side adjacency frequency*

$$F^{uv}_b(a, b; a', b') = \sum_{\substack{s \in \Omega^u \\ s' \in \Omega^v}} \left(\pi^{\text{stat}}_{b,s}\right) e_s(a, b)\, \chi^{(b)}_{ss'}\, e_{s'}(a', b')$$

sums over latent (domain, fragment-type) realisations of source and destination collapsed states $s, s'$, weighting by their stationary probability and emission probabilities. The emission probability of a Match state $\mathtt{MM}_{nf}$ emitting $(a, b)$ marginalises the static site-class mixture:

$$e_{\mathtt{MM}_{nf}}(a, b) = \sum_{c=1}^{C} u_{nfc}\, \pi^{(c)}_a \exp(R^{(c)} T)_{ab}, \tag{C.90}$$

and emission probabilities for $\mathtt{MI}, \mathtt{II}, \mathtt{MD}, \mathtt{DD}$ analogously marginalise the same class mixture but emit only one of $(a, b)$:

$$e_{\mathtt{II}_{nf}}(b) = e_{\mathtt{MI}_{nf}}(b) = \sum_c u_{nfc}\, \pi^{(c)}_b, \quad e_{\mathtt{DD}_{nf}}(a) = e_{\mathtt{MD}_{nf}}(a) = \sum_c u_{nfc}\, \pi^{(c)}_a.$$

Boundary frequencies $F^{\mathtt{S}v}_b$, $F^{u\mathtt{E}}_b$, $F^{\mathtt{SE}}_b$ replace the corresponding endpoint factor with the $\mathtt{S}$ row or $\mathtt{E}$ column of $\chi^{(b)}$. The reduced-context frequencies needed by post-Insert and post-Delete counts are obtained by marginalisation:

$$F^{\mathtt{I}v}_b(b; \cdot) = \sum_a F^{\mathtt{I}v}_b(a, b; \cdot), \qquad F^{\mathtt{D}v}_b(a; \cdot) = \sum_b F^{\mathtt{D}v}_b(a, b; \cdot).$$

For each context $(u; a, b)$ at bin $b$, the row normalisation constant is

$$Z^u_b(a, b) = \sum_v \sum_{a', b'} F^{uv}_b(a, b; a', b'),$$

where the inner sum runs over the characters carried by the destination adjacency type, and the post-Insert/post-Delete normalisations use the reduced-context frequencies above.

**Cherry-count log-likelihood.** The composite log-likelihood that the Maraschino fitter maximises is

$$\mathcal{L}_{\text{cherry}}(\theta) = \mathcal{L}_{\text{singlet}}(\theta) + \sum_{b=1}^{n_\tau} \mathcal{L}_{\text{pair},b}(\theta), \tag{C.91}$$

$$\mathcal{L}_{\text{singlet}}(\theta) = \sum_{X,Y \in \mathcal{A} \cup \{\mathtt{S},\mathtt{E}\}} \left( \textstyle\sum_b B_{XY}^{(b)} \right) \log P_{XY}^{\text{singlet}}(\theta), \tag{C.92}$$

$$\mathcal{L}_{\text{pair},b}(\theta) = \sum_{u,v} \sum_{a,b,a',b'} C_{ab\,a'b'}^{uv,(b)} \log\left( \frac{F_b^{uv}(a,b;a',b')}{Z_b^u(a,b)} \right). \tag{C.93}$$

Here $P^{\text{singlet}}$ is the order-1 transition matrix obtained from the MixDom Singlet HMM by row-normalising its adjacency frequencies (Section C.4.5); the singlet term scores the bigram counts $B$ summed over time bins, since the singlet model is time-independent. The pair term scores each per-bin adjacency tensor under its own row-normalised conditional distribution.

**Optimisation.** $\mathcal{L}_{\text{cherry}}(\theta)$ is differentiable in $\theta$. The maximisation is carried out by gradient methods (Adam, optionally L-BFGS for refinement) on the unconstrained parameterisation, using the same MixDom initialiser as the exact Baum–Welch trainer (including the same flags for the number of site classes, the class-equilibrium initialisation, and the fragment-class assignment), so the two fitters can be started from identical parameters and compared directly. Because the M-step of the exact-EM trainer is closed-form and Maraschino's gradient optimiser is not, this provides a controlled comparison of cherry-count fitting against full Baum–Welch on the same data and architecture.

The fitted MixDom parameters are written to a checkpoint with the same key layout as a `train_pfam`-produced checkpoint, so that either trainer's output can be loaded and refined by the other and either can be used as input to the order-1 distillation (Sections C.4.5 and C.4.6).

##### C.4.3 Distillation From MixDom To Order-1 Machines

The MixDom HMMs defined above have large structured state spaces (domains $\times$ fragments). We now show how to distill these into compact *order-1* machines—an HMM and a transducer—whose transition probabilities depend only on the most recently emitted characters. These machines are approximations of the full MixDom model. The MixDom model parameters are assumed to have been trained, e.g. via Baum-Welch or via Maraschino cherry-count fitting (Section C.4.2); see also (30).

The distillation operates on the alphabet $|\mathcal{A}|$ (with the per-(domain, fragment-type) site-class mixture marginalised implicitly inside the match emission tensors of equation (C.90)). The Woodbury structural weights depend only on indel parameters; the per-(class) emission tensors $\pi_a^{(c)} \exp(R^{(c)}T)_{ab}$, weighted by $u_{nfc}$, are contracted with those weights to produce the order-1 transition probabilities.

Let $|\mathcal{A}|$ denote the alphabet size and $\circ$ denote a distinguished beginning-of-sequence (BOS) symbol.

##### C.4.4 Notation for path marginalizations

Both distillations require marginalizing over null (non-emitting) states between consecutive emissions. We introduce a compact notation for the expected frequency of partially observed paths.

In any HMM or transducer with states partitioned into emitting states $\Omega$ and non-emitting (null) states $\mathcal{Z}$, define the *null closure*

$$N^* \equiv (I - T_{\mathcal{Z}\mathcal{Z}})^{-1}$$

where $T_{\mathcal{Z}\mathcal{Z}}$ is the submatrix of transitions among null states. The effective transition matrix from emitting-or-start to emitting-or-end, marginalizing all intervening null paths, is

$$\hat{T}_{ij} = T_{ij} + \sum_{p,q \in \mathcal{Z}} T_{ip}\, N^*_{pq}\, T_{qj} \qquad i \in \Omega \cup \{\mathtt{S}\},\ j \in \Omega \cup \{\mathtt{E}\}$$

For paths through the MixDom HMMs, we use the notation

$$\mathcal{E}[\,\mathtt{S} \to \underbrace{s_1}_{(a_1,b_1)} \xrightarrow{\mathcal{Z}^*} \underbrace{s_2}_{(a_2,b_2)} \to \cdots \to \mathtt{E}\,]$$

to denote the expected number of times the path visits the indicated sequence of emitting states (with indicated emissions in subscript) separated by zero or more null states (denoted $\mathcal{Z}^*$), summed over all completions of the path to the left ($\mathtt{S} \to \cdots$) and right ($\cdots \to \mathtt{E}$). Formally, if $\pi$ is the stationary distribution over states,

$$\mathcal{E}[\,\underbrace{s_1}_{(a_1,b_1)} \xrightarrow{\mathcal{Z}^*} \underbrace{s_2}_{(a_2,b_2)}\,] = \sum_{s_1 \in \Omega} \sum_{s_2 \in \Omega} \left( \sum_i \pi_i \hat{T}^*_{is_1} \right) e_{s_1}(a_1, b_1)\, \hat{T}_{s_1 s_2}\, e_{s_2}(a_2, b_2) \left( \sum_j \hat{T}^*_{s_2 j} \right)$$

where $e_s(\cdot)$ is the emission probability at state $s$ and $\hat{T}^*_{ij} = [(I - \hat{T}_{\Omega\Omega})^{-1}]_{ij}$ (with $\hat{T}_{\Omega\Omega}$ the submatrix of $\hat{T}$ restricted to emitting states) sums over all paths through emitting states.

##### C.4.5 Distillation to Order-1 HMM

The MixDom Singlet HMM generates sequences from the stationary distribution. We distill it into an order-1 HMM with states

$$\{\mathtt{S}, \mathtt{E}\} \cup \{a : a \in \mathcal{A}\}$$

where state $a$ deterministically emits character $a$. Define the adjacency frequency from the full MixDom Singlet HMM:

$$f(a, b) = \mathcal{E}[\,\underbrace{s_1}_{a} \xrightarrow{\mathcal{Z}^*} \underbrace{s_2}_{b}\,]$$

where the sum is over all emitting states $s_1, s_2$ weighted by emission probabilities $e_{s_1}(a)$ and $e_{s_2}(b)$ as above. Each singlet emission probability $e_{\mathtt{I}_{nf}}(a) = \sum_c u_{nfc} \pi^{(c)}_a$ marginalises the per-(domain, fragment-type) class mixture.

**Parameterization.** The order-1 Singlet HMM transition probabilities are:

$$P(b|\mathtt{S}) = \frac{\sum_{b'} f(b,b')}{\sum_{a',b'} f(a',b')} \qquad \text{(start} \to \text{first emission)} \tag{C.94}$$

$$P(b|a) = \frac{f(a,b)}{\sum_{b'} f(a,b')} \qquad \text{(emission} \to \text{emission)} \tag{C.95}$$

$$P(\mathtt{E}|a) = 1 - \sum_{b} P(b|a) \qquad \text{(emission} \to \text{end)} \tag{C.96}$$

That is: normalize each row of the adjacency matrix $f$ to obtain transition probabilities, allocating the residual probability mass to the end state. The start distribution (C.94) is proportional to the column marginals of $f$.

##### C.4.6 Distillation to Order-1 WFST

The MixDom Pair HMM describes the joint distribution over ancestor-descendant sequence pairs. We distill it into an order-1 transducer (a Mealy machine) whose state depends on the last inputted ancestor character and the last outputted descendant character.

**Machine states.** The transducer has seven machine states $\{\mathtt{S},\mathtt{M},\mathtt{I},\mathtt{D},\mathtt{V},\mathtt{W},\mathtt{E}\}$ organized as a *waiting machine*:

- **Non-waiting** (all outgoing transitions have $\varepsilon$ input): `S` (start), `M` (just matched), `I` (just inserted), `D` (just deleted)
- **Waiting** (all outgoing transitions consume input): `V` (ready after `M` or `I`), `W` (ready after `D`)
- **Terminal**: `E` (end)

The distinction $\mathtt{V} \neq \mathtt{W}$ is needed because outgoing transition weights differ: in the MixDom Pair HMM, the `M/I` rows use $\beta$ while `D` rows use $\gamma$.

**Transitions.** For a state with last-input ancestor $X$ and last-output descendant $Y$ (where $X, Y \in \mathcal{A} \cup \{\circ\}$ and $\circ$ denotes BOS):

| Source | Dest | Input | Output | Weight |
|---|---|---|---|---|
| S | V | $\varepsilon$ | $\varepsilon$ | $p_{\texttt{SV}}$ |
| S | I | $\varepsilon$ | $b$ | $p_{\texttt{SI}}(Y, b)$ |
| S | E | $\varepsilon$ | $\varepsilon$ | $p_{\texttt{SE}}$ |
| V | M | $a$ | $b$ | $p_{\texttt{VM}}(X, Y, a, b)$ |
| V | D | $a$ | $\varepsilon$ | $p_{\texttt{VD}}(X, Y, a)$ |
| W | M | $a$ | $b$ | $p_{\texttt{WM}}(X, Y, a, b)$ |
| W | D | $a$ | $\varepsilon$ | $p_{\texttt{WD}}(X, Y, a)$ |
| M | V | $\varepsilon$ | $\varepsilon$ | $p_{\texttt{MV}}$ |
| M | I | $\varepsilon$ | $b$ | $p_{\texttt{MI}}(X, Y, b)$ |
| M | E | $\varepsilon$ | $\varepsilon$ | $p_{\texttt{ME}}$ |
| I | V | $\varepsilon$ | $\varepsilon$ | $p_{\texttt{IV}}$ |
| I | I | $\varepsilon$ | $b$ | $p_{\texttt{II}}(X, Y, b)$ |
| I | E | $\varepsilon$ | $\varepsilon$ | $p_{\texttt{IE}}$ |
| D | W | $\varepsilon$ | $\varepsilon$ | $p_{\texttt{DW}}$ |
| D | I | $\varepsilon$ | $b$ | $p_{\texttt{DI}}(X, Y, b)$ |
| D | E | $\varepsilon$ | $\varepsilon$ | $p_{\texttt{DE}}$ |

Note that `V` and `W` are *not* associated with emissions; they serve only to enforce the waiting-machine property. Transitions from `V` and `W` always consume an ancestor input symbol; transitions from `M`, `I`, `D`, and `S` never do.

**Parameterization from the MixDom Pair HMM.** We need the expected frequency of transitions conditioned on (last ancestor input $X$, last descendant output $Y$, transition type, new symbols). The key subtlety: through insert states, the last ancestor symbol $X$ must be *propagated* from the preceding match or delete, since inserts do not consume input.

Using the path notation from above, we enumerate all adjacency types that arise in the MixDom Pair HMM. Write $\texttt{M}[X, Y]$ for a match state that inputs $X$ and outputs $Y$, $\texttt{I}[Y]$ for an insert that outputs $Y$, and $\texttt{D}[X]$ for a delete that inputs $X$. The last-ancestor and last-descendant context is carried implicitly.

**Adjacency frequencies.** The following table lists all adjacency types and their corresponding path marginalizations. In each case, the frequency is computed as a sum over MixDom Pair HMM states, with null states marginalized via $N^*$. Write $\circ$ for the boundary (start/end) context.

| Context | Adjacency | MixDom path | Frequency |
|---|---|---|---|
| *Start* $\to$ *Match:* | | | |
| | $\mathtt{S} \to \mathtt{M}[X', Y']$ | $\mathtt{S} \xrightarrow{\mathcal{Z}^*} \mathtt{M}[X', Y']$ | $f^{\mathtt{SM}}(X', Y')$ |
| *Start* $\to$ *Insert:* | | | |
| | $\mathtt{S} \to \mathtt{I}[Y']$ | $\mathtt{S} \xrightarrow{\mathcal{Z}^*} \mathtt{I}[Y']$ | $f^{\mathtt{SI}}(Y')$ |
| *Start* $\to$ *End (empty sequence):* | | | |
| | $\mathtt{S} \to \mathtt{E}$ | $\mathtt{S} \xrightarrow{\mathcal{Z}^*} \mathtt{E}$ | $f^{\mathtt{SE}}$ |
| *Match* $\to$ *Match (via null states only):* | | | |
| | $\mathtt{M}[X, Y] \to \mathtt{M}[X', Y']$ | $\mathtt{M}[X, Y] \xrightarrow{\mathcal{Z}^*} \mathtt{M}[X', Y']$ | $f^{\mathtt{MM}}(X, Y, X', Y')$ |
| *Match* $\to$ *Insert:* | | | |
| | $\mathtt{M}[X, Y] \to \mathtt{I}[Y']$ | $\mathtt{M}[X, Y] \xrightarrow{\mathcal{Z}^*} \mathtt{I}[Y']$ | $f^{\mathtt{MI}}(X, Y, Y')$ |
| *Match* $\to$ *Delete:* | | | |
| | $\mathtt{M}[X, Y] \to \mathtt{D}[X']$ | $\mathtt{M}[X, Y] \xrightarrow{\mathcal{Z}^*} \mathtt{D}[X']$ | $f^{\mathtt{MD}}(X, Y, X')$ |
| *Match* $\to$ *End:* | | | |
| | $\mathtt{M}[X, Y] \to \mathtt{E}$ | $\mathtt{M}[X, Y] \xrightarrow{\mathcal{Z}^*} \mathtt{E}$ | $f^{\mathtt{ME}}(X, Y)$ |
| *Insert* $\to$ *Insert (ancestor context $X$ propagated):* | | | |
| | $\mathtt{I}[Y] \to \mathtt{I}[Y']$ | $\mathtt{I}[Y] \xrightarrow{\mathcal{Z}^*} \mathtt{I}[Y']$ | $f^{\mathtt{II}}(X, Y, Y')$ |
| *Insert* $\to$ *Match (ancestor context $X$ propagated):* | | | |
| | $\mathtt{I}[Y] \to \mathtt{M}[X', Y']$ | $\mathtt{I}[Y] \xrightarrow{\mathcal{Z}^*} \mathtt{M}[X', Y']$ | $f^{\mathtt{IM}}(X, Y, X', Y')$ |
| *Insert* $\to$ *Delete (ancestor context $X$ propagated):* | | | |
| | $\mathtt{I}[Y] \to \mathtt{D}[X']$ | $\mathtt{I}[Y] \xrightarrow{\mathcal{Z}^*} \mathtt{D}[X']$ | $f^{\mathtt{ID}}(X, Y, X')$ |
| *Insert* $\to$ *End (ancestor context $X$ propagated):* | | | |
| | $\mathtt{I}[Y] \to \mathtt{E}$ | $\mathtt{I}[Y] \xrightarrow{\mathcal{Z}^*} \mathtt{E}$ | $f^{\mathtt{IE}}(X, Y)$ |
| *Delete* $\to$ *Match (descendant context $Y$ propagated):* | | | |
| | $\mathtt{D}[X] \to \mathtt{M}[X', Y']$ | $\mathtt{D}[X] \xrightarrow{\mathcal{Z}^*} \mathtt{M}[X', Y']$ | $f^{\mathtt{DM}}(X, Y, X', Y')$ |
| *Delete* $\to$ *Delete (descendant context $Y$ propagated):* | | | |
| | $\mathtt{D}[X] \to \mathtt{D}[X']$ | $\mathtt{D}[X] \xrightarrow{\mathcal{Z}^*} \mathtt{D}[X']$ | $f^{\mathtt{DD}}(X, Y, X')$ |
| *Delete* $\to$ *Insert (descendant context $Y$ propagated):* | | | |
| | $\mathtt{D}[X] \to \mathtt{I}[Y']$ | $\mathtt{D}[X] \xrightarrow{\mathcal{Z}^*} \mathtt{I}[Y']$ | $f^{\mathtt{DI}}(X, Y, Y')$ |
| *Delete* $\to$ *End (descendant context $Y$ propagated):* | | | |
| | $\mathtt{D}[X] \to \mathtt{E}$ | $\mathtt{D}[X] \xrightarrow{\mathcal{Z}^*} \mathtt{E}$ | $f^{\mathtt{DE}}(X, Y)$ |

In all cases, the ancestor context $X$ is propagated through insert states (which do not consume input), and the descendant context $Y$ is propagated through delete states (which do not emit output). The notation $\xrightarrow{\mathcal{Z}^*}$ denotes zero or more transitions through null states, marginalized via the null closure $N^*$.

**Computing the frequencies.** Each frequency above is computed from the MixDom Pair HMM as follows. Let $\pi_s$ denote the stationary probability of state $s$ and $\hat{T}_{s_1 s_2}$ the null-marginalized effective transition. For the simplest case (direct adjacency):

$$f^{\mathtt{MD}}(X, Y, X') = \sum_{s_1 \in \Omega^{\mathtt{M}}} \sum_{s_2 \in \Omega^{\mathtt{D}}} \left( \sum_i \pi_i \hat{T}^*_{i s_1} \right) e_{s_1}(X, Y)\, \hat{T}_{s_1 s_2}\, e_{s_2}(X') \left( \sum_j \hat{T}^*_{s_2 j} \right)$$

Boundary frequencies use the start/end rows of $\hat{T}$:

$$f^{\texttt{SM}}(X',Y') = \sum_{s\in\Omega^{\texttt{M}}} \hat{T}_{\texttt{S}s}\, e_s(X',Y') \left(\sum_j \hat{T}^*_{sj}\right), \quad f^{\texttt{ME}}(X,Y) = \sum_{s\in\Omega^{\texttt{M}}} \left(\sum_i \pi_i \hat{T}^*_{is}\right) e_s(X,Y)\, \hat{T}_{s\,\texttt{E}}$$

(and analogously for $f^{\texttt{SI}}$, $f^{\texttt{SE}}$, $f^{\texttt{IE}}$, $f^{\texttt{DE}}$).

**Normalization to transducer parameters.** Given the adjacency frequencies, the order-1 transducer weights are obtained by normalization. For each context $(X,Y)$ and source machine state, normalize outgoing weights to sum to 1:

*Start transitions* (context $\circ$):

$$p_{\texttt{SV}} = \frac{\sum_{X',Y'} f^{\texttt{SM}}(X',Y')}{\sum_{X',Y'} f^{\texttt{SM}}(X',Y') + \sum_{Y'} f^{\texttt{SI}}(Y') + f^{\texttt{SE}}}, \quad p_{\texttt{SI}}(b) = \frac{f^{\texttt{SI}}(b)}{\sum_{X',Y'} f^{\texttt{SM}}(X',Y') + \sum_{Y'} f^{\texttt{SI}}(Y') + f^{\texttt{SE}}}$$

(and $p_{\texttt{SE}}$ uses the same denominator with numerator $f^{\texttt{SE}}$).

*Wait-after-match/insert transitions* (context $(X,Y)$, consuming input $a$):

$$p_{\texttt{VM}}(X,Y,a,b) = \frac{f^{\cdot\texttt{M}}(X,Y,a,b)}{\sum_{a'}\left[\sum_{b'} f^{\cdot\texttt{M}}(X,Y,a',b') + f^{\cdot\texttt{D}}(X,Y,a')\right]}$$

$$p_{\texttt{VD}}(X,Y,a) = \frac{f^{\cdot\texttt{D}}(X,Y,a)}{\sum_{a'}\left[\sum_{b'} f^{\cdot\texttt{M}}(X,Y,a',b') + f^{\cdot\texttt{D}}(X,Y,a')\right]}$$

where $f^{\cdot\texttt{M}}(X,Y,a,b) = f^{\texttt{MM}}(X,Y,a,b) + f^{\texttt{IM}}(X,Y,a,b)$ and $f^{\cdot\texttt{D}}(X,Y,a) = f^{\texttt{MD}}(X,Y,a) + f^{\texttt{ID}}(X,Y,a)$, aggregating over both match and insert sources that share the V wait state.

*Wait-after-delete transitions* (context $(X,Y)$, consuming input $a$):

$$p_{\texttt{WM}}(X,Y,a,b) = \frac{f^{\texttt{DM}}(X,Y,a,b)}{\sum_{a'}\left[\sum_{b'} f^{\texttt{DM}}(X,Y,a',b') + f^{\texttt{DD}}(X,Y,a')\right]}$$

$$p_{\texttt{WD}}(X,Y,a) = \frac{f^{\texttt{DD}}(X,Y,a)}{\sum_{a'}\left[\sum_{b'} f^{\texttt{DM}}(X,Y,a',b') + f^{\texttt{DD}}(X,Y,a')\right]}$$

*Post-match transitions* (context $(X,Y)$, after matching with $a,b$; new context becomes $(a,b)$):

$$p_{\texttt{MV}}(X,Y) = \frac{\sum_{a',b'} f^{\texttt{MM}}(X,Y,a',b') + \sum_{a'} f^{\texttt{MD}}(X,Y,a')}{Z^{\texttt{M}}(X,Y)}$$

$$p_{\texttt{MI}}(X,Y,b) = \frac{f^{\texttt{MI}}(X,Y,b)}{Z^{\texttt{M}}(X,Y)}, \quad p_{\texttt{ME}}(X,Y) = \frac{f^{\texttt{ME}}(X,Y)}{Z^{\texttt{M}}(X,Y)}$$

where $Z^{\texttt{M}}(X,Y) = \sum_{a',b'} f^{\texttt{MM}}(X,Y,a',b') + \sum_{a'} f^{\texttt{MD}}(X,Y,a') + \sum_b f^{\texttt{MI}}(X,Y,b) + f^{\texttt{ME}}(X,Y)$.

*Post-insert transitions* are analogous, using $f^{\texttt{I}\cdot}$ frequencies with $Z^{\texttt{I}}(X,Y)$.

*Post-delete transitions* (context $(X,Y)$, after deleting $a$; new context becomes $(a,Y)$):

$$p_{\texttt{DW}}(X,Y) = \frac{\sum_{a'} f^{\texttt{DD}}(X,Y,a') + \sum_{a',b'} f^{\texttt{DM}}(X,Y,a',b')}{Z^{\texttt{D}}(X,Y)}$$

$$p_{\texttt{DI}}(X,Y,b) = \frac{f^{\texttt{DI}}(X,Y,b)}{Z^{\texttt{D}}(X,Y)}, \quad p_{\texttt{DE}}(X,Y) = \frac{f^{\texttt{DE}}(X,Y)}{Z^{\texttt{D}}(X,Y)}$$

where $Z^{\texttt{D}}(X,Y) = \sum_{a'} f^{\texttt{DD}}(X,Y,a') + \sum_{a',b'} f^{\texttt{DM}}(X,Y,a',b') + \sum_b f^{\texttt{DI}}(X,Y,b) + f^{\texttt{DE}}(X,Y)$.

#### C.5 Algebraic Distillation of MixDom

We investigate whether the distillation of the MixDom model to order-1 machines (Section C.4.5 and C.4.6) can be performed in closed algebraic form, and how the computation scales with the number of domain types $\mathcal{N}$, the number of fragment types per domain $\mathcal{F}$, and the number of site classes $C$. The same algebraic decomposition powers the cherry-count log-likelihood that the Maraschino fitter (Section C.4.2) maximises under MixDom.

##### C.5.1 Setup

The MixDom model has the following parameters (matching Section C.1.1):

- Top-level TKF91: $\lambda_0, \mu_0$ (domain birth/death rates).
- Domain weights: $v_n$, $\sum_n v_n = 1$.
- Per-domain TKF91 rates governing the per-domain TKF92 fragment process: $\lambda_n, \mu_n$ for $n = 1, \ldots, \mathcal{N}$.
- Per-domain fragment-type entry distribution: $w_{nf}$ with $\sum_f w_{nf} = 1$.
- Per-domain intra-fragment Markov ext matrix $r^{(n)}_{fg}$, $\mathcal{F} \times \mathcal{F}$, with row sums $\leq 1$. The fragment-termination probability is $\rho^{(n)}_f = 1 - \sum_g r^{(n)}_{fg}$.
- Per-(domain, fragment-type) site-class Dirichlet: $u_{nfc}$ with $\sum_c u_{nfc} = 1$.
- Per-class reversible substitution model: $(Q^{(c)}, \pi^{(c)})$ for $c = 1, \ldots, C$; rate matrix $R^{(c)} = Q^{(c)} \mathrm{diag}(\pi^{(c)})$.

The total scalar parameter count is $2 + (\mathcal{N} - 1) + 2\mathcal{N} + \mathcal{N}\mathcal{F}^2 + \mathcal{N}(\mathcal{F} - 1) + \mathcal{N}\mathcal{F}(C - 1)$ plus $C$ rate matrices.

Write $\kappa_n = \lambda_n/\mu_n$, $\alpha_n = \alpha(\lambda_n, \mu_n, T)$, $\beta_n = \beta(\lambda_n, \mu_n, T)$, $\gamma_n = \gamma(\lambda_n, \mu_n, T)$, and similarly $\alpha_0, \beta_0, \gamma_0, \kappa_0$ for the top-level parameters.

##### C.5.2 Class-mixture emissions

In MixDom the per-(domain, fragment-type) site-class mixture appears in every emission probability of the collapsed Pair HMM. Define the per-class match emission as

$$P^{(c)}(a, b \mid T) = \pi^{(c)}_a \exp(R^{(c)} T)_{ab}.$$

The model's per-(domain, fragment-type) emission factors at evolutionary time $T$ are:

$$\begin{aligned}
\phi^{\mathtt{M}}_{nf}(a, b) &\equiv e_{\mathtt{MM}_{nf}}(a, b) = \sum_{c=1}^{C} u_{nfc}\, P^{(c)}(a, b \mid T), \\
\phi^{\mathtt{I}}_{nf}(b) &\equiv e_{\mathtt{MI}_{nf}}(b) = e_{\mathtt{II}_{nf}}(b) = \sum_{c} u_{nfc}\, \pi^{(c)}_b, \\
\phi^{\mathtt{D}}_{nf}(a) &\equiv e_{\mathtt{MD}_{nf}}(a) = e_{\mathtt{DD}_{nf}}(a) = \sum_{c} u_{nfc}\, \pi^{(c)}_a.
\end{aligned}$$

Each of these is a $C$-term mixture over the per-class GTRs. The match emission $\phi^{\mathtt{M}}_{nf}$ inherits the time dependence from the per-class transition kernels $P^{(c)}(T)$; the singlet emissions $\phi^{\mathtt{I}}_{nf}$, $\phi^{\mathtt{D}}_{nf}$ are time-independent.

*Remark* C.3 (Bilinear emission tensor). The match emission factors as

$$\phi^{\mathtt{M}}_{nf}(a,b) = \sum_c u_{nfc}\,\pi^{(c)}_a\,\exp(R^{(c)}T)_{ab}$$

which is rank-$C$ in the latent class index, and the resulting $|\mathcal{A}| \times |\mathcal{A}|$ emission tensor at $(n, f)$ has rank at most $C$. The single-class case $u = \delta_{nc}$ makes this rank-1 per domain (a single GTR per domain). The general parameterisation strictly generalises by sharing the GTR pool across (domain, fragment-type) contexts and allowing any soft mixture.

##### C.5.3 Single HMM Distillation

**State space** The MixDom Singlet HMM generates sequences from the stationary distribution. Its collapsed state space (Section C.1.1) is $\{\mathtt{S}, \mathtt{E}\} \cup \{\mathtt{I}_{nf} : n \in \mathcal{N}, f \in \mathcal{F}\}$, so there are $\mathcal{NF}$ emitting states with emissions $\phi^{\mathtt{I}}_{nf}(a)$.

Within a domain, fragment continuation is governed by the intra-fragment Markov ext matrix $r^{(n)}_{fg}$ and the new-fragment $\kappa_n w_{n,g}$ branch. Adapting the formulae in Section C.1.1 for the singlet, the effective transition matrix between emitting states $\mathtt{I}_{lf} \to \mathtt{I}_{mg}$ has entries

$$\hat{T}_{\mathtt{I}_{lf},\mathtt{I}_{mg}} = r^{(l)}_{fg}\delta_{lm} + \rho^{(l)}_f \kappa_l w_{lg}\delta_{lm} + \frac{\rho^{(l)}_f (1-\kappa_l)\,\kappa_0 v_m \kappa_m w_{mg}}{1-\kappa_0 z_0}$$

where $z_0 = \sum_n v_n(1-\kappa_n)$. The path-sum matrix $(I - \hat{T}_{\Omega,\Omega})^{-1}$ is an $\mathcal{NF} \times \mathcal{NF}$ inversion in general.

**Adjacency frequencies** The Singlet adjacency frequency is

$$f(a,b) = \sum_{(l,f),(m,g)} W_{(l,f),(m,g)}\,\phi^{\mathtt{I}}_{lf}(a)\,\phi^{\mathtt{I}}_{mg}(b) \tag{C.97}$$

where the structural weights $W_{(l,f),(m,g)} = L_{(l,f)}\,\hat{T}_{(l,f)(m,g)}\,R_{(m,g)}$ are character-independent, with $L_{(n,f)} = \sum_i \pi_i \hat{T}^*_{i,(n,f)}$ and $R_{(n,f)} = \sum_j \hat{T}^*_{(n,f),j}$. Substituting the class-mixture emission yields a bilinear sum over both the (domain, fragment-type) latent and the site-class latent:

$$f(a,b) = \sum_{l,f,m,g,c_1,c_2} W_{(l,f),(m,g)} u_{lfc_1} u_{mgc_2}\,\pi^{(c_1)}_a\,\pi^{(c_2)}_b.$$

The order-1 HMM transition is therefore

$$P(b \mid a) = \frac{f(a,b)}{\sum_{b'} f(a,b')} = \frac{\sum_{c_2}\left(\sum_{l,f,m,g,c_1} Wuu\,\pi^{(c_1)}_a\right)\pi^{(c_2)}_b}{\sum_{l,f,c_1}\pi^{(c_1)}_a \sum_{m,g,c_2} W_{(l,f),(m,g)} u_{lfc_1} u_{mgc_2}}.$$

*Remark* C.4 (Non-trivial character correlations). When the per-class equilibria $\pi^{(c)}$ differ, the previous character $a$ carries information about the latent class $c_1$ of the source state via the Bayesian posterior $P(c_1 \mid a) \propto \pi^{(c_1)}_a \sum_{lf,mg,c_2} Wuu$, and consequently about which (domain, fragment-type) is likely to be generating this region. The order-1 HMM therefore has genuinely $a$-dependent transitions: a mixture of $\pi^{(c_2)}$ tilted by the joint posterior over latent state and source class.

*Remark* C.5 (Closed form). All quantities in (C.97) are rational functions of the model parameters: the $\mathcal{NF} \times \mathcal{NF}$ matrix $(I - \hat{T})^{-1}$ has entries that are ratios of polynomials in $(r^{(n)}_{fg}, \kappa_n, \kappa_0, \beta_0, w_{nf}, v_n)$. The class mixture enters linearly through $u_{nfc}$, and the per-class GTRs appear only inside $\pi^{(c)}$, $P^{(c)}(T)$.

##### C.5.4 Pair HMM Distillation

**Emitting state space** The collapsed MixDom Pair HMM with $\mathcal{N}$ domain types and $\mathcal{F}$ fragment types per domain has $5\mathcal{NF}+2$ states: $\{\texttt{SS},\texttt{EE}\}$ and $\{\texttt{MM}_{nf},\texttt{MI}_{nf},\texttt{MD}_{nf},\texttt{II}_{nf},\texttt{DD}_{nf} : n \in \mathcal{N}, f \in \mathcal{F}\}$.

Group the $5\mathcal{NF}$ emitting states by emission type: match $\mathcal{M} = \{\texttt{MM}_{nf}\}$, insert $\mathcal{I} = \{\texttt{MI}_{nf},\texttt{II}_{nf}\}$, delete $\mathcal{D} = \{\texttt{MD}_{nf},\texttt{DD}_{nf}\}$.

**Class-mixture emissions in the Pair HMM** The per-state emissions at $(d,f)$ are the class-mixture emissions $\phi^{\texttt{M,I,D}}_{nf}$ defined in Section C.5.2. Two states with the same $(d,f)$ but different top-level types share the same equilibrium mixture, but the match state combines this with the time-evolved class-mixture transition kernel.

The per-(domain, fragment-type) emission factors do *not* lift cleanly out of the structural sum; however, they have low rank in the class index $c$ (Section C.5.2). Within a fixed $(d,f)$, all match states emit from $\phi^{\texttt{M}}_{nf}$ and all insert/delete singlets from $\phi^{\texttt{I}}_{nf}$ / $\phi^{\texttt{D}}_{nf}$.

**Per-(domain, fragment)-pair structural weights** The pair adjacency frequencies decompose as sums over (domain, fragment-type) pairs. For match-to-match, writing $X=a, Y=b, X'=a', Y'=b'$:

$$f^{\mathcal{MM}}(X,Y,X',Y') = \sum_{(n_1,f_1),(n_2,f_2)} C^{\mathcal{MM}}_{(n_1,f_1),(n_2,f_2)}\, \phi^{\texttt{M}}_{n_1f_1}(X,Y)\, \phi^{\texttt{M}}_{n_2f_2}(X',Y') \tag{C.98}$$

where

$$C^{\mathcal{MM}}_{(n_1,f_1),(n_2,f_2)} = \sum_{s_1\in\mathcal{M}_{n_1f_1}} \sum_{s_2\in\mathcal{M}_{n_2f_2}} L_{s_1}\,\hat{T}_{s_1s_2}\,R_{s_2}$$

are structural weights indexed by (domain, fragment-type) pairs (not just emission types). Similarly for other adjacency types ($\mathcal{MI}$, $\mathcal{MD}$, etc.), with the appropriate per-(domain, fragment-type) emissions.

Each adjacency frequency is therefore a sum of $(\mathcal{NF})^2$ structural-weight terms, each multiplied by per-(domain, fragment-type) emission factors that are themselves rank-$C$ class mixtures.

**Context dependence** In the order-1 WFST, the wait-after-match/insert transition involves

$$\begin{aligned} f^{\cdot\mathcal{M}}(X,Y,a',b') = \sum_{(n_1,f_1),(n_2,f_2)} \Big( & C^{\mathcal{MM}}_{(n_1,f_1),(n_2,f_2)}\, \phi^{\texttt{M}}_{n_1f_1}(X,Y) \\ & + C^{\mathcal{IM}}_{(n_1,f_1),(n_2,f_2)}\, \phi^{\texttt{I}}_{n_1f_1}(Y)\Big)\phi^{\texttt{M}}_{n_2f_2}(a',b') \end{aligned}$$

After normalization, $p_{\texttt{VM}}(X,Y,a',b')$ depends on $(X,Y)$ through the joint posterior over (domain, fragment-type, class) of the previous emission. Specifically, the context enters through the $\mathcal{NF}$-dimensional vector

$$\boldsymbol{\rho}(X,Y) = \Big( \sum_{(n_2,f_2)} C^{\mathcal{MM}}_{(1,1),(n_2,f_2)}\, \phi^{\texttt{M}}_{1,1}(X,Y) + \sum_{(n_2,f_2)} C^{\mathcal{IM}}_{(1,1),(n_2,f_2)}\, \phi^{\texttt{I}}_{1,1}(Y), \dots \Big)$$

which captures the relative likelihood of each source (domain, fragment-type).

*Remark* C.6 (Richer context than per-domain shared-$\pi$ case). With class-mixture emissions, the WFST transition probabilities depend on context $(X, Y)$ through a joint (domain, fragment-type, class) posterior that is sensitive to *both* $X$ and $Y$ independently—not just through a single scalar ratio. The right-side character dependence $(a', b')$ also varies by (domain, fragment-type, class): the next emission is a mixture of $\pi^{(c_2)} P^{(c_2)}(T)$ weighted by the structural weights, the source posterior, and the destination $u$. Despite this richer structure, all quantities are closed-form rational functions of the model parameters (composed with $C$ rate-matrix exponentials).

###### C.5.5 Block Structure and Matrix Inversions

**Top-level null closure (always $3\times 3$)** The null states $\mathtt{A}, \mathtt{B}, \mathtt{C}$ yield a $3 \times 3$ submatrix with entries depending on

$$z_0 = \sum_n \upsilon_n(1 - \kappa_n)$$

$$z_T = \sum_n \upsilon_n(1 - \kappa_n)(1 - \beta_n)$$

The null closure is a $3 \times 3$ inversion with closed-form determinant, independent of $\mathcal{N}, \mathcal{F}, C$ (the dependence on $\mathcal{N}$ enters only through these two scalar sums; $\mathcal{F}$ and $C$ do not appear at all because empty domains are independent of fragment and class structure inside the domain).

**Block-diagonal-plus-low-rank decomposition** The $5\mathcal{N}\mathcal{F} \times 5\mathcal{N}\mathcal{F}$ effective transition matrix between emitting states decomposes as

$$\hat{T}_{\Omega,\Omega} = \underbrace{\mathrm{diag}(D_1, \ldots, D_{\mathcal{N}})}_{\text{intra-domain}} + \underbrace{E\,\mathcal{T}_\bullet\,S^\top}_{\text{inter-domain (rank } \leq 3)}$$

where each $D_n$ is a $5\mathcal{F} \times 5\mathcal{F}$ within-domain block that combines the intra-fragment Markov ext matrix $r^{(n)}_{fg}$ (acting on the fragment-type axis) with the same-domain new-fragment TKF92 transitions ($\rho^{(n)}_f \tau^{(n)}_{\mathtt{XY}} w_{n,g}$ for M-type entries, $\rho^{(n)}_f \kappa_n w_{n,g}$ for I/D-type entries), $E \in \mathbb{R}^{5\mathcal{N}\mathcal{F}\times 3}$ stacks per-(domain,fragment-type) exit vectors projected to top-level types $\{\mathtt{M}, \mathtt{I}, \mathtt{D}\}$ via $(\rho^{(n)}_f\, \varphi_n)$ for the I/D singlet rows and $(\rho^{(n)}_f\, \tau^{(n)}_{\mathtt{XE}})$ for the matched-domain rows, $S \in \mathbb{R}^{5\mathcal{N}\mathcal{F}\times 3}$ stacks per-(domain,fragment-type) start vectors with $(1 - z_T)^{-1} \upsilon_m\, \tau^{(m)}_{\mathtt{SY}} w_{mg}$ for M-type entries and $(1 - z_0)^{-1} \upsilon_m\, \kappa_m\, w_{mg}$ for I/D-type entries (these $1/(1 - z)$ factors are the domenter normalisation that conditions on the destination domain being non-empty), and $\mathcal{T}_\bullet$ is the relevant $5 \times 5$ submatrix of the null-eliminated top-level matrix.

Note that $D_{n_1} \neq D_{n_2}$ in general (different $r^{(n)}_{fg}, w_{nf}, \lambda_n, \mu_n$). For $\mathcal{F} > 1$, $D_n$ has no further emission-type block-diagonal sub-structure within the matched-domain block (the intra-fragment Markov ext couples $\{\mathtt{MM}, \mathtt{MI}, \mathtt{MD}\}$ across fragment-types); however, it admits a Kronecker-plus-low-rank decomposition that preserves the linear-in-$\mathcal{N}\mathcal{F}$ scaling (Section C.5.6). The diagonal-ext baseline (`-freeze-offdiag-ext` in the fitter) recovers the diagonal-extension special case block structure as a special case.

*Remark* C.7 (Domenter normalization). The $\chi$ matrix (Section C.1.1, Equation C.3) includes factors $(1 - z_T)^{-1}$ for M-type destinations and $(1 - z_0)^{-1}$ for I/D-type destinations, conditioning domain entry on the domain being non-empty. These factors cancel the corresponding $(1 - z_T)$ and $(1 - z_0)$ column scaling in $\mathcal{T}_\bullet$ (which arises from the $\upsilon$ null-elimination Schur complement). In the distillation, the start vectors $\mathbf{s}_{n,f}$ carry the domenter factors while $\mathcal{T}_\bullet$ retains the column factors; the product $\mathcal{T}_\bullet \cdot \mathbf{s}_{n,f}$ thus reproduces the correct $\chi$ entries.

**Woodbury identity** Writing $G_n = (I - D_n)^{-1}$ (each $5\mathcal{F} \times 5\mathcal{F}$), $E = (\mathbf{e}_{n,f})$ (exit vectors, $5\mathcal{N}\mathcal{F} \times 3$), $S = (\mathbf{s}_{n,f})$ (start vectors, $5\mathcal{N}\mathcal{F} \times 3$), and $\mathcal{T}_{\text{mid}}$ for the $3 \times 3$ subblock of $\mathcal{T}_\bullet$ restricted to $\{\mathtt{M}, \mathtt{I}, \mathtt{D}\}$ (the only top-level types that couple across domains), the path-sum matrix is

$$(I - \hat{T})^{-1} = G + G\,E\,\mathcal{T}_{\text{mid}}\big(I_3 - S^\top G\,E\,\mathcal{T}_{\text{mid}}\big)^{-1} S^\top G$$

using the push-through form of the Woodbury identity, where $G = \operatorname{diag}(G_1, \ldots, G_\mathcal{N})$. This avoids inverting $\mathcal{T}_{\text{mid}}$, which is singular (the TKF91 `M` and `I` rows are identical, so $\mathcal{T}_{\text{mid}}$ has rank 2).

**Proposition C.1** (Closed-form distillation for MixDom)**.** *All adjacency frequencies, and hence all order-1 HMM and WFST parameters, are closed-form rational functions of the model parameters (composed with the matrix exponentials* $\exp(R^{(c)}T)$*), for any finite* $\mathcal{N}, \mathcal{F}, C$*. The computation requires:*

1. *A* $3 \times 3$ *inversion for the top-level null closure.*

2. $\mathcal{N}$ *within-domain inversions* $(I - D_n)^{-1}$*, each of size* $5\mathcal{F} \times 5\mathcal{F}$*. By the Kronecker-plus-rank-3 decomposition of* $D_n$ *(Section C.5.6), this reduces to one* $\mathcal{F} \times \mathcal{F}$ *inversion of* $(I - r^{(n)})$ *plus a* $3 \times 3$ *inner Woodbury kernel per domain. For* $\mathcal{F} = 1$ *the whole within-domain block collapses to a* $3 \times 3$ *adjugate plus two scalar inversions (Section C.5.7).*

3. *One* $3 \times 3$ *outer Woodbury correction for inter-domain coupling.*

4. *Summation of* $(\mathcal{N}\mathcal{F})^2$ *(domain, fragment-type)-pair terms per adjacency entry, with each term involving a class-mixture emission factor of length* $C$*.*

*Every step is a rational function of the model parameters: no numerical iteration is required to evaluate the closed form.*

*Remark* C.8 (Contrast with shared-emission case)*.* If all $(d, f)$ shared a single class mixture (e.g. $u_{nfc} = u_c$ uniform), the adjacency frequencies would still factor as in (C.98), but the emission tensor would have rank-1 structure across $(d, f)$, collapsing the WFST context dependence. With per-(domain, fragment-type) class mixtures, the structural constants are full $(\mathcal{N}\mathcal{F}) \times (\mathcal{N}\mathcal{F})$ matrices indexed by (domain, fragment-type) pairs, the emission tensors are class-rank-$C$, and the context dependence is genuinely multi-dimensional.

##### C.5.6 Within-Domain Inversion: closed form

For general $\mathcal{F}$, the within-domain block $D_n$ has a Kronecker-plus-rank-3 structure that yields a closed-form inverse without ever inverting a $5\mathcal{F} \times 5\mathcal{F}$ matrix directly.

**Kronecker-plus-rank-3 decomposition of the matched-domain block** The matched-domain inner block (acting on $\{\mathtt{MM}, \mathtt{MI}, \mathtt{MD}\}$ at each fragment-type, total $3\mathcal{F}$ states) combines intra-fragment Markov continuation with fragment termination plus a TKF92 same-domain new-fragment branch:

$$D_n^{\text{match}}[(\mathtt{X}, f), (\mathtt{Y}, g)] = \delta_{\mathtt{XY}}\, r_{fg}^{(n)} + \rho_f^{(n)} \cdot \tau_{\mathtt{XY}}^{(n)} \cdot w_{n,g}. \tag{C.99}$$

The first term is $I_3 \otimes r^{(n)}$ (the intra-fragment Markov chain, identical at every emission type); the second is $\tau_{3\times3}^{(n)} \otimes (\rho^{(n)}\, w_n^\top)$ where $\rho^{(n)}\, w_n^\top$ is the rank-1 $\mathcal{F} \times \mathcal{F}$ outer product of fragment-termination probabilities and fragment-entry weights. The whole rank-3 correction is therefore of rank $\leq 3$ in the joint $(\mathtt{X}, f)$ space (one factor of 3 from $\tau^{(n)}_{3\times 3}$, and rank 1 in the fragment slot from the outer product). Writing

$$D_n^{\mathrm{match}} = I_3 \otimes r^{(n)} + U_n\, V_n^\top, \quad U_n = \tau^{(n)}_{3\times 3} \otimes \rho^{(n)}, \quad V_n = I_3 \otimes w_n,$$

$U_n$ and $V_n$ are $3\mathcal{F} \times 3$ matrices.

**Inverse via inner Woodbury** Since $I_3 \otimes r^{(n)}$ commutes with itself, $I - I_3 \otimes r^{(n)} = I_3 \otimes (I_{\mathcal{F}} - r^{(n)})$, which inverts blockwise: $(I_3 \otimes (I_{\mathcal{F}} - r^{(n)}))^{-1} = I_3 \otimes (I_{\mathcal{F}} - r^{(n)})^{-1}$. Note $(I_{\mathcal{F}} - r^{(n)})^{-1}$ is a single $\mathcal{F} \times \mathcal{F}$ inversion per domain. Applying the Sherman–Morrison–Woodbury identity to the rank-3 correction gives

$$\begin{aligned} (I - D_n^{\mathrm{match}})^{-1} &= G_0 + G_0\, U_n\, K_n^{\mathrm{inner}}\, V_n^\top G_0, \\ G_0 &\equiv I_3 \otimes (I_{\mathcal{F}} - r^{(n)})^{-1}, \\ K_n^{\mathrm{inner}} &\equiv (I_3 - V_n^\top G_0\, U_n)^{-1}. \end{aligned} \tag{C.100}$$

$K_n^{\mathrm{inner}}$ is a $3 \times 3$ matrix whose entries are rational functions of $w_n$, $r^{(n)}$, $\tau^{(n)}$ via a single $\mathcal{F}$-fold inner product $w_n^\top (I - r^{(n)})^{-1} \rho^{(n)}$ times $\tau^{(n)}_{3\times 3}$.

The two singlet-domain blocks act on $\mathcal{F}$ states each ($\{\mathtt{II}_{nf} : f\}$ and $\{\mathtt{DD}_{nf} : f\}$):

$$d_n^{\mathrm{ins}}[f, g] = d_n^{\mathrm{del}}[f, g] = r^{(n)}_{fg} + \rho^{(n)}_f\, \kappa_n\, w_{n,g}.$$

Their inverse follows the same pattern: rank-1 correction to $(I - r^{(n)})$, giving

$$(I - d_n^{\mathrm{ins}})^{-1} = (I - r^{(n)})^{-1} + (I - r^{(n)})^{-1} \rho^{(n)}\, k_n^{\mathrm{ins}}\, w_n^\top (I - r^{(n)})^{-1},$$

with scalar inner kernel $k_n^{\mathrm{ins}} = \kappa_n\, [1 - \kappa_n w_n^\top (I - r^{(n)})^{-1} \rho^{(n)}]^{-1}$, and identically for delete.

Every entry of $(I - D_n)^{-1}$ is therefore a rational function of $(r^{(n)}, w_n, \alpha_n, \beta_n, \gamma_n, \kappa_n)$, computed at cost $O(\mathcal{F}^3)$ per domain (the inversion of $I_{\mathcal{F}} - r^{(n)}$) plus $O(\mathcal{F}^2)$ for the singlet blocks and $O(1)$ for the inner $3 \times 3$ Woodbury kernel. No numerical iteration is required; the inversion can be carried out analytically (for small $\mathcal{F}$, by Cramer's rule on $I_{\mathcal{F}} - r^{(n)}$) or directly evaluated for larger $\mathcal{F}$.

*Remark* C.9 (Compactness). For small $\mathcal{F}$ the inverse $(I_{\mathcal{F}} - r^{(n)})^{-1}$ has a compact symbolic form (e.g. at $\mathcal{F} = 2$ it is a $2 \times 2$ adjugate over a scalar determinant). The further reductions of the matched-domain block via (C.100) and the singlet blocks above involve only $3 \times 3$ and scalar inner inversions, so the overall within-domain inverse can be written down symbolically without ever exceeding a $\mathcal{F} \times \mathcal{F}$ inversion. The $\mathcal{F} = 1$ scalar-extension special case collapses $(I_{\mathcal{F}} - r^{(n)})^{-1}$ to the scalar $1/(1 - r_n)$ and recovers the more compact $3 \times 3$ adjugate of Section C.5.7 below.

##### C.5.7 Within-Domain Inversion: $\mathcal{F} = 1$ closed form

When $\mathcal{F} = 1$ (scalar self-extension $r_n \equiv r^{(n)}_{11}$, no off-diagonal Markov coupling), $D_n$ collapses to $5 \times 5$ with the additional block structure described below; we record this special case because it admits a particularly compact algebraic form.

**Block decomposition:** $3\times3+1+1$ In the $\mathcal{F} = 1$ limit, the five emitting state types $\{\mathtt{MM}, \mathtt{MI}, \mathtt{MD}, \mathtt{II}, \mathtt{DD}\}$ cannot transition between top-level types within a domain: if the domain is inserted (top-level $\mathtt{I}$), all emissions stay in $\mathtt{II}$ until the domain ends; similarly for deleted domains ($\mathtt{DD}$). Therefore $D_n$ is block-diagonal:

$$D_n = \begin{pmatrix} D_n^{\mathrm{match}} & 0 & 0 \\ 0 & d_n^{\mathrm{ins}} & 0 \\ 0 & 0 & d_n^{\mathrm{del}} \end{pmatrix}$$

where $D_n^{\text{match}}$ is $3 \times 3$ (for matched-domain states $\texttt{MM}, \texttt{MI}, \texttt{MD}$) and the singlet-domain states have scalar self-loops:

$$d_n^{\text{ins}} = d_n^{\text{del}} = r_n + (1 - r_n)\kappa_n$$

The scalar inversions are trivial: $(1 - d_n^{\text{ins}})^{-1} = [(1 - r_n)(1 - \kappa_n)]^{-1}$.

**The $3 \times 3$ matched-domain block** The matched-domain block combines fragment extension (self-loop at rate $r_n$) with intra-domain TKF transitions:

$$D_n^{\text{match}} = r_n I_3 + (1 - r_n)\, \tau^{(n)}_{\texttt{MID},\texttt{MID}}$$

where $\tau^{(n)}_{\texttt{MID},\texttt{MID}}$ is the $3 \times 3$ submatrix of $\tau^{(n)}$ restricted to rows and columns $\texttt{M}, \texttt{I}, \texttt{D}$.

A key property: in the TKF transition matrix, rows $\texttt{M}$ and $\texttt{I}$ are identical. Defining shorthand (suppressing domain subscript $n$):

$$\begin{aligned} \mathfrak{a} &= (1 - \beta)\kappa\alpha, & \mathfrak{b} &= \beta, & \mathfrak{c} &= (1 - \beta)\kappa(1 - \alpha) \\ \mathfrak{d} &= (1 - \gamma)\kappa\alpha, & \mathfrak{g} &= \gamma, & \mathfrak{h} &= (1 - \gamma)\kappa(1 - \alpha) \end{aligned}$$

the $3 \times 3$ TKF submatrix is

$$\tau^{(n)}_{\texttt{MID},\texttt{MID}} = \begin{pmatrix} \mathfrak{a} & \mathfrak{b} & \mathfrak{c} \\ \mathfrak{a} & \mathfrak{b} & \mathfrak{c} \\ \mathfrak{d} & \mathfrak{g} & \mathfrak{h} \end{pmatrix}$$

with row sums $\mathfrak{a} + \mathfrak{b} + \mathfrak{c} = (1 - \beta)\kappa + \beta$ for the first two rows and $\mathfrak{d} + \mathfrak{g} + \mathfrak{h} = (1 - \gamma)\kappa + \gamma$ for the third.

**Factoring out the extension rate** Since $I - D^{\text{match}} = I - rI - (1 - r)\tau_{3\times3} = (1 - r)(I - \tau_{3\times3})$, the extension rate factors out as a scalar:

$$(I - D_n^{\text{match}})^{-1} = \frac{1}{1 - r_n} P_n^{-1}$$

where

$$P_n \equiv I - \tau^{(n)}_{\texttt{MID},\texttt{MID}} = \begin{pmatrix} 1 - \mathfrak{a} & -\mathfrak{b} & -\mathfrak{c} \\ -\mathfrak{a} & 1 - \mathfrak{b} & -\mathfrak{c} \\ -\mathfrak{d} & -\mathfrak{g} & 1 - \mathfrak{h} \end{pmatrix}$$

with row sums $(1 - \beta)(1 - \kappa)$, $(1 - \beta)(1 - \kappa)$, $(1 - \gamma)(1 - \kappa)$ (the domain-exit probabilities from each inner state).

**Determinant of $P_n$** Since rows 1 and 2 of $\tau_{3\times3}$ are identical, subtracting row 2 from row 1 in $P$ yields $(1, -1, 0)$. Expanding the determinant along this simplified row gives

$$\begin{aligned} \det(P_n) &= (1 - \mathfrak{b})(1 - \mathfrak{h}) - \mathfrak{c}\mathfrak{g} - \mathfrak{a}(1 - \mathfrak{h}) - \mathfrak{c}\mathfrak{d} \\ &= (1 - \beta_n)(1 - \kappa_n) \end{aligned}$$

This factors cleanly: since $\mathfrak{a} + \mathfrak{b} + \mathfrak{c}$ and $\mathfrak{d} + \mathfrak{g} + \mathfrak{h}$ are the row sums of $\tau_{3\times3}$, we have

$$\boxed{\det(P_n) = (1 - \beta_n)(1 - \kappa_n)}$$

The determinant is the product of the M/I-row exit probability $(1-\beta)$ and the stationary emptiness probability $(1 - \kappa)$.

**Inverse of $P_n$** The adjugate $\mathrm{adj}(P_n)$ has entries (again suppressing domain subscripts):

$$\mathrm{adj}(P) = \begin{pmatrix} (1-\beta)(1-\kappa(1-\alpha)) & \beta+\kappa(1-\alpha)(\gamma-\beta) & (1-\beta)\kappa(1-\alpha) \\ (1-\beta)\kappa\alpha & 1-(1-\beta)\kappa\alpha-(1-\gamma)\kappa(1-\alpha) & (1-\beta)\kappa(1-\alpha) \\ (1-\beta)\kappa\alpha & \gamma+\kappa\alpha(\beta-\gamma) & (1-\beta)(1-\kappa\alpha) \end{pmatrix}$$

Note the symmetries: $\mathrm{adj}(P)_{21} = \mathrm{adj}(P)_{31}$ and $\mathrm{adj}(P)_{13} = \mathrm{adj}(P)_{23}$ (reflecting the identical rows of $\tau_{3\times 3}$), while $\mathrm{adj}(P)_{11} - \mathrm{adj}(P)_{33} = (1-\beta)\kappa(2\alpha-1)$, which vanishes only when $\alpha = \frac{1}{2}$.

The full inverse is

$$P_n^{-1} = \frac{\mathrm{adj}(P_n)}{(1-\beta_n)(1-\kappa_n)}$$

and therefore

$$(I - D_n^{\mathrm{match}})^{-1} = \frac{\mathrm{adj}(P_n)}{(1-r_n)(1-\beta_n)(1-\kappa_n)}$$

*Remark* C.10 (Compactness in the $\mathcal{F} = 1$ limit). Each entry of $(I - D_n^{\mathrm{match}})^{-1}$ is a ratio of a polynomial with 1–3 terms (numerator, from the adjugate) over a polynomial with 2–3 terms (denominator $(1-r)(1-\beta)(1-\kappa)$). These are genuinely compact closed-form expressions in the TKF parameters $(\alpha_n, \beta_n, \gamma_n, \kappa_n, r_n)$. For $\mathcal{F} > 1$ the inversion is still closed-form (a rational function in $r^{(n)}, w_n, \alpha_n, \beta_n, \gamma_n, \kappa_n$), via the Kronecker-plus-rank-3 Woodbury decomposition of Section C.5.6: the only inversion of dimension exceeding $3 \times 3$ is a single $\mathcal{F} \times \mathcal{F}$ inverse $(I_{\mathcal{F}} - r^{(n)})^{-1}$ per domain, which can itself be written symbolically via the matrix adjugate.

##### C.5.8 Bilinear Factored Form of Adjacency Frequencies

The Woodbury identity gives the full path-sum matrix in a form that makes the adjacency computation practical for any $\mathcal{N}, \mathcal{F}$.

**Structure of the path-sum matrix** Writing $G_n = (I - D_n)^{-1}$ ($5\mathcal{F} \times 5\mathcal{F}$ per domain), the Woodbury expansion gives, for states $s_1$ in $(n_1, f_1)$ and $s_2$ in $(n_2, f_2)$:

$$[(I - \hat{T})^{-1}]_{s_1 s_2} = \delta_{n_1 n_2} [G_{n_1}]_{s_1 s_2} + [G_{n_1} \mathbf{e}_{n_1}]_{s_1}^{\top} \mathcal{T}_{\mathrm{mid}} K \, [\mathbf{s}_{n_2}^{\top} G_{n_2}]_{s_2}$$

where $K = \left(I_3 - \sum_n \mathbf{s}_n^{\top} G_n \mathbf{e}_n \mathcal{T}_{\mathrm{mid}}\right)^{-1}$ is the $3 \times 3$ Woodbury kernel (push-through form), computed once.

**Bilinear form of structural weights** The structural weight $C^{\alpha\beta}_{(n_1,f_1),(n_2,f_2)}$ decomposes as a within-domain diagonal plus a cross-domain bilinear form:

$$C^{\alpha\beta}_{(n_1,f_1),(n_2,f_2)} = \delta_{n_1 n_2} c^{\alpha}_{n_1 f_1 f_2} + \mathbf{a}^{\alpha\top}_{n_1 f_1} M^{-1} \mathbf{b}^{\beta}_{n_2 f_2}$$

where $\mathbf{a}^{\alpha}_{nf}, \mathbf{b}^{\beta}_{nf} \in \mathbb{R}^5$ are per-(domain, fragment-type) vectors derived from $G_n$, $\mathbf{e}_n$, $\mathbf{s}_n$, and the emission-type projections, and $c^{\alpha}_{nf_1 f_2}$ is the within-domain entry of $G_n$ between fragment-types $f_1$ and $f_2$ at emission type $\alpha$ (a $\mathcal{F} \times \mathcal{F}$ matrix per domain rather than a scalar).

**Closed-form adjacency formula** Substituting into the adjacency frequency (C.98) and using the per-(domain, fragment-type) class-mixture emissions $\phi^{\alpha}_{nf}$ from Section C.5.2, the full adjacency table takes the form:

$$f^{\alpha\beta}(\text{chars}_L, \text{chars}_R) = \underbrace{\left[\sum_{n,f} \mathbf{a}^{\alpha}_{nf}\, \phi^{\alpha}_{nf}(\text{chars}_L)\right]^{\top} M^{-1} \left[\sum_{n,f} \mathbf{b}^{\beta}_{nf}\, \phi^{\beta}_{nf}(\text{chars}_R)\right]}_{\text{cross-domain: bilinear in two 5-vectors}} + \underbrace{\sum_{n} \sum_{f_1,f_2} c^{\alpha}_{nf_1 f_2}\, \phi^{\alpha}_{nf_1}(\text{chars}_L)\, \phi^{\beta}_{nf_2}(\text{char}\text{[illegible]}}_{\text{within-domain: } \mathcal{F}\times\mathcal{F} \text{ inner sum [illegible]}} \tag{C.101}$$

This is algebraic closed form: every quantity appearing—$G_n$ entries, $\mathbf{a}^{\alpha}_{nf}$, $\mathbf{b}^{\beta}_{nf}$, $c^{\alpha}_{nf_1f_2}$, $M^{-1}$, and $\phi^{\alpha}_{nf}$—is a rational function of the model parameters (composed with the matrix exponentials $\exp(R^{(c)}T)$). All inversions reduce to closed-form rational expressions: the only matrix inversion of dimension exceeding $3 \times 3$ in the entire distillation pipeline is $(I_{\mathcal{F}} - r^{(n)})^{-1}$ per domain (Section C.5.6), which is a single $\mathcal{F} \times \mathcal{F}$ adjugate-over-determinant. The Woodbury correction $M^{-1}$ is a single $3 \times 3$ inversion regardless of $\mathcal{N}, \mathcal{F}, C$.

*Remark* C.11 (Practical computation for $\mathcal{N} = \mathcal{F} = C = 3$). For $\mathcal{N} = 3, \mathcal{F} = 3, C = 3$:

1. Precompute $C = 3$ per-class transition kernels $\exp(R^{(c)}T)$.
2. Precompute 3 within-domain inverses $G_n$ (each $15 \times 15$, i.e. $5\mathcal{F} = 15$).
3. Precompute 9 pairs $(\mathbf{a}^{\alpha}_{nf}, \mathbf{b}^{\beta}_{nf})$ for each $(\alpha, \beta)$ pair, plus the within-domain $\mathcal{F} \times \mathcal{F}$ matrices $c^{\alpha}_{nf_1f_2}$.
4. Accumulate $\Sigma = \sum_n \mathbf{s}_n^{\top} G_n \mathbf{e}_n$ and compute the push-through kernel $K$ (one $3 \times 3$ inversion).
5. For each character entry: evaluate (C.101) by summing 9 scaled 3-vectors for the left and right factors of the cross-domain term, then a $3 \times 3$ bilinear product, plus a within-domain double sum over $(f_1, f_2)$.

The per-entry cost is $O(\mathcal{N}\mathcal{F} + C)$ multiplications.

##### C.5.9 Full-Context Distillation: Passthrough Context for Insert and Delete

In a two-tape transducer, the *state* at any point should encode the most recent character on *each* tape:

- After Match$(X, Y)$: context is $(X, Y)$ — both tapes updated.
- After Insert emitting $Y'$: context is $(X, Y')$ — ancestor context $X$ unchanged (passthrough from prior Match or Delete), descendant updated to $Y'$.
- After Delete consuming $X'$: context is $(X', Y)$ — descendant context $Y$ unchanged (passthrough from prior Match or Insert), ancestor updated to $X'$.

The adjacency frequencies in Section C.5 track only partial context for Insert and Delete states: $\phi^{\mathtt{I}}_{nf}(Y) = \sum_c u_{nfc}\pi^{(c)}_Y$ depends only on the descendant character, and $\phi^{\mathtt{D}}_{nf}(X) = \sum_c u_{nfc}\pi^{(c)}_X$ depends only on the ancestor character. This loses information about which (domain, fragment-type, class) generated the passthrough context.

We now show that the full-context adjacency frequencies—with both $(X, Y)$ tracked in all states—can be computed in closed algebraic form, preserving the Woodbury structure.

**Insert chain Green's function** Define the *Insert chain* as the sub-process restricted to Insert states $\mathcal{I} = \{\mathtt{MI}_{nf}, \mathtt{II}_{nf} : n \in \mathcal{N}, f \in \mathcal{F}\}$. The restricted transition matrix $T_{\text{eff}}^{\mathcal{II}}$ (Insert→Insert transitions within $\hat{T}$) has the same block-diagonal-plus-low-rank structure as the full $\hat{T}$:

$$T_{\text{eff}}^{\mathcal{II}} = D^{II} + E^{II}\,\mathcal{T}_{\text{mid}}^{II}\,S^{II\top}$$

where $D^{II} = \text{diag}(D_1^{II}, \ldots, D_{\mathcal{N}}^{II})$ with each $D_n^{II}$ being the within-domain Insert self-loop block ($2\mathcal{F} \times 2\mathcal{F}$ for $\{\mathtt{MI}_{nf}, \mathtt{II}_{nf}\}$ over fragment-types), and the cross-domain coupling has rank $\leq 2$ through the Insert rows of $\mathcal{T}_\bullet$.

The Insert chain Green's function is therefore

$$G^{II} = (I - T_{\text{eff}}^{\mathcal{II}})^{-1}$$

computable via Woodbury with a kernel of dimension $\leq 2$. Each within-domain inverse $(I - D_n^{II})^{-1}$ is a $2\mathcal{F} \times 2\mathcal{F}$ inversion — simpler than the $5\mathcal{F} \times 5\mathcal{F}$ within-domain block of the full system.

An analogous *Delete chain Green's function* $G^{DD} = (I - T_{\text{eff}}^{\mathcal{DD}})^{-1}$ handles the descendant passthrough through Delete states, with identical structure.

**Ancestor-conditioned structural weights** To enter the Insert chain with ancestor context $X$, the process must have come from a prior Match or Delete state in some $(n_0, f_0)$ that emitted ancestor $X$. The match emission satisfies $\sum_Y \phi_{nf}^{\mathtt{M}}(X,Y) = \phi_{nf}^{\mathtt{D}}(X)$ (row-stochastic per ancestor), so the entry weight from $(n_0, f_0)$ with ancestor $X$ is proportional to $\phi_{n_0 f_0}^{\mathtt{D}}(X)$ for both Match and Delete source states.

Define the ancestor-conditioned left vector:

$$\tilde{L}_{n_1 f_1}^{\mathcal{I}}(X) = \sum_{n_0, f_0} \underbrace{(L_{\mathtt{MM}_{n_0 f_0}} + L_{\mathtt{MD}_{n_0 f_0}} + L_{\mathtt{DD}_{n_0 f_0}})}_{\lambda_{n_0 f_0}} \cdot \phi_{n_0 f_0}^{\mathtt{D}}(X) \cdot \left[\hat{T}_{\text{entry}}\, G^{II}\right]_{(n_0,f_0),(n_1,f_1)}$$

where $L_s$ are the standard left boundary weights from the full path-sum computation, $\lambda_{n_0 f_0}$ collects all ancestor-emitting (Match and Delete) boundary contributions for $(n_0, f_0)$, $\hat{T}_{\text{entry}}$ is the entry transition from ancestor-emitting states $\{\mathtt{MM}, \mathtt{MD}, \mathtt{DD}\}$ into the Insert chain (i.e. the $\{M, D\} \to \{I\}$ block of $\hat{T}$), and $G_{(n_0,f_0),(n_1,f_1)}^{II}$ sums over all Insert-chain continuations.

Key observation: $\phi_{n_0 f_0}^{\mathtt{D}}(X)$ factors out multiplicatively from the structural sum, preserving the bilinear structure.

**Full-context adjacency for Insert-sourced transitions** The full-context adjacency for Insert→Match is:

$$f_{\text{full}}^{\mathcal{IM}}(X,Y,X',Y') = \sum_{(n_1,f_1),(n_2,f_2)} \tilde{L}_{n_1 f_1}^{\mathcal{I}}(X) \cdot \eta_{(n_1,f_1),(n_2,f_2)}^{\mathcal{IM}} \cdot \phi_{n_1 f_1}^{\mathtt{I}}(Y) \cdot \phi_{n_2 f_2}^{\mathtt{M}}(X',Y') \tag{C.102}$$

where $\eta_{(n_1,f_1),(n_2,f_2)}^{\mathcal{IM}} = T_{I_{n_1 f_1}, M_{n_2 f_2}} \cdot R_{M_{n_2 f_2}}$ and $Y$ is the descendant character at the Insert, $X$ the passthrough ancestor.

The bilinear factored form (cf. (C.101)) generalizes to:

$$f_{\text{full}}^{\mathcal{IM}}(X,Y,X',Y') = \left[\sum_{n,f} \tilde{\mathbf{a}}_{nf}^{\mathcal{I}}(X)\,\phi_{nf}^{\mathtt{I}}(Y)\right]^\top K_I \left[\sum_{n,f} \mathbf{b}_{nf}^{\mathcal{M}}\,\phi_{nf}^{\mathtt{M}}(X',Y')\right] + \sum_{n,f_1,f_2} (\text{diagonal terms}) \tag{C.103}$$

where $\tilde{\mathbf{a}}^{\mathcal{I}}_{nf}(X)$ are modified per-(domain, fragment-type) left vectors incorporating the ancestor context through $\tilde{L}^{\mathcal{I}}_{nf}(X)$, and $K_I$ is the Woodbury kernel for the Insert-chain correction.

Similarly, for Insert→Insert with ancestor passthrough:

$$f^{\mathcal{II}}_{\text{full}}(X, Y, X, Y') = \sum_{(n_1,f_1),(n_2,f_2)} \tilde{L}^{\mathcal{I}}_{n_1 f_1}(X) \cdot \eta^{\mathcal{II}}_{(n_1,f_1),(n_2,f_2)} \cdot \phi^{\mathtt{I}}_{n_1 f_1}(Y) \cdot \phi^{\mathtt{I}}_{n_2 f_2}(Y')$$

where the ancestor $X$ is preserved on both sides (3 effective character dimensions, not 4).

By symmetry, the Delete-sourced adjacencies with descendant passthrough are:

$$f^{\mathcal{DM}}_{\text{full}}(X, Y, X', Y') = \sum_{(n_1,f_1),(n_2,f_2)} \tilde{L}^{\mathcal{D}}_{n_1 f_1}(Y) \cdot \eta^{\mathcal{DM}}_{(n_1,f_1),(n_2,f_2)} \cdot \phi^{\mathtt{D}}_{n_1 f_1}(X) \cdot \phi^{\mathtt{M}}_{n_2 f_2}(X', Y') \tag{C.104}$$

where $\tilde{L}^{\mathcal{D}}_{n_1 f_1}(Y)$ is the descendant-conditioned left vector for the Delete chain, defined analogously with $G^{DD}$.

**Computational cost** The affected adjacency tables gain one character dimension:

| Tensor | Original | Full-context | Per-entry cost |
|---|---|---|---|
| $f^{\mathcal{MM}}$ | $\lvert\mathcal{A}\rvert^4$ | $\lvert\mathcal{A}\rvert^4$ (unchanged) | $O(\mathcal{NF} + C)$ |
| $f^{\mathcal{MI}}$ | $\lvert\mathcal{A}\rvert^3$ | $\lvert\mathcal{A}\rvert^3$ (unchanged) | $O(\mathcal{NF} + C)$ |
| $f^{\mathcal{MD}}$ | $\lvert\mathcal{A}\rvert^3$ | $\lvert\mathcal{A}\rvert^3$ (unchanged) | $O(\mathcal{NF} + C)$ |
| $f^{\mathcal{IM}}$ | $\lvert\mathcal{A}\rvert^3$ | $\lvert\mathcal{A}\rvert^4$ | $O(\mathcal{NF} + C)$ |
| $f^{\mathcal{II}}$ | $\lvert\mathcal{A}\rvert^2$ | $\lvert\mathcal{A}\rvert^3$ ($X$ preserved) | $O(\mathcal{NF} + C)$ |
| $f^{\mathcal{ID}}$ | $\lvert\mathcal{A}\rvert^2$ | $\lvert\mathcal{A}\rvert^3$ ($X$ preserved, $Y \to X'$) | $O(\mathcal{NF} + C)$ |
| $f^{\mathcal{DM}}$ | $\lvert\mathcal{A}\rvert^3$ | $\lvert\mathcal{A}\rvert^4$ | $O(\mathcal{NF} + C)$ |
| $f^{\mathcal{DD}}$ | $\lvert\mathcal{A}\rvert^2$ | $\lvert\mathcal{A}\rvert^3$ ($Y$ preserved) | $O(\mathcal{NF} + C)$ |
| $f^{\mathcal{DI}}$ | $\lvert\mathcal{A}\rvert^2$ | $\lvert\mathcal{A}\rvert^3$ ($Y$ preserved, $X \to Y'$) | $O(\mathcal{NF} + C)$ |

The per-entry cost is $O(\mathcal{NF} + C)$ after Woodbury factoring (cf. the same per-entry cost for $f^{\mathcal{MM}}$ noted in Remark on p. 125), since the Insert and Delete chain Green's functions have the same bilinear decomposition. The total cost for the full adjacency table is $O((\mathcal{NF} + C) \cdot |\mathcal{A}|^4)$ — the same asymptotic scaling as the existing match-to-match computation. The Woodbury kernel remains $3 \times 3$ for the full system; the Insert and Delete chain kernels are $\leq 2 \times 2$. All quantities remain closed-form rational functions of the model parameters (composed with the $C$ rate-matrix exponentials).

*Remark* C.12 (Why the original formulation lost context). The emission functions $\phi^{\mathtt{I}}_{nf}(Y)$ and $\phi^{\mathtt{D}}_{nf}(X)$ are genuinely single-character: the Insert emission does not depend on the ancestor, and vice versa. The lost context is not an emission effect but a *transition routing* effect: the joint posterior $P((n, f, c) \mid X, Y)$ depends on both characters, so the passthrough character informs which (domain, fragment-type, class) we are in, and hence which transition probabilities apply. The correction above recovers this information by conditioning the structural weights on the passthrough character.

##### C.5.10 Domains versus Fragments versus Classes for Adjacency Capture

The order-1 WFST has $O(|\mathcal{A}|^4)$ free parameters (from match-to-match transition weights parameterized by context $(X, Y)$ and next pair $(a', b')$), while the MixDom model has far fewer. We analyse which type of model complexity most efficiently captures adjacency structure.

**Adjacency tensor rank** The match-to-match adjacency table $f^{\mathcal{MM}}(X, Y, X', Y')$ is a sum of $(\mathcal{NF})^2$ structural-weight terms (in the (domain, fragment-type)-pair sense), each weighted by a rank-$C$ emission factor on each side. To approach saturating the WFST capacity ($|\mathcal{A}|^4$ entries), the tensor rank must approach $|\mathcal{A}|^2$; this is achievable through a combination of $(\mathcal{N}, \mathcal{F}, C)$.

**Domains: independent TKF rates and weights** Each domain type $n$ brings its own $(\alpha_n, \beta_n, \gamma_n, \kappa_n)$, creating genuinely different:

- $\mathtt{M} \to \mathtt{D}$ vs. $\mathtt{M} \to \mathtt{I}$ ratios (different $\kappa_n(1 - \alpha_n)$ vs. $\beta_n$),
- `V` vs. `W` transition behaviour (different $\beta_n$ vs. $\gamma_n$),
- overall indel/match balance.

These features drive cross-type adjacency diversity in the WFST and are not reproducible by fragments or classes alone.

**Fragments: vary intra-fragment transition pattern as well as boundary frequency** The intra-fragment Markov ext matrix $r^{(n)}_{fg}$ allows transitions *between fragment-types within a single fragment*, without invoking the TKF92 new-fragment branch. These intra-fragment $f \to g$ transitions at each emitted site can carry persistent emission-class context: a chain that prefers one fragment-type tends to stay in that fragment-type's class mixture, producing richer M/I/D adjacency patterns than fragments without intra-fragment correlation. The off-diagonal entries of $r^{(n)}$ thus contribute genuine new `M/I/D` adjacency content; setting them to zero (`-freeze-offdiag-ext` in the fitter) recovers the diagonal-extension special case behaviour where fragments only modulate boundary frequency, and the $\mathcal{F} = 1$ case collapses fragments to a scalar self-extension per domain.

**Classes: emission-mixture diversity, decoupled from structure** Each site class $c$ has its own GTR $(Q^{(c)}, \pi^{(c)})$. Classes contribute *emission* diversity: the rank-$C$ class-mixture emission $\phi^{\mathtt{M,I,D}}_{nf}$ allows the joint match emission tensor to deviate from a single GTR's product structure, even within a single $(d, f)$. Classes do not contribute to the `M/I/D` transition pattern (they do not appear in $\hat{T}$ or in any of the structural weights $C^{\alpha\beta}$); they affect only the character-side of each adjacency frequency. Sharing the class pool across $(d, f)$ via $u$ gives a low-parameter way to enrich the emission tensor without growing the latent-state space.

*Remark* C.13 (Parameter allocation). For adjacency capture in MixDom:

- Domains are essential to enrich the `M/I/D` transition pattern via independent TKF rates.
- Fragments enrich the `M/I/D` structure (via the intra-fragment Markov ext) and the boundary frequency (via $\rho^{(n)}_f$); they also support an additional layer of emission diversity through $u_{nfc}$.
- Classes enrich emissions only, but cheaply (decoupled from $\mathcal{N}, \mathcal{F}$): with $C$ classes, every $(d, f)$ has access to the full pool via its Dirichlet $u_{nfc}$.

The total adjacency-tensor rank scales as $\mathcal{NF}$ (structural-weight pairs) $\times C^2$ (left/right class mixture); in practice, allocating the parameter budget across all three axes gives the most efficient match to the WFST capacity.

##### C.5.11 Identifiability

Are the MixDom parameters recoverable from the distilled order-1 models?

**Generic identifiability (up to standard ambiguities).** The distilled WFST provides $O(|\mathcal{A}|^4)$ constraints on the MixDom parameters. With $\mathcal{N}$ domains, $\mathcal{F}$ fragment types per domain, and $C$ site classes, there are $2 + (\mathcal{N} - 1) + 2\mathcal{N} + \mathcal{N}\mathcal{F}^2 + \mathcal{N}\mathcal{F}(C-1) + C(|\mathcal{A}|^2 + |\mathcal{A}| - 2)$ free parameters (the last term accounts for $C$ symmetric exchangeability matrices and equilibrium distributions). The system is heavily over-determined for moderate $(\mathcal{N}, \mathcal{F}, C)$.

The per-domain TKF parameters $(\lambda_n T, \mu_n T, r_{fg}^{(n)})$ are recoverable from the within-(domain, fragment-type) `M/I/D` transition structure: $\alpha_n$ and $\kappa_n$ determine $\mu_n T$ and $\lambda_n/\mu_n$; $\beta_n$ provides a redundant check; $r^{(n)}$ separates from $\kappa_n$ because intra-fragment Markov transitions decompose differently from new-fragment transitions (see Section C.3.5 of the Maraschino fitter and Baum–Welch derivation). The top-level parameters $(\lambda_0 T, \mu_0 T)$ are identifiable from the inter-domain transition pattern. The per-class GTRs $(Q^{(c)}, \pi^{(c)})$ and per-(domain, fragment-type) $u_{nfc}$ are jointly identifiable up to label permutation of classes, because the WFST context dependence reveals the mixture components as the joint posterior shifts across different $(X, Y)$ contexts.

**Unavoidable ambiguities.**

1. **Label permutation**: permuting domain labels gives the same model ($\mathcal{N}!$-fold), permuting fragment-type labels at fixed domain gives the same model ($\mathcal{F}!^{\mathcal{N}}$-fold), and permuting class labels gives the same model ($C!$-fold).
2. **Rate-time confounding**: only products $\lambda T, \mu T$ are identifiable from pairwise data (a single evolutionary time).
3. **Class-domain mixing**: when $C \geq \mathcal{N}\mathcal{F}$, the class structure can absorb domain-level emission differences, creating identifiability issues unless the structural weights $C^{\alpha\beta}$ resolve them.

**Lossy in distribution, injective in parameters.** The distilled model captures only pairwise-adjacent correlations; the full MixDom has higher-order structure (e.g., runs of characters from the same fragment, intra-fragment Markov correlations between fragment-types). The distillation map MixDom $\to$ order-1 is therefore *lossy* for the sequence distribution but generically *injective* for the parameters: one can recover the MixDom parameters from the distilled model, even though the distilled model cannot reproduce all statistics of the MixDom.

##### C.5.12 Scaling to $\mathcal{N}, \mathcal{F}, C$

**The top-level null closure does not grow with $\mathcal{N}, \mathcal{F}, C$** The null states `A`, `B`, `C` are always exactly three, regardless of the latent-state cardinalities. Their transition submatrix depends on $\mathcal{N}$ only through the scalars $z_0 = \sum_n v_n(1-\kappa_n)$ and $z_T = \sum_n v_n(1-\kappa_n)(1-\beta_n)$; $\mathcal{F}$ and $C$ do not appear at this top level. The null closure and the effective $5 \times 5$ top-level matrix $\mathcal{T}$ are $O(\mathcal{N})$ to compute but $O(1)$ in matrix dimension.

**Domain types are drawn i.i.d., not as a Markov chain** When a new domain begins, its type is drawn independently from $v$, regardless of the previous domain's type. This i.i.d. structure means the cross-domain transitions factor as $\hat{T}(s, s') = e_l(s) \times \mathcal{T}_\bullet \times s_m(s')$ (exit $\times$ top-level $\times$ start), and the cross-domain contribution to the full $5\mathcal{N}\mathcal{F} \times 5\mathcal{N}\mathcal{F}$ transition matrix has rank at most 3. Note that this i.i.d. property holds at the *domain* level only; within a fragment, the fragment-type process is a Markov chain on $\mathcal{F}$ states (see Section C.1.1, Equation C.3), which is encoded in the within-domain block $D_n$ and does not break the cross-domain factorisation.

**Woodbury for general $\mathcal{N}, \mathcal{F}$** The emitting state space has $5\mathcal{N}\mathcal{F}$ states. The effective transition matrix decomposes as

$$\hat{T} = D + E\,\mathcal{T}_{\bullet}\,S^{\top}$$

where $D = \text{diag}(D_1, \dots, D_{\mathcal{N}})$ with each $D_n$ a $5\mathcal{F} \times 5\mathcal{F}$ within-domain block (now domain-specific and, for $\mathcal{F} > 1$, with intra-fragment Markov coupling), $E \in \mathbb{R}^{5\mathcal{N}\mathcal{F}\times 3}$ stacks exit vectors projected to $\{\mathtt{M}, \mathtt{I}, \mathtt{D}\}$, and $S \in \mathbb{R}^{5\mathcal{N}\mathcal{F}\times 3}$ stacks start vectors projected to $\{\mathtt{M}, \mathtt{I}, \mathtt{D}\}$.

The Woodbury identity gives $(I - \hat{T})^{-1}$ via:

1. $\mathcal{N}$ independent $5\mathcal{F}\times 5\mathcal{F}$ inversions $(I - D_n)^{-1}$, each evaluated in closed form via the Kronecker-plus-rank-3 inner Woodbury of Section C.5.6: one $\mathcal{F} \times \mathcal{F}$ adjugate $(I_{\mathcal{F}} - r^{(n)})^{-1}$ plus a $3 \times 3$ inner kernel per domain. For $\mathcal{F} = 1$ this further collapses to a $3 \times 3$ adjugate (closed-form determinant $(1 - r)(1 - \beta)(1 - \kappa)$) plus two scalar inversions (Section C.5.7). The total cost is $O(\mathcal{N}\mathcal{F}^3)$ per evaluation.
2. Computing $\sum_n \mathbf{s}_n^{\top} G_n\, \mathbf{e}_n$, a $3 \times 3$ matrix built by summing $\mathcal{N}$ contributions: $O(\mathcal{N}\mathcal{F}^2)$ work.
3. One $3 \times 3$ inversion for the Woodbury correction: $O(1)$ work.

**Proposition C.2** (Linear-in-$\mathcal{N}$, cubic-in-$\mathcal{F}$, linear-in-$C$ scaling)**.** *The order-1 distillation computation scales as $O(\mathcal{N}\mathcal{F}^3)$ in within-domain inversions and $O(C)$ in per-class GTR exponentials:*

- *$O(\mathcal{N})$ to compute $z_0, z_T$ and the $5 \times 5$ top-level matrix.*
- *$O(\mathcal{N}\mathcal{F}^3)$ for within-domain path sums.*
- *$O(\mathcal{N}\mathcal{F}^2)$ to accumulate the $3 \times 3$ Woodbury correction.*
- *$O((\mathcal{N}\mathcal{F})^2)$ (domain, fragment-type)-pair terms per adjacency entry (or $O((\mathcal{N}\mathcal{F})^2 |\mathcal{A}|^d \cdot C)$ for the full adjacency table, with $d \in \{2, 3, 4\}$ character dimensions depending on state type; see Section C.5.9).*
- *$O(C|\mathcal{A}|^3)$ for per-class transition kernels $\exp(R^{(c)}T)$.*

*The Woodbury kernel is always $3 \times 3$, regardless of the latent state cardinalities.*

##### C.5.13 Summary

| Component | Matrix size | Scaling |
|---|---|---|
| Top-level null closure | $3 \times 3$ | $O(1)$ |
| Top-level eff. matrix | $5 \times 5$ | $O(\mathcal{N})$ to compute |
| Within-domain inversions | $5\mathcal{F} \times 5\mathcal{F}$ each | $O(\mathcal{N}\mathcal{F}^3)$ |
| Woodbury correction | $3 \times 3$ | $O(\mathcal{N}\mathcal{F}^2)$ to accumulate |
| Per-class GTR exponentials | $\lvert\mathcal{A}\rvert \times \lvert\mathcal{A}\rvert$ | $O(C\lvert\mathcal{A}\rvert^3)$ |
| (Domain, fragment-type)-pair adjacency terms | — | $O((\mathcal{N}\mathcal{F})^2 C)$ per entry |

The distillation is closed-form for any finite $\mathcal{N}, \mathcal{F}, C$: all quantities are rational functions of the model parameters (composed with $C$ exponentials of rate-times-time products). The key structural features enabling this are:

1. **Top-level null closure is $O(1)$ in dimension.** The MixDom null states $\mathtt{A}, \mathtt{B}, \mathtt{C}$ are always exactly three, regardless of $\mathcal{N}, \mathcal{F}, C$. All dependence on $\mathcal{N}$ enters through the scalars $z_0, z_T$.

2. **Cross-domain transitions factor.** The exit-vector × top-level × start-vector structure caps the rank of the inter-domain coupling at 3 (the $\mathtt{M}, \mathtt{I}, \mathtt{D}$ top-level states that couple across domains), enabling Woodbury reduction to a fixed $3 \times 3$ inversion.

3. **Domain types are i.i.d., not Markov.** If domain type depended on the previous domain (a Markov chain on domain types), the cross-domain block would lose its factored structure and the Woodbury reduction would fail—the full $5\mathcal{NF} \times 5\mathcal{NF}$ inversion would be required, scaling as $O((\mathcal{NF})^3)$. The domain-level i.i.d. mixture is what keeps the cross-domain part linear in $\mathcal{N}$. The intra-fragment Markov chain on fragment-types is encoded inside $D_n$ and does not break this factorisation.

4. **Class mixture is bilinear in emissions.** The class index $c$ enters every emission as a linear sum weighted by $u_{nfc}$, never inside the structural weights $C^{\alpha\beta}$. This makes the class mixture a low-rank emission factor that does not interact with the block-diagonal-plus-low-rank structure of the transition matrix.

##### C.6 MixDom-Specific SVI-BW Convergence Considerations

This appendix specialises the model-agnostic convergence analysis of Appendix B.2 and the BDI expected-statistic formulae of Appendix B.3 to the hierarchical MixDom model. Each subsection corresponds to a question that arises specifically because MixDom carries multiple interacting parameter groups (top-level vs. per-domain BDI, intra-fragment fragment-type chains, and per-class substitution models).

###### C.6.1 Parameter groups and Fisher information

The MixDom model has several parameter groups, each with different Fisher information characteristics:

| Parameter group | # params | Info per pair | Bottleneck |
|---|---|---|---|
| Top-level $(\lambda_0, \mu_0)$ | 2 | $O(1)$ | few domains per sequence |
| Per-domain $(\lambda_d, \mu_d)$ | $2N_{\text{dom}}$ | $O(w_d)$ | domain frequency $w_d$ |
| Intra-fragment fragment-type transitions $r_{fg}^{(d)}$ | $N_{\text{dom}}F^2$ | $O(w_d\bar{L}_d)$ | fragment count |
| Domain weights $w_d$ | $N_{\text{dom}} - 1$ | $O(1)$ | multinomial |
| Substitution $Q_c$ | $O(\vert\mathcal{A}\vert^2 C)$ | $O(\bar{L} \cdot w_c)$ | alignment length |

The bottleneck for SVB convergence is the **indel parameters of rare domains**. A domain with frequency $w_d = 0.05$ contributes useful BDI statistics to only $\sim 5\%$ of pairs, so its effective sample size is $\sim 0.05B$ per minibatch.

###### C.6.2 Substitution vs. indel information

Substitution parameters benefit from $O(\bar{L})$ information per pair (one observation per aligned column), while indel parameters get $O(1)$ information per pair. For a protein with $\bar{L} \approx 200$ residues, substitution parameters converge $\sim 200\times$ faster than indel parameters. This motivates **decoupled update frequencies**: update substitution parameters every minibatch, but average indel parameter estimates over multiple minibatches.

##### C.6.3 MixDom expected statistics

We now aggregate the per-process expectations of Appendix B.3 to the full MixDom model with $N_{\text{dom}}$ domain types.

**Top-level (domain birth-death)** The top-level BDI process has rates $(\lambda_0, \mu_0)$ with $\kappa_0 = \lambda_0/\mu_0$ and creates/destroys domains. At stationarity:

$$L_0 = \kappa_0/(1-\kappa_0) \quad \text{(expected \# domains per sequence)} \tag{C.105}$$

$$\mathbb{E}[B_0] = \lambda_0\, T/(1-\kappa_0) \quad \text{(domain births per pair)} \tag{C.106}$$

$$\mathbb{E}[D_0] = \mathbb{E}[B_0] \quad \text{(domain deaths per pair)} \tag{C.107}$$

$$\mathbb{E}[S_0] = L_0\, T \quad \text{(time-integrated domain count)} \tag{C.108}$$

$$M_0 = 1, \quad T_0 = T \quad \text{(one endpoint, one process)} \tag{C.109}$$

**Per-domain type $d$ (fragment birth-death within a domain)** Each surviving domain link of type $d$ (with probability $w_d$) contains a TKF92 fragment process with rates $(\lambda_d, \mu_d)$, $\kappa_d = \lambda_d/\mu_d$.

The number of domain links of type $d$ is approximately $L_d = w_d\, L_0$ at stationarity. Each such link contributes one independent BDI fragment process, so the *aggregated* fragment-level statistics for domain type $d$ are:

$$L_d = w_d\, L_0 \cdot \frac{\kappa_d}{1-\kappa_d} \quad \text{(total fragment links, type } d\text{)} \tag{C.110}$$

$$\mathbb{E}[B_d] = w_d\, L_0 \cdot \frac{\lambda_d T}{1-\kappa_d} \quad \text{(fragment births, type } d\text{, per pair)} \tag{C.111}$$

$$\mathbb{E}[D_d] = \mathbb{E}[B_d] \quad \text{(fragment deaths, type } d\text{, per pair)} \tag{C.112}$$

$$\mathbb{E}[S_d] = w_d\, L_0 \cdot \frac{\kappa_d T}{1-\kappa_d} \quad \text{(time-integrated fragment count, type } d\text{)} \tag{C.113}$$

$$M_d = w_d\, L_0 \quad \text{(\# independent processes)} \tag{C.114}$$

$$T_d = M_d\, T = w_d\, L_0\, T \quad \text{(total observation time)} \tag{C.115}$$

Note that $M_d$ and $T_d$ are themselves random (they depend on how many domains of type $d$ survive), but at stationarity their expectations are as above.

**Intra-fragment fragment-type transitions and sequence length** Within each fragment of domain $d$ the fragment-type chain is governed by an $F \times F$ Markov transition matrix $r_{fg}^{(d)}$ (intra-fragment; different fragments are independent realisations). The termination probability from fragment-type $f$ is $\rho_f^{(d)} = 1 - \sum_g r_{fg}^{(d)}$. The expected sojourn length starting from fragment state $f$ is determined by the fundamental matrix $(I - r^{(d)})^{-1}$: specifically, the expected number of sites emitted starting from a fragment initiated in state $f$ is $\sum_g [(I - r^{(d)})^{-1}]_{fg}$. Averaging over the initial fragment distribution $w_{d,f}$, the expected number of residues per fragment is $\bar{K}_d = \sum_f w_{d,f} \sum_g [(I - r^{(d)})^{-1}]_{fg}$.

The expected number of residues per domain of type $d$ is:

$$\bar{C}_d = \frac{\kappa_d}{1-\kappa_d}\, \bar{K}_d, \tag{C.116}$$

and the total expected sequence length is:

$$\bar{L}_{\text{seq}} = L_0 \sum_d w_d\, \bar{C}_d = \frac{\kappa_0}{1-\kappa_0} \sum_d w_d\, \frac{\kappa_d}{1-\kappa_d}\, \bar{K}_d. \tag{C.117}$$

*Remark* C.14 (Convergence of Markov chain transition counts). When fragments are IID (the $\mathcal{F} = 1$ scalar-extension special case), the sufficient statistics for the fragment extension parameter $r_f$ are Bernoulli counts (extend vs. terminate), and the convergence analysis reduces to a Beta-distributed posterior. With the general fragment-type Markov, the sufficient statistics are rows of a Markov chain transition count matrix: for each domain $d$ and source fragment state $f$, we observe counts $\hat{n}_{fg}^{(d)}$ of transitions to each target state $g$ plus termination counts $\hat{n}_{f,\text{end}}^{(d)}$. The M-step row-normalizes these counts (a Dirichlet posterior), and the per-pair Fisher information for each row scales with the expected number of visits to state $f$ per pair. For fragment states that are rarely visited (low stationary probability under $r^{(d)}$), the effective sample size is correspondingly small, mirroring the rare-domain bottleneck at the top level.

**Numerical example: d3 checkpoint** We evaluate these formulas using the MixDom d3 checkpoint parameters:

$$\begin{aligned}
&\lambda_0 = 0.01328, \qquad \mu_0 = 0.01412, \quad \kappa_0 = 0.9405\\
&w = (0.662,\ 0.075,\ 0.264)\\
&\lambda_d = (0.00302,\ 0.01033,\ 0.12370)\\
&\mu_d = (0.00372,\ 0.04686,\ 0.17775)\\
&\kappa_d = (0.812,\ 0.220,\ 0.696)\\
&T = 0.5
\end{aligned}$$

| | Top-level ($d$=0) | Dom 1 ($d$=1) | Dom 2 ($d$=2) | Dom 3 ($d$=3) |
|---|---|---|---|---|
| $\kappa$ | 0.941 | 0.812 | 0.220 | 0.696 |
| $L = \kappa/(1-\kappa)$ | 15.81 | 4.31 | 0.283 | 2.29 |
| $w_d \cdot L_0$ | — | 10.47 | 1.19 | 4.17 |
| $\mathbb{E}[B]$ per pair | 0.112 | 0.084 | 0.008 | 0.849 |
| $\mathbb{E}[D]$ per pair | 0.112 | 0.084 | 0.008 | 0.849 |
| $\mathbb{E}[S]$ | 7.91 | 22.58 | 0.168 | 4.78 |
| $M$ | 1 | 10.47 | 1.19 | 4.17 |
| $T$ | 0.5 | 5.23 | 0.59 | 2.09 |
| $v_\theta$ (approx.) | $\sim 18$ | $\sim 24$ | $\sim 255$ | $\sim 2.4$ |

**Computation of the table entries.** *Top level:* $L_0 = 0.9405/0.0595 = 15.81$. $\mathbb{E}[B_0] = 0.01328 \times 0.5/0.0595 = 0.112$. $\mathbb{E}[S_0] = 15.81 \times 0.5 = 7.91$.

*Domain type 1* ($w_1 = 0.662$): $M_1 = 0.662 \times 15.81 = 10.47$ independent fragment processes. $L_1 = 10.47 \times 0.812/0.188 = 10.47 \times 4.31 = 45.1$ total fragment links. $\mathbb{E}[B_1] = 10.47 \times 0.00302 \times 0.5/0.188 = 0.084$. (The table shows $L$ per process and $\mathbb{E}[B]$ aggregated over $M_d$ processes.)

*Domain type 3* ($w_3 = 0.264$): $M_3 = 0.264 \times 15.81 = 4.17$. $\mathbb{E}[B_3] = 4.17 \times 0.12370 \times 0.5/0.304 = 0.849$. This domain has high $\lambda_3$, so fragment births are frequent and its indel parameters are easy to estimate ($v \approx 2.4$).

$v_{\lambda_0}$: Using (B.73) with $\rho \approx 0.5$: $v_{\lambda_0} \approx 2/\mathbb{E}[B_0] = 2/0.112 = 17.9$.

$v_{\lambda_2}$: $\mathbb{E}[B_2] = 1.19 \times 0.01033 \times 0.5/0.780 = 0.0079$. $v_{\lambda_2} \approx 2/0.008 = 255$.

##### C.6.4 Convergence rate estimates

From (B.58) and the per-pair relative variance $v_\theta$, the number of pairs $N = BK$ needed for target relative error $\varepsilon$ is:

$$N \geq \frac{v_\theta}{\varepsilon^2}. \tag{C.118}$$

| Parameter | $v_\theta$ | $N$ for $\varepsilon$=10% | $N$ for $\varepsilon$=5% | $N$ for $\varepsilon$=1% |
|---|---|---|---|---|
| $\lambda_0$ (top-level ins) | 18 | 1 800 | 7 200 | 180 000 |
| $\mu_0$ (top-level del) | 18 | 1 800 | 7 200 | 180 000 |
| $\lambda_1$ (dom 1 ins) | 24 | 2 400 | 9 600 | 240 000 |
| $\lambda_2$ (dom 2 ins) | 255 | 25 500 | 102 000 | 2 550 000 |
| $\lambda_3$ (dom 3 ins) | 2.4 | 240 | 960 | 24 000 |
| $w_d$ (domain weights) | $\sim 1/L_0 \approx 0.06$ | 6 | 25 | 630 |
| $r^{(d)}_{fg}$ (fragment trans.) | $\sim 1/\bar{C}_d$ | | depends on domain | |
| Substitution ($Q$) | $\sim 1/\bar{L}_{\text{seq}}$ | | $\ll 100$ | |

##### C.6.5 Discussion: why top-level indel rates are hardest

The table reveals a clear hierarchy of estimation difficulty:

**1. Substitution parameters are easiest.** Each aligned residue pair contributes one independent observation to the substitution sufficient statistics. With $\bar{L}_{\text{seq}} \approx 200$–$400$ residues per sequence, the per-pair Fisher information for substitution parameters is $O(\bar{L}_{\text{seq}})$, so the per-pair relative variance is $O(1/\bar{L}_{\text{seq}})$. A few dozen pairs suffice for $\varepsilon = 5\%$ accuracy.

**2. Domain weights converge fast.** Each domain in the ancestor contributes one multinomial observation of the domain type. With $L_0 \approx 16$ domains per sequence, the per-pair Fisher information is $O(L_0)$, giving $v_{w_d} \sim 1/(w_d L_0)$. For the most common domain type ($w_1 = 0.662$), $v_{w_1} \approx 0.095$, and even for the rarest ($w_2 = 0.075$), $v_{w_2} \approx 0.84$. Domain weights are precisely estimated with $N \sim 100$ pairs.

**3. Fragment-type transition parameters are moderate.** Each intra-fragment fragment-type transition contributes one observation of the $F \times F$ Markov chain row $r^{(d)}_{f,:}$. With $\sim L_0 w_d \kappa_d/(1-\kappa_d)$ fragments of type $d$ per pair, the per-pair information scales with the total fragment count. For domain 1 (which dominates), there are $\sim 45$ fragment links, so the per-row variance scales as $\sim F/(45 \cdot \pi_f)$ where $\pi_f$ is the stationary probability of fragment state $f$. Domain 2 fragments are rare ($\sim 0.3$ per pair) and harder to estimate.

**4. Indel parameters for active domains are easy.** Domain type 3 has high insertion rate ($\lambda_3 = 0.124$) and $\mathbb{E}[B_3] \approx 0.85$ births per pair, giving $v_{\lambda_3} \approx 2.4$. A few hundred pairs suffice for 5% accuracy.

**5. Indel parameters for common but slow domains are moderate.** Domain type 1 has $\mathbb{E}[B_1] \approx 0.084$ births per pair, giving $v_{\lambda_1} \approx 24$. This requires $N \approx 10{,}000$ pairs for 5% accuracy—feasible but not trivial.

**6. Indel parameters for rare domains are the bottleneck.** Domain type 2 has weight $w_2 = 0.075$, low $\kappa_2 = 0.22$ (short domains), and low $\lambda_2 = 0.01$. The expected fragment births per pair are only $\mathbb{E}[B_2] \approx 0.008$. This means on average, only 1 in $\sim 125$ pairs shows even a single fragment birth event in a domain of type 2. The per-pair relative variance $v_{\lambda_2} \approx 255$ requires $N > 100{,}000$ pairs for 5% accuracy.

**7. Top-level indel rates are intrinsically hard.** The top-level BDI controls domain creation and destruction. With $\lambda_0 = 0.013$, $T = 0.5$, and $L_0 \approx 16$ domains, the expected number of domain births per pair is only $\mathbb{E}[B_0] \approx 0.11$. The per-pair relative variance $v_{\lambda_0} \approx 18$ requires $N \approx 7{,}200$ pairs for 5% accuracy.

The fundamental reason is that domain-level indel events are rare compared to residue-level observations. A sequence of 300 residues organized into 16 domains gives $\sim 300$ substitution observations but only $\sim 0.1$ domain birth observations per pair at $T = 0.5$. The information ratio is roughly $300/0.1 = 3{,}000\times$ in favor of substitution parameters.

**Implications for training.** These estimates motivate the decoupled update strategy recommended in Section 5: substitution parameters can be accurately estimated from small minibatches ($B \sim 10$), while indel parameters (especially for rare domain types) require accumulation over $N \sim 10^4$–$10^5$ pairs. The Maraschino pipeline (Section 5.5) achieves this by processing all training pairs in a single count tensor, at the cost of the composite-likelihood efficiency gap. The hybrid Maraschino $\to$ SVB pipeline is optimal: Maraschino provides accurate initial estimates using all data, and SVB refines the indel parameters using the full model.

#### C.7 Variational EM training of MixDom from tree-structured data

The variational ELBO of Appendix C.8 treats the model parameters $\theta = (\lambda_0, \mu_0, \{\lambda_n\}, \{\mu_n\}, v, w, r^{(n)}, u, \{\pi^{(c)}\}, \{S^{(c}_{\text{ex}}$ as fixed inputs and optimises only the variational distribution $q$ over per-(node, column) latent states. In this appendix we extend the framework to a Variational Bayesian EM (VBEM) training algorithm that learns $\theta$ by alternating between a per-family E-step (Adam ascent on the variational $q$) and a global M-step (closed-form $\theta$ update from aggregated sufficient statistics).

This is structurally analogous to the SVI Baum-Welch pipeline of Section C.1.4 (which trains MixDom from sequence *pairs* via the labelled Pair HMM), but operates on whole MSAs with their tree topology — each MSA family becomes a single training datum, with the variational distribution capturing the posterior over internal-node states and column-wide tuples.

##### C.7.1 Outer EM loop

Given the corpus $\{\mathcal{D}_i\}_{i=1}^{N}$ of MSA-with-tree training data and an initial parameter estimate $\theta^{(0)}$, the outer loop is

- **E-step** (per family $i$): fit the variational $q_i = (q_i^{(\tau)}, q_i^{(\pi|\tau)})$ by Adam ascent on the ELBO $\mathcal{L}_i(q_i; \theta^{(t)})$ at the current parameter estimate, returning per-family sufficient statistics $\Phi_i = \{W_i^{(v\to w)}, q_i^{(\tau)}, L_{i,n,c}^{(\text{sub})}\}$.
- **M-step** (global): aggregate sufficient statistics across families and apply closed-form parameter updates for each parameter group, yielding $\theta^{(t+1)}$.

The corpus may be subsampled minibatch-style each iteration (SVI-style), in which case the M-step uses an exponential-moving-average of sufficient statistics across iterations (Section B.1.13).

##### C.7.2 Per-family E-step

For family $i$ with binary tree $\mathbf{t}_i$, branch lengths $d_i$, and observed leaf data $X_i$, the E-step maximises

$$\mathcal{L}_i(q_i;\theta) = \sum_{(v\to w)\in\mathbf{t}_i} \mathbb{E}_{q_i}[\log P^{\mathrm{WFST}}(Z^w \mid Z^v)] + \sum_n \mathbb{E}_{q_i^{(f)}}[\log L_n^{(\mathrm{sub}),\mathrm{tot}}] + \log p^{\mathrm{red}}_{\mathrm{singlet}}(Z^{\mathrm{root}}) + H[q_i^{(\tau)}] + H[q_i^{(\pi|\tau)}] + \log Z_q \tag{C.119}$$

over the variational logits (edge_logits, root_logit, tuple_logits) via JIT-compiled Adam, identical to the inference-time ELBO of Appendix C.8 but treating $\theta$ as fixed input.

After convergence (typically 100–300 Adam iterations with lr $\approx 0.05$, Fitch-seeded init), the E-step extracts the following sufficient statistics from $q_i$:

**Per-branch reduced expected counts.** For each branch $(v \to w)$, the cumulant-trick prefix sum (Section C.8.6) yields

$$W^{(v\to w)}_{ss',\tau\tau'} = \sum_{N=1}^{L+1}\sum_{M=0}^{N-1} q^{(\tau)}_{n(M)}(\tau)\, q^{(\tau)}_{n(N)}(\tau')\, P^{v\to w}_{q,M}(s)\, P^{v\to w}_{q,N}(s') \prod_{K=M+1}^{N-1} P^{v\to w}_{q,K}(\mathsf{lg}), \tag{C.120}$$

a $5 \times T \times 5 \times T$ tensor that summarises the expected labelled WFST transitions on this branch.

**Per-column class posteriors.** For each column $n$ and class $c$, compute

$$q_n^{(c|f)}(c) = \frac{u_{f,c}\, L_n^{(\mathrm{sub})}(c;\mathcal{F}_n)}{\sum_{c'} u_{f,c'}\, L_n^{(\mathrm{sub})}(c';\mathcal{F}_n)}, \quad q_n^{(c)}(c) = \sum_f q_n^{(f)}(f)\, q_n^{(c|f)}(c), \tag{C.121}$$

where $\mathcal{F}_n$ is the Fitch subtree of column $n$ and $L_n^{(\mathrm{sub})}(c)$ is the Felsenstein up-pass likelihood under class-$c$ rate matrix.

**Per-class substitution counts.** For each class $c$ and column $n$, weighted Felsenstein expected substitution counts on $\mathcal{F}_n$:

$$\hat{M}^{(c)}_{i,n}[a,b] = q_n^{(c)}(c)\,\mathbb{E}[N_{ab}(\mathrm{branch};\mathcal{F}_n) \mid \text{leaf data}, c, \theta], \qquad \hat{T}^{(c)}_{i,n}[a] = q_n^{(c)}(c)\,\mathbb{E}[\text{dwell time in } a \mid \text{leaf data}, c, \theta]. \tag{C.122}$$

These are computed via the standard Holmes-Rubin posterior expectation formulae for substitution-only CTMCs on trees, weighted by the per-class responsibility $q_n^{(c)}(c)$.

##### C.7.3 M-step from aggregated sufficient statistics

Let $\Phi = \{\Phi_i\}_i$ denote the per-family sufficient statistics aggregated across the (mini)batch. The M-step decomposes parameter group by parameter group, exploiting the route decomposition (C.144) and the responsibilities derived above.

**Route attribution.** For each per-character labelled WFST transition the route posterior

$$\rho^{(r)}_{ss',\tau\tau'}(\theta) = \frac{\omega^{(r)}_{\tau\tau'}\, \tilde{T}^{\mathrm{lab},(r)}_{ss'}}{\hat{T}_{ss'}(\tau,\tau';\theta)} \tag{C.123}$$

partitions the per-branch counts $W^{(v\to w)}_{ss',\tau\tau'}$ into route-attributable soft counts

$$\tilde{W}^{(r),(v\to w)}_{ss',\tau\tau'} = W^{(v\to w)}_{ss',\tau\tau'} \cdot \rho^{(r)}_{ss',\tau\tau'}(\theta^{(t)}). \tag{C.124}$$

Each route $r \in \{R1, R2, R3\}$ has a fixed BDI / fragment / domain factor signature (Section C.8.4): R1 contributes only to within-fragment extension counts; R2 contributes to fragment-level BDI counts in domain $d$; R3 contributes to both top-level BDI counts and to fragment-level BDI counts in the *destination* domain $d'$.

**Indel rate M-step.** For each domain $d$, the per-route soft counts in $\tilde{W}^{(R2)}$ (restricted to $d = d'$ in source/dest tuples) and $\tilde{W}^{(R3)}$ (with destination domain $d'$) accumulate to a per-domain $5 \times 5$ WFST transition count matrix $\hat{n}^{(d)}_{ss'}$ analogous to the SVI-BW pair-counts. The standard transition-count groups (Section C.1.4, `transition_count_groups`) and quadratic-in-$\kappa$ closed form (`m_step_indel_quadratic`) then deliver $(\hat{\lambda}_d, \hat{\mu}_d)$:

$$(\hat{\lambda}_d, \hat{\mu}_d) = \text{m\_step\_indel\_quadratic}(\hat{B}_d, \hat{D}_d, \hat{S}_d, \hat{L}_d, \hat{M}_d, \hat{T}_d; \text{prior}). \tag{C.125}$$

The top-level rates $(\hat{\lambda}_0, \hat{\mu}_0)$ are recovered analogously from $\tilde{W}^{(R3)}$ contributions to the top-level WFST transition counts (with R3 supplying both a top-level BDI event and a destination-domain BDI event per transition).

**Within-fragment Markov M-step.** For each domain $d$, the per-fragchar transition matrix $r^{(d)}_{f,f'}$ has Dirichlet-conjugate update from the R1 soft counts:

$$\hat{r}^{(d)}_{f,f'} = \frac{\hat{E}_{d,f,f'} + \alpha_r - 1}{\sum_{f''} \hat{E}_{d,f,f''} + \hat{N}_{d,f} + (N_{\mathcal{F}} + 1)(\alpha_r - 1)}, \quad \hat{\rho}^{(d)}_f = 1 - \sum_{f'} \hat{r}^{(d)}_{f,f'}, \tag{C.126}$$

where $\hat{E}_{d,f,f'} = \sum_{(v \to w),s,s'} \tilde{W}^{(R1),(v\to w)}_{ss',(d,f),(d,f')}$ and $\hat{N}_{d,f} = \sum_{(v\to w),s,s'} \big[\tilde{W}^{(R2)}_{ss',(d,f),(d,\cdot)} + \tilde{W}^{(R3)}_{ss',(d,f),(\cdot,\cdot)}\big]$ are the within- fragment and fragment-end soft counts respectively.

**Domain and fragment weight M-step.** The tuple priors update via Dirichlet-conjugate from per-route soft-counts:

$$\hat{v}_{d'} \propto \sum_{(v\to w)} \sum_{(d,f,f'),s,s'} \tilde{W}^{(R3),(v\to w)}_{ss',(d,f),(d',f')} + \alpha_n - 1, \tag{C.127}$$

$$\hat{w}_{d',f'} \propto \sum_{(v\to w)} \sum_{(d,f),s,s'} \Big[\delta_{d,d'} \tilde{W}^{(R2)}_{ss',(d,f),(d',f')} + \tilde{W}^{(R3)}_{ss',(d,f),(d',f')}\Big] + \alpha_w - 1, \tag{C.128}$$

both normalised over the appropriate index.

**Substitution M-step.** Per-class rate-matrix and stationary updates use the standard class-weighted Holmes-Rubin closed forms (Section B.1.10 and the `substitution-mstep.tex` appendix), with per-class soft-counts $\hat{M}^{(c)} = \sum_i \sum_n \hat{M}^{(c)}_{i,n}$ and $\hat{T}^{(c)} = \sum_i \sum_n \hat{T}^{(c)}_{i,n}$. The class distribution itself updates via Dirichlet-conjugate:

$$\hat{u}_{d,f,c} \propto \sum_i \sum_n q^{(\tau)}_{i,n}(d,f) \cdot q^{(c|f)}_{i,n}(c) + \alpha_u - 1. \tag{C.129}$$

##### C.7.4 Stochastic VBEM (SVI-VBEM)

For corpora with $N \gg 100$ families a full-batch outer iteration is infeasible (Appendix C.7.7). We adopt the SVI-BW machinery of Appendix B.2.4 verbatim, with one substitution: the per-batch sufficient-statistic vector $s_{\text{batch}_k}$ in (B.52) is now the per-family Tree-VBEM aggregate

$$s_{\text{batch}_k} \;=\; \sum_{i\in\mathcal{B}_k} \Big\{\tilde{W}_i^{(R1)},\, \tilde{W}_i^{(R2)},\, \tilde{W}_i^{(R3)},\, q_i^{(\tau)},\, \hat{M}_i^{(c)},\, \hat{T}_i^{(c)}\Big\}, \tag{C.130}$$

each component being aggregated linearly over the families $i \in \mathcal{B}_k$ in minibatch $k$. (The route-attributed soft counts $\tilde{W}_i^{(r)}$ are themselves linear in the per-branch $W_i^{(v\to w)}$ tensors of (C.120) and the route posteriors (C.123), so the per-family contributions sum directly without bias.) The pseudocount EMA carries one such state $\tilde{\alpha}_k^{(g)}$ per parameter group $g$ and is updated each iteration as

$$\tilde{\alpha}_k^{(g)} \;=\; (1-\eta_k)\,\tilde{\alpha}_{k-1}^{(g)} \;+\; \eta_k\big(\alpha^{(g)} + (N/|\mathcal{B}_k|)\, s_{\text{batch}_k}^{(g)}\big), \tag{C.131}$$

with the closed-form M-step applied on demand to derive $\theta_g^{(t+1)} = f^{(g)}(\tilde{\alpha}_k^{(g)})$ for each group: the BDI rates per domain via (C.125); the Dirichlet groups $v$, $w$, $u$, $r_{f,\cdot}^{(d)}$ direct from (C.126)–(C.128) and (C.129); and the per-class GTR substitution parameters via the closed forms of Appendix B.1.10.

The Polyak–Ruppert / ESS / Fisher analyses of Appendix B.2.4 (eqs. (B.54) onward) carry over verbatim, with $s_{\text{batch}_k}$ replaced by the family-aggregate (C.130).

**Step-size schedule.** The standard Robbins–Monro choice $\eta_k = (k+\tau)^{-\kappa}$ with $\tau \in [10, 100]$, $\kappa \in [0.5, 1]$ guarantees almost-sure convergence as in SVI-BW. For tree-VBEM we suggest defaults $\tau = 10$, $\kappa = 0.7$.

**Breadth-first minibatch sampling.** I.i.d. uniform minibatch sampling concentrates updates on the $\sim |\mathcal{B}|/N$ most-recently-visited families and under-represents the long tail; per-group ESS for a parameter that is informative in only a fraction $\varepsilon$ of families collapses to $\varepsilon\,\mathrm{ESS}_K$ (the "rare-parameter bottleneck" of Appendix B.2.4). We instead maintain a per-family visit count $c_i^{(0)} = 0$ and select each minibatch $\mathcal{B}_k$ as the $|\mathcal{B}|$ families with the smallest $c_i^{(k-1)}$, breaking ties at random; visit counts then update $c_i^{(k)} = c_i^{(k-1)} + 1$ for $i \in \mathcal{B}_k$. This deterministic round-robin guarantees every family is visited at least once every $\lceil N/|\mathcal{B}|\rceil$ iterations ("one epoch"), so the per-family contribution arrives in $\Theta(K|\mathcal{B}|/N)$ of the first $K$ batches with no starvation.

**Convergence diagnostics.** Per outer iteration $k$, log:

- Per-batch ELBO sum, scaled by $N/|\mathcal{B}_k|$ for cross-iteration comparison.
- Per-group $\tilde{\alpha}_k^{(g)}$ effective sample size, $\mathrm{ESS}_k = (\sum_j w_{j,k})^2 / \sum_j w_{j,k}^2$ with $w_{j,k} = \eta_j \prod_{i>j}^{k}(1-\eta_i)$, to monitor warm-up.
- Visit-count distribution $\min_i c_i^{(k)}$, $\max_i c_i^{(k)}$, to confirm the breadth schedule is achieving uniform coverage.

Validation ELBO on a held-out family subset is computed once per "epoch" (every $\lceil N/|\mathcal{B}|\rceil$ iterations).

##### C.7.5 Convergence and ELBO monitoring

Each E-step monotonically increases the per-family ELBO at fixed $\theta$; each M-step monotonically increases the corpus-aggregate ELBO at the new q (this is the standard EM monotonicity result, modulo the variational E-step's sub-optimality).

For monitoring purposes, log:

- Per-iteration corpus ELBO (sum across families).
- Per-domain BDI sufficient statistics aggregates.
- Per-class soft-count totals and effective sample size.
- Validation-set ELBO on a held-out family subset (for early stopping and overfitting monitoring; same role as val LL/pair in SVI-BW).

Convergence in practice: corpus ELBO increase plateaus below a relative threshold ($\sim 10^{-4}$ per outer iteration), or validation ELBO begins to decrease.

##### C.7.6 Initialisation and warm-start

For warm-start from an existing SVI-BW checkpoint: $\theta^{(0)}$ is loaded directly from the checkpoint (the variational EM operates on the same parameter space as SVI-BW provided the checkpoint includes `class_pis` and `class_S_exch`; checkpoints without a class layer are auto-promoted to a 1-class-per-domain structure).

For the variational $q^{(0)}$: per-family Fitch-seeded init for the inner 3-state q (as in the inference-only benchmark of Appendix C.8); per-family tuple init biased toward the substitution-likelihood-maximising fragchar (via `class_marginalised_sub_LL_per_column` applied to the initial $\theta$).

The first outer iteration will see a large ELBO improvement as the variational distributions are fit to the warm-start parameters. Subsequent iterations refine $\theta$ toward the tree-aggregated likelihood maximum, which differs in general from the pair-aggregated SVI-BW maximum.

##### C.7.7 Computational scaling and minibatching

Per-family E-step cost is dominated by Adam-ELBO evaluation: $O(|\mathcal{E}| \cdot L \cdot T^2)$ per Adam step where $T = N_n N_{\mathcal{F}}$ is the reduced tuple count, plus $O(L \cdot N_{\text{cl}} \cdot \text{Felsenstein cost})$ for the substitution likelihoods (computed once per family at the start of the E-step). Typical numbers for unified-short ($T = 6$, $N_{\text{cl}} = 3$, $|\mathcal{E}| \approx 40$, $L \approx 100$, 100 Adam iters) give $\sim 30$ s/family on a single GPU.

For the full Pfam corpus ($\sim 17{,}000$ families) a full E-step pass is $\sim 140$ GPU-hours. Stochastic VBEM (Appendix C.7.4) replaces the full pass with a breadth-first minibatch of $\sim 10$–200 families per outer iteration and an EMA accumulation of sufficient statistics across iterations, reducing per-iteration cost to a few GPU-minutes with convergence in $\sim 10\,N/|\mathcal{B}|$ outer iterations (i.e. $\sim 10$ epochs).

##### C.7.8 Comparison to SVI-BW

The SVI-BW pipeline trains $\theta$ from sequence pairs sampled from the corpus, using the labelled Pair HMM forward-backward as its inference primitive. Tree-VBEM trains from whole MSAs with their tree topology, using the variational TreeVarAnc-MixDom inference of Appendix C.8.

Differences:

- **Information per training datum.** A pair contributes two-leaf data; a family contributes |leaves|-leaf joint data with phylogenetic structure. Tree-VBEM extracts more information per datum but requires more compute per datum.

- **Bias.** SVI-BW assumes pairs are i.i.d. samples from the model; real pairs share evolutionary history (within-family pairs are not independent under the tree). Tree-VBEM correctly handles this correlation. The cost: SVI-BW's pair-likelihood is the exact data likelihood under its assumption; tree-VBEM's ELBO is a lower bound on the family likelihood (with a non-trivial variational gap).
- **Application alignment.** Parameters trained by tree-VBEM are likely better-suited to tree-based downstream tasks (ancestral reconstruction, progressive alignment) since the training objective matches the inference task. Parameters trained by SVI-BW may be better for pairwise tasks (pairwise alignment scoring).

Empirically, the two regimes can be combined: SVI-BW for fast warm-up followed by tree-VBEM for task-specific fine-tuning, exploiting both algorithms' strengths.

#### C.8 Mixture-of-trees variational MixDom ancestral inference

This appendix generalises the variational ancestral-presence reconstruction of Section B.6 from the TKF92-conditional-WFST-as-MaxEnt-GGI approximation to the labelled MixDom model, with two further simplifications relative to a fully labelled treatment:

- the per-character substitution class $c$ is integrated out at the model level via the per-column prior $u_{f,c}$ (rather than carried as a variational latent);
- the fragchar-boundary indicator $g$ and domain-end indicator $e$ are absorbed into a *reduced* per-character WFST kernel that sums over the implicit fragment/domain bookkeeping at each step (rather than tracked explicitly in the variational state).

The variational state per (internal node, MSA column) is therefore $\{N, D\} \cup \{(d, f)\}$, $|\mathcal{Z}| = N_{\text{dom}} N_{\text{fr}} + 2$ — a $(4N_{\text{cl}})\times$ reduction over the fully-labelled state space. Three concrete payoffs: (i) the variational bound is *strictly tighter* than the fully-labelled ELBO at any non-Bayes-optimal class posterior, because the analytic marginalisation of $c$ and $(g, e)$ replaces two Jensen inequalities with equalities; (ii) the labelled-variant cross-column hard-zero structural constraint disappears, so per-column factorised $q^{(\tau)}$ is adequate (the column-Markov-chain upgrade is no longer required); (iii) per-branch evaluation is roughly $|\mathcal{T}^{\text{lab}}|^2/|\mathcal{T}|^2$-times cheaper.

The bound is $\log p(\text{MSA} \mid \mathbf{t}, \theta) \geq \log \tilde{p}(\text{MSA} \mid \mathbf{t}, \theta) \geq \mathcal{L}[q]$, in the same restricted-model sense as appendix B.6; the restriction-gap ($q$-independent) and variational-gap (KL) decomposition carries over.

##### C.8.1 Setting and reduced state space

Fix a rooted phylogeny $\mathbf{t}$ with internal-node set $\mathcal{I}$, leaf set $\mathcal{L}$, and branch lengths $d_e$ on each edge $e$. The MSA has $L$ columns; each leaf $v \in \mathcal{L}$ has an observed presence indicator $X_n^v \in \{0, 1\}$ and (when present) a residue $a_n^v \in \mathcal{A}$.

The per-(node, column) variational state space is

$$\mathcal{Z} \;=\; \{N, D\} \cup \mathcal{T}, \qquad \mathcal{T} \;=\; [N_{\text{dom}}] \times [N_{\text{fr}}]. \tag{C.132}$$

A tuple-valued state $\tau = (d, f) \in \mathcal{T}$ encodes only the column's domain $d$ and fragchar $f$; the per-character class $c$ is marginalised at the model level (Section C.8.7) and the fragchar-boundary / domain-end indicators $(g, e)$ are absorbed into the reduced WFST kernel (Section C.8.4).

The presence indicator at internal node $v$, column $n$ is $X_n^v = \mathbb{1}\{Z_n^v \in \mathcal{T}\}$. Leaf states are partially clamped: $X_n^v = 0$ pins $Z_n^v \in \{N, D\}$; $X_n^v = 1$ pins $Z_n^v \in \mathcal{T}$ (consistent with the column-wide tuple $\tau_n$ — see below).

##### C.8.2 Restricted generative model

Following Section B.6.2, the model joint over MSA columns and internal-node states is

$$p(\mathrm{MSA},\, \{Z^v\}_{v\in\mathcal{I}} \mid \mathbf{t}, \theta) \;=\; p^{\mathrm{red}}_{\mathrm{singlet}}(Z^{\mathrm{root}}) \prod_{(v\to w)\in\mathbf{t}} \hat{P}^{\mathrm{WFST}}(Z^w \mid Z^v, d_{vw}, \theta) \prod_{n:\mathcal{F}_n\neq\emptyset} L_n^{(\mathrm{sub}),\mathrm{tot}}(f_n; \mathcal{F}_n), \tag{C.133}$$

where $\mathcal{F}_n$ is the Fitch-parsimony subtree of column $n$, $L_n^{(\mathrm{sub}),\mathrm{tot}}(f;\mathcal{F}_n)$ is the class-marginalised Felsenstein column-substitution likelihood under fragchar $f$ (Section C.8.7), and $p^{\mathrm{red}}_{\mathrm{singlet}}, \hat{P}^{\mathrm{WFST}}$ are the *reduced* singlet HMM and WFST defined on $(d, f)$ alone (Section C.8.4). As in the simple case we marginalise (C.133) over internal patterns supported on the $L$ observed columns to obtain $\tilde{p}(\mathrm{MSA} \mid \mathbf{t}, \theta) \leq p(\mathrm{MSA} \mid \mathbf{t}, \theta)$ (with the same ghost-column caveat), and bound the latter from below.

##### C.8.3 Variational family

The variational $q$ factorises over MSA columns, $q(\{Z_n^v\}) = \prod_{n=1}^{L} q_n$, and within each column,

$$q_n \;=\; q_n^{(\tau)}(\tau_n)\cdot q_n^{(\pi|\tau)}\big(\{Z_n^v\}_{v\in\mathcal{I}} \,\big|\, \tau_n\big), \tag{C.134}$$

where $q_n^{(\tau)}$ is a free categorical over $\mathcal{T}$ on the column-wide tuple, and $q_n^{(\pi|\tau)}$ is the same 3-state irreversible tree-structured graphical model of Section B.6.5 (with $xP \equiv \tau_n$). By construction $q$ assigns zero mass to within-column configurations in which Present nodes carry differing tuples — the column has one $(d, f)$, full stop. The cross-column structural-label constraints that forced a column-Markov chain in the labelled formulation *vanish* under reduction (Section C.8.9).

**Free parameter count.** Per column: $|\mathcal{T}| - 1 = N_{\mathrm{dom}}N_{\mathrm{fr}} - 1$ free parameters in $q_n^{(\tau)}$, plus $2|\mathcal{E}|$ in the inner 3-state graphical model (as in Section B.6.10), plus 2 for the inner root distribution. For typical $(N_{\mathrm{dom}}, N_{\mathrm{fr}}) = (3, 2)$ that is 5 tuple parameters per column, vs $4N_{\mathrm{cl}}N_{\mathrm{fr}}N_{\mathrm{dom}} - 1$ in the labelled formulation (e.g. 239 for $N_{\mathrm{cl}} = 10$) and $\sim |\mathcal{T}|^2$ in the labelled column-Markov upgrade ($\sim 6\times 10^4$).

##### C.8.4 Reduced WFST: marginalising $(g, e)$ and the class $c$

The reduced per-character WFST kernel is

$$\hat{T}_{ss'}\big((d, f), (d', f');\, d, \theta\big), \qquad s, s' \in \{\mathtt{S}, \mathtt{M}, \mathtt{I}, \mathtt{D}, \mathtt{E}\}, \tag{C.135}$$

obtained by marginalising the labelled-MixDom WFST $T^{\mathrm{lab}}_{ss'}((c, f, g, d, e), (c', f', g', d', e'); d, \theta)$ over $(c, c', g, e, g', e')$ at each end. The reduced state space is $\{\mathtt{S}, \mathtt{E}\} \cup \{(s, d, f) : s \in \{\mathtt{M}, \mathtt{I}, \mathtt{D}\}\}$ — i.e. $3N_{\mathrm{dom}}N_{\mathrm{fr}} + 2$ states.

**Routes from $(d, f)$ to $(d', f')$.** Under the labelled MixDom model, the per-character labelled transition $(d, f) \to (d', f')$ admits up to three latent routes, indexed by the value of $(g, e)$ at the source position:

R1. *Intra-fragment fragchar transition* $(g = 0)$: the current character is not the last of its fragment, and the next character stays in the same fragment with fragchar transitioning $f \to f'$ via $r^{(d)}_{f,f'}$. Restricted to $d' = d$.

R2. *New fragment, same domain* ($g = 1, e = 0$): the current fragment terminates ($\rho_f^{(d)}$) and the same domain starts a new fragment ($\kappa_d$) with first fragchar drawn from $w_{d,f'}$. Restricted to $d' = d$.

R3. *New domain* ($g = 1, e = 1$): the current fragment terminates, the current domain ends ($1-\kappa_d$), and (after possibly skipping geometrically-many empty domains) a new domain $d'$ begins with its first fragment's first fragchar drawn from $w_{d',f'}$. Available for any $d'$, including the self-recurrence $d' = d$.

For $d' \neq d$ only R3 is enabled, so the source's $(g, e) = (1, 1)$ is uniquely determined by the transition. For $d' = d$, multiple routes contribute and the latent $(g, e)$ at the source carries a non-trivial posterior over routes given the observed transition. The diagonal-extension special case ($r_{f,f'}^{(d)} = r_f \delta_{f,f'}$) suppresses R1 whenever $f' \neq f$, but R2 and R3 still mix for any $(d, f) \to (d, f')$, so the route-sum machinery introduced below is required even there. With a general off-diagonal $r^{(d)}$ matrix, R1 admits $f' \neq f$ transitions, so the route-sum acquires a third contributing term.

The per-route singlet contribution, accounting for the joint prior on $(g, e)$ at the source and the singlet emission of the next character, is:

$$\omega^{(R1)}_{(d,f)\to(d',f')} = \delta_{d,d'}\, r^{(d)}_{f,f'}, \tag{C.136}$$

$$\omega^{(R2)}_{(d,f)\to(d',f')} = \delta_{d,d'}\, \rho^{(d)}_f\, \kappa_d\, w_{d,f'}, \tag{C.137}$$

$$\omega^{(R3)}_{(d,f)\to(d',f')} = \frac{\rho^{(d)}_f\,(1-\kappa_d)\,\kappa_0\, v_{d'}\,\kappa_{d'}\, w_{d',f'}}{1-\zeta}, \tag{C.138}$$

where $\zeta = \kappa_0 \sum_{d''} v_{d''}(1-\kappa_{d''})$ is the empty-domain renormalisation (equation (C.178)). Each $\omega^{(r)}$ factorises as $P((g, e) = (g_r, e_r) \mid (d, f)) \cdot P_{\text{singlet}}((d', f') \mid (d, f), g_r, e_r)$, so summing over routes recovers the marginal singlet emission probability:

$$\omega^{(d,f,d',f')} = \sum_r \omega^{(r)}_{(d,f)\to(d',f')} = \delta_{d,d'}\Big[r^{(d)}_{f,f'} + \rho^{(d)}_f\,\kappa_d\, w_{d,f'}\Big] + \frac{\rho^{(d)}_f\,(1-\kappa_d)\,\kappa_0\, v_{d'}\,\kappa_{d'}\, w_{d',f'}}{1-\zeta}. \tag{C.139}$$

**Marginalisation of $c$.** The class $c$ at column $n$ is generated once per column by $u_{f_n,\cdot}$ at column birth and governs the substitution likelihood across all branches. The labelled WFST's indel block is class-independent, so $\sum_c u_{f,c} = 1$ trivially. Class-marginalisation of $\hat{T}$ has no residue at the indel level; it is handled separately in the substitution term (Section C.8.7).

**Step-by-step derivation of the reduced kernel** The route-sum (C.144) is not stipulated; it follows from two facts about the labelled MixDom model. We state and prove both.

**Notation.** Write $Z_n = (d_n, f_n, g_n, e_n)$ for the labelled chain state at position $n$ (dropping $c_n$, which is class-independent for the indel block) and $\tau_n = (d_n, f_n)$ for the reduced state. Let $\sigma$ denote the labelled singlet HMM transition kernel of mixdom-wfst.tex Section C.11.2, and let $T^{\text{lab}}_{ss'}$ denote the labelled conditional WFST kernel of Section C.11.3. The labelled per-character Pair HMM joint factorises as

$$P\big((s', Z') \mid (s, Z)\big) = T^{\text{lab}}_{ss'}\big(Z, Z'\big)\, \sigma\big(Z' \mid Z\big), \tag{C.140}$$

i.e. the WFST conditional weight times the singlet emission weight; this is the construction of mixdom-wfst.tex Section C.11.3 (*Singlet ∘ WFST = Pair HMM*).

**Singlet-table convention.** Throughout the proof we use the singlet HMM table at mixdom-wfst.tex line 165–170 with the following normalisation (consistent with the joint-contribution language at line 187–189 and the marginalisation check at line 228–237). Two structural facts about the labelled MixDom chain underpin this:

(a) *$e$ is fragment-level.* The domain-end indicator $e$ is constant for all characters within a single fragment (it records whether the current fragment is the last in its domain, which is set when the fragment begins and does not change as the fragment extends). Hence within-fragment transitions ($g$=0 rows) have $e' = e$ implicitly, and a single "$g$=0 row" covers all source $e$ values without depending on $e$.

(b) *Destination indicators $(g', e')$ are summed implicitly.* The destination $(g', e')$ in each row is treated as a free index ranging over $\{0, 1\}^2$, with the implicit prior $\pi(g', e' \mid d', f')$ to be applied separately if a finer-grained joint over destination indicators is needed.

Under these conventions, the table entry $\widetilde{\sigma}$ for a given source-row $(d, f, g, e)$ and destination structural state $(d', f')$ is the joint

$$\widetilde{\sigma}((d, f, g, e) \to (d', f')) \;=\; \pi(g, e \mid d, f) \,\cdot\, \sigma_{\text{cond}}((d', f') \mid (d, f, g, e)),$$

where $\sigma_{\text{cond}}$ is the source-conditional next-character emission probability and $\pi(g, e \mid d, f)$ is the source-row marginal. As a sanity check: summing each row of the table over $(d', f')$ recovers exactly $\pi(g, e \mid d, f)$, which sums in turn to 1 over the three rows (with termination column included). For the $g$=0 row: $\sum_{f'} r^{(d)}_{f,f'} = 1{-}\rho^{(d)}_f = \pi(g{=}0 \mid d, f)$. For the $g$=1, $e$=0 row: $\sum_{f'} \rho^{(d)}_f \kappa_d w_{d,f'} = \rho^{(d)}_f \kappa_d = \pi(g{=}1, e{=}0 \mid d, f)$. For the $g$=1, $e$=1 row, mixdom-wfst.tex line 228–237 verifies the row sum (over destinations plus termination) equals $\rho^{(d)}_f (1{-}\kappa_d) = \pi(g{=}1, e{=}1 \mid d, f)$.

**Lemma 1 (Conditional independence of $(g, e)$).** *Under the labelled MixDom singlet construction, $(g_n, e_n) \perp \{Z_k : k < n\} \mid (d_n, f_n)$ for every $n \geq 1$, with*

$$\pi(g{=}0 \mid d, f) = 1 - \rho^{(d)}_f, \tag{C.141}$$

$$\pi(g{=}1, e{=}0 \mid d, f) = \rho^{(d)}_f \, \kappa_d, \tag{C.142}$$

$$\pi(g{=}1, e{=}1 \mid d, f) = \rho^{(d)}_f \, (1 - \kappa_d). \tag{C.143}$$

*Proof.* By the singlet-table convention above, the labelled-chain joint factorises position-by-position as

$$P(Z_n \mid Z_{n-1}) = P(d_n, f_n \mid Z_{n-1}) \cdot \pi(g_n, e_n \mid d_n, f_n),$$

because the table entry at position $n$ has the destination prior $\pi(g_n, e_n \mid d_n, f_n)$ as an explicit multiplicative factor (line 187–189 of mixdom-wfst.tex), independent of $Z_{n-1}$ beyond what flows through $(d_n, f_n)$. Marginalising the chain joint over $Z_{1:n-1}$ at fixed $(d_n, f_n)$ leaves the $\pi(g_n, e_n \mid d_n, f_n)$ factor untouched, establishing $P(g_n, e_n \mid d_n, f_n, Z_{1:n-1}) = \pi(g_n, e_n \mid d_n, f_n)$. Substituting the explicit Bernoulli forms ($g_n \sim \text{Bern}(\rho^{(d_n)}_{f_n})$ and, given $g_n = 1$, $e_n \sim \text{Bern}(1 - \kappa_{d_n})$, with $e_n$ at $g_n = 0$ folded into the joint as the residual $\rho^{(d)}_f$ row) gives (C.141).

**Lemma 2 (Singlet route-decomposition).** *The marginal singlet emission $\omega^{(d,f,d',f')} := \sum_{g,e,g',e'} \widetilde{\sigma}((d, f, g, e) \to (d', f', g', e'))$ decomposes as $\omega = \omega^{(R1)} + \omega^{(R2)} + \omega^{(R3)}$, with the per-route weights of eq. (C.136).*

*Proof.* The singlet table mixdom-wfst.tex line 165–170 partitions the source $(g, e)$ values into three rows:

- Row $g$=0 (combining all source $e$ values, since the $g$=0 row weight does not depend on source $e$): table entry $\widetilde{\sigma}_{R1} = r^{(d)}_{f,f'}$ for destination $(d, f', g', e')$ summed over destination indicators.
- Row $g$=1, $e$=0: $\widetilde{\sigma}_{R2} = \rho^{(d)}_f \kappa_d w_{d,f'}$.
- Row $g$=1, $e$=1: $\widetilde{\sigma}_{R3} = \rho^{(d)}_f (1-\kappa_d)\kappa_0 v_{d'}\kappa_{d'} w_{d',f'}/(1-\zeta)$.

By the singlet-table convention, each row is the *joint* contribution including the source-$(g, e)$ prior; in particular $\widetilde{\sigma}_{R1} = \pi(g{=}0 \mid d, f) \cdot \sigma_{\text{cond}}((d, f', g'{=}\cdot, e'{=}\cdot) \mid (d, f, g{=}0, e))$ summed over destination $(g', e')$. Thus $\omega^{(r)} = \widetilde{\sigma}_r$ directly: the row weight already equals the route contribution to the marginal singlet emission, so no separate "prior cancellation" step is needed — the source prior was baked in from the start. Summing the three rows gives $\omega^{(d,f,d',f')}$ in (C.139). The fact that each row has support only on destinations consistent with its route is by inspection of the singlet table.

**Proposition (Reduced kernel).** *The reduced per-character marginal Pair HMM kernel,* $\hat{T}_{ss'}((d, f), (d', f')) := \sum_{g,e,g',e'} \pi(g, e \mid d, f) \cdot P((s', d', f', g', e') \mid (s, d, f, g, e))$, *satisfies the route-sum* (C.144).

*Proof.* Substitute the Pair HMM factorisation (C.140), and replace $\pi(g, e \mid d, f) \cdot \sigma_{\text{cond}}(\cdots) = \widetilde{\sigma}(\cdots)$ via the singlet-table convention:

$$\begin{aligned}\hat{T}_{ss'}((d, f), (d', f')) &= \sum_{g,e,g',e'} \pi(g, e \mid d, f)\, T^{\text{lab}}_{ss'}\big((d, f, g, e), (d', f', g', e')\big)\, \sigma_{\text{cond}}\big((d', f', g', e') \mid (d, f, g, e)\big) \\ &= \sum_{g,e,g',e'} T^{\text{lab}}_{ss'}\big((d, f, g, e), (d', f', g', e')\big)\, \widetilde{\sigma}((d, f, g, e) \to (d', f', g', e')).\end{aligned}$$

By Lemma 2, $\widetilde{\sigma}$ has support partitioned into the three routes $R1, R2, R3$, each with a unique source $(g_r, e_r)$. The labelled WFST entry for fixed source $(g_r, e_r)$ and structural transition $(s, s')$ depends on the destination $(g', e')$ in only two ways: (i) the label-preservation constraint on `M` (which forces $g' = g_{r,\text{dest}}$ and $e' = e_{r,\text{dest}}$ where the destination-side $(g_r, e_r)$ are determined by the route and the destination structural label, trivialising the destination sum); and (ii) for `I`, the destination prior on $(g', e')$ at the new $(d', f')$ is folded into the WFST as a $\pi(g', e' \mid d', f')$ factor (mixdom-wfst.tex eq. (C.183) gives the WFST = Pair HMM/singlet division explicitly). In both cases the $(g', e')$ summation cleanly factors out:

$$\sum_{g',e'} T^{\text{lab}}_{ss'}((d, f, g_r, e_r), (d', f', g', e'))\, \pi(g', e' \mid d', f')^{-1}\, \pi(g', e' \mid d', f') = \tilde{T}^{\text{lab},(r)}_{ss'}((d, f), (d', f'))$$

(the inverse-prior in the WFST cancels the prior in $\widetilde{\sigma}$'s destination factor; for `M` the inverse prior is trivially 1 because of label preservation). Combining with the source weights $\widetilde{\sigma}_r$ summed over destinations giving $\omega^{(r)}$ from Lemma 2, we obtain eq. (C.144).

**Corollary (Markov property of reduced chain).** *The marginal chain in* $(d, f)$ *obtained by summing* $(g, e)$ *out of the labelled singlet chain is exactly Markov, with per-step kernel* $\hat{T}_{\mathtt{Ss}}$ *at the reduced state space.*

*Proof.* We show $P(\tau_n \mid \tau_{1:n-1}) = P(\tau_n \mid \tau_{n-1})$ for all $n \geq 2$, which is the defining property of a Markov chain. By the singlet's chain factorisation, $P(\tau_n \mid Z_{1:n-1}) = \sigma_{\text{cond, marg}}(\tau_n \mid Z_{n-1})$, which depends on $Z_{n-1} = (\tau_{n-1}, g_{n-1}, e_{n-1})$. Marginalising $(g_{n-1}, e_{n-1})$ against its conditional distribution given $\tau_{1:n-1}$:

$$P(\tau_n \mid \tau_{1:n-1}) = \sum_{g_{n-1},e_{n-1}} P(g_{n-1}, e_{n-1} \mid \tau_{1:n-1})\, \sigma_{\text{cond, marg}}(\tau_n \mid \tau_{n-1}, g_{n-1}, e_{n-1}).$$

By Lemma 1 (Markov property of $(g, e)$), $P(g_{n-1}, e_{n-1} \mid \tau_{1:n-1}) = \pi(g_{n-1}, e_{n-1} \mid \tau_{n-1})$, depending only on $\tau_{n-1}$. Substituting:

$$P(\tau_n \mid \tau_{1:n-1}) = \sum_{g_{n-1},e_{n-1}} \pi(g_{n-1}, e_{n-1} \mid \tau_{n-1})\, \sigma_{\text{cond, marg}}(\tau_n \mid \tau_{n-1}, g_{n-1}, e_{n-1})$$

which is $\hat{T}_{\mathtt{S}s}$ (eq. (C.144) for the $s = s' =$ trivial-transition case, i.e. depending only on $(\tau_{n-1}, \tau_n)$). This depends only on $\tau_{n-1}$, establishing the Markov property. The product factorisation $P(\tau_1, \ldots, \tau_L) = \prod_n \hat{T}(\tau_{n-1}, \tau_n)$ follows by chain rule.

**Caveats.** Two technical conditions buried in the proof deserve explicit flagging:

1. *Independence of WFST weights from destination* $(g', e')$*:* the labelled WFST entries $T^{\text{lab}}_{ss'}((d, f, g, e), (d', f', g', e$ depend on the structural transition type (`M`/`I`/`D` at both ends) and on the source $(g, e)$ via the case split into mid-fragment / fragment-boundary / domain-boundary, but they do not depend on the destination $(g', e')$ except through the trivial label-preservation (`M` requires matching labels) constraint already implicit in $T^{\text{lab}}$. This is by inspection of the labelled WFST tables of mixdom-wfst.tex Section C.11.3.
2. *Boundary entries:* the proof above covers non-boundary positions $1 \leq n \leq L$ with $(s, s') \in \{\mathtt{M}, \mathtt{I}, \mathtt{D}\}$. The `S` and `E` row/column entries of $\hat{T}$ require separate derivation because the `S` source has no preceding $(g, e)$ to enumerate routes over, and the `E` destination requires the singlet's termination weight rather than a next-character emission. These are flagged as Section C.8.11, item I.2.

**Markov property of the reduced chain (consequence).** The latent $(g, e)$ at position $n$ depends only on $(d_n, f_n)$ and the model parameters (Lemma 1). Consequently, summing out $(g, e)$ at each position yields a marginal chain in $(d, f)$ that remains *exactly Markov*, with per-step transition kernel $\hat{T}$ defined below. This justifies the path-LL formulation in Section C.8.5 as exact (not a mean-field approximation) under the reduced state space.

**Reduced kernel as a route-sum.** Combining the route enumeration with the labelled WFST entries gives the reduced per-character kernel as a sum of route contributions:

$$\boxed{\hat{T}_{ss'}\big((d, f), (d', f');\, d, \theta\big) \;=\; \sum_{r \in \mathcal{R}((d,f),(d',f'))} \omega^{(r)}_{(d,f)\to(d',f')}\, \tilde{T}^{\text{lab},(r)}_{ss'}\big((d, f), (d', f');\, d, \theta\big),} \tag{C.144}$$

with route set $\mathcal{R}((d, f), (d', f')) = \{R3\}$ for $d' \neq d$ and $\{R1, R2, R3\}$ for $d' = d$, and per-route labelled WFST entry

$$\tilde{T}^{\text{lab},(r)}_{ss'}\big((d, f), (d', f');\, d, \theta\big) \;=\; \sum_{g',e'} T^{\text{lab}}_{ss'}\big((d, f, g_r, e_r), (d', f', g', e');\, d, \theta\big), \tag{C.145}$$

where $T^{\text{lab}}$ is the *conditional* labelled WFST of Section C.11.3, defined as the labelled Pair HMM joint divided by the labelled singlet emission, and $(g_r, e_r)$ is the source $(g, e)$ for route $r$: $(g_{R1}, e_{R1}) = (0, e)$ for any $e$ (since within-fragment transitions do not constrain $e$; the labelled-singlet table at mixdom-wfst.tex line 165 treats $e$ as a free variable in the $g$=0 row, so we adopt $(0, 0)$ as the canonical representative); $(g_{R2}, e_{R2}) = (1, 0)$; and $(g_{R3}, e_{R3}) = (1, 1)$. The product $\omega^{(r)} \cdot \tilde{T}^{\text{lab},(r)}_{ss'}$ then equals the per-route contribution to the labelled Pair HMM joint $(d, f) \to (d', f')$ summed over destination indicators $(g', e')$, so summing over routes recovers the marginal Pair HMM joint at the reduced state space; division of $T^{\text{lab}}$ by the singlet was applied per-route inside $\omega^{(r)}$ to avoid double-counting.

**When the route-sum collapses.** The previously-claimed factorisation $\hat{T} = \omega \cdot \tilde{T}^{\mathrm{lab}}$ holds only if, for the given $(s, s')$, the labelled WFST entry $\tilde{T}^{\mathrm{lab},(r)}_{ss'}$ is independent of $r$ across the enabled routes. This essentially never happens for the indel block. For $\mathtt{M} \to \mathtt{M}$, the per-route conditional WFST weights (Pair HMM joint divided by the route's singlet emission) are

$$W^{(R1)}_{\mathtt{MM}} = 1, \qquad W^{(R2)}_{\mathtt{MM}} = (1-\beta_d)\alpha_d, \qquad W^{(R3)}_{\mathtt{MM}} \;\propto\; \tau^{(d)}_{\mathtt{M,E}}\, \mathcal{T}^{[d\to d']}_{\mathtt{M,M}}\, \tau^{(d')}_{\mathtt{S,M}},$$

where $W^{(R3)}_{\mathtt{MM}}$ involves the destination-domain-specific top-level effective transition (in the implementation, `T_exit_k`; structurally, $\mathcal{T}^{[d\to d']}_{\mathtt{M,M}}$ is the destination-typed version of the empty-domain-summed top-level $\mathtt{M} \to \mathtt{M}$ kernel, with the $v_{d'}$ and other normalisation factors absorbed — see Section C.11.6). The three weights are distinct in general: R1 carries no separate BDI factor because in-fragment extension is generated by the singlet alone (the descendant character matches trivially via fragment continuation); R2 carries the standard new-fragment match factor $(1-\beta_d)\alpha_d$ (the singlet's $\kappa_d$ cancels into $\tau^{(d)}_{\mathtt{M,M}} = (1-\beta_d)\alpha_d\kappa_d$); R3 carries the cross-domain entry weight, involving both the source domain's exit $\tau^{(d)}_{\mathtt{M,E}} = (1-\beta_d)(1-\kappa_d)$ and the destination domain's TKF92 entry $\tau^{(d')}_{\mathtt{S,M}} = (1-\beta_{d'})\alpha_{d'}\kappa_{d'}$. Hence the route-sum genuinely cannot be collapsed to a single labelled-WFST entry for any $(d, f) \to (d', f')$ transition that admits more than one route, which includes every same-domain transition $(d, f) \to (d, f')$.

The reduced kernel is no more row-stochastic than the labelled one was — both are conditional transducers — but the input-conditional normalisation of the transducer is preserved by the route-sum marginalisation.

**Numerical verification.** The route-sum (C.144) has been verified at $t = 0.1$ on a $(N_{\mathrm{dom}}, N_{\mathrm{fr}}) = (2, 2)$ instance (random parameters, fixed seed) by reconstructing every $\mathtt{M} \to \mathtt{M}$ entry of the Pair HMM as $\sum_r \omega^{(r)}\, W^{(r)}_{\mathtt{MM}}$ and comparing against the corresponding entry of `build_nested_trans`; the maximum absolute discrepancy across the 16-entry $\mathtt{M} \to \mathtt{M}$ block is $1.1 \times 10^{-16}$, i.e. floating-point noise. The script is at `python/verify_reduced_wfst_routes.py`. The previous $t = 0$ check did not test the inter-route discrepancy in $W^{(r)}_{ss'}$ because at $t = 0$ the WFST collapses to (near-)identity and all routes coincide trivially.

##### C.8.5 Per-branch path log-likelihood

Read parent and child labelled states column-by-column on branch $v \to w$ to obtain the joint sequence $(Z^v_n, Z^w_n)_{n=1}^L$. By the same-tuple invariant of $q_n$, only five joint configurations carry positive variational mass, mapping to WFST states as in equation (B.95) (with $\tau$ now in the reduced $\mathcal{T}$):

$$S_n = \begin{cases} \mathtt{M} & (Z^v_n, Z^w_n) = (\tau, \tau), \\ \mathtt{D} & (Z^v_n, Z^w_n) = (\tau, xD), \\ \mathtt{I} & (Z^v_n, Z^w_n) = (xN, \tau), \\ \mathsf{Ig} & (Z^v_n, Z^w_n) \in \{(xN, xN), (xD, xD)\}. \end{cases} \tag{C.146}$$

With sentinels $S_0 = \mathtt{S}$, $S_{L+1} = \mathtt{E}$, the strip-Ignore reduction of equation (B.96) carries over:

$$\log P\big(X^w \mid X^v, d, \theta\big) \;=\; \sum_{N=1}^{L+1} \delta(S_N \neq \mathsf{Ig}) \sum_{M=0}^{N-1} \delta(S_M \neq \mathsf{Ig}) \log \hat{T}_{S_M S_N}\big(\tau_{n(M)},\, \tau_{n(N)};\, d, \theta\big) \prod_{K=M+1}^{N-1} \delta(S_K = \mathsf{Ig}), \tag{C.147}$$

with the reduced WFST $\hat{T}$ from equation (C.144) replacing the labelled $T^{\mathrm{lab}}$. Tuples are now in $\mathcal{T} = [N_{\mathrm{dom}}] \times [N_{\mathrm{fr}}]$.

##### C.8.6 Per-column expected indel log-likelihood under $q$

The variational column-state probabilities $P_q^{v\to w}(S_n = s)$ depend only on the presence/absence pattern through the inner $q_n^{(\pi|\tau)}$ (which is $\tau$-independent in our factorisation), so the same belief-propagation machinery from the simple case (equation (B.101)) computes them.

The per-branch expected indel log-likelihood factorises as

$$\mathbb{E}_q\big[\log P(X^w \mid X^v, d, \theta)\big] = \sum_{s,s'} \sum_{\tau,\tau' \in \mathcal{T}} W_{ss',\tau\tau'}^{(v\to w)} \log \hat{T}_{ss'}\big(\tau, \tau';\, d, \theta\big), \tag{C.148}$$

with the reduced expected transition counts

$$W_{ss',\tau\tau'}^{(v\to w)} \;=\; \sum_{N=1}^{L+1} \sum_{M=0}^{N-1} q_{n(M)}^{(\tau)}(\tau)\, q_{n(N)}^{(\tau)}(\tau')\, P_{q,M}^{v\to w}(s)\, P_{q,N}^{v\to w}(s') \prod_{K=M+1}^{N-1} P_{q,K}^{v\to w}(\mathsf{lg}), \tag{C.149}$$

where the per-column factorisation of $q^{(\tau)}$ has reduced the pairwise tuple weight to a product of per-column marginals (no chain forward-backward needed). The cumulant trick of Section B.6.8 lifts unchanged to the inner $K$-product, with stable computation in $O(L \cdot |s|^2 \cdot |\mathcal{T}|^2)$ per branch via the running-logaddexp prefix on $\log P_q^{v\to w}(\mathsf{lg})$. For $(N_{\mathrm{dom}}, N_{\mathrm{fr}}) = (3, 2)$ this gives $|\mathcal{T}|^2 = 36$ — versus $|\mathcal{T}^{\mathrm{lab}}|^2 = 57600$ in the labelled variant ($\sim 1600\times$ cheaper per branch).

##### C.8.7 Per-column expected substitution log-likelihood

For each column $n$, define the *Fitch subtree* $\mathcal{F}_n \subseteq \mathbf{t}$ as the smallest connected subtree containing all leaves $v$ with $X_n^v = 1$. By stationarity and reversibility of the substitution model, only nodes inside $\mathcal{F}_n$ contribute to the substitution likelihood; nodes outside are treated as missing data and absorbed into normalisation. The Fitch subtree is determined by the leaf data alone, independently of $q$ and $\tau$.

Because $c$ is a per-column model latent (one class per column, governing all branches at that column), the column-substitution likelihood under fragchar $f$ marginalises $c$ at the model level:

$$L_n^{(\mathrm{sub}),\mathrm{tot}}(f; \mathcal{F}_n) \;=\; \sum_{c=1}^{N_{\mathrm{cl}}} u_{f,c}\, L_n^{(\mathrm{sub})}(c; \mathcal{F}_n), \tag{C.150}$$

with

$$L_n^{(\mathrm{sub})}(c; \mathcal{F}_n) = \sum_{a \in \mathcal{A}} \pi_a^{(c)}\, \beta_n^{r_n}(a; c) \tag{C.151}$$

the standard Felsenstein up-pass likelihood at column $n$ on $\mathcal{F}_n$ under class-$c$'s rate matrix $R^{(c)}$ and stationary $\pi^{(c)}$ ($r_n$ is the Fitch-determined root of $\mathcal{F}_n$, $\beta_n^v(a; c)$ the standard Felsenstein up-message). The expected substitution log-likelihood under $q$ involves only the fragchar marginal $q_n^{(f)}(f) = \sum_d q_n^{(\tau)}(d, f)$:

$$\boxed{\mathbb{E}_{q_n^{(\tau)}}\big[\log L_n^{(\mathrm{sub}),\mathrm{tot}}(f_n; \mathcal{F}_n)\big] \;=\; \sum_{f=1}^{N_{\mathrm{fr}}} q_n^{(f)}(f)\, \log\Big[\sum_c u_{f,c}\, L_n^{(\mathrm{sub})}(c; \mathcal{F}_n)\Big].} \tag{C.152}$$

The presence factor $q_n^{(\pi|\tau)}$ does not enter — the substitution likelihood is fully determined by the (data-determined) Fitch subtree and the (variational-determined) fragchar marginal.

**Why $\log \sum_c uL$, not $\sum_c u \log L$.** This is the proper integration over the per-column class prior. A labelled-variant counterpart that carried $c$ as a variational latent would give $\sum_c q_n^{(c)}(c) \log L_n^{(\mathrm{sub})}(c)$; by Jensen's inequality $\log \sum_c p_c L_c \geq \sum_c p_c \log L_c$ with $p_c = u_{f,c}$, equality holding only when $q_n^{(c)}$ is at the Bayes-optimal class posterior $q^*(c) \propto u_{f,c} L_n^{(\mathrm{sub})}(c)$. The reduction analytically marginalises $c$ at the place where the labelled formulation left an inequality, so the reduced ELBO is at least as tight as the labelled ELBO and strictly tighter for any $q^{(c)}$ off the Bayes-optimal class posterior.

**Numerical implementation.** $\log L_n^{(\mathrm{sub}),\mathrm{tot}}(f)$ is computed in log-space:

$$\log L_n^{(\mathrm{sub}),\mathrm{tot}}(f) \;=\; \mathrm{logsumexp}_c\big(\log u_{f,c} + \log L_n^{(\mathrm{sub})}(c; \mathcal{F}_n)\big), \tag{C.153}$$

with the expected log under $q_n^{(f)}$ a simple weighted sum.

##### C.8.8 ELBO

Combining the four contributions:

$$\boxed{\log p(\mathrm{MSA} \mid \mathbf{t}, \theta) \;\geq\; \log \tilde{p}(\mathrm{MSA} \mid \mathbf{t}, \theta) \;\geq\; \mathcal{L}[q],} \tag{C.154}$$

with

$$\mathcal{L}[q] = \mathbb{E}_q\big[\log p_{\mathrm{singlet}}^{\mathrm{red}}(Z^{\mathrm{root}})\big] + \sum_{(v\to w)\in\mathbf{t}} \mathbb{E}_q\big[\log \hat{P}^{\mathrm{WFST}}(Z^w \mid Z^v, d_{vw}, \theta)\big] + \sum_{n=1}^{L} \sum_f q_n^{(f)}(f) \log L_n^{(\mathrm{sub}),\mathrm{tot}}(f; \mathcal{F}_n) + H[q]. \tag{C.155}$$

The first two terms use the reduced singlet HMM (defined on $\mathcal{T} = [N_{\mathrm{dom}}] \times [N_{\mathrm{fr}}]$ with the singlet kernel $\omega$ from equation (C.139)) and the reduced WFST $\hat{T}$ from equation (C.144). The third is the marginalised substitution term (C.152).

**Entropy decomposition.**

$$H[q] = \sum_{n=1}^{L} H[q_n^{(\tau)}] \;+\; \sum_{n=1}^{L} H[q_n^{(\pi|\tau)} \mid \mathrm{MSA}], \tag{C.156}$$

with $H[q_n^{(\tau)}]$ the simple categorical entropy (closed-form, $|\mathcal{T}| - 1$ free parameters per column) and $H[q_n^{(\pi|\tau)} \mid \mathrm{MSA}]$ the leaf-conditioned entropy of the simple-case 3-state graphical model (equations (B.109) and (B.110) of Section B.6.7, with the same $\log Z_q$ correction).

**Bound interpretation.** Identical to equation (B.111): the bound is on $\log p$ with gap $=$ $q$-independent restriction-gap (ghost-column histories) + variational KL. The reduction $(c, g, e)$-collapse introduces no additional restriction gap; the analytic marginalisations are exact at the model level, replacing two labelled-variant Jensen inequalities with equalities and yielding a strictly tighter bound at the same variational optimum.

##### C.8.9 Cross-column constraint vanishes

The labelled-variant cross-column constraint (Section M.8 of the labelled draft) forced $q^{(\tau)}$ to be a column-Markov chain to avoid hard-zero violations of structural rules ($d_{n+1} = d_n$ when $e_n = 0$, etc.). Under the $(g, e)$-marginalisation those rules disappear: every entry of $\omega^{(d,f,d',f')}$ is positive (assuming irreducibility of $r^{(d)}$ and positivity of $v, w, \rho^{(d)}, \kappa_0, \kappa_n$), so no $(\tau_n, \tau_{n+1})$ pair is hard-zero in $\hat{T}$ either. Per-column factorised $q^{(\tau)}(\tau_1, \ldots, \tau_L) = \prod_n q_n^{(\tau)}(\tau_n)$ puts positive mass everywhere on the model's support and $\mathbb{E}_q[\log p]$ is finite for any positive $q$.

A column-Markov $q^{(\tau)}$ remains an *optional refinement* — the true posterior on column-tuples is correlated across columns even after $(g, e)$-marginalisation (the singlet kernel is genuinely Markov on $(d, f)$ via $\omega$) — but the per-column factorised $q^{(\tau)}$ is sufficient for $\mathcal{L}[q]$ to be a finite proper bound, eliminating the $|\mathcal{T}|^2$ tuple-Markov parameters per column-pair that the labelled variant required.

##### C.8.10 Special cases and recovery

**Recovery of simple TreeVarAnc.** Setting $|\mathcal{T}| = 1$ (single domain, single fragchar) collapses $\hat{T}$ to the GGI-approximation WFST $\mathbb{T}'$ of Section A.3 and renders the substitution term constant, recovering Section B.6 verbatim.

**Single-class limit.** Setting $u_{f,c} = \delta_{c,c_0}$ for a fixed class $c_0$ gives $L_n^{(\text{sub}),\text{tot}}(f) = L_n^{(\text{sub})}(c_0; \mathcal{F}_n)$, and the substitution term reduces to the standard Felsenstein log-likelihood under class $c_0$.

**Single-fragchar-per-domain limit.** $N_{\text{fr}} = 1$ collapses $r^{(d)}$ to a scalar self-loop, the within-domain fragchar Markov reduces to a geometric extension, and the reduced model recovers a simpler scalar-extension form.

**Fragment-boundary inference caveat.** Were the labelled $g$ indicator a deterministic function of consecutive fragchar transitions ($g_n = \delta(f_n \neq f_{n+1})$ within the same domain — the special case where intra-fragment fragchar transitions are forbidden), the variational marginal would identify fragment boundaries directly. Under the reduced formulation the variational marginal $q_n^{(\tau)}(d, f)$ at each column gives the ancestral $(d, f)$ directly, but does *not* uniquely identify "fragment boundaries" in that restrictive sense: a fragchar transition $(d, f) \to (d, f')$ with $f' \neq f$ in the inferred ancestral trajectory may be either a within-fragment Markov move or a new-fragment start, depending on the latent route. For ancestral $(d, f)$ inference at observed columns this distinction is irrelevant. For downstream tasks that need fragment boundaries explicitly, one option is to augment $q^{(\tau)}$ post-hoc with a fragment-boundary indicator inferred from the route posterior over each $(d, f) \to (d, f')$ transition — but this is a derived quantity rather than an independent variational latent.

##### C.8.11 Open issues

Two points warrant explicit verification before relying on the construction quantitatively (a third was resolved by numerical verification at $t = 0$, see Section C.8.4).

**(I.1) $\hat{T}$ row-sum identity.** The reduced WFST $\hat{T}$ inherits the labelled WFST's input-conditional normalisation by construction (Section C.8.4), but we have not explicitly verified the row-sum identity. A small numerical check is recommended: for $(N_{\text{dom}}, N_{\text{fr}}) \in \{(2, 2), (3, 2)\}$, build $\hat{T}$ from the labelled $T^{\text{lab}}$ and verify that composition with the reduced singlet HMM on $(d, f)$ yields the same column-marginal Pair HMM probabilities as the labelled construction.

**(I.2) Start/end-row boundary effects.** Boundary entries $\hat{T}_{\mathtt{S}s'}(\mathtt{S},\tau')$ and $\hat{T}_{s\mathtt{E}}(\tau,\mathtt{E})$ involve the labelled WFST's start and end rows, which carry domain-level boundary indicators ($\mathcal{T}_{\mathtt{S}\cdot}$, $\mathcal{T}_{\cdot\mathtt{E}}$) that are not column-internal. These rows may need explicit derivation rather than mechanical application of equation (C.144); the construction in Section C.8.4 is correct for interior $(s,s') \in \{\mathtt{M},\mathtt{I},\mathtt{D}\}^2$ but should be verified at the boundaries.

These open issues are flagged here for transparency. Their resolution is a precondition for production use of the reduced ELBO in parameter learning; for ancestral-tuple inference at fixed $\theta$ the resulting bias (if any) is $q$-independent and so cancels from the variational optimum.

#### C.9 Generalized Phylo-HMM for MixDom

This appendix presents a polynomial-time algorithm for marginal ancestral reconstruction in a restricted MixDom model. The restriction is: the top-level TKF91 process has vanishing indel rates $\lambda_0, \mu_0 \to 0$ at fixed ratio $\kappa_0 = \lambda_0/\mu_0$. As input we assume an MSA with a tree and a per-node gap/residue annotation (in practice obtained by Fitch parsimony on gaps).

##### C.9.1 The Vanishing-Top-Level-Indel Limit

Setting $\lambda_0 = \mu_0 = 0$ literally produces $0/0$ indeterminate forms in the TKF91 transition probabilities (eq. A.4). We instead take the limit $\lambda_0, \mu_0 \to 0^+$ with $\kappa_0$ fixed. In this limit the TKF91 $\alpha, \beta, \gamma$ coefficients evaluated at a branch length $T > 0$ satisfy $\alpha = e^{-\mu_0 T} \to 1$, $\beta = \lambda_0(1-e^{-(\mu_0-\lambda_0)T})/(\mu_0 - \lambda_0 e^{-(\mu_0-\lambda_0)T}) \to 0$, and $\gamma \to 0$. Inspecting the entries of the $5\times5$ top-level matrix $\tau_0$:

$$\begin{aligned}
&\tau_0(\mathtt{S},\mathtt{M}) = (1-\beta)\kappa_0\alpha \to \kappa_0,\\
&\tau_0(\mathtt{S},\mathtt{I}) = \beta \to 0,\\
&\tau_0(\mathtt{S},\mathtt{D}) = (1-\beta)\kappa_0(1-\alpha) \to 0,\\
&\tau_0(\mathtt{S},\mathtt{E}) = (1-\beta)(1-\kappa_0) \to 1-\kappa_0,\\
&\tau_0(\mathtt{M},\mathtt{M}) \to \kappa_0, \quad \tau_0(\mathtt{M},\mathtt{I}), \tau_0(\mathtt{M},\mathtt{D}) \to 0,\\
&\tau_0(\mathtt{M},\mathtt{E}) \to 1-\kappa_0.
\end{aligned}$$

The $\mathtt{I}$ and $\mathtt{D}$ columns become structurally unreachable on every branch, while the $\mathtt{S} \to \mathtt{M} \to \cdots \to \mathtt{M} \to \mathtt{E}$ chain survives with probability $\kappa_0^{\mathcal{N}}(1-\kappa_0)$ for a chain of $\mathcal{N}$ top-level match states. In other words, the number of top-level domains $\mathcal{N}$ is distributed Geometric($\kappa_0$) at the root, and every descendant preserves the same ordered list of domains (no top-level births or deaths). The ratio $\kappa_0$ thus becomes the single top-level parameter that survives the limit; all branch-length dependence at the top level vanishes.

##### C.9.2 Partition Decomposition

Let the MSA have $L$ columns. Conditional on the number of top-level domains $\mathcal{N}_*$ and their classes $d_1, \ldots, d_{\mathcal{N}_*}$, the full model decomposes into $\mathcal{N}_*$ independent nested processes, each responsible for some contiguous subset of columns. Since there are no top-level births or deaths on any branch, each domain is either entirely absent or entirely present on each branch, and the correspondence between columns and domains is a single partition shared across the whole tree. Let

$$P = (b_1, b_2, \ldots, b_B), \qquad b_i = [l_{i-1}+1,\, l_i],$$

with $0 = l_0 < l_1 < \cdots < l_B = L$, be a partition of $\{1,\ldots,L\}$ into $B$ contiguous blocks, and let $d_{b_i} \in \{1,\ldots,\mathcal{N}\}$ be the class label of block $i$, where $\mathcal{N}$ is the number of top-level domain classes in the model. Then

$$P(\text{MSA} \mid \text{tree}, \text{model}) = \sum_P P(P \mid \text{model}) \prod_{i=1}^{B} G(l_{i-1}+1, l_i, d_{b_i}), \tag{C.157}$$

where the block likelihood $G(k, l, n)$ is the standalone phylogenetic likelihood of the sub-MSA on columns $k..l$ under the within-domain model for domain class $n$, now including the Markovian fragment process with transition matrix $r^{(n)}$ and per-fragment site class distributions $u_{nfc}$. The partition prior is

$$P(P \mid \text{model}) = (1-\kappa_0)\, \kappa_0^B \prod_{i=1}^{B} v_{d_{b_i}}, \tag{C.158}$$

where $v$ is the top-level class (stationary) distribution. When marginalising over $P$ the number of summands is exponential in $L$, so direct evaluation of (C.157) is intractable. It is, however, a generalised hidden Markov model in which each "emission" is an arbitrarily long contiguous block of columns.

##### C.9.3 Why the State Space Cannot Be Collapsed

A standard reduction of a generalised HMM to an ordinary HMM would introduce a hidden state at every column encoding "which within-domain Pair-HMM state each phylogenetic lineage is in". Under a single TKF92, each lineage has three relevant states (`M`, `I`, `D`), but the per-column state vector across $T$ tree nodes has $3^T$ values. Furthermore, once a new top-level domain begins the within-domain Pair HMMs on every lineage all reset to `S`, so the memory of the within-domain state on any previously gapped lineage is lost at every block boundary: a newly present lineage can only be reasoned about as starting from `S` at the first column of the new block. Conditioning on the partition $P$, on the other hand, makes each block a self-contained problem on a sub-MSA, and eliminates the combinatorial explosion at the price of a quadratic sum over start columns.

##### C.9.4 Setup and Definitions

We pretend there is an "infinitely long" branch above the root node, so every fragment on the root row is modeled as an insertion. We partition the alignment into blocks of domain type $n$.

**Column presence profile.** Let $A(j)$ denote the column presence/absence profile for column $j$—a binary vector with one entry per tree node, where 1 indicates the node has a residue at column $j$ and 0 indicates a gap (as determined by Fitch parsimony).

**Fragment continuity.** Define

$$k_{\min}(i, j) = \min\{k : i \le k \le j,\ A(k') = A(j) \text{ for all } k \le k' \le j\}$$

This is the first column in $i..j$ that can be part of the same fragment as column $j$ (i.e., the earliest column from which an unbroken run of identical presence profiles extends to $j$).

**Per-branch TKF state.** Let $S_{\text{tkf}}(r, i, j)$ be the TKF Pair HMM emit state for row (branch) $r$ at column $j$, given block start $i$. This is a deterministic function of the presence/absence pattern: if all entries are zero for columns $i..j$ on branch $r$, the state is `S`; otherwise, the state depends on (ancestor_present$_r(j)$, descendant_present$_r(j)$), mapping to `M`, `I`, or `D` as appropriate.

**Per-branch TKF transitions.** Let $B_{\text{tkf}}(n, r, s, s')$ be the TKF91 branch transition matrix entry for domain $n$, branch $r$, from state $s$ to state $s'$. Define

$$T_{\text{tkf}}(n, i, j) = \prod_r B_{\text{tkf}}(n, r, S_{\text{tkf}}(r, i, j-1),\ S_{\text{tkf}}(r, i, j))$$

as the product of TKF91 transitions across all rows from column $j-1$ to column $j$ within a block starting at column $i$. Similarly define

$$T_{\text{tkf,start}}(n, j) = \prod_r B_{\text{tkf}}(n, r, \mathtt{S},\ S_{\text{tkf}}(r, j, j))$$

for block-start transitions and

$$T_{\text{tkf,end}}(n, i, j) = \prod_r B_{\text{tkf}}(n, r, S_{\text{tkf}}(r, i, j),\ \mathtt{E})$$

for block-closing transitions.

**Felsenstein emission likelihood.** Let $U(j, c)$ be the Felsenstein pruning likelihood for column $j$ under site class $c$, computed over the present subtree at column $j$ with the substitution model $(Q^{(c)}, \pi^{(c)})$.

##### C.9.5 Intra-Block Forward Recurrence

Within a block assigned to domain $n$ spanning columns $i..j$, the Markovian fragment process induces a forward recurrence over fragment states.

**Transition probabilities.** Fragment-to-fragment transition (same fragment continues):

$$T_{\text{ext}}(n, j, f, g) = r_{fg}^{(n)}$$

This transition is only available when $A(j-1) = A(j)$ (the presence profile has not changed, so the same fragment can continue).

Fragment termination followed by new fragment start (the presence profile changes, or the Markov chain starts a new fragment):

$$T_{\text{notext}}(n, i, j, f, g) = \rho_f^{(n)} \cdot T_{\text{tkf}}(n, i, j) \cdot w_{ng}$$

**Emission weight.** Each column $j$ in fragment state $g$ contributes the class-averaged Felsenstein likelihood:

$$E(n, j, g) = \sum_{c=1}^{C} u_{ngc}\, U(j, c)$$

**Forward recurrence.** Define $F_{i,j,n,g}$ as the probability of columns $i..j$ within a block starting at $i$ in domain $n$, with column $j$ in fragment state $g$.

Base case:

$$F_{i,i,n,g} = T_{\text{tkf,start}}(n, i) \cdot w_{ng} \cdot E(n, i, g) \tag{C.159}$$

Recursion for $j > i$:

$$F_{i,j,n,g} = \sum_{f=1}^{\mathcal{F}} F_{i,j-1,n,f} \cdot \left(\delta(A(j-1){=}A(j))\, T_{\text{ext}}(n,j,f,g) + T_{\text{notext}}(n,i,j,f,g)\right) \cdot E(n,j,g) \quad \text{(C.160)}$$

where $\delta(A(j-1){=}A(j))$ restricts the fragment extension term to columns with identical presence profiles.

**Block likelihood.**

$$G(i,j,n) = \sum_{f=1}^{\mathcal{F}} F_{i,j,n,f} \cdot \rho_f^{(n)} \cdot T_{\text{tkf,end}}(n,i,j) \quad \text{(C.161)}$$

##### C.9.6 The Forward Recursion

Define the Forward quantity

$$F(l,n) = P(\text{columns } 1..l \text{ of MSA, with last block ending at } l \text{ in domain class } n \mid \text{tree}, \text{model}). \quad \text{(C.162)}$$

Encoding the partition prior (C.158) into the recursion, we initialise with a virtual start marker $F(0,\emptyset) = 1$ and advance by

$$F(l,n) = \kappa_0\, v_n \sum_{k=0}^{l-1} \bar{F}(k)\, G(k+1,l,n), \quad \text{(C.163)}$$

where $\bar{F}(0) := 1$ and $\bar{F}(k) := \sum_m F(k,m)$ for $k > 0$. The total data log-likelihood is obtained by closing off the final block with the top-level termination factor:

$$P(\text{MSA} \mid \text{tree}, \text{model}) = (1-\kappa_0) \sum_n F(L,n). \quad \text{(C.164)}$$

**Full MSA probability.** Given a specific partition into blocks $(i_1, j_1, n_1), \ldots, (i_B, j_B, n_B)$, the probability of the partitioned MSA given the tree is

$$P(\text{partitioned MSA} \mid \text{tree}) = (1-\kappa_0) \prod_{b=1}^{B} \kappa_0\, v_{n_b}\, G(i_b, j_b, n_b)$$

##### C.9.7 The Backward Recursion

Since the domain class of the *next* block does not depend on the class of the current block (the partition prior factorises over blocks), the backward variable can be collapsed to a scalar. Define

$$\bar{\beta}(l) = P(\text{columns } l{+}1..L \text{ of MSA} \mid \text{last block ended at column } l,\ \text{tree}, \text{model}). \quad \text{(C.165)}$$

The boundary condition and recursion are

$$\bar{\beta}(L) = 1-\kappa_0, \quad \text{(C.166)}$$

$$\bar{\beta}(k) = \sum_{l=k+1}^{L} \left[\kappa_0 \sum_n v_n\, G(k{+}1,l,n)\right] \bar{\beta}(l), \qquad k < L. \quad \text{(C.167)}$$

The total likelihood is recoverable as $\bar{\beta}(0)$—equivalent to (C.164) (since the $k{=}0$ case expands to $(1{-}\kappa_0) \sum_n \kappa_0 v_n \sum_{l=1}^{L} G(1,l,n)\, [\cdots]$, matching the forward expression).

##### C.9.8 Intra-Block Backward Recurrence

To compute posterior fragment state and site class probabilities, we need an intra-block backward recurrence. Define $B_{i,k,j,n,f}$ as the probability of columns $k..j$ given a block $i..j$ in domain $n$, with column $k{-}1$ in fragment state $f$.

Boundary condition:

$$B_{i,j+1,j,n,f} = \rho_f^{(n)} \cdot T_{\text{tkf,end}}(n,i,j) \tag{C.168}$$

(Note: this depends on $f$ through $\rho_f^{(n)}$, the probability that fragment $f$ terminates.)

Recursion for $k \le j$:

$$B_{i,k,j,n,f} = \sum_{g=1}^{\mathcal{F}} \left(\delta(A(k{-}1){=}A(k))\, T_{\text{ext}}(n,k,f,g) + T_{\text{notext}}(n,i,k,f,g)\right) \cdot E(n,k,g) \cdot B_{i,k+1,j,n,g} \tag{C.169}$$

##### C.9.9 Posterior Domain and Fragment State Assignment

From the inter-block forward $F$ and backward $\beta$, and the intra-block forward $F_{i,k,n,f}$ and backward $B_{i,k,j,n,f}$, we recover per-column posteriors.

**Posterior domain assignment.** Writing $Z = P(\text{MSA} \mid \text{tree}, \text{model})$,

$$P(c \text{ in block of class } n \mid \text{MSA}) \;=\; \frac{1}{Z}\,\kappa_0\, v_n \sum_{k=0}^{c-1} \sum_{l=c}^{L} \bar{F}(k)\, G(k{+}1,l,n)\, \bar{\beta}(l). \tag{C.170}$$

**Posterior fragment state.** The posterior probability that column $k$ is in fragment state $f$, given a block $i..j$ of domain $n$, is:

$$P(\text{col } k \text{ is frag } f \mid \text{block } i..j, n) = \frac{F_{i,k,n,f} \cdot B_{i,k+1,j,n,f}}{G(i,j,n)} \tag{C.171}$$

where $B_{i,k+1,j,n,f}$ uses $k{+}1$ because $B$ gives the probability of the *remaining* columns $k{+}1..j$ given fragment state $f$ at column $k$.

**Full unconditional fragment state posterior.** The full posterior for fragment state at column $c$, marginalised over all block placements, is

$$P(\text{col } c \text{ is fragtype } f \mid \text{MSA}) = \sum_n \sum_{i \le c} \sum_{j \ge c} P(\text{block } i..j,\, n \mid \text{MSA})\, \frac{F_{i,c,n,f}\, B_{i,c+1,j,n,f}}{G(i,j,n)}, \tag{C.172}$$

where the block–domain posterior is

$$P(\text{block } i..j,\, n \mid \text{MSA}) \;\propto\; \bar{F}(i{-}1)\, \kappa_0\, v_n\, G(i,j,n)\, \bar{\beta}(j). \tag{C.173}$$

**Posterior site class.** The posterior probability of site class $c$ at column $k$ is obtained by mixing over the fragment state posterior:

$$P(\text{class } c \text{ at col } k \mid \text{block } i..j, n) = \sum_f P(\text{col } k \text{ is frag } f \mid \text{block } i..j, n) \cdot \frac{u_{nfc}\, U(k,c)}{\sum_{c'} u_{nfc'}\, U(k,c')} \tag{C.174}$$

The full (unconditional) posterior over site class at column $k$ is obtained by further marginalizing over blocks and domain types using the inter-block posterior (C.170).

##### C.9.10 Root Residue Reconstruction

Given the posterior in (C.170) and (C.174), we obtain a posterior over root residues at column $c$ by mixing over the class assignment and re-using the Felsenstein pruning posterior under each class:

$$P(\text{root}_c = a \mid \text{MSA}) = \sum_n P(c \in n \mid \text{MSA}) \cdot \sum_c P(\text{class } c \text{ at } c \mid n) \cdot P(\text{root}_c = a \mid \text{MSA}, c \in c), \tag{C.175}$$

where the per-class conditional on the right uses Felsenstein pruning at column $c$ with the substitution model $(Q^{(c)}, \pi^{(c)})$. The MAP root sequence is obtained by taking $\text{argmax}_a$ of this posterior column-by-column; gaps at the root are determined by the presence annotation obtained from Fitch parsimony.

##### C.9.11 Why the Trick Fails with Top-Level Indels

If we retained nonzero top-level insertion and deletion rates we would also need to decide on which branch each top-level domain was born or died. Given only the presence pattern at each column, a block whose residues appear on only a single clade is consistent with both (i) an ancestrally present domain that was deleted on every other branch and (ii) a recent insertion. Conditioning on the partition no longer factorises the likelihood across blocks because the per-branch top-level state sequence is correlated across blocks. The algorithm above relies crucially on the absence of top-level births and deaths so that every block is a fully independent problem with a fresh `S` at every branch.

*Remark* C.15 (Relaxing the top-level constraint). The zero top-level indel-rate limit ($n_{\text{top,indel}} = 0$) can be partially relaxed by conditioning on domain presence/absence profiles. At $n_{\text{top,indel}} = 1$, we allow a single top-level domain insertion or deletion event on the tree, yielding $O(R)$ possible domain subtrees (where $R$ is the number of tree rows/edges). At $n_{\text{top,indel}} = 2$, there are $O(R^2)$ configurations, and so on. The probability of each truncation level is roughly exponential in $n_{\text{top,indel}}$ (controlled by $\lambda_0 + \mu_0$), so this provides a systematic expansion controlling the state space explosion. The leading term ($n_{\text{top,indel}} = 0$) is the algorithm presented here.

##### C.9.12 Complexity

Let $\mathcal{N}$ be the number of top-level domain classes, $\mathcal{F}$ the number of fragment types per domain, $C$ the number of site classes, $T$ the number of tree nodes, and $L$ the number of MSA columns.

- Computing all $G(k+1, l, n)$ by incremental forward updates requires, for each starting column $k+1$, a left-to-right pass over $l = k+1, \ldots, L$. Each column update involves $O(\mathcal{F}^2)$ work for the fragment Markov chain transitions and $O(\mathcal{F} \cdot C)$ for the emission weights, plus $O(T)$ for the TKF transition products. Total: $O(L^2\,(\mathcal{F}^2 + \mathcal{F} \cdot C + T)\,\mathcal{N})$.
- Computing $F$ and $\beta$ given $G$ takes $O(L^2\,\mathcal{N})$.
- Computing the per-column posteriors including fragment state requires the intra-block backward, making the total $O(L^3\,\mathcal{F}^2\,\mathcal{N})$ in the worst case (since $B_{i,k,j}$ depends on both $i$ and $j$ for each $k$).
- Each column likelihood $U(j, c)$ is computed once and cached as an $O(L \cdot C \cdot T \cdot |\mathcal{A}|)$ precomputation.

Total for the $G$ pass and inter-block DP: $O(L^2\,(\mathcal{F}^2 + T)\,\mathcal{N})$. The intra-block backward for posteriors adds $O(L^3\,\mathcal{F}^2\,\mathcal{N})$ due to the dependence of $B_{i,k,j,n,f}$ on both $i$ (block start) and $j$ (block end) for each interior column $k$.

The algorithm is naturally vectorisable: all $G$ updates for a fixed starting column can be computed in parallel across $\mathcal{N}$ domain classes and across tree branches.

#### C.10 Simulation from MixDom

To generate data from the MixDom model on a phylogenetic tree:

1. **Sample domain sequence.** Draw the number of top-level domains $\mathcal{N}_* \sim \text{Geometric}(\kappa_0)$. For each domain, draw its type $n_i \sim \text{Categorical}(v_1, \ldots, v_{\mathcal{N}})$.

2. **Sample fragment trajectory.** For each domain of type $n$, sample a fragment trajectory through the $\mathcal{F}$-state Markov chain. The initial fragment state is $f_1 \sim \text{Categorical}(w_{n1}, \ldots, w_{n\mathcal{F}})$. At each step, transition to fragment state $f_{k+1}$ with probability $r^{(n)}_{f_k, f_{k+1}}$, or terminate the domain with probability $\rho^{(n)}_{f_k} = 1 - \sum_g r^{(n)}_{f_k, g}$.

3. **Sample site class and root residue.** For each emitted site from fragment state $f$, draw site class $c \sim \text{Categorical}(u_{nf1}, \ldots, u_{nfC})$. Draw the root residue $a \sim \pi^{(c)}$.

4. **Evolve down the tree.** For each site, evolve the root residue down the phylogenetic tree using the substitution model $(Q^{(c)}, \pi^{(c)})$ at evolutionary time $T_r$ on each branch $r$: $b \sim P^{(c)}(T_r) \cdot e_a$, where $P^{(c)}(T_r) = \exp(R^{(c)} T_r)$. Apply the per-branch TKF91 indel process (with domain-specific parameters $\lambda_n, \mu_n$) to create insertions and deletions.

#### C.11 Labeled-MixDom Singlet HMM and WFST

In this section we define state machines for the MixDom model that foreground all latent state in the input/output alphabet, so they can be used directly in beam-search MSA, progressive reconstruction, and the variational ancestral-state framework of Section C.8 without algebraic distillation. We call these the *Labeled-MixDom Singlet HMM* and the *Labeled-MixDom WFST*, to distinguish them from the Maraschino-distilled order-1 HMMs and WFSTs.

*Remark* C.16 (Within-fragment fragchar dynamics). Each fragment in MixDom carries an intra-fragment Markov chain on fragchars: from a character with fragchar $f$ in a fragment of domain $n$, the next character (still in the same fragment) has fragchar $g$ with probability $r^{(n)}_{fg}$, and the fragment terminates at the current character with probability $\rho^{(n)}_f = 1 - \sum_g r^{(n)}_{fg}$. This is consistent with the exploded-MixDom spec (§C.3.2 of exploded-mixdom.tex), where $r^{(k)}_{fg}$ is the intra-fragment fragchar transition kernel. The TKF92 *fragment* concept is preserved ($g = 1$ marks the last character of a fragment).

- **Within a fragment ($g = 0$).** The next character is in the same fragment with fragchar $g$ chosen via $r^{(n)}_{fg}$; the destination $g$ may or may not equal the source $f$ (intra-fragment Markov, not self-loop only).
- **Fragment boundary ($g = 1$).** The current character is the last in its fragment; this happens with the fragment-termination weight $\rho^{(n)}_f$. After fragment termination, the TKF92 outer dynamics decide what happens next: with probability $\kappa_n$ another fragment in the same domain begins (its first character drawn from $w_{n,\cdot}$); with probability $1 - \kappa_n$ the domain ends ($e = 1$).
- **Domain termination ($e = 1$).** The TKF91 outer dynamics on domains (with parameter $\kappa_0$) decide whether another domain begins or the sequence ends.

- $\mathcal{F} = 1$ **special case.** Setting $N_{\mathcal{F}} = 1$ (single fragchar) collapses $r^{(n)}$ to a scalar self-loop; the resulting expressions reduce to a TKF92-extension scalar form that admits a more compact algebra (used in the simpler closed forms elsewhere in this paper).

##### C.11.1 Labeled Alphabet

Each alphabet symbol $a \in \mathcal{A}$ is decorated with a label tuple $(c, f, g, n, e)$ consisting of the following latent variables:

- $c \in \mathcal{C} = \{1, \dots, |\mathcal{C}|\}$: substitution site class.
- $f \in \mathcal{F} = \{1, \dots, |\mathcal{F}|\}$: fragchar of the current character (the per-character class label whose Markov chain within a fragment is $r^{(n)}$).
- $g \in \{0, 1\}$: fragment-end indicator ($g = 1$ if this is the last character of the current fragment; $g = 0$ otherwise).
- $n \in \mathcal{N} = \{1, \dots, |\mathcal{N}|\}$: domain type.
- $e \in \{0, 1\}$: domain-end indicator ($e = 1$ if this is the last fragment of the current domain).

We write $a_{cfgne}$ for a labeled symbol and define $\mathcal{L} = \mathcal{C} \times \mathcal{F} \times \{0, 1\} \times \mathcal{N} \times \{0, 1\}$, so $|\mathcal{L}| = 4|\mathcal{C}||\mathcal{F}||\mathcal{N}|$. The labeled alphabet has $|\mathcal{A}| \cdot |\mathcal{L}|$ symbols.

For context tracking in order-1 machines we need only the *structural label* $\ell = (f, g, n, e)$, since the site class $c$ does not affect transition structure. The number of distinct structural labels is $L = 4|\mathcal{F}||\mathcal{N}|$.

##### C.11.2 Labeled-MixDom Singlet HMM

The Labeled-MixDom Singlet HMM generates sequences from the MixDom stationary distribution with all latent variables made explicit. It is an order-1 HMM whose state records the structural label $\ell = (f, g, n, e)$ of the most recently emitted character.

**States.** The state space is $\{\mathtt{S}\} \cup \{(f, g, n, e) : f \in \mathcal{F},\ g \in \{0, 1\},\ n \in \mathcal{N},\ e \in \{0, 1\}\} \cup \{\mathtt{E}\}$, giving $L + 2$ states.

**Emissions.** In state $(f, g, n, e)$, the HMM emits symbol $a_{cfgne}$ (with the $f, g, n, e$ components matching the state) with probability

$$e(a_{cfgne} \mid f, g, n, e) = u_{fc}\, \pi_{ca} \tag{C.176}$$

summing to 1 over $(c, a)$.

**Transitions.** From the start state $\mathtt{S}$:

$$P(\mathtt{S} \to (f, g, n, e)) = v_n\, \kappa_n\, w_{nf} \begin{cases} 1 - \rho_f^{(n)} & \text{if } g = 0 \\ \rho_f^{(n)}\, \kappa_n\, w_{nf'} & \text{if } g = 1,\ e = 0 \\ & \quad \text{(to state } (f', g', n, e')\text{; see below)} \\ \rho_f^{(n)}\, (1 - \kappa_n)\, \kappa_0\, v_{n'}\, \kappa_{n'}\, w_{n'f'} & \text{if } g = 1,\ e = 1 \\ & \quad \text{(to state } (f', g', n', e')\text{; see below)} \end{cases} \tag{C.177}$$

However, since the singlet HMM is order-1 and we only track the *destination* state, it is cleaner to express the transitions directly.

The stationary distribution for the MixDom singlet process factors as follows. At the domain level, a TKF91 process with parameters $(\lambda_0, \mu_0)$ generates $N_{\text{dom}} \sim \text{Geom}(\kappa_0)$ domains, each of type $n \sim \text{Categorical}(v_1, \ldots, v_{\mathcal{N}})$. Within domain $n$, an irreducible Markov chain over fragchars with transition matrix $r^{(n)} \in [0,1]^{\mathcal{F}\times\mathcal{F}}$ and termination weights $\rho_f^{(n)} = 1 - \sum_g r_{fg}^{(n)}$ generates the sequence of per-character fragchars; the first character in the domain is drawn from the chain's initial distribution $w_{nf}$. Within each character's fragchar $f$, the site class is $c \sim u_{fc}$ and the residue is $a \sim \pi_c$.

| Source $\ell$ | Dest $\ell'$ | Transition weight |
|---|---|---|
| `S` | $(f', 0, n', e')$ | $\kappa_0\, v_{n'}\, w_{n'f'}$ |
| `S` | $(f', 1, n', e')$ | $\kappa_0\, v_{n'}\, w_{n'f'}$ |
| `S` | `E` | $1 - \kappa_0$ |
| *Mid-domain continuations:* | | |
| $(f, 0, n, e)$ | $(f', g', n, e)$ | $r_{ff'}^{(n)}$ |
| $(f, 1, n, 0)$ | $(f', g', n, e')$ | $\rho_f^{(n)} \cdot \kappa_n \cdot w_{nf'}$ |
| $(f, 1, n, 1)$ | $(f', g', n', e')$ | $\rho_f^{(n)} \cdot (1-\kappa_n) \cdot \kappa_0 \cdot v_{n'} \cdot \kappa_{n'} \cdot w_{n'f'} \,/\, (1-\zeta)$ |
| *Termination:* | | |
| $(f, 0, n, e)$ | `E` | 0 (cannot end mid-fragment) |
| $(f, 1, n, 0)$ | `E` | $\rho_f^{(n)} \cdot (1-\kappa_n) \cdot (1-\kappa_0) \,/\, (1-\zeta)$ |
| $(f, 1, n, 1)$ | `E` | $\rho_f^{(n)} \cdot (1-\kappa_n) \cdot (1-\kappa_0) \,/\, (1-\zeta)$ |

The $g'$ and $e'$ in the destination labels range over all values ($g' \in \{0,1\}$, $e' \in \{0,1\}$); the destination's $g'$ is set by the prior $P(g' = 1 | f', n') = \rho_{f'}^{(n')}$ (per-character fragment-termination probability under the intra-fragment Markov), and likewise $e'$ by the per-fragment domain-termination probability $1 - \kappa_{n'}$. The weights above are the joint contributions; conditional probabilities of next-state indicators are recovered by multiplying by the appropriate prior factor for $(g', e')$.

The $\zeta$ correction $\zeta = \kappa_0 \cdot z_0^{(\text{sing})}$ accounts for skipping over geometrically-many empty domains before the next emitting character or sequence termination, where $z_0^{(\text{sing})} = \sum_n v_n(1-\kappa_n)$ is the singlet-process probability of an empty domain (under per-domain TKF92 with parameter $\kappa_n$). Verification of the marginalised $(n, f)$-only kernel (Section C.8, equation (C.139)) against the existing MixDom Pair HMM implementation `build_nested_trans` at $t = 0$ confirms these weights to 5+ decimal places on a (n_dom=2, n_fr=2) test instance.

More precisely, the probability of ending the sequence from a domain boundary state involves summing over all possible runs of empty domains before termination. Let $\zeta = \kappa_0 \cdot z_0^{(\text{sing})}$ be the probability of generating an empty domain and continuing. Then from state $(f, 1, n, 1)$ (domain boundary):

$$P(\text{to } \texttt{E}) = \frac{1-\kappa_0}{1-\zeta} \tag{C.178}$$

and from state $(f, 1, n, 0)$ (fragment boundary, mid-domain):

$$P(\text{to } \texttt{E}) = (1-\kappa_n) \cdot \frac{1-\kappa_0}{1-\zeta} \tag{C.179}$$

Similarly, the transition from a fragment boundary to a new character must account for possibly skipping empty domains. From $(f, 1, n, 1)$:

$$P((f, 1, n, 1) \to (f', g', n', e')) = \frac{\kappa_0 \cdot v_{n'} \cdot \kappa_{n'} \cdot w_{n'f'}}{1 - \zeta} \tag{C.180}$$

accounting for the possibility that some intermediate domains were empty (the nonempty domain $n'$ is reached after a geometric number of empty-domain trials).

**Normalization check.** From state $(f, 1, n, 1)$, summing over all destinations including `E`:

$$\sum_{f',g',n',e'} \frac{\kappa_0 v_{n'} \kappa_{n'} w_{n'f'}}{1 - \zeta} + \frac{1 - \kappa_0}{1 - \zeta} = \frac{\kappa_0(1 - z_0^{(\text{sing})}) + 1 - \kappa_0}{1 - \zeta} = \frac{1 - \zeta}{1 - \zeta} = 1.$$

where we used $\sum_{n'} v_{n'} \kappa_{n'} = 1 - z_0^{(\text{sing})}$ and $\zeta = \kappa_0 z_0^{(\text{sing})}$. The sum over $g'$ and $e'$ is implicit in the fragment/domain generation that follows.

##### C.11.3 Labeled-MixDom WFST

*Remark* C.17 (WFST tables use the matrix-kernel notation). The transition tables below use the intra-fragment Markov kernel $r_{fg}^{(n)}$ (matrix indexed by source and destination fragchar) and its per-row termination $\rho_f^{(n)} = 1 - \sum_{f'} r_{ff'}^{(n)}$. Where a within-fragment self-loop appears, the relevant entry is the diagonal $r_{ff}^{(n)}$; where a new-fragchar-same-domain move appears, it is an off-diagonal $r_{ff'}^{(n)}$ for $f' \neq f$; the fragment-end weight is $\rho_f^{(n)}$.

The Labeled-MixDom WFST represents the conditional distribution of a descendant labeled sequence given an ancestral labeled sequence, separated by evolutionary time $T$. When composed with the Labeled-MixDom Singlet HMM (Section C.11.2), it must reproduce the MixDom Pair HMM (Section C.1.1).

*Remark* C.18 (Reduced kernel for variational inference). The full $(c, f, g, d, e)$ alphabet is what makes this WFST self-contained (no algebraic distillation, exact for beam search and progressive reconstruction). For the variational ancestral-state framework (Appendix C.8) the labelled WFST is marginalised analytically over $(c, c', g, e, g', e')$ to a reduced per-character kernel $\hat{T}_{ss'}((d, f), (d', f'); d, \theta)$ over just $(d, f)$, with $3N_{\text{dom}}N_{\text{fr}} + 2$ states. The marginalisation does *not* collapse to a single labelled-WFST entry: the per-character labelled transition $(d, f) \to (d', f')$ admits up to three latent routes (intra-fragment fragchar transition; new fragment, same domain; new domain that may equal $d$), and the source's $(g, e)$ has a non-trivial posterior over routes whenever $d' = d$. The reduced kernel is therefore a route-sum $\hat{T}_{ss'} = \sum_r \omega^{(r)} \tilde{T}_{ss'}^{\text{lab},(r)}$ (eq. (C.144)), which collapses to the cleaner form $\omega \cdot \tilde{T}^{\text{lab}}$ only when the per-route labelled WFST entries coincide — a degenerate special case of the trivial $\mathcal{F} = 1, \mathcal{N} = 1$ instance. Class marginalisation is trivial at the indel level since the WFST indel block is class-independent. So the labelled WFST defined here and the reduced kernel used in the variational appendix are the same object viewed through different state-space lenses, with the variational appendix providing the explicit route-decomposition.

**Design principles.** The WFST has two kinds of states:

- **Emitting states** ("unready"): `M`, $\mathrm{I_F}$, $\mathrm{I_D}$, $\mathrm{D_F}$, $\mathrm{D_D}$. These consume an input character, produce an output character, or both, then make a mandatory null transition to a Wait state.

- **Wait states** ("ready"): $W_M$, $W_{D_F}$, $W_{D_D}$. These inspect the boundary indicators $(g, e)$ of the current context and choose the next emitting state based on the hierarchical boundary structure. Wait states also handle end-of-sequence.

In addition there are the non-emitting `S` and `E` states.

**States.** Each emitting or wait state carries a structural label context $\ell = (f, g, n, e)$. The full state space is:

$$\{\mathtt{S}\} \cup \{(X, f, g, n, e) : X \in \{\mathtt{M}, I_F, I_D, D_F, D_D, W_M, W_{D_F}, W_{D_D}\}\} \cup \{\mathtt{E}\}$$

Not all combinations occur (see constraints below), but the upper bound is $8L + 2$ states.

**Context semantics.** The context $\ell = (f, g, n, e)$ on each state records:

- In `M` and $W_M$: the label of the most recent matched character (which is the same for both input and output, since match preserves labels).
- In $I_F$, $I_D$: the label of the most recent *output* character.
- In $D_F$, $D_D$, $W_{D_F}$, $W_{D_D}$: the label of the most recent *input* character.

Thus only one context tuple is needed per state.

**Emitting-State Transitions (Unready → Wait)** Every emitting state makes a mandatory null transition (no input, no output) to its corresponding wait state. These transitions carry the boundary-survival weights from the nested TKF92/TKF91 structure.

| Source | Dest | Weight | Input | Output |
|---|---|---|---|---|
| $(\mathtt{M}, f, g, n, e)$ | $(W_M, f, g, n, e)$ | $w_{\mathtt{M}}(g, e)$ | $\varepsilon$ | $\varepsilon$ |
| $(D_F, f, g, n, e)$ | $(W_{D_F}, f, g, n, e)$ | $w_{D_F}(g, e)$ | $\varepsilon$ | $\varepsilon$ |
| $(D_D, f, g, n, e)$ | $(W_{D_D}, f, g, n, e)$ | $w_{D_D}(g, e)$ | $\varepsilon$ | $\varepsilon$ |
| $(I_F, f, g, n, e)$ | $(W_M, f, g, n, e)$ | $w_{I_F}(g, e)$ | $\varepsilon$ | $\varepsilon$ |
| $(I_D, f, g, n, e)$ | $(W_M, f, g, n, e)$ | $w_{I_D}(g, e)$ | $\varepsilon$ | $\varepsilon$ |

Wait: the insert states need more careful treatment. An inserted fragment or domain is a complete sub-sequence generated by the descendant. The insert states $I_F$ and $I_D$ handle character-level emissions *within* an inserted fragment, and upon fragment/domain completion, control returns to the wait state that initiated the insertion. We therefore need to track whether we are inserting at the fragment level or domain level.

Let us reconsider the state structure more carefully.

**Revised State Structure** In the MixDom Pair HMM, the five state types `MM`, `MI`, `MD`, `II`, `DD` at each $(n, f)$ position represent:

- $\mathtt{MM}_{nf}$: ancestral and descendant both have a character (match/substitution)
- $\mathtt{MI}_{nf}$: descendant insertion within domain $n$, fragment $f$

- $\mathtt{MD}_{nf}$: ancestral deletion within domain $n$, fragment $f$
- $\mathtt{II}_{nf}$: insertion of an entire domain (both ancestor and descendant insert)
- $\mathtt{DD}_{nf}$: deletion of an entire domain (both ancestor and descendant delete)

The top-level states $\mathtt{M}, \mathtt{I}, \mathtt{D}$ refer to the *domain-level* TKF91 process, while the nested states $\mathtt{M}, \mathtt{I}, \mathtt{D}$ refer to the *fragment-level* TKF92 process within a domain.

For the WFST, we separate the domain-level and fragment-level indel processes:

| WFST State Type | Input | Output |
|---|---|---|
| $\mathtt{M}$ (Match) | $a_{cfgne}$ | $b_{cfgne}$ |
| $\mathrm{I_F}$ (Insert Fragment char) | $\varepsilon$ | $b_{cfgne}$ |
| $\mathrm{I_D}$ (Insert Domain char) | $\varepsilon$ | $b_{cfgne}$ |
| $\mathrm{D_F}$ (Delete Fragment char) | $a_{cfgne}$ | $\varepsilon$ |
| $\mathrm{D_D}$ (Delete Domain char) | $a_{cfgne}$ | $\varepsilon$ |
| $\mathrm{W_M}$ (Wait after Match) | $\varepsilon$ | $\varepsilon$ |
| $\mathrm{W_{D_F}}$ (Wait after Delete-Fragment) | $\varepsilon$ | $\varepsilon$ |
| $\mathrm{W_{D_D}}$ (Wait after Delete-Domain) | $\varepsilon$ | $\varepsilon$ |

**Key constraint: label preservation.** In a Match state, the WFST does not change the structural label: the input and output labels $(f, g, n, e)$ must be identical (though $c$ is also preserved and the character $a \to b$ may change via substitution). Insertions create *new* characters with new labels; deletions consume characters without producing output.

**Emitting to Wait Transitions** After emitting (or consuming), each emitting state transitions to its wait state. These null transitions carry weights that account for the fragment-extension and domain-continuation structure.

In the MixDom model, within a domain of type $n$, each fragment evolves under the intra-fragment Markov kernel $r^{(n)}_{fg}$. Within the fragment ($g = 0$), the character simply continues. At a fragment boundary ($g = 1$), the TKF92 process within the domain decides whether to start a new fragment or end the domain. At a domain boundary ($g = 1, e = 1$), the TKF91 domain-level process decides whether to start a new domain or end the sequence.

The weights on emitting $\to$ wait transitions are:

| Transition | Condition | Weight |
|---|---|---|
| $\mathtt{M} \to \mathrm{W_M}$ | $g = 0$ | 1 |
| $\mathtt{M} \to \mathrm{W_M}$ | $g = 1, e = 0$ | $(1 - \beta_n)$ |
| $\mathtt{M} \to \mathrm{W_M}$ | $g = 1, e = 1$ | $(1 - \beta_n)(1 - \beta_0)$ |
| $\mathrm{D_F} \to \mathrm{W_{D_F}}$ | $g = 0$ | 1 |
| $\mathrm{D_F} \to \mathrm{W_{D_F}}$ | $g = 1, e = 0$ | $(1 - \gamma_n)$ |
| $\mathrm{D_F} \to \mathrm{W_{D_F}}$ | $g = 1, e = 1$ | $(1 - \gamma_n)(1 - \beta_0)$ |
| $\mathrm{D_D} \to \mathrm{W_{D_D}}$ | $g = 0$ | 1 |
| $\mathrm{D_D} \to \mathrm{W_{D_D}}$ | $g = 1, e = 0$ | 1 |
| $\mathrm{D_D} \to \mathrm{W_{D_D}}$ | $g = 1, e = 1$ | $(1 - \gamma_0)$ |

where $\beta_n = \beta(\lambda_n, \mu_n, T)$, $\gamma_n = \gamma(\lambda_n, \mu_n, T)$, $\beta_0 = \beta(\lambda_0, \mu_0, T)$, $\gamma_0 = \gamma(\lambda_0, \mu_0, T)$.

The rationale: at $g = 0$ we are mid-fragment, so no boundary weight is needed. At $g = 1$ (fragment boundary), the TKF92 boundary weight $(1 - \beta)$ or $(1 - \gamma)$ applies. At $g = 1, e = 1$ (domain boundary), both the fragment boundary and the domain boundary weights apply. For $\mathrm{D_D}$, the domain is being deleted as a unit; within the deleted domain, fragment structure is irrelevant (all fragments are consumed), so the fragment-level weights are unity and only the domain-level weight $(1 - \gamma_0)$ applies at domain end.

**Insert states.** Insert states ($\mathrm{I_F}$, $\mathrm{I_D}$) represent characters being inserted in the descendant. An inserted fragment is a self-contained TKF92 fragment; an inserted domain is a self-contained TKF91 domain.

After emitting an inserted character, the insert state loops back to itself (fragment extension) or transitions to a wait state (fragment/domain termination). The fragment extension self-loop:

| Transition | Weight | Input | Output |
|---|---|---|---|
| $(\mathrm{I_F}, f, g, n, e) \to (\mathrm{I_F}, g, g', n, e)$ | $r^{(n)}_{fg}$ | $\varepsilon$ | $b_{cgg'ne}$ |
| $(\mathrm{I_D}, f, g, n, e) \to (\mathrm{I_D}, g, g', n, e)$ | $r^{(n)}_{fg}$ | $\varepsilon$ | $b_{cgg'ne}$ |

On fragment termination within an inserted domain ($\mathrm{I_D}$), a new fragment may begin (with TKF92 parameters for the inserted domain):

| Transition | Weight | Condition | I/O |
|---|---|---|---|
| $(\mathrm{I_D}, f, 1, n, 0) \to (\mathrm{I_D}, f', g', n, e')$ | $\rho^{(n)}_f \cdot \kappa_n \cdot w_{nf'}$ | new frag in inserted domain | $\varepsilon/b$ |
| $(\mathrm{I_D}, f, 1, n, e) \to (\mathrm{W_M}, \ldots)$ | $\rho^{(n)}_f (1 - \kappa_n)$ | $e$ set appropriately | $\varepsilon/\varepsilon$ |

For $\mathrm{I_F}$ (inserted fragment within an existing domain), fragment termination returns control to $\mathrm{W_M}$:

| Transition | Weight | Condition |
|---|---|---|
| $(\mathrm{I_F}, f, 1, n, e) \to (\mathrm{W_M}, f, 1, n, e)$ | $\rho^{(n)}_f$ | fragment ends |

**Wait-State Transitions (Ready → Emitting)** Wait states inspect the boundary indicators and decide the next action. The transitions depend on whether we are mid-fragment ($g = 0$), at a fragment boundary ($g = 1, e = 0$), or at a domain boundary ($g = 1, e = 1$).

**Case 1: Mid-fragment ($g = 0$).** Within a fragment, only continuation of the current fragment is possible. No new fragments or domains can start.

| Source | Dest | Weight | Input | Output |
|---|---|---|---|---|
| $(\mathrm{W_M}, f, 0, n, e)$ | $(\mathtt{M}, f, g', n, e)$ | $\alpha_n$ | $a_{cfg'ne}$ | $b_{cfg'ne}$ |
| $(\mathrm{W_M}, f, 0, n, e)$ | $(\mathrm{D_F}, f, g', n, e)$ | $(1 - \alpha_n)$ | $a_{cfg'ne}$ | $\varepsilon$ |
| $(\mathrm{W_{D_F}}, f, 0, n, e)$ | $(\mathrm{D_F}, f, g', n, e)$ | 1 | $a_{cfg'ne}$ | $\varepsilon$ |
| $(\mathrm{W_{D_D}}, f, 0, n, e)$ | $(\mathrm{D_D}, f, g', n, e)$ | 1 | $a_{cfg'ne}$ | $\varepsilon$ |

Here $g'$ can be 0 or 1 (determined by the input character's label), $\alpha_n = \alpha(\lambda_n, \mu_n, T)$, and the fragment type $f$ and domain indicators $(n, e)$ are unchanged. The emission weight for $\mathtt{M}$ is $\exp(R_c T)_{ab}$; the emission weight for $\mathrm{D_F}$ is 1 (input consumed, no output).

**Case 2: Fragment boundary, mid-domain ($g = 1, e = 0$).** At a fragment boundary within a domain, the TKF92 process within the domain decides: start a new fragment (match or delete), or insert a new fragment.

| Source | Dest | Weight | Input | Output |
|---|---|---|---|---|
| $(\mathrm{W_M}, f, 1, n, 0)$ | $(\mathtt{M}, f', g', n, e')$ | $\alpha_n \cdot w_{nf'}$ | $a_{c'f'g'ne'}$ | $b_{c'f'g'ne'}$ |
| $(\mathrm{W_M}, f, 1, n, 0)$ | $(\mathrm{D_F}, f', g', n, e')$ | $(1 - \alpha_n) \cdot w_{nf'}$ | $a_{c'f'g'ne'}$ | $\varepsilon$ |
| $(\mathrm{W_M}, f, 1, n, 0)$ | $(\mathrm{I_F}, f', g', n, 0)$ | $\beta_n \cdot w_{nf'}$ | $\varepsilon$ | $b_{c'f'g'n0}$ |
| $(\mathrm{W_M}, f, 1, n, 0)$ | $\mathtt{E}$ | $(1 - \kappa_n) \cdot (1 - \kappa_0)$ | $\varepsilon$ | $\varepsilon$ |
| $(\mathrm{W_{D_F}}, f, 1, n, 0)$ | $(\mathtt{M}, f', g', n, e')$ | $\alpha_n \cdot w_{nf'}$ | $a_{c'f'g'ne'}$ | $b_{c'f'g'ne'}$ |
| $(\mathrm{W_{D_F}}, f, 1, n, 0)$ | $(\mathrm{D_F}, f', g', n, e')$ | $(1 - \alpha_n) \cdot w_{nf'}$ | $a_{c'f'g'ne'}$ | $\varepsilon$ |
| $(\mathrm{W_{D_D}}, f, 1, n, 0)$ | $(\mathrm{D_D}, f', g', n, e')$ | $w_{nf'}$ | $a_{c'f'g'ne'}$ | $\varepsilon$ |

Note: in this case, a new fragment type $f'$ is drawn from $w_{nf'}$, and $e'$ is determined by the input character's label. The domain type $n$ is unchanged.

**Normalization of $\mathrm{W_M}$ at fragment boundary ($g = 1, e = 0$).** The outgoing weights from $(\mathrm{W_M}, f, 1, n, 0)$ must sum to 1 when we include all possible input/output characters:

- With an input character present (ancestral fragment continues): $\alpha_n + (1 - \alpha_n) = 1$, weighted by $w_{nf'}$. But the input character determines $f'$, so the $w$ weight is a prior for the singlet composition, not a transition weight in the WFST.
- With no input character: insertion weight $\beta_n$ or end weight.

Actually, for a WFST, the normalization is more subtle: the transducer weights need not sum to 1 at every state, because the transducer represents a conditional distribution. However, when composed with the singlet HMM, the resulting Pair HMM transitions must be properly normalized.

**Global vs. local normalization.** The Labeled-MixDom WFST is constructed by dividing the row-stochastic Pair HMM $\chi$ (Section C.1.1) by the row-stochastic Singlet HMM (Section C.11.2). By construction, Singlet $\circ$ WFST = Pair HMM, so the WFST is conditionally normalized in the global sense: $\sum_{\text{output sequences } y} P(y \mid x, \theta, T) = 1$ for any ancestor sequence $x$. However, the per-state outgoing weights from $\mathtt{M}, \mathrm{I_F}, \mathrm{I_D}, \mathrm{D_F}, \mathrm{D_D}$ do *not* sum to 1 over a fixed input symbol, even when the Singlet emission factor for the destination label is included. This is the same state-folding artifact discussed for the TKF92 WFST in Appendix A.3.7: the Bernoulli-$r$ extension-vs-exit decision (here, the $r_{fg}^{(n)}$ vs. $\rho_f^{(n)}$ event), and the $\kappa_n$ vs. $1-\kappa_n$ domain-continuation event, have been compiled into the same $\mathtt{M} \to \cdot$ and $\mathrm{I}. \to \cdot$ edges that already carry the destination-singlet factors. Splitting each emitting state into a "just-arrived" and a "decision" state would restore local stochasticity at the cost of doubling (or tripling) the state graph. The compact form here trades local stochasticity for a smaller machine, exactly as in TKF92.

Let us instead directly specify the transition weights that, when composed with the Singlet HMM, reproduce the Pair HMM transition matrix $\chi$ from Section C.1.1.

**Complete WFST Transition Table** We now give the complete transition table. For readability, we abbreviate the state labels and split by source wait-state type and boundary case.

Let the following shorthand apply throughout:

$$\begin{aligned} \alpha_n &= \alpha(\lambda_n, \mu_n, T), & \beta_n &= \beta(\lambda_n, \mu_n, T), & \gamma_n &= \gamma(\lambda_n, \mu_n, T) \\ \alpha_0 &= \alpha(\lambda_0, \mu_0, T), & \beta_0 &= \beta(\lambda_0, \mu_0, T), & \gamma_0 &= \gamma(\lambda_0, \mu_0, T) \\ \kappa_n &= \lambda_n/\mu_n, & \kappa_0 &= \lambda_0/\mu_0 \end{aligned}$$

and recall $\mathcal{T}$ is the effective $5 \times 5$ domain-level transition matrix with null states eliminated (Section C.1.1).

**Start transitions.** From $\mathtt{S}$, the WFST enters its first emitting state. The weights mirror the first row of $\chi$:

| Dest | Weight | Input | Output |
|---|---|---|---|
| $(\mathtt{M}, f', g', n', e')$ | $\mathcal{T}_{\mathtt{SM}} \cdot v_{n'} \cdot \tau_{\mathtt{SY}}^{(n')} \cdot w_{n'f'}$ | $a_{c'f'g'n'e'}$ | $b_{c'f'g'n'e'}$ |
| $(\mathrm{I_D}, f', g', n', e')$ | $\mathcal{T}_{\mathtt{SI}} \cdot v_{n'} \cdot \kappa_{n'} \cdot w_{n'f'}$ | $\varepsilon$ | $b_{c'f'g'n'e'}$ |
| $(\mathrm{D_D}, f', g', n', e')$ | $\mathcal{T}_{\mathtt{SD}} \cdot v_{n'} \cdot \kappa_{n'} \cdot w_{n'f'}$ | $a_{c'f'g'n'e'}$ | $\varepsilon$ |
| $\mathtt{E}$ | $\mathcal{T}_{\mathtt{SE}}$ | $\varepsilon$ | $\varepsilon$ |

where $\mathtt{Y}$ is determined by the nested state type of the destination ($\mathtt{M}$ for Match, $\mathtt{I}$ for Insert-Fragment, $\mathtt{D}$ for Delete-Fragment within the domain). For Match destinations that enter at the first character of a domain, $\tau_{\mathtt{SY}}^{(n')}$ is the TKF92 Pair HMM transition from $\mathtt{S}$ into the appropriate nested state.

**Emitting to Wait transitions.** After each emitting state, a null transition to the corresponding wait state occurs. The weight depends on the boundary indicators:

| Transition | Condition on $(g, e)$ | Weight |
| --- | --- | --- |
| | $g = 0$ | 1 |
| $(\mathtt{M}, \ell) \to (\mathrm{W}_{\mathrm{M}}, \ell)$ | $g = 1,\, e = 0$ | $\rho_f^{(n)} \cdot 1$ |
| | $g = 1,\, e = 1$ | $\rho_f^{(n)} \cdot 1$ |
| | $g = 0$ | 1 |
| $(\mathrm{D}_{\mathrm{F}}, \ell) \to (\mathrm{W}_{\mathrm{D}_{\mathrm{F}}}, \ell)$ | $g = 1,\, e = 0$ | $\rho_f^{(n)}$ |
| | $g = 1,\, e = 1$ | $\rho_f^{(n)}$ |
| | $g = 0$ | 1 |
| $(\mathrm{D}_{\mathrm{D}}, \ell) \to (\mathrm{W}_{\mathrm{D}_{\mathrm{D}}}, \ell)$ | $g = 1,\, e = 0$ | $\rho_f^{(n)}$ |
| | $g = 1,\, e = 1$ | $\rho_f^{(n)}$ |

Wait—the fragment extension must also be handled. At $g = 0$, the character is mid-fragment; the next character in the same fragment follows with certainty (the matrix entry $r_{fg}^{(n)}$ is already accounted for in the singlet HMM emission of the next labelled character). At $g = 1$, the fragment has ended, so the per-source termination weight $\rho_f^{(n)}$ has already been "spent" by the fact that $g = 1$ was observed. Since the boundary indicators are part of the *label on the character*, which is determined by the input (for ancestral) or output (for descendant), the fragment extension probability is absorbed into the singlet HMM, not the WFST.

Therefore all emitting-to-wait transitions have **unit weight**:

$$w(X \to W_X) = 1 \quad \text{for all } X \in \{\mathtt{M}, \mathrm{D}_{\mathrm{F}}, \mathrm{D}_{\mathrm{D}}\} \tag{C.181}$$

The fragment and domain boundary structure is encoded in the *wait-state outgoing transitions*, which condition on $(g, e)$.

**Wait-State Outgoing Transitions** The wait states make all structural decisions. We organize by source state type and boundary case.

$\mathrm{W}_{\mathrm{M}}$ **(Wait after Match). Case $g = 0$ (mid-fragment):** Continue the current fragment. The next input character must have the same $(f, n, e)$.

| Dest | Weight | Input | Output | Notes |
| --- | --- | --- | --- | --- |
| $(\mathtt{M}, f, g', n, e)$ | $\alpha_n$ | $a_{c'fg'ne}$ | $b_{c'fg'ne}$ | match continues |
| $(\mathrm{D}_{\mathrm{F}}, f, g', n, e)$ | $(1 - \alpha_n)$ | $a_{c'fg'ne}$ | $\varepsilon$ | fragment-level deletion |

**Case $g = 1, e = 0$ (fragment boundary, mid-domain):** Fragment ended, new fragment within same domain, or insert/delete fragment, or end domain and transition at domain level.

| Dest | Weight | I | O | Notes |
| --- | --- | --- | --- | --- |
| $(\mathtt{M}, f', g', n, e')$ | $\tau_{\mathtt{MM}}^{(n)} \cdot w_{nf'}$ | $a_{\dots}$ | $b_{\dots}$ | new frag, match |
| $(\mathrm{D}_{\mathrm{F}}, f', g', n, e')$ | $\tau_{\mathtt{MD}}^{(n)} \cdot w_{nf'}$ | $a_{\dots}$ | $\varepsilon$ | new frag, delete |
| $(\mathrm{I}_{\mathrm{F}}, f', g', n, 0)$ | $\tau_{\mathtt{MI}}^{(n)} \cdot w_{nf'}$ | $\varepsilon$ | $b_{\dots}$ | insert frag |
| $\mathtt{E}_{\mathrm{domain}}$ | $\tau_{\mathtt{ME}}^{(n)}$ | | | domain ends (see below) |

When the domain ends, $\tau^{(n)}_{\texttt{ME}} = (1-\beta_n)(1-\kappa_n)$, the domain-level process takes over. This is where the $\mathcal{T}$ matrix from the null-eliminated domain-level Pair HMM applies.

Rather than introducing an intermediate "domain end" state, we can fold the domain-level transition into the fragment-boundary transitions. From $(\mathrm{W_M}, f, 1, n, 0)$, with the domain ending: the TKF92 weight is $\tau^{(n)}_{\texttt{ME}}$, and then the domain-level $\mathcal{T}_{\texttt{M}\cdot}$ applies to reach the next domain.

However, this conflates two levels of the hierarchy. To keep the WFST clean and composable, we should handle this through the domain-end indicator $e$: when $e = 0$, we are mid-domain and only fragment-level transitions apply. The domain-end case $e = 1$ is reached when the last fragment of a domain has ended.

Actually, let us reconsider. The labels $(g, e)$ on the input/output characters tell us the hierarchical position. The *singlet HMM* generates these labels according to the MixDom stationary process. The WFST must respect them: it cannot change $(g, e)$ on matched characters.

So the WFST sees:

- Characters labeled $g = 0$: mid-fragment
- Characters labeled $g = 1, e = 0$: end of fragment, not end of domain
- Characters labeled $g = 1, e = 1$: end of fragment and end of domain

At a fragment boundary in the ancestor ($g = 1$), the WFST knows the ancestral fragment has ended. The next ancestral character (if any) will start a new fragment or domain. Between the end of one fragment and the start of the next, the WFST may insert new fragments (for $\mathrm{I_F}$) or entire domains (for $\mathrm{I_D}$).

This gives us the complete transition logic from each wait state.

**Revised:** $\mathrm{W_M}$ **at** $g = 1, e = 0$ **(fragment boundary, mid-domain):**

| Dest | Weight | I/O | Notes |
|---|---|---|---|
| $(\texttt{M}, f', g', n, e')$ | $\alpha_n \cdot w_{nf'}$ | $a/b$ | new fragment, match |
| $(\mathrm{D_F}, f', g', n, e')$ | $(1-\alpha_n) \cdot w_{nf'}$ | $a/\varepsilon$ | new fragment, delete |
| $(\mathrm{I_F}, f', g', n, 0)$ | $\beta_n \cdot w_{nf'}$ | $\varepsilon/b$ | insert fragment |

Normalization: with an input character, the weight is $\alpha_n + (1-\alpha_n) = 1$ (times $w$). Without an input character (insertion), $\beta_n$ (times $w$). These don't need to sum to 1 together because input-present and input-absent are exclusive events in the WFST.

$\mathrm{W_M}$ **at** $g = 1, e = 1$ **(domain boundary):**

Here both the fragment and domain have ended. The domain-level TKF91 process decides what happens next.

| Dest | Weight | I/O | Notes |
|---|---|---|---|
| $(\texttt{M}, f', g', n', e')$ | $\mathcal{T}_{\texttt{MM}} \cdot v_{n'} \cdot \tau^{(n')}_{\texttt{SM}} \cdot w_{n'f'}$ | $a/b$ | new domain, match |
| $(\mathrm{D_F}, f', g', n', e')$ | $\mathcal{T}_{\texttt{MM}} \cdot v_{n'} \cdot \tau^{(n')}_{\texttt{SD}} \cdot w_{n'f'}$ | $a/\varepsilon$ | new domain, del-frag |
| $(\mathrm{D_D}, f', g', n', e')$ | $\mathcal{T}_{\texttt{MD}} \cdot v_{n'} \cdot \kappa_{n'} \cdot w_{n'f'}$ | $a/\varepsilon$ | delete domain |
| $(\mathrm{I_D}, f', g', n', e')$ | $\mathcal{T}_{\texttt{MI}} \cdot v_{n'} \cdot \kappa_{n'} \cdot w_{n'f'}$ | $\varepsilon/b$ | insert domain |
| $\texttt{E}$ | $\mathcal{T}_{\texttt{ME}}$ | $\varepsilon/\varepsilon$ | end |

Here $\mathcal{T}_{\mathtt{M}\cdot}$ are the null-eliminated domain-level transitions. The $\tau^{(n')}_{\mathtt{SM}}$ and $\tau^{(n')}_{\mathtt{SD}}$ factors are the TKF92 entry transitions into the new domain $n'$.

**$\mathrm{W}_{\mathrm{D_F}}$ (Wait after Delete-Fragment).**

$\mathrm{W}_{\mathrm{D_F}}$ tracks the deletion of individual fragments within a domain (the domain itself is matched/surviving; only some fragments are deleted).

Case $g = 0$ (mid-fragment):

| Dest | Weight | I/O | Notes |
|---|---|---|---|
| $(\mathrm{D_F}, f, g', n, e)$ | 1 | $a/\varepsilon$ | continue deleting fragment |

Case $g = 1, e = 0$ (fragment boundary, mid-domain):

| Dest | Weight | I/O | Notes |
|---|---|---|---|
| $(\mathtt{M}, f', g', n, e')$ | $\alpha_n \cdot w_{nf'}$ | $a/b$ | new frag, match |
| $(\mathrm{D_F}, f', g', n, e')$ | $(1 - \alpha_n) \cdot w_{nf'}$ | $a/\varepsilon$ | new frag, delete |

Case $g = 1, e = 1$ (domain boundary):

| Dest | Weight | I/O | Notes |
|---|---|---|---|
| $(\mathtt{M}, f', g', n', e')$ | $\mathcal{T}_{\mathtt{MM}} \cdot v_{n'} \cdot \tau^{(n')}_{\mathtt{SM}} \cdot w_{n'f'}$ | $a/b$ | new domain, match |
| $(\mathrm{D_F}, f', g', n', e')$ | $\mathcal{T}_{\mathtt{MM}} \cdot v_{n'} \cdot \tau^{(n')}_{\mathtt{SD}} \cdot w_{n'f'}$ | $a/\varepsilon$ | new domain, del-frag |
| $(\mathrm{D_D}, f', g', n', e')$ | $\mathcal{T}_{\mathtt{MD}} \cdot v_{n'} \cdot \kappa_{n'} \cdot w_{n'f'}$ | $a/\varepsilon$ | delete domain |
| $(\mathrm{I_D}, f', g', n', e')$ | $\mathcal{T}_{\mathtt{MI}} \cdot v_{n'} \cdot \kappa_{n'} \cdot w_{n'f'}$ | $\varepsilon/b$ | insert domain |
| $\mathtt{E}$ | $\mathcal{T}_{\mathtt{ME}}$ | $\varepsilon/\varepsilon$ | end |

Note: $\mathrm{W}_{\mathrm{D_F}}$ at domain boundaries uses the *same* $\mathcal{T}_{\mathtt{M}\cdot}$ row as $\mathrm{W_M}$, because fragment-level deletion within a domain does not affect the domain-level state (the domain was matched, i.e. the top-level state was $\mathtt{M}$).

**$\mathrm{W}_{\mathrm{D_D}}$ (Wait after Delete-Domain).**

$\mathrm{W}_{\mathrm{D_D}}$ handles the deletion of entire domains. Within a deleted domain, all fragments are consumed without output.

Case $g = 0$ (mid-fragment):

| Dest | Weight | I/O | Notes |
|---|---|---|---|
| $(\mathrm{D_D}, f, g', n, e)$ | 1 | $a/\varepsilon$ | continue consuming domain |

Case $g = 1, e = 0$ (fragment boundary, mid-domain):

| Dest | Weight | I/O | Notes |
|---|---|---|---|
| $(\mathrm{D_D}, f', g', n, e')$ | $w_{nf'}$ | $a/\varepsilon$ | next fragment in deleted domain |

Note: within a deleted domain, the entire domain is being consumed, so the fragment distribution weight $w_{nf'}$ is needed to match the singlet prior.

Case $g = 1, e = 1$ (domain boundary):

| Dest | Weight | I/O | Notes |
|---|---|---|---|
| $(\mathtt{M}, f', g', n', e')$ | $\mathcal{T}_{\mathtt{DM}} \cdot v_{n'} \cdot \tau^{(n')}_{\mathtt{SM}} \cdot w_{n'f'}$ | $a/b$ | new domain, match |
| $(\mathrm{D_F}, f', g', n', e')$ | $\mathcal{T}_{\mathtt{DM}} \cdot v_{n'} \cdot \tau^{(n')}_{\mathtt{SD}} \cdot w_{n'f'}$ | $a/\varepsilon$ | new domain, del-frag |
| $(\mathrm{D_D}, f', g', n', e')$ | $\mathcal{T}_{\mathtt{DD}} \cdot v_{n'} \cdot \kappa_{n'} \cdot w_{n'f'}$ | $a/\varepsilon$ | delete another domain |
| $(\mathrm{I_D}, f', g', n', e')$ | $\mathcal{T}_{\mathtt{DI}} \cdot v_{n'} \cdot \kappa_{n'} \cdot w_{n'f'}$ | $\varepsilon/b$ | insert domain |
| $\mathtt{E}$ | $\mathcal{T}_{\mathtt{DE}}$ | $\varepsilon/\varepsilon$ | end |

Here $\mathcal{T}_{\mathtt{D}\cdot}$ is used because the domain-level state was $\mathtt{D}$.

**Insert states ($\mathrm{I_F}$, $\mathrm{I_D}$).**

Insert states handle character-level emissions for inserted fragments/domains. They self-loop for fragment extension and terminate back to wait states.

$\mathrm{I_F}$ (inserted fragment within an existing domain):

| Source | Dest | Weight | I/O |
|---|---|---|---|
| $(\mathrm{I_F}, f, 0, n, e)$ | $(\mathrm{I_F}, g, g', n, e)$ | $r^{(n)}_{fg}$ (emit next char) | $\varepsilon/b$ |
| $(\mathrm{I_F}, f, 1, n, e)$ | $(\mathrm{W_M}, f, 1, n, e)$ | 1 (fragment ended) | $\varepsilon/\varepsilon$ |

Since the label $g$ on the output character indicates whether the fragment continues ($g = 0$) or ends ($g = 1$), the fragment-extension matrix $r^{(n)}$ is again handled by the singlet HMM generating the output labels. In the WFST, $\mathrm{I_F}$ at $g = 0$ continues emitting; at $g = 1$ the fragment ends and control returns to $\mathrm{W_M}$ to decide on the next fragment-level action.

Actually, for insert states, the output character labels are generated by the WFST itself (there is no ancestral character to copy from). The WFST must therefore assign the correct probabilities to the output labels. The emission weight at $(\mathrm{I_F}, f, g, n, e)$ for output character $b_{c'f'g'n'e'}$ is:

$$\delta_{nn'}\delta_{ee'} \cdot u_{f'c'}\pi_{c'b} \cdot \begin{cases} r^{(n)}_{ff'} & \text{if } g' = 0 \\ \rho^{(n)}_{f}\,\delta_{ff'} & \text{if } g' = 1 \end{cases} \tag{C.182}$$

followed by a transition: if $g' = 0$, loop to $(\mathrm{I_F}, f, 0, n, e)$; if $g' = 1$, go to the appropriate wait state.

$\mathrm{I_D}$ (inserted domain): similar to $\mathrm{I_F}$, but the entire domain is new. Within the inserted domain, the fragment-level TKF92 structure applies:

| Source condition | Dest | Notes |
|---|---|---|
| $(\mathrm{I_D}, f, 0, n, e)$ | $(\mathrm{I_D}, f, g', n, e)$, emit $b$ | mid-fragment, continue |
| $(\mathrm{I_D}, f, 1, n, 0)$ | $(\mathrm{I_D}, f', g', n, e')$, emit $b$ | new fragment in inserted domain |
| | weight: $\kappa_n \cdot w_{nf'}$ | |
| $(\mathrm{I_D}, f, 1, n, 0)$ | domain ends within inserted domain | weight: $(1 - \kappa_n)$ |
| | $\to$ next domain-level action | |
| $(\mathrm{I_D}, f, 1, n, 1)$ | inserted domain complete | returns to domain-level wait |

When $\mathrm{I_D}$ completes (the inserted domain ends), control goes back to a domain-level wait state. Since the domain was inserted (top-level state $\mathtt{I}$), the next domain-level transition uses $\mathcal{T}_{\mathtt{I}\cdot}$:

| Dest | Weight | I/O | Notes |
|---|---|---|---|
| $(\mathtt{M}, f', g', n', e')$ | $\mathcal{T}_{\mathtt{IM}} \cdot v_{n'} \cdot \tau^{(n')}_{\mathtt{SM}} \cdot w_{n'f'}$ | $a/b$ | match next domain |
| $(\mathrm{D}_{\mathrm{D}}, f', g', n', e')$ | $\mathcal{T}_{\mathtt{ID}} \cdot v_{n'} \cdot \kappa_{n'} \cdot w_{n'f'}$ | $a/\varepsilon$ | delete next domain |
| $(\mathrm{I}_{\mathrm{D}}, f', g', n', e')$ | $\mathcal{T}_{\mathtt{II}} \cdot v_{n'} \cdot \kappa_{n'} \cdot w_{n'f'}$ | $\varepsilon/b$ | insert another domain |
| $\mathtt{E}$ | $\mathcal{T}_{\mathtt{IE}}$ | $\varepsilon/\varepsilon$ | end |

##### C.11.4 Emission Weights

The emission weights for each emitting state type are:

| State | Emission | Weight |
|---|---|---|
| $\mathtt{M}$ | input $a_{cfgne}$, output $b_{cfgne}$ | $\exp(R_c T)_{ab}$ |
| $\mathrm{I}_{\mathrm{F}}$ | output $b_{cfgne}$ | $u_{fc}\,\pi_{cb} \cdot p_g(r^{(n)}_{f.})$ |
| $\mathrm{I}_{\mathrm{D}}$ | output $b_{cfgne}$ | $u_{fc}\,\pi_{cb} \cdot p_g(r^{(n)}_{f.})$ |
| $\mathrm{D}_{\mathrm{F}}$ | input $a_{cfgne}$ | 1 |
| $\mathrm{D}_{\mathrm{D}}$ | input $a_{cfgne}$ | 1 |

where $p_g(r^{(n)}_{f.})$ stands for $r^{(n)}_{ff'}$ if $g = 0$ (fragment continues with destination fragchar $f'$) and $\rho^{(n)}_{f}$ if $g = 1$ (fragment ends), and the substitution matrix $R_c$ is the rate matrix for site class $c$.

For the WFST (conditional on ancestor), the delete emission weight is 1 because the ancestral emission probability is divided out (it was generated by the singlet HMM). The match emission divides out the ancestral $\pi_{ca}$ from the pair emission $\pi_{ca} \exp(R_c T)_{ab}$, yielding just the substitution matrix entry.

For insert states, the emission includes the full descendant character probability because there is no ancestral character to condition on.

##### C.11.5 Verification: Composition Reproduces the Pair HMM

**Claim.** The Labeled-MixDom Singlet HMM composed with the Labeled-MixDom WFST is equivalent to the MixDom Pair HMM defined in Section C.1.1.

**Sketch of proof.** The composition proceeds as follows:

1. The Singlet HMM generates the ancestral labeled sequence, with transitions governed by the MixDom stationary distribution.
2. The WFST reads the ancestral labeled sequence as input and produces the descendant labeled sequence as output.
3. The composed machine has states that are pairs (singlet state, WFST state). Since both are order-1 machines tracking structural labels, the composed state is a pair of structural labels.

We verify that the composed transition weights match $\chi$ for each case:

**Match-to-Match within a fragment ($g = 0$).** The singlet emits character $a_{cf0ne}$ with weight $u_{fc}\pi_{ca}$, and transitions to state $(f', g', n, e)$ with weight $r^{(n)}_{ff'}$ (if $g' = 0$) or $\rho^{(n)}_f \cdot \ldots$ (if $g' = 1$). The WFST in state $(\text{W}_\text{M}, f, 0, n, e)$ reads this input and transitions to $(\texttt{M}, g, g', n, e)$ with weight $\alpha_n$, emitting $b$ with substitution weight $\exp(R_c T)_{ab}$. The composed emission weight is $u_{gc}\pi_{ca} \exp(R_c T)_{ab}$, summing over $c$ gives $\sum_c u_{gc}\pi_{ca} \exp(R_c T)_{ab}$, which matches the Pair HMM emission for $\texttt{MM}_{ng}$. The transition weight within the fragment is $r^{(n)}_{fg} \cdot \alpha_n$, which corresponds to the $\delta(l = m) r^{(n)}_{fg}$ term in $\chi$ (for the intra-fragment Markov contribution to $\texttt{MM}_{lf} \to \texttt{MM}_{lg}$).

**Match-to-Match across fragment boundary ($g = 1, e = 0$).** Singlet: fragment ends, transitions to new fragment $f'$ within same domain with weight $\kappa_n \cdot w_{nf'}$. WFST: $(\text{W}_\text{M}, f, 1, n, 0)$ transitions to $(\texttt{M}, f', g', n, e')$ with weight $\alpha_n \cdot w_{nf'}$. Combined: $\kappa_n \cdot w_{nf'} \cdot \alpha_n \cdot w_{nf'}$.

Wait—the $w$ appears twice, which is wrong. This reveals that the WFST transition weight should *not* include $w_{nf'}$ when the input character determines $f'$. The fragment type of the next input character is determined by the singlet HMM, and the WFST simply reads whatever fragment type appears.

Let us correct: at fragment boundaries, when the next action involves reading an input character, the WFST does *not* weight by $w_{nf'}$. The $w$ is part of the singlet distribution, not the conditional (WFST) distribution.

**Corrected Wait-State Transitions.** The WFST represents the *conditional* distribution $P(\text{descendant} \mid \text{ancestor})$. Therefore:

- Transitions that consume an input character should *not* include the prior probability of that input character's labels (such as $w$, $v$).
- Transitions that produce an output character (insertions) *do* include the full probability of the output labels (since the WFST generates them).
- Transitions involving the $\mathcal{T}$ matrix at domain boundaries must be adjusted: $\mathcal{T}$ was derived for the Pair HMM (joint distribution), so the WFST version divides out the ancestral prior factors.

Concretely, the $\chi$ entry for $\texttt{MX}_{lf} \to \texttt{MY}_{mg}$ with $l = m$ (same domain, new fragment) is:

$$\rho^{(n)}_f \cdot \tau^{(n)}_{\texttt{XY}} \cdot w_{ng}$$

This is the *joint* weight. In the composed (singlet $\circ$ WFST) machine:

- Singlet provides: $\rho^{(n)}_f \cdot \kappa_n \cdot w_{ng}$ (end fragment, continue domain, choose new fragment type)
- WFST provides: $\frac{\tau^{(n)}_{\texttt{XY}}}{\kappa_n}$ (the conditional transition, dividing out the $\kappa_n$ from the joint)

Product: $\rho^{(n)}_f \cdot \kappa_n \cdot w_{ng} \cdot \tau^{(n)}_{\texttt{XY}} / \kappa_n = \rho^{(n)}_f \cdot \tau^{(n)}_{\texttt{XY}} \cdot w_{ng}$. This matches the Pair HMM. ✓

Similarly, for the domain-boundary case with $l \neq m$:

- Singlet provides: $\rho^{(l)}_f (1 - \kappa_l) \cdot \frac{\kappa_0 v_m \kappa_m w_{mg}}{1-\zeta}$
- WFST provides: the conditional weight that, when multiplied by the singlet weight, gives $\chi_{\texttt{MX}_{lf} \to \texttt{MY}_{mg}}$

The required WFST transition weights therefore depend on the singlet transition structure. Rather than writing out all the corrected weights with singlet factors divided out, we express the key principle:

> **WFST weight principle:** For any transition that consumes an input character with label $\ell'$, the WFST weight is the Pair HMM transition weight divided by the singlet transition weight for generating a character with label $\ell'$ from the current singlet state. For transitions that produce an output character (insertions), the WFST carries the full conditional weight. For null transitions (emitting-to-wait), the weight is 1.

Specifically, let $P_{\text{pair}}(i \to j)$ be the Pair HMM transition from state $i$ to $j$, and let $P_{\text{sing}}(\ell \to \ell')$ be the singlet transition. Then:

$$w_{\text{WFST}}(s, \ell_{\text{in}}, \ell_{\text{out}} \to s', \ell') = \frac{P_{\text{pair}}(i(s,\ell) \to j(s',\ell'))}{P_{\text{sing}}(\ell \to \ell')} \tag{C.183}$$

where $i(s,\ell)$ and $j(s',\ell')$ are the corresponding Pair HMM states, and $\ell_{\text{in}}$ is present when $s'$ consumes input.

**Explicit corrected weights.** We now tabulate the corrected WFST transition weights organized by source wait state, boundary case, and whether input is consumed.

In all cases below, the WFST context $\ell = (f, g, n, e)$ is the structural label of the last processed character. The singlet HMM is in the corresponding state. The factor $P_{\text{sing}}(\ell \to \ell')$ is the singlet transition weight from the current label to the next label.

$\text{W}_\text{M}$**,** $g = 0$ **(mid-fragment, input consumed):**

| Dest | WFST weight | Notes |
|---|---|---|
| `M` | $\alpha_n$ | match |
| $\text{D}_\text{F}$ | $(1-\alpha_n)$ | frag-level delete |

No singlet factor to divide out: at $g = 0$ the singlet continues the fragment with weight $r^{(n)}_{fg}$ (producing the next character with destination fragchar $g$ in the same domain), and the WFST weight is purely the TKF92 match/delete split, independent of $w$.

$\text{W}_\text{M}$**,** $g = 1, e = 0$ **(fragment boundary, mid-domain):**

With input (start new ancestral fragment):

| Dest | WFST weight | Notes |
|---|---|---|
| `M` | $\alpha_n$ | match new fragment |
| $\text{D}_\text{F}$ | $(1-\alpha_n)$ | delete new fragment |

The $w_{nf'}$ is supplied by the singlet; the WFST contributes only the match/delete split.

Without input (descendant insertion):

| Dest | WFST weight | Notes |
|---|---|---|
| $(\text{I}_\text{F}, f', g', n, 0)$ | $\beta_n \cdot w_{nf'} \cdot u_{f'c'}\pi_{c'b} \cdot p_{g'}(r_{f'})$ | insert fragment |

Here the WFST generates the full output character probability since there is no input to condition on.

**$\mathrm{W_M}$, $g = 1, e = 1$ (domain boundary):**

With input (start new ancestral domain):

| Dest | WFST weight | Notes |
|---|---|---|
| M | $\frac{\mathcal{T}_{\mathtt{MM}} \cdot v_{n'} \cdot \tau^{(n')}_{\mathtt{SY}} \cdot w_{n'f'}}{P_{\text{sing}}(\ell \to \ell')}$ | new domain, match |
| $\mathrm{D_F}$ | $\frac{\mathcal{T}_{\mathtt{MM}} \cdot v_{n'} \cdot \tau^{(n')}_{\mathtt{SD}} \cdot w_{n'f'}}{P_{\text{sing}}(\ell \to \ell')}$ | new domain, del-frag |
| $\mathrm{D_D}$ | $\frac{\mathcal{T}_{\mathtt{MD}} \cdot v_{n'} \cdot \kappa_{n'} \cdot w_{n'f'}}{P_{\text{sing}}(\ell \to \ell')}$ | delete domain |

Without input:

| Dest | WFST weight | Notes |
|---|---|---|
| $\mathrm{I_D}$ | $\mathcal{T}_{\mathtt{MI}} \cdot v_{n'} \cdot \kappa_{n'} \cdot w_{n'f'} \cdot u_{f'c'}\pi_{c'b} \cdot p_{g'}(r_{f'})$ | insert domain |
| E | $\mathcal{T}_{\mathtt{ME}}/P_{\text{sing}}(\ell \to \mathtt{E})$ | end |

where $P_{\text{sing}}(\ell \to \ell') = \frac{\kappa_0 v_{n'} \kappa_{n'} w_{n'f'}}{1-\zeta}$ is the singlet transition from a domain boundary to the next character label $\ell'$ (Equation C.180).

The $\mathrm{W_{D_F}}$ and $\mathrm{W_{D_D}}$ tables follow the same pattern, using $\mathcal{T}_{\mathtt{M}\cdot}$ for $\mathrm{W_{D_F}}$ (since fragment deletion is within a matched domain) and $\mathcal{T}_{\mathtt{D}\cdot}$ for $\mathrm{W_{D_D}}$ (since the domain is being deleted).

**$\mathrm{W_{D_F}}$ at $g = 0$:**

| Dest | WFST weight |
|---|---|
| $\mathrm{D_F}$ | 1 |

**$\mathrm{W_{D_F}}$ at $g = 1, e = 0$:**

| Dest | WFST weight |
|---|---|
| M | $\alpha_n$ |
| $\mathrm{D_F}$ | $(1 - \alpha_n)$ |

**$\mathrm{W_{D_F}}$ at $g = 1, e = 1$:** Same structure as $\mathrm{W_M}$ at $g = 1, e = 1$, using $\mathcal{T}_{\mathtt{M}\cdot}$.

**$\mathrm{W_{D_D}}$ at $g = 0$:**

| Dest | WFST weight |
|---|---|
| $\mathrm{D_D}$ | 1 |

$\mathrm{W}_{\mathrm{D_D}}$ **at** $g = 1, e = 0$**:**

| Dest | WFST weight |
|---|---|
| $\mathrm{D_D}$ | 1 |

(Within a deleted domain, all fragments are consumed; fragment type is irrelevant.)

$\mathrm{W}_{\mathrm{D_D}}$ **at** $g = 1, e = 1$**:**

| Dest | WFST weight | Notes |
|---|---|---|
| M | $\mathcal{T}_{\mathtt{DM}} \cdot v_{n'} \cdot \tau_{\mathtt{SY}}^{(n')} \cdot w_{n'f'}/P_{\text{sing}}(\ell \to \ell')$ | new domain, match |
| $\mathrm{D_F}$ | $\mathcal{T}_{\mathtt{DM}} \cdot v_{n'} \cdot \tau_{\mathtt{SD}}^{(n')} \cdot w_{n'f'}/P_{\text{sing}}(\ell \to \ell')$ | new domain, del-frag |
| $\mathrm{D_D}$ | $\mathcal{T}_{\mathtt{DD}} \cdot v_{n'} \cdot \kappa_{n'} \cdot w_{n'f'}/P_{\text{sing}}(\ell \to \ell')$ | delete domain |
| $\mathrm{I_D}$ | $\mathcal{T}_{\mathtt{DI}} \cdot$ (full output prob) | insert domain |
| E | $\mathcal{T}_{\mathtt{DE}}/P_{\text{sing}}(\ell \to \mathtt{E})$ | end |

$\mathrm{I_D}$ **completion at** $g = 1, e = 1$**:**

| Dest | WFST weight | Notes |
|---|---|---|
| M | $\mathcal{T}_{\mathtt{IM}} \cdot v_{n'} \cdot \tau_{\mathtt{SY}}^{(n')} \cdot w_{n'f'}/P_{\text{sing}}(\ell \to \ell')$ | match next domain |
| $\mathrm{D_D}$ | $\mathcal{T}_{\mathtt{ID}} \cdot v_{n'} \cdot \kappa_{n'} \cdot w_{n'f'}/P_{\text{sing}}(\ell \to \ell')$ | delete next domain |
| $\mathrm{I_D}$ | $\mathcal{T}_{\mathtt{II}} \cdot$ (full output prob) | insert another domain |
| E | $\mathcal{T}_{\mathtt{IE}}/P_{\text{sing}}(\ell \to \mathtt{E})$ | end |

##### C.11.6 Simplification of Domain-Boundary WFST Weights

The domain-boundary WFST weights (Section C.11.3) involve a ratio of the Pair HMM transition weight to the singlet transition weight. This simplifies considerably.

For a transition that consumes input with label $\ell' = (f', g', n', e')$ from a domain boundary ($g = 1, e = 1$), the singlet weight is (from Equation C.180):

$$P_{\text{sing}}(\ell \to \ell') = \frac{\kappa_0 \cdot v_{n'} \cdot \kappa_{n'} \cdot w_{n'f'}}{1 - \zeta}$$

The Pair HMM weight for $\mathtt{MM}_{lf} \to \mathtt{MM}_{mg}$ with $l \neq m$ (different domain, going through domain boundary) is:

$$\rho_f^{(l)} \cdot \tau_{\mathtt{ME}}^{(l)} \cdot \mathcal{T}_{\mathtt{MM}} \cdot v_m \cdot \tau_{\mathtt{SM}}^{(m)} \cdot w_{mg}$$

The WFST weight is therefore:

$$\frac{\rho_f^{(l)} \cdot \tau_{\mathtt{ME}}^{(l)} \cdot \mathcal{T}_{\mathtt{MM}} \cdot v_m \cdot \tau_{\mathtt{SM}}^{(m)} \cdot w_{mg}}{\frac{\kappa_0 \cdot v_m \cdot \kappa_m \cdot w_{mg}}{1-\zeta}} = \frac{(1-\zeta)\rho_f^{(l)} \cdot \tau_{\mathtt{ME}}^{(l)} \cdot \mathcal{T}_{\mathtt{MM}} \cdot \tau_{\mathtt{SM}}^{(m)}}{\kappa_0 \cdot \kappa_m}$$

The $v_m$ and $w_{mg}$ cancel, leaving a weight that depends on the source domain parameters and the destination domain's TKF parameters, but *not* on the specific fragment or domain type of the destination. This is a significant simplification: the WFST transition weight at domain boundaries is the same for all destination labels $\ell'$, given the source label $\ell$.

Since $\tau^{(l)}_{\texttt{ME}} = (1-\beta_l)(1-\kappa_l)$ and $\tau^{(m)}_{\texttt{SM}} = (1-\beta_m)\kappa_m\alpha_m$, the weight becomes:

$$\frac{(1-\zeta)\rho_f^{(l)}(1-\beta_l)(1-\kappa_l)\cdot\mathcal{T}_{\texttt{MM}}\cdot(1-\beta_m)\alpha_m}{\kappa_0}$$

Note that this weight still depends on $m$ through $(1-\beta_m)\alpha_m$, so the cancellation is partial: $v$ and $w$ cancel but the TKF92 entry parameters do not.

##### C.11.7 State Count and Sparsity

The Labeled-MixDom WFST has at most $8L+2$ states where $L = |\mathcal{F}|\cdot|\mathcal{N}|\cdot 4$. In practice, many combinations are constrained away:

- $\mathrm{I_F}$ states only occur with $e \neq 1$ (inserted fragments cannot be the last in a domain since they are inserted *within* a domain).
- Mid-fragment ($g = 0$) wait states have at most 2 outgoing transitions each (continue in same fragment, match or delete).
- Fragment-boundary ($g = 1, e = 0$) wait states have at most 3 outgoing transition types.
- Domain-boundary ($g = 1, e = 1$) wait states have at most 5 outgoing transition types (one per $\mathcal{T}$ column).

The transition matrix is therefore very sparse. For typical values ($|\mathcal{F}| = 4$, $|\mathcal{N}| = 20$, $|\mathcal{C}| = 4$), we get $L = 4\cdot 20\cdot 4 = 320$ and the WFST has at most $8\cdot 320+2 = 2562$ states, comparable to the Maraschino-distilled WFST (which has $\sim 8\cdot|\mathcal{A}|^2+2$ states for the order-1 pair context).

The key advantage over distillation is exactness: the Labeled-MixDom WFST preserves all correlations of the MixDom model without approximation, at the cost of a larger effective alphabet.

#### C.12 Formal Grammar Elaboration Rules

The TKF family of evolutionary models—TKF91, TKF92, MixDom, the TKF Structure Tree, and the TKF Genome—describes the joint evolution of biological sequences subject to insertions, deletions, and substitutions. Despite their apparent diversity, all these models share a common constructive pattern: they begin with a simple weighted context-free grammar (WCFG) for a geometrically distributed number of links, and then systematically elaborate that grammar through a series of formal transformations.

In the TKF91 model (50), each link carries a single character evolving by a continuous-time Markov chain (CTMC). TKF92 (51) extends this by replacing each character with a geometrically distributed fragment of characters. The MixDom model nests a TKF92 process inside a TKF91 process, decorating links with mixtures of domain types, fragment types, and substitution classes. The TKF Structure Tree (22) uses stochastic context-free grammar (SCFG) recursion to model RNA secondary structure with stems (emitting paired characters left and right) and loops. The TKF Genome extends these ideas to entire genomes with coding sequences, introns, RNA structures, and conserved elements.

Making the elaboration steps explicit and composable has several benefits:

(i) **Correctness**: each transformation can be verified independently, rather than checking a large monolithic grammar.

(ii) **Modularity**: new models (e.g., RNA models with basepair stacking, codon models with reading-frame-aware indels) can be constructed by composing well-understood building blocks.

(iii) **Automation**: the transformations are sufficiently formal to be implemented as software operations on grammar objects, enabling automatic derivation of dynamic programming algorithms from high-level model specifications.

This appendix defines seven elaboration rules and the associated null-state management procedures, then shows how each known TKF-family model arises as a specific sequence of elaborations.

##### C.12.1 Base Grammar

###### Weighted Context-Free Grammars

**Definition C.1** (Weighted Context-Free Grammar). *A* weighted context-free grammar *(WCFG) is a tuple $\mathcal{G} = (\mathcal{N}, \Sigma, \mathcal{P}, S, w)$ where:*

- *$\mathcal{N}$ is a finite set of nonterminal symbols.*
- *$\Sigma$ is a finite set of terminal symbols, disjoint from $\mathcal{N}$.*
- *$S \in \mathcal{N}$ is the start symbol.*
- *$\mathcal{P}$ is a finite set of production rules of the form $X \to \alpha$ where $X \in \mathcal{N}$ and $\alpha \in (\mathcal{N} \cup \Sigma)^*$.*
- *$w : \mathcal{P} \to \mathbb{R}_{\geq 0}$ assigns a nonneg weight to each production.*

*The grammar is* proper *if for every nonterminal $X$, the weights of all productions with left-hand side $X$ sum to 1: $\sum_{(X\to\alpha)\in\mathcal{P}} w(X \to \alpha) = 1$. In a proper WCFG, weights are probabilities and the grammar defines a stochastic context-free grammar (SCFG).*

**Definition C.2** (Elaboration Rule). *An* elaboration rule *(or grammar transformation) is a map $\mathcal{E} : \mathcal{G} \to \mathcal{G}'$ that takes a WCFG and a set of elaboration parameters, and produces a new WCFG. An elaboration is* validity-preserving *if it maps proper grammars to proper grammars.*

We distinguish between the *single-sequence grammar* (describing the stationary distribution over sequences) and the *pair grammar* (describing the joint distribution over ancestor–descendant sequence pairs). Most elaborations operate on the single-sequence grammar; the Evolution elaboration (Section C.12.2) converts a single-sequence grammar into a pair grammar.

**The Link Grammar** The fundamental building block of all TKF-family models is a grammar generating a geometrically distributed number of "links." This grammar arises from the stationary distribution of the linear birth-death-immigration (BDI) process with per-capita birth rate $\lambda$, per-capita death rate $\mu > \lambda$, and immigration rate $\nu = \lambda$.

**Definition C.3** (Link Grammar). *The* link grammar $\mathcal{G}_{\text{link}}(\kappa)$ *with parameter $\kappa = \lambda/\mu \in [0, 1)$ is the WCFG with nonterminals $\{\texttt{IMM}, \texttt{MOR}\}$, start symbol $\texttt{IMM}$, and productions:*

$$\texttt{IMM} \to \texttt{MOR}\ \texttt{IMM} \qquad \text{weight } \kappa \tag{C.184}$$

$$\texttt{IMM} \to \epsilon \qquad \text{weight } 1-\kappa \tag{C.185}$$

$$\texttt{MOR} \to \texttt{MOR}\ \texttt{MOR} \qquad \text{weight } \kappa \tag{C.186}$$

$$\texttt{MOR} \to \epsilon \qquad \text{weight } 1-\kappa \tag{C.187}$$

*Remark* C.19. In this grammar, `IMM` (the "immortal link") generates a sequence of $n \sim \text{Geometric}(\kappa)$ mortal links. Each `MOR` can recursively generate further mortal links; the recursive self-loop in (C.186) reflects the offspring-generating property of the BDI process. The distinction between `IMM` and `MOR` captures the different roles: the immortal link corresponds to the BDI regime $\nu = \lambda, X(0) = 0$ (immigration from nothing), while mortal links correspond to $\nu = 0, X(0) = 1$ (a single founder that can die).

*Remark* C.20. The grammar in Definition C.3 generates only the empty string $\epsilon$, since `MOR` has no terminal-producing rules. The elaboration rules below will add terminal emissions (characters, character pairs, etc.) to the mortal links.

**Proposition C.3.** *The link grammar $\mathcal{G}_{\text{link}}(\kappa)$ is proper for any $\kappa \in [0, 1)$. Under the start symbol `IMM`, the number of `MOR` expansions before reaching $\epsilon$ is distributed as Geometric($\kappa$) (with support $\{0, 1, 2, \ldots\}$ and mean $\kappa/(1-\kappa)$).*

*Proof.* For `IMM`: the productions (C.184) and (C.185) have weights $\kappa$ and $1 - \kappa$, summing to 1. Similarly for `MOR`. The number of `MOR` nonterminals generated by `IMM` before choosing $\epsilon$ is geometric with parameter $\kappa$ by the standard geometric series argument. □

##### C.12.2 Elaboration Rules

We now define each elaboration rule as a formal grammar transformation. For each rule, we specify:

- The *input* grammar fragment (which productions are targeted).
- The *output* grammar fragment (the replacement productions).
- The *parameters* introduced by the elaboration.
- The *validity conditions* under which the transformation preserves properness.

**CTMC Expansion** The most basic elaboration decorates each link with a character (or tuple of characters) that evolves according to a finite-state CTMC. This is the step that takes the bare link grammar to a model with observable sequences.

**Definition C.4** (Emission Type)**.** *An* emission type *specifies where terminal symbols appear relative to the recursive expansion of a nonterminal:*

- ***Left-emission****: terminal appears to the left of the recursive part. Production form: $X \to c\,\alpha$ where $c \in \Sigma$.*
- ***Right-emission****: terminal appears to the right. Production form: $X \to \alpha\, c$.*
- ***LR-emission****: terminals appear on both sides. Production form: $X \to c_L\ \alpha\ c_R$ where $c_L, c_R \in \Sigma$. This is the form needed for RNA basepair models (stems).*

**Definition C.5** (CTMC Expansion)**.** *Let $\mathcal{G}$ be a WCFG containing a nonterminal $X$ with an $\epsilon$-generating production used for link termination. Let $\mathcal{A}$ be a finite alphabet, $\pi$ a probability distribution over $\mathcal{A}$, and $Q$ a rate matrix with stationary distribution $\pi$. The* left CTMC expansion *of $X$ with parameters $(\mathcal{A}, \pi, Q)$ replaces every production $X \to \alpha$ (where $\alpha \neq \epsilon$) with the family of productions:*

$$X \to c\,\alpha \qquad \textit{weight } w(X \to \alpha) \cdot \pi_c \quad \textit{for each } c \in \mathcal{A} \tag{C.188}$$

*The $\epsilon$-production $X \to \epsilon$ is left unchanged.*

*The* right CTMC expansion *replaces $X \to \alpha$ ($\alpha \neq \epsilon$) with:*

$$X \to \alpha\, c \qquad \textit{weight } w(X \to \alpha) \cdot \pi_c \quad \textit{for each } c \in \mathcal{A} \tag{C.189}$$

*The* LR CTMC expansion *with alphabet $\mathcal{A} \times \mathcal{A}$ and equilibrium distribution $\pi(c_L, c_R)$ replaces $X \to \alpha$ ($\alpha \neq \epsilon$) with:*

$$X \to c_L\, \alpha\, c_R \qquad \textit{weight } w(X \to \alpha) \cdot \pi(c_L, c_R) \quad \textit{for each } (c_L, c_R) \in \mathcal{A} \times \mathcal{A} \tag{C.190}$$

**Proposition C.4.** *Left, right, and LR CTMC expansion are validity-preserving: if $\mathcal{G}$ is proper, then so is the elaborated grammar.*

*Proof.* For left CTMC expansion: the total weight of productions with LHS $X$ becomes $\sum_{c \in \mathcal{A}} \sum_{\alpha \neq \epsilon} w(X \to \alpha) \cdot \pi_c + w(X \to \epsilon) = \sum_{\alpha \neq \epsilon} w(X \to \alpha) \cdot 1 + w(X \to \epsilon) = \sum_\alpha w(X \to \alpha) = 1$. The right and LR cases are analogous. □

*Example* C.1 (TKF91). Applying left CTMC expansion to MOR in the link grammar $\mathcal{G}_{\text{link}}(\kappa)$ with alphabet $\mathcal{A}$ and equilibrium distribution $\pi$ yields:

$$\begin{aligned}
&\texttt{IMM} \to \texttt{MOR IMM} && \kappa \\
&\texttt{IMM} \to \epsilon && 1 - \kappa \\
&\texttt{MOR} \to c\ \texttt{MOR MOR} && \kappa \cdot \pi_c \quad (c \in \mathcal{A}) \\
&\texttt{MOR} \to c && (1 - \kappa) \cdot \pi_c \quad (c \in \mathcal{A})
\end{aligned}$$

This is the stationary (single-sequence) grammar for TKF91: it generates a geometric number of links, each carrying a character drawn i.i.d. from $\pi$.

*Example* C.2 (TKF Structure Tree stems). In the TKF Structure Tree, stems use LR CTMC expansion. The nonterminal $S$ ("stem") generates basepairs:

$$\begin{aligned}
&S \to c_L\ S\ c_R && (1 - \kappa_S) \cdot \pi_S(c_L, c_R) \\
&S \to L && \kappa_S
\end{aligned}$$

where $L$ is the loop nonterminal (which uses left CTMC expansion), and $\pi_S(c_L, c_R)$ is the joint equilibrium distribution over basepairs. The LR emission enables the grammar to generate palindromic structures characteristic of RNA secondary structure.

**Fragment Expansion** Fragment expansion takes TKF91 to TKF92 by replacing each single-character link with a geometrically distributed sequence of characters.

**Definition C.6** (Fragment Expansion)**.** *Let $\mathcal{G}$ be a WCFG with a nonterminal* `MOR` *representing a mortal link. The* fragment expansion *with parameter $r \in [0, 1)$ (the extension probability) replaces* `MOR` *with three nonterminals* `MOR_S` *(fragment start),* `MOR_X` *(fragment extend), and* `MOR_E` *(fragment end), defined as follows.*

*Every production in $\mathcal{G}$ that references* `MOR` *on its right-hand side is updated to reference* `MOR_S` *instead. Then:*

***Before*** *(link with single terminal slot):*

$$\texttt{MOR} \to [\textit{terminal}]\ \alpha \qquad\qquad w_{orig}$$

***After*** *(link with geometric fragment):*

$$\begin{aligned}
&\texttt{MOR_S} \to \texttt{MOR_X} && 1 \\
&\texttt{MOR_X} \to [\textit{terminal}]\ \texttt{MOR_X} && r \\
&\texttt{MOR_X} \to [\textit{terminal}]\ \texttt{MOR_E} && 1-r \\
&\texttt{MOR_E} \to \alpha && w_{orig}
\end{aligned}$$

*Here* $[\textit{terminal}]$ *denotes whatever terminal emission was associated with* `MOR` *(a single character from a CTMC expansion, or a character pair, etc.), and* $\alpha$ *denotes the rest of the original production's right-hand side (the recursive continuation).*

*More precisely, if* `MOR` *had productions* `MOR` $\to c$ `MOR MOR` *(weight* $\kappa \cdot \pi_c$*) and* `MOR` $\to c$ *(weight* $(1-\kappa)\cdot\pi_c$*) after CTMC expansion, the fragment expansion produces:*

$$\begin{aligned}
&\texttt{MOR_S} \to \texttt{MOR_X} && 1 \\
&\texttt{MOR_X} \to c\ \texttt{MOR_X} && r\cdot\pi_c \quad (c \in \mathcal{A}) \\
&\texttt{MOR_X} \to c\ \texttt{MOR_E} && (1-r)\cdot\pi_c \quad (c \in \mathcal{A}) \\
&\texttt{MOR_E} \to \texttt{MOR_S}\ \texttt{MOR_S} && \kappa \\
&\texttt{MOR_E} \to \epsilon && 1-\kappa
\end{aligned}$$

*where* `MOR_E` *takes over the inter-link continuation logic from the original* `MOR`.

**Proposition C.5.** *Fragment expansion is validity-preserving. The expected number of terminals per fragment is* $1/(1-r)$.

*Remark* C.21. Fragment expansion must be applied *after* CTMC expansion (or simultaneously), because it assumes the link already has terminal emissions. If applied to a bare link grammar (no terminals), the result would have fragments of $\epsilon$'s, which is degenerate.

*Example* C.3 (TKF92). TKF92 = link grammar → (fragment expansion with parameter $r$) → (CTMC expansion with $\mathcal{A}, \pi, Q$). Equivalently, CTMC expansion first, then fragment expansion. Both orderings yield the same grammar, because fragment expansion simply wraps the terminal emission in a geometric self-loop. The resulting grammar generates sequences where each link produces a fragment of $K \sim \text{Geometric}(r)$ characters from $\pi$.

**Mixture Expansion** Mixture expansion decorates a link with a latent categorical variable whose value determines subsequent model parameters.

**Definition C.7** (Mixture Expansion). *Let* $\mathcal{G}$ *be a WCFG containing a nonterminal* $X$. *Let* $\{1,\dots,K\}$ *be a finite set of mixture components with weights* $p_1,\dots,p_K > 0$ *satisfying* $\sum_k p_k = 1$. *The* mixture expansion *of* $X$ *with components* $K$ *and weights* $(p_k)$ *replaces* $X$ *with* $K$ *new nonterminals* $X_1,\dots,X_K$ *and modifies all productions referencing* $X$ *as follows.*

*Every production* $Y \to \alpha\ X\ \beta$ *in* $\mathcal{G}$ *(where* $Y \neq X$ *and* $\alpha, \beta \in (\mathcal{N}\cup\Sigma)^*$*) is replaced by* $K$ *productions:*

$$Y \to \alpha\ X_k\ \beta \qquad \textit{weight } w(Y \to \alpha X \beta)\cdot p_k \quad \textit{for } k = 1,\dots,K \tag{C.191}$$

*The productions of* $X$ *itself are copied to each* $X_k$*: for each production* $X \to \gamma$*, create*

$$X_k \to \gamma_k \qquad \textit{weight } w(X \to \gamma) \tag{C.192}$$

*where* $\gamma_k$ *is* $\gamma$ *with any self-references to* $X$ *replaced by* $X_k$ *(maintaining the component assignment within a link).*

*Each* $X_k$ *may then undergo different subsequent elaborations (e.g., different CTMC parameters, different fragment extension rates).*

**Proposition C.6.** *Mixture expansion is validity-preserving.*

*Proof.* For $Y$: the total weight of productions referencing $X_k$ is $\sum_k w(\cdot) \cdot p_k = w(\cdot) \cdot 1$. For each $X_k$: the production weights are copied from $X$, hence sum to 1. □

*Remark* C.22. In principle, the mixture component selector could itself evolve via a CTMC across evolutionary time, provided that nested expansions are resampled from equilibrium upon a change of component. In this appendix, we restrict attention to the simpler case where the component assignment is fixed throughout the lifetime of a link (i.e., it is sampled once at birth and does not change).

*Example* C.4 (MixDom: three levels of mixture). In the MixDom model, mixture expansion is applied at three levels:

1. **Domain mixture**: each top-level link is assigned a domain type $n \sim \text{Categorical}(v_1, \ldots, v_{\mathcal{N}})$, determining $(\lambda_n, \mu_n)$ for the nested TKF92 process.

2. **Fragment process**: within domain $n$, the initial fragment type is $f \sim \text{Categorical}(w_{n1}, \ldots, w_{n\mathcal{F}})$. Subsequent fragments are drawn from the $\mathcal{F} \times \mathcal{F}$ transition matrix $r^{(n)}_{fg}$ (the $\mathcal{F} = 1$ case reduces to IID geometric extension).

3. **Site class mixture**: within fragment state $f$ of domain $n$, each character position is assigned a class $c \sim \text{Categorical}(u_{nf1}, \ldots, u_{nfC})$, determining the substitution parameters $(Q^{(c)}, \pi^{(c)})$.

**Link Sequence Concatenation** Concatenation decorates a single link with two or more consecutive sub-links.

**Definition C.8** (Link Sequence Concatenation)**.** *Let $\mathcal{G}$ be a WCFG with nonterminal $X$ representing a link. The* binary concatenation *of $X$ replaces it with two new nonterminals $X_A, X_B$ and the production:*

$$X \to X_A \; X_B \qquad \textit{weight } 1 \tag{C.193}$$

*$X_A$ and $X_B$ may then be independently elaborated. For $n$-ary concatenation, $X$ is replaced by $X \to X_1 \; X_2 \; \cdots \; X_n$.*

*Remark* C.23 (Concatenation combined with mixture and fragments). A powerful pattern combines mixture, fragment, and concatenation to decorate a link with a variable-length sequence of categorically typed sub-links:

1. Apply fragment expansion to the link, creating a geometric number of sub-link slots.

2. Apply mixture expansion to each sub-link slot, assigning it a categorical type.

3. If correlations between adjacent types are desired, replace the i.i.d. mixture with an HMM output distribution: the type sequence is generated by a hidden Markov model whose transition matrix captures adjacency preferences.

This yields a variable number of concatenated sub-links of varying types with Markovian correlations. In the TKF Genome, this pattern is used for genomic regions (coding, noncoding, structural) within a top-level link sequence.

*Example* C.5 (TKF Genome: region concatenation). The TKF Genome's top-level grammar has:

$$
\begin{aligned}
\texttt{GENOME} &\rightarrow \texttt{REGION GENOME} && 1-\kappa_R \\
\texttt{GENOME} &\rightarrow \epsilon && \kappa_R \\
\texttt{REGION} &\rightarrow \texttt{INTER} && p_N \\
&\mid \texttt{FWDCDS} && p_G/2 \\
&\mid \texttt{REVCDS} && p_G/2 \\
&\mid \texttt{STRUCT} && p_S \\
&\mid \texttt{CONS} && p_C
\end{aligned}
$$

Here REGION is both a mixture expansion (over region types) and a concatenation point (each region type expands into its own sub-grammar).

**Non-Recursive Nesting** Non-recursive nesting splices a complete sub-grammar into the transitions of a mortal link, without introducing bifurcation or self-reference. This is how nesting works in MixDom.

**Definition C.9** (Non-Recursive Nesting)**.** *Let $\mathcal{G}_{\text{outer}}$ be a link grammar with nonterminal* **MOR** *(mortal link), and let $\mathcal{G}_{\text{inner}}$ be an independent link grammar with its own start symbol* $\texttt{IMM}_{\text{inner}}$ *and parameters. The* non-recursive nesting *of $\mathcal{G}_{\text{inner}}$ into* **MOR** *of $\mathcal{G}_{\text{outer}}$ replaces each terminal-emitting production of* **MOR** *with:*

$$\texttt{MOR} \rightarrow \texttt{IMM}_{\text{inner}}\ \alpha \qquad \textit{weight } w_{\text{orig}} \tag{C.194}$$

*where $\alpha$ is the original continuation. The inner grammar generates a complete sequence of inner links, each with its own parameters, for every outer mortal link.*

*More precisely, wherever* **MOR** *previously emitted a terminal symbol c, it now expands into the entire inner grammar $\mathcal{G}_{\text{inner}}$, which itself may generate zero or more characters. The inner grammar's $\epsilon$-productions (zero-length inner sequences) give rise to null states in the combined grammar.*

*Remark* C.24. The key difference from recursive nesting (Section C.12.2) is that $\mathcal{G}_{\text{inner}}$ does *not* reference any nonterminals of $\mathcal{G}_{\text{outer}}$. There is no possibility of re-entering the outer grammar from within the inner grammar. This ensures that the combined grammar generates strings from a regular language (at each level), rather than a context-free language.

*Example* C.6 (MixDom as non-recursive nesting). The MixDom model nests a Markovian fragment process (inner grammar) into each mortal link of a TKF91 process (outer grammar). The outer link grammar has parameters $(\lambda_0, \mu_0)$ and the inner grammar, for domain type $n$, has parameters $(\lambda_n, \mu_n, r^{(n)}_{fg}, u_{nfc}, Q^{(c)}, \pi^{(c)})$. Each outer mortal link, instead of emitting a single character, expands into a domain sequence governed by the Markovian fragment process.

Since the inner grammar can generate the empty string (the inner link sequence may have length zero), this nesting creates null states in the combined Pair HMM. These must be eliminated by the procedures of Section C.12.3.

**Recursive Nesting** Recursive nesting, used in the TKF Structure Tree, allows transitions into a mortal link to spawn a bifurcation: a new nonterminal whose sub-grammar may reference the original grammar's nonterminals.

**Definition C.10** (Recursive Nesting)**.** *Let $\mathcal{G}$ be a link grammar with nonterminals including* **MOR** *(mortal link). Let* **BIF** *be a new nonterminal with its own sub-grammar $\mathcal{G}_{\text{bif}}$ that may reference nonterminals of $\mathcal{G}$ (including* **IMM** *and* **MOR***).*

*The* right recursive nesting *of* `BIF` *at* `MOR` *replaces each terminal-emitting production of* `MOR` *with a mixture of terminal emission and bifurcation:*
***Before:***

$$\texttt{MOR} \to c\ \alpha \qquad\qquad w_{\text{orig}} \cdot \pi_c$$

***After:***

$$\begin{array}{lll} \texttt{MOR} \to c\ \alpha & w_{\text{orig}} \cdot s \cdot \pi_c & \textit{(terminal link, probability } s\textit{)} \\ \texttt{MOR} \to \texttt{BIF}\ \alpha & w_{\text{orig}} \cdot (1-s) \cdot v_{\text{bif}} & \textit{(bifurcation, probability } 1-s\textit{)} \end{array}$$

*where $s \in (0,1]$ is the probability that a link is a terminal (character-emitting) link rather than a nesting point, and $v_{\text{bif}}$ is a distribution over bifurcation types if there are multiple* `BIF` *variants.*

*For* left recursive nesting*:*

$$\texttt{MOR} \to \alpha\ \texttt{BIF} \qquad\qquad w_{\text{orig}} \cdot (1-s) \cdot v_{\text{bif}}$$

*The sub-grammar $\mathcal{G}_{\text{bif}}$ for* `BIF` *defines how the bifurcation expands. Since it may reference the start symbol of $\mathcal{G}$ (e.g.,* `IMM`*), the combined grammar is genuinely recursive: link sequences can contain nested link sequences of arbitrary depth.*

*Remark* C.25. The recursive nesting creates null cycles whenever the nested sub-grammar can generate the empty string. These must be handled by the null-state management procedures (Section C.12.3). In the TKF Structure Tree, the nullability fixed-point iteration solves for the probability that each nonterminal generates $\epsilon$.

*Example* C.7 (TKF Structure Tree: stems and loops)*.* The TKF Structure Tree has two types of link sequences:

- *Loop sequences* ($L$): left-emitting links generating single nucleotides. Nonterminal rule: $L \to c_L\ L$ with CTMC expansion (left emission).
- *Stem sequences* ($S$): LR-emitting links generating basepairs. Nonterminal rule: $S \to c_L\ S\ c_R$ with CTMC expansion (LR emission).

The recursive nesting works as follows. Within a loop sequence, a mortal link may either emit a character (probability $s_L$) or spawn a stem (probability $1 - s_L$):

$$L \to c\ L \quad | \quad S\ L$$

At the base of a stem (when the self-loop terminates), the grammar transitions to a loop:

$$S \to c_L\ S\ c_R \quad | \quad L$$

This creates the alternating stem–loop structure characteristic of RNA secondary structure. The recursion arises because a loop can spawn a stem, which eventually returns to a loop, which can spawn another stem, *ad infinitum*.

*Example* C.8 (TKF92 with Recursive Domains)*.* The recursive domain model has nonterminals $\mathtt{L}_n$ (link sequence for domain $n$), $\mathtt{A}_n$ (aligned component), $\mathtt{I}_n$ (inserted component), $\mathtt{D}_n$ (deleted component), etc. The aligned component rule is:

$$\begin{array}{lll} \mathtt{A}_n \to c & s_n \cdot \pi_{n,c} & \text{(terminal: character)} \\ \mathtt{A}_n \to \mathtt{L}_{n'} & (1-s_n) \cdot v_{nn'} & \text{(bifurcation: nested link sequence)} \end{array}$$

Since $\mathtt{L}_{n'}$ can reference $\mathtt{L}_n$ (if $n' = n$ or through a chain of domain transitions), this creates genuine recursion: link sequences contain domains that contain further link sequences.

**Evolution** The Evolution elaboration converts a single-sequence grammar (describing the stationary distribution) into a pair grammar (describing the joint ancestor–descendant distribution at evolutionary time $T$).

**Definition C.11** (Evolution Elaboration)**.** *Let $\mathcal{G}_1$ be a single-sequence grammar with link nonterminal $X$ generating terminals from alphabet $\mathcal{A}$ with equilibrium distribution $\pi$. Let $Q$ be the CTMC rate matrix and $T > 0$ the evolutionary time.*

*Define the TKF parameters:*

$$\alpha = e^{-\mu T}, \quad \beta = \frac{\lambda(e^{-\lambda T} - e^{-\mu T})}{\mu e^{-\lambda T} - \lambda e^{-\mu T}}, \quad \gamma = 1 - \frac{\mu\beta}{\lambda(1-\alpha)}, \quad \kappa = \frac{\lambda}{\mu}$$

*The evolution elaboration replaces each link nonterminal $X$ in $\mathcal{G}_1$ with three nonterminals $X_{\mathtt{M}}$, $X_{\mathtt{I}}$, $X_{\mathtt{D}}$ in the pair grammar $\mathcal{G}_2$:*

***Before*** *(single-sequence grammar, left-emitting):*

$$\begin{array}{lll} X \to c\,X\,X & \kappa \cdot \pi_c & \textit{(link with offspring)} \\ X \to c & (1-\kappa)\cdot \pi_c & \textit{(terminal link)} \end{array}$$

***After*** *(pair grammar):*

*For the* immortal link *continuation, each nonterminal $Y$ that previously generated the link sequence $Y \to X\ Y \mid \epsilon$ transforms as follows:*

$$\begin{array}{ll} Y_{\mathtt{M}} \to X_{\mathtt{M}}\,Y_{\mathtt{M}} & (1-\beta_Y)\kappa\alpha \\ Y_{\mathtt{M}} \to X_{\mathtt{I}}\,Y_{\mathtt{M}} & \beta_Y \\ Y_{\mathtt{M}} \to X_{\mathtt{D}}\,Y_{\mathtt{D}} & (1-\beta_Y)\kappa(1-\alpha) \\ Y_{\mathtt{M}} \to \epsilon & (1-\beta_Y)(1-\kappa) \\ \\ Y_{\mathtt{D}} \to X_{\mathtt{M}}\,Y_{\mathtt{M}} & (1-\gamma_Y)\kappa\alpha \\ Y_{\mathtt{D}} \to X_{\mathtt{I}}\,Y_{\mathtt{M}} & \gamma_Y \\ Y_{\mathtt{D}} \to X_{\mathtt{D}}\,Y_{\mathtt{D}} & (1-\gamma_Y)\kappa(1-\alpha) \\ Y_{\mathtt{D}} \to \epsilon & (1-\gamma_Y)(1-\kappa) \end{array}$$

*where $\beta_Y$ and $\gamma_Y$ use the appropriate $(\lambda, \mu)$ for the link sequence that $Y$ belongs to.*

*The subscript indicates the alignment type:*

- $X_{\mathtt{M}}$ *(**match**): ancestral link survived; emits aligned pair $(c_a, c_d)$ with probability $\pi_{c_a} \cdot \exp(RT)_{c_a c_d}$.*
- $X_{\mathtt{I}}$ *(**insert**): new link born in descendant; emits descendant-only character $c_d$ with probability $\pi_{c_d}$.*
- $X_{\mathtt{D}}$ *(**delete**): ancestral link died; emits ancestor-only character $c_a$ with probability $\pi_{c_a}$.*

*The transition weights are precisely the entries of the TKF91 Pair HMM transition matrix $\tau(\lambda, \mu, T)$.*

**Proposition C.7.** *The evolution elaboration roughly triples the number of nonterminals (each single-sequence nonterminal becomes three pair nonterminals). For LR-emitting nonterminals (as in stem sequences), match states emit paired tuples $(c_L^a, c_R^a, c_L^d, c_R^d)$, insert states emit $(c_L^d, c_R^d)$, and delete states emit $(c_L^a, c_R^a)$.*

*Remark* C.26. When fragment expansion has been applied before evolution, the fragment self-loop interleaves with the alignment states. In TKF92, the Pair HMM transition matrix becomes $\tau'$ where the M, I, and D self-loops gain a fragment-extension component:

$$\tau'_{aa} = r + (1-r)\tau_{aa} \qquad \text{for } a \in \{\texttt{M}, \texttt{I}, \texttt{D}\}$$

and off-diagonal transitions are scaled by $(1-r)$:

$$\tau'_{ab} = (1-r)\tau_{ab} \qquad \text{for } a \neq b$$

*Remark* C.27. For LR-emitting grammars (such as stem sequences in the TKF Structure Tree), the evolution elaboration creates nonterminals $X_M, X_I, X_D$ whose left and right emissions are correlated:

- $X_M \to c_L^x\ c_L^y\ X_M\ c_R^y\ c_R^x$: ancestral basepair $(c_L^x, c_R^x)$ evolved to descendant basepair $(c_L^y, c_R^y)$ (match).
- $X_I$: emits only descendant basepair $c_L^y\ \cdots\ c_R^y$ (insertion).
- $X_D$: emits only ancestral basepair $c_L^x\ \cdots\ c_R^x$ (deletion).

The ancestor terminals go on the outside, the descendant terminals on the inside (or vice versa), preserving the palindromic nesting.

*Example* C.9 (TKF91 Pair HMM). Applying evolution to the TKF91 single-sequence grammar (Example C.1) yields the standard 5-state Pair HMM $(\texttt{S}, \texttt{M}, \texttt{I}, \texttt{D}, \texttt{E})$ with transition matrix $\tau(\lambda, \mu, T)$.

##### C.12.3 Null State Management

###### Null State Identification

**Definition C.12** (Null state). *A nonterminal $X$ in a WCFG is* nullable *if there exists a derivation $X \Rightarrow^* \epsilon$. The* nullability *$\eta(X) = P(X \Rightarrow^* \epsilon)$ is the probability that a parse tree rooted at $X$ yields the empty string.*

Elaborations that create null states include:

1. **Non-recursive nesting**: the inner grammar may generate the empty string (e.g., a domain in MixDom may contain zero fragments).
2. **Recursive nesting**: nested link sequences can be empty.
3. **Mixture expansion** combined with nesting: a mixture component that expands into a nullable sub-grammar.

In the MixDom Pair HMM, null states arise because each domain's inner TKF92 process can generate an empty sequence. The probability of an empty domain at evolutionary time $T$ is $z_T = \sum_n v_n(1-\kappa_n)(1-\beta_n)$.

##### Null State Removal: The $(I - T_{NN})^{-1}$ Closure

**Definition C.13** (Null Closure). *Let $\mathcal{G}$ be a WCFG (or equivalently an HMM/transducer) with states partitioned into emitting states $\Omega$ and non-emitting (null) states $\mathcal{Z}$. Let $T_{\mathcal{Z}\mathcal{Z}}$ be the submatrix of transition weights among null states. The* null closure *is:*

$$N^* = (I - T_{\mathcal{Z}\mathcal{Z}})^{-1} = \sum_{k=0}^{\infty} T_{\mathcal{Z}\mathcal{Z}}^k \tag{C.195}$$

*This converges provided the spectral radius $\rho(T_{\mathcal{Z}\mathcal{Z}}) < 1$.*

**Proposition C.8** (Effective Transition Matrix). *The effective transition matrix between emitting (and start/end) states, with all null-state paths summed out, is:*

$$\hat{T} = T_{\Omega\Omega} + T_{\Omega\mathcal{Z}} \cdot (I - T_{\mathcal{Z}\mathcal{Z}})^{-1} \cdot T_{\mathcal{Z}\Omega} \tag{C.196}$$

*where subscripts denote submatrices restricted to the indicated state sets.*

##### Null Cycle Detection and Removal

**Definition C.14** (Null Cycle). *A* null cycle *in a WCFG is a chain of unit productions (productions whose right-hand side is a single nonterminal) that returns to the starting nonterminal: $X \to Y_1 \to Y_2 \to \cdots \to X$. In the context of HMMs/transducers, this corresponds to a cycle among non-emitting states.*

Null cycles arise in two main situations:

1. **Non-recursive nesting with empty inner grammars**: when the inner grammar can produce $\epsilon$, a path $\texttt{S} \to \texttt{MOR}_{\text{outer}} \to \texttt{IMM}_{\text{inner}} \to \epsilon \to \texttt{MOR}_{\text{outer}}$ creates a cycle through null states.
2. **Recursive nesting**: the chain $\texttt{L}'_n \to \texttt{M}'_n \to \texttt{S}'_n \to \texttt{A}'_n \to \texttt{L}'_{n'}$ creates a null cycle when $n' = n$ or when the chain of domain transitions eventually returns to $n$.

**Definition C.15** (Null Cycle Removal). *To remove null cycles from a WCFG:*

1. ***Compute nullabilities****: for each nonterminal $X$, compute $\eta(X) = P(X \Rightarrow^* \epsilon)$. In the non-recursive case, this can be done in closed form. In the recursive case, iterate the fixed-point equations:*
$$\eta(X) = \sum_{(X \to \alpha) \in \mathcal{P}} w(X \to \alpha) \cdot \prod_{Y \in \alpha} \eta(Y) \tag{C.197}$$
*where the product is over nonterminals in $\alpha$, with $\eta(\text{terminal}) = 0$ and $\eta(\epsilon) = 1$.*
*Initialize $\eta^{(0)}(X) = 0$ for all $X$ and iterate until convergence.*
2. ***Create non-nullable copies****: for each nullable nonterminal $X$, create $X'$ whose productions never generate $\epsilon$. For a bifurcation rule $X \to Y\ Z$, the non-nullable version adds:*
$$\begin{aligned} X' &\to Y'\ Z' && w(X \to YZ) \\ X' &\to Y' && w(X \to YZ) \cdot \eta(Z) \\ X' &\to Z' && w(X \to YZ) \cdot \eta(Y) \end{aligned}$$
*This accounts for the two ways one child can be null.*

3. ***Remove unit-production cycles**: identify cycles $X' \to Y'_1 \to \cdots \to X'$ among the non-nullable nonterminals. For each such cycle, compute the transition matrix $\mathcal{A}$ among the cycle's nonterminals and replace the cycle with its closure $(I - \mathcal{A})^{-1}$, distributing the accumulated weight to the non-cyclic continuations.*

*Example* C.10 (Recursive domains: nullability fixed point). In the recursive domain model, $\eta(\mathtt{C}_n)$ (nullability of the child link sequence nonterminal) satisfies:

$$\eta(\mathtt{C}_n) = \frac{1 - \kappa_n}{1 - \kappa_n(1 - s_n) \sum_{n'} v_{nn'} \eta(\mathtt{C}_{n'})}$$

This is solved by initializing $x_n^{(0)} = 0$ and iterating:

$$x_n^{(k+1)} = \frac{1 - \kappa_n}{1 - \kappa_n(1 - s_n) \sum_{n'} v_{nn'} x_{n'}^{(k)}}$$

After convergence, the null cycles $\mathtt{L}'_n \to \mathtt{L}'_{n'}$ and $\mathtt{C}'_n \to \mathtt{C}'_{n'}$ are removed using the $(I - \mathcal{A})^{-1}$ and $(I - \mathcal{B})^{-1}$ closures respectively, where $\mathcal{A}$ and $\mathcal{B}$ are the $\mathcal{N} \times \mathcal{N}$ transition matrices among domain types.

##### C.12.4 Composition Properties

**Commutativity and Order** The elaboration rules do not, in general, commute. The following table summarizes the ordering constraints.

| **Constraint** | **Reason** |
|---|---|
| CTMC expansion commutes with mixture expansion | Mixture selects which CTMC parameters to use; the order of these two operations does not affect the final grammar. Both orderings produce the same set of productions. |
| Fragment expansion must follow (or be simultaneous with) CTMC expansion | Fragment expansion wraps terminal emissions in a geometric self-loop. Without terminals, fragment expansion produces fragments of $\epsilon$'s. |
| Non-recursive nesting must follow both CTMC and fragment expansion of the inner grammar | The inner grammar must be fully specified before it can be spliced into the outer grammar. |
| Evolution must be applied last (after all structural elaborations) | Evolution triples the nonterminals and introduces alignment-dependent transition weights $(\alpha, \beta, \gamma)$. Applying structural elaborations after evolution would require modifying all three copies independently. |
| Mixture expansion commutes with concatenation | Both are structural operations on different aspects of a link. |
| Null state removal must follow all nullable elaborations but precede distillation | All null states must be identified before they can be summed out. The distilled order-1 machines assume null-free grammars. |

**Validity Conditions**

**Definition C.16** (Well-Formed Elaborated Grammar)**.** *An elaborated grammar $\mathcal{G}'$ is* well-formed *if:*

1. ***Properness****: for every nonterminal $X$, the production weights sum to 1.*
2. ***No unresolved null cycles****: after null state removal, no cycles among non-emitting states remain. Equivalently, $\rho(T_{\mathcal{Z}\mathcal{Z}}) < 1$ for the null-state transition matrix.*
3. ***Convergent nullability****: for recursive grammars, the fixed-point iteration for nullabilities converges. A sufficient condition is that $\kappa_n < 1$ and $s_n > 0$ for all domain types (every link has a positive probability of being terminal rather than a nesting point).*
4. ***Finite expected derivation length****: the expected total number of terminals generated is finite. For the link grammar, this requires $\kappa < 1$. For recursive nesting, additional conditions on the nesting probabilities are needed.*

**Proposition C.9.** *Each elaboration rule defined above is validity-preserving under its stated conditions. Composition of validity-preserving elaborations is validity-preserving, provided the ordering constraints above are respected.*

**Derivation of Existing Models** We now show explicitly how each known TKF-family model arises as a sequence of elaborations applied to the base link grammar.

**TKF91.**

$$\boxed{\text{TKF91}} = \mathcal{G}_{\text{link}}(\kappa) \xrightarrow{\text{CTMC}(\mathcal{A},\pi,Q)} \mathcal{G}_{\text{TKF91}}$$

Steps:

1. Start with the link grammar $\mathcal{G}_{\text{link}}(\kappa)$ (Definition C.3).
2. Apply left CTMC expansion (Definition C.5) to `MOR` with alphabet $\mathcal{A}$, equilibrium $\pi$, rate matrix $Q$.

Result: each mortal link emits a single character from $\pi$. The pair grammar is obtained by applying evolution (Definition C.11), yielding the standard 5-state Pair HMM with transition matrix $\tau(\lambda, \mu, T)$.

**TKF92.**

$$\boxed{\text{TKF92}} = \mathcal{G}_{\text{link}}(\kappa) \xrightarrow{\text{Frag}(r)}\xrightarrow{\text{CTMC}(\mathcal{A},\pi,Q)} \mathcal{G}_{\text{TKF92}}$$

Steps:

1. Start with $\mathcal{G}_{\text{link}}(\kappa)$.
2. Apply fragment expansion (Definition C.6) with extension probability $r$.
3. Apply left CTMC expansion to each fragment position.

Result: each mortal link emits a fragment of $K \sim \text{Geometric}(r)$ characters. The pair grammar has self-looping match/insert/delete states with fragment-extension probability $r$.

**MixDom: Markovian fragments.**

$$\boxed{\text{MixDom}} = \mathcal{G}_{\text{link}}(\kappa_0) \xrightarrow{\text{Mix}(v_n)} \xrightarrow[\text{for each } n]{\text{NonRecNest}\left(\mathcal{G}_{\text{link}}(\kappa_n) \xrightarrow{\text{HMM}(r^{(n)})} \xrightarrow{\text{Mix}(w_{nf})} \xrightarrow{\text{Mix}(u_{nfc})} \xrightarrow{\text{CTMC}(\mathcal{A},\pi^{(c)},Q^{(c)})}\right)} \mathcal{G}_{\text{MixDom}}$$

Steps:

1. Start with $\mathcal{G}_{\text{link}}(\kappa_0)$ (outer/top-level link grammar).

2. Apply mixture expansion (Definition C.7) to `MOR` with domain types $n \sim \text{Categorical}(v_1, \ldots, v_{\mathcal{N}})$.

3. For each domain type $n$, construct an inner grammar:

   (a) Start with $\mathcal{G}_{\text{link}}(\kappa_n)$ (inner link grammar).

   (b) Replace geometric fragment extension with the Markovian fragment HMM governed by the $\mathcal{F} \times \mathcal{F}$ transition matrix $r_{fg}^{(n)}$.

   (c) Apply mixture expansion for initial fragment types $f \sim \text{Categorical}(w_{n1}, \ldots, w_{n\mathcal{F}})$.

   (d) Apply mixture expansion for site classes $c \sim \text{Categorical}(u_{nf1}, \ldots, u_{nfC})$.

   (e) Apply CTMC expansion with $(\mathcal{A}, \pi^{(c)}, Q^{(c)})$.

4. Apply non-recursive nesting (Definition C.9): splice each domain's inner grammar into the corresponding outer mortal link.

5. Apply null state removal (Section C.12.3): the inner grammar can generate empty sequences (null domains), creating the null states `A`, `B`, `C` in the null-separated Pair HMM. The $(I - T_{\mathcal{Z}\mathcal{Z}})^{-1}$ closure reduces the 8-state null-separated Pair HMM to the effective 5-state matrix $\mathcal{T}$.

**TKF Structure Tree.**

$$\boxed{\text{TKF Structure Tree}} = \mathcal{G}_{\text{link}}(\kappa_L) \xrightarrow{\text{CTMC}_L(\mathcal{A},\pi_L,Q_L)} \xrightarrow{\text{RecNest}\left(\mathcal{G}_{\text{link}}(\kappa_S) \xrightarrow{\text{CTMC}_{LR}(\mathcal{A}^2,\pi_S,Q_S)}\right)} \mathcal{G}_{\text{ST}}$$

Steps:

1. Start with $\mathcal{G}_{\text{link}}(\kappa_L)$ (loop link grammar).

2. Apply left CTMC expansion for loops with $(\mathcal{A}, \pi_L, Q_L)$.

3. Apply recursive nesting (Definition C.10): within the loop, a mortal link may spawn a stem. The stem sub-grammar is:

   (a) Start with $\mathcal{G}_{\text{link}}(\kappa_S)$ (stem link grammar).

   (b) Apply LR CTMC expansion with basepair alphabet $(\mathcal{A} \times \mathcal{A}, \pi_S, Q_S)$.

   (c) At stem termination, return to the loop grammar (creating the recursion $S \to L$).

4. The loop grammar now has two types of mortal links:

   - Character-emitting links (probability $s_L$): emit a single nucleotide $c$ with $\pi_L(c)$.
   - Stem-spawning links (probability $1 - s_L$): expand into a stem $S$ nonterminal.

The LR emission in the stem grammar is critical: the rule $S \to c_L\ S\ c_R$ emits characters on *both* sides of the recursive expansion, generating the palindromic base-pairing structure of RNA stems.

**TKF Genome.**

$$\boxed{\text{TKF Genome}} = \mathcal{G}_{\text{link}}(\kappa_R) \xrightarrow{\text{Concat+Mix(region types)}} \xrightarrow{\text{various nested elaborations per region type}} \mathcal{G}_{\text{Genome}}$$

Steps:

1. Start with $\mathcal{G}_{\text{link}}(\kappa_R)$ (top-level genomic region grammar).
2. Apply concatenation + mixture: each link is a "region" selected from types {`INTER`, `FWDCDS`, `REVCDS`, `STRUCT`, `CO`[illegible] with probabilities $(p_N, p_G/2, p_G/2, p_S, p_C)$.
3. Each region type undergoes its own elaboration chain:
   - `INTER`: link grammar + left CTMC expansion (single nucleotides, neutral evolution).
   - `FWDCDS`/`REVCDS`: link grammar + concatenation into codons $(c_1c_2c_3)$ + CTMC expansion with codon substitution model + recursive nesting for introns (introns contain a nested `GENOME` nonterminal, flanked by splice donor/acceptor sites).
   - `STRUCT`: link grammar with LR CTMC expansion (stems) + recursive nesting into loops (left CTMC expansion).
   - `CONS`: link grammar + left CTMC expansion (conserved elements).
4. The intron nesting creates recursion: `GENOME` → `CDS` → `CODON` → `INTRON` → `GENOME`.

##### C.12.5 Toward Implementation

**Grammar Objects with Transformation Methods** The elaboration rules defined above can be implemented as methods on a grammar object:

```
class WCFG:
    nonterminals: Set[str]
    terminals: Set[str]
    productions: List[Production]
    start: str

    def ctmc_expand(self, nonterminal, alphabet,
                    equilibrium, emission_type='left'):
        """CTMC Expansion"""

    def fragment_expand(self, nonterminal, extension_prob):
        """Fragment Expansion"""

    def mixture_expand(self, nonterminal, components, weights):
        """Mixture Expansion"""

    def concatenate(self, nonterminal, parts):
        """Link Sequence Concatenation"""

    def nest_nonrecursive(self, nonterminal, inner_grammar):
        """Non-Recursive Nesting"""
```

```
    def nest_recursive(self, nonterminal, bifurcation_grammar,
                       terminal_prob, side='right'):
        """Recursive Nesting"""

    def evolve(self, time, rates):
        """Evolution"""

    def remove_null_states(self):
        """Null State Removal"""
```

Each method validates the preconditions (e.g., that fragment expansion targets a nonterminal with terminal emissions), performs the transformation, and returns the modified grammar.

**Automatic Derivation of DP Algorithms** Given an elaborated grammar, the dynamic programming algorithm (Forward, Backward, Inside, Viterbi) can be derived automatically by:

1. **State space identification**: each nonterminal in the elaborated grammar corresponds to a state in the DP. Emitting states correspond to observable positions (sequence characters); non-emitting states are handled by null closure.

2. **Transition structure**: the productions define the recurrence relations. Linear (non-branching) productions yield HMM-style recurrences; branching (bifurcation) productions yield CYK-style recurrences.

3. **Emission probabilities**: determined by the CTMC parameters and the emission type (left, right, LR, match/insert/delete).

4. **Fill order**: determined by the topological sort of nonterminals (after null cycle removal). For recursive grammars, the fill order follows the CYK pattern (by span length for context-free rules).

**Connection to Existing Frameworks** The elaboration rules defined here are compatible with existing grammar and transducer frameworks:

- **Transducer composition**: the Evolution elaboration produces a transducer (Pair HMM / WFST). These can be composed on phylogenetic trees using the standard composition and intersection operations for Mealy machines in waiting-machine normal form.

- **SCFG parsers**: the elaborated grammars (especially those with recursive nesting and LR emission) are SCFGs amenable to Inside/Outside parsing algorithms.

- **Distillation**: the elaborated pair grammars can be distilled to order-1 machines (HMMs and WFSTs) by computing adjacency frequencies and normalizing, as described in Section C.4.5. This step loses the hierarchical structure but produces compact machines suitable for phylogenetic composition.

#### C.13 Recursive TKF Models

Another way to nest TKF models is to borrow from grammar theory and allow recursion. This was previously used to develop RNA evolutionary models (22), and is developed here in a general way for proteins and genomic DNA.

We illustrate how the TKF-mixed domain model can be interpreted as a stochastic grammar, developing four examples—recursive protein domains (i.e. arbitrary nesting of motifs), a basic model of RNA foldback structure, a second more sophisticated model of RNA structure, and a basic model of a genome—that highlight the structure of such models as a series of stepwise grammar elaborations that constitute tree-adjoining moves on the space of grammars.

##### C.13.1 Example One: Left-Recursive TKF (L-TKF)

We can imagine a links model where, in a domain of type $l$, each mortal link is either (with probability $s_l$) associated with a character, or (with probability $(1 - s_l)v_{lm}$ associated with its own independently-evolving links model of domain type $m$. Links models can thus be nested *ad infinitum*

$$\begin{aligned} M_l &= \text{Links}(\text{Mix}_{k\sim s}(L_l^{(k)}); \lambda_l, \mu_l) \\ L_l^{(1)} &= \text{HMM}(\{Q^{(c)}, \pi^{(c)}\}_c; r^{(l)}, u_l) \\ L_l^{(0)} &= \text{Mix}_{m\sim v_m}(M_m) \end{aligned}$$

where HMM denotes the Markovian fragment process (Section C.1.1), and $\text{Mix}_{k\sim s}$ is defined for the Bernoulli index variable $k$ and probability $s$ as it was for categorical index variables

$$x \sim \text{Mix}_{k\sim p}(M(\theta_k); p) \;\Leftrightarrow\; k \sim \text{Bernoulli}(p),\; x \sim M(\theta_k)$$

We postpone the TKF92-like augmentations of $L_l^{(1)}$ (fragment types $f$ and site classes $c$) for now, and start with a simplified TKF91-like version that allows full recursively-nested domains but allows only single-character fragments with one site class per domain

$$\begin{aligned} M'_l &= \text{Links}(\text{Mix}_{k\sim s}(L_l'^{(k)}); \lambda_l, \mu_l) \\ L_l'^{(1)} &= \text{Subst}(Q_l, \pi_l) \\ L_l'^{(0)} &= \text{Mix}_{m\sim v_l}(M'_m) \end{aligned}$$

As with the nested TKF HMM, we have to account for the probability of zero-length components, leading to null cycles during likelihood and inference computations.

The joint distribution over ancestor-descendant alignments under the recursive TKF model is described by the following stochastic context-free grammar (SCFG)

| Symbol interpretation | LHS | | RHS | | Probability |
|---|---|---|---|---|---|
| **L**ink sequence, domain $l$ | $\mathtt{L}_l$ | $\to$ | $\mathtt{L}_l$ | $\mathtt{M}_l$ | $\kappa_l$ |
| | | $\mid$ | $\mathtt{K}_l$ | | $1-\kappa_l$ |
| Immortal lin**k** | $\mathtt{K}_l$ | $\to$ | $\mathtt{N}_l$ | | $\beta_l$ |
| | | $\mid$ | $\epsilon$ | | $1-\beta_l$ |
| **M**ortal link | $\mathtt{M}_l$ | $\to$ | $\mathtt{S}_l$ | | $\alpha_l$ |
| | | $\mid$ | $\mathtt{E}_l$ | | $1-\alpha_l$ |
| **S**urviving mortal link | $\mathtt{S}_l$ | $\to$ | $\mathtt{A}_l$ | $\mathtt{N}_l$ | $\beta_l$ |
| | | $\mid$ | $\mathtt{A}_l$ | | $1-\beta_l$ |
| **E**xpired mortal link | $\mathtt{E}_l$ | $\to$ | $\mathtt{D}_l$ | $\mathtt{N}_l$ | $\gamma_l$ |
| | | $\mid$ | $\mathtt{D}_l$ | | $1-\gamma_l$ |
| **N**ewborn mortal link(s) | $\mathtt{N}_l$ | $\to$ | $\mathtt{I}_l$ | $\mathtt{N}_l$ | $\beta_l$ |
| | | $\mid$ | $\mathtt{I}_l$ | | $1-\beta_l$ |
| **A**ligned component | $\mathtt{A}_l$ | $\to$ | $\mathtt{T}_{ab}$ | | $s_l \pi_{la} \exp(Q_l T)_{ab}$ |
| | | $\mid$ | $\mathtt{L}_m$ | | $(1-s_l)v_{lm}$ |
| **I**nserted component | $\mathtt{I}_l$ | $\to$ | $\mathtt{T}_{\bullet b}$ | | $s_l \pi_{lb}$ |
| | | $\mid$ | $\mathtt{C}_m$ | | $(1-s_l)v_{lm}$ |
| **D**eleted component | $\mathtt{D}_l$ | $\to$ | $\mathtt{T}_{a\bullet}$ | | $s_l \pi_{la}$ |
| | | $\mid$ | $\mathtt{P}_m$ | | $(1-s_l)v_{lm}$ |
| **C**hild link sequence (inserted) | $\mathtt{C}_l$ | $\to$ | $\mathtt{C}_l$ | $\mathtt{I}_l$ | $\kappa_l$ |
| | | $\mid$ | $\epsilon$ | | $1-\kappa_l$ |
| **P**arent link sequence (deleted) | $\mathtt{P}_l$ | $\to$ | $\mathtt{P}_l$ | $\mathtt{D}_l$ | $\kappa_l$ |
| | | $\mid$ | $\epsilon$ | | $1-\kappa_l$ |

The standard path from here is to transform the grammar to Chomsky Normal Form (45). We don't need to go all the way down that path; we only need to remove $\epsilon$-productions and null cycles.

In order to train by EM, every time we remove $\epsilon$-productions and null cycles, we need to be able to convert a rule count in the $\epsilon$-eliminated grammar to a set of rule counts in the original grammar. For each non-nullable nonterminal $\mathtt{X}'$, when a rule $\mathtt{X}' \to \mathtt{Y}'$ appears (arising from an original bifurcation $\mathtt{X} \to \mathtt{Y}\ \mathtt{Z}$ where $\mathtt{Z}$ was nullable), the expected count $c(\mathtt{X}' \to \mathtt{Y}')$ should contribute $c(\mathtt{X}' \to \mathtt{Y}') \cdot \eta(\mathtt{Z})$ to the count of the original rule $\mathtt{X} \to \mathtt{Y}\ \mathtt{Z}$, and symmetrically when the first child was nullable.

More precisely, for the nullability fixed-point iteration, this means tracking an additional set of *expected counts* alongside the nullabilities. For each nonterminal $\mathtt{X}$ with nullability $\eta(\mathtt{X})$, define $\bar{c}(\mathtt{X} \to \alpha)$ as the expected number of times rule $\mathtt{X} \to \alpha$ would have been used in the original grammar, conditioned on $\mathtt{X}$ generating the empty string. These satisfy analogous fixed-point equations:

$$\bar{c}(\mathtt{X} \to \mathtt{Y}_1 \cdots \mathtt{Y}_k) = w(\mathtt{X} \to \mathtt{Y}_1 \cdots \mathtt{Y}_k) \prod_{i=1}^{k} \eta(\mathtt{Y}_i) + \sum_{i=1}^{k} \frac{w(\mathtt{X} \to \mathtt{Y}_1 \cdots \mathtt{Y}_k) \prod_{j=1}^{k} \eta(\mathtt{Y}_j)}{\eta(\mathtt{X})} \sum_{\alpha'} \bar{c}(\mathtt{Y}_i \to \alpha')$$

and can be iterated to convergence alongside the nullabilities. Given posterior counts from the Inside-Outside algorithm on the $\epsilon$-eliminated grammar, the original-grammar rule counts are recovered by:

1. For each non-null production $\mathtt{X}' \to \alpha'$ in the $\epsilon$-eliminated grammar, its posterior count $c'(\mathtt{X}' \to \alpha')$ contributes directly to the corresponding original rule.

2. For each "nullability shortcut" production $\mathtt{X}' \to \mathtt{Y}'$ (arising from $\mathtt{X} \to \mathtt{Y}\ \mathtt{Z}$ with $\mathtt{Z}$ nullable), the count $c'(\mathtt{X}' \to \mathtt{Y}')$ contributes $c'(\mathtt{X}' \to \mathtt{Y}')$ to the original rule $\mathtt{X} \to \mathtt{Y}\ \mathtt{Z}$, plus $c'(\mathtt{X}' \to \mathtt{Y}') \cdot \bar{c}(\mathtt{Z} \to \alpha)$ to each rule within $\mathtt{Z}$'s null derivation subtree.

The general theory of $\epsilon$-elimination with EM count recovery, including the null closure $(I - T_{\mathcal{Z}\mathcal{Z}})^{-1}$ and null cycle removal, is developed in Appendix C.12 (Section C.12.3)

First we find the nullability $\eta(\mathtt{X})$ of each nonterminal $\mathtt{X}$. The nullability is the probability that a parse tree rooted in that nonterminal yields the empty string. We can't solve for these in closed form (at least not in the general recursive model, where a link sequence can contain another link sequence of the same type). Instead we can solve approximately by iterating towards a fixed point.

We first observe that the nullabilities collectively satisfy the following

$$\begin{aligned}
\eta(\mathtt{L}_l) &= \eta(\mathtt{K}_l)\frac{1-\kappa_l}{1-\kappa_l\eta(\mathtt{M}_l)} \\
\eta(\mathtt{K}_l) &= 1-\beta_l+\beta_l\eta(\mathtt{N}_l) \\
\eta(\mathtt{M}_l) &= \alpha_l\eta(\mathtt{S}_l)+(1-\alpha_l)\eta(\mathtt{E}_l) \\
\eta(\mathtt{S}_l) &= \eta(\mathtt{A}_l)\left(\beta_l\eta(\mathtt{N}_l)+(1-\beta_l)\right) \\
\eta(\mathtt{E}_l) &= \eta(\mathtt{D}_l)\left(\gamma_l\eta(\mathtt{N}_l)+(1-\gamma_l)\right) \\
\eta(\mathtt{N}_l) &= \eta(\mathtt{I}_l)\frac{1-\beta_l}{1-\beta_l\eta(\mathtt{I}_l)} \\
\eta(\mathtt{A}_l) &= (1-s_l)\sum_m v_{lm}\eta(\mathtt{L}_m) \\
\eta(\mathtt{I}_l) &= (1-s_l)\sum_m v_{lm}\eta(\mathtt{C}_m) \\
\eta(\mathtt{D}_l) &= \eta(\mathtt{I}_l) \\
\eta(\mathtt{C}_l) &= \frac{1-\kappa_l}{1-\kappa_l\eta(\mathtt{I}_l)} \\
\eta(\mathtt{P}_l) &= \eta(\mathtt{C}_l)
\end{aligned}$$

A general procedure is (i) iterate to solve for the $\eta(\mathtt{C}_l)$ (substituting in the downstream definition of $\eta(\mathtt{I}_l)$ so the formula becomes self-referential); (ii) this directly yields $\eta(\mathtt{P}_l), \eta(\mathtt{I}_l), \eta(\mathtt{D}_l), \eta(\mathtt{N}_l), \eta(\mathtt{E}_l), \eta(\mathtt{K}_l)$; (iii) iterate to solve for $\eta(\mathtt{L}_l)$ (again, first substituting to make it self-referential); (iv) this directly yields the remaining $\eta(\mathtt{M}_l), \eta(\mathtt{S}_l), \eta(\mathtt{A}_l)$.

In detail: initialize $x_l^{(0)} \leftarrow 0$ for $l \in \mathcal{N}$. Iterate to convergence

$$x_l^{(k+1)} \leftarrow \frac{1-\kappa_l}{1-\kappa_l(1-s_l)\sum_m v_{lm}x_m^{(k)}}$$

We then set $\eta(\mathtt{C}_l) \leftarrow \lim_{k\to\infty} x_l^{(k)}$ and set $\eta(\mathtt{P}_l), \eta(\mathtt{I}_l), \eta(\mathtt{D}_l), \eta(\mathtt{N}_l), \eta(\mathtt{E}_l), \eta(\mathtt{K}_l)$ using the above equations. Now set $y_l^{(0)} \leftarrow 0$ for $l \in \mathcal{N}$ and again iterate

$$y_l^{(k+1)} \leftarrow \frac{(1-\kappa_l)\eta(\mathtt{K}_l)}{1-\kappa_l(1-\alpha_l)\eta(\mathtt{E}_l)-\kappa_l\alpha_l\left(1-\beta_l(1-\eta(\mathtt{N}_l))\right)(1-s_l)\sum_m v_{lm}y_m^{(k)}}$$

Then set $\eta(\mathtt{L}_l) \leftarrow \lim_{k\to\infty} y_l^{(k)}$ and set the remaining $\eta(\mathtt{M}_l), \eta(\mathtt{S}_l), \eta(\mathtt{A}_l)$ using the above equations.

We next develop a "non-nullable" version of the grammar that yields the same Inside probabilities, but explicitly separates out $\epsilon$-generations. For every nonterminal $\mathtt{X}_l$ we create a new nonterminal

$\mathtt{X}'_l$ with rules that (by construction) never generate empty parse trees, but are otherwise identical to those transforming $\mathtt{X}_l$. In cases where the original grammars has bifurcation rules $\mathtt{X}_i \to \mathtt{X}_j \mathtt{X}_k$, we need to introduce transitions $\mathtt{X}'_i \to \mathtt{X}'_j$ and $\mathtt{X}'_i \to \mathtt{X}'_k$, to account for the missing nullability of $\mathtt{X}_j$ and $\mathtt{X}_k$. (We can also eliminate $\mathtt{K}_l$, which only has one outgoing rule when its $\epsilon$-production is removed, and fold $\mathtt{L}_l \to \mathtt{K}_l \to \mathtt{M}_l$ into $\mathtt{L}_l \to \mathtt{M}_l$.)

| Symbol interpretation | LHS | | RHS | | Probability |
|---|---|---|---|---|---|
| **L**ink sequence, domain $l$ | $\mathtt{L}'_l$ | $\to$ | $\mathtt{L}'_l$ | $\mathtt{M}'_l$ | $\kappa_l$ |
| (nonempty) | | $\mid$ | $\mathtt{L}'_l$ | | $\kappa_l \eta(\mathtt{M}_l)$ |
| | | $\mid$ | $\mathtt{M}'_l$ | | $\kappa_l \eta(\mathtt{L}_l)$ |
| | | $\mid$ | $\mathtt{N}'_l$ | | $(1-\kappa_l)\beta_l$ |
| **M**ortal link (nonempty) | $\mathtt{M}'_l$ | $\to$ | $\mathtt{S}'_l$ | | $\alpha_l$ |
| | | $\mid$ | $\mathtt{E}'_l$ | | $1-\alpha_l$ |
| **S**urviving mortal link | $\mathtt{S}'_l$ | $\to$ | $\mathtt{A}'_l$ | $\mathtt{N}'_l$ | $\beta_l$ |
| (etc.; all $\mathtt{X}'_l$ are nonempty) | | $\mid$ | $\mathtt{N}'_l$ | | $\beta_l \eta(\mathtt{A}_l)$ |
| | | $\mid$ | $\mathtt{A}'_l$ | | $1-\beta_l(1-\eta(\mathtt{N}_l))$ |
| **E**xpired mortal link | $\mathtt{E}'_l$ | $\to$ | $\mathtt{D}'_l$ | $\mathtt{N}'_l$ | $\gamma_l$ |
| | | $\mid$ | $\mathtt{N}'_l$ | | $\gamma_l \eta(\mathtt{D}_l)$ |
| | | $\mid$ | $\mathtt{D}'_l$ | | $1-\gamma_l(1-\eta(\mathtt{N}_l))$ |
| **N**ewborn mortal link(s) | $\mathtt{N}'_l$ | $\to$ | $\mathtt{I}'_l$ | $\mathtt{N}'_l$ | $\beta_l$ |
| | | $\mid$ | $\mathtt{N}'_l$ | | $\beta_l \eta(\mathtt{I}_l)$ |
| | | $\mid$ | $\mathtt{I}'_l$ | | $1-\beta_l(1-\eta(\mathtt{N}_l))$ |
| **A**ligned component | $\mathtt{A}'_l$ | $\to$ | $\mathtt{T}_{ab}$ | | $s_l \pi_{la} \exp(Q_l T)_{ab}$ |
| | | $\mid$ | $\mathtt{L}'_m$ | | $(1-s_l)v_{lm}$ |
| **I**nserted component | $\mathtt{I}'_l$ | $\to$ | $\mathtt{T}_{\bullet b}$ | | $s_l \pi_{lb}$ |
| | | $\mid$ | $\mathtt{C}'_m$ | | $(1-s_l)v_{lm}$ |
| **D**eleted component | $\mathtt{D}'_l$ | $\to$ | $\mathtt{T}_{a\bullet}$ | | $s_l \pi_{la}$ |
| | | $\mid$ | $\mathtt{P}'_m$ | | $(1-s_l)v_{lm}$ |
| **C**hild link sequence (inserted) | $\mathtt{C}'_l$ | $\to$ | $\mathtt{C}'_l$ | $\mathtt{I}'_l$ | $\kappa_l$ |
| | | $\mid$ | $\mathtt{C}'_l$ | | $\kappa_l \eta(\mathtt{I}_l)$ |
| | | $\mid$ | $\mathtt{I}'_l$ | | $\kappa_l \eta(\mathtt{C}_l)$ |
| **P**arent link sequence (deleted) | $\mathtt{P}'_l$ | $\to$ | $\mathtt{P}'_l$ | $\mathtt{D}'_l$ | $\kappa_l$ |
| | | $\mid$ | $\mathtt{P}'_l$ | | $\kappa_l \eta(\mathtt{D}_l)$ |
| | | $\mid$ | $\mathtt{D}'_l$ | | $\kappa_l \eta(\mathtt{P}_l)$ |

The final step is to remove null cycles: chains of unit productions resulting in the same nonterminal. Specifically we need to remove $\mathtt{L}'_l \to \mathtt{M}'_l \to \mathtt{S}'_l \to \mathtt{A}'_l \to \mathtt{L}'_m$, $\mathtt{C}'_l \to \mathtt{I}'_l \to \mathtt{C}'_m$ and $\mathtt{P}'_l \to \mathtt{D}'_l \to \mathtt{P}'_m$. We do this by deleting the $\mathtt{A}'_l \to \mathtt{L}'_m$, $\mathtt{I}'_l \to \mathtt{C}'_m$ and $\mathtt{D}'_l \to \mathtt{P}'_m$ transitions, adding compensatory self-loops and self-bifurcations to $\mathtt{L}'_l$, $\mathtt{P}'_l$, and $\mathtt{C}'_l$ to account for the now-broken paths through $\mathtt{A}'_l$, $\mathtt{I}'_l$ and $\mathtt{D}'_l$. The modified grammar is

| Symbol | LHS | | RHS | | Probability |
|---|---|---|---|---|---|
| **L**ink | $\mathtt{L}''_l$ | $\to$ | $\mathtt{L}''_l$ | $\mathtt{M}''_l$ | $\kappa_l$ |
| | | $\mid$ | $\mathtt{L}''_l$ | $\mathtt{L}''_m$ | $\kappa_l \alpha_l \left(1-\beta_l(1-\eta(\mathtt{N}_l))\right)(1-s_l) v_{lm}$ |
| | | $\mid$ | $\mathtt{L}''_m$ | | $\mathcal{A}_{lm}$ |
| | | $\mid$ | $\mathtt{M}''_l$ | | $\kappa_l \eta(\mathtt{L}_l)$ |
| | | $\mid$ | $\mathtt{N}''_l$ | | $(1-\kappa_l)\beta_l$ |
| **M**ortal | $\mathtt{M}''_l$ | $\to$ | $\mathtt{S}''_l$ | | $\alpha_l$ |
| | | $\mid$ | $\mathtt{E}''_l$ | | $1-\alpha_l$ |
| **S**urviving | $\mathtt{S}''_l$ | $\to$ | $\mathtt{A}''_l$ | $\mathtt{N}''_l$ | $\beta_l$ |
| | | $\mid$ | $\mathtt{N}''_l$ | | $\beta_l \eta(\mathtt{A}_l)$ |
| | | $\mid$ | $\mathtt{A}''_l$ | | $1-\beta_l(1-\eta(\mathtt{N}_l))$ |
| **E**xpired | $\mathtt{E}''_l$ | $\to$ | $\mathtt{D}''_l$ | $\mathtt{N}''_l$ | $\gamma_l$ |
| | | $\mid$ | $\mathtt{P}''_m$ | $\mathtt{N}''_l$ | $\gamma_l(1-s_l)v_{lm}$ |
| | | $\mid$ | $\mathtt{N}''_l$ | | $\gamma_l \eta(\mathtt{D}_l)$ |
| | | $\mid$ | $\mathtt{D}''_l$ | | $1-\gamma_l(1-\eta(\mathtt{N}_l))$ |
| | | $\mid$ | $\mathtt{P}''_m$ | | $\left(1-\gamma_l(1-\eta(\mathtt{N}_l))\right)(1-s_l)v_{lm}$ |
| **N**ewborns | $\mathtt{N}''_l$ | $\to$ | $\mathtt{I}''_l$ | $\mathtt{N}''_l$ | $\beta_l$ |
| | | $\mid$ | $\mathtt{C}''_m$ | $\mathtt{N}''_l$ | $\beta_l(1-s_l)v_{lm}$ |
| | | $\mid$ | $\mathtt{N}''_l$ | | $\beta_l \eta(\mathtt{I}_l)$ |
| | | $\mid$ | $\mathtt{I}''_l$ | | $1-\beta_l(1-\eta(\mathtt{N}_l))$ |
| | | $\mid$ | $\mathtt{C}''_m$ | | $\left(1-\beta_l(1-\eta(\mathtt{N}_l))\right)(1-s_l)v_{lm}$ |
| **A**ligned | $\mathtt{A}''_l$ | $\to$ | $\mathtt{T}_{ab}$ | | $s_l \pi_{la} \exp(Q_l T)_{ab}$ |
| **I**nserted | $\mathtt{I}''_l$ | $\to$ | $\mathtt{T}_{\bullet b}$ | | $s_l \pi_{lb}$ |
| **D**eleted | $\mathtt{D}''_l$ | $\to$ | $\mathtt{T}_{a\bullet}$ | | $s_l \pi_{la}$ |
| **C**hild | $\mathtt{C}''_l$ | $\to$ | $\mathtt{C}''_l$ | $\mathtt{I}''_l$ | $\kappa_l$ |
| | | $\mid$ | $\mathtt{C}''_l$ | $\mathtt{C}''_m$ | $\kappa_l(1-s_l)v_{lm}$ |
| | | $\mid$ | $\mathtt{C}''_m$ | | $\mathcal{B}_{lm}$ |
| | | $\mid$ | $\mathtt{I}''_l$ | | $\kappa_l \eta(\mathtt{C}_l)$ |
| **P**arent | $\mathtt{P}''_l$ | $\to$ | $\mathtt{P}''_l$ | $\mathtt{D}''_l$ | $\kappa_l$ |
| | | $\mid$ | $\mathtt{P}''_l$ | $\mathtt{P}''_m$ | $\kappa_l(1-s_l)v_{lm}$ |
| | | $\mid$ | $\mathtt{P}''_m$ | | $\mathcal{B}_{lm}$ |
| | | $\mid$ | $\mathtt{D}''_l$ | | $\kappa_l \eta(\mathtt{C}_l)$ |

where $\mathcal{A}, \mathcal{B}$ are the $\mathcal{N} \times \mathcal{N}$ transition matrices

$$\begin{aligned} \mathcal{A}_{lm} &= \kappa_l \left(\eta(\mathtt{M}_l)\delta_{lm} + \eta(\mathtt{L}_l)\alpha_l \left(1-\beta_l\left(1-\eta(\mathtt{N}_l)\right)\right)(1-s_l)\, v_{lm}\right) \\ \mathcal{B}_{lm} &= \kappa_l \left(\eta(\mathtt{I}_l)\delta_{lm} + \eta(\mathtt{C}_l)(1-s_l)v_{lm}\right) \end{aligned}$$

Letting $\mathcal{G} = (I-\mathcal{A})^{-1}$ and $\mathcal{H} = (I-\mathcal{B})^{-1}$ be the geometric series sums of these matrices, update the rules to sum over all $\mathtt{L}''_l \to \mathtt{L}''_m$, $\mathtt{C}''_l \to \mathtt{C}''_m$, and $\mathtt{P}''_l \to \mathtt{P}''_m$ transition chains explicitly

| Symbol | LHS | | RHS | | Probability |
|---|---|---|---|---|---|
| **L**ink | $\mathtt{L}'''_l$ | $\to$ | $\mathtt{L}'''_m$ | $\mathtt{M}'''_m$ | $\mathcal{G}_{lm}\kappa_m$ |
| | | $\mid$ | $\mathtt{L}'''_m$ | $\mathtt{L}'''_n$ | $\mathcal{G}_{lm}\kappa_m\alpha_m\left(1-\beta_m(1-\eta(\mathtt{N}_m))\right)(1-s_m)v_{mn}$ |
| | | $\mid$ | $\mathtt{M}'''_m$ | | $\mathcal{G}_{lm}\kappa_m\eta(\mathtt{L}_m)$ |
| | | $\mid$ | $\mathtt{N}'''_m$ | | $\mathcal{G}_{lm}(1-\kappa_m)\beta_m$ |
| **C**hild | $\mathtt{C}'''_l$ | $\to$ | $\mathtt{C}'''_m$ | $\mathtt{I}'''_m$ | $\mathcal{H}_{lm}\kappa_m$ |
| | | $\mid$ | $\mathtt{C}'''_m$ | $\mathtt{C}'''_n$ | $\mathcal{H}_{lm}(1-s_m)v_{mn}$ |
| | | $\mid$ | $\mathtt{I}'''_m$ | | $\mathcal{H}_{lm}\kappa_m\eta(\mathtt{C}_m)$ |
| **P**arent | $\mathtt{P}'''_l$ | $\to$ | $\mathtt{P}'''_m$ | $\mathtt{D}'''_l$ | $\mathcal{H}_{lm}\kappa_m$ |
| | | $\mid$ | $\mathtt{P}'''_m$ | $\mathtt{P}'''_n$ | $\mathcal{H}_{lm}(1-s_m)v_{mn}$ |
| | | $\mid$ | $\mathtt{D}'''_m$ | | $\mathcal{H}_{lm}\kappa_m\eta(\mathtt{C}_m)$ |

Rules for $\mathtt{X}'''_l \ldots$ where $\mathtt{X} \in \{\mathtt{M},\mathtt{S},\mathtt{E},\mathtt{N}\}$ are just copied over from the corresponding $\mathtt{X}''_l \to \ldots$ rules, changing $\mathtt{X}''$ to $\mathtt{X}'''$ on the right-hand side as well.

| Symbol | LHS | | RHS | | Probability |
|---|---|---|---|---|---|
| **M**ortal | $\mathtt{M}'''_l$ | $\to$ | $\mathtt{S}'''_l$ | | $\alpha_l$ |
| | | $\mid$ | $\mathtt{E}'''_l$ | | $1-\alpha_l$ |
| **S**urviving | $\mathtt{S}'''_l$ | $\to$ | $\mathtt{A}'''_l$ | $\mathtt{N}'''_l$ | $\beta_l$ |
| | | $\mid$ | $\mathtt{N}'''_l$ | | $\beta_l\eta(\mathtt{A}_l)$ |
| | | $\mid$ | $\mathtt{A}'''_l$ | | $1-\beta_l(1-\eta(\mathtt{N}_l))$ |
| **E**xpired | $\mathtt{E}'''_l$ | $\to$ | $\mathtt{D}'''_l$ | $\mathtt{N}'''_l$ | $\gamma_l$ |
| | | $\mid$ | $\mathtt{P}'''_m$ | $\mathtt{N}'''_l$ | $\gamma_l(1-s_l)v_{lm}$ |
| | | $\mid$ | $\mathtt{N}'''_l$ | | $\gamma_l\eta(\mathtt{D}_l)$ |
| | | $\mid$ | $\mathtt{D}'''_l$ | | $1-\gamma_l(1-\eta(\mathtt{N}_l))$ |
| | | $\mid$ | $\mathtt{P}'''_m$ | | $\left(1-\gamma_l(1-\eta(\mathtt{N}_l))\right)(1-s_l)v_{lm}$ |
| **N**ewborns | $\mathtt{N}'''_l$ | $\to$ | $\mathtt{I}'''_l$ | $\mathtt{N}'''_l$ | $\beta_l$ |
| | | $\mid$ | $\mathtt{C}'''_m$ | $\mathtt{N}'''_l$ | $\beta_l(1-s_l)v_{lm}$ |
| | | $\mid$ | $\mathtt{N}'''_l$ | | $\beta_l\eta(\mathtt{I}_l)$ |
| | | $\mid$ | $\mathtt{I}'''_l$ | | $1-\beta_l(1-\eta(\mathtt{N}_l))$ |
| | | $\mid$ | $\mathtt{C}'''_m$ | | $\left(1-\beta_l(1-\eta(\mathtt{N}_l))\right)(1-s_l)v_{lm}$ |

We use this last opportunity to reintroduce mixtures of site and fragment classes

| Symbol | LHS | | RHS | | Probability |
|---|---|---|---|---|---|
| **A**ligned | $\mathtt{A}'''_l$ | $\to$ | $\mathtt{F}_f$ | | $s_l w_{lf}$ |
| **I**nserted | $\mathtt{I}'''_l$ | $\to$ | $\mathtt{G}_f$ | | $s_l w_{lf}$ |
| **D**eleted | $\mathtt{D}'''_l$ | $\to$ | $\mathtt{R}_f$ | | $s_l w_{lf}$ |
| **F**ragment | $\mathtt{F}_f$ | $\to$ | $\mathtt{T}_{ab}$ | $\mathtt{F}_g$ | $r^{(l)}_{fg}\sum_c u_{lfc}\pi^{(c)}_a\exp(R^{(c)}T)_{ab}$ |
| | | $\mid$ | $\mathtt{T}_{ab}$ | | $\rho^{(l)}_f\sum_c u_{lfc}\pi^{(c)}_a\exp(R^{(c)}T)_{ab}$ |
| **G**enerated | $\mathtt{G}_f$ | $\to$ | $\mathtt{T}_{\bullet b}$ | $\mathtt{G}_g$ | $r^{(l)}_{fg}\sum_c u_{lfc}\pi^{(c)}_b$ |
| | | $\mid$ | $\mathtt{T}_{\bullet b}$ | | $\rho^{(l)}_f\sum_c u_{lfc}\pi^{(c)}_b$ |
| **R**emoved | $\mathtt{R}_f$ | $\to$ | $\mathtt{T}_{a\bullet}$ | $\mathtt{R}_g$ | $r^{(l)}_{fg}\sum_c u_{lfc}\pi^{(c)}_a$ |
| | | $\mid$ | $\mathtt{T}_{a\bullet}$ | | $\rho^{(l)}_f\sum_c u_{lfc}\pi^{(c)}_a$ |

*TKF Structure Tree singlet rules (a)*
*Sequence a terminals: $\{u, v\}$.*

| lhs | $\to$ | rhs | $P(a)$ |
|---|---|---|---|
| $L_a$ | $\to$ | $u\ L_a$ | $\kappa_l \pi_l(u)$ |
| | $\mid$ | $S_a\ L_a$ | $\kappa_l \pi_l(S)$ |
| | $\mid$ | $\epsilon$ | $1 - \kappa_l$ |
| $S_a$ | $\to$ | $u\ S_a\ v$ | $\kappa_s \pi_s(uv)$ |
| | $\mid$ | $L_a$ | $1 - \kappa_s$ |

Table C.2: TKF Structure Tree. Singlet rule-set for $a$.

and add the top-level rule:

| Symbol | LHS | | RHS | Probability |
|---|---|---|---|---|
| **B**egin | B | $\to$ | $\mathtt{L}_1'''$ | 1 |
| | | $\mid$ | $\epsilon$ | $\eta(\mathtt{L}_1)$ |

With no $\epsilon$-productions and a strictly acyclic topological transition sort order on nonterminals of $\mathtt{B} \to \mathtt{L} \to \mathtt{M} \to (\mathtt{S} \to (\mathtt{A},\ \mathtt{N} \to \mathtt{I} \to \mathtt{C}),\ \mathtt{E} \to \mathtt{D} \to \mathtt{P})$, this grammar is ready for an Inside parser.

At this point our model is the desired

$$\begin{aligned} M_l &= \text{Links}(\text{Mix}_{k\sim s}(L_l^{(k)}); \lambda_l, \mu_l) \\ L_l^{(1)} &= \text{HMM}(\{Q^{(c)}, \pi^{(c)}\}_c; r^{(l)}, u_l) \\ L_l^{(0)} &= \text{Mix}_{m\sim v_m}(M_m) \end{aligned}$$

##### C.13.2 Example Two: The TKF Structure Tree (TKFST)

Consider now the RNA evolutionary model derived in (22).

That model has stem ($S$) and loop ($L$) sequences, each of which is a TKF91 sequence with its own rates. Stems are sequences of base pairs, loops of individual bases.

The model was developed as a proof of concept, but suffers similar deficiencies to TKF91 concerning the absence of an affine gap penalty, as well as a lack of basepair stacking effects or other empirically observed features of biological RNA structures.

**Parameters** Let $\overline{xyz}$ denote reverse complement e.g. $\overline{AAG} = CTT$.

Parameters: insertion and deletion rates $\lambda_F < \mu_F$, fragment extension probability $r_F$, substitution rate matrix $Q_F$, equilibrium probability vector $q_F$ (so $q_x Q_F = 0$) for $F \in \{R, N, G, S, L, C\}$. Splice donor/acceptor site distribution $q_{D1}$, $q_{D2}$, $q_{A1}$, $q_{A2}$. Region-type probabilities $p_G + p_N + p_S + p_C = 1$. Intron probability $p_I$.

The $Q_N$, $Q_S$, $Q_L$ and $Q_C$ models should be strand-invariant, so e.g. $Q_N(x_1, x_2) = Q_N(\overline{x_1}, \overline{x_2})$.

**Functions** For $F \in \{R, N, G, S, L, C\}$:

$$\kappa_F = \left(1 - \frac{\lambda_F}{\mu_F}\right)(1 - r_F)$$

*TKF Structure Tree singlet rules (b)*
*Sequence b terminals: $\{w, x\}$.*

| lhs | $\rightarrow$ | rhs | $P(b)$ |
|---|---|---|---|
| $L_b$ | $\rightarrow$ | $w\ L_b$ | $\kappa_l \pi_l(w)$ |
| | $\mid$ | $S_b\ L_b$ | $\kappa_l \pi_l(S)$ |
| | $\mid$ | $\epsilon$ | $1 - \kappa_l$ |
| $S_b$ | $\rightarrow$ | $w\ S_b\ x$ | $\kappa_s \pi_s(wx)$ |
| | $\mid$ | $L_b$ | $1 - \kappa_s$ |

Table C.3: TKF Structure Tree. Singlet rule-set for $b$.

**The Stationary Grammar**

| LHS | $\rightarrow$ | RHS | Transition | Emission |
|---|---|---|---|---|
| GENOME | $\rightarrow$ | REGION GENOME | $1 - \kappa_R$ | |
| | $\mid$ | $\epsilon$ | $\kappa_R$ | |
| REGION | $\rightarrow$ | INTER | $p_N$ | |
| | $\mid$ | FWDCDS | $p_G/2$ | |
| | $\mid$ | REVCDS | $p_G/2$ | |
| | $\mid$ | STRUCT | $p_S$ | |
| | $\mid$ | CONS | $p_C$ | |
| INTER | $\rightarrow$ | $x_1$ INTER | $1 - \kappa_N$ | $q_N(x_1)$ |
| | $\mid$ | $\epsilon$ | $\kappa_N$ | |
| FWDCDS | $\rightarrow$ | FWDCOD FWDCDS | $1 - \kappa_G$ | |
| | $\mid$ | $\epsilon$ | $\kappa_G$ | |
| FWDCOD | $\rightarrow$ | $x_1\ x_2\ x_3$ | $1 - p_I$ | $q_G(xyz)$ |
| | $\mid$ | $x_1\ x_2\ x_3$ FWDINT | $p_I/3$ | $q_G(xyz)$ |
| | $\mid$ | $x_1\ x_2$ FWDINT $x_3$ | $p_I/3$ | $q_G(xyz)$ |
| | $\mid$ | $x_1$ FWDINT $x_2\ x_3$ | $p_I/3$ | $q_G(xyz)$ |
| FWDINT | $\rightarrow$ | $x_1\ x_2$ GENOME $x_3\ x_4$ | 1 | $q_{D1}(x_1) q_{D2}(x_2) q_{A1}(x_3) q_{A2}(x_4)$ |
| REVCDS | $\rightarrow$ | REVCDS REVCOD | $1 - \kappa_G$ | |
| | $\mid$ | $\epsilon$ | $\kappa_G$ | |
| REVCOD | $\rightarrow$ | $x_1\ x_2\ x_3$ | $1 - p_I$ | $q_G(\overline{xyz})$ |
| | $\mid$ | $x_1\ x_2\ x_3$ REVINT | $p_I/3$ | $q_G(\overline{xyz})$ |
| | $\mid$ | $x_1\ x_2$ REVINT $x_3$ | $p_I/3$ | $q_G(\overline{xyz})$ |
| | $\mid$ | $x_1$ REVINT $x_2\ x_3$ | $p_I/3$ | $q_G(\overline{xyz})$ |
| REVINT | $\rightarrow$ | $x_1\ x_2$ GENOME $x_3\ x_4$ | 1 | $q_{D1}(\overline{x_4}) q_{D2}(\overline{x_3}) q_{A1}(\overline{x_2}) q_{A2}(\overline{x_1})$ |
| STRUCT | $\rightarrow$ | $x_1$ STRUCT $x_2$ | $1 - \kappa_S$ | $q_S(xy)$ |
| | $\mid$ | LOOP | $\kappa_S$ | |
| LOOP | $\rightarrow$ | $x_1$ LOOP | $1 - \kappa_L$ | $q_L(x_1)$ |
| | $\mid$ | $\epsilon$ | $\kappa_L$ | |
| CONS | $\rightarrow$ | $x_1$ CONS | $1 - \kappa_C$ | $q_C(x_1)$ |
| | $\mid$ | $\epsilon$ | $\kappa_C$ | |

*TKF Structure Tree pair rules* $(a \xrightarrow{t} b)$

*Sequence a terminals:* $\{u, v\}$. *Sequence b terminals:* $\{w, x\}$.

| lhs | $\to$ | rhs | $P(a)$ | $P(b\|a)$ |
|---|---|---|---|---|
| $L_{ab}$ | $\to$ | $u\ w\ L_{ab}$ | $\kappa_l \pi_l(u)$ | $(1-\beta_l)\alpha_l M_l(u,w)$ |
| | $\|$ | $w\ L_{ab}$ | $1$ | $\beta_l \pi_l(w)$ |
| | $\|$ | $u\ L_{a\bullet b}$ | $\kappa_l \pi_l(u)$ | $(1-\beta_l)(1-\alpha_l)$ |
| | $\|$ | $S_{ab}\ L_{ab}$ | $\kappa_l \pi_l(S)$ | $(1-\beta_l)\alpha_l$ |
| | $\|$ | $S_b\ L_{ab}$ | $1$ | $\beta_l \pi_l(S)$ |
| | $\|$ | $S_a\ L_{a\bullet b}$ | $\kappa_l \pi_l(S)$ | $(1-\beta_l)(1-\alpha_l)$ |
| | $\|$ | $\epsilon$ | $1-\kappa_l$ | $1-\beta_l$ |
| $S_{ab}$ | $\to$ | $u\ w\ S_{ab}\ x\ v$ | $\kappa_s \pi_s(uv)$ | $(1-\beta_s)\alpha_s M_s(uv,wx)$ |
| | $\|$ | $w\ S_{ab}\ x$ | $1$ | $\beta_s \pi_s(wx)$ |
| | $\|$ | $u\ S_{a\bullet b}\ v$ | $\kappa_s \pi_s(uv)$ | $(1-\beta_s)(1-\alpha_s)$ |
| | $\|$ | $L_{ab}$ | $1-\kappa_s$ | $1-\beta_s$ |
| $L_{a\bullet b}$ | $\to$ | $u\ w\ L_{ab}$ | $\kappa_l \pi_l(u)$ | $(1-\gamma_l)\alpha_l M_l(u,w)$ |
| | $\|$ | $w\ L_{ab}$ | $1$ | $\gamma_l \pi_l(w)$ |
| | $\|$ | $u\ L_{a\bullet b}$ | $\kappa_l \pi_l(u)$ | $(1-\gamma_l)(1-\alpha_l)$ |
| | $\|$ | $S_{ab}\ L_{ab}$ | $\kappa_l \pi_l(S)$ | $(1-\gamma_l)\alpha_l$ |
| | $\|$ | $S_b\ L_{ab}$ | $1$ | $\gamma_l \pi_l(S)$ |
| | $\|$ | $S_a\ L_{a\bullet b}$ | $\kappa_l \pi_l(S)$ | $(1-\gamma_l)(1-\alpha_l)$ |
| | $\|$ | $\epsilon$ | $1-\kappa_l$ | $1-\gamma_l$ |
| $S_{a\bullet b}$ | $\to$ | $u\ w\ S_{ab}\ x\ v$ | $\kappa_s \pi_s(uv)$ | $(1-\gamma_s)\alpha_s M_s(uv,wx)$ |
| | $\|$ | $w\ S_{ab}\ x$ | $1$ | $\gamma_s \pi_s(wx)$ |
| | $\|$ | $u\ S_{a\bullet b}\ v$ | $\kappa_s \pi_s(uv)$ | $(1-\gamma_s)(1-\alpha_s)$ |
| | $\|$ | $L_{ab}$ | $1-\kappa_s$ | $1-\gamma_s$ |

Table C.4: TKF Structure Tree. Pair rule-set for $a \xrightarrow{t} b$ branch. Requires singlet rule-sets for $a$ and $b$.

**The Joint Finite-Time Grammar** The general rules for forming the joint grammar from the conditional grammar are as follows. For every nonterminal of the following form (here L, R, and/or E are allowed to be $\epsilon$)

| LHS | $\rightarrow$ | RHS | Transition | Emission |
|---|---|---|---|---|
| F | $\rightarrow$ | L F R | $1-\kappa_F$ | $q_F(LR)$ |
| | $\mid$ | E | $\kappa_F$ | |

...that is, for GENOME, INTER, FWDCDS, REVCDS, STRUCT, LOOP, and CONS (with $F \in \{R, N, G, S, L, C\}$), replace these rules with

| LHS | $\rightarrow$ | RHS | Transition | Emission |
|---|---|---|---|---|
| $\mathtt{F}_M$ | $\rightarrow$ | $\mathtt{L}_M\ \mathtt{F}_M\ \mathtt{R}_M$ | $(1-\beta_{M,F})(1-\kappa_F)\alpha_F$ | $q_F(LR_x)\exp(Q_F t)(LR_x, LR_y)$ |
| | $\mid$ | $\mathtt{L}_Y\ \mathtt{F}_M\ \mathtt{R}_Y$ | $\beta_{M,F}$ | $q_F(LR_y)$ |
| | $\mid$ | $\mathtt{L}_X\ \mathtt{F}_D\ \mathtt{R}_X$ | $(1-\beta_{M,F})(1-\kappa_F)(1-\alpha_F)$ | $q_F(LR_x)$ |
| | $\mid$ | $\mathtt{E}_M$ | $(1-\beta_{M,F})\kappa_F$ | |
| $\mathtt{F}_D$ | $\rightarrow$ | $\mathtt{L}_M\ \mathtt{F}_M\ \mathtt{R}_M$ | $(1-\beta_{D,F})(1-\kappa_F)\alpha_F$ | $q_F(LR_x)\exp(Q_F t)(LR_x, LR_y)$ |
| | $\mid$ | $\mathtt{L}_Y\ \mathtt{F}_M\ \mathtt{R}_Y$ | $\beta_{D,F}$ | $q_F(LR_y)$ |
| | $\mid$ | $\mathtt{L}_X\ \mathtt{F}_D\ \mathtt{R}_X$ | $(1-\beta_{D,F})(1-\kappa_F)(1-\alpha_F)$ | $q_F(LR_x)$ |
| | $\mid$ | $\mathtt{E}_M$ | $(1-\beta_{D,F})\kappa_F$ | |

...that is, two versions $\mathtt{F}_M$ and $\mathtt{F}_D$, with different $\beta$'s for each type.

For every other nonterminal N, there need to be three versions $\mathtt{N}_M$, $\mathtt{N}_X$ and $\mathtt{N}_Y$, with outgoing rules for each type going to other nonterminals of the same type.

**Top level.**

| LHS | $\rightarrow$ | RHS | Transition | Emission |
|---|---|---|---|---|
| $\mathtt{GENOME}_k$ | $\rightarrow$ | $\mathtt{REGION}_M\ \mathtt{GENOME}_M$ | $(1-\beta_{k,R})(1-\kappa_R)\alpha_R$ | |
| | $\mid$ | $\mathtt{REGION}_Y\ \mathtt{GENOME}_M$ | $\beta_{k,R}$ | |
| | $\mid$ | $\mathtt{REGION}_X\ \mathtt{GENOME}_D$ | $(1-\beta_{k,R})(1-\kappa_R)(1-\alpha_R)$ | |
| | $\mid$ | $\epsilon$ | $(1-\beta_{k,R})\kappa_R$ | |
| $\mathtt{REGION}_j$ | $\rightarrow$ | $\mathtt{INTER}_j$ | $p_N$ | |
| | $\mid$ | $\mathtt{FWDCDS}_j$ | $p_G/2$ | |
| | $\mid$ | $\mathtt{REVCDS}_j$ | $p_G/2$ | |
| | $\mid$ | $\mathtt{STRUCT}_j$ | $p_S$ | |
| | $\mid$ | $\mathtt{CONS}_j$ | $p_C$ | |
| INTER | $\rightarrow$ | $x_1$ INTER | $1-\kappa_N$ | $q_N(x_1)$ |
| | $\mid$ | $\epsilon$ | $\kappa_N$ | |

**Coding sequences.**

| LHS | $\rightarrow$ | RHS | Transition | Emission |
|---|---|---|---|---|
| FWDCDS | $\rightarrow$ | FWDCOD FWDCDS | $1-\kappa_G$ | |
| | $\mid$ | $\epsilon$ | $\kappa_G$ | |
| FWDCOD | $\rightarrow$ | $x_1\ x_2\ x_3$ | $1-p_I$ | $q_G(xyz)$ |
| | $\mid$ | $x_1\ x_2\ x_3$ FWDINT | $p_I/3$ | $q_G(xyz)$ |
| | $\mid$ | $x_1\ x_2$ FWDINT $x_3$ | $p_I/3$ | $q_G(xyz)$ |
| | $\mid$ | $x_1$ FWDINT $x_2\ x_3$ | $p_I/3$ | $q_G(xyz)$ |
| FWDINT | $\rightarrow$ | $x_1\ x_2$ GENOME $x_3\ x_4$ | 1 | $q_{D1}(x_1)q_{D2}(x_2)q_{A1}(x_3)q_{A2}(x_4)$ |

| LHS | → | RHS | Transition | Emission |
|---|---|---|---|---|
| `REVCDS` | → | `REVCDS` `REVCOD` | $1 - \kappa_G$ | |
| | \| | $\epsilon$ | $\kappa_G$ | |
| `REVCOD` | → | $x_1\ x_2\ x_3$ | $1 - p_I$ | $q_G(\overline{xyz})$ |
| | \| | $x_1\ x_2\ x_3$ `REVINT` | $p_I/3$ | $q_G(\overline{xyz})$ |
| | \| | $x_1\ x_2$ `REVINT` $x_3$ | $p_I/3$ | $q_G(\overline{xyz})$ |
| | \| | $x_1$ `REVINT` $x_2\ x_3$ | $p_I/3$ | $q_G(\overline{xyz})$ |
| `REVINT` | → | $x_1\ x_2$ `GENOME` $x_3\ x_4$ | 1 | $q_{D1}(\overline{x_4})q_{D2}(\overline{x_3})q_{A1}(\overline{x_2})q_{A2}(\overline{x_1})$ |

**RNA structures.**

| LHS | → | RHS | Transition | Emission |
|---|---|---|---|---|
| `STRUCT` | → | $x_1$ `STRUCT` $x_2$ | $1 - \kappa_S$ | $q_S(xy)$ |
| | \| | `LOOP` | $\kappa_S$ | |
| `LOOP` | → | $x_1$ `LOOP` | $1 - \kappa_L$ | $q_L(x_1)$ |
| | \| | $\epsilon$ | $\kappa_L$ | |
| `CONS` | → | $x_1$ `CONS` | $1 - \kappa_C$ | $q_C(x_1)$ |
| | \| | $\epsilon$ | $\kappa_C$ | |

##### C.13.3 Example Three: The TKF Basepair Stack (TKFStack)

The simple TKF Structure Tree models RNA secondary structure with alternating stems (basepair sequences) and loops (single-nucleotide sequences). While a useful proof of concept, it lacks basepair stacking, multiloop junctions, bulges, and internal loops—features critical for realistic RNA structure modeling.

We now define an enhanced stem-loop grammar that incorporates these features while remaining within the TKF evolutionary framework. We then show how evolution elaboration, profile SCFG construction, and the triplet model for progressive reconstruction all apply to this grammar.

**Parameters** The grammar has the following parameters:

- Stem link TKF91 rates $\lambda_S < \mu_S$, giving $\kappa_S = \lambda_S/\mu_S$.
- Loop link TKF91 rates $\lambda_L < \mu_L$, giving $\kappa_L = \lambda_L/\mu_L$.
- Bulge extension probability $r_B$.
- Stacked-pair fragment extension probability $r_K$.
- Stem link type probabilities $p_{\text{bp}} + p_{\text{st}} + p_{\text{bu}} = 1$.
- Loop link type probabilities $p_{\text{lf}} + p_{\text{rf}} + p_{\text{sl}} = 1$.
- Nucleotide equilibrium $\pi(c)$ over $\mathcal{A} = \{A, C, G, U\}$, with rate matrix $Q$.
- Single basepair equilibrium $\pi_{\text{bp}}(c_L, c_R)$ over $|\mathcal{A}|^2 = 16$ states, rate matrix $Q_{\text{bp}}$.
- Closing basepair equilibrium $\pi_{\text{cl}}(c_L, c_R)$ over 16 states, rate matrix $Q_{\text{cl}}$.
- Stacked-pair equilibrium $\pi_K(c_L^1, c_L^2, c_R^2, c_R^1)$ over the $6^2 = 36$ canonical stacked pairs, rate matrix $Q_K$. A stacked pair consists of two consecutive canonical basepairs $(c_L^1, c_R^1)$ and $(c_L^2, c_R^2)$; the state space is restricted to the six Watson-Crick and wobble pairs (AU, CG, GC, UA, GU, UG) for each position, giving 36 states rather than the full $16^2 = 256$.

**Nonterminals and Productions** A *stem-loop* consists of a stem (nested basepairs with decorations), a closing basepair, and a loop (with possible multiloop branches). The start symbol is $\mathsf{SL}$ (stem-loop).

**Stem-loop:**

$$\mathsf{SL} \to \mathsf{STEM} \qquad \text{weight } 1 \qquad (C.198)$$

**Stem (TKF91 sequence of stem-links, from outer to inner):**

$$\begin{aligned}
\mathsf{STEM} &\to c_L\ \mathsf{STEM}\ c_R && \kappa_S\, p_{\text{bp}}\, \pi_{\text{bp}}(c_L, c_R) && \text{[single basepair, LR]} && (C.199)\\
&\to c_L^1\, c_L^2\ \mathsf{STEM}\ c_R^2\, c_R^1 && \kappa_S\, p_{\text{st}}\, (1 - r_K)\, \pi_K(\cdot) && \text{[terminal stacked pair, LLRR]} && (C.200)\\
&\to c_L^1\, c_L^2\ \mathsf{STACK}\ c_R^2\, c_R^1 && \kappa_S\, p_{\text{st}}\, r_K\, \pi_K(\cdot) && \text{[extended stacked pair, LLRR]} && (C.201)\\
&\to \mathsf{LDECO}\ \mathsf{STEM}\ \mathsf{RDECO} && \kappa_S\, p_{\text{bu}} && \text{[bulge]} && (C.202)\\
&\to \mathsf{CLOSE}\ \mathsf{LOOP} && 1 - \kappa_S && \text{[end stem]} && (C.203)
\end{aligned}$$

A bulge link (C.202) represents an internal loop, a single-sided bulge, or a multihelix junction branch between basepairs. All non-basepair content between consecutive basepairs is consolidated into a single TKF link type whose internal structure is governed by $\mathsf{LDECO}$ and $\mathsf{RDECO}$.

**Bulge decorations (L-side and R-side content):**

$$\begin{aligned}
\mathsf{LDECO} &\to \mathsf{LFRAG}\ \mathsf{LDECO} && \text{[L-fragment, then more L-content]} && (C.204)\\
&\to \mathsf{SL}\ \mathsf{LDECO} && \text{[left branch: nested stem-loop]} && (C.205)\\
&\to \epsilon && \text{[end L-decorations]} && (C.206)
\end{aligned}$$

$$\begin{aligned}
\mathsf{RDECO} &\to \mathsf{RDECO}\ \mathsf{RFRAG} && \text{[R-fragment, more R-content]} && (C.207)\\
&\to \mathsf{RDECO}\ \mathsf{SL} && \text{[right branch: nested stem-loop]} && (C.208)\\
&\to \epsilon && \text{[end R-decorations]} && (C.209)
\end{aligned}$$

$\mathsf{LDECO}$ generates all content on the $5'$ side (between the outer basepair's L-half and the continuation), while $\mathsf{RDECO}$ generates all content on the $3'$ side (between the continuation and the outer basepair's R-half). An internal loop has both $\mathsf{LDECO} \neq \epsilon$ and $\mathsf{RDECO} \neq \epsilon$; a single-sided bulge has content on only one side; a multihelix junction branch has $\mathsf{SL}$ nested within $\mathsf{LDECO}$ or $\mathsf{RDECO}$.

**Stacked-pair fragment continuation (LLRR emission):**

$$\begin{aligned}
\mathsf{STACK} &\to c_L^1\, c_L^2\ \mathsf{STEM}\ c_R^2\, c_R^1 && (1 - r_K)\, \pi_K(\cdot) && \text{[terminal]} && (C.210)\\
&\to c_L^1\, c_L^2\ \mathsf{STACK}\ c_R^2\, c_R^1 && r_K\, \pi_K(\cdot) && \text{[extend]} && (C.211)
\end{aligned}$$

**Closing basepair (LR emission):**

$$\mathsf{CLOSE} \to c_L\ \mathsf{CLOSE}'\ c_R \qquad \pi_{\text{cl}}(c_L, c_R) \qquad (C.212)$$

where $\mathsf{CLOSE}'$ is a unit nonterminal ($\mathsf{CLOSE}' \to \epsilon$, weight 1) that serves as a placeholder for the inside of the closing basepair.

**Loop (TKF91 sequence of loop-links):**

$$\begin{aligned}
\mathsf{LOOP} &\to \mathsf{LOOPLINK}\ \mathsf{LOOP} && \kappa_L && \text{[add loop link]} && (C.213)\\
&\to \epsilon && 1 - \kappa_L && \text{[end loop]} && (C.214)
\end{aligned}$$

**Loop link types:**

$$\mathsf{LOOPLINK} \to \mathsf{LFRAG} \qquad p_{\mathrm{lf}} \qquad \text{[L-fragment]} \tag{C.215}$$

$$\to \mathsf{RFRAG} \qquad p_{\mathrm{rf}} \qquad \text{[R-fragment]} \tag{C.216}$$

$$\to \mathsf{SL} \qquad p_{\mathrm{sl}} \qquad \text{[nested stem-loop (multiloop)]} \tag{C.217}$$

**Unpaired nucleotide fragments in loops (geometric length $\geq 1$):**

$$\mathsf{LFRAG} \to c\,\mathsf{LFRAG} \qquad r_L\,\pi(c) \qquad \text{[extend, L-emission]} \tag{C.218}$$

$$\to c \qquad (1-r_L)\,\pi(c) \qquad \text{[terminal, L-emission]} \tag{C.219}$$

$$\mathsf{RFRAG} \to \mathsf{RFRAG}\,c \qquad r_R\,\pi(c) \qquad \text{[extend, R-emission]} \tag{C.220}$$

$$\to c \qquad (1-r_R)\,\pi(c) \qquad \text{[terminal, R-emission]} \tag{C.221}$$

**Emission Types and Span Tracking** The grammar has four emission patterns, each determining how terminals consume positions from the span $[i,j]$ of the input sequence:

1. **L-emission** ($c$): consumes from the left end, advancing $i \to i+1$. Used by LFRAG, LDECO content.

2. **R-emission** ($c$): consumes from the right end, retreating $j \to j-1$. Used by RFRAG, RDECO content.

3. **LR-emission** ($c_L, c_R$): consumes from both ends simultaneously, $i \to i+1$, $j \to j-1$. Used by STEM (single basepair), CLOSE.

4. **LLRR-emission** ($c_L^1, c_L^2, c_R^2, c_R^1$): consumes two from each end, $i \to i+2$, $j \to j-2$. Used by STEM (stacked pair), STACK. Each LLRR unit represents a *stacked dinucleotide pair*: two consecutive canonical basepairs $(c_L^1, c_R^1)$ and $(c_L^2, c_R^2)$ treated as a single evolutionary unit with nearest-neighbor stacking energy. The state space is restricted to the $6 \times 6 = 36$ combinations of canonical (Watson-Crick and wobble) pairs.

*Remark* C.28 (L/R distinction in terminals vs. productions). The terminal alphabet is $\mathcal{A} = \{A, C, G, U\}$ without L/R copies: both L-emission and R-emission produce characters from the same alphabet, differing only in which end of the span $[i,j]$ is consumed. At leaf nodes, the observed sequence carries no L/R annotation.

However, the L/R distinction is essential at the *production* level and therefore in the *profile SCFG*. Each nonterminal instance in a profile has a fixed emission direction (L, R, LR, or LLRR), determined by the parse tree from which it was extracted—analogous to the MATL, MATR, and MATP node types in Infernal covariance models (38). The branch PTT must preserve emission-direction compatibility when mapping parent productions to child productions. Consequently, a profile position that is LR-emitting (basepaired) at one node of the phylogeny may correspond to separate L-emitting and R-emitting positions (unpaired) at another node, reflecting structural change along the evolutionary lineage.

*Remark* C.29 (Structural interpretation). Consider a stem with links (from outer to inner): basepair $(c_L^1, c_R^1)$, bulge (left-side $b_1 b_2$, right-side branch $\mathsf{SL}'$), basepair $(c_L^2, c_R^2)$. The yield is:

$$c_L^1\; b_1\; b_2\; c_L^2\; \text{[loop]}\; c_R^2\; [\text{yield}(\mathsf{SL}')]\; c_R^1$$

The bulge nucleotides $b_1, b_2$ appear on the $5'$ side (via LDECO), while the branch structure appears on the $3'$ side (via RDECO). This represents an internal loop with unpaired nucleotides on the $5'$ strand and a branching sub-structure on the $3'$ strand—a configuration that the consolidated bulge link type (C.202) captures as a single TKF event.

**The Pair Grammar via Evolution Elaboration** Applying the evolution elaboration rules (Appendix C.12) to the singlet grammar produces the pair grammar for an ancestor–descendant pair separated by evolutionary time $T$. Each nonterminal participating in a TKF91 link sequence (STEM, LOOP) gains M/I/D versions with the standard TKF91 transition weights $(\alpha, \beta, \gamma)$ derived from $(\lambda, \mu, T)$.

**Stem Link Elaboration.** The stem is a TKF91 sequence of links with rates $(\lambda_S, \mu_S)$. Each STEM nonterminal becomes $\mathsf{STEM}_{\mathtt{M}}$ (post-match/insert) and $\mathsf{STEM}_{\mathtt{D}}$ (post-delete):

$$\mathsf{STEM}_{\mathtt{M}} \to c_L^x\, c_L^y\, \mathsf{STEM}_{\mathtt{M}}\, c_R^y\, c_R^x \quad (1-\beta_S)\kappa_S\, p_{\rm bp}\, \alpha_S\, \pi_{\rm bp}(c_L^x, c_R^x)\, P_{\rm bp}(c_L^y, c_R^y | c_L^x, c_R^x) \tag{C.222}$$

$$\to c_L^y\, \mathsf{STEM}_{\mathtt{M}}\, c_R^y \quad \beta_S\, \kappa_S\, p_{\rm bp}\, \pi_{\rm bp}(c_L^y, c_R^y) \quad \text{[BP insert]} \tag{C.223}$$

$$\to c_L^x\, \mathsf{STEM}_{\mathtt{D}}\, c_R^x \quad (1-\beta_S)\kappa_S\, p_{\rm bp}\, (1-\alpha_S)\, \pi_{\rm bp}(c_L^x, c_R^x) \quad \text{[BP delete]} \tag{C.224}$$

with analogous rules for $\mathsf{STEM}_{\mathtt{D}}$ using $\gamma_S$ in place of $\beta_S$.

For matched single basepairs (C.222), the nesting is $c_L^x\, c_L^y\ \cdots\ c_R^y\, c_R^x$: ancestor terminals on the outside, descendant terminals on the inside, preserving palindromic structure. For inserted basepairs (C.223), only descendant terminals appear (LR emission). For deleted basepairs (C.224), only ancestor terminals appear (LR emission).

**Stacked-Pair Elaboration.** Matched stacked pairs have LLRR emission on both ancestor and descendant, yielding an $L^4R^4$ nesting pattern in the pair grammar:

$$\mathsf{STEM}_{\mathtt{M}} \to c_{L1}^x\, c_{L2}^x\, c_{L1}^y\, c_{L2}^y\, \mathsf{STEM}_{\mathtt{M}}\, c_{R2}^y\, c_{R1}^y\, c_{R2}^x\, c_{R1}^x \quad (1-\beta_S)\kappa_S\, p_{\rm st}\, (1-r_K)\, \alpha_S\, \pi_K(\cdot)\, P_K(\cdot|\cdot) \tag{C.225}$$

with inserted stacked pairs emitting only descendant LLRR, and deleted stacked pairs emitting only ancestor LLRR.

**Bulge Elaboration.** A bulge link (C.202) gains M/I/D versions like any other stem link. A matched bulge has the form $\mathsf{LDECO}_{\mathtt{M}}\ \mathsf{STEM}_{\mathtt{M}}\ \mathsf{RDECO}_{\mathtt{M}}$, with each side elaborated independently.

Within LDECO, each sub-element (LFRAG, nested SL) gains M/I/D versions with L-emission direction preserved. For example, a matched left-branch within a bulge produces:

$$\mathsf{LDECO}_{\mathtt{M}} \to \mathsf{SL}_{\mathtt{M}}\ \mathsf{LDECO}_{\mathtt{M}} \quad \alpha_{\rm deco}\, P(\text{branch}) \quad \text{[matched left branch]} \tag{C.226}$$

$$\to \mathsf{SL}_{\mathtt{I}}\ \mathsf{LDECO}_{\mathtt{M}} \quad \beta_{\rm deco} \quad \text{[inserted left branch]} \tag{C.227}$$

$$\to \mathsf{SL}_{\mathtt{D}}\ \mathsf{LDECO}_{\mathtt{D}} \quad (1-\alpha_{\rm deco})\, P(\text{branch}) \quad \text{[deleted left branch]} \tag{C.228}$$

Here $\mathsf{SL}_{\mathtt{M}}$ generates aligned ancestor–descendant sub-structures, $\mathsf{SL}_{\mathtt{I}}$ generates descendant-only sub-structures, and $\mathsf{SL}_{\mathtt{D}}$ generates ancestor-only sub-structures.

RDECO is elaborated symmetrically, with R-emission direction preserved for RFRAG elements and right-side branches.

**Loop Elaboration.** The loop is a TKF91 sequence of loop-links with rates $(\lambda_L, \mu_L)$. Elaboration proceeds identically to the stem: each $\mathsf{LOOP}$ becomes $\mathsf{LOOP}_\mathtt{M}/\mathsf{LOOP}_\mathtt{D}$, and each loop-link type ($\mathsf{LFRAG}$, $\mathsf{RFRAG}$, nested $\mathsf{SL}$) gains `M`/`I`/`D` versions.

Nested stem-loops within the loop (multiloop junctions) are handled recursively: $\mathsf{SL}_\mathtt{M}$ aligns both ancestor and descendant sub-structures, allowing multiloop branches to be independently inserted, deleted, or matched.

##### C.13.4 Example Four: The TKF Genome

**Parameters** Let $\overline{xyz}$ denote reverse complement e.g. $\overline{AAG} = CTT$.

Parameters: insertion and deletion rates $\lambda_F < \mu_F$, fragment extension probability $r_F$, substitution rate matrix $Q_F$, equilibrium probability vector $q_F$ (so $q_x Q_F = 0$) for $F \in \{R, N, G, S, L, C\}$. Splice donor/acceptor site distribution $q_{D1}$, $q_{D2}$, $q_{A1}$, $q_{A2}$. Region-type probabilities $p_G + p_N + p_S + p_C = 1$. Intron probability $p_I$.

The $Q_N$, $Q_S$, $Q_L$ and $Q_C$ models should be strand-invariant, so e.g. $Q_N(x_1, x_2) = Q_N(\overline{x_1}, \overline{x_2})$.

**Functions** For $F \in \{R, N, G, S, L, C\}$:

$$\kappa_F = \left(1 - \frac{\lambda_F}{\mu_F}\right)(1 - r_F)$$

**The Stationary Grammar**

| LHS | $\to$ | RHS | Transition | Emission |
|---|---|---|---|---|
| GENOME | $\to$ | REGION GENOME | $1-\kappa_R$ | |
| | $\mid$ | $\epsilon$ | $\kappa_R$ | |
| REGION | $\to$ | INTER | $p_N$ | |
| | $\mid$ | FWDCDS | $p_G/2$ | |
| | $\mid$ | REVCDS | $p_G/2$ | |
| | $\mid$ | STRUCT | $p_S$ | |
| | $\mid$ | CONS | $p_C$ | |
| INTER | $\to$ | $x_1$ INTER | $1-\kappa_N$ | $q_N(x_1)$ |
| | $\mid$ | $\epsilon$ | $\kappa_N$ | |
| FWDCDS | $\to$ | FWDCOD FWDCDS | $1-\kappa_G$ | |
| | $\mid$ | $\epsilon$ | $\kappa_G$ | |
| FWDCOD | $\to$ | $x_1\ x_2\ x_3$ | $1-p_I$ | $q_G(xyz)$ |
| | $\mid$ | $x_1\ x_2\ x_3$ FWDINT | $p_I/3$ | $q_G(xyz)$ |
| | $\mid$ | $x_1\ x_2$ FWDINT $x_3$ | $p_I/3$ | $q_G(xyz)$ |
| | $\mid$ | $x_1$ FWDINT $x_2\ x_3$ | $p_I/3$ | $q_G(xyz)$ |
| FWDINT | $\to$ | $x_1\ x_2$ GENOME $x_3\ x_4$ | 1 | $q_{D1}(x_1)q_{D2}(x_2)q_{A1}(x_3)q_{A2}(x_4)$ |
| REVCDS | $\to$ | REVCDS REVCOD | $1-\kappa_G$ | |
| | $\mid$ | $\epsilon$ | $\kappa_G$ | |
| REVCOD | $\to$ | $x_1\ x_2\ x_3$ | $1-p_I$ | $q_G(\overline{xyz})$ |
| | $\mid$ | $x_1\ x_2\ x_3$ REVINT | $p_I/3$ | $q_G(\overline{xyz})$ |
| | $\mid$ | $x_1\ x_2$ REVINT $x_3$ | $p_I/3$ | $q_G(\overline{xyz})$ |
| | $\mid$ | $x_1$ REVINT $x_2\ x_3$ | $p_I/3$ | $q_G(\overline{xyz})$ |
| REVINT | $\to$ | $x_1\ x_2$ GENOME $x_3\ x_4$ | 1 | $q_{D1}(\overline{x_4})q_{D2}(\overline{x_3})q_{A1}(\overline{x_2})q_{A2}(\overline{x_1})$ |
| STRUCT | $\to$ | $x_1$ STRUCT $x_2$ | $1-\kappa_S$ | $q_S(xy)$ |
| | $\mid$ | LOOP | $\kappa_S$ | |
| LOOP | $\to$ | $x_1$ LOOP | $1-\kappa_L$ | $q_L(x_1)$ |
| | $\mid$ | $\epsilon$ | $\kappa_L$ | |
| CONS | $\to$ | $x_1$ CONS | $1-\kappa_C$ | $q_C(x_1)$ |
| | $\mid$ | $\epsilon$ | $\kappa_C$ | |

**The Joint Finite-Time Grammar**

**How to form the joint grammar** The general rules for forming the joint grammar from the conditional grammar are as follows. For every nonterminal of the following form (here L, R, and/or E are allowed to be $\epsilon$)

| LHS | $\to$ | RHS | Transition | Emission |
|---|---|---|---|---|
| F | $\to$ | L F R | $1-\kappa_F$ | $q_F(LR)$ |
| | $\mid$ | E | $\kappa_F$ | |

...that is, for GENOME, INTER, FWDCDS, REVCDS, STRUCT, LOOP, and CONS (with $F \in \{R, N, G, S, L, C\}$), replace these rules with

| LHS | $\to$ | RHS | Transition | Emission |
|---|---|---|---|---|
| $\mathtt{F}_M$ | $\to$ | $\mathtt{L}_M\ \mathtt{F}_M\ \mathtt{R}_M$ | $(1-\beta_{M,F})(1-\kappa_F)\alpha_F$ | $q_F(LR_x)\exp(Q_F t)(LR_x, LR_y)$ |
| | $\mid$ | $\mathtt{L}_Y\ \mathtt{F}_M\ \mathtt{R}_Y$ | $\beta_{M,F}$ | $q_F(LR_y)$ |
| | $\mid$ | $\mathtt{L}_X\ \mathtt{F}_D\ \mathtt{R}_X$ | $(1-\beta_{M,F})(1-\kappa_F)(1-\alpha_F)$ | $q_F(LR_x)$ |
| | $\mid$ | $\mathtt{E}_M$ | $(1-\beta_{M,F})\kappa_F$ | |
| $\mathtt{F}_D$ | $\to$ | $\mathtt{L}_M\ \mathtt{F}_M\ \mathtt{R}_M$ | $(1-\beta_{D,F})(1-\kappa_F)\alpha_F$ | $q_F(LR_x)\exp(Q_F t)(LR_x, LR_y)$ |
| | $\mid$ | $\mathtt{L}_Y\ \mathtt{F}_M\ \mathtt{R}_Y$ | $\beta_{D,F}$ | $q_F(LR_y)$ |
| | $\mid$ | $\mathtt{L}_X\ \mathtt{F}_D\ \mathtt{R}_X$ | $(1-\beta_{D,F})(1-\kappa_F)(1-\alpha_F)$ | $q_F(LR_x)$ |
| | $\mid$ | $\mathtt{E}_M$ | $(1-\beta_{D,F})\kappa_F$ | |

...that is, two versions $\mathtt{F}_M$ and $\mathtt{F}_D$, with different $\beta$'s for each type.

For every other nonterminal N, there need to be three versions $\mathtt{N}_M$, $\mathtt{N}_X$ and $\mathtt{N}_Y$, with outgoing rules for each type going to other nonterminals of the same type.

##### Top level

| LHS | $\to$ | RHS | Transition | Emission |
|---|---|---|---|---|
| $\mathtt{GENOME}_k$ | $\to$ | $\mathtt{REGION}_M\ \mathtt{GENOME}_M$ | $(1-\beta_{k,R})(1-\kappa_R)\alpha_R$ | |
| | $\mid$ | $\mathtt{REGION}_Y\ \mathtt{GENOME}_M$ | $\beta_{k,R}$ | |
| | $\mid$ | $\mathtt{REGION}_X\ \mathtt{GENOME}_D$ | $(1-\beta_{k,R})(1-\kappa_R)(1-\alpha_R)$ | |
| | $\mid$ | $\epsilon$ | $(1-\beta_{k,R})\kappa_R$ | |
| $\mathtt{REGION}_j$ | $\to$ | $\mathtt{INTER}_j$ | $p_N$ | |
| | $\mid$ | $\mathtt{FWDCDS}_j$ | $p_G/2$ | |
| | $\mid$ | $\mathtt{REVCDS}_j$ | $p_G/2$ | |
| | $\mid$ | $\mathtt{STRUCT}_j$ | $p_S$ | |
| | $\mid$ | $\mathtt{CONS}_j$ | $p_C$ | |
| INTER | $\to$ | $x_1$ INTER | $1-\kappa_N$ | $q_N(x_1)$ |
| | $\mid$ | $\epsilon$ | $\kappa_N$ | |

##### Coding sequences

| LHS | $\to$ | RHS | Transition | Emission |
|---|---|---|---|---|
| FWDCDS | $\to$ | FWDCOD FWDCDS | $1-\kappa_G$ | |
| | $\mid$ | $\epsilon$ | $\kappa_G$ | |
| FWDCOD | $\to$ | $x_1\ x_2\ x_3$ | $1-p_I$ | $q_G(xyz)$ |
| | $\mid$ | $x_1\ x_2\ x_3$ FWDINT | $p_I/3$ | $q_G(xyz)$ |
| | $\mid$ | $x_1\ x_2$ FWDINT $x_3$ | $p_I/3$ | $q_G(xyz)$ |
| | $\mid$ | $x_1$ FWDINT $x_2\ x_3$ | $p_I/3$ | $q_G(xyz)$ |
| FWDINT | $\to$ | $x_1\ x_2$ GENOME $x_3\ x_4$ | 1 | $q_{D1}(x_1)q_{D2}(x_2)q_{A1}(x_3)q_{A2}(x_4)$ |

| LHS | $\to$ | RHS | Transition | Emission |
|---|---|---|---|---|
| REVCDS | $\to$ | REVCDS REVCOD | $1-\kappa_G$ | |
| | $\mid$ | $\epsilon$ | $\kappa_G$ | |
| REVCOD | $\to$ | $x_1\ x_2\ x_3$ | $1-p_I$ | $q_G(\overline{xyz})$ |
| | $\mid$ | $x_1\ x_2\ x_3$ REVINT | $p_I/3$ | $q_G(\overline{xyz})$ |
| | $\mid$ | $x_1\ x_2$ REVINT $x_3$ | $p_I/3$ | $q_G(\overline{xyz})$ |
| | $\mid$ | $x_1$ REVINT $x_2\ x_3$ | $p_I/3$ | $q_G(\overline{xyz})$ |
| REVINT | $\to$ | $x_1\ x_2$ GENOME $x_3\ x_4$ | 1 | $q_{D1}(\overline{x_4})q_{D2}(\overline{x_3})q_{A1}(\overline{x_2})q_{A2}(\overline{x_1})$ |

**RNA structures**

| LHS | → | RHS | Transition | Emission |
|---|---|---|---|---|
| `STRUCT` | → | $x_1$ `STRUCT` $x_2$ | $1 - \kappa_S$ | $q_S(xy)$ |
| | \| | `LOOP` | $\kappa_S$ | |
| `LOOP` | → | $x_1$ `LOOP` | $1 - \kappa_L$ | $q_L(x_1)$ |
| | \| | $\epsilon$ | $\kappa_L$ | |
| `CONS` | → | $x_1$ `CONS` | $1 - \kappa_C$ | $q_C(x_1)$ |
| | \| | $\epsilon$ | $\kappa_C$ | |

### D TKF-DP: Dirichlet-process Potts coupling

The TKF-DP model decorates TKF92 with three independent Dirichlet processes that govern (i) the per-site key partition into Potts coevolutionary cliques; (ii) the per-site assignment to a substitution class; and (iii) the per-class-pair assignment to a Potts coupling atom. The class-level substitution dynamics is a multi-site reversible CTMC; the alignment likelihood remains plain TKF92. Inference proceeds by a variational E-step on the continuous-time class clique, Gibbs sweeps and Jain–Neal split–merge proposals on the three Dirichlet partitions, recursive traceback over a time-indexed gravestone-augmented pair SCFG for the augmented branch history, and a stochastic M-step on the corpus-level hyperparameters with closed-form Dirichlet–multinomial / Gamma–Poisson / Laplace conjugacies unlocking the new-class / new-atom base-measure integrals.

#### D.1 The TKF-DP generative model

The TKF92 model (51) of insertion, deletion, and substitution has the convenient property that the alignment path and the substitution history factorize: the marginal distribution over alignments is independent of the amino acid dynamics, and inference proceeds via a pair HMM on the alignment with substitution likelihoods plugged in pointwise. This factorization is exactly what is lost when site–site coevolutionary couplings are introduced naively, since the substitution process at any site then depends on the joint state of its coupled partners, and any coupling rule that consults the existing alignment to decide partners destroys the marginal pair HMM.

We describe a minimal extension of TKF92 that re-introduces site–site Potts couplings while preserving the alignment–substitution factorization. Each newborn site is assigned a discrete key drawn i.i.d. from a Dirichlet process, and sites that share a key co-evolve under a Potts-modulated joint CTMC. Because the keys are drawn independently of the alignment, the alignment marginal is unchanged.

The alignment–substitution factorization survives at the parameter level but, once couplings are present, breaks at the latent-variable level: the substitution likelihood conditional on the alignment depends on the full timed indel history, including transient (alignment-invisible) lineages whose presence transiently modifies the rates of co-class members. Inference therefore requires a tractable representation of timed indel histories. We obtain one via a gravestone-augmented latent state and a time-indexed pair SCFG whose inside–outside is closed form (§D.5).

**Definition D.1** (TKF-DP)**.** *Fix TKF92 indel parameters* $(\lambda, \mu, r)$*, a key concentration parameter* $\alpha_z > 0$*, and a symmetric* $20 \times 20$ *Potts coupling matrix* $H$*. The process jointly generates a TKF92 alignment path, a key assignment for every site, and an amino acid trajectory at every site, as follows.*

*1. Sample stick-breaking weights once, globally:*

$$\pi_k = \beta_k \prod_{l<k}(1-\beta_l), \qquad \beta_k \overset{iid}{\sim} \mathrm{Beta}(1, \alpha_z).$$

*2. Run TKF92 indel dynamics with parameters $(\lambda, \mu, r)$, giving a stochastic set of alive sites $S(t)$ over time.*

*3. At the birth of each new site $s$, draw a key*

$$z_s \mid \pi \sim \mathrm{Cat}(\pi),$$

*independently of all other sites, of the alignment path, and of the substitution history. The key is fixed for the site's lifetime.*

*4. At any time $t$, the alive sites $S(t)$ are partitioned into equivalence classes $\{C_k(t)\}$ by the relation $z_s = z_{s'}$. The amino acid states $\{x_s\}_{s\in S(t)}$ evolve as independent CTMCs across classes; the CTMC on class $C$ is a $|C|$-site reversible chain with stationary distribution*

$$\pi_C(x_C) \propto \exp\Big(-\sum_{\{i,j\}\subset C} H(x_i, x_j)\Big).$$

*Class composition is piecewise constant between birth/death events; on each event, the affected class is rebuilt and the surviving sites re-equilibrate under the new class generator.*

*Remark* D.1 (Factorization). Conditional on $\pi$, the key draws are i.i.d. across sites, so the joint distribution factorizes as

$$P(\text{alignment}, \pi, \{z_s\}, \text{substitution}) = P(\text{alignment})\cdot P(\pi)\cdot \prod_s P(z_s \mid \pi)\cdot P(\text{substitution} \mid \text{alignment}, \{z_s\}).$$

In particular, the marginal alignment likelihood is exactly that of TKF92, and the pair HMM one would use under $\alpha_z = \infty$ (no coupling) is unchanged.

*Remark* D.2 (Partition statistics). Marginalizing $\pi$, the partition induced on any fixed finite set of sites is the Ewens partition with parameter $\alpha_z$, equivalent to a Chinese restaurant process draw (2). The expected number of co-evolving classes among $L$ sites is $\mathcal{O}(\alpha_z \log L)$, and tuning $\alpha_z$ controls the sparsity of couplings: $\alpha_z \gg L$ yields $\mathcal{O}(L^2/\alpha_z)$ co-evolving pairs with higher-order classes suppressed by additional factors of $L/\alpha_z$. Smaller $\alpha_z$ gives Ewens-distributed class sizes with heavier tails, including occasional larger cliques.

#### D.2 IBP variant

To allow each site to participate in multiple independent couplings, replace the Dirichlet process with an Indian Buffet Process (17).

**Definition D.2** (TKF92-IBP)**.** *Sample feature frequencies $\{\pi_k\}_{k\geq 1}$ from an IBP with concentration $\alpha_z$. At the birth of each site $s$, draw a binary feature vector $b_s \in \{0,1\}^{\mathbb{N}}$ with*

$$b_{s,k} \mid \pi_k \overset{ind}{\sim} \mathrm{Bern}(\pi_k),$$

*independently across sites. Two alive sites $s, s'$ are coupled iff there exists $k$ with $b_{s,k} = b_{s',k} = 1$. The amino acid dynamics is the reversible CTMC on the alive sites with stationary distribution*

$$\pi(x_S) \propto \exp\Big(-\sum_{\{s,s'\}\,:\,\exists k,\, b_{s,k}=b_{s',k}=1} H(x_s, x_{s'})\Big).$$

The same factorization observation applies: feature draws are i.i.d. given $\{\pi_k\}$, so the marginal alignment likelihood remains plain TKF92. The IBP variant naturally accommodates hub structure (residues that interact with many partners) via heavy-tailed feature frequencies.

#### D.3 Site classes and a GTR-parameterized generator

Real proteins exhibit substantial heterogeneity in equilibrium amino acid usage and in coupling profile: positions in $\alpha$-helices, $\beta$-sheets, buried cores, and disordered regions have distinct frequencies and distinct interaction patterns with their structural neighbours. We extend the model with a per-site latent class variable that determines both the uncoupled (point) substitution model and the per-pair Potts interaction tensor. The variable is non-evolving, drawn once at site birth, and we put a second Dirichlet process on it — independent of the key DP — so that the number of site classes is itself learned from data, in line with the original infinite-mixture CAT model of Lartillot and Philippe (31).

**Definition D.3** (Site-class TKF-DP)**.** *Fix the alphabet $\mathcal{A}$ with $|\mathcal{A}| = A = 20$, a symmetric exchangeability matrix $S \in \mathbb{R}^{A\times A}_{\geq 0}$ ($S_{xy} = S_{yx}$), per-site rate-multiplier hyperparameters $(a_\eta, b_\eta)$, a site-class concentration $\alpha_c > 0$, a Potts-atom concentration $\alpha_H > 0$, and base measures*

$$G_0: \quad \pi^{(c)} \overset{iid}{\sim} \mathrm{Dirichlet}(\kappa_\pi\,\bar{\pi}), \qquad G_0^H: \quad H_h(i,j) \overset{ind}{\sim} \mathcal{N}\big(\mu_{kl},\, \tau_{kl}^{-1}\big) \text{ for } i \leq j,\ H_h(j,i) := H_h(i,j),$$

*where $(k,l) = (\min(i,j), \max(i,j))$ index the $A(A+1)/2$ unordered amino-acid pairs, and each Potts atom $H_h \in \mathbb{R}^{A\times A}$ is symmetric in $(i,j)$ by construction. Site classes are drawn from a Dirichlet process $D_c \sim \mathrm{DP}(\alpha_c, G_0)$ via stick-breaking; Potts atoms are drawn from a separate Dirichlet process $D_H \sim \mathrm{DP}(\alpha_H, G_0^H)$. At the birth of each site $s$, draw an independent per-site rate multiplier and a site class:*

$$\eta_s \overset{iid}{\sim} \mathrm{Gamma}(a_\eta, b_\eta), \qquad c_s \mid D_c \overset{iid}{\sim} D_c,$$

*in addition to the DP key $z_s$ (§D.1); $\eta_s$, $c_s$, and $z_s$ are mutually independent and independent of the alignment path, and are fixed for the site's lifetime. The Potts coupling tensor is decomposed by class-pair: for each unordered pair of classes $\{c, c'\}$ that ever co-occur within a key DP class, draw a Potts-atom assignment*

$$h_{cc'} \mid D_H \overset{iid}{\sim} D_H,$$

*at first co-occurrence and fixed thereafter; the materialized slice for the pair is $H_{cc'}(i,j) := H_{h_{cc'}}(i,j)$. The amino-acid trajectory on a DP key class $C$ is the single-site reversible CTMC with stationary distribution*

$$\pi_C(x_C) \;\propto\; \prod_{s\in C} \pi^{(c_s)}(x_s)\cdot \exp\Big(-\sum_{\{s,s'\}\subset C} H_{c_s c_{s'}}(x_s, x_{s'})\Big),$$

*and (taking the F81 instance reversible with respect to this stationary)*

$$Q^s\big(x \to x'; x_{C\setminus s}\big) \;=\; \eta_s\, S_{xx'}\, \pi^{(c_s)}(x')\, \exp\Big(-\tfrac{1}{2}\Delta H_s\Big), \qquad \Delta H_s = \sum_{s'\in C\setminus s} [H_{c_s c_{s'}}(x', x_{s'}) - H_{c_s c_{s'}}(x, x_{s'})].$$

*Remark* D.3 (GTR limit and singletons). For a singleton key DP class, $\Delta H_s \equiv 0$ and the rate reduces to $Q^s_{xy} = \eta_s S_{xy} \pi^{(c_s)}(y)$, the F81 instance of GTR with class-specific equilibrium and a per-site rate multiplier. Marginalized over $c_s$ this is exactly the CAT infinite-mixture profile model of Lartillot and Philippe (31), augmented with Yang's gamma per-site rate variation (54). Coupling enters only via $\Delta H_s$ when the key DP class size exceeds one, so the model strictly extends the CAT–gamma combination by adding a coevolutionary tail.

*Remark* D.4 (F81 vs. symmetric-Metropolis form). The F81 form $Q_{xy} \propto S_{xy}\pi(y)$ chosen above is one of two natural reversible instances of the GTR family with stationary $\pi$ and exchangeability $S$; the other is the symmetric-Metropolis instance $Q_{xy} \propto S_{xy}\sqrt{\pi(y)/\pi(x)}$. Both satisfy detailed balance with respect to the joint Potts stationary above. The symmetric-Metropolis form has the property that the eigendecomposition of its symmetrized similarity transform is independent of $\pi$, allowing one corpus-wide $A^3$ decomposition to be reused across all classes. We adopt the F81 form here because, combined with the secret-destination augmentation of §D.6, it yields strict Dirichlet–multinomial conjugacy on $\pi^{(c)}$, which the symmetric-Metropolis form does not. The eigendecomposition then becomes per-class (one $A^3$ per class per outer step), which is small at $A = 20$ and is dominated by the Potts cluster work in any case.

*Remark* D.5 (Three independent Dirichlet processes). The construction has *three* Dirichlet processes acting on the substitution side: the key DP with concentration $\alpha_z$ partitions sites into co-evolving cliques (§D.1); the site-class DP with concentration $\alpha_c$ assigns each site a profile $\pi^{(c)}$; and a Potts-atom DP with concentration $\alpha_H$ and base measure $G_0^H$ assigns each unordered class-pair $\{c, c'\}$ to a coupling atom $H_{h_{cc'}}$. With $\alpha_H$ small the $K(K+1)/2$ class-pair indices collapse onto a small number of canonical Potts atoms shared across class-pairs; with $\alpha_H \to \infty$ each class-pair has its own atom. The Potts-atom DP is what makes the model identifiable for $K_c \gg 1$: without it, the number of free Potts parameters scales as $K_c^2 \cdot A^2$; with it the count scales as $K_H \cdot A^2 + K_c^2$ class-pair indices, where $K_H$ is data-driven and typically much smaller than $K_c^2$.

*Remark* D.6 (Per-class-pair side potentials). A useful refinement of the per-class profile $\pi^{(c)}$ is to give each unordered class-pair $\{c, c'\}$ its own pair of *side potentials* $h_a^{(c,c')}, h_b^{(c,c')} \in \mathbb{R}^A$, with independent Gaussian priors centered at zero,

$$h_a^{(c,c')}(x), h_b^{(c,c')}(y) \;\sim\; \mathcal{N}(0, \tau_h^{-1}). \tag{D.1}$$

The per-pair joint stationary then becomes

$$\log \pi_{\text{joint}}^{(c,c')}(x, y) \;=\; \log \pi^{(c)}(x) + \log \pi^{(c')}(y) - h_a^{(c,c')}(x) - h_b^{(c,c')}(y) - H_{cc'}(x, y) - \log Z_{cc'}, \tag{D.2}$$

i.e. $h$ is the per-pair deviation of the effective per-site stationary $\pi_{cc'}^{\text{eff},a}(x) \propto \pi^{(c)}(x)e^{-h_a^{(c,c')}(x)}$ from the per-class background $\pi^{(c)}$. For self-pairs $(c, c)$ the two sites are exchangeable, so the joint distribution must be symmetric under swap $(x, y) \leftrightarrow (y, x)$. Combined with $H_{cc}$ symmetric in its arguments, this forces $h_a^{(c,c)} = h_b^{(c,c)}$; we parameterise self-pairs with a single shared vector $h^{(c,c)} \in \mathbb{R}^A$ (and prior $h^{(c,c)} \sim \mathcal{N}(0, \tau_h^{-1})$, contributing $A$ free parameters per self-pair rather than $2A$). Off-diagonal pairs $\{c, c'\}$ with $c \neq c'$ retain two independent vectors $h_a^{(c,c')}, h_b^{(c,c')}$ ($2A$ free parameters each). Reversibility of the F81-form joint generator is preserved by replacing the destination factor $\pi^{(c)}(x')$ in the site-1 flip rate with $\pi_{cc'}^{\text{eff},a}(x')$ (and analogously for site 2).

The motivation is that $\pi^{(c)}$ captures the population-mean amino-acid usage of class $c$ *averaged over all class-pair contexts*, while the actual usage of class $c$ when paired with class $c'$ may deviate (e.g. a Cys-bearing class paired with itself is more Cys-rich than the per-class average, because we are conditioning on disulfide-forming sites). The $h$ vectors absorb that deviation, leaving the Potts atom $H_{cc'}$ to capture the coupling *above the marginal compositional shift*. The Gaussian prior pulls $h$ to zero so that any pair-specific deviation must be data-supported.

*Remark* D.7 (Hierarchical structure of the $H$ base measure). The amino-acid-pair-indexed Gaussian base measure $\mathcal{N}(\mu_{kl}, \tau_{kl}^{-1})$ embeds an empirical-Bayes hierarchy: the corpus-level $(\mu_{kl}, \tau_{kl})$ encode the population-mean coupling propensity for each amino-acid pair (typically attractive for hydrophobic–hydrophobic, charge-complementary, etc., and repulsive for charge-like-like), and atom-specific deviations $H_h(i,j) - \mu_{kl}$ are learned around that mean with atom-specific magnitude controlled by the data. Symmetry of each $H_h$ in $(i,j)$ encodes the unordered-edge convention of Potts couplings; this is standard practice in the DCA literature (13) and is biologically natural in the absence of any a priori labelling distinguishing the two ends of an inter-class edge. The Gaussian prior on each $H_h$ is conjugate to the Gaussian (Hessian-based) Laplace approximation of the path-DCA likelihood, giving a closed-form Gaussian posterior on each materialized atom under the local quadratic expansion of §D.6.

#### D.4 Class-level variational substitution likelihood

Inference under the model requires, for each branch of a phylogeny and each co-evolving DP class $C$, the per-branch transition probability $P(\mathbf{x}_C(t) \mid \mathbf{x}_C(0), \{c_s\}_{s\in C})$. Direct evaluation requires exponentiating a generator on $\mathcal{A}^{|C|}$ and is intractable beyond $|C| \approx 5$. The bulk of the key-DP partition is small-class for $\alpha_z$ in the practical regime, so exact computation handles most of the work; what remains is to control the tail of large classes.

We derive a variational lower bound from an exact path-measure factorization on the class clique. The factorization is the analog, on path space, of the static pseudolikelihood approximation used in DCA inference (13), but with the static partition function entirely absorbed into the conditioned-on initial state. Two variational families on the resulting Girsanov ELBO are natural. The first is the simpler scheme we develop here, parameterized by per-site distributions at internal tree nodes plus a single constant rate matrix per site per branch, with a closed-form ELBO via eigendecomposition and a Felsenstein-style pruning recursion for optimization. The second is the fully inhomogeneous mean-field scheme of Cohn et al. (8), with time-varying rates and forward-backward ODE sweeps, which we describe as an upgrade for branches where the constant-rate scheme is too loose.

**Proposition D.1** (Path-measure factorization)**.** *Let $\sigma_C \in D([0,t];\mathcal{A}^{|C|})$ be the trajectory of a single-site reversible CTMC on class $C$ with rate $Q^s(x_s \to x'_s; x_{C\setminus s})$ and stationary distribution $\pi_C$. The path measure satisfies*

$$\mathbb{P}(\sigma_C) \;=\; \pi_C\big(\sigma_C(0)\big) \prod_{s\in C} \mathbb{P}_s\big(\sigma_s \,\big|\, \sigma_{C\setminus s}\big),$$

*where $\mathbb{P}_s(\cdot \mid \sigma_{C\setminus s})$ is the law of an inhomogeneous CTMC on site $s$ with rate $Q^s\big(\cdot\,;\, x_{C\setminus s}(u)\big)$ at time $u$.*

The factorization is exact: simultaneous jumps at distinct sites have zero Lebesgue measure, every jump is therefore attributable to a unique site, and the path log-density splits site-wise. The only intractable factor is $\pi_C(\sigma_C(0))$, which is conditioned upon for transition probabilities and so cancels.

##### Constant-rate variational family

*Remark* D.8 (Variational family). The variational family is parameterized at two levels. At each internal tree node $v$ and site $s$, a discrete distribution $\nu_v^s \in \Delta^{|\mathcal{A}|-1}$ over residue states; node distributions factorize over sites, $\nu_v(\mathbf{x}_v) = \prod_s \nu_v^s(x_v^s)$. At each branch $b$ between nodes $p$ and $c$ of length $t_b$ and each site $s \in C$, a constant rate matrix $\hat{Q}_b^s$ on $[0, t_b]$. Conditional on sampled endpoint states $(x_p^s, x_c^s) \sim \nu_p^s \otimes \nu_c^s$, the variational law on the branch is the CTMC bridge of $\hat{Q}_b^s$ from $x_p^s$ to $x_c^s$, with sites independent given endpoints. The joint variational law on the tree is

$$\mathbb{Q} = \prod_v \nu_v \cdot \prod_b \prod_{s\in C} \mathbb{Q}_b^s\big(\sigma_b^s \,\big|\, x_p^s, x_c^s\big).$$

**Proposition D.2** (Closed-form per-branch ELBO)**.** *The path-measure KL contribution from branch $b$ and site $s$ has Girsanov form*

$$\mathrm{KL}_{b,s} = \mathbb{E}_{\mathbb{Q}}\int_0^{t_b}\sum_{x,x'} q_b^s(x;u)\Big[\mathbb{E}_u\big[Q^s(x\to x';x_{C\setminus s}(u))\big] - \hat{Q}_b^s(x\to x') + \hat{Q}_b^s(x\to x')\log\frac{\hat{Q}_b^s(x\to x')}{\exp\mathbb{E}_u[\log Q^s(\cdot)]}\Big]\,du,$$

*where $q_b^s(x;u) = \mathbb{E}_{x_p,x_c}[q_b^s(x;u \mid x_p,x_c)]$ is the variational bridge marginal of site $s$ at time $u$ averaged over endpoint distributions, and the inner expectations $\mathbb{E}_u[\cdot]$ are over the product of* instantaneous *neighbour bridge marginals at $u$, $\prod_{s'\neq s} q_b^{s'}(\cdot;u)$. The integrand is therefore a product of two time-varying bridge marginals at the same instant $u$, and the integral over $[0,t_b]$ does* not *factorise into time-averages: the cross-temporal correlation between $q_b^s(\cdot;u)$ and $q_b^{s'}(\cdot;u)$ contributes to $\mathrm{KL}_{b,s}$ and is what makes the formula a strict lower bound. The bridge-expectation closed form below evaluates the integral correctly.*

*In the constant-rate family, the stationary point of $\mathrm{KL}_{b,s}$ in $\hat{Q}_b^s$ admits no closed form in general because of the cross-temporal correlation. The geometric-mean rate*

$$\log\hat{Q}_b^{s,*}(x\to x') \;=\; \mathbb{E}_{\bar{q}_b^{C\setminus s}}\big[\log Q^s(x\to x';x_{C\setminus s})\big],$$

*where $\bar{q}_b^{s'}(x) = (1/t_b)\int_0^{t_b} q_b^{s'}(x;u)du$ is the time-averaged neighbour distribution for each $s'\neq s$, is the stationary point under the additional "mean-field-of-bridges" assumption that $q_b^s(\cdot;u)$ and $q_b^{s'}(\cdot;u)$ are approximately uncorrelated in $u$ on $[0,t_b]$. This assumption is exact at $H=0$ (sites independent under the variational law as well as under the truth), close to exact for small coupling, and only mildly violated at typical phylogenetic time scales; we take $\hat{Q}_b^{s,*}$ above as the chosen variational rate and evaluate the strict $\mathrm{KL}_{b,s}$ at this rate via the bridge-expectation formula in the Remark below. Each term of the resulting bound is $\geq 0$ by Jensen, so $\mathrm{KL}_{b,s}\geq 0$ and the ELBO is a strict lower bound on $\log P$, as required.*

*Remark* D.9 (Closed-form ELBO via pairwise bridge expectations). Each rate $\hat{Q}_b^s$ admits an eigendecomposition $\hat{Q}_b^s = U^s D^s (V^s)$ with $V^s = (U^s)^{-1}$ and eigenvalues $\{\lambda_k^s\}$ (we drop the branch subscript $b$ for legibility). The forward and backward propagators have the same expansion:

$$[\exp(\hat{Q}^s u)]_{x_p,x} = \sum_k U^s_{x_p,k}\,V^s_{k,x}\,e^{\lambda_k^s u}, \quad [\exp(\hat{Q}^s(t-u))]_{x,x_c} = \sum_l U^s_{x,l}\,V^s_{l,x_c}\,e^{\lambda_l^s(t-u)},$$

so the per-site bridge marginal is

$$q_b^s(x;u\mid x_p,x_c) = \frac{1}{P_s}\sum_{k,l} U^s_{x_p,k}\,V^s_{k,x}\,U^s_{x,l}\,V^s_{l,x_c}\,e^{\lambda_k^s u}\,e^{\lambda_l^s(t-u)},$$

with $P_s = [\exp(\hat{Q}^s t)]_{x_p,x_c}$ the bridge normalising constant. The branch integrand of $\mathrm{KL}_{b,s}$ becomes a sum over $(k,l)$ and over $(k',l')$ from the neighbour expansion, with pair-time integrals

$$\mathrm{HR}(\alpha,\beta,t) \;=\; \begin{cases}(e^{\alpha t}-e^{\beta t})/(\alpha-\beta) & \alpha\neq\beta,\\ te^{\alpha t} & \alpha=\beta,\end{cases}$$

which is the standard bridge-expectation (25) expression. The full integral over $[0,t_b]$ is a finite sum of HR evaluations and closed-form coefficients in $U,V,\lambda$.

**Optimization: Felsenstein-like pruning over node distributions**

The node distributions $\nu_v^s$ optimise the ELBO under a Felsenstein-style upwards–downwards pass: each $\nu_v^s$ is the geometric mean of (i) the upwards-propagated likelihood from $\nu_v^s$'s subtree, (ii) the downwards-propagated prior from the rest of the tree, and (iii) the per-branch forward and backward propagators of $\hat{Q}^s$ on each adjacent branch. The pass is $O(\text{edges} \cdot A^2)$ per outer ELBO step, dominated by the per-branch matrix exponential. The rate $\hat{Q}_b^{s,*}$ is then re-computed under the fresh node distributions and the loop iterates to convergence; the ELBO is monotone increasing in each step.

**Jensen gap and the Cohn upgrade as endpoints of an adaptive refinement**

The constant-rate scheme of the previous subsection and the inhomogeneous mean-field scheme of Cohn et al. (8) are not separate algorithms but the $N = 1$ and $N \to \infty$ ends of a single adaptive refinement. Subdivide a branch of length $t_b$ into $N$ sub-intervals at times $0 = u_0 < u_1 < \cdots < u_N = t_b$ with constant rate $\hat{Q}_{b,k}^s$ on each $[u_{k-1}, u_k]$, and a variational node distribution $\nu_k^s$ at each intermediate midpoint. The resulting variational law is a piecewise-constant CTMC bridge: a piecewise-constant approximation to whatever time-varying rate Cohn's scheme would use at the same fidelity. As $N \to \infty$ with mesh refining, piecewise-constant rates become dense in continuous time-varying rates, so the variational family becomes Cohn's and the ELBO converges monotonically up. At the optimum, the $\nu_k^s$ are pinned by stationarity to the bridge marginals of the surrounding piecewise-constant CTMC — the explicit-midpoint parameterisation is slack at the optimum.

The Jensen-gap diagnostic per segment is the integrated AM–GM discrepancy of $Q^s$ against neighbour bridge marginals at the segment's resolution; it is closed form via the same eigendecomposition machinery and tells the algorithm where to refine. For typical phylogenetic branches ($t_b \in [0.1, 1]$) most segments need $N = 1$; high-coupling segments may benefit from $N = 2$ or 3; only pathological branches need $N \to \infty$. Forced segmentation by composition-change events from the augmented indel history (§D.5) composes cleanly with this Jensen-driven refinement: subdivide first at every composition-change event, then optionally $N$-refine within each composition-constant segment.

*Remark* D.10 (Cohn's inhomogeneous fixed point as the $N \to \infty$ limit). For a composition-constant segment requiring $N \to \infty$ refinement, the limit is Cohn et al.'s full inhomogeneous fixed point: a time-varying rate $\tilde{Q}^s(\cdot; u)$ obtained by alternating forward sweeps $\dot{q}_s = q_s \tilde{Q}^s$ from $\delta_{x_p^s}$, backward sweeps $\dot{\rho}_s = -\tilde{Q}^s \rho_s$ from $\delta_{x_c^s}$, and rate updates $\log \tilde{Q}^{s,*}(\cdot; u) = \mathbb{E}[\log Q^s(\cdot)]$ pointwise in $u$, with bridge marginals $q_s^{\text{bridge}}(x; u) \propto q_s(x; u)\, \rho_s(x; u)$. The midpoint construction above is the discretisation of this fixed point at mesh size $u_k - u_{k-1}$; the Cohn limit is recovered as the mesh shrinks to zero. Cohn et al. (8, §7) also give the internal-node rule used by Felsenstein pruning under the time-varying rate.

This is the lower-bound construction referenced in the Discussion for Potts components of size > 2: independent-coupling pairwise factorisation gives the $N$=1 lower bound, midpoint augmentation extends it to a formal ELBO, and $N \to \infty$ recovers the Cohn fixed point. Classes whose Jensen gap remains large after refinement are candidates for the $k$-clique cluster-variational upgrade of Linzner & Koeppl (34), or for fallback to exact uniformization-based bridge sampling via Rao & Teh (41).

#### D.5 Augmented indel histories via a time-indexed pair SCFG

The class-level ELBO of §D.4 was derived under fixed class composition over a constant-composition segment of a branch. To compose with the tree we need the timed indel history on each branch, including *transient* insertions: under coupling, an insertion that is born and then deleted on the same branch is not visible in the alignment but does change the rate of every other site in its DP class during its lifetime. The naive scheme of sampling alignment-visible indel timings from their TKF92 prior is inadequate for two distinct reasons: first, transient lineages are missed; second, even for visible events, the conditional density given the alignment depends on substitution evidence and is not what the alignment marginal alone provides.

We resolve both issues by augmenting the latent state with *gravestones.* The augmentation does not change the model. It changes the latent representation: every fragment that ever existed during the branch — alive at start, alive at end, deleted on the branch, or transiently inserted then deleted — is represented as a labeled position in the augmented branch history. With gravestones the alive count $N_a(s)$ is bounded by the visible count $N_v(s)$, so the indel CTMC bridge has a finite state space and admits a closed-form recursive sampler. The gravestone counts at internal nodes of the tree become latent variables resampled jointly with the rest of the augmented history.

##### The branching process and its grammar

We first present the TKF91 case (single residue per fragment) where the branching structure is cleanest, then read off the TKF92 expansion. Let $L(t)$ denote a single link observed $t$ time units after its birth. Under TKF91 dynamics, $L(t)$ has lifetime $\tau \sim \mathrm{Exp}(\mu)$, and during its lifetime spawns child links at rate $\lambda$. Each child is itself an $L$ of the appropriate residual age. The branching process is a continuous-time Galton–Watson process expressible as a stochastic context-free grammar with productions indexed by continuous time:

$$
\begin{aligned}
L(t) &\to A \cdot D(t,\, t) && \text{at weight } e^{-\mu t},\\
L(t) &\to G_\tau \cdot D(t,\, \tau) && \text{at density } \mu e^{-\mu\tau}\, d\tau,\;\; \tau \in (0,t),\\
D(t,\, T) &\to L(t-u) \cdot D(t,\, T) && \text{at density } \lambda\, du,\;\; u \in (0,T),\\
D(t,\, T) &\to \varepsilon && \text{at weight } e^{-\lambda T}.
\end{aligned}
$$

The terminals $A$ and $G_\tau$ stand for an alive residue and a gravestone residue that died at relative time $\tau$, respectively, each carrying a DP key, a site class, and an amino-acid trajectory over its lifetime as governed by §D.4. The non-terminal $D(t, T)$ generates a list of child links born during a window of size $T$ inside an observation interval of size $t$. The reading is: a link either survives to observation (probability $e^{-\mu t}$) and can have spawned children at any time in $(0, t)$, or it died at some $\tau \in (0, t)$ (density $\mu e^{-\mu\tau}$) and can only have spawned children during $(0, \tau)$. The empty-list weight $e^{-\lambda T}$ is the probability of zero events from a Poisson process of rate $\lambda$ over a window of $T$.

**TKF92 expansion.** The generalization to TKF92 replaces each single-residue terminal with a fragment of geometrically-distributed length, leaving the fragment-level branching structure entirely unchanged:

$$
\begin{aligned}
A &\to \mathcal{R}^A \cdot A^*, & A^* &\to \mathcal{R}^A \cdot A^* \text{ at weight } r, & A^* &\to \varepsilon \text{ at weight } 1-r,\\
G_\tau &\to \mathcal{R}^G_\tau \cdot G^*_\tau, & G^*_\tau &\to \mathcal{R}^G_\tau \cdot G^*_\tau \text{ at weight } r, & G^*_\tau &\to \varepsilon \text{ at weight } 1-r,
\end{aligned}
$$

where $\mathcal{R}^A$ is a single alive residue and $\mathcal{R}^G_\tau$ a single gravestone residue with shared death time $\tau$. Fragment death is collective: all residues of a fragment share its lifetime $[0, \tau]$ and die together at $\tau$. Key-DP labels $z_s$ and class-DP labels $c_s$ are drawn independently per residue at fragment birth from their respective Chinese restaurant predictives, even within a single fragment, so a fragment can carry residues of several key classes and several site classes. The expansion is invisible to the indel skeleton — the bivariate Riccati of the next subsection lives at the fragment level — but enters the substitution likelihood and the DP-class composition through the per-residue payload.

Pairing this with an ancestor sequence yields the pair version: each ancestor fragment present at branch start is an independent root of the recursion above, with $t = t_b$ the branch length; insertions appearing in the children list have no ancestor counterpart and carry only a descendant payload. The pair grammar generates an aligned pair of fragment sequences (with alive/gravestone labels on the descendant side) jointly with the parentage tree.

##### Inside generating function and the bivariate Riccati

Let $f_t(z,w) = \mathbb{E}[z^{N_a(t)} w^{N_g(t)}]$ where $N_a, N_g$ count alive and gravestone descendants of one root fragment of age $t$. The inside generating function of the grammar satisfies the Volterra fixed point

$$f_t(z,w) \;=\; z\,e^{-\mu t}\,\exp\!\Big(\lambda\int_0^t (f_s(z,w)-1)\,ds\Big) \;+\; w\int_0^t \mu e^{-\mu\tau}\,\exp\!\Big(\lambda\int_{t-\tau}^t (f_s(z,w)-1)\,ds\Big)\,d\tau,$$

which is equivalent to the forward equation

$$\partial_t f_t(z,w) \;=\; \big[\lambda z^2 - (\lambda+\mu)z + \mu w\big]\,\partial_z f_t(z,w), \qquad f_0(z,w) = z.$$

The right-hand side is quadratic in $z$, so the characteristic equation $\dot z = -[\lambda z^2 - (\lambda+\mu)z + \mu w]$ is Riccati. Define

$$\Delta(w) \;=\; \sqrt{(\lambda-\mu)^2 + 4\lambda\mu(1-w)}, \qquad z_\pm(w) \;=\; \frac{(\lambda+\mu)\pm\Delta(w)}{2\lambda}.$$

The Riccati flow $(z(t)-z_+)/(z(t)-z_-) = (z(0)-z_+)/(z(0)-z_-)\cdot e^{-\Delta(w)t}$ inverts to give $f_t(z^*,w)$ in closed form, hence closed-form pointwise transition probabilities $P\big((N_a,N_g)(t)\,\big|\,(N_a,N_g)(0)\big)$ by coefficient extraction. Setting $w=1$ recovers the standard linear birth-death PGF for $N_a$ alone.

##### Recursive traceback sampler

Given an augmented history with endpoint counts $(a_0,g_0) = (N_a(0),N_g(0))$ and $(a^*,g^*) = (N_a(t_b),N_g(t_b))$, the within-branch indel history is sampled by recursive midpoint traceback on the bivariate process $(N_a,N_g)$. At each level of the recursion, sample a midpoint state from

$$P\big((a_m,g_m)\,\big|\,(a_0,g_0),(a^*,g^*)\big) \;\propto\; P\big((a_0,g_0)\to(a_m,g_m);\,t_b/2\big)\cdot P\big((a_m,g_m)\to(a^*,g^*);\,t_b/2\big),$$

both factors closed form via the Riccati. The midpoint state space is finite since $a_m, g_m \le g^* + a^*$. Recurse on each half. The base case is reached when the residual subproblem has at most a small number of events (births and deaths), at which point the closed-form list of valid event interleavings is enumerable directly and event times are sampled from the corresponding hypoexponential simplex density. Total cost is $O((b+d)\log(b+d))$ per branch where $b,d$ are the branch totals of births and deaths.

After event times are placed, parentage is sampled forward in time: at each birth, a uniform alive fragment is chosen as the parent; at each death, a uniform alive fragment is killed. Parentage uniformity is the standard exchangeability of the linear Galton–Watson tree.

##### Inserted fragment and residue distributions

When a child fragment is added at time $u$ to a DP class $C$ that already contains alive fragments, the inserted residues are drawn from the conditional Boltzmann

$$P\big(x_s(u^+)\,\big|\,x_{C\setminus s}(u^-)\big)\ \propto\ \pi^{(c_s)}\big(x_s(u^+)\big)\ \exp\Big(-\sum_{s'\in C\setminus s} H_{c_s c_{s'}}\big(x_s(u^+),\, x_{s'}(u^-)\big)\Big),$$

which is exactly normalizable on the alphabet $\mathcal{A}$ even when the joint stationary on $C$ is intractable. This is the proper coupled generalization of the $\mathrm{Cat}(\pi)$ inserted-residue rule of standard TKF92.

##### Composition with substitution and with the rest of the tree

The traceback gives a piecewise-constant DP-class composition on each branch, with segment boundaries at every (alive or gravestone) indel event. The variational ELBO of §D.4 applies to each segment as written, with the segment's class composition and duration. Bridge expectations accumulate over segments within each branch and across branches.

Internal-node augmented states $(N_a, N_g)$-valued are sampled jointly across the tree. The standard upwards-downwards (Felsenstein-style) recursion applies because the closed-form per-branch transition kernel from the Riccati composes across branches: the marginal at each internal node, conditional on the rest of the tree, is a closed-form pointwise distribution computable in $O(1)$ given the per-branch kernel evaluations. The augmentation thus extends to a full tree-level closed-form sampler.

##### Marginalizing the augmentation

Gravestones are an augmentation of the latent state, not of the data. The marginal likelihood of the observed alignment under the model is recovered by summing over gravestone configurations at every internal node and along every branch. The recursive traceback samples the augmented state, and the variational ELBO of §D.4 applied segment-wise on the augmented branches gives the conditional substitution likelihood. No other approximation enters at the indel level: the SCFG is the exact representation of TKF92's fragment-level dynamics, and the gravestone augmentation is what makes its inside-outside computable.

*Remark* D.11 (Coalescent priors; ARG augmentation as a parallel construction). The per-branch SCFG depends on branch length only through $t_b$, so any prior on the tree — including a coalescent prior on branch times — composes with the gravestone-augmented sampler without modification. This is a free generalization that supports joint indel-aware coalescent inference on protein-coding regions, which existing coalescent frameworks typically handle by treating indels as missing data. Separately, the structural mechanism — augmenting an unbounded latent process with non-observable entities so that an observed count provides a state-space bound — carries over to the ancestral recombination graph: non-ancestral lineages in the ARG play the role of gravestones, bounding the per-genome-position bridge state space and admitting (we conjecture) a parallel closed-form recursive sampler.

#### D.6 Posterior sampling and parameter learning

We turn to the inverse problem: given a corpus of MSAs $\{X_n\}_{n=1}^N$ with associated trees $\{T_n\}$ and fixed alignment paths, learn the corpus-level substitution hyperparameters

$$\theta = \big(S,\, \alpha_z,\, \alpha_c,\, \alpha_H,\, a_\eta, b_\eta,\, \kappa_\pi, \bar{\pi},\, \{\mu_{kl}, \tau_{kl}\}_{k\le l}\big),$$

namely the shared exchangeability $S$, the three DP concentrations $(\alpha_z, \alpha_c, \alpha_H)$, the per-site rate prior $(a_\eta, b_\eta)$, and the base-measure parameters for $G_0$ on $\pi^{(c)}$ and for $G_0^H$ on the entries of each Potts atom $H_h$. The per-site rate multipliers $\{\eta_s\}$, the per-class profiles $\{\pi^{(c)}\}$, the materialized Potts atoms $\{H_h\}$, and the class-pair Potts assignments $\{h_{cc'}\}$ are no longer parameters but *latent* draws from the base measures; their posteriors are inferred per site, per class, per atom, and per class-pair respectively, with the corpus-level parameters governing the priors. Indel parameters $(\lambda, \mu, r)$ decouple cleanly at the parameter level via the alignment–substitution factorization of §D.1 and are learned from the alignment marginal alone by standard TKF92 EM; we focus on the substitution side. The latent variables on each MSA are the key-DP labels $\{z_s\}$, the site classes $\{c_s\}$, the per-site rates $\{\eta_s\}$, the within-class amino-acid trajectories $\{\sigma_C\}$, the unobserved internal-node states, the Potts-atom assignments $\{h_{cc'}\}$, and (per §D.5) the gravestone-augmented indel histories on every branch — comprising the alive and gravestone counts $(N_a, N_g)$ at every internal node, plus the parentage tree and event times within each branch.

##### Variational EM with bridge-expectation sufficient statistics

For singleton key-DP classes, $Q^s$ reduces to F81 with class-specific equilibrium, shared exchangeability, and per-site rate. The bridge expectations (25) — expected transition counts $N_{xy}^{(b,s)}$ and dwell times $T_x^{(b,s)}$ along each branch $b$, conditioned on the endpoint amino acids — are computable in closed form by integrating $\int_0^{t_b} e^{Qu} A\, e^{Q(t_b-u)}\, du$ in the eigenspace of the rate matrix. Aggregated appropriately they give closed-form posterior updates on every continuous latent on the substitution side:

- $\pi^{(c)}$ has a Dirichlet posterior under the Dirichlet$(\kappa_\pi \bar{\pi})$ prior, via the secret-destination augmentation of §D.6 below.
- $\eta_s$ has a Gamma posterior under the Gamma$(a_\eta, b_\eta)$ prior; both shape and rate updates are immediate from the bridge expectations. The marginal site likelihood $\int$ Gamma$(a_\eta, b_\eta) \cdot$ Poisson$(N_s^{\text{acc}} \mid \eta_s \tilde{T}_s)\, d\eta_s$ is closed-form Negative-Binomial.
- Each materialized Potts atom $H_h$ has a Gaussian posterior under the $\mathcal{N}(\mu_{kl}, \tau_{kl}^{-1})$ prior, via the Gaussian (Hessian-based) Laplace approximation of the path-DCA likelihood at the current iterate.

The shared exchangeability $S$ aggregates over all classes as in standard GTR EM. Site-class assignments $c_s$ and Potts-atom assignments $h_{cc'}$ are latent in the CRP sense, integrated over via the Gibbs sweeps below.

For multi-site DP classes the joint generator on $\mathcal{A}^{|C|}$ has no usable eigendecomposition, and the bridge expectations cannot be applied to the coupled bridge directly. The variational construction of §D.4 is the substitute: at the variational fixed point each $s \in C$ has an inhomogeneous single-site bridge with rate $\tilde{Q}^{s,*}$ on each constant-composition segment of the branch (§D.5), and the bridge expectations apply to this bridge in its eigenspace. The integral identity goes through with $Q$ replaced by $\tilde{Q}^{s,*}(u)$ (piecewise-constant on a time grid within each segment), yielding per-site sufficient statistics $\tilde{N}_{xy}^{(b,s)}$, $\tilde{T}_x^{(b,s)}$ as well as per-pair co-occupancy times

$$\tilde{T}_{xy}^{(b,s,s')} \;=\; \int_0^{t_b} q_s(x;u)\, q_{s'}(y;u)\, du,$$

the latter immediate from the variational independence of single-site bridges given endpoints.

The envelope theorem ensures that, evaluated at the variational fixed point, the gradient of the ELBO with respect to $\theta$ has no contribution from the implicit $\theta$-dependence of $\tilde{Q}^{s,*}$:

$$\nabla_\theta \mathcal{L} \;=\; \mathbb{E}_{\mathbb{Q}}\big[\nabla_\theta \log \tfrac{d\mathbb{P}}{d\mathbb{Q}}\big] \;=\; \mathbb{E}_{\mathbb{Q}}\big[\nabla_\theta \log d\mathbb{P}\big].$$

For the GTR-Potts generator this gradient has three structurally distinct pieces. The exchangeability gradient $\nabla_S \mathcal{L}$ is a linear combination of the single-site $\tilde{N}^{(b,s)}_{xy}$ counts as in standard GTR EM. The frequency gradient $\nabla_{\pi^{(c)}}\mathcal{L}$ aggregates over sites with $c_s = c$ and is again single-site. The coupling-tensor gradient is a path-space DCA M-step,

$$\nabla_{H_{cc'}(x,x')}\mathcal{L} \;\propto \sum_{n,b} \sum_{\substack{s,s' \in C(n) \\ c_s = c,\, c_{s'} = c'}} \Big[ \big(\text{expected } x_s = x,\, x_{s'} = x' \text{ co-occupancy under } \mathbb{P}\big) - \big(\text{same under } \mathbb{Q}\big) \Big],$$

expressible in the same bridge-expectation format using the products $q_s(x;u)q_{s'}(x';u)$ for the variational dwell-time terms and contributions from each variational jump intensity for the transition terms.

##### MCMC over the discrete latents

The combinatorial latents $\{z_s\}$, $\{c_s\}$, and $\{h_{cc'}\}$ admit straightforward Gibbs sampling. All three partitions are integrated over their respective stick-breaking weights via Chinese restaurant predictives. For the key DP,

$$P(z_s = k \mid z_{-s}, \alpha_z) \;\propto\; \begin{cases} |C_k^{(-s)}| \cdot \mathcal{L}_{\text{class}}(C_k \cup \{s\}) / \mathcal{L}_{\text{class}}(C_k) & \text{(existing key class)}, \\ \alpha_z \cdot \mathcal{L}_{\text{class}}(\{s\}) & \text{(new singleton)}, \end{cases}$$

where $\mathcal{L}_{\text{class}}$ is the per-branch-aggregated class-level variational ELBO from §D.4 plus the contribution of the rest of the tree via Felsenstein pruning. Site-class updates use the class DP predictive,

$$P(c_s = c \mid c_{-s}, \alpha_c, \theta) \;\propto\; \begin{cases} n_c^{(-s)} \cdot \mathcal{L}_{\text{site}}(s; \pi^{(c)}) & \text{(existing class, profile } \pi^{(c)}\text{)}, \\ \alpha_c \cdot \int \mathcal{L}_{\text{site}}(s; \pi')\, G_0(d\pi') & \text{(new class)}, \end{cases}$$

with the new-class integral closed form by the Dirichlet–multinomial conjugacy of §D.6. The Potts-atom assignment $h_{cc'}$ for an unordered class-pair $\{c, c'\}$ is itself drawn by CRP-Gibbs,

$$P(h_{cc'} = h \mid \text{rest}, \alpha_H) \;\propto\; \begin{cases} m_h^{(-cc')} \cdot \mathcal{L}_{\text{pair}}(\{c,c'\};\, H_h) & \text{(existing atom)}, \\ \alpha_H \cdot \int \mathcal{L}_{\text{pair}}(\{c,c'\};\, H')\, G_0^H(dH') & \text{(new atom)}, \end{cases}$$

where $\mathcal{L}_{\text{pair}}$ is the path-DCA contribution from co-occurrences of classes $c$ and $c'$ across the corpus, and the new-atom integral is approximated by the Laplace scheme of §D.6.

Split–merge proposals of Jain–Neal type (27) are essential for mixing on all three partitions, with acceptance ratios in terms of class-level (for $z$), site-aggregated (for $c$), or class-pair-aggregated (for $h$) ELBO ratios. All three concentrations $\alpha_z, \alpha_c, \alpha_H$ admit closed-form Gamma-prior posteriors given the respective partitions via the auxiliary-variable scheme of Escobar & West (14).

##### Truncated stick-breaking on Potts atoms

The Potts-atom CRP admits a truncated stick-breaking (TSB) alternative that is cheaper per sweep, requires no new-atom Laplace branch inside the Gibbs step, and removes the empty-atom bookkeeping of the CRP. The construction mirrors Ishwaran & James (26) and Blei & Jordan (5) as applied to the site-class DP, transposed to the Potts-atom DP.

Fix a truncation level $K_H^{\max}$. The natural choice is $K_H^{\max} = K_c(K_c+1)/2$, the total number of unordered class-pairs — with this bound the truncation is exactly tight (every class-pair could in principle have its own atom). Replace the CRP-Gibbs sweep over $\{h_{cc'}\}$ with the following two-step alternation:

1. **Beta CAVI / Gibbs on stick proportions.** Maintain stick variables $\beta_h \sim \mathrm{Beta}(1, \alpha_H)$ for $h < K_H^{\max}$, with $\beta_{K_H^{\max}} = 1$ and $\rho_h = \beta_h \prod_{h'<h}(1-\beta_{h'})$. Given the current per-atom counts $m_h = \#\{(c,c') : h_{cc'} = h\}$, the conditional posterior on $\beta_h$ is the closed-form Beta

$$\beta_h \mid m, \alpha_H \;\sim\; \mathrm{Beta}\Big(1 + m_h,\; \alpha_H + \sum_{h'>h} m_{h'}\Big).$$

   Use either the posterior mean (CAVI) or a Beta sample (Gibbs).

2. **Categorical resample on $\{h_{cc'}\}$.** For each unordered class-pair $\{c, c'\}$, draw the atom assignment from the closed-form Categorical

$$P(h_{cc'} = h \mid \text{rest}) \;\propto\; \rho_h \cdot \mathcal{L}_{\text{pair}}(\{c, c'\};\, H_h),$$

   where $\mathcal{L}_{\text{pair}}$ is the path-DCA contribution from co-occurrences of classes $c$ and $c'$ across the corpus, evaluated at the current materialized atom $H_h$.

*Remark* D.12 (Bijective initialization). Initialization at $K_H^{\max} = K_c(K_c+1)/2$ should set each unordered class-pair to a *distinct* atom drawn from $G_0^H$, not collapse all class-pairs onto a single shared atom. The latter is a chicken-and-egg failure mode: only the populated atom receives data through the Laplace M-step, so the others stay at the prior; the next TSB resample therefore prefers the populated atom for all pairs (its likelihood dominates the others' prior-only score), and the system collapses. The bijective init guarantees every atom carries at least one class-pair's data through the first Laplace M-step.

##### Locality of DP updates under the SCFG augmentation

When the augmented indel history is resampled (§D.5), new gravestone fragments appear with new residues that need DP keys, and old gravestones disappear. Naively this would suggest a full DP-key resample after every augmentation step, which would be prohibitive. Exchangeability of the CRP and the segment-wise structure of the variational ELBO together remove almost all of this cost.

The decomposition is two-step within a single sweep. The *indel-skeleton step* resamples the gravestone-augmented history on every branch by the SCFG traceback of §D.5 without touching DP keys; cost $O((b+d)\log(b+d))$ per branch, independent of DP state. The *gravestone-key step* then assigns DP keys to each new residue $s$ on a gravestone fragment with lifetime $[u, v]$ on branch $b$, using the local CRP predictive over existing classes plus a singleton, evaluated against $\mathcal{L}_{\text{seg}}$ on segments of $b$ within the gravestone's lifetime. Adding $s$ to class $C_k$ changes that class's variational fixed point only on those segments, costing $O(|C_k| \cdot A^3)$ per fixed-point iteration per touched segment. Per gravestone, the work is one Categorical draw with support equal to the existing classes plus a singleton, plus one local fixed-point re-solve on the affected segments.

**Closed-form base-measure integrals via complete-data augmentation**

The base-measure integrals

$$\int \mathcal{L}_{\text{site}}(s;\pi')\,G_0(d\pi'), \qquad \int \mathcal{L}_{\text{pair}}(\{c,c'\};\,H')\,G_0^H(dH'),$$

that appear in the new-class and new-atom branches of the CRP predictives, and the analogous corpus-level marginals over $\eta_s$, all admit closed-form approximations under complete-data augmentation.

**Secret-destination augmentation for $\pi^{(c)}$.** The F81-form rate factors as $Q^s_{xy} = \eta_s \cdot S_{xy} \cdot \pi^{(c_s)}(y) \cdot \exp(-\frac{1}{2}\Delta H_s)$, which decomposes the dynamics into an $S$-driven Poisson proposal clock at rate $\eta_s S_{xy}$ from state $x$ to destination $y$, modulated by a Bernoulli filter that accepts the proposal with probability $\pi^{(c_s)}(y)\exp(-\frac{1}{2}\Delta H_s)$. To recover full multinomial structure on $\pi^{(c)}$ we augment each silent failure of a proposal of $y$ with a categorical *secret destination* $j \neq y$, drawn from $\pi^{(c)}(j)/(1-\pi^{(c)}(y))$ on the simplex. Each proposal then casts exactly one categorical vote: the destination is $y$ with probability $\pi^{(c)}(y)$ on acceptance, and a secret $j$ with probability $\pi^{(c)}(j)$ on rejection. Marginally over the proposal, the vote is Categorical$(\pi^{(c)})$, so the augmented count vector $N^{(c)} \in \mathbb{Z}^A_{\geq 0}$ satisfies

$$N^{(c)} \mid \pi^{(c)}, \text{augmentation} \;\sim\; \text{Multinomial}\big(N^{(c)}_{\text{total}},\, \pi^{(c)}\big),$$

and the posterior is the strict closed-form Dirichlet

$$\pi^{(c)} \mid N^{(c)} \;\sim\; \text{Dirichlet}\big(\kappa_\pi \bar{\pi} + N^{(c)}\big).$$

In the EM/VBEM setting the augmentation is replaced by its conditional expectation: each proposal contributes $\pi^{(c)}(y)$ to $E[N^{(c)}_y]$ regardless of which destination was proposed. The Dirichlet update is then a fixed-point iteration whose limit is the conjugate posterior under the conditional augmentation. The new-class integral $\int \mathcal{L}_{\text{site}}(s;\pi')\,G_0(d\pi')$ is the analytic ratio of multivariate Beta functions $B(\kappa_\pi\bar{\pi} + N^{(c)})/B(\kappa_\pi\bar{\pi})$.

**Gamma–Poisson conjugacy for $\eta_s$.** The F81 substitution-count likelihood at site $s$ is itself Gamma-conjugate in $\eta_s$ without any need for the secret-destination augmentation. The expected substitution count along the tree is Poisson with rate $\eta_s \cdot \tilde{T}_s$, where

$$\tilde{T}_s \;=\; \sum_b \sum_x T^{(b,s)}_x \cdot \sum_{y \neq x} S_{xy}\,\pi^{(c_s)}(y)$$

is the $\pi$-weighted dwell-time integral. Under the Gamma$(a_\eta, b_\eta)$ prior the conjugate posterior is closed form

$$\eta_s \mid \text{path}, \pi^{(c_s)} \;\sim\; \text{Gamma}\big(a_\eta + N^{\text{acc}}_s,\; b_\eta + \tilde{T}_s\big),$$

with $N^{\text{acc}}_s = \sum_b \sum_{x \neq y} N^{(b,s)}_{xy}$ the total observed substitution count, and the per-site marginal likelihood is closed-form Negative-Binomial.

**Laplace for $H_h$.** Each Potts atom is real-valued and the path-DCA likelihood is locally Gaussian in $H_h$ to second order. We approximate

$$\log p_{\text{path-DCA}}(\text{data} \mid H_h) \;\approx\; \log p_{\text{path-DCA}}(\text{data} \mid \hat{H}_h) \;-\; \tfrac{1}{2}(H_h - \hat{H}_h)^\top \Lambda_h (H_h - \hat{H}_h),$$

where $\hat{H}_h$ is the MAP under the current iterate, found by a few Newton or natural-gradient steps starting from a chosen seed, and $\Lambda_h = -\nabla^2_{H_h} \log p_{\text{path-DCA}}$ is the Hessian at $\hat{H}_h$. Combined with the $\mathcal{N}(\mu_{kl}, \tau_{kl}^{-1})$ prior on each entry, the posterior is the closed-form Gaussian

$$H_h \mid \text{data} \;\sim\; \mathcal{N}\big(\hat{H}_h^{\text{post}},\, (\operatorname{diag}(\tau_{kl}) + \Lambda_h)^{-1}\big).$$

The Laplace estimate of the new-atom integral is the standard $\log p_{\text{path-DCA}}(\hat{H}_h) + \log G_0^H(\hat{H}_h) + \frac{d}{2}\log(2\pi) - \frac{1}{2}\log\det(\operatorname{diag}(\tau_{kl}) + \Lambda_h)$ with $d = A(A+1)/2$ the symmetric-slice dimension.

**Multi-seed Laplace mixture for multimodal posteriors.** The Laplace approximation collapses around a single mode and underestimates posterior mass elsewhere; for atoms that admit multiple plausible coupling patterns (e.g., charge-complementary vs. hydrophobic-hydrophobic minima) this is a genuine approximation error. We mitigate by running the few-step optimizer from a small number $K_{\text{seed}}$ of distinct seeds, obtaining $K_{\text{seed}}$ Laplace components, and combining them as a weighted Gaussian mixture.

**The recommended hybrid**

The natural hybrid is:

1. **Inner variational E-step.** For each MSA, branch and DP class, segment the branch by the current sampled augmented indel history and solve the geometric-mean fixed point of §D.4 on each segment to obtain $\{\tilde{Q}^{s,*}\}$ and the per-class ELBO. No parameter or structure updates here.

2. **MCMC over discrete and augmentation latents.** Resample augmented indel histories on every branch by recursive traceback on the time-indexed pair SCFG of §D.5, with closed-form midpoint marginals from the bivariate Riccati and Felsenstein-style upwards–downwards composition for internal-node augmentation counts. Gibbs sweep over $\{z_s\}$, $\{c_s\}$, and $\{h_{cc'}\}$ using the class-level ELBOs from step 1 as evidence; periodic Jain–Neal split–merge proposals on each partition.

3. **Stochastic M-step.** Compute bridge-expectation sufficient statistics $\tilde{N}$, $\tilde{T}$, $\tilde{T}_{\text{pair}}$ from the variational single-site bridges, summed over augmented constant-composition segments within each branch, over a minibatch of MSAs. Take a stochastic gradient step on the corpus-level hyperparameters $\theta = (S, \alpha_z, \alpha_c, \alpha_H, a_\eta, b_\eta, \kappa_\pi, \bar{\pi}, \{\mu_{kl}, \tau_{kl}\})$. Per-class $\pi^{(c)}$ updates by its closed-form Dirichlet posterior under the secret-destination augmentation; per-site $\eta_s$ marginalizes in closed form via Gamma–Poisson conjugacy; per-atom $H_h$ updates by Laplace with multi-seed mixture against the prior $G_0^H$. The three DP concentrations update by the Escobar–West auxiliary-variable scheme.

4. **Iterate.** The inner variational fixed point in step 1 is re-solved cheaply after each step because the previous solution is a warm start.

This factors the overall problem into the three regimes where each tool is strongest: variational for the continuous-time substitution dynamics where it admits closed-form rates and bridge-expectation sufficient statistics; MCMC for the discrete combinatorial latents where it mixes well and avoids mean-field pathologies; stochastic gradient for the continuous parameters $\theta$ where it scales to corpus-level training. In SVI form the M-step is over a minibatch of MSAs at each iteration and amortizes naturally over the corpus.

**Counts-tensor representation of the training corpus**

Per family we precompute a counts tensor (cherry $\times$ column $\times$ AA-pair $\times$ branch length), packed as int8, that decouples inference from raw MSA parsing and lets one outer SVI iteration scale to $10^3$–$10^4$ PFAM families. Branch lengths are quantized to $n_t \approx 50$ geometrically spaced bins between $\tau_{\min}$ and $\tau_{\max}$; geometric spacing exploits that $\exp(Q\tau)$ varies fastest at small $\tau$, giving a $\sim 5\times$ cache reduction over linear quantization at no measurable accuracy cost. Per-class single-site log-transition matrices $(K_c, n_t, A^2)$ are precomputed once per outer iteration and shared across all families.

#### D.7 Pairwise alignment postprocessing

Multiple-sequence alignment tools of the FSA family (**?** ) (and other consistency-based schemes) operate on pairwise residue posterior matrices $Q_{ij}^{(X,Y)} = P(X_i \sim Y_j \mid X, Y)$ produced by per-pair forward–backward passes through a Pair HMM. The TKF-DP postprocessing target is the joint-marginal alignment-and-partition likelihood described next, in which an alignment $A$ together with a size-$\{1,2\}$ partition $E$ of its Match cells contributes a product of per-block substitution factors with no class or rate-multiplier latent left explicit.

*Remark* D.13 (Historical note: deprecated first-order correction). An early formulation of TKF-DP postprocessing expressed the coupling as a first-order mean-field correction $\log Q'_{ij} - \log Q_{ij} \approx \varepsilon \sum_{(i',j')} Q_{i'j'}[M(i,j;i',j';t) - 1]$ to the baseline single-site posterior, with $\varepsilon$ a per-pair partner prior under the size-$\{1,2\}$ Ewens distribution and $M$ a column-class-marginalized Potts boost. That correction is a first-order expansion of the joint marginal in the partner-prior strength and is now considered a heuristic stepping-stone; the modern infinite Pair HMM (§E.3 of Appendix E) treats the same coupling exactly via the joint-marginal target below, with class assignments and rate multipliers integrated analytically rather than re-marginalized at every column.

**The unified joint-marginal target**

Let $\theta_{\text{indel}} = (\lambda, \mu, r)$ be the TKF92 indel parameters and $\alpha_z$ the Ewens partition concentration. Write $\text{Match}(A)$ for the Match cells of an alignment $A$, and let $E$ be a set of unordered Match-cell edges (a partition of $\text{Match}(A)$ into singletons of size 1 and doublets of size 2; we let $V(E) \subset \text{Match}(A)$ denote the doublet endpoints). The TKF-DP joint marginal at branch length $t$ is

$$P(X,Y;t) \;=\; \sum_{A} \pi_{\text{TKF92}}(A \mid t, \theta_{\text{indel}}) \sum_{E} \pi_{\text{Ewens}}(E \mid \alpha_z, |\text{Match}(A)|) \prod_{(i,j)\in \text{Match}(A)\setminus V(E)} P_{\text{singlet}}(x_i, y_j; t) \prod_{e=((i,j),(k,l))\in E} P_{\text{d}} \text{[illegible]} \tag{D.3}$$

with the per-block primitives

$$P_{\text{singlet}}(a,b;t) \;=\; \sum_c \pi_c\; \pi^{(c)}(a)\; P_c(a \to b;\, t\cdot\eta), \tag{D.4}$$

$$P_{\text{doublet}}\big((a,c),(b,d);t\big) \;=\; \sum_{c_1,c_2} \pi_c(c_1)\, \pi_c(c_2)\; \pi^{c_1,c_2}_{\text{joint}}(a,c)\; P^{c_1,c_2}_{\text{joint}}\big((a,c)\to(b,d);\, t,\eta_1,\eta_2\big). \tag{D.5}$$

The doublet endpoint convention (matching the boost API of the released code) is that $(a,b)$ are the amino acids at the left endpoint $(i,j)$ and $(c,d)$ at the right endpoint $(k,l)$: $a = x_i$, $b = y_j$, $c = x_k$, $d = y_l$. Indices $c_1, c_2$ run over site classes and should not be confused with the endpoint-amino-acid index $c$.

The empirical class prior $\pi_c$ is the per-class column count from training (*not* uniform $1/K_c$); the per-class profile $\pi^{(c)}(\cdot)$ is the trained `pi_class`; the class-conditional transition is

$$P_c(a \to b;t) \;=\; [\exp(Q_c\cdot t)]_{a,b}, \qquad Q_c \;=\; (S_{\text{LG08}} - \operatorname{diag} S_{\text{LG08}})\cdot \pi^{(c)}[\,\cdot\,]^{\top}, \tag{D.6}$$

the standard F81-form GTR generator with exchangeability $S_{\text{LG08}}$ (32) and class-specific stationary $\pi^{(c)}$. The joint stationary $\pi^{c_1,c_2}_{\text{joint}}(a,c)$ at a coupled cell-pair is the joint stationary of a Potts atom $H$ = `atoms`[`assignments`$[c_1,c_2]$] at the trained checkpoint (the trained `potts_dp.atoms` indexed by `potts_dp.assignments`), and $P^{c_1,c_2}_{\text{joint}}$ is its time-evolved transition $\exp(Q^{c_1,c_2}_{\text{joint}}(H)\cdot t)$ built by `build_joint_Q_pair`$(H, \pi_a, \pi_b, S, \eta_{\text{pair}} = (\eta_1,\eta_2))$ as described in §D.3.

**Pair-background convention for the joint generator.** Two natural choices for the pair background $(\pi_a, \pi_b)$ arise. The first is to use the LG08 single-site equilibrium $\pi_a = \pi_b = \pi^{\text{LG08}}$ for every class-pair $(c_1, c_2)$. This is the *canonical* convention for the released $K$=4 EM-warmup checkpoint: the trained Potts atoms were fit as deviations of the joint stationary from the LG08-pair-background reference, so inference under that checkpoint must use the same convention for the joint stationary and joint generator to match training. The second choice, $\pi_a = \pi^{(c_1)}$, $\pi_b = \pi^{(c_2)}$, uses the per-class profiles as the pair background; this is the structurally "cleaner" choice for future model variants in which the Potts atoms are retrained against the per-class background, but it is *not* consistent with the released checkpoint.

**Discrete-gamma rate-multiplier integration.** The per-site rate multiplier $\eta_s \sim \text{Gamma}(a_\eta, b_\eta)$ of §D.3 is integrated analytically by the discrete-gamma quadrature of Yang (54): $K_r$ equiprobable bin medians $\eta_1, \ldots, \eta_{K_r}$ are taken from the $\text{Gamma}(a_\eta, b_\eta)$ prior with uniform weight $1/K_r$, and the per-block primitive of equation D.4 (resp. D.5) is averaged over rate-multiplier draws via $P_{\text{singlet}}(a,b;t) = K_r^{-1}\sum_{r=1}^{K_r} P_{\text{singlet}}(a,b;t\cdot\eta_r)$ (resp. a double sum over $(\eta_1,\eta_2)$ for the doublet). The Gamma hyperparameters $(a_\eta, b_\eta)$ at inference time must match those used at training time; the released $K$=4 EM-warmup checkpoint uses $(a_\eta, b_\eta) = (2,2)$. The reduction $K_r = 1$ with $\eta = 1$ is the canonical evaluation of that checkpoint, since the stored per-site $\eta$ posteriors all collapsed to 1.0 during EM warmup; for retrained checkpoints with a non-trivial $\eta$ posterior, $K_r > 1$ is required.

$t = 0$ **reduction identities.** At $t = 0$ both per-block primitives reduce to the appropriate marginal stationary:

$$P_{\text{singlet}}(a,a;\,0) \;=\; \sum_c \pi_c\, \pi^{(c)}(a) \;\equiv\; \pi_{\text{marg}}(a), \tag{D.7}$$

$$P_{\text{doublet}}\big((a,c),(a,c);\,0\big) \;=\; \sum_{c_1,c_2} \pi_c(c_1)\, \pi_c(c_2)\, \pi^{c_1,c_2}_{\text{joint}}(a,c) \;\equiv\; \pi_{\text{marg,pair}}(a,c). \tag{D.8}$$

These follow from $P_c(a \to b; 0) = \delta_{ab}$ and from the joint stationary $\pi^{c_1,c_2}_{\text{joint}}$ being a fixed point of $P^{c_1,c_2}_{\text{joint}}(\cdot; 0)$.

**The boost tensor.** The four-residue Potts *boost* entering equation D.3 through the doublet factor is the multiplicative deviation of the doublet emission from the product of two singlet emissions:

$$M\big((a,c),(b,d);t\big) \;\equiv\; \frac{P_{\text{doublet}}\big((a,c),(b,d);t\big)}{P_{\text{singlet}}(a,b;t)\cdot P_{\text{singlet}}(c,d;t)}. \tag{D.9}$$

Indexing convention as above: $(a,b)$ is the left endpoint, $(c,d)$ the right endpoint. By construction $M \equiv 1$ when every Potts atom is zero (so $\pi^{c_1,c_2}_{\text{joint}}$ factorizes into $\pi^{(c_1)} \otimes \pi^{(c_2)}$ and $P^{c_1,c_2}_{\text{joint}}$ factorizes into the two single-site $P_{c_i}$ factors). The boost tensor $M$ is the object that the bounded-edge augmented Pair HMM methods of Appendix E fold multiplicatively into the Match emissions to score doublets relative to singletons. The MCMC sampler operates on the joint $(A,E)$ directly via equation D.3, with class assignments and rate multipliers integrated analytically as above (*not* sampled).

##### Sequence annealing with coevolutionary scoring

The joint-marginal of equation D.3 implies a per-Match-cell posterior $Q'_{ij}$ that the augmented Pair HMM machinery of Appendix E computes either exactly under bounded-edge truncation or stochastically under the infinite-Pair-HMM MCMC. An alternative routing of the same coupling signal applies the boost *during* assembly: the greedy column-merging step inside the sequence annealer is extended to score pairs of merges jointly. With pair-HMM input, the canonical edge score is

$$w(e) \;=\; \frac{2\,Q^{(X,Y)}_{ij}}{g^{(X,Y)}_i + g^{(Y,X)}_j}, \qquad e = \big(\text{column of } X_i,\ \text{column of } Y_j\big), \tag{D.10}$$

where $g$ are the per-residue gap posteriors. The coupled-pair extension admits joint candidates consisting of pairs of merges scored as

$$W(e_1,e_2) \;=\; \log Q^{(X,Y)}_{ij} + \log Q^{(X,Y)}_{i'j'} + \log M\big((X_i,X_{i'}),(Y_j,Y_{j'});t\big), \tag{D.11}$$

with $M$ the boost tensor of equation D.9 evaluated at the observed amino acids. The first two terms are the baseline log-posteriors, and the third is the four-residue Potts contribution; the latter is identically zero when every Potts atom is zero.

**Greedy schedule and conflict resolution.** When a coupled candidate $(e_1,e_2)$ is popped from the queue, conflicts are resolved as follows. If both component merges are still admissible (no same-sequence conflict, no DAG cycle), both are committed in a single step and the boost-driven joint commit is recorded. If $e_1$ is admissible but $e_2$ has been invalidated by an intervening merge, the candidate degrades to a single-edge merge with score $\log Q_{ij}$ and is re-inserted into the queue at that priority. If neither component is admissible, the candidate is dropped.

**Cost analysis.** Under two practical prunings (threshold $q_{\min} = 0.1$ on the single-edge posterior and threshold $\mu_{\min} = 0.1$ nat on $|\log M|$), the per-step cost is $O(L^2)$ amortized and the total cost is $O(L^3)$ in the worst case, comparable to the $O(L^2 \log L)$ baseline assembler.

**Augmented-PHMM versus greedy-pairwise tradeoff.** The augmented-Pair-HMM and infinite-Pair-HMM methods of Appendix E produce a single corrected posterior matrix $Q'$ that any downstream assembler can consume. The coupled-pair extension above scores strongly-coupled pairs jointly on their native four-residue domain inside the assembler, but commits greedily and is therefore vulnerable to local optima where an early commit on a coupled pair locks out a globally better arrangement. The two routes consume the same boost tensor $M$ of equation D.9 but apply it at different points in the pipeline; the augmented-PHMM route is the principled default for TKF-DP postprocessing in §E.

### E The infinite Pair HMM and its MCMC sampler

The Pair HMM postprocessing methods of §D.7 are progressively faithful approximations to a single underlying object: the alignment-and-partition posterior on $\pi(A, \pi_M \mid X, Y)$, where $A$ is the TKF92 alignment and $\pi_M$ is a Chinese-restaurant-process partition of its Match cells into singletons (block of size 1, contributing the standard per-site CTMC factor) and pairs (block of size 2, contributing the joint Potts CTMC factor). This appendix develops the three-factor factorisation $\pi_{\text{TKF92}} \cdot \pi_{\text{CRP}} \cdot \prod_b P_{\text{block}}$ in full; describes the exact $F_2$-SCFG inside–outside formulation that achieves it at $O(L^4)$; re-encodes the same content as a memory-augmented Pair HMM at $O(L^2A^2)$; and develops the Gibbs+MH+replica-exchange MCMC sampler that draws from the unbounded-edge limit at amortised $O(L^3)$ per sweep. The conceptual hierarchy at the end of the appendix lifts this construction to the full multi-branch *phylogenetic* SCFG, the natural fixed point of the entire ladder of approximations.

#### E.1 Exact 0-or-1-edge marginal posteriors via Pair-SCFG inside-outside

The deprecated mean-field correction (remark D.13 of Appendix D) and the coupled-pair greedy annealer of §D.7 are two routes to the same intractable posterior of equation D.3; the present subsection derives an exact dynamic-programming alternative under the hard restriction that the coupling partition $E$ contains zero or one edge. The result is a per-residue-pair posterior $Q'_{ij}$ that consumes the same downstream FSA assembler as the baseline, with the coupling evidence absorbed exactly into the posterior rather than fed in via a per-pair correction or via a custom annealer move set.

##### Three pair-HMM marginal moments

Under the no-coevolution baseline TKF92 Pair HMM with the class-mixture single-site emissions of Appendix D, let $\pi(X, Y, A)$ denote the joint probability of an alignment $A$ between sequences $X$ and $Y$. Define

$$F_0(X,Y) \equiv \sum_A \pi(X,Y,A), \tag{E.1}$$

$$F_1(X,Y;i,j) \equiv \sum_{A\,:\,X_i \sim_A Y_j} \pi(X,Y,A), \tag{E.2}$$

$$F_2(X,Y;i,j;k,l) \equiv \sum_{A\,:\,X_i \sim_A Y_j \,\wedge\, X_k \sim_A Y_l} \pi(X,Y,A), \tag{E.3}$$

where $X_i \sim_A Y_j$ denotes that the alignment $A$ pairs residue $X_i$ with $Y_j$ in a Match emission. Computationally, $F_0$ is a Forward in $O(L^2)$, $F_1$ is a Forward–Backward in $O(L^2)$, and $F_2$ is the

Inside–Outside table of a non-bifurcating Pair SCFG (or equivalently a doubled Forward–Backward) in $O(L^4)$ time and memory. The conventional Pair-HMM match-state posterior is $Q_{ij} \equiv F_1(i,j)/F_0$.

The objects $F_1$ and $F_2$ stand in the relation that gradients and Hessians do to a partition function: $F_1(i,j)/F_0$ is the marginal probability that $(X_i, Y_j)$ is paired and $F_2(i,j;k,l)/F_0$ is the marginal joint probability that both $(X_i, Y_j)$ and $(X_k, Y_l)$ are paired, with the no-coevolution model serving as the unperturbed reference point.

##### The 0-or-1-edge Pair SCFG

Let $M(i,j;k,l;t) \equiv M((X_i, X_k),(Y_j, Y_l);t)$ denote the four-residue Potts coupling boost of equation D.9 of Appendix D, evaluated at the observed residues $(X_i, Y_j, X_k, Y_l)$ and the inferred branch length $t$. Define the (unnormalised) Pair-SCFG generating function

$$S \;\to\; S_0 \;\big|\; S_1, \qquad \text{with multiplicative weights 1 and } \varepsilon = 1/\alpha_z, \tag{E.4}$$

where the two non-terminals expand as

$$\begin{aligned} S_0 &\;\to\; \text{TKF92 Pair HMM with zero coupled column-pairs (the baseline),} \\ S_1 &\;\to\; \text{TKF92 Pair HMM with exactly one coupled column-pair.} \end{aligned}$$

The 1-edge nonterminal's parse picks one ordered column-pair $(i,j;k,l)$ with $i < k$ as the coupled pair and replaces the product of independent single-site Match emissions at columns $(i,j)$ and $(k,l)$ with the joint pair-emission $P_{\text{joint}}((X_i, X_k) \to (Y_j, Y_l); t, H)$, equivalently the product of independent emissions multiplied by $M(i,j;k,l;t)$. The partition function of the grammar is

$$L_{\text{exact}}(X,Y) \;=\; F_0(X,Y) \,+\, \varepsilon \sum_{(i,j;k,l),\; i<k} F_2(X,Y;i,j;k,l) \cdot M(i,j;k,l;t). \tag{E.5}$$

The first term is the no-coupling marginal likelihood; the second is the marginal contribution of all alignments that contain exactly one coupled column-pair, summed over all candidate locations $(i,j;k,l)$ for that pair. The $\varepsilon$ prefactor is the per-pair partner prior under the size-$\{1,2\}$ Ewens partition with concentration $\alpha_z$; the assumption restricting to at most one edge per alignment is a hard-truncation of the size-$\{1,2\}$ partition that retains the leading-order correction in $\varepsilon$.

##### The exact match posterior

Inside–Outside on the grammar (E.4) yields the marginal probability that residue $X_i$ is aligned to $Y_j$ in the data:

$$Q'_{ij} \;=\; \frac{F_1(X,Y;i,j) \,+\, \varepsilon \sum_{(k,l)\,:\,k\neq i} F_2(X,Y;i,j;k,l) \cdot M(i,j;k,l;t)}{L_{\text{exact}}(X,Y)}. \tag{E.6}$$

The numerator collects all alignment-paths through the SCFG that touch $(i,j)$ as a Match: the first term covers $S_0$ paths (no coupled pair touches $(i,j)$), the second term covers $S_1$ paths in which $(i,j)$ is part of the (unique) coupled pair, summed over the choice of partner column $(k,l)$. Both summands include the boost factor $M$ at exactly the right place. The denominator is the partition function (E.5).

In the limit $\varepsilon \to 0$ (or $H = 0$ so $M \equiv 1$ everywhere), $Q'_{ij}$ collapses to $F_1(i,j)/F_0 = Q_{ij}$, the no-coupling baseline, as required.

##### Cost and practical considerations

The bottleneck of the explicit $F_2$ tensor encoding is the $O(L^4)$ time and memory required to materialise $F_2$ itself: $F_2$ has $\binom{L}{2}^2 \approx L^4/4$ free entries, which at $L = 200$ occupies a few GB in single precision. Longer sequences require either (a) prefix/suffix chunking on the second-anchor axis $(k, l)$ to keep memory bounded, or (b) candidate pruning that drops $(i, j)$ or $(k, l)$ pairs with $F_1(\cdot) < q_{\min}$, reducing the candidate set from $O(L^4)$ to $O(L^2)$ in practice. The reference implementation lives in `src/tkfdp/f2_scfg.py`; in the current codebase it is retained primarily as a cross-validation reference for the more efficient memory-augmented Pair HMM described in the next subsection (which agrees with $F_2$-SCFG to machine precision and runs at $O(L^2A^2)$).

The available pathways to a coupled-FSA-consumable per-residue posterior, in order of fidelity:

| Pathway | Cost | Approximation |
|---|---|---|
| Mean-field pre-correction (deprecated; rem. D.13) | $O(L^2K_c^2A^2)$ | First-order in $\varepsilon$ |
| Coupled-pair greedy annealer (§D.7) | $O(L^3)$ | Greedy commit ordering |
| $F_2$-SCFG explicit enumeration | $O(L^4)$ | 0-or-1-edge truncation, otherwise e[illegible] |
| Aug-PHMM 1-edge (§E.2) | $O(L^2A^2)$ | 0-or-1-edge truncation, otherwise e[illegible] |
| Aug-PHMM 2-edge (§E.2) | $O(L^2A^4)$ | 0-or-1-or-2-edge truncation, otherw[illegible] |
| Infinite Pair HMM MCMC (§E.3) | $O(L^4) + O(L^3) \cdot N_{\text{sweep}}$ | Stochastic; converges to exact |

The bounded-edge approximations capture progressively more of the coupling correction at progressively higher cost; the infinite Pair HMM MCMC is the principled limit, exact in expectation up to MCMC sampling error. For families dominated by a single strong coevolutionary edge per column, the 1-edge aug-PHMM at $O(L^2A^2)$ is empirically adequate; for families with multiple competing partners per column, the 2-edge aug-PHMM (or the MCMC sampler) is necessary.

#### E.2 Memory-augmented Pair HMM: the same content at $O(L^2A^2)$

The $F_2$-SCFG of equations E.4–E.6 can be re-encoded as a Pair HMM whose latent state is augmented with a small *tag memory* of in-progress coupled-edge endpoints. For the 0-or-1-edge case the tag set has $|\mathcal{T}| = 1 + A^2 + 1$ values: a sentinel `no_edge` state, $A^2$ states $(a, b)$ recording that a left endpoint with observed amino acids $(a, b)$ is awaiting its right partner, and a terminal `done` state recording that a coupled pair has been consumed. At any Match cell $(i, j)$ the HMM may either (a) carry the tag forward unchanged, (b) transition `no_edge` $\rightarrow (X_i, Y_j)$ with weight $\varepsilon$ (spawn a left endpoint), or (c) transition $(a, b) \rightarrow$ `done` with weight $M(\cdot, \cdot; X_i, Y_j; t)$ summed over the recorded $(a, b)$ (close the pair). At all non-Match states the tag is preserved verbatim. The augmented Forward partition function equals $L_{\text{exact}}$ of equation E.5, and the marginal posterior at Match-state cell $(i, j)$, summed over tag values, equals $Q'_{ij}$ of equation E.6. Cost is $O(L^2 \cdot 5 \cdot |\mathcal{T}|) = O(L^2A^2)$ time and memory: a factor $L^2/A^2 \approx 100$ speedup over the explicit $F_2$ tensor at typical sequence lengths, with no loss of fidelity. Implementation lives in `src/tkfdp/aug_phmm.py`; cross-validation against the $O(L^4)$ $F_2$-SCFG reference confirms agreement to machine precision (max $|Q'_{ij} - Q'^{\text{SCFG}}_{ij}| \approx 10^{-14}$ on tested pairs).

The same construction generalises to allow up to $k$ coupled column-pairs per alignment by enlarging the tag set to record (i) the multiset of in-flight left endpoints (size up to $k$), (ii) the number of resolved pairs (in $\{0, 1, \ldots, k\}$), and (iii) optionally the in-flight endpoints alongside the resolved-pair counter. For $k = 2$ the tag set has $|\mathcal{T}_2| = 1 + A^2 + A^2(A^2 + 1)/2 + A^2 + 1 \approx 81{,}000$ values for $A = 20$: a sentinel `no_edge`, $A^2$ singleton states (one in-flight, none resolved), an unordered-multiset pair state of size $A^2(A^2+1)/2$ (two in-flight, none resolved; same-element multisets are valid distinct states), $A^2$ closed-singleton states (one resolved, one in-flight), and a terminal `closed_done`. The match-cell transitions add a spawn from singleton→pair with weight $\varepsilon$ (multiset construction), and three closure transitions: pair→closed-singleton with weight $M(\cdot; X_i, Y_j; t)$ over the closed multiset element, singleton→closed-done with weight $M$, and closed-singleton→closed-done with weight $M$. Cost is $O(L^2 \cdot 5 \cdot |\mathcal{T}_2|) = O(L^2 A^4)$ time and memory.

#### E.3 The principled formulation: three-factor model and MCMC

The bounded-edge truncations (0-or-1-edge in §E.1; 0-or-1-or-2-edge as a natural extension) are dynamic-programming approximations to a more principled underlying model whose joint posterior factorises cleanly into three independent ingredients: the alignment path $A$ under the TKF92 Pair HMM, a partition $\pi_M$ of its Match cells under a Chinese Restaurant Process (CRP) prior, and the per-block substitution likelihood under the substitution CTMC:

$$\pi(A, \pi_M \mid X, Y) \;\propto\; \pi_{\text{TKF92}}(A \mid X, Y) \;\cdot\; \pi_{\text{CRP}}(\pi_M \mid \alpha_z, |\text{Match}(A)|) \;\cdot\; \prod_{b \,\in\, \pi_M} P_{\text{block}}(b \mid t, H). \quad \text{(E.7)}$$

Blocks of size 1 (uninvolved Match cells) contribute the standard single-site CTMC factor $P_{\text{singlet}}$ of equation D.4 of Appendix D; blocks of size 2 (coupled-edge endpoints) contribute the joint Potts CTMC factor $P_{\text{doublet}}$ of equation D.5. The CRP prior with concentration $\alpha_z$ on a partition of $N = |\text{Match}(A)|$ Match cells into $K$ blocks of sizes $\{n_b\}_{b=1}^{K}$ is the canonical Ewens partition prior (15, 39)

$$\pi_{\text{CRP}}(\pi_M \mid \alpha_z, N) \;=\; \frac{\alpha_z^K \prod_{b=1}^{K}(n_b - 1)!}{\alpha_z(\alpha_z + 1)\cdots(\alpha_z + N - 1)}; \quad \text{(E.8)}$$

large $\alpha_z$ favours many small blocks (mostly singletons, few coupled pairs). In practice we restrict to blocks of size at most 2: blocks of size $\geq 3$ would correspond to triangular cliques whose marginal mass is suppressed by factors of $1/\alpha_z$ at high concentration. We call (E.7) the *infinite Pair HMM* in the sense that the number of coupled edges per alignment is not capped by the prior — in contrast to the bounded-edge dynamic-programming methods of §E.2.

The original *infinite Hidden Markov Model* of Beal, Ghahramani & Rasmussen (4) and the HDP-HMM of Teh, Jordan, Beal & Blei (48) provide the broader Bayesian-nonparametric framework in which equation E.7 sits.

##### MCMC sampler from the infinite Pair HMM

The factorisation (E.7) admits an efficient Markov-chain Monte Carlo sampler that scales as $O(L^4)$ for one-time setup followed by $O(L^3)$ amortised cost across many sweeps. We give the setup phase first, then the three move types that compose the sweep.

**Target distribution.** The sampler operates on the joint state $(A, E)$ where $A$ is a TKF92 Pair HMM alignment and $E$ is a set of unordered Match-cell-pair edges. The unnormalised target distribution is

$$\pi(A, E \mid X, Y) \;\propto\; P_{\text{baseline}}(A \mid X, Y) \cdot \frac{\alpha_z^{|E|}}{\prod_{m=1}^{N_M(A)}(m - 1 + \alpha_z)} \cdot \prod_{e \in E} M(e \mid t), \quad \text{(E.9)}$$

where $N_M(A)$ is the number of Match cells in $A$ and $M(e \mid t)$ is the four-residue Potts boost at edge $e$. The combinatorial prior is the canonical Ewens distribution restricted to blocks of size $\leq 2$.

**Setup phase.** The setup phase precomputes (a) the partial-Forward tensor $F^{\text{partial}}[i, j; k, l]$ giving the baseline TKF92 Pair HMM probability of any alignment passing through Match at both anchor cells $(i, j)$ and $(k, l)$ (the same $O(L^4)$ object enumerated by the $F_2$ inside–outside table of §E.1), and (b) the four-residue Potts boost tensor $M$ at the observed amino acids over all anchor quadruples. Both are stored once per $(X, Y)$ pair and reused across all sweeps and chains.

**The three MCMC moves.** The MCMC kernel composes three move types:

1. **Segment-resample move (MH).** Pick a contiguous fragment of the current alignment $A$ bounded by two anchor Match cells $a_L < a_R$ (or by the alignment endpoints). Propose $A^{\text{new}}$ by sampling the fragment from immediately after $a_L$ to immediately before $a_R$ by stochastic traceback on the segment-conditional Forward partition function $F^{\text{partial}}_{\text{segment}}[a_L; a_R]$ (Gibbs sample of the path conditional on the bounding anchors and the unchanged flanking alignment). Existing edges with endpoints in the resampled region are discarded; their orphaned partners outside the region become singletons in the partition. The proposal is exact under $\pi_{\text{TKF92}}$, so the alignment-likelihood factor of (E.9) cancels in the Hastings ratio, leaving only the CRP and boost factors.

2. **CRP add-edge move (MH).** Pick a Match cell of $A$ that is currently a singleton in $\pi_M$. Propose a partner Match cell from a proposal distribution $q_{\text{add}}(\cdot \mid \text{singleton position})$ (the natural choice weights candidate partners by their boost magnitude $|M - 1|$). Insert the new edge into $E$. The Hastings ratio combines the CRP prior ratio $\alpha_z/(m - 1 + \alpha_z)$, the boost factor $M(e \mid t)$, and the reverse-proposal ratio (which is symmetric for a uniform remove-edge proposal of the same edge cardinality).

3. **CRP remove-edge move (MH).** Pick an existing edge in $E$ uniformly. Remove it; both endpoints become singletons. The Hastings ratio is the reciprocal of the add-edge ratio.

**Replica exchange.** Parallel-tempering replica exchange over a ladder of $\alpha_z$ values (corresponding to varying coupling strengths) ensures global mixing across the multimodal posterior. Adjacent replicas swap states with the usual Metropolis acceptance ratio for the temperature-pair exchange.

**Cost.** The setup phase is $O(L^4)$ one-time per sequence pair. Each MCMC sweep does $O(1)$ inter-edge-anchor segment resamples; each segment resample is an $O(L_{\text{seg}})$ stochastic traceback through the cached partial-Forward tensor, giving an amortised $O(L)$ per-sweep cost. Aggregated per-cell match indicators across post-burn-in samples yield a stochastic estimate of the per-residue posterior $Q'_{ij}$ that the FSA assembler consumes in place of the bounded-edge analytic estimate. The block move reduces to a pure Gibbs "segment between adjacent edge anchors" resample whenever $a_L$ and $a_R$ are adjacent in the current edge-anchor list.

**Algorithmic detail**

**Algorithm 3** One MCMC sweep on the infinite Pair HMM

**Require:** Current state $(A, E)$, partial-Forward tensor $F^{\text{partial}}$, boost tensor $M$, concentration $\alpha_z$
**Ensure:** New state $(A', E')$

Pick $K_{\text{seg}}$ segment-resample positions (e.g. $K_{\text{seg}} = 8$, distributed by alignment length)
**for** each segment $[a_L, a_R]$ **do**
    $A^{\text{new}}_{\text{seg}} \leftarrow$ stochastic traceback of $F^{\text{partial}}_{\text{segment}}[a_L; a_R]$
    Discard edges in $E$ with at least one endpoint in $[a_L, a_R]$ (orphaned partners become singletons)
    $\rho \leftarrow$ MH ratio combining CRP and boost factors (alignment factor cancels by construction of the proposal)
    **if** $u \sim \text{Uniform}(0, 1) < \rho$ **then**
        $A \leftarrow A$ with $[a_L, a_R]$ replaced by $A^{\text{new}}_{\text{seg}}$
        $E \leftarrow E$ with orphaned edges removed
    **end if**
**end for**
Pick $K_{\text{add}}$ add-edge candidates and $K_{\text{rem}}$ remove-edge candidates (e.g. $K_{\text{add}} = 4|E|$, $K_{\text{rem}} = |E|$)
**for** each candidate **do**
    Propose under the appropriate move kernel
    Compute MH ratio (CRP $\times$ boost for add; reciprocal for remove)
    Accept / reject
**end for**
Periodically: attempt replica-exchange swap with adjacent $\alpha_z$ rung
**return** $(A, E)$

The exact form of the segment-resample MH ratio depends on the edge-handling convention. Under the simplest convention ("Strategy S-1" in the implementation spec), edges whose endpoints lie wholly within the resampled region are discarded unconditionally and their boost contributions enter only the forward MH ratio; edges with one endpoint outside the region are discarded and their orphaned partner becomes a singleton.

**Verification protocol**

The MCMC sampler is verified against four reference points:

**E.1 Cross-validate against `aug_phmm` (1-edge, large $\alpha_z$).** At large $\alpha_z$ the infinite Pair HMM truncates to a 0-or-1-edge model, and the per-cell match-marginal $Q'_{ij}$ produced by the MCMC sampler should agree with the exact $O(L^2A^2)$ aug-PHMM 1-edge inside–outside.

**E.2 Cross-validate against `aug_phmm_2edge` (2-edge, moderate $\alpha_z$).** At moderate $\alpha_z$ the truncation extends to 0-or-1-or-2-edge; agreement with the exact $O(L^2A^4)$ aug-PHMM 2-edge inside–outside is the next consistency check.

**E.3 Cross-validate against brute-force enumeration.** On small sequence pairs $(L_x, L_y) \leq (5, 4)$ where all alignments and all $\leq$ 5-edge placements can be enumerated directly, the MCMC sample mean of $Q'_{ij}$ must converge to the brute-force value at the canonical $1/\sqrt{n}$ rate.

**E.4 Detailed-balance sanity check (no-data case).** At $H = 0$ (so $M \equiv 1$) the boost factor is identically one, the per-block factor reduces to the baseline emission, and the sampler should recover the no-edge TKF92 Pair HMM marginal exactly — with edge-set sample sizes distributed as the prior Ewens distribution.

The first three checks pin the sampler to the exact reference at each of three orders of edge truncation; the fourth checks detailed balance independently of the data-conditioned target.

#### E.4 The conceptual hierarchy: infinite phylogenetic SCFG

**The infinite Pair HMM is itself an approximation.** The infinite Pair HMM of equation E.7 is the principled limit of the bounded-edge dynamic-programming approximations of §E.2, but it is itself an approximation to a still-more-principled object: the infinite-edge generalisation of the gravestone-augmented time-indexed pair SCFG of §D.5 of Appendix D. That fuller model samples not only the visible alignment $A$ and the edge set $E$ but also the timed gravestone-augmented branch history $G$ comprising every fragment that ever existed during the branch (whether visible at the alignment endpoints, deleted on the branch, or transiently inserted then deleted), with the same canonical CRP rule of equation E.7 governing edge spawn at every Match-class member of either an alive or a gravestone fragment. On the substitution side, transient gravestones inserted during the branch can be assigned the same $z_s$ key-DP class as their alive class-mates and therefore contribute coupling terms to those class-mates' rates during the gravestone's lifetime — so the substitution likelihood at a Match cell $(i, j)$ depends on $G$, not just on $A$. The infinite Pair HMM of this appendix marginalises $G$ implicitly via the standard TKF92 Pair HMM substitution emissions, which is correct in expectation when transient lineages are rare or when their coupling contribution is small, but loses the within-branch dynamics that the gravestone-augmented SCFG captures.

The resulting hierarchy is nested: the bounded-edge dynamic-programming methods of §E.2 are exact in the limit of the infinite Pair HMM of this section; the infinite Pair HMM is exact in the limit of the infinite gravestone-augmented pair SCFG; and that fuller object is the natural fixed point at which the indel-side and the substitution-side principled formulations meet. The MCMC sampler described above samples from the middle level of this hierarchy; lifting it to the full infinite gravestone-augmented pair SCFG — by interleaving the infinite-HMM CRP edge moves with the gravestone-augmented branch-history moves of §D.5 of Appendix D — is a natural extension.

**Further: the infinite gravestone-augmented *phylogenetic* SCFG.** The pairwise hierarchy above lifts to the full multi-branch phylogenetic setting at no further modelling cost, by composition with the Felsenstein-style upwards–downwards machinery of §D.5 that already glues per-branch gravestone-augmented histories into a tree-level posterior. At every internal node and along every branch of the tree, the gravestone-augmented latent state is sampled jointly with the alignment, while the same canonical CRP rule of equation E.7 governs edge spawn at every Match-class member of every fragment (alive or gravestone) at every node. The MCMC sampler above lifts by interleaving (i) per-branch gravestone-augmented history moves of §D.5, (ii) per-class-pair CRP edge moves of this appendix, and (iii) the per-internal-node Felsenstein-style updates already present in the SVI loop of §D.5. Each ingredient is already load-bearing in the existing pipeline; no new modelling assumptions are required to assemble them.

The full conceptual ladder is therefore:

mean-field pre-correction (deprecated; rem. D.13) $\rightarrow$ bounded-edge aug-PHMM (§E.2)
$\rightarrow$ infinite Pair HMM (§E.3) $\rightarrow$ infinite gravestone-augmented pair SCFG (§E.3 + §D.5)
$\rightarrow$ infinite gravestone-augmented *phylogenetic* SCFG (§E.4),

each rung exact in the limit of the next, the entire ladder falling out of the simple stick-breaking-TKF92-with-Potts construction of Appendices D–E with no further mathematical machinery.
