## Supplementary material for "Variational inference in coupled models of amino acid substitution": Combined manuscript + Supplemental Appendices A-E (from another version of this preprint: version 1, 2026-05-19)

### Insertions, deletions, and exchangeable couplings: a Dirichlet process over TKF92 domains and sites

Department of Bioengineering, University of California, Berkeley
{annabel_large, ihh}@berkeley.edu

**Abstract**

The TKF92 model of molecular evolution—a linear birth-death process for indels, with finite-state continuous-time Markov chain substitutions—is exchangeable in residue identity at every site: the generative process treats amino acids symmetrically, conditional on a single substitution rate matrix. To introduce local heterogeneity, evolutionary models are often equipped with site-class mixtures, preserving this symmetry in the sense of de Finetti: conditional on the latent class, residues are still exchangeable. In a four-step theoretical ladder, we show how long-range structure such as couplings between distant sites can also be introduced exchangeably by using a Dirichlet process to partition sites into co-evolving classes. Our first step is a thorough analysis of TKF92 to establish sufficient statistics, limiting behavior, and inferential tools. We then lift the pairwise TKF92 hidden Markov model, in the limit of small time, to a *time-indexed gravestone-augmented pair stochastic context-free grammar*, and from there to its phylogenetic generalisation. This framing allows trajectories to be sampled exactly by Inside-Outside recursion. The third step places a Dirichlet process over the alive sites and asks co-keyed sites to evolve under a sparse Potts interaction — an exchangeably-partitioned hidden direct-coupling model whose marginal alignment likelihood is unchanged from plain TKF92. The fourth rung of the ladder develops inference machinery: a Gibbs–Metropolis sampler that alternates alignment resamples, key-partition resamples, and stochastic parameter updates. We close several gaps along the way — exact closed-form sufficient statistics for the linear birth–death–immigration component, the resolvable L'Hôpital limit at $\lambda{=}\mu$, and a closed-form M-step for a recursive generalisation of TKF92 — and we report a 1,000-family PFAM fit with $K{=}4$ site classes whose Potts atoms carry $\sim$0.54 nats of covariation per class-pair on top of a substantial single-site substitution model. Supplementary material, including full source code for inference, may be found at https://tkfdp.net/.

#### 1 Exchangeability of indel models

The dominant frameworks of protein-sequence statistics are each limited in their own way. Substitution-only phylogenetic models (LG08, CAT, gamma-rates (24, 23, 37)) treat pre-aligned sites as independent given the tree; site–site epistasis is fitted away into an across-site rate-multiplier, not as a structural interaction. Direct-coupling analyses (plmDCA, mfDCA (25, 27, 9)) recover sharp contact predictions from joint amino-acid frequencies, but they treat the alignment as data and rely on multiple-sequence alignment tools whose own assumptions are broken by the same coevolution they want to detect. Indel birth-death models (TKF91/TKF92 (33, 34, 10) and the Poisson indel process (2)) sit somewhere between, with a cleanly separable framework for insertions, deletions, and point substitutions but no mechanism for coevolution between surviving sites.

**The full potential of exchangeability is going unused.** TKF92 is exchangeable in residue identity: the indel process treats amino acids symmetrically (insertions are drawn i.i.d. from a fixed distribution) and the substitution generator factorises over sites. A site-class mixture (23) breaks this exchangeability in a controlled way — two latent classes have two different stationary distributions, but the dynamics within a class are still identity-exchangeable. The Mixture-of-Domains generalisation of TKF92 (22) (MixDom) breaks it again by nesting geometric fragments inside a top-level domain HMM. Both elaborations preserve a key feature: *conditional on the latent class / domain / fragment, residues are still exchangeable.*

This is our starting point for indel-aware coevolution. Each newly inserted residue is assigned a random "key": shared keys indicate coupling partners. The key assignment is exchangeable in the sense of de Finetti: the key labels are latent, and the distribution over them is invariant to permutation of the sites. The same is true of the site-class labels that govern the single-site substitution dynamics. Once those labels are sampled, the substitution rates are determined symbolically, and indels are independent of all of them.

The contributions are organised as a four-rung ladder:

1. **Rung 1 (Section 2.1).** *Exchangeably covariant TKF92.* Closed-form BDI sufficient statistics and closed-form Baum–Welch M-steps for TKF91, TKF92, and the singly-recursive MixDom; the L'Hôpital limit at $\lambda=\mu$ closes the numerical singularity.

2. **Rung 2 (Section 2.2).** *Posterior over evolutionary trajectories.* Gravestone-augmented TKF92 in its infinitesimal-time limit is a continuous-time-indexed pair SCFG; its Volterra fixed-point is a bivariate Riccati on alive and gravestone counts, closed-form inside-outside, lifting to a *phylogenetic* SCFG via Felsenstein-style upwards–downwards.

3. **Rung 3 (Section 2.3).** *Exchangeable partitioning via Dirichlet processes.* A Chinese-restaurant draw partitions alive sites into co-keyed cliques; clique-mates share a Potts coupling on top of a per-class GTR generator, governed by three independent DPs (keys, classes, Potts atoms). The marginal alignment likelihood is unchanged from rung 1.

4. **Rung 4 (Section 2.4).** *Inference by variational EM and MCMC.* Gibbs-within-stochastic-VI: inner E-step for the clique CTMC; Jain–Neal sweeps over the three partition latents; recursive-traceback resample of the gravestone-augmented branch history; closed-form M-step into profile, Potts atom, and rate multiplier.

### 2 Methods at a glance

#### 2.1 Rung 1: closed-form TKF foundations

**The BDI process and its sufficient statistics.** TKF91 places one linear birth–death–immigration process on the sequence length, with insertion rate $\lambda$, deletion rate $\mu$, and immigration coincident with insertion. The endpoint-conditioned expected sufficient statistics — the birth count $B$, the death count $D$, and the time-integrated alive count $S$ — have a closed form via the Fisher score identity applied to the Kendall–Karlin–McGregor transition density (20). The sufficient statistics implicitly account for the gravestone lineages and other transient trajectory segments unobserved at the endpoints (Figure 1). Writing $p(j \mid i; t) = \mathbb{E}[\mathbf{1}\{N(t) = j\} \mid N(0) = i]$ for the BDI transition density, differentiating $\log p$ with respect to $\lambda$ and $\mu$ gives two score identities involving $\mathbb{E}[B]$, $\mathbb{E}[D]$, and $\mathbb{E}[S]$; the Karlin–McGregor count-balance $j - i = \mathbb{E}[B] - \mathbb{E}[D]$ closes the system. Solving:

$$\begin{aligned}
\mathbb{E}[S \mid i, j, t] \;&=\; \frac{(j-i) \;+\; \mu\,\partial_\mu \log p(j \mid i; t) \;-\; \lambda\,\partial_\lambda \log p(j \mid i; t) \;-\; \lambda t}{\lambda - \mu}, \\
\mathbb{E}[B \mid i, j, t] \;&=\; \lambda\,\partial_\lambda \log p(j \mid i; t) + \lambda\,\mathbb{E}[S \mid i, j, t] + \lambda t, \\
\mathbb{E}[D \mid i, j, t] \;&=\; \mu\,\partial_\mu \log p(j \mid i; t) + \mu\,\mathbb{E}[S \mid i, j, t].
\end{aligned} \tag{1}$$

The $\mathbb{E}[S]$ formula has a $0/0$ singularity at $\lambda = \mu$: the $(\lambda - \mu)$ denominator vanishes, and the numerator vanishes too as an automatic consequence of the same count-balance identity that closed the system. Earlier work (7, 12) patched the singularity with an $\varepsilon$-clamp; it is in fact a resolvable L'Hôpital limit recovered exactly by differentiating numerator and denominator a second time in $\lambda - \mu$, matching Gillespie simulations to Monte-Carlo precision at $10^7$ trajectories per regime (Figure 2). The same identity applied to the TKF92 fragment-level process supplies the closed-form sufficient statistics for the indel side of TKF92 and of MixDom.

**Closed-form Baum–Welch for TKF, including recursive variants.** With sufficient statistics in hand, the M-step for TKF91 and TKF92 is closed form: an exact quadratic equation in $(\lambda, \mu, r)$ from the per-cherry counts, plus a standard GTR update on the substitution side from bridge expectations (14). Recursive variants—such as the previously-described singly-recursive MixDom model of proteins (22) and fully-recursive TKF Structure Tree models of RNA structure (11), or other models of RNA, protein, and genome evolution described in the appendix—present a separate subtlety: their potentially-empty nested substructures require null state elimination by Schur complement. Naively, the M-step then needs second derivatives through the eliminated null blocks. We show how to restore eliminated counts exactly by a systematic backward elimination chain: the Forward–Backward (Inside–Outside) counts on the collapsed pair HMM (or pair SCFG) are pushed backward to per-null counts on the exploded model, and a single sum-by-parameter-group recovers unbiased BDI sufficient statistics for every parameter. The algebra is verified against autodiff at $10^{-6}$ relative tolerance for MixDom; the practical consequence is that rate-space Baum–Welch is exact at every level (top-level domain, per-domain fragment, per-fragment class, weights).

**Single-site substitution.** The substitution generator is the F81 instance of GTR, $Q^{(c)}_{xy} = \eta_s S_{xy} \pi^{(c)}(y)$, with a shared exchangeability matrix $S \in \mathbb{R}^{20\times 20}$, a per-class stationary distribution $\pi^{(c)} \in \Delta^{19}$, and a per-site gamma rate multiplier $\eta_s$. The F81 form is one of two natural reversible instances; we adopt it (over the symmetric Metropolis form) because it admits strict Dirichlet–multinomial conjugacy on $\pi^{(c)}$ via a secret-destination augmentation of the embedded random walk (Section 2.4). Class assignments are drawn from a Dirichlet process with concentration $\alpha_c$; the limit reproduces the CAT profile mixture (23) and, augmented with the $\eta_s$ multiplier, the Yang gamma-rates model (37).

#### 2.2 Rung 2: the time-indexed gravestone-augmented pair SCFG

**From pair HMM to time-indexed pair SCFG.** TKF91, viewed as a Markov process on sequence length and observed through a pair HMM at finite time $t$, has a continuous-time embedding that is not itself a pair HMM but a stochastic context-free grammar with productions indexed by time. Let $L(t)$ denote one link of age $t$; under TKF91 dynamics, $L(t)$ has lifetime $\tau \sim \mathrm{Exp}(\mu)$ and spawns child links at rate $\lambda$ during its lifetime. The branching structure is a continuous-time

Galton–Watson process expressible as

$$
\begin{aligned}
L(t) &\to A \cdot D(t,t) && \text{weight } e^{-\mu t}, \\
L(t) &\to G_\tau \cdot D(t,\tau) && \text{density } \mu e^{-\mu\tau}\,d\tau, \quad \tau \in (0,t), \\
D(t,T) &\to L(t-u) \cdot D(t,T) && \text{density } \lambda\,du, \quad u \in (0,T), \\
D(t,T) &\to \varepsilon && \text{weight } e^{-\lambda T}.
\end{aligned}
$$

The terminals $A$ and $G_\tau$ are an *alive* residue and a *gravestone* residue that died on the branch at relative time $\tau$; $D(t,T)$ generates a list of child links born during a window of size $T$ inside an observation interval of size $t$. The TKF92 expansion is mechanical: replace each single-residue terminal with a geometric fragment of shared death time, leaving the fragment-level branching grammar unchanged. The pair version adds an ancestor strand on the left of each $A/G_\tau$ terminal; insertions have no ancestor counterpart. The grammar parse tree is the continuous-time embedding of the TKF trajectory (Figure 1): each $L(t)$ non-terminal expansion corresponds to one branch of the Galton–Watson process, and the $A/G_\tau$ terminals are the alive and gravestone residues whose lifetimes the parse tree records.

**Why the augmentation matters: transient lineages.** Under any model with site–site coupling, an alignment-invisible fragment — born and deleted on the same branch — still changes the substitution dynamics of every site it shares a coupling partner with during its lifetime. The naive scheme of sampling only the alignment-visible indel events misses these transient lineages. The augmentation places them explicitly in the grammar's parse trees, as $G_\tau$ residues with finite lifetimes, so that the substitution likelihood can sum over them. The instantaneous alive count $N_a(t)$ is bounded pathwise by the visible-at-$T$ count plus the gravestone count, so the indel CTMC bridge has a finite state space and admits a closed-form recursive sampler.

**The bivariate Riccati.** Let $f_t(z,w) = \mathbb{E}[z^{N_a(t)} w^{N_g(t)}]$ be the joint generating function for alive count $N_a$ and gravestone count $N_g$. The Volterra fixed point

$$
f_t(z,w) \;=\; z\,e^{-\mu t}\,\exp\!\Big(\lambda\int_0^t (f_s(z,w)-1)\,ds\Big) \;+\; w\int_0^t \mu e^{-\mu\tau}\,\exp\!\Big(\lambda\int_{t-\tau}^t (f_s(z,w)-1)\,ds\Big)\,d\tau, \quad (2)
$$

is equivalent to the forward equation $\partial_t f_t = [\lambda z^2 - (\lambda+\mu)z + \mu w]\,\partial_z f_t$ with $f_0 = z$, a Riccati in $z$ whose flow closes algebraically. The Volterra fixed-point above is the Laplace transform of the Inside algorithm of stochastic context-free grammars (21, 30, 15): the SCFG production weights of the time-indexed grammar enter as their integrated kernels (survival $e^{-\mu t}$ and rate-density $\mu e^{-\mu\tau}\,d\tau$), and inside-outside in the $(z,w)$ generating-function domain is the fixed-point iteration of the Volterra equation. Defining $\Delta(w) = \sqrt{(\lambda-\mu)^2 + 4\lambda\mu(1-w)}$ and $z_\pm(w) = ((\lambda+\mu) \pm \Delta(w))/(2\lambda)$, the flow $(z(t)-z_+)/(z(t)-z_-) = (z(0)-z_+)/(z(0)-z_-) \cdot e^{-\Delta(w)t}$ inverts to give pointwise transition probabilities for $(N_a, N_g)$ at any branch length. Setting $w = 1$ recovers the standard linear birth–death PGF for $N_a$ alone.

**Phylogenetic lift.** The per-branch closed-form transition kernel from the Riccati composes across branches: the marginal at each internal node, conditional on the rest of the tree, is a closed-form pointwise distribution computable in $O(1)$ given the per-branch kernel evaluations. Felsenstein-style upwards–downwards then gives a tree-level closed-form sampler for the gravestone-augmented history — a *phylogenetic* SCFG of the same time-indexed form. Earlier work on pair SCFGs (32, 21, 15, 30) operates at fixed time; the present construction is the infinitesimal limit, and it lifts to trees without modification.

##### 2.3 Rung 3: Dirichlet-process partitioning and Potts couplings

**The TKF-DP model.** Fix TKF92 indel parameters $(\lambda, \mu, r)$, a key-class concentration $\alpha_z > 0$, a site-class concentration $\alpha_c > 0$, a Potts-atom concentration $\alpha_H > 0$, and base measures $G_0$ on $\pi^{(c)}$ (Dirichlet) and $G_0^H$ on entries of each Potts atom $H_h \in \mathbb{R}^{20\times 20}$ (Gaussian).

**Definition 1** (TKF-DP)**.** Sample stick-breaking weights for three independent Dirichlet processes (one over key classes, one over site classes, one over Potts atoms). Run TKF92 dynamics with parameters $(\lambda, \mu, r)$. At each fragment-birth event, draw $(z_s, c_s, \eta_s)$ from $(\mathrm{Cat}(\pi^z), \mathrm{Cat}(\pi^c), \mathrm{Gamma}(a_\eta, b_\eta))$ for each new residue; the labels are fixed for the residue's lifetime. At each first co-occurrence of an unordered class-pair $\{c, c'\}$, assign it a Potts-atom index $h_{cc'} \sim \mathrm{Cat}(\pi^H)$, materialising an atom $H_{h_{cc'}}$ from $G_0^H$ if new. The amino-acid trajectory on each key class $C$ is the single-site reversible CTMC with rate

$$Q^s(x \to x';\, x_{C\setminus s}) \;=\; \eta_s\, S_{xx'}\, \pi^{(c_s)}(x')\, \exp\Big(-\tfrac{1}{2} \sum_{s' \in C\setminus s} [\, H_{c_s c_{s'}}(x', x_{s'}) - H_{c_s c_{s'}}(x, x_{s'})\,]\Big),$$

which is reversible with respect to the joint Potts stationary

$$\pi_C(x_C) \propto \prod_{s\in C} \pi^{(c_s)}(x_s) \cdot \exp(-\sum_{\{s,s'\}\subset C} H_{c_s c_{s'}}(x_s, x_{s'})).$$

**Alignment–substitution factorisation is preserved.** The key DP draws are i.i.d. across sites at birth, independent of the alignment path and of the substitution history. The joint distribution therefore factorises as

$$P(\text{align}, \mathcal{P}, \{z_s\}, \text{subst}) \;=\; P_{\text{TKF}}(\text{align}) \cdot P(\mathcal{P} \mid L_A) \cdot \prod_s P(z_s \mid \mathcal{P}) \cdot P(\text{subst} \mid \text{align}, \{z_s\}), \quad (3)$$

where $L_A := |\text{Alive(align)}|$ is the number of alive sites the alignment exposes (for the pairwise object of Equation (5): the Match cells); the partition prior depends only on this count, not on alignment geometry. In particular: the marginal alignment likelihood is exactly TKF92's at every $\alpha_z$; the $\alpha_z = \infty$ all-singletons limit recovers TKF92 + per-site GTR pointwise. The coupling enters only through the substitution factor.

**Partition statistics and identifiability.** Marginalising $\mathcal{P}^z$, the partition on any finite set of $L$ sites is the Ewens partition with parameter $\alpha_z$, equivalent to a Chinese restaurant process draw. The expected total number of classes among $L$ sites is $O(\alpha_z \log L)$, with $k$-cliques suppressed by $1/\alpha_z^{k-1}$ relative to the all-singleton baseline; the prior is naturally sparse. The three DPs play distinct roles. $\alpha_c$ controls the number of materialised site classes; $\alpha_z$ controls the per-MSA partition into Potts cliques; and $\alpha_H$ controls identifiability of the Potts side. Without an atom DP, the free parameter count is $O(K_c^2 \cdot A^2)$ for $K_c$ site classes and $A$-state alphabet; with it, the count is $O(K_H \cdot A^2 + K_c^2)$ class-pair indices, where $K_H$ is data-driven and typically much smaller than $K_c^2$. Distinct class-pairs $\{c, c'\}, \{d, d'\}$ may share an atom index $h_{cc'} = h_{dd'}$; $K_H$ counts distinct atoms, not distinct class-pairs, and the atom DP's reinforcement is what drives this sharing. We use $K_H \le K_c(K_c + 1)/2$ truncated stick-breaking by default; at $K_H = K_c(K_c + 1)/2$ the truncation is tight (every class-pair could in principle have its own atom). Dirichlet processes have previously been explored to partition MSA columns into site classes and rows into functional classes (17, 16, 4, 28, 31). The CRP draws on key classes $\{z_s\}$ are the "lottery tickets" of Figure 1: each residue draws a key at birth, residues sharing a key are coupled, and the Ewens prior on the resulting partition is what selects which columns enter a Potts clique.

**Connection to direct-coupling analysis.** Direct-coupling analyses (25, 27, 9) fit a Potts model on the joint amino-acid frequencies of aligned columns of a single protein family. The TKF-DP is the indel-aware, phylogenetically-explicit, exchangeably-clustered generalisation: it conditions on a generative model of how aligned columns came to exist (TKF92 indels through time on a tree), it accumulates coevolutionary evidence across a corpus through a shared Potts-atom DP rather than per-family, and the columns it couples are drawn from an Ewens-distributed cluster prior rather than fitted as $O(L^2)$ explicit dyadic edges. The Wilburn–Eddy hidden Potts model (36), designed for remote homology search, is the closest single antecedent: it pairs a profile-HMM-style indel architecture with a Potts model over residue identities and makes the same architectural commitment that coevolution and indels must be scored jointly rather than as post-hoc rescoring.

#### 2.4 Rung 4: inference

The inference loop combines four ingredients in a stochastic variational EM scheme. We give each in one paragraph; full derivations are in the supplement.

**Inner variational E-step.** For each branch and each key DP class, the joint substitution dynamics on $\mathcal{A}^{|C|}$ has no usable eigendecomposition for $|C| > 1$, and direct matrix exponentiation is infeasible for $|C| > 2$. We approximate by the path-measure mean-field of Cohn et al. (6), restricted to a constant per-site rate matrix $\hat{Q}_b^s$ on each composition-constant segment. The geometric-mean fixed point $\log \hat{Q}_b^{s,*}(x \to x') = \log \eta_s + \log S_{xx'} + \log \pi^{(c_s)}(x') - \frac{1}{2} \sum_{s' \in C \setminus s} \mathbb{E}_{\bar{q}_b^{s'}}[H_{c_s c_{s'}}(x', x_{s'}) - H_{c_s c_{s'}}(x, x_{s'})]$ admits a closed-form ELBO via pairwise bridge expectations ($\mathrm{Br}(\alpha, \beta, t) = (e^{\alpha t} - e^{\beta t})/(\alpha - \beta)$ for $\alpha \neq \beta$, $= te^{\alpha t}$ otherwise). The constant-rate scheme is the $N = 1$ end of an adaptive refinement that converges to Cohn's full inhomogeneous family as $N \to \infty$ sub-segments; the gravestone augmentation of rung 2 supplies a forced segmentation at every composition-change event.

**MCMC over the discrete and augmentation latents.** Three combinatorial latents alternate. Augmented branch histories can be resampled by recursive midpoint traceback on the time-indexed pair SCFG of rung 2, with closed-form midpoint marginals from the bivariate Riccati and Felsenstein-style upwards–downwards across the tree. Key DP labels $z_s$ flip by Chinese-restaurant Gibbs with class-level ELBO ratios from the previous variational step; site classes $c_s$ flip by analogous CRP-Gibbs with single-site likelihoods; Potts-atom indices $h_{cc'}$ resample by truncated stick-breaking sweeps that share atoms across class-pairs. Jain–Neal split–merge proposals (18) on all three partitions provide global mixing. Pairwise alignment to the Infinite Pair HMM (1) uses a per-MSA block-resample on site partitions, with a setup-phase $O(L^4)$ precompute of the two-anchor partial-Forward tensor and per-sweep cost $O(L)$ (amortised: each sweep does $O(1)$ inter-edge-anchor segment resamples, each an $O(L_{\mathrm{seg}})$ stochastic traceback through the cached partial-Forward).

**Closed-form base-measure integrals.** The new-class predictives in the three CRPs require integrals against the base measures. Three structural conjugacies make them closed form: a *secret-destination augmentation* on $\pi^{(c)}$ (the F81 rate factors as an $S$-driven Poisson proposal clock with a $\pi^{(c)}$-driven Bernoulli filter, and silent-failure votes—including transitions back to source as an unobserved destination—complete the multinomial structure) gives Dirichlet–multinomial conjugacy on the per-class profile; a Gamma–Poisson conjugacy on $\eta_s$ gives closed-form Negative-Binomial site marginals; and a Gaussian-Laplace approximation around the path-DCA MAP gives closed-form Gaussian posteriors on each Potts atom, with a small multi-seed Laplace mixture for atoms with multimodal basins.

**Pairwise posteriors for decision-theoretic multiple alignment.** For downstream consensus-MSA assembly via sequence annealing (3), we expose a per-Match-cell posterior $Q'_{ij}$ derived from the joint-marginal alignment-and-partition target of Equation (5): an alignment $A$ together with a size-$\{1,2\}$ partition $E$ of its Match cells contributes a product of per-block factors — $P_{\text{singlet}}(x_i, y_j; t)$ at unpartnered Match cells and $P_{\text{doublet}}((x_i, x_k), (y_j, y_l); t)$ at coupled doublet endpoints — with class assignments and rate multipliers integrated analytically rather than re-marginalized at every column. The supplement details the per-block primitives, the augmented-Pair-HMM bounded-edge family ($O(L^2A^2)$ for 1-edge, $O(L^2A^4)$ for 2-edge), and the infinite-Pair-HMM MCMC sampler that targets the unbounded-edge limit.

**Counts-tensor training corpus.** Per-family int8 (cherry × column × AA-pair × branch-bin) count tensors decouple inference from MSA parsing, letting one SVI outer iteration scale to $10^3$–$10^4$ Pfam families.

### 3 Results

#### 3.1 Closed-form sufficient statistics match Gillespie simulation

The BDI sufficient-statistic formulae of Equation (1) agree with event-driven Gillespie simulation to within Monte-Carlo precision at $n_{\text{sim}} = 10^7$ trajectories per $(\lambda, \mu, t, i_0)$ regime. Figure 2 panels (a)–(c) show closed-form score-identity estimates against Gillespie sample means for the birth count $B$, death count $D$, and time-integrated alive count $S$ across a grid spanning $\kappa = \lambda/\mu \in \{< 1, 1, > 1\}$. The L'Hôpital limit at $\lambda = \mu$ has no $\varepsilon$-clamp degeneracy at the diagonal and matches Gillespie to Monte-Carlo precision. Lifting to the TKF91/TKF92 process and to the full forward–backward E-step, panel (d) shows the relative Gillespie residual scaling as the canonical $1/\sqrt{n}$ across three representative regimes, with 800 replicate seeds per $n_{\text{sim}}$ and no detectable bias offset — confirming both unbiasedness of the score-identity expressions and statistical correctness of the Gillespie simulator.

#### 3.2 Pfam training signals: covariation and rate heterogeneity

**Covariation signal.** A $K_c$=4, $K_H$=10 TKF-DP fit to 1,000 Pfam families (26) (indel rates pre-trained by Maraschino cherry-distillation (29) and fixed; 500 EM warmup iterations on the Dirichlet–multinomial class assignment, then 55 stochastic VI outer iterations with the full coupled loop; MSAs as the only supervision, no PDB) recovers a covariation signal visible directly in the stationary class-pair log-odds heatmap (Figure 3): a +2.51-bit Cys–Cys disulfide signature on the diagonal, a +1.66-bit off-diagonal Met–Tyr attraction consistent with the methionine-aromatic motif, and a band of Chou–Fasman-consistent proline aversions on the negative side.

**Indel and compositional heterogeneity.** Separately, a three-domain single-fragment MixDom model (`d3f1`) fit to the same corpus via SVI–BW with the exact closed-form chain-restoration M-step (Section 2.1; a Maraschino warm-start was tried but did not improve held-out LL) separates its three domains cleanly: one high-indel ($\lambda$≈0.11, $\mu$≈0.17, weight 0.32, mean fragment length 2.8) with the classic disordered / loop signature (enriched in S, P; depleted in W, C, M, H, Y; mean Kyte–Doolittle −0.82), and two low-indel domains (∼20× smaller rates, mean KD +0.07 and −0.22; enriched in L, V, I — the buried-core hydrophobic residues absent from the high-indel domain's enriched set). Indel rate tracks hydrophilicity in the direction biological intuition predicts

— mobile loops drift in length faster than buried cores — though most of the contrast is carried by the high-indel domain.

##### 3.3 Pairwise alignment accuracy on structural benchmarks

**Evaluation by expected pair-confusion.** We score pairwise alignment accuracy through the soft confusion matrix on the residue-pair cell grid: for each pair of sequences in a reference MSA guided by experimentally-determined structures, we read off the gold match set $T = \{(r_i, r_j)\}$ from the curated alignment and compute, against the model's $L_i \times L_j$ match-posterior matrix $P$, the sufficient statistics

$$e_{\mathrm{TP}} = \sum_{(r_i,r_j)\in T} P[r_i, r_j], \qquad \text{total mass} = \sum_{r_i,r_j} P[r_i, r_j] = e_{\mathrm{TP}} + e_{\mathrm{FP}}, \qquad |T| = e_{\mathrm{TP}} + e_{\mathrm{FN}}. \quad (4)$$

These four numbers are additive across pairs and across families and suffice to compute soft precision, recall, and expected pair F1. The measurement is exact given the posterior; no Viterbi decoding is required, and no MSA reconstruction is performed.

**BAliBase results.** Table 1 summarises pairwise expected-$F_1$ results on the 22-family BAliBase BAli3pdbm $L < 150$ subset (187 sequence pairs) for the canonical pipeline: TKF92 baseline (single-component, corpus-fitted indel rates), TKF92-$K$=20 indel mixture, CherryML-$C$=20 per-site substitution-class mixture (single TKF92, marginalised over $C$=20 emission components), MixDom-d3f1, external aligners MAFFT (19) and MUSCLE (8), and the infinite-Pair-HMM (TKF-DP $K$=4 Potts-coupled) MCMC sampler. The MixDom-d3f1 model ($D = 3$ domains, $F = 1$ fragment per domain) is amongst the smallest MixDom models that meaningfully improve over plain TKF92 on this benchmark; adding fragments or classes beyond this scale gives diminishing returns. The infinite-PHMM-$K$=4 sampler uses the released checkpoint trained on ∼1,000 Pfam families with EM warmup (Section 3.2); its column entries are the running-mean cell posterior $Q'$ from the MCMC sampler (the 22-family $L < 150$ subset is the practical eligibility ceiling for the $O(L^4)$ partial-Forward precompute on an 11 GiB GPU; see Section 3.2 for the calibration). The TKF-DP coevolutionary coupling is weak in the pairwise case, and appears to sharpen alignment accuracy only slightly, with gains lost by the sequence-annealing algorithm for MSA assembly (which does not, at present, share coupling signal across MSA columns, unlike the composite-likelihood method discussed in Section 3.4 and Figure 5).

##### 3.4 The infinite Pair HMM as the natural fixed point

The methods of Table 1 together span the $|E| = 0$ truncation (the soft-DP rows, which factor over Match cells and ignore Potts coupling between sites) and the $|E| \to \infty$ limit (the infinite-Pair-HMM sampler row, which couples sites via the size-$\{1, 2\}$ Ewens prior at the trained $\alpha_z$); they are progressively faithful approximations to a single underlying object: the alignment-and-partition posterior

$$\pi(A, \mathcal{P}_M \mid X, Y) \propto \pi_{\mathrm{TKF92}}(A \mid X, Y) \cdot \pi_{\mathrm{CRP}}(\mathcal{P}_M \mid \alpha_z, |\mathrm{Match}(A)|) \cdot \prod_{b\in\mathcal{P}_M} P_{\mathrm{block}}(b \mid t, H), \quad (5)$$

where $\mathcal{P}_M$ partitions the Match cells into singletons (block of size 1, contributing the standard per-site CTMC factor) and pairs (block of size 2, contributing the joint Potts CTMC factor), with the size-$\{1, 2\}$ truncation of the Ewens prior. The factorisation makes the alignment, partition, and

| **Method** | $E_{\text{pairs}}(Q')$ $F_1$ | $e_{\text{TP}}$ | $E_{\text{pairs}}$(opt-acc) $F_1$ | $e_{\text{TP}}$ | $E_{\text{pairs}}$ in MSA $F_1$ | MSA score SP | TC |
|---|---|---|---|---|---|---|---|
| TKF92 | 0.450 | 4839 | 0.475 | 5202 | 0.514 | 0.724 | 0.631 |
| MixDom-d3f1 | 0.462 | 4946 | 0.489 | 5372 | 0.513 | 0.726 | 0.638 |
| TKF92-$K$=20 | 0.458 | 4921 | 0.486 | 5324 | 0.515 | 0.732 | 0.637 |
| CherryML-$C$=20 | 0.457 | 4893 | 0.482 | 5270 | 0.515 | 0.723 | 0.629 |
| inf-PHMM-$K$=4 (RE)* | 0.450 | 4908 | 0.475 | 5232 | 0.509 | 0.705 | 0.609 |
| MAFFT (FFT-NS-2) | — | — | — | — | 0.509 | 0.691 | 0.558 |
| MUSCLE (default) | — | — | — | — | 0.512 | 0.760 | 0.662 |

Table 1: **Alignment accuracy on the 22-family BAliBase BAli3pdbm L < 150 subset (187 pairs).** The L < 150 ceiling reflects the practical memory headroom for the infinite-Pair-HMM $O(L^4)$ sampler on an 11 GiB GPU (max-seq-len < 150; see Section 3.2). $E_{\text{pairs}}(Q')$ columns are the raw posterior pool; $E_{\text{pairs}}$(opt-acc) is decision-theoretic optimal-accuracy DP applied to the same posterior (13); $E_{\text{pairs}}$ in MSA is the cell-indicator pool from the downstream FSA-merged MSA. * The infinite-PHMM-$K$=4 row uses the released `K4-emwarm-top1000-2026-05-09` TKF-DP checkpoint ($K_c$=4 classes, $K_H$=10 Potts atoms), with $\alpha_z$=100 and corpus-fitted indel rates ($\lambda$=0.04581, $\mu$=0.04680, $r$=0.6835), under replica-exchange (6-rung $\alpha_z$ ladder $\{100, 178, 316, 562, 10^3, 5\times10^3\}$, swap every 10 sweeps; 8000 sweeps + 2000 burn-in per rung; 2 RE replicates per pair).

| **Method** | $E_{\text{pairs}}(Q')$ $F_1$ | $e_{\text{TP}}$ | $E_{\text{pairs}}$(opt-acc) $F_1$ | $e_{\text{TP}}$ | $E_{\text{pairs}}$ in MSA $F_1$ | MSA score SP | TC |
|---|---|---|---|---|---|---|---|
| TKF92 | 0.289 | 77,284 | 0.309 | 84,913 | 0.377 | 0.776 | 0.622 |
| MixDom-d3f1 | 0.316 | 81,324 | 0.330 | 89,284 | 0.384 | 0.807 | 0.662 |
| TKF92-$K$=20 | 0.292 | 78,140 | 0.313 | 85,940 | 0.378 | 0.784 | 0.633 |
| CherryML-$C$=20 | 0.297 | 78,633 | 0.315 | 86,220 | 0.384 | 0.791 | 0.645 |
| MAFFT (FFT-NS-2) | — | — | — | — | 0.365 | 0.797 | 0.653 |
| MUSCLE (default) | — | — | — | — | 0.373 | 0.841 | 0.715 |

Table 2: **Alignment accuracy on the full BAliBase bali3pdbm corpus (120 families, all sequence lengths).** Same methods and column definitions as Table 1 (the $L < 150$ subset), except the infinite-Pair-HMM-$K$=4 row is omitted due to GPU memory constraints on the $O(L^4)$ partial-Forward precompute.

substitution factors independent given each other; the block-resample MCMC kernel of Section 2.4 samples from Equation (5) exactly. Equation (5) is itself the principled single-branch marginal of the infinite gravestone-augmented phylogenetic SCFG (rung 2 of the ladder): in the full multi-branch object, the same CRP rule governs edge spawn at every site of every fragment — alive or gravestone — at every internal node, with the substitution likelihood depending on the full augmented history rather than only the alignment-visible fragments (Figure 1).

**Worked example: divergent disulfide-rich pair from a held-out family (Figures 4 and 5).** We ran the canonical infinite Pair HMM sampler on a single sequence pair from human **LAMA1** (laminin subunit $\alpha$-1, UniProt `P25391`), a member of the PFAM Laminin EGF-like family `PF00053` (*held out from training*): two EGF-like repeats of LAMA1, residues 1452–1506 and 1090–1147, with $L_x$=55, $L_y$=58, $\tau$=2.15 and only 21% sequence identity — diverged enough that the TKF92 baseline alignment is no longer near-deterministic. The joint sampler concentrates posterior mass at Cys–Cys cells at ~17× the background rate on the X axis and ~12× on Y, against ~5× for the single-sequence baselines (Figure 4). The per-pair coupling bonus is weak but detectable; pooled across the 72-sequence family MSA, the composite-likelihood posterior *perfectly* rank-separates all 28 Cys–Cys cell-pairs from the non-Cys background (Figure 5).

### 4 Discussion

TKF-DP introduces exchangeable *nonlocal* coupling structure — DP-clustered Potts interactions between residues that may sit anywhere on the sequence — on top of a single TKF92 indel backbone. The companion MixDom construction (22) and its recursive cousins introduce a different, *local* kind of structure, positional and sequence-ordered, with composition and indel rate varying across a small mixture of latent domains. A realistic generative model of protein evolution should have both, with couplings restricted to within a domain or to specific local contexts: buried-core residues in a globular protein, base-paired residues in an RNA stem, or residues sharing a secondary-structure element. Combining nonlocally- and locally-structured models is a larger technical challenge but well within reach: the recursive grammars of Appendix C provide the natural scaffolding for within-domain or within-element coupling structure.

We made several approximations in our first evaluation of the TKF-DP. Two are worth calling out in detail. (1) We truncated the Ewens prior of Equation (5) to size-$\{1, 2\}$ components; larger Potts components admit an independent-coupling lower bound ($N$=1 in the midpoint refinement of Appendix D) that extends to a formal ELBO by mesh-refining the midpoint construction, recovering the Cohn fixed point as midpoint spacing tends to zero. (2) We used the standard simplification of treating substitution as conditional on the alignment-visible fragments alone; a more careful treatment would account for the substitution effect on alive sites of transient ghost / gravestone lineages during their lifetimes. We conjecture that this effect can be absorbed into a mean-field correction using the expected trajectory sufficient statistics of Section 2.2, with a heavier importance-sampling treatment available via MCMC from the trajectory SCFG itself.

The hierarchical-DP schema we present here — one DP each for site classes, Potts-atom indices, and key classes — is one design among many; alternative atom-sharing structures, hierarchical class groupings, or supervised training schemes remain unexplored. Further, our use of the DP to induce coupling is transferable to other indel models. For example, the Poisson Indel Process (2) yields a column-independent PIP-DP variant while preserving the DP latent structure.

### Acknowledgements

We thank Jeff Thorne, Sean Eddy, Jotun Hein, Debora Marks, Yun Song, Marc Suchard, Gerton Lunter, Nick Goldman, Nicola De Maio, and Elena Rivas for helpful discussions. Selected mathematical derivations and exposition were generated by Anthropic's Claude under the supervision, and following the intuition, of the human authors. The work was funded by NIH grants HG004483, GM080203, and HG013117.

### Author contributions

I.H. led theory and software development, experimental evaluation, and writing. A.L. developed independent JAX implementations of the mixture-of-sites, -fragments, and -domains generalisations of TKF92 with Adam-based fitting pipelines. Both authors approved the text.

#### Appendix: roadmap

The technical body of this work — BDI sufficient statistics, exact closed-form Baum–Welch for TKF91/TKF92/MixDom, Maraschino cherry-distillation, the time-indexed gravestone-augmented pair SCFG, the TKF-DP generative graph and inference loop, the infinite Pair HMM MCMC kernel, and the bounded-edge alignment postprocessing family — is collected into five appendices whose contents are largely quoted verbatim from the two companion manuscripts. The roadmap is:

**Appendix A.** BDI and TKF foundations. The finite-state CTMC and linear birth–death–immigration process, the closed-form bridge-expectation and Fisher-score sufficient statistics for TKF91 and TKF92, the singlet-division WFST derivation, the L'Hôpital limits at $\lambda{=}\mu$, and a discussion of how TKF92 relates to the latent-boundary-free General Geometric Indel (GGI) model.

**Appendix B.** EM, composite likelihoods, and variational inference. Closed-form substitution M-steps for the many GTR specialisations; the stochastic-variational Baum–Welch loop for TKF91/TKF92 with its ELBO derivation, natural-gradient updates, and convergence theorem; Maraschino, the TKF92 cherry-distilled generalisation of CherryML, and the tree-level inference algorithms it composes with (FSA, BeamASR, VarAnc, svi-VarAnc); and a structural-bias analysis of the BP cumulant under a column-factorised variational field.

**Appendix C.** Recursive TKF. The hierarchical-mixture-of-domains generalisation of TKF92 (MixDom), its exact closed-form M-step via six-step chain-restoration through the fully exploded null-state model, the order-1 Maraschino distillation, the algebraic-distillation family, the SVI-BW convergence considerations specific to MixDom, the tree-level VEM and ancestral-reconstruction algorithms, the generalised phylo-HMM, the labeled-MixDom Singlet and WFST, and the recursive-grammar-elaboration rules that lift these constructions to arbitrary nested-TKF models.

**Appendix D.** TKF-DP: the Dirichlet-process Potts coupling. The generative model of Definition 1, the three-DP latent structure, the class-level path-measure factorisation and constant-rate variational ELBO via pairwise bridge expectations, the augmented indel histories via the time-indexed pair SCFG, the SVI loop, and the pairwise alignment postprocessing landscape (joint-marginal target, coupled annealer, augmented-Pair-HMM family).

**Appendix E.** The infinite Pair HMM and its block-resample MCMC sampler. The three-factor factorisation of Equation (5); the setup-phase $O(L^4)$ partial-Forward precompute; the per-sweep $O(L)$ block-resample kernel; and the projected lift to the full gravestone-augmented phylogenetic SCFG.

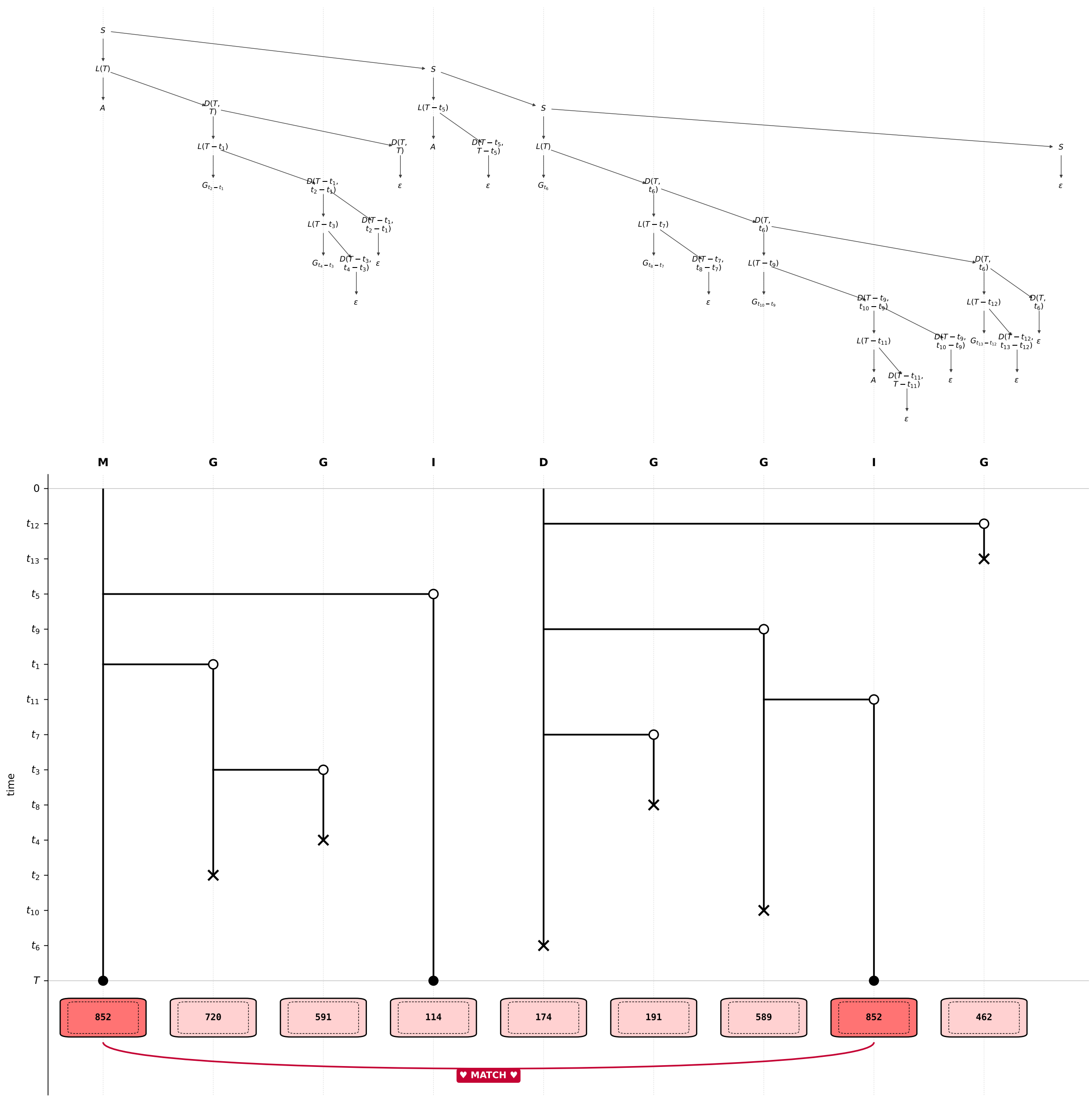
M
G
G
I
D
G
G
I
G
time
0
$t_{12}$
$t_{13}$
$t_5$
$t_9$
$t_1$
$t_{11}$
$t_7$
$t_3$
$t_8$
$t_4$
$t_2$
$t_{10}$
$t_6$
$T$
852
720
591
114
174
191
589
852
462
♥ MATCH ♥

Figure 1: **TKF91 indel trajectory as a Galton–Watson branching process and its gravestone-augmented SCFG parse tree.** *Top:* the gravestone-augmented SCFG parse tree, with each non-terminal annotated by its symbolic time arguments $(t, T)$ in terms of the event indices $t_1, \ldots, t_{13}$ that label the middle panel's vertical axis. Each $L(\cdot)$ leaf-terminal ($A$ for surviving lineages, $G_\tau$ for transient ones) sits directly above its corresponding trajectory column; the faint dotted vertical guides carry the eye from each parse-tree terminal down through the trajectory column and into its lottery ticket. Epsilon leaves ($\varepsilon$) terminate $D$-recursion and $S$-recursion and are placed mid-way between their column-anchored neighbours. The event labels $t_k$ are assigned in fixed pre-order traversal of the tree, so the parse-tree subscripts are deterministic; the random topological sort over event times permutes only the vertical axis ordering of the middle panel. *Middle:* one realisation of the continuous-time TKF91 process on a branch of length $T$. Columns are residue lineages, time runs downward from 0 to $T$. Open circles ($\circ$) mark birth events; filled circles ($\bullet$) mark survival to $T$; crosses ($\times$) mark death events. The ancestral marker M (col. 1) survives to $T$ and spawns a transient insert G (col. 2; lifetime $t_1 \to t_2$), which itself spawns a deeper transient G (col. 3; $t_3 \to t_4$). The insertion I (col. 4) is born at $t_5$ and survives. The ancestral site D (col. 5) dies at $t_6$ after spawning three transient inserts (cols. 6, 7, 9; lifetimes $t_7 \to t_8$, $t_9 \to t_{10}$, $t_{12} \to t_{13}$); the second of these transients itself spawns the surviving insertion I (col. 8) at $t_{11}$. The horizontal arrows depict the SCFG branching production $D(t, T) \to L(t - u) \cdot D(t, T)$ firing at each child link's birth; the immortal link (dashed) anchors the leftmost lineage. *Bottom:* a row of red lottery tickets cartooning the TKF-DP Potts-coupling layer: each surviving column draws a random integer "key" (here a three-digit number) from the Chinese restaurant process partition. Most keys are unique, but two distant alive columns happen to share a key (the MATCH), which is precisely the event in which two sites end up at the same CRP table and acquire a Potts coupling under the K=4 model of Section 2.3.

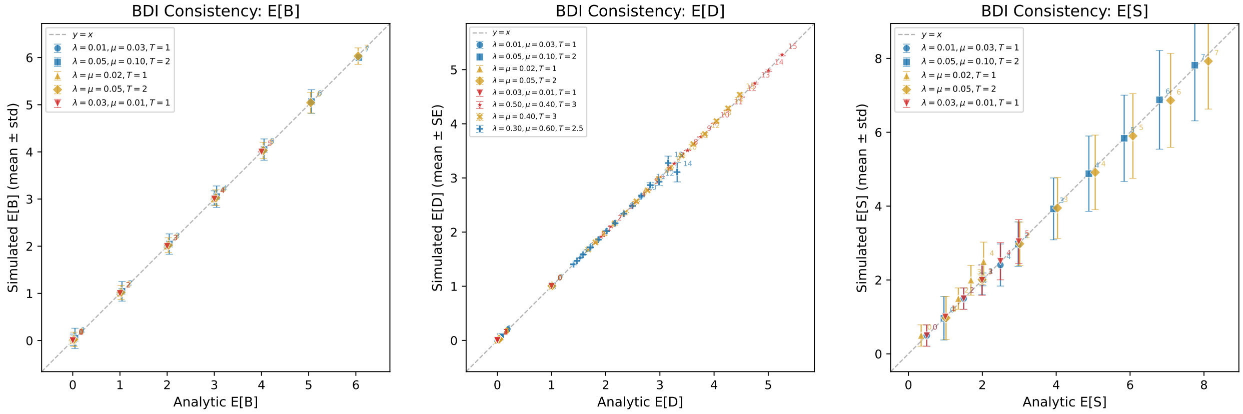

TKF BDI Gillespie vs analytic moments (n_sims = 20000000). Diamonds: L'Hôpital regime (λ ≈ μ).

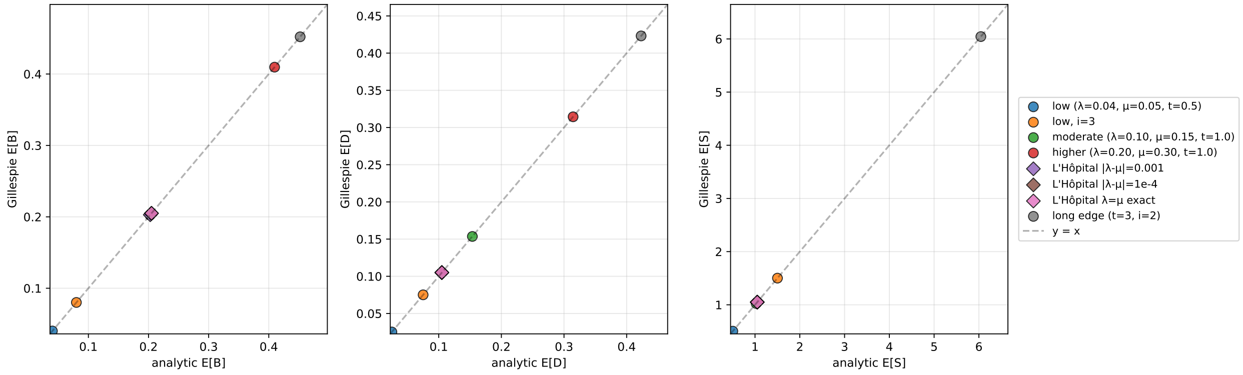

Gillespie BDI residuals: mean ± SE across 800 replicate runs per n_sims. Curves track 1/√n with no systematic offset.

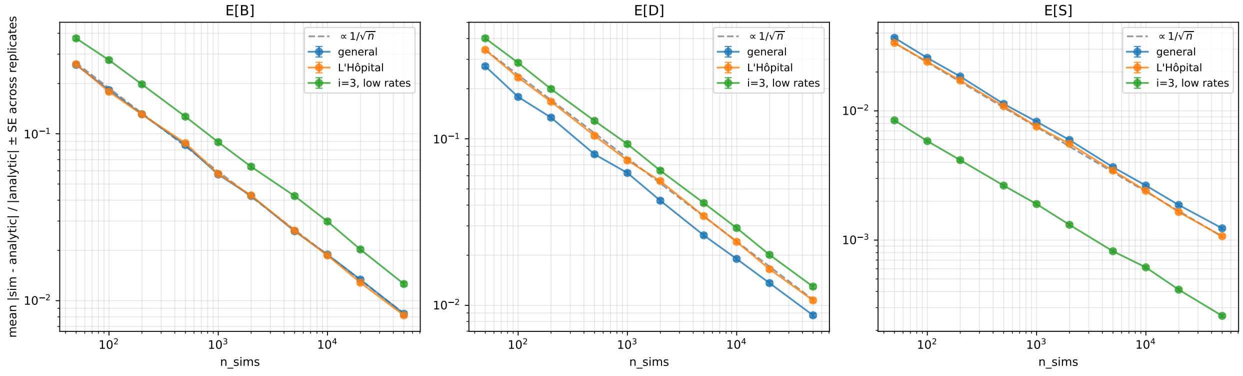

Figure 2: **BDI / TKF sufficient-statistic validation.** Top row, panels (a)–(c): closed-form score-identity expectations (analytic) plotted against Gillespie sample means (simulation, $10^7$ trajectories per regime at low rates; $10^6$ trajectories at higher rates) for births $B$ (left), deaths $D$ (middle), and time-integrated alive count $S$ (right). Each point is one $(\lambda, \mu, t, i_0)$ regime; the identity-line scatter is below plotting precision across $\kappa < 1$, $\kappa = 1$ (L'Hôpital diamonds), and $\kappa > 1$. The five low-rate regimes ($\lambda, \mu \sim 0.01$–$0.10$, $t \sim 1$) sweep $\mathbb{E}[D]$ over $\sim$0–1 (most trajectories see at most one death); three additional higher-rate regimes ($\lambda, \mu \sim 0.3$–$0.6$, $t \sim 2.5$–$3$) extend the D panel through $\mathbb{E}[D] \approx 5$, exercising the score identity across an order of magnitude in event count without bias. Numerals beside each point are the descendant count $j$ that bin represents; residual scatter at the largest $j$ reflects per-bin MC noise that shrinks as $1/\sqrt{n_{\text{bin}}}$. Error bars are standard errors of the mean (SE $= \sigma_{\text{sim}}/\sqrt{n_{\text{bin}}}$). Panel (d): TKF91/TKF92 BDI moments via Gillespie ($2 \times 10^7$ sims per regime, 5 regimes) against the closed-form expressions; tight identity-line. Panel (e): relative Gillespie residual scaling as $1/\sqrt{n_{\text{sim}}}$ over 800 replicate seeds per $n_{\text{sim}}$, three representative regimes (general, L'Hôpital, low-rate $i_0$=3); no detectable offset.

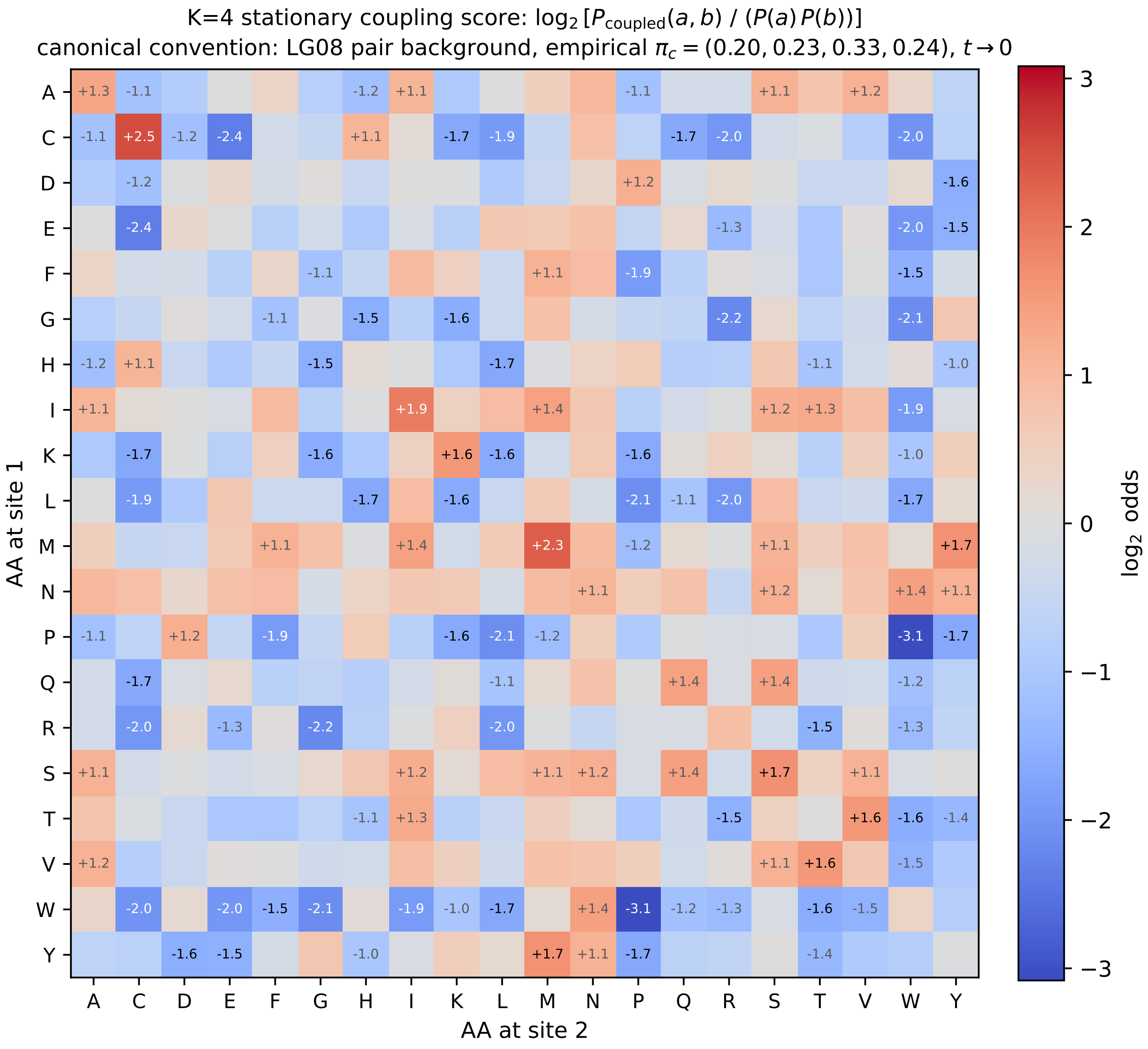
K=4 stationary coupling score: $\log_2 [P_{\text{coupled}}(a,b) \,/\, (P(a)\,P(b))]$
canonical convention: LG08 pair background, empirical $\pi_c = (0.20, 0.23, 0.33, 0.24)$, $t \to 0$
AA at site 1
AA at site 2
$\log_2$ odds

Figure 3: **Stationary coupling log-odds at a coupled site-pair in the K=4 TKF-DP model.** Cell $(a, b)$ shows $\log_2[\pi_{\text{joint}}(a, b)/(\pi_{\text{singlet}}(a)\,\pi_{\text{singlet}}(b))]$ in the $t \to 0$ limit, under the canonical convention (LG08 pair-stationary background + empirical class prior $\pi_c = (0.20, 0.23, 0.33, 0.24)$ from SVI training-set column assignments). *Red* cells are enriched at coupled sites relative to the singlet product (positive log-odds); *blue* cells are depleted. The strongest positive cells are C–C (+2.51 bits; the disulfide-bridge signature: cysteine pairs co-occur far more often at structurally coupled positions than background frequencies predict), M–M (+2.32), and I–I (+1.93); the strongest off-diagonal positive is M–Y (+1.66), consistent with the methionine–aromatic (sulfur–$\pi$) stabilising interaction documented in folded structures (35). The strongest negative cells include several pairs whose Chou–Fasman secondary-structure propensities are antagonistic — P–W (−3.08), L–P (−2.12), K–P (−1.63) all pair the helix/strand breaker proline with strong helix or strand formers (5) — though the pattern is not uniformly explained by secondary-structure preferences alone: top empirical negatives also include C–E (−2.36), G–R (−2.21), and a Trp column with broad aversion that the low Trp background frequency ($\pi(\text{W}) \approx 0.011$ in LG08) likely inflates. Mean per-class-pair MI under this convention is ∼0.54 nats / 0.78 bits (atom-averaged).

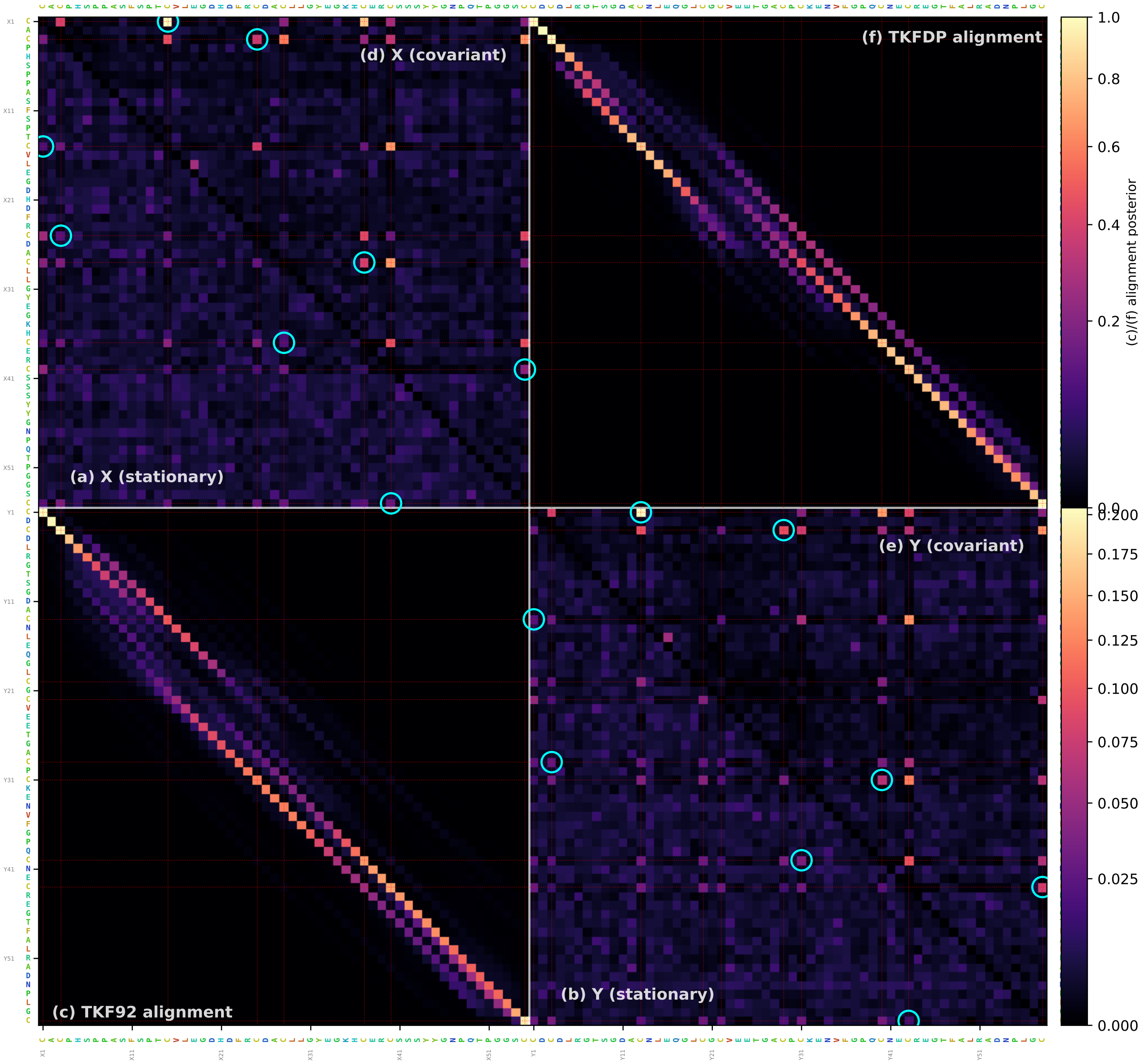
(d) X (covariant)
(f) TKFDP alignment
(a) X (stationary)
(e) Y (covariant)
(c) TKF92 alignment
(b) Y (stationary)
(c)/(f) alignment posterior

Figure 4: **Coevolutionary signal boosts Cys–Cys disulfide bond identification on a sequence pair from a held-out test family.** Sequences X, Y are two EGF-like repeats of human **LAMA1** (laminin subunit $\alpha$-1, UniProt `P25391`; PFAM Laminin EGF-like family `PF00053`, held out from K=4 training). X is LAMA1 residues 1452–1506 ($L_x$=55, 8 Cys forming 4 UniProt-annotated disulfides (circled) at seq-pos $(1, 15), (3, 25), (28, 37), (40, 55)$); Y is the LAMA1 repeat two positions earlier, residues 1090–1147 ($L_y$=58, 10 Cys: the same 4 disulfides at seq-pos $(1, 13), (3, 29), (31, 40), (43, 58)$ plus 2 *free* Cys at LAMA1 positions 1109 and 1111 not in any annotated disulfide). *Panels (c, f):* TKF92 baseline alignment posterior (lower-left, lower-right) vs the TKF-DP alignment posterior after coupled-edge resampling (upper-right); the diagonal is sharpened by the coupling boost on conserved disulfide cells. *Panels (a, b):* single-sequence X/Y edge posteriors (no homolog evidence; Cys–Cys cells receive only the per-pair model boost and the matching-constraint prior). *Panels (d, e):* joint sampler edge posteriors at the same Cys positions, with the homolog providing positional discrimination across the diverged alignment. The joint sampler concentrates posterior mass at Cys–Cys cells at $\sim$17$\times$ the background rate on the X axis (0.074 vs 0.004) and $\sim$12$\times$ on Y (0.047 vs 0.004); the single-sequence baselines give only $\sim$5$\times$ each. Sampler: joint pair replica-exchange ladder $\alpha_z \in \{100, 250, 700, 2000, 10^4\}$, 8000 sweeps + 1500 burn-in; single-sequence ladder $\alpha_z \in \{100, 500, 2000, 10^4\}$, 10000 sweeps + 1000 burn-in.

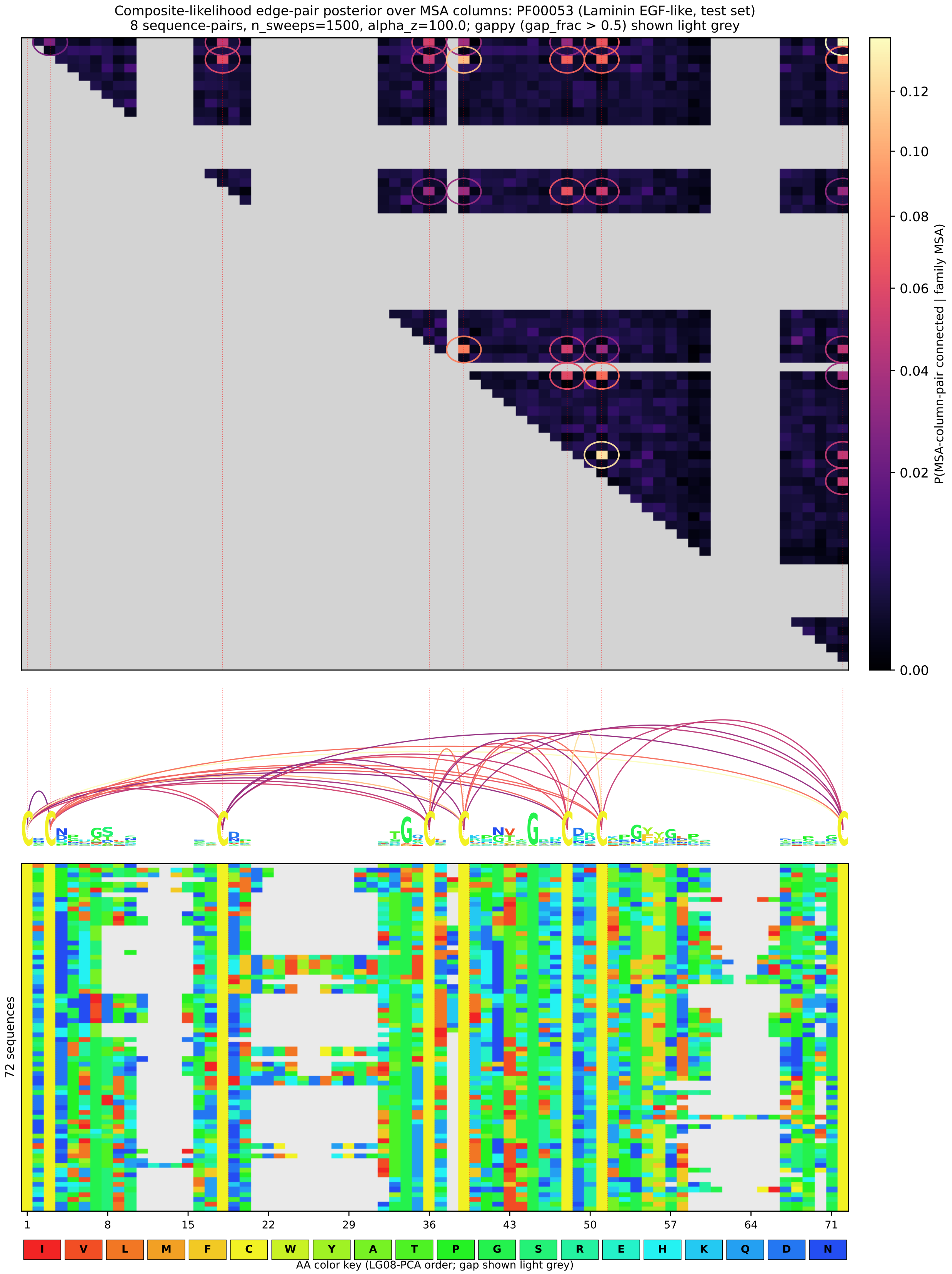

Composite-likelihood edge-pair posterior over MSA columns: PF00053 (Laminin EGF-like, test set)
8 sequence-pairs, n_sweeps=1500, alpha_z=100.0; gappy (gap_frac > 0.5) shown light grey
P(MSA-column-pair connected | family MSA)
0.12
0.10
0.08
0.06
0.04
0.02
0.00
72 sequences
1
8
15
22
29
36
43
50
57
64
71
I
V
L
M
F
C
W
Y
A
T
P
G
S
R
E
H
K
Q
D
N
AA color key (LG08-PCA order; gap shown light grey)

Figure 5: **Composite-likelihood MSA-column edge-pair posterior for the held-out PF00053 (Laminin EGF-like) family.** Layout from Holmes (11) Fig. 14. *Top:* triangular heatmap $P(\text{cols } (k_1, k_2) \text{ connected} \mid \text{family MSA})$ aggregated over 8 sequence pairs by the infinite Pair HMM sampler at the canonical convention. Gappy columns (gap-fraction $> 0.5$) are masked light grey. *Bottom:* the 72-sequence PF00053 MSA coloured by AA chemistry (gap = light grey); red arcs connect the top-scoring column pairs in the triangle above. Cys columns (yellow) at MSA positions $\{1, 3, 18, 36, 39, 48, 51, 72\}$ — the canonical Laminin EGF-like 8-cysteine pattern. The signal is striking: the chain achieves *perfect rank-separation* of Cys–Cys cells from background. With 8 Cys columns there are $\binom{8}{2} = 28$ candidate Cys–Cys pairs, and the top-28 cells in the composite posterior are exactly those 28 pairs — every single Cys–Cys pair ranks above every non-Cys cell. Notably, this perfect rank-separation is a property of the *multi-pair composite*; the underlying single-pair posteriors do not achieve it. On the single LAMA1 pair of Figure 4, the minimum-$N$ to cover all Cys–Cys cells is $N$=124 for panel (a), $N$=291 for (b), $N$=1402 for (d), and $N$=1500 for (e) — far above the perfect $N$=28 / $N$=45 that the respective C-Cys counts would allow. The composite likelihood averages out per-pair alignment noise; only consistent across-pair Cys–Cys signal survives. Mean Cys–Cys posterior $\langle P \rangle = 0.060$ vs 0.0037 at non-Cys non-gappy background ($\sim 16\times$ enrichment) — identical ratio to the single-pair LAMA1 figure (Figure 4). The chain distributes mass across all 28 candidate Cys–Cys pairs without discriminating the 4 *true* disulfide bonds from the other 24. This is the model's expected behaviour: the per-pair coupling boost $M_{\text{solo}}[a, c]$ is position-agnostic (it depends only on the AA pair $(a, c) = (C, C)$, not on the columns), so additional discrimination would have to come from phylogenetic conservation patterns the present composite-likelihood does not yet exploit. Sampler: 1500 sweeps + 300 burn-in per pair, 5-rung RE ladder $\alpha_z \in \{100, 250, 700, 2000, 10^4\}$.

with $\mathtt{M}$ aligning pairs of characters $(i,j)$ with probability $\exp(RT)_{ij}$, $\mathtt{I}$ emitting unaligned descendant characters $(\epsilon, i)$ with $i \sim \pi$, and $\mathtt{D}$ consuming unaligned ancestral characters $(i,\epsilon)$ without any additional weight penalty.

This transducer can be interpreted as a decomposition of each mortal-link BDI trajectory into founder survival/death, possible survival of the youngest orphan (which, by insertion order, is the leftmost surviving descendant when the founder dies), and a geometric tail of further offspring. This yields the parameters $\alpha, \beta, \gamma$ appearing in the WFST transition matrix and determines which transition counts correspond to birth and death events.

**TKF91 Pair HMM** The joint distribution of ancestor and descendant is given by the transducer composition product of the stationary HMM and the transition WFST. This product can itself be represented as a Pair HMM with states $\{\mathtt{S},\mathtt{M},\mathtt{I},\mathtt{D},\mathtt{E}\}$ that models the joint distribution, as opposed to the conditional distribution represented by the WFST. The transition matrix $\tau$ of the Pair HMM is similar to the transition matrix $\mathbb{T}$ of the transducer, with the differences that incoming transitions to $\mathtt{M}$ and $\mathtt{D}$ states acquire a factor of $\kappa$ for the extension of the ancestral sequence and transitions to $\mathtt{E}$ states acquire a factor of $1-\kappa$ for the termination of the ancestral sequence.

For concreteness, because we will refer to it frequently, the transition matrix of the TKF91 Pair HMM is

$$\tau(\lambda,\mu,T) = \left(\begin{array}{c|ccccc} & \mathtt{S} & \mathtt{M} & \mathtt{I} & \mathtt{D} & \mathtt{E} \\ \hline \mathtt{S} & 0 & (1-\beta)\alpha\kappa & \beta\kappa & (1-\beta)(1-\alpha)\kappa & (1-\beta)(1-\kappa) \\ \mathtt{M} & 0 & (1-\beta)\alpha\kappa & \beta\kappa & (1-\beta)(1-\alpha)\kappa & (1-\beta)(1-\kappa) \\ \mathtt{I} & 0 & (1-\beta)\alpha\kappa & \beta\kappa & (1-\beta)(1-\alpha)\kappa & (1-\beta)(1-\kappa) \\ \mathtt{D} & 0 & (1-\gamma)\alpha\kappa & \gamma\kappa & (1-\gamma)(1-\alpha)\kappa & (1-\gamma)(1-\kappa) \\ \mathtt{E} & 0 & 0 & 0 & 0 & 0 \end{array}\right)$$

The emission weights of the Pair HMM are similarly related to the input/output emission weights of the WFST, with $\mathtt{M}$- and $\mathtt{I}$-state emissions acquiring a factor of $\pi_i$ for the ancestral character $i$.

$$\mathbb{E}[B \mid x, y] - \mathbb{E}[D \mid x, y] = \hat{n}_{\mathtt{SI}} + \hat{n}_{\mathtt{MI}} + \hat{n}_{\mathtt{II}} - \hat{n}_{\mathtt{SD}} - \hat{n}_{\mathtt{MD}} - \hat{n}_{\mathtt{DD}} - \hat{n}_{\mathtt{ID}} \tag{A.13}$$

$$= \sum_{i,j} \hat{n}_{ij} \, \mathcal{C}^{\text{cons}}_{ij} \tag{A.14}$$

where $\mathcal{C}^{\text{cons}}_{ij} = \delta(j = \mathtt{I}) - \delta(j = \mathtt{D})$ counts whether the WFST transition $i \to j$ corresponds to a birth, death, or neither. Substituting into the BDI score identities gives

$$\mathbb{E}[S \mid x, y] = \frac{\sum_{i,j} \hat{n}_{ij} \left( \mathcal{C}^{\text{cons}}_{ij} - \mathcal{C}^{\lambda}_{ij} + \mathcal{C}^{\mu}_{ij} \right) - \lambda T}{\lambda - \mu} \tag{A.15}$$

$$\mathbb{E}[B \mid x, y] = \lambda \sum_{i,j} \hat{n}_{ij} \, \mathcal{C}^{\lambda}_{ij}(\lambda, \mu, t) + \lambda \, \mathbb{E}[S \mid x, y] + \lambda T \tag{A.16}$$

$$\mathbb{E}[D \mid x, y] = \mu \sum_{i,j} \hat{n}_{ij} \, \mathcal{C}^{\mu}_{ij}(\lambda, \mu, t) + \mu \, \mathbb{E}[S \mid x, y] \tag{A.17}$$

**M-step.** The goal is to maximize $\ell(\theta) = \ell_1(\lambda, \mu) + \ell_2(Q, \pi)$ where

$$\begin{aligned}
\ell_1(\lambda,\mu) &= (B+L)\log\lambda + (D-L-M)\log\mu + M\log(\mu-\lambda) - (\lambda+\mu)\,S - \lambda T \quad \text{(A.18)}\\
\ell_2(Q,\pi) &= \sum_i V_i' \log \pi_i + \sum_{j>i}(U_{ij}+U_{ji})\log Q_{ij} - \sum_{j>i} Q_{ij}\,(W_i\pi_j + W_j\pi_i) \quad \text{(A.19)}
\end{aligned}$$

using the reversible parameterization $\kappa = \lambda/\mu$ and $R_{ij} = Q_{ij}\pi_j$ and letting $V_i' = V_i + \sum_{j\neq i} U_{ji}$ for notational convenience. Differentiating and setting to zero yields equations the MLEs must satisfy. For $\lambda, \mu$ these are simultaneous quadratics

$$\begin{aligned}
\partial_\lambda \ell_1 &= \frac{B+L}{\lambda} - \frac{M}{\mu-\lambda} - S - T = 0 \quad \text{(A.20)}\\
\partial_\mu \ell_1 &= \frac{D-L-M}{\mu} + \frac{M}{\mu-\lambda} - S = 0 \quad \text{(A.21)}
\end{aligned}$$

which reduce to a single quadratic in $\kappa$. Setting $a = B + L$ and $b = D - L - M$

$$\kappa^2(S+T)b - \kappa\,(S(a+b+2M) + T(b+M)) + Sa = 0. \quad \text{(A.22)}$$

For the coefficients $A_\kappa = (S+T)b$, $B_\kappa = -\,(S(a+b+2M) + T(b+M))$, and $C_\kappa = Sa$, the root $\kappa \in (0,1)$ will be the smaller of the two roots

$$\begin{aligned}
\hat\kappa(B,D,L,M,S,T) &= \frac{-B_\kappa - \sqrt{B_\kappa^2 - 4A_\kappa C_\kappa}}{2A_\kappa} \quad \text{(A.23)}\\
\hat\mu(B,D,L,M,S,T) &= \frac{1}{S}\left(b + \frac{M}{1-\hat\kappa}\right) \quad \text{(A.24)}\\
\hat\lambda(B,D,L,M,S,T) &= \hat\kappa\hat\mu \quad \text{(A.25)}
\end{aligned}$$

For $Q, \pi$ the MLE equations are

$$\begin{aligned}
\partial_{Q_{ij}} \ell_2 &= \frac{U_{ij}+U_{ji}}{Q_{ij}} - (W_i\pi_j + W_j\pi_i) = 0 \quad \text{(A.26)}\\
\partial_{\pi_i} \ell_2 &= \frac{V_i'}{\pi_i} - \sum_{j\neq i} Q_{ij}W_j + \eta = 0 \quad \text{(A.27)}
\end{aligned}$$

**Row-stochasticity.** For each non-E source row, the row sum equals

$$r + (1-r)\Big[\beta + (1-\beta)\kappa\alpha + (1-\beta)\kappa(1-\alpha) + (1-\beta)(1-\kappa)\Big] = r + (1-r)\big[\beta + (1-\beta)\big] = 1$$

(replacing $\beta$ with $\gamma$ for the D row), confirming that $\tau''$ is a valid Markov transition matrix.

For benchmarks dominated by alignments without leading inserts (typical when the ancestor's $N$-terminus is well-conserved), the numerical effect of the discrepancy is small per pair. However, the $5\times5$ form does not represent the true conditional distribution $P(\text{descendant} \mid \text{ancestor}, \theta)$, and the $6\times6$ WFST in (A.29) is the construction we recommend.

Derivation of (A.50): the numerator is $T\mu\eta + \alpha = \alpha(se^u + 1) = \alpha\big((1{+}s) + s\varepsilon T + O(\varepsilon^2)\big)$ and the denominator is $\delta = \alpha\varepsilon\big((1{+}s) + s\varepsilon T/2 + O(\varepsilon^2)\big)$ from (A.47). So:

**Complementary parameters.** For any parameter $\xi$: $\frac{\partial \log(1-\xi)}{\partial \theta} = -\frac{\xi}{1-\xi} \frac{\partial \log \xi}{\partial \theta}$.

##### A.5.2 Observed-data score

Table A.1: Correspondence between transition counts, complete-data log-likelihood terms, and lineage fates.

| Log-likelihood term | Coefficient | Interpretation |
|---|---|---|
| $\log \alpha$ | $n_{SM} + n_{MM} + n_{DM} + n_{IM}$ | Founder survival (match) |
| $\log(1 - \alpha)$ | $n_{SD} + n_{MD} + n_{DD} + n_{ID}$ | Founder death (delete) |
| $\log(1 - \gamma)$ | $n_{DM} + n_{DD} + n_{DE}$ | Lineage extinction (no $\texttt{D} \to \texttt{I}$) |
| $\log \gamma$ | $n_{DI}$ | Youngest orphan ($\texttt{D} \to \texttt{I}$) |
| $\log \beta$ | $n_{SI} + n_{MI} + n_{II}$ | Further offspring (insertions continue) |
| $\log(1 - \beta)$ | $n_{IM} + n_{ID} + n_{IE}$ | No further offspring (insertions end) |

$$\sigma^2_{S,\mathrm{outer}} = \psi^2\,T^2\,\frac{\kappa}{(1-\kappa)^2} \tag{B.69}$$

$$\frac{\sigma^2_{S,\mathrm{outer}}}{\mathbb{E}_\pi[S]^2} = \frac{\psi^2}{\kappa\,/(1-\kappa)^2 \cdot T^2} \cdot \frac{\kappa}{(1-\kappa)^2} \;=\; \frac{\psi^2\,(1-\kappa)^2}{\kappa} \;\cdot\; \frac{1}{1}$$

The relative variance of $\hat{\lambda}$ from a single pair becomes (using the outer-variance approximation in (B.50), and noting that the $B$–$S$ covariance term partially cancels the $B$ and $S$ terms):

$$v_\lambda \;\approx\; \frac{\psi^2\,(1-\kappa)^2}{\kappa}\left[1 + \frac{\lambda^2}{(\lambda + 1/T)^2} - \frac{2\lambda}{\lambda + 1/T}\right] \;=\; \frac{\psi^2\,(1-\kappa)^2}{\kappa\,(1+\lambda T)^2} \tag{B.70}$$

This uses $\mathbb{E}[S] + T = (L+1)\,T = T/(1-\kappa)$, so $\lambda/(\lambda + 1/T) = \lambda T/(1+\lambda T) = \kappa\delta T/(\delta + \delta/(\kappa \cdots))$— more directly, substitute into (B.50):

$$\begin{aligned}
v_\lambda &= \frac{\lambda^2\sigma^2_{S,\mathrm{outer}}}{\mathbb{E}[B]^2} + \frac{\sigma^2_{S,\mathrm{outer}}}{(\mathbb{E}[S]+T)^2} - \frac{2\lambda\sigma^2_{S,\mathrm{outer}}}{\mathbb{E}[B](\mathbb{E}[S]+T)} \\
&= \sigma^2_{S,\mathrm{outer}}\left(\frac{\lambda^2(1-\kappa)^2}{\lambda^2 T^2} + \frac{(1-\kappa)^2}{T^2} - \frac{2\lambda(1-\kappa)^2}{\lambda T^2}\right) \\
&= \frac{\psi^2\kappa}{(1-\kappa)^2} \cdot \frac{(1-\kappa)^2}{T^2} \cdot \big(1 + 1 - 2\big) \\
&= 0.
\end{aligned}$$

The Maraschino composite log-likelihood for TKF92 is the sum, over divergence-time bins $b$ and adjacency types $\mathtt{X} \to \mathtt{Y}$, of the cherry-count multiplied by the model log-probability of that adjacency at that bin's representative time:

$$\mathcal{L}_{\text{cherry}}^{\text{TKF92}}(\lambda, \mu, r, Q, \pi) = \sum_{b=1}^{n_\tau} \Bigg[ \sum_{\mathtt{X},\mathtt{Y}} n_{\mathtt{X}\to\mathtt{Y}}^{(b)} \log \tau'_{\mathtt{X}\to\mathtt{Y}}(\lambda, \mu, \bar{T}_b, r) + \sum_{a,b} n_{a,b}^{\mathtt{M},(b)} \log \left[\pi_a \exp(R\bar{T}_b)_{ab}\right] + \sum_b n_b^{\mathtt{I},(b)} \log \pi_b + \sum_a n_a^{\mathtt{D},(b)} \log \pi_a \Bigg] \quad \text{(B.75)}$$

where $n_{\mathtt{X}\to\mathtt{Y}}^{(b)}$ are the binned transition counts, $n_{a,b}^{\mathtt{M},(b)}, n_b^{\mathtt{I},(b)}, n_a^{\mathtt{D},(b)}$ are the binned per-state emission counts, $\tau'$ is the TKF92 Pair HMM transition matrix from Section A.2.2, and $\bar{T}_b$ is the representative time of bin $b$ (typically the geometric mean of the bin endpoints). The first term scores indel/fragment behaviour; the remaining three score the substitution model. Boundary contributions ($\mathtt{S} \to \cdot$, $\cdot \to \mathtt{E}$, $\mathtt{S} \to \mathtt{E}$ for empty alignments) are scored against the corresponding $\mathtt{S}$-row and $\mathtt{E}$-column entries of $\tau'$.

**Algorithm 1** EM-around-Maraschino for a mixture of $K$ TKF92 components.

Initialize component parameters $\theta_1^{(0)}, \ldots, \theta_K^{(0)}$ and mixture weights $\pi_1^{(0)}, \ldots, \pi_K^{(0)}$.
**for** $t = 0, 1, \ldots$ **do**
  **E-step.** For each cherry $c$ in the training set, compute the responsibility $\gamma_{c,k}^{(t)} \propto \pi_k^{(t)} P(c \mid \theta_k^{(t)})$ (the per-component cherry-count likelihood normalised across components). Each $P(c \mid \theta_k)$ is a single forward pass through the cherry-count likelihood at the cherry's counts subtensor.
  **Component-weighted count accumulation.** For each component $k$, accumulate the responsibility-weighted cherry-count tensor $C_k^{(t+1)} = \sum_c \gamma_{c,k}^{(t)} C_c$.
  **Component M-step (Maraschino).** For each component $k$, run Maraschino (Section B.4.1) on $C_k^{(t+1)}$ to obtain $\theta_k^{(t+1)}$.
  **Mixture-weight M-step.** $\pi_k^{(t+1)} \propto \sum_c \gamma_{c,k}^{(t)}$.
**end for**

normalised across $k$ for each $(m, c)$, and the responsibility-weighted Holmes–Rubin sufficient statistics for class $k$ are

$$\begin{aligned}
\hat{W}^{(t+1,k)}_i &= \sum_{m,c} \gamma^{(t)}_{m,c}(k) \sum_{p\,:\,\mathtt{M}\text{ at }c} \mathcal{C}^W_i\big(a_{p,c}, b_{p,c}, t_p\big),\\
\hat{U}^{(t+1,k)}_{ij} &= \sum_{m,c} \gamma^{(t)}_{m,c}(k) \sum_{p\,:\,\mathtt{M}\text{ at }c} \mathcal{C}^U_{ij}\big(a_{p,c}, b_{p,c}, t_p\big),\\
\hat{V}^{(t+1,k)}_i &= \sum_{m,c} \gamma^{(t)}_{m,c}(k) \sum_{p\,:\,\mathtt{M}\text{ at }c} \delta\big(a_{p,c} = i\big),
\end{aligned} \tag{B.78}$$

**Peeling (postorder).** For each node $v$ in postorder, compute the conditional likelihood:

$$\mathrm{CL}_v(a) = \psi_v(a) \prod_{\text{children } c} \left[ \sum_{a_c} \phi_{(v,c)}(a, a_c) \, \mathrm{CL}_c(a_c) \right] \tag{B.89}$$

with $\mathrm{CL}_\ell(a) = \delta(a = y_\ell)$ for observed leaves. The log-partition function is $\log Z_c = \log \sum_a \pi(a) \, \mathrm{CL}_{\text{root}}(a)$.

##### B.6.2 Restricted generative model

The model joint over leaf and internal-node presence patterns at the $L$ MSA columns is

$$p(\mathrm{MSA},\ \{X^v\}_{v\in\mathcal{I}} \mid \mathbf{t},\theta) = p_{\mathrm{singlet}}\big(X^{\mathrm{root}} \mid \theta\big) \prod_{(v\to w)\in\mathbf{t}} P\big(X^w \mid X^v, d_{vw}, \theta\big), \tag{B.92}$$

where $p_{\mathrm{singlet}}$ is the TKF stationary distribution (Section B.6.3, below) and each $P(X^w \mid X^v, t)$ is the per-branch presence-pattern conditional defined by the WFST path log-likelihood (Section B.6.4). Marginalising (B.92) over internal patterns supported on the $L$ observed MSA columns gives a quantity we denote $\tilde{p}(\mathrm{MSA} \mid \mathbf{t},\theta)$. The full TKF92 marginal $p(\mathrm{MSA} \mid \mathbf{t},\theta)$ also sums over evolutionary histories with ancestral residues that get inserted and then fully deleted on every leaf-bound lineage before reaching any leaf — *ghost columns* that leave no MSA trace and so are not representable as values of $X_n^v$ at observed columns $1,\ldots,L$. Since each ghost-column history adds a non-negative probability to $p$ that is absent from $\tilde{p}$,

$$\tilde{p}(\mathrm{MSA} \mid \mathbf{t},\theta) \ \le\ p(\mathrm{MSA} \mid \mathbf{t},\theta). \tag{B.93}$$

The gap $\log p - \log \tilde{p} \ge 0$ depends on the model parameters and the tree but not on the variational distribution $q$ introduced below. For the ancestral-presence reconstruction task — fix $\theta$, optimise $q$ to recover internal-node marginals at the observed columns — this gap is a $q$-independent constant and the restricted-model posterior at observed columns coincides with the full-model posterior conditioned on the same support, so all inferences below proceed unchanged.

##### B.6.3 Singlet HMM at the root

The TKF92 stationary HMM (Section A.2.2) emits a sequence of length $L_{\mathrm{root}}$ with

$$p_{\mathrm{singlet}}(L_{\mathrm{root}}) = \begin{cases} 1-\kappa & L_{\mathrm{root}} = 0, \\ \kappa\, p^{L_{\mathrm{root}}-1}\,(1-r)(1-\kappa) & L_{\mathrm{root}} \ge 1, \end{cases} \tag{B.94}$$

where $\kappa = \lambda/\mu$ and $p = r + (1-r)\kappa$. Identifying $L_{\mathrm{root}}$ with the number of MSA columns at which the root is in state $xP$ ($L_{\mathrm{root}} = \sum_n \delta(X_n^{\mathrm{root}} = 1)$) gives the root prior in the joint (B.92). The two-cases form arises because the TKF92 singlet HMM has distinct "from S" (probability $\kappa$ to emit, $1-\kappa$ to end) and "from emitting state" (probability $p$ to extend, $(1-r)(1-\kappa)$ to end) transition contexts.

$$\mathcal{T}^{(\mathrm{eqm})}_{\mathtt{I}_{lf}, \mathtt{I}_{mg}} = \frac{\rho_f^{(l)} \varphi_l \cdot \kappa_0 \cdot v_m \kappa_m w_{mg}}{1 - \kappa_0 z_0} + \delta(l{=}m)\, \rho_f^{(l)} \kappa_l w_{lg} + \delta(l{=}m)\, r_{fg}^{(l)}$$

where $\rho_f^{(n)} = 1 - \sum_g r_{fg}^{(n)}$ is the fragment termination probability, and the three terms represent (respectively) inter-domain transitions via the null-corrected top-level geometric process, same-domain new-fragment transitions, and same-domain intra-fragment Markov transitions on fragment-types. The $\mathtt{S}$ row is $\mathcal{T}^{(\mathrm{eqm})}_{\mathtt{S}, \mathtt{I}_{mg}} = \kappa_0 v_m \kappa_m w_{mg} / (1 - \kappa_0 z_0)$ and $\mathcal{T}^{(\mathrm{eqm})}_{\mathtt{S}, \mathtt{E}} = (1 - \kappa_0)/(1 - \kappa_0 z_0)$; the $\mathtt{E}$ column is $\mathcal{T}^{(\mathrm{eqm})}_{\mathtt{I}_{lf}, \mathtt{E}} = \rho_f^{(l)} \varphi_l (1 - \kappa_0)/(1 - \kappa_0 z_0)$.

The transition matrix $\chi(\theta, T)$ for this collapsed Pair HMM has entries

| Source $i$ | Destination $j$ | $\chi_{ij} = p_{\text{out}} \times \mathcal{T}_{\texttt{UV}} \times p_{\text{in}} + \delta(\texttt{U} = \texttt{V})\delta(l = m)(p_{\text{SameDom}} + \delta(\texttt{X} = \texttt{Y}) p_{\text{SameFrag}})$ | | | | |
|---|---|---|---|---|---|---|
| $(\texttt{UX}_{lf})$ | $(\texttt{VY}_{mg})$ | $p_{\text{out}}(\texttt{U}, \texttt{X}, l, f)$ | $\mathcal{T}_{\texttt{UV}}$ | $p_{\text{in}}(\texttt{V}, \texttt{Y}, m, g)$ | $p_{\text{SameDom}}(\texttt{X}, \texttt{Y}, l, f, g)$ | $p_{\text{SameFrag}}(l, f, g)$ |
| $\texttt{SS}$ | $\texttt{MY}_{mg}$ | $1$ | $\mathcal{T}_{\texttt{SM}}$ | $(1-z_T)^{-1} v_m \tau^{(m)}_{\texttt{SY}} w_{mg}$ | $0$ | $0$ |
| | $\texttt{II}_{mg}$ | $1$ | $\mathcal{T}_{\texttt{SI}}$ | $(1-z_0)^{-1} v_m \kappa_m w_{mg}$ | $0$ | $0$ |
| | $\texttt{DD}_{mg}$ | $1$ | $\mathcal{T}_{\texttt{SD}}$ | $(1-z_0)^{-1} v_m \kappa_m w_{mg}$ | $0$ | $0$ |
| | $\texttt{EE}$ | $1$ | $\mathcal{T}_{\texttt{SE}}$ | $1$ | $0$ | $0$ |
| $\texttt{MX}_{lf}$ | $\texttt{MY}_{mg}$ | $\rho^{(l)}_f \tau^{(l)}_{\texttt{XE}}$ | $\mathcal{T}_{\texttt{MM}}$ | $(1-z_T)^{-1} v_m \tau^{(m)}_{\texttt{SY}} w_{mg}$ | $\rho^{(l)}_f \tau^{(l)}_{\texttt{XY}} w_{lg}$ | $r^{(l)}_{fg}$ |
| | $\texttt{II}_{mg}$ | $\rho^{(l)}_f \tau^{(l)}_{\texttt{XE}}$ | $\mathcal{T}_{\texttt{MI}}$ | $(1-z_0)^{-1} v_m \kappa_m w_{mg}$ | $0$ | $0$ |
| | $\texttt{DD}_{mg}$ | $\rho^{(l)}_f \tau^{(l)}_{\texttt{XE}}$ | $\mathcal{T}_{\texttt{MD}}$ | $(1-z_0)^{-1} v_m \kappa_m w_{mg}$ | $0$ | $0$ |
| | $\texttt{EE}$ | $\rho^{(l)}_f \tau^{(l)}_{\texttt{XE}}$ | $\mathcal{T}_{\texttt{ME}}$ | $1$ | $0$ | $0$ |
| $\texttt{II}_{lf}$ | $\texttt{MY}_{mg}$ | $\rho^{(l)}_f \varphi_l$ | $\mathcal{T}_{\texttt{IM}}$ | $(1-z_T)^{-1} v_m \tau^{(m)}_{\texttt{SY}} w_{mg}$ | $0$ | $0$ |
| | $\texttt{II}_{mg}$ | $\rho^{(l)}_f \varphi_l$ | $\mathcal{T}_{\texttt{II}}$ | $(1-z_0)^{-1} v_m \kappa_m w_{mg}$ | $\rho^{(l)}_f \kappa_l w_{lg}$ | $r^{(l)}_{fg}$ |
| | $\texttt{DD}_{mg}$ | $\rho^{(l)}_f \varphi_l$ | $\mathcal{T}_{\texttt{ID}}$ | $(1-z_0)^{-1} v_m \kappa_m w_{mg}$ | $0$ | $0$ |
| | $\texttt{EE}$ | $\rho^{(l)}_f \varphi_l$ | $\mathcal{T}_{\texttt{IE}}$ | $1$ | $0$ | $0$ |
| $\texttt{DD}_{lf}$ | $\texttt{MY}_{mg}$ | $\rho^{(l)}_f \varphi_l$ | $\mathcal{T}_{\texttt{DM}}$ | $(1-z_T)^{-1} v_m \tau^{(m)}_{\texttt{SY}} w_{mg}$ | $0$ | $0$ |
| | $\texttt{II}_{mg}$ | $\rho^{(l)}_f \varphi_l$ | $\mathcal{T}_{\texttt{DI}}$ | $(1-z_0)^{-1} v_m \kappa_m w_{mg}$ | $0$ | $0$ |
| | $\texttt{DD}_{mg}$ | $\rho^{(l)}_f \varphi_l$ | $\mathcal{T}_{\texttt{DD}}$ | $(1-z_0)^{-1} v_m \kappa_m w_{mg}$ | $\rho^{(l)}_f \kappa_l w_{lg}$ | $r^{(l)}_{fg}$ |
| | $\texttt{EE}$ | $\rho^{(l)}_f \varphi_l$ | $\mathcal{T}_{\texttt{DE}}$ | $1$ | $0$ | $0$ |

, (C.3)

where $\tau^{(n)} \equiv \tau(\lambda_n, \mu_n, T)$, $\rho^{(n)}_f \equiv 1 - \sum_g r^{(n)}_{fg}$ is the fragment termination probability for fragment state $f$ in domain $n$, and (as before) $\varphi_l \equiv 1 - \kappa_l$.

The substitution parameters $(Q^{(c)}, \pi^{(c)})$ do not appear in the transition matrix, but in the emission probabilities of the various states. The probability of emitting token $(a, b)$ from state $\texttt{MM}_{nf}$ is

$$P(a, b \mid \texttt{MM}_{nf}, T) = \sum_{c=1}^{C} u_{nfc}\, \pi^{(c)}_a \exp(R^{(c)} T)_{a,b} \tag{C.4}$$

where $R^{(c)} = Q^{(c)} \cdot \operatorname{diag}(\pi^{(c)})$ is the rate matrix for site class $c$, and $u_{nfc}$ is the probability that fragment state $f$ in domain $n$ generates site class $c$. The probability of emitting ancestral token $a$ from states $\{\texttt{MD}_{nf}, \texttt{DD}_{nf}\}$, or descendant token $a$ from states $\{\texttt{MI}_{nf}, \texttt{II}_{nf}\}$, is $\sum_c u_{nfc}\, \pi^{(c)}_a$.

*Per-class equilibrium:* $\pi_a^{(c)} \propto V_a^{(c)} + \alpha_\pi - 1$. (This is the standard empirical-frequency estimator; the exact EM M-step couples $\pi$ with $Q$ through dwell-time statistics, but the approximation is standard practice for GTR models.)

*Per-class exchangeability:* The rate is $Q_{a,a'}^{(c)} \cdot \pi_{a'}^{(c)}$. The exchangeability is estimated from Holmes–Rubin transition and dwell counts:

$$Q_{a,a'}^{(c)} = \frac{U_{a,a'}^{(c)} + U_{a',a}^{(c)}}{W_a^{(c)} \cdot \pi_{a'}^{(c)} + W_{a'}^{(c)} \cdot \pi_a^{(c)}}$$

At each internal node $v$ with children $l, r$ and observed descendant sequences $c_l, c_r$, we seek

$$\hat{a}_v = \operatorname*{argmax}_a \bigl[\log P(a, c_l \mid B_l) + \log P(a, c_r \mid B_r) - \log P(a \mid R)\bigr] \tag{C.9}$$

where $P(a, c_k \mid B_k)$ is the pair HMM forward probability on branch $k$ and $P(a \mid R)$ is the singlet probability under the root generator, subtracted to avoid double-counting the prior on $a$.

**Unpeeling (preorder).** Propagate top-down to obtain the posterior marginal at each node:

$$q_c^{(v)}(a) \propto \mathrm{CL}_v(a) \cdot \mathrm{msg}_{\text{parent}\to v}(a) \tag{C.15}$$

and the pairwise marginal on each edge:

$$q_c^{(u,v)}(a_u, a_v) \propto \mathrm{msg}_{\text{above}\, u}(a_u) \cdot \phi_{(u,v)}(a_u, a_v) \cdot \mathrm{CL}_v(a_v) \tag{C.16}$$

$$\texttt{DelFragType}[k,f] \xrightarrow{(a,\epsilon)} \texttt{DelEmit}[k,f] : \quad \sum_{c=1}^{C} u_{kfc}\, \pi_a^{(c)} \quad \text{(delete } a\text{)} \tag{C.73}$$

$$\texttt{IFragType}[k,f] \xrightarrow{(\epsilon,b)} \texttt{IEmit}[k,f] : \quad \sum_{c=1}^{C} u_{kfc}\, \pi_b^{(c)} \quad \text{(insert } b\text{)} \tag{C.74}$$

$$\texttt{DFragType}[k,f] \xrightarrow{(a,\epsilon)} \texttt{DEmit}[k,f] : \quad \sum_{c=1}^{C} u_{kfc}\, \pi_a^{(c)} \quad \text{(delete } a\text{)} \tag{C.75}$$

**Intra-fragment fragment-type transition vs. fragment termination** Within each fragment of domain $k$ the fragment-type process is a Markov chain on $\mathcal{F}+2$ states (start, end, and $\mathcal{F}$ fragment-type states): from the current emit state with fragment-type $f$, the chain either advances within the fragment to fragment-type $g$ with probability $r_{fg}^{(k)}$, or terminates the fragment (transition to the end state) with probability $\rho_f^{(k)} = 1 - \sum_g r_{fg}^{(k)}$. Different fragments are statistically independent realisations of this chain. The transition from each emit state goes to any fragment-type's type-selection state within the current fragment (not just the same type), or to the fragment end:

$$\texttt{MatEmit}[k,f] \to \texttt{MatFragType}[k,g] : \quad r_{fg}^{(k)} \quad \text{(intra-fragment type transition } f \to g\text{)} \tag{C.76}$$

$$\texttt{MatEmit}[k,f] \to \texttt{MatFragEnd}[k,f] : \quad \rho_f^{(k)} \quad \text{(fragment termination)} \tag{C.77}$$

$$\texttt{InsEmit}[k,f] \to \texttt{InsFragType}[k,g] : \quad r_{fg}^{(k)} \tag{C.78}$$

$$\texttt{InsEmit}[k,f] \to \texttt{InsFragEnd}[k,f] : \quad \rho_f^{(k)} \tag{C.79}$$

$$\texttt{DelEmit}[k,f] \to \texttt{DelFragType}[k,g] : \quad r_{fg}^{(k)} \tag{C.80}$$

$$\texttt{DelEmit}[k,f] \to \texttt{DelFragEnd}[k,f] : \quad \rho_f^{(k)} \tag{C.81}$$

$$\texttt{IEmit}[k,f] \to \texttt{IFragType}[k,g] : \quad r_{fg}^{(k)} \tag{C.82}$$

$$\texttt{IEmit}[k,f] \to \texttt{IFragEnd}[k,f] : \quad \rho_f^{(k)} \tag{C.83}$$

$$\texttt{DEmit}[k,f] \to \texttt{DFragType}[k,g] : \quad r_{fg}^{(k)} \tag{C.84}$$

$$\texttt{DEmit}[k,f] \to \texttt{DFragEnd}[k,f] : \quad \rho_f^{(k)} \tag{C.85}$$

where $g$ ranges over all $\mathcal{F}$ fragment types. For $\mathcal{F} = 1$, this reduces to a scalar self-extension with $r_{11}^{(k)} = r_{k1}$ and $\rho_1^{(k)} = 1 - r_{k1}$.

##### C.3.3 Null State Classification

Every state except `S`, `E`, and the five emit states $\texttt{MatEmit}[k,f]$, $\texttt{InsEmit}[k,f]$, $\texttt{DelEmit}[k,f]$, $\texttt{IEmit}[k,f]$, $\texttt{DEmit}[k,f]$ is a **null state** (non-emitting). The collapsed model retains only `S`, `E`, and the $5\mathcal{N}\mathcal{F}$ emit states.

Within the entry, the path `MatDomType`$[k] \to$ `MatFrag`$[k]$ uses $\tau_k[\mathtt{S}, \cdot]$. The phantom path `MatDomType`$[k] \to$ `MatDomEnd` has probability $\tau_k[\mathtt{S}, \mathtt{E}] = (1 - \beta_k)(1 - \kappa_k)$ and contributes a phantom birth-death event to domain $k$'s BDI statistics.

| Tensor | Shape | Meaning |
|---|---|---|
| $B$ | $n_\tau \times (\lvert\mathcal{A}\rvert + 2)^2$ | Singlet bigrams (incl. `S`/`E`) |
| $C^{\mathtt{MM}}$ | $n_\tau \times \lvert\mathcal{A}\rvert^4$ | Match→Match: $(a, b, a', b')$ |
| $C^{\mathtt{MI}}$ | $n_\tau \times \lvert\mathcal{A}\rvert^3$ | Match→Insert: $(a, b, b')$ |
| $C^{\mathtt{MD}}$ | $n_\tau \times \lvert\mathcal{A}\rvert^3$ | Match→Delete: $(a, b, a')$ |
| $C^{\mathtt{IM}}$ | $n_\tau \times \lvert\mathcal{A}\rvert^3$ | Insert→Match: $(b, a', b')$ |
| $C^{\mathtt{II}}$ | $n_\tau \times \lvert\mathcal{A}\rvert^2$ | Insert→Insert: $(b, b')$ |
| $C^{\mathtt{ID}}$ | $n_\tau \times \lvert\mathcal{A}\rvert^2$ | Insert→Delete: $(b, a')$ |
| $C^{\mathtt{DM}}$ | $n_\tau \times \lvert\mathcal{A}\rvert^3$ | Delete→Match: $(a, a', b')$ |
| $C^{\mathtt{DD}}$ | $n_\tau \times \lvert\mathcal{A}\rvert^2$ | Delete→Delete: $(a, a')$ |
| $C^{\mathtt{DI}}$ | $n_\tau \times \lvert\mathcal{A}\rvert^2$ | Delete→Insert: $(a, b')$ |
| $C^{\mathtt{SM}}$ | $n_\tau \times \lvert\mathcal{A}\rvert^2$ | Start→Match: $(a', b')$ |
| $C^{\mathtt{SI}}$ | $n_\tau \times \lvert\mathcal{A}\rvert$ | Start→Insert: $(b')$ |
| $C^{\mathtt{SD}}$ | $n_\tau \times \lvert\mathcal{A}\rvert$ | Start→Delete: $(a')$ |
| $C^{\mathtt{ME}}$ | $n_\tau \times \lvert\mathcal{A}\rvert^2$ | Match→End: $(a, b)$ |
| $C^{\mathtt{IE}}$ | $n_\tau \times \lvert\mathcal{A}\rvert$ | Insert→End: $(b)$ |
| $C^{\mathtt{DE}}$ | $n_\tau \times \lvert\mathcal{A}\rvert$ | Delete→End: $(a)$ |
| $C^{\mathtt{SE}}$ | $n_\tau$ | Start→End (empty alignment) |

| Source | Dest | Input | Output | Weight |
|---|---|---|---|---|
| S | V | $\varepsilon$ | $\varepsilon$ | $p_{\mathtt{SV}}$ |
| S | I | $\varepsilon$ | $b$ | $p_{\mathtt{SI}}(Y, b)$ |
| S | E | $\varepsilon$ | $\varepsilon$ | $p_{\mathtt{SE}}$ |
| V | M | $a$ | $b$ | $p_{\mathtt{VM}}(X, Y, a, b)$ |
| V | D | $a$ | $\varepsilon$ | $p_{\mathtt{VD}}(X, Y, a)$ |
| W | M | $a$ | $b$ | $p_{\mathtt{WM}}(X, Y, a, b)$ |
| W | D | $a$ | $\varepsilon$ | $p_{\mathtt{WD}}(X, Y, a)$ |
| M | V | $\varepsilon$ | $\varepsilon$ | $p_{\mathtt{MV}}$ |
| M | I | $\varepsilon$ | $b$ | $p_{\mathtt{MI}}(X, Y, b)$ |
| M | E | $\varepsilon$ | $\varepsilon$ | $p_{\mathtt{ME}}$ |
| I | V | $\varepsilon$ | $\varepsilon$ | $p_{\mathtt{IV}}$ |
| I | I | $\varepsilon$ | $b$ | $p_{\mathtt{II}}(X, Y, b)$ |
| I | E | $\varepsilon$ | $\varepsilon$ | $p_{\mathtt{IE}}$ |
| D | W | $\varepsilon$ | $\varepsilon$ | $p_{\mathtt{DW}}$ |
| D | I | $\varepsilon$ | $b$ | $p_{\mathtt{DI}}(X, Y, b)$ |
| D | E | $\varepsilon$ | $\varepsilon$ | $p_{\mathtt{DE}}$ |

Note that `V` and `W` are *not* associated with emissions; they serve only to enforce the waiting-machine property. Transitions from `V` and `W` always consume an ancestor input symbol; transitions from `M`, `I`, `D`, and `S` never do.

**Parameterization from the MixDom Pair HMM.** We need the expected frequency of transitions conditioned on (last ancestor input $X$, last descendant output $Y$, transition type, new symbols). The key subtlety: through insert states, the last ancestor symbol $X$ must be *propagated* from the preceding match or delete, since inserts do not consume input.

Using the path notation from above, we enumerate all adjacency types that arise in the MixDom Pair HMM. Write $\mathtt{M}[X, Y]$ for a match state that inputs $X$ and outputs $Y$, $\mathtt{I}[Y]$ for an insert that outputs $Y$, and $\mathtt{D}[X]$ for a delete that inputs $X$. The last-ancestor and last-descendant context is carried implicitly.

**Adjacency frequencies.** The following table lists all adjacency types and their corresponding path marginalizations. In each case, the frequency is computed as a sum over MixDom Pair HMM states, with null states marginalized via $N^*$. Write $\circ$ for the boundary (start/end) context.

| Context | Adjacency | MixDom path | Frequency |
|---|---|---|---|
| *Start → Match:* | | | |
| | $\mathtt{S} \to \mathtt{M}[X', Y']$ | $\mathtt{S} \xrightarrow{\mathcal{Z}^*} \mathtt{M}[X', Y']$ | $f^{\mathtt{SM}}(X', Y')$ |
| *Start → Insert:* | | | |
| | $\mathtt{S} \to \mathtt{I}[Y']$ | $\mathtt{S} \xrightarrow{\mathcal{Z}^*} \mathtt{I}[Y']$ | $f^{\mathtt{SI}}(Y')$ |
| *Start → End (empty sequence):* | | | |
| | $\mathtt{S} \to \mathtt{E}$ | $\mathtt{S} \xrightarrow{\mathcal{Z}^*} \mathtt{E}$ | $f^{\mathtt{SE}}$ |
| *Match → Match (via null states only):* | | | |
| | $\mathtt{M}[X, Y] \to \mathtt{M}[X', Y']$ | $\mathtt{M}[X, Y] \xrightarrow{\mathcal{Z}^*} \mathtt{M}[X', Y']$ | $f^{\mathtt{MM}}(X, Y, X', Y')$ |
| *Match → Insert:* | | | |
| | $\mathtt{M}[X, Y] \to \mathtt{I}[Y']$ | $\mathtt{M}[X, Y] \xrightarrow{\mathcal{Z}^*} \mathtt{I}[Y']$ | $f^{\mathtt{MI}}(X, Y, Y')$ |
| *Match → Delete:* | | | |
| | $\mathtt{M}[X, Y] \to \mathtt{D}[X']$ | $\mathtt{M}[X, Y] \xrightarrow{\mathcal{Z}^*} \mathtt{D}[X']$ | $f^{\mathtt{MD}}(X, Y, X')$ |
| *Match → End:* | | | |
| | $\mathtt{M}[X, Y] \to \mathtt{E}$ | $\mathtt{M}[X, Y] \xrightarrow{\mathcal{Z}^*} \mathtt{E}$ | $f^{\mathtt{ME}}(X, Y)$ |
| *Insert → Insert (ancestor context $X$ propagated):* | | | |
| | $\mathtt{I}[Y] \to \mathtt{I}[Y']$ | $\mathtt{I}[Y] \xrightarrow{\mathcal{Z}^*} \mathtt{I}[Y']$ | $f^{\mathtt{II}}(X, Y, Y')$ |
| *Insert → Match (ancestor context $X$ propagated):* | | | |
| | $\mathtt{I}[Y] \to \mathtt{M}[X', Y']$ | $\mathtt{I}[Y] \xrightarrow{\mathcal{Z}^*} \mathtt{M}[X', Y']$ | $f^{\mathtt{IM}}(X, Y, X', Y')$ |
| *Insert → Delete (ancestor context $X$ propagated):* | | | |
| | $\mathtt{I}[Y] \to \mathtt{D}[X']$ | $\mathtt{I}[Y] \xrightarrow{\mathcal{Z}^*} \mathtt{D}[X']$ | $f^{\mathtt{ID}}(X, Y, X')$ |
| *Insert → End (ancestor context $X$ propagated):* | | | |
| | $\mathtt{I}[Y] \to \mathtt{E}$ | $\mathtt{I}[Y] \xrightarrow{\mathcal{Z}^*} \mathtt{E}$ | $f^{\mathtt{IE}}(X, Y)$ |
| *Delete → Match (descendant context $Y$ propagated):* | | | |
| | $\mathtt{D}[X] \to \mathtt{M}[X', Y']$ | $\mathtt{D}[X] \xrightarrow{\mathcal{Z}^*} \mathtt{M}[X', Y']$ | $f^{\mathtt{DM}}(X, Y, X', Y')$ |
| *Delete → Delete (descendant context $Y$ propagated):* | | | |
| | $\mathtt{D}[X] \to \mathtt{D}[X']$ | $\mathtt{D}[X] \xrightarrow{\mathcal{Z}^*} \mathtt{D}[X']$ | $f^{\mathtt{DD}}(X, Y, X')$ |
| *Delete → Insert (descendant context $Y$ propagated):* | | | |
| | $\mathtt{D}[X] \to \mathtt{I}[Y']$ | $\mathtt{D}[X] \xrightarrow{\mathcal{Z}^*} \mathtt{I}[Y']$ | $f^{\mathtt{DI}}(X, Y, Y')$ |
| *Delete → End (descendant context $Y$ propagated):* | | | |
| | $\mathtt{D}[X] \to \mathtt{E}$ | $\mathtt{D}[X] \xrightarrow{\mathcal{Z}^*} \mathtt{E}$ | $f^{\mathtt{DE}}(X, Y)$ |

In all cases, the ancestor context $X$ is propagated through insert states (which do not consume input), and the descendant context $Y$ is propagated through delete states (which do not emit output). The notation $\xrightarrow{\mathcal{Z}^*}$ denotes zero or more transitions through null states, marginalized via the null closure $N^*$.

**Computing the frequencies.** Each frequency above is computed from the MixDom Pair HMM as follows. Let $\pi_s$ denote the stationary probability of state $s$ and $\hat{T}_{s_1 s_2}$ the null-marginalized effective transition. For the simplest case (direct adjacency):

$$f^{\mathtt{MD}}(X, Y, X') = \sum_{s_1 \in \Omega^{\mathtt{M}}} \sum_{s_2 \in \Omega^{\mathtt{D}}} \left( \sum_i \pi_i \hat{T}^*_{i s_1} \right) e_{s_1}(X, Y) \, \hat{T}_{s_1 s_2} \, e_{s_2}(X') \left( \sum_j \hat{T}^*_{s_2 j} \right)$$

Boundary frequencies use the start/end rows of $\hat{T}$:

$$f^{\mathtt{SM}}(X',Y') = \sum_{s\in\Omega^{\mathtt{M}}} \hat{T}_{\mathtt{S}s}\, e_s(X',Y') \left(\sum_j \hat{T}^*_{sj}\right), \quad f^{\mathtt{ME}}(X,Y) = \sum_{s\in\Omega^{\mathtt{M}}} \left(\sum_i \pi_i \hat{T}^*_{is}\right) e_s(X,Y)\, \hat{T}_{s\,\mathtt{E}}$$

(and analogously for $f^{\mathtt{SI}}$, $f^{\mathtt{SE}}$, $f^{\mathtt{IE}}$, $f^{\mathtt{DE}}$).

**Normalization to transducer parameters.** Given the adjacency frequencies, the order-1 transducer weights are obtained by normalization. For each context $(X,Y)$ and source machine state, normalize outgoing weights to sum to 1:

*Start transitions* (context $\circ$):

$$p_{\mathtt{SV}} = \frac{\sum_{X',Y'} f^{\mathtt{SM}}(X',Y')}{\sum_{X',Y'} f^{\mathtt{SM}}(X',Y') + \sum_{Y'} f^{\mathtt{SI}}(Y') + f^{\mathtt{SE}}}, \quad p_{\mathtt{SI}}(b) = \frac{f^{\mathtt{SI}}(b)}{\sum_{X',Y'} f^{\mathtt{SM}}(X',Y') + \sum_{Y'} f^{\mathtt{SI}}(Y') + f^{\mathtt{SE}}}$$

(and $p_{\mathtt{SE}}$ uses the same denominator with numerator $f^{\mathtt{SE}}$).

*Wait-after-match/insert transitions* (context $(X,Y)$, consuming input $a$):

$$p_{\mathtt{VM}}(X,Y,a,b) = \frac{f^{\cdot\mathtt{M}}(X,Y,a,b)}{\sum_{a'}\left[\sum_{b'} f^{\cdot\mathtt{M}}(X,Y,a',b') + f^{\cdot\mathtt{D}}(X,Y,a')\right]}$$
$$p_{\mathtt{VD}}(X,Y,a) = \frac{f^{\cdot\mathtt{D}}(X,Y,a)}{\sum_{a'}\left[\sum_{b'} f^{\cdot\mathtt{M}}(X,Y,a',b') + f^{\cdot\mathtt{D}}(X,Y,a')\right]}$$

where $f^{\cdot\mathtt{M}}(X,Y,a,b) = f^{\mathtt{MM}}(X,Y,a,b) + f^{\mathtt{IM}}(X,Y,a,b)$ and $f^{\cdot\mathtt{D}}(X,Y,a) = f^{\mathtt{MD}}(X,Y,a) + f^{\mathtt{ID}}(X,Y,a)$, aggregating over both match and insert sources that share the $\mathtt{V}$ wait state.

*Wait-after-delete transitions* (context $(X,Y)$, consuming input $a$):

$$p_{\mathtt{WM}}(X,Y,a,b) = \frac{f^{\mathtt{DM}}(X,Y,a,b)}{\sum_{a'}\left[\sum_{b'} f^{\mathtt{DM}}(X,Y,a',b') + f^{\mathtt{DD}}(X,Y,a')\right]}$$
$$p_{\mathtt{WD}}(X,Y,a) = \frac{f^{\mathtt{DD}}(X,Y,a)}{\sum_{a'}\left[\sum_{b'} f^{\mathtt{DM}}(X,Y,a',b') + f^{\mathtt{DD}}(X,Y,a')\right]}$$

*Post-match transitions* (context $(X,Y)$, after matching with $a,b$; new context becomes $(a,b)$):

$$p_{\mathtt{MV}}(X,Y) = \frac{\sum_{a',b'} f^{\mathtt{MM}}(X,Y,a',b') + \sum_{a'} f^{\mathtt{MD}}(X,Y,a')}{Z^{\mathtt{M}}(X,Y)}$$
$$p_{\mathtt{MI}}(X,Y,b) = \frac{f^{\mathtt{MI}}(X,Y,b)}{Z^{\mathtt{M}}(X,Y)}, \quad p_{\mathtt{ME}}(X,Y) = \frac{f^{\mathtt{ME}}(X,Y)}{Z^{\mathtt{M}}(X,Y)}$$

where $Z^{\mathtt{M}}(X,Y) = \sum_{a',b'} f^{\mathtt{MM}}(X,Y,a',b') + \sum_{a'} f^{\mathtt{MD}}(X,Y,a') + \sum_b f^{\mathtt{MI}}(X,Y,b) + f^{\mathtt{ME}}(X,Y)$.

*Post-insert transitions* are analogous, using $f^{\mathtt{I}\cdot}$ frequencies with $Z^{\mathtt{I}}(X,Y)$.

*Post-delete transitions* (context $(X,Y)$, after deleting $a$; new context becomes $(a,Y)$):

$$p_{\mathtt{DW}}(X,Y) = \frac{\sum_{a'} f^{\mathtt{DD}}(X,Y,a') + \sum_{a',b'} f^{\mathtt{DM}}(X,Y,a',b')}{Z^{\mathtt{D}}(X,Y)}$$
$$p_{\mathtt{DI}}(X,Y,b) = \frac{f^{\mathtt{DI}}(X,Y,b)}{Z^{\mathtt{D}}(X,Y)}, \quad p_{\mathtt{DE}}(X,Y) = \frac{f^{\mathtt{DE}}(X,Y)}{Z^{\mathtt{D}}(X,Y)}$$

The model's per-(domain, fragment-type) emission factors at evolutionary time $T$ are:

$$\phi_{nf}^{\mathtt{M}}(a, b) \equiv e_{\mathtt{MM}_{nf}}(a, b) = \sum_{c=1}^{C} u_{nfc} \, P^{(c)}(a, b \mid T),$$

$$\phi_{nf}^{\mathtt{I}}(b) \equiv e_{\mathtt{MI}_{nf}}(b) = e_{\mathtt{II}_{nf}}(b) = \sum_{c} u_{nfc} \, \pi_b^{(c)},$$

$$\phi_{nf}^{\mathtt{D}}(a) \equiv e_{\mathtt{MD}_{nf}}(a) = e_{\mathtt{DD}_{nf}}(a) = \sum_{c} u_{nfc} \, \pi_a^{(c)}.$$

Each of these is a $C$-term mixture over the per-class GTRs. The match emission $\phi_{nf}^{\mathtt{M}}$ inherits the time dependence from the per-class transition kernels $P^{(c)}(T)$; the singlet emissions $\phi_{nf}^{\mathtt{I}}$, $\phi_{nf}^{\mathtt{D}}$ are time-independent.

##### C.5.4 Pair HMM Distillation

**Emitting state space** The collapsed MixDom Pair HMM with $\mathcal{N}$ domain types and $\mathcal{F}$ fragment types per domain has $5\mathcal{N}\mathcal{F}+2$ states: $\{\mathtt{SS},\mathtt{EE}\}$ and $\{\mathtt{MM}_{nf},\mathtt{MI}_{nf},\mathtt{MD}_{nf},\mathtt{II}_{nf},\mathtt{DD}_{nf} : n\in\mathcal{N}, f\in\mathcal{F}\}$.

Group the $5\mathcal{N}\mathcal{F}$ emitting states by emission type: match $\mathcal{M}=\{\mathtt{MM}_{nf}\}$, insert $\mathcal{I}=\{\mathtt{MI}_{nf},\mathtt{II}_{nf}\}$, delete $\mathcal{D}=\{\mathtt{MD}_{nf},\mathtt{DD}_{nf}\}$.

**Class-mixture emissions in the Pair HMM** The per-state emissions at $(d,f)$ are the class-mixture emissions $\phi^{\mathtt{M,I,D}}_{nf}$ defined in Section C.5.2. Two states with the same $(d,f)$ but different top-level types share the same equilibrium mixture, but the match state combines this with the time-evolved class-mixture transition kernel.

The per-(domain, fragment-type) emission factors do *not* lift cleanly out of the structural sum; however, they have low rank in the class index $c$ (Section C.5.2). Within a fixed $(d,f)$, all match states emit from $\phi^{\mathtt{M}}_{nf}$ and all insert/delete singlets from $\phi^{\mathtt{I}}_{nf}$ / $\phi^{\mathtt{D}}_{nf}$.

**Per-(domain, fragment)-pair structural weights** The pair adjacency frequencies decompose as sums over (domain, fragment-type) pairs. For match-to-match, writing $X=a, Y=b, X'=a', Y'=b'$:

$$f^{\mathcal{M}\mathcal{M}}(X,Y,X',Y') = \sum_{(n_1,f_1),(n_2,f_2)} C^{\mathcal{M}\mathcal{M}}_{(n_1,f_1),(n_2,f_2)}\,\phi^{\mathtt{M}}_{n_1f_1}(X,Y)\,\phi^{\mathtt{M}}_{n_2f_2}(X',Y') \tag{C.98}$$

where

$$C^{\mathcal{M}\mathcal{M}}_{(n_1,f_1),(n_2,f_2)} = \sum_{s_1\in\mathcal{M}_{n_1f_1}}\sum_{s_2\in\mathcal{M}_{n_2f_2}} L_{s_1}\,\hat{T}_{s_1s_2}\,R_{s_2}$$

are structural weights indexed by (domain, fragment-type) pairs (not just emission types). Similarly for other adjacency types ($\mathcal{M}\mathcal{I}$, $\mathcal{M}\mathcal{D}$, etc.), with the appropriate per-(domain, fragment-type) emissions.

Each adjacency frequency is therefore a sum of $(\mathcal{N}\mathcal{F})^2$ structural-weight terms, each multiplied by per-(domain, fragment-type) emission factors that are themselves rank-$C$ class mixtures.

**Context dependence** In the order-1 WFST, the wait-after-match/insert transition involves

$$\begin{aligned} f^{\cdot\mathcal{M}}(X,Y,a',b') = \sum_{(n_1,f_1),(n_2,f_2)} \Big( & C^{\mathcal{M}\mathcal{M}}_{(n_1,f_1),(n_2,f_2)}\,\phi^{\mathtt{M}}_{n_1f_1}(X,Y) \\ & + C^{\mathcal{I}\mathcal{M}}_{(n_1,f_1),(n_2,f_2)}\,\phi^{\mathtt{I}}_{n_1f_1}(Y)\Big)\phi^{\mathtt{M}}_{n_2f_2}(a',b') \end{aligned}$$

After normalization, $p_{\mathtt{VM}}(X,Y,a',b')$ depends on $(X,Y)$ through the joint posterior over (domain, fragment-type, class) of the previous emission. Specifically, the context enters through the $\mathcal{N}\mathcal{F}$-dimensional vector

$$\boldsymbol{\rho}(X,Y) = \Big(\sum_{(n_2,f_2)} C^{\mathcal{M}\mathcal{M}}_{(1,1),(n_2,f_2)}\,\phi^{\mathtt{M}}_{1,1}(X,Y) + \sum_{(n_2,f_2)} C^{\mathcal{I}\mathcal{M}}_{(1,1),(n_2,f_2)}\,\phi^{\mathtt{I}}_{1,1}(Y),\dots\Big)$$

which captures the relative likelihood of each source (domain, fragment-type).

$$\begin{aligned} (I - D_n^{\text{match}})^{-1} &= G_0 + G_0\, U_n\, K_n^{\text{inner}}\, V_n^{\top} G_0, \\ G_0 &\equiv I_3 \otimes (I_{\mathcal{F}} - r^{(n)})^{-1}, \\ K_n^{\text{inner}} &\equiv (I_3 - V_n^{\top} G_0\, U_n)^{-1}. \end{aligned} \tag{C.100}$$

$K_n^{\text{inner}}$ is a $3 \times 3$ matrix whose entries are rational functions of $w_n$, $r^{(n)}$, $\tau^{(n)}$ via a single $\mathcal{F}$-fold inner product $w_n^{\top}(I - r^{(n)})^{-1}\rho^{(n)}$ times $\tau^{(n)}_{3\times3}$.

The two singlet-domain blocks act on $\mathcal{F}$ states each ($\{\mathtt{II}_{nf} : f\}$ and $\{\mathtt{DD}_{nf} : f\}$):

$$d_n^{\text{ins}}[f, g] = d_n^{\text{del}}[f, g] = r^{(n)}_{fg} + \rho^{(n)}_f\, \kappa_n\, w_{n,g}.$$

Their inverse follows the same pattern: rank-1 correction to $(I - r^{(n)})$, giving

$$D_n = \begin{pmatrix} D_n^{\text{match}} & 0 & 0 \\ 0 & d_n^{\text{ins}} & 0 \\ 0 & 0 & d_n^{\text{del}} \end{pmatrix}$$

where $D_n^{\text{match}}$ is $3 \times 3$ (for matched-domain states $\texttt{MM}, \texttt{MI}, \texttt{MD}$) and the singlet-domain states have scalar self-loops:

**Domains: independent TKF rates and weights** Each domain type $n$ brings its own $(\alpha_n, \beta_n, \gamma_n, \kappa_n)$, creating genuinely different:

- $\texttt{M} \to \texttt{D}$ vs. $\texttt{M} \to \texttt{I}$ ratios (different $\kappa_n(1-\alpha_n)$ vs. $\beta_n$),
- `V` vs. `W` transition behaviour (different $\beta_n$ vs. $\gamma_n$),
- overall indel/match balance.

**Within-fragment Markov M-step.** For each domain $d$, the per-fragchar transition matrix $r_{f,f'}^{(d)}$ has Dirichlet-conjugate update from the R1 soft counts:

$$\hat{r}_{f,f'}^{(d)} = \frac{\hat{E}_{d,f,f'} + \alpha_r - 1}{\sum_{f''} \hat{E}_{d,f,f''} + \hat{N}_{d,f} + (N_{\mathcal{F}} + 1)(\alpha_r - 1)}, \quad \hat{\rho}_f^{(d)} = 1 - \sum_{f'} \hat{r}_{f,f'}^{(d)}, \tag{C.126}$$

where $\hat{E}_{d,f,f'} = \sum_{(v \to w),s,s'} \tilde{W}_{ss',(d,f),(d,f')}^{(R1),(v \to w)}$ and $\hat{N}_{d,f} = \sum_{(v \to w),s,s'} \left[\tilde{W}_{ss',(d,f),(d,\cdot)}^{(R2)} + \tilde{W}_{ss',(d,f),(\cdot,\cdot)}^{(R3)}\right]$ are the within- fragment and fragment-end soft counts respectively.

**Domain and fragment weight M-step.** The tuple priors update via Dirichlet-conjugate from per-route soft-counts:

$$\hat{v}_{d'} \propto \sum_{(v \to w)} \sum_{(d,f,f'),s,s'} \tilde{W}_{ss',(d,f),(d',f')}^{(R3),(v \to w)} + \alpha_n - 1, \tag{C.127}$$

$$\hat{w}_{d',f'} \propto \sum_{(v \to w)} \sum_{(d,f),s,s'} \left[\delta_{d,d'} \tilde{W}_{ss',(d,f),(d',f')}^{(R2)} + \tilde{W}_{ss',(d,f),(d',f')}^{(R3)}\right] + \alpha_w - 1, \tag{C.128}$$

##### C.8.2 Restricted generative model

Following Section B.6.2, the model joint over MSA columns and internal-node states is

$$p(\text{MSA}, \{Z^v\}_{v\in\mathcal{I}} \mid \mathbf{t}, \theta) \;=\; p^{\text{red}}_{\text{singlet}}(Z^{\text{root}}) \prod_{(v\to w)\in\mathbf{t}} \hat{P}^{\text{WFST}}(Z^w \mid Z^v, d_{vw}, \theta) \prod_{n:\mathcal{F}_n\neq\emptyset} L^{(\text{sub}),\text{tot}}_n(f_n; \mathcal{F}_n), \tag{C.133}$$

where $\mathcal{F}_n$ is the Fitch-parsimony subtree of column $n$, $L^{(\text{sub}),\text{tot}}_n(f; \mathcal{F}_n)$ is the class-marginalised Felsenstein column-substitution likelihood under fragchar $f$ (Section C.8.7), and $p^{\text{red}}_{\text{singlet}}, \hat{P}^{\text{WFST}}$ are the *reduced* singlet HMM and WFST defined on $(d, f)$ alone (Section C.8.4). As in the simple case we marginalise (C.133) over internal patterns supported on the $L$ observed columns to obtain $\tilde{p}(\text{MSA} \mid \mathbf{t}, \theta) \leq p(\text{MSA} \mid \mathbf{t}, \theta)$ (with the same ghost-column caveat), and bound the latter from below.

##### C.8.3 Variational family

The variational $q$ factorises over MSA columns, $q(\{Z^v_n\}) = \prod_{n=1}^{L} q_n$, and within each column,

$$q_n \;=\; q^{(\tau)}_n(\tau_n) \cdot q^{(\pi|\tau)}_n\big(\{Z^v_n\}_{v\in\mathcal{I}} \,\big|\, \tau_n\big), \tag{C.134}$$

where $q^{(\tau)}_n$ is a free categorical over $\mathcal{T}$ on the column-wide tuple, and $q^{(\pi|\tau)}_n$ is the same 3-state irreversible tree-structured graphical model of Section B.6.5 (with $xP \equiv \tau_n$). By construction $q$ assigns zero mass to within-column configurations in which Present nodes carry differing tuples — the column has one $(d, f)$, full stop. The cross-column structural-label constraints that forced a column-Markov chain in the labelled formulation *vanish* under reduction (Section C.8.9).

**Free parameter count.** Per column: $|\mathcal{T}| - 1 = N_{\text{dom}} N_{\text{fr}} - 1$ free parameters in $q^{(\tau)}_n$, plus $2|\mathcal{E}|$ in the inner 3-state graphical model (as in Section B.6.10), plus 2 for the inner root distribution. For typical $(N_{\text{dom}}, N_{\text{fr}}) = (3, 2)$ that is 5 tuple parameters per column, vs $4N_{\text{cl}}N_{\text{fr}}N_{\text{dom}} - 1$ in the labelled formulation (e.g. 239 for $N_{\text{cl}} = 10$) and $\sim |\mathcal{T}|^2$ in the labelled column-Markov upgrade ($\sim 6 \times 10^4$).

##### C.8.4 Reduced WFST: marginalising $(g, e)$ and the class $c$

The reduced per-character WFST kernel is

$$\hat{T}_{ss'}\big((d, f), (d', f');\, d, \theta\big), \qquad s, s' \in \{\texttt{S}, \texttt{M}, \texttt{I}, \texttt{D}, \texttt{E}\}, \tag{C.135}$$

obtained by marginalising the labelled-MixDom WFST $T^{\text{lab}}_{ss'}((c, f, g, d, e), (c', f', g', d', e'); d, \theta)$ over $(c, c', g, e, g', e')$ at each end. The reduced state space is $\{\texttt{S}, \texttt{E}\} \cup \{(s, d, f) : s \in \{\texttt{M}, \texttt{I}, \texttt{D}\}\}$ — i.e. $3N_{\text{dom}}N_{\text{fr}} + 2$ states.

**Routes from $(d, f)$ to $(d', f')$.** Under the labelled MixDom model, the per-character labelled transition $(d, f) \to (d', f')$ admits up to three latent routes, indexed by the value of $(g, e)$ at the source position:

R1. *Intra-fragment fragchar transition* $(g = 0)$: the current character is not the last of its fragment, and the next character stays in the same fragment with fragchar transitioning $f \to f'$ via $r^{(d)}_{f,f'}$. Restricted to $d' = d$.

R2. *New fragment, same domain* ($g = 1, e = 0$): the current fragment terminates ($\rho_f^{(d)}$) and the same domain starts a new fragment ($\kappa_d$) with first fragchar drawn from $w_{d,f'}$. Restricted to $d' = d$.

R3. *New domain* ($g = 1, e = 1$): the current fragment terminates, the current domain ends ($1-\kappa_d$), and (after possibly skipping geometrically-many empty domains) a new domain $d'$ begins with its first fragment's first fragchar drawn from $w_{d',f'}$. Available for any $d'$, including the self-recurrence $d' = d$.

For $d' \neq d$ only R3 is enabled, so the source's $(g, e) = (1, 1)$ is uniquely determined by the transition. For $d' = d$, multiple routes contribute and the latent $(g, e)$ at the source carries a non-trivial posterior over routes given the observed transition. The diagonal-extension special case ($r_{f,f'}^{(d)} = r_f \delta_{f,f'}$) suppresses R1 whenever $f' \neq f$, but R2 and R3 still mix for any $(d, f) \to (d, f')$, so the route-sum machinery introduced below is required even there. With a general off-diagonal $r^{(d)}$ matrix, R1 admits $f' \neq f$ transitions, so the route-sum acquires a third contributing term.

The per-route singlet contribution, accounting for the joint prior on $(g, e)$ at the source and the singlet emission of the next character, is:

$$\omega_{(d,f)\to(d',f')}^{(R1)} = \delta_{d,d'}\, r_{f,f'}^{(d)}, \tag{C.136}$$

$$\omega_{(d,f)\to(d',f')}^{(R2)} = \delta_{d,d'}\, \rho_f^{(d)}\, \kappa_d\, w_{d,f'}, \tag{C.137}$$

The per-branch expected indel log-likelihood factorises as

$$\mathbb{E}_q\big[\log P(X^w \mid X^v, d, \theta)\big] = \sum_{s,s'} \sum_{\tau,\tau' \in \mathcal{T}} W^{(v\to w)}_{ss',\tau\tau'} \log \hat{T}_{ss'}\big(\tau, \tau';\, d, \theta\big), \tag{C.148}$$

with the reduced expected transition counts

$$W^{(v\to w)}_{ss',\tau\tau'} \;=\; \sum_{N=1}^{L+1} \sum_{M=0}^{N-1} q^{(\tau)}_{n(M)}(\tau)\, q^{(\tau)}_{n(N)}(\tau')\, P^{v\to w}_{q,M}(s)\, P^{v\to w}_{q,N}(s') \prod_{K=M+1}^{N-1} P^{v\to w}_{q,K}(\mathsf{Ig}), \tag{C.149}$$

where the per-column factorisation of $q^{(\tau)}$ has reduced the pairwise tuple weight to a product of per-column marginals (no chain forward-backward needed). The cumulant trick of Section B.6.8 lifts unchanged to the inner $K$-product, with stable computation in $O(L \cdot |s|^2 \cdot |\mathcal{T}|^2)$ per branch via the running-logaddexp prefix on $\log P_q^{v\to w}(\mathsf{Ig})$. For $(N_{\text{dom}}, N_{\text{fr}}) = (3, 2)$ this gives $|\mathcal{T}|^2 = 36$ — versus $|\mathcal{T}^{\text{lab}}|^2 = 57600$ in the labelled variant ($\sim 1600\times$ cheaper per branch).

##### C.8.7 Per-column expected substitution log-likelihood

For each column $n$, define the *Fitch subtree* $\mathcal{F}_n \subseteq \mathbf{t}$ as the smallest connected subtree containing all leaves $v$ with $X_n^v = 1$. By stationarity and reversibility of the substitution model, only nodes inside $\mathcal{F}_n$ contribute to the substitution likelihood; nodes outside are treated as missing data and absorbed into normalisation. The Fitch subtree is determined by the leaf data alone, independently of $q$ and $\tau$.

Because $c$ is a per-column model latent (one class per column, governing all branches at that column), the column-substitution likelihood under fragchar $f$ marginalises $c$ at the model level:

$$L_n^{(\text{sub}),\text{tot}}(f; \mathcal{F}_n) \;=\; \sum_{c=1}^{N_{\text{cl}}} u_{f,c}\, L_n^{(\text{sub})}(c; \mathcal{F}_n), \tag{C.150}$$

with

$$L_n^{(\text{sub})}(c; \mathcal{F}_n) = \sum_{a \in \mathcal{A}} \pi_a^{(c)}\, \beta_n^{r_n}(a; c) \tag{C.151}$$

the standard Felsenstein up-pass likelihood at column $n$ on $\mathcal{F}_n$ under class-$c$'s rate matrix $R^{(c)}$ and stationary $\pi^{(c)}$ ($r_n$ is the Fitch-determined root of $\mathcal{F}_n$, $\beta_n^v(a; c)$ the standard Felsenstein up-message). The expected substitution log-likelihood under $q$ involves only the fragchar marginal $q_n^{(f)}(f) = \sum_d q_n^{(\tau)}(d, f)$:

$$\boxed{\mathbb{E}_{q_n^{(\tau)}}\big[\log L_n^{(\text{sub}),\text{tot}}(f_n; \mathcal{F}_n)\big] \;=\; \sum_{f=1}^{N_{\text{fr}}} q_n^{(f)}(f)\, \log\Big[\sum_c u_{f,c}\, L_n^{(\text{sub})}(c; \mathcal{F}_n)\Big].} \tag{C.152}$$

The presence factor $q_n^{(\pi|\tau)}$ does not enter — the substitution likelihood is fully determined by the (data-determined) Fitch subtree and the (variational-determined) fragchar marginal.

**Why $\log \sum_c uL$, not $\sum_c u \log L$.** This is the proper integration over the per-column class prior. A labelled-variant counterpart that carried $c$ as a variational latent would give $\sum_c q_n^{(c)}(c) \log L_n^{(\text{sub})}(c)$; by Jensen's inequality $\log \sum_c p_c L_c \geq \sum_c p_c \log L_c$ with $p_c = u_{f,c}$, equality holding only when $q_n^{(c)}$ is at the Bayes-optimal class posterior $q^*(c) \propto u_{f,c} L_n^{(\text{sub})}(c)$. The reduction analytically marginalises $c$ at the place where the labelled formulation left an inequality, so the reduced ELBO is at least as tight as the labelled ELBO and strictly tighter for any $q^{(c)}$ off the Bayes-optimal class posterior.

**Numerical implementation.** $\log L_n^{(\text{sub}),\text{tot}}(f)$ is computed in log-space:

$$\log L_n^{(\text{sub}),\text{tot}}(f) \;=\; \operatorname{logsumexp}_c\!\big(\log u_{f,c} + \log L_n^{(\text{sub})}(c; \mathcal{F}_n)\big), \tag{C.153}$$

with the expected log under $q_n^{(f)}$ a simple weighted sum.

##### C.8.8 ELBO

Combining the four contributions:

$$\boxed{\log p(\text{MSA} \mid \mathbf{t}, \theta) \;\geq\; \log \tilde{p}(\text{MSA} \mid \mathbf{t}, \theta) \;\geq\; \mathcal{L}[q],} \tag{C.154}$$

with

$$\mathcal{L}[q] = \mathbb{E}_q\big[\log p^{\text{red}}_{\text{singlet}}(Z^{\text{root}})\big] + \sum_{(v \to w) \in \mathbf{t}} \mathbb{E}_q\big[\log \hat{P}^{\text{WFST}}(Z^w \mid Z^v, d_{vw}, \theta)\big] + \sum_{n=1}^{L} \sum_f q_n^{(f)}(f) \, \log L_n^{(\text{sub}),\text{tot}}(f; \mathcal{F}_n) + H[q]. \tag{C.155}$$

**Single-class limit.** Setting $u_{f,c} = \delta_{c,c_0}$ for a fixed class $c_0$ gives $L_n^{(\mathrm{sub}),\mathrm{tot}}(f) = L_n^{(\mathrm{sub})}(c_0; \mathcal{F}_n)$, and the substitution term reduces to the standard Felsenstein log-likelihood under class $c_0$.

Base case:

$$F_{i,i,n,g} = T_{\text{tkf,start}}(n, i) \cdot w_{ng} \cdot E(n, i, g) \tag{C.159}$$

Recursion for $j > i$:

$$F_{i,j,n,g} = \sum_{f=1}^{\mathcal{F}} F_{i,j-1,n,f} \cdot (\delta(A(j-1)=A(j))\, T_{\text{ext}}(n,j,f,g) + T_{\text{notext}}(n,i,j,f,g)) \cdot E(n,j,g) \quad \text{(C.160)}$$

where we used $\sum_{n'} v_{n'}\kappa_{n'} = 1 - z_0^{(\mathrm{sing})}$ and $\zeta = \kappa_0 z_0^{(\mathrm{sing})}$. The sum over $g'$ and $e'$ is implicit in the fragment/domain generation that follows.

##### C.11.3 Labeled-MixDom WFST

*Remark* C.17 (WFST tables use the matrix-kernel notation). The transition tables below use the intra-fragment Markov kernel $r^{(n)}_{fg}$ (matrix indexed by source and destination fragchar) and its per-row termination $\rho^{(n)}_f = 1 - \sum_{f'} r^{(n)}_{ff'}$. Where a within-fragment self-loop appears, the relevant entry is the diagonal $r^{(n)}_{ff}$; where a new-fragchar-same-domain move appears, it is an off-diagonal $r^{(n)}_{ff'}$ for $f' \neq f$; the fragment-end weight is $\rho^{(n)}_f$.

- **Wait states** ("ready"): $\mathrm{W_M}$, $\mathrm{W_{D_F}}$, $\mathrm{W_{D_D}}$. These inspect the boundary indicators $(g, e)$ of the current context and choose the next emitting state based on the hierarchical boundary structure. Wait states also handle end-of-sequence.

In addition there are the non-emitting `S` and `E` states.

**States.** Each emitting or wait state carries a structural label context $\ell = (f, g, n, e)$. The full state space is:

$$\{\mathtt{S}\} \cup \{(X, f, g, n, e) : X \in \{\mathtt{M}, \mathrm{I_F}, \mathrm{I_D}, \mathrm{D_F}, \mathrm{D_D}, \mathrm{W_M}, \mathrm{W_{D_F}}, \mathrm{W_{D_D}}\}\} \cup \{\mathtt{E}\}$$

Not all combinations occur (see constraints below), but the upper bound is $8L + 2$ states.

**Context semantics.** The context $\ell = (f, g, n, e)$ on each state records:

- In `M` and $\mathrm{W_M}$: the label of the most recent matched character (which is the same for both input and output, since match preserves labels).
- In $\mathrm{I_F}$, $\mathrm{I_D}$: the label of the most recent *output* character.
- In $\mathrm{D_F}$, $\mathrm{D_D}$, $\mathrm{W_{D_F}}$, $\mathrm{W_{D_D}}$: the label of the most recent *input* character.

Thus only one context tuple is needed per state.

**Emitting-State Transitions (Unready → Wait)** Every emitting state makes a mandatory null transition (no input, no output) to its corresponding wait state. These transitions carry the boundary-survival weights from the nested TKF92/TKF91 structure.

| Source | Dest | Weight | Input | Output |
|---|---|---|---|---|
| $(\mathtt{M}, f, g, n, e)$ | $(\mathrm{W_M}, f, g, n, e)$ | $w_{\mathtt{M}}(g, e)$ | $\varepsilon$ | $\varepsilon$ |
| $(\mathrm{D_F}, f, g, n, e)$ | $(\mathrm{W_{D_F}}, f, g, n, e)$ | $w_{\mathrm{D_F}}(g, e)$ | $\varepsilon$ | $\varepsilon$ |
| $(\mathrm{D_D}, f, g, n, e)$ | $(\mathrm{W_{D_D}}, f, g, n, e)$ | $w_{\mathrm{D_D}}(g, e)$ | $\varepsilon$ | $\varepsilon$ |
| $(\mathrm{I_F}, f, g, n, e)$ | $(\mathrm{W_M}, f, g, n, e)$ | $w_{\mathrm{I_F}}(g, e)$ | $\varepsilon$ | $\varepsilon$ |
| $(\mathrm{I_D}, f, g, n, e)$ | $(\mathrm{W_M}, f, g, n, e)$ | $w_{\mathrm{I_D}}(g, e)$ | $\varepsilon$ | $\varepsilon$ |

Wait: the insert states need more careful treatment. An inserted fragment or domain is a complete sub-sequence generated by the descendant. The insert states $\mathrm{I_F}$ and $\mathrm{I_D}$ handle character-level emissions *within* an inserted fragment, and upon fragment/domain completion, control returns to the wait state that initiated the insertion. We therefore need to track whether we are inserting at the fragment level or domain level.

Let us reconsider the state structure more carefully.

**Revised State Structure** In the MixDom Pair HMM, the five state types `MM`, `MI`, `MD`, `II`, `DD` at each $(n, f)$ position represent:

- $\mathtt{MM}_{nf}$: ancestral and descendant both have a character (match/substitution)
- $\mathtt{MI}_{nf}$: descendant insertion within domain $n$, fragment $f$

- $\texttt{MD}_{nf}$: ancestral deletion within domain $n$, fragment $f$
- $\texttt{II}_{nf}$: insertion of an entire domain (both ancestor and descendant insert)
- $\texttt{DD}_{nf}$: deletion of an entire domain (both ancestor and descendant delete)

The top-level states $\texttt{M}, \texttt{I}, \texttt{D}$ refer to the *domain-level* TKF91 process, while the nested states $\texttt{M}, \texttt{I}, \texttt{D}$ refer to the *fragment-level* TKF92 process within a domain.

For the WFST, we separate the domain-level and fragment-level indel processes:

| Dest | Weight | Input | Output | Notes |
|---|---|---|---|---|
| $(\mathtt{M}, f, g', n, e)$ | $\alpha_n$ | $a_{c' f g' n e}$ | $b_{c' f g' n e}$ | match continues |
| $(\mathrm{D}_{\mathrm{F}}, f, g', n, e)$ | $(1 - \alpha_n)$ | $a_{c' f g' n e}$ | $\varepsilon$ | fragment-level deletion |

**Case $g = 1, e = 0$ (fragment boundary, mid-domain):** Fragment ended, new fragment within same domain, or insert/delete fragment, or end domain and transition at domain level.

| Dest | Weight | I | O | Notes |
|---|---|---|---|---|
| $(\mathtt{M}, f', g', n, e')$ | $\tau_{\mathtt{MM}}^{(n)} \cdot w_{nf'}$ | $a_{\ldots}$ | $b_{\ldots}$ | new frag, match |
| $(\mathrm{D}_{\mathrm{F}}, f', g', n, e')$ | $\tau_{\mathtt{MD}}^{(n)} \cdot w_{nf'}$ | $a_{\ldots}$ | $\varepsilon$ | new frag, delete |
| $(\mathrm{I}_{\mathrm{F}}, f', g', n, 0)$ | $\tau_{\mathtt{MI}}^{(n)} \cdot w_{nf'}$ | $\varepsilon$ | $b_{\ldots}$ | insert frag |
| $\mathtt{E}_{\text{domain}}$ | $\tau_{\mathtt{ME}}^{(n)}$ | | | domain ends (see below) |

##### C.11.4 Emission Weights

The emission weights for each emitting state type are:

| State | Emission | Weight |
|---|---|---|
| $\mathtt{M}$ | input $a_{cfgne}$, output $b_{cfgne}$ | $\exp(R_c T)_{ab}$ |
| $\mathrm{I_F}$ | output $b_{cfgne}$ | $u_{fc}\, \pi_{cb} \cdot p_g(r^{(n)}_{f\cdot})$ |
| $\mathrm{I_D}$ | output $b_{cfgne}$ | $u_{fc}\, \pi_{cb} \cdot p_g(r^{(n)}_{f\cdot})$ |
| $\mathrm{D_F}$ | input $a_{cfgne}$ | 1 |
| $\mathrm{D_D}$ | input $a_{cfgne}$ | 1 |

where $p_g(r^{(n)}_{f\cdot})$ stands for $r^{(n)}_{ff'}$ if $g = 0$ (fragment continues with destination fragchar $f'$) and $\rho^{(n)}_f$ if $g = 1$ (fragment ends), and the substitution matrix $R_c$ is the rate matrix for site class $c$.

For the WFST (conditional on ancestor), the delete emission weight is 1 because the ancestral emission probability is divided out (it was generated by the singlet HMM). The match emission divides out the ancestral $\pi_{ca}$ from the pair emission $\pi_{ca} \exp(R_c T)_{ab}$, yielding just the substitution matrix entry.

For insert states, the emission includes the full descendant character probability because there is no ancestral character to condition on.

##### C.11.5 Verification: Composition Reproduces the Pair HMM

**Claim.** The Labeled-MixDom Singlet HMM composed with the Labeled-MixDom WFST is equivalent to the MixDom Pair HMM defined in Section C.1.1.

**Sketch of proof.** The composition proceeds as follows:

1. The Singlet HMM generates the ancestral labeled sequence, with transitions governed by the MixDom stationary distribution.

2. The WFST reads the ancestral labeled sequence as input and produces the descendant labeled sequence as output.

3. The composed machine has states that are pairs (singlet state, WFST state). Since both are order-1 machines tracking structural labels, the composed state is a pair of structural labels.

We verify that the composed transition weights match $\chi$ for each case:

**Match-to-Match within a fragment ($g = 0$).** The singlet emits character $a_{cf0ne}$ with weight $u_{fc}\pi_{ca}$, and transitions to state $(f', g', n, e)$ with weight $r^{(n)}_{ff'}$ (if $g' = 0$) or $\rho^{(n)}_f \cdot \ldots$ (if $g' = 1$). The WFST in state $(\mathrm{W_M}, f, 0, n, e)$ reads this input and transitions to $(\mathtt{M}, g, g', n, e)$ with weight $\alpha_n$, emitting $b$ with substitution weight $\exp(R_cT)_{ab}$. The composed emission weight is $u_{gc}\pi_{ca}\exp(R_cT)_{ab}$, summing over $c$ gives $\sum_c u_{gc}\pi_{ca}\exp(R_cT)_{ab}$, which matches the Pair HMM emission for $\mathtt{MM}_{ng}$. The transition weight within the fragment is $r^{(n)}_{fg} \cdot \alpha_n$, which corresponds to the $\delta(l = m)r^{(n)}_{fg}$ term in $\chi$ (for the intra-fragment Markov contribution to $\mathtt{MM}_{lf} \to \mathtt{MM}_{lg}$).

**Match-to-Match across fragment boundary ($g = 1, e = 0$).** Singlet: fragment ends, transitions to new fragment $f'$ within same domain with weight $\kappa_n \cdot w_{nf'}$. WFST: $(\mathrm{W_M}, f, 1, n, 0)$ transitions to $(\mathtt{M}, f', g', n, e')$ with weight $\alpha_n \cdot w_{nf'}$. Combined: $\kappa_n \cdot w_{nf'} \cdot \alpha_n \cdot w_{nf'}$.

Wait—the $w$ appears twice, which is wrong. This reveals that the WFST transition weight should *not* include $w_{nf'}$ when the input character determines $f'$. The fragment type of the next input character is determined by the singlet HMM, and the WFST simply reads whatever fragment type appears.

Let us correct: at fragment boundaries, when the next action involves reading an input character, the WFST does *not* weight by $w_{nf'}$. The $w$ is part of the singlet distribution, not the conditional (WFST) distribution.

**Corrected Wait-State Transitions.** The WFST represents the *conditional* distribution $P(\text{descendant} \mid \text{ancestor})$. Therefore:

- Transitions that consume an input character should *not* include the prior probability of that input character's labels (such as $w$, $v$).
- Transitions that produce an output character (insertions) *do* include the full probability of the output labels (since the WFST generates them).
- Transitions involving the $\mathcal{T}$ matrix at domain boundaries must be adjusted: $\mathcal{T}$ was derived for the Pair HMM (joint distribution), so the WFST version divides out the ancestral prior factors.

Concretely, the $\chi$ entry for $\mathtt{MX}_{lf} \to \mathtt{MY}_{mg}$ with $l = m$ (same domain, new fragment) is:

$$\rho^{(n)}_f \cdot \tau^{(n)}_{\mathtt{XY}} \cdot w_{ng}$$

This is the *joint* weight. In the composed (singlet $\circ$ WFST) machine:

- Singlet provides: $\rho^{(n)}_f \cdot \kappa_n \cdot w_{ng}$ (end fragment, continue domain, choose new fragment type)
- WFST provides: $\frac{\tau^{(n)}_{\mathtt{XY}}}{\kappa_n}$ (the conditional transition, dividing out the $\kappa_n$ from the joint)

Product: $\rho^{(n)}_f \cdot \kappa_n \cdot w_{ng} \cdot \tau^{(n)}_{\mathtt{XY}}/\kappa_n = \rho^{(n)}_f \cdot \tau^{(n)}_{\mathtt{XY}} \cdot w_{ng}$. This matches the Pair HMM. ✓

Similarly, for the domain-boundary case with $l \neq m$:

- Singlet provides: $\rho^{(l)}_f(1 - \kappa_l) \cdot \frac{\kappa_0 v_m \kappa_m w_{mg}}{1-\zeta}$
- WFST provides: the conditional weight that, when multiplied by the singlet weight, gives $\chi_{\mathtt{MX}_{lf} \to \mathtt{MY}_{mg}}$

**$\mathrm{W}_{\mathrm{M}}$, $g = 1, e = 1$ (domain boundary):**
With input (start new ancestral domain):

| Dest | WFST weight | Notes |
|---|---|---|
| `M` | $\dfrac{\mathcal{T}_{\mathtt{MM}} \cdot v_{n'} \cdot \tau^{(n')}_{\mathtt{SY}} \cdot w_{n'f'}}{P_{\text{sing}}(\ell \to \ell')}$ | new domain, match |
| $\mathrm{D}_{\mathrm{F}}$ | $\dfrac{\mathcal{T}_{\mathtt{MM}} \cdot v_{n'} \cdot \tau^{(n')}_{\mathtt{SD}} \cdot w_{n'f'}}{P_{\text{sing}}(\ell \to \ell')}$ | new domain, del-frag |
| $\mathrm{D}_{\mathrm{D}}$ | $\dfrac{\mathcal{T}_{\mathtt{MD}} \cdot v_{n'} \cdot \kappa_{n'} \cdot w_{n'f'}}{P_{\text{sing}}(\ell \to \ell')}$ | delete domain |

Without input:

| Dest | WFST weight | Notes |
|---|---|---|
| $\mathrm{I}_{\mathrm{D}}$ | $\mathcal{T}_{\mathtt{MI}} \cdot v_{n'} \cdot \kappa_{n'} \cdot w_{n'f'} \cdot u_{f'c'}\pi_{c'b} \cdot p_{g'}(r_{f'})$ | insert domain |
| `E` | $\mathcal{T}_{\mathtt{ME}}/P_{\text{sing}}(\ell \to \mathtt{E})$ | end |

where $P_{\text{sing}}(\ell \to \ell') = \frac{\kappa_0 v_{n'} \kappa_{n'} w_{n'f'}}{1-\zeta}$ is the singlet transition from a domain boundary to the next character label $\ell'$ (Equation C.180).

The $\mathrm{W}_{\mathrm{D}_{\mathrm{F}}}$ and $\mathrm{W}_{\mathrm{D}_{\mathrm{D}}}$ tables follow the same pattern, using $\mathcal{T}_{\mathtt{M}\cdot}$ for $\mathrm{W}_{\mathrm{D}_{\mathrm{F}}}$ (since fragment deletion is within a matched domain) and $\mathcal{T}_{\mathtt{D}\cdot}$ for $\mathrm{W}_{\mathrm{D}_{\mathrm{D}}}$ (since the domain is being deleted).

**$\mathrm{W}_{\mathrm{D}_{\mathrm{F}}}$ at $g = 0$:**

| Dest | WFST weight |
|---|---|
| $\mathrm{D}_{\mathrm{F}}$ | 1 |

**$\mathrm{W}_{\mathrm{D}_{\mathrm{F}}}$ at $g = 1, e = 0$:**

| Dest | WFST weight |
|---|---|
| `M` | $\alpha_n$ |
| $\mathrm{D}_{\mathrm{F}}$ | $(1 - \alpha_n)$ |

**$\mathrm{W}_{\mathrm{D}_{\mathrm{F}}}$ at $g = 1, e = 1$:** Same structure as $\mathrm{W}_{\mathrm{M}}$ at $g = 1, e = 1$, using $\mathcal{T}_{\mathtt{M}\cdot}$.

**$\mathrm{W}_{\mathrm{D}_{\mathrm{D}}}$ at $g = 0$:**

| Dest | WFST weight |
|---|---|
| $\mathrm{D}_{\mathrm{D}}$ | 1 |

$W_{D_D}$ **at** $g = 1, e = 0$**:**

| Dest | WFST weight |
|---|---|
| $D_D$ | 1 |

(Within a deleted domain, all fragments are consumed; fragment type is irrelevant.)

$W_{D_D}$ **at** $g = 1, e = 1$**:**

| Dest | WFST weight | Notes |
|---|---|---|
| M | $\mathcal{T}_{\mathtt{DM}} \cdot v_{n'} \cdot \tau^{(n')}_{\mathtt{SY}} \cdot w_{n'f'} / P_{\text{sing}}(\ell \to \ell')$ | new domain, match |
| $D_F$ | $\mathcal{T}_{\mathtt{DM}} \cdot v_{n'} \cdot \tau^{(n')}_{\mathtt{SD}} \cdot w_{n'f'} / P_{\text{sing}}(\ell \to \ell')$ | new domain, del-frag |
| $D_D$ | $\mathcal{T}_{\mathtt{DD}} \cdot v_{n'} \cdot \kappa_{n'} \cdot w_{n'f'} / P_{\text{sing}}(\ell \to \ell')$ | delete domain |
| $I_D$ | $\mathcal{T}_{\mathtt{DI}} \cdot$ (full output prob) | insert domain |
| E | $\mathcal{T}_{\mathtt{DE}} / P_{\text{sing}}(\ell \to \mathtt{E})$ | end |

$I_D$ **completion at** $g = 1, e = 1$**:**

| Dest | WFST weight | Notes |
|---|---|---|
| M | $\mathcal{T}_{\mathtt{IM}} \cdot v_{n'} \cdot \tau^{(n')}_{\mathtt{SY}} \cdot w_{n'f'} / P_{\text{sing}}(\ell \to \ell')$ | match next domain |
| $D_D$ | $\mathcal{T}_{\mathtt{ID}} \cdot v_{n'} \cdot \kappa_{n'} \cdot w_{n'f'} / P_{\text{sing}}(\ell \to \ell')$ | delete next domain |
| $I_D$ | $\mathcal{T}_{\mathtt{II}} \cdot$ (full output prob) | insert another domain |
| E | $\mathcal{T}_{\mathtt{IE}} / P_{\text{sing}}(\ell \to \mathtt{E})$ | end |

##### C.11.6 Simplification of Domain-Boundary WFST Weights

The domain-boundary WFST weights (Section C.11.3) involve a ratio of the Pair HMM transition weight to the singlet transition weight. This simplifies considerably.

For a transition that consumes input with label $\ell' = (f', g', n', e')$ from a domain boundary ($g = 1, e = 1$), the singlet weight is (from Equation C.180):

$$P_{\text{sing}}(\ell \to \ell') = \frac{\kappa_0 \cdot v_{n'} \cdot \kappa_{n'} \cdot w_{n'f'}}{1 - \zeta}$$

The Pair HMM weight for $\mathtt{MM}_{lf} \to \mathtt{MM}_{mg}$ with $l \neq m$ (different domain, going through domain boundary) is:

$$\rho^{(l)}_f \cdot \tau^{(l)}_{\mathtt{ME}} \cdot \mathcal{T}_{\mathtt{MM}} \cdot v_m \cdot \tau^{(m)}_{\mathtt{SM}} \cdot w_{mg}$$

The WFST weight is therefore:

$$\frac{\rho^{(l)}_f \cdot \tau^{(l)}_{\mathtt{ME}} \cdot \mathcal{T}_{\mathtt{MM}} \cdot v_m \cdot \tau^{(m)}_{\mathtt{SM}} \cdot w_{mg}}{\frac{\kappa_0 \cdot v_m \cdot \kappa_m \cdot w_{mg}}{1-\zeta}} = \frac{(1-\zeta)\rho^{(l)}_f \cdot \tau^{(l)}_{\mathtt{ME}} \cdot \mathcal{T}_{\mathtt{MM}} \cdot \tau^{(m)}_{\mathtt{SM}}}{\kappa_0 \cdot \kappa_m}$$

*Example* C.1 (TKF91). Applying left CTMC expansion to `MOR` in the link grammar $\mathcal{G}_{\text{link}}(\kappa)$ with alphabet $\mathcal{A}$ and equilibrium distribution $\pi$ yields:

| | |
|---|---|
| $\texttt{IMM} \to \texttt{MOR}\ \texttt{IMM}$ | $\kappa$ |
| $\texttt{IMM} \to \epsilon$ | $1 - \kappa$ |
| $\texttt{MOR} \to c\ \texttt{MOR}\ \texttt{MOR}$ | $\kappa \cdot \pi_c \quad (c \in \mathcal{A})$ |
| $\texttt{MOR} \to c$ | $(1 - \kappa) \cdot \pi_c \quad (c \in \mathcal{A})$ |

*Every production $Y \to \alpha\ X\ \beta$ in $\mathcal{G}$ (where $Y \neq X$ and $\alpha, \beta \in (\mathcal{N} \cup \Sigma)^*$) is replaced by $K$ productions:*

$$Y \to \alpha\ X_k\ \beta \qquad \text{weight } w(Y \to \alpha X \beta)\cdot p_k \quad \text{for } k = 1,\ldots,K \tag{C.191}$$

*Example* C.5 (TKF Genome: region concatenation). The TKF Genome's top-level grammar has:

$$
\begin{array}{ll}
\texttt{GENOME} \to \texttt{REGION GENOME} & 1-\kappa_R \\
\texttt{GENOME} \to \epsilon & \kappa_R \\
\texttt{REGION} \to \texttt{INTER} & p_N \\
\quad\quad\quad\;\mid \texttt{FWDCDS} & p_G/2 \\
\quad\quad\quad\;\mid \texttt{REVCDS} & p_G/2 \\
\quad\quad\quad\;\mid \texttt{STRUCT} & p_S \\
\quad\quad\quad\;\mid \texttt{CONS} & p_C
\end{array}
$$

*The* right recursive nesting *of* $\texttt{BIF}$ *at* $\texttt{MOR}$ *replaces each terminal-emitting production of* $\texttt{MOR}$ *with a mixture of terminal emission and bifurcation:*
***Before***:

$$\texttt{MOR} \to c\ \alpha \qquad\qquad w_{\text{orig}} \cdot \pi_c$$

***After***:

$$\begin{array}{lll} \texttt{MOR} \to c\ \alpha & w_{\text{orig}} \cdot s \cdot \pi_c & \textit{(terminal link, probability } s\textit{)} \\ \texttt{MOR} \to \texttt{BIF}\ \alpha & w_{\text{orig}} \cdot (1-s) \cdot v_{\text{bif}} & \textit{(bifurcation, probability } 1-s\textit{)} \end{array}$$

*where* $s \in (0,1]$ *is the probability that a link is a terminal (character-emitting) link rather than a nesting point, and* $v_{\text{bif}}$ *is a distribution over bifurcation types if there are multiple* $\texttt{BIF}$ *variants.*

*For* left recursive nesting*:*

$$\texttt{MOR} \to \alpha\ \texttt{BIF} \qquad\qquad w_{\text{orig}} \cdot (1-s) \cdot v_{\text{bif}}$$

*The sub-grammar* $\mathcal{G}_{\text{bif}}$ *for* $\texttt{BIF}$ *defines how the bifurcation expands. Since it may reference the start symbol of* $\mathcal{G}$ *(e.g.,* $\texttt{IMM}$*), the combined grammar is genuinely recursive: link sequences can contain nested link sequences of arbitrary depth.*

*Remark* C.25. The recursive nesting creates null cycles whenever the nested sub-grammar can generate the empty string. These must be handled by the null-state management procedures (Section C.12.3). In the TKF Structure Tree, the nullability fixed-point iteration solves for the probability that each nonterminal generates $\epsilon$.

*Example* C.7 (TKF Structure Tree: stems and loops). The TKF Structure Tree has two types of link sequences:

- *Loop sequences* ($L$): left-emitting links generating single nucleotides. Nonterminal rule: $L \to c_L\ L$ with CTMC expansion (left emission).
- *Stem sequences* ($S$): LR-emitting links generating basepairs. Nonterminal rule: $S \to c_L\ S\ c_R$ with CTMC expansion (LR emission).

The recursive nesting works as follows. Within a loop sequence, a mortal link may either emit a character (probability $s_L$) or spawn a stem (probability $1 - s_L$):

$$L \to c\ L \quad | \quad S\ L$$

At the base of a stem (when the self-loop terminates), the grammar transitions to a loop:

$$S \to c_L\ S\ c_R \quad | \quad L$$

This creates the alternating stem–loop structure characteristic of RNA secondary structure. The recursion arises because a loop can spawn a stem, which eventually returns to a loop, which can spawn another stem, *ad infinitum*.

*Example* C.8 (TKF92 with Recursive Domains). The recursive domain model has nonterminals $\texttt{L}_n$ (link sequence for domain $n$), $\texttt{A}_n$ (aligned component), $\texttt{I}_n$ (inserted component), $\texttt{D}_n$ (deleted component), etc. The aligned component rule is:

$$\begin{array}{lll} \texttt{A}_n \to c & s_n \cdot \pi_{n,c} & \text{(terminal: character)} \\ \texttt{A}_n \to \texttt{L}_{n'} & (1-s_n) \cdot v_{nn'} & \text{(bifurcation: nested link sequence)} \end{array}$$

Since $\texttt{L}_{n'}$ can reference $\texttt{L}_n$ (if $n' = n$ or through a chain of domain transitions), this creates genuine recursion: link sequences contain domains that contain further link sequences.

1. For each non-null production $\mathtt{X}' \to \alpha'$ in the $\epsilon$-eliminated grammar, its posterior count $c'(\mathtt{X}' \to \alpha')$ contributes directly to the corresponding original rule.

2. For each "nullability shortcut" production $\texttt{X}' \to \texttt{Y}'$ (arising from $\texttt{X} \to \texttt{Y Z}$ with $\texttt{Z}$ nullable), the count $c'(\texttt{X}' \to \texttt{Y}')$ contributes $c'(\texttt{X}' \to \texttt{Y}')$ to the original rule $\texttt{X} \to \texttt{Y Z}$, plus $c'(\texttt{X}' \to \texttt{Y}') \cdot \bar{c}(\texttt{Z} \to \alpha)$ to each rule within $\texttt{Z}$'s null derivation subtree.

The general theory of $\epsilon$-elimination with EM count recovery, including the null closure $(I - T_{\mathcal{Z}\mathcal{Z}})^{-1}$ and null cycle removal, is developed in Appendix C.12 (Section C.12.3)

First we find the nullability $\eta(\texttt{X})$ of each nonterminal $\texttt{X}$. The nullability is the probability that a parse tree rooted in that nonterminal yields the empty string. We can't solve for these in closed form (at least not in the general recursive model, where a link sequence can contain another link sequence of the same type). Instead we can solve approximately by iterating towards a fixed point.

We first observe that the nullabilities collectively satisfy the following

$$
\begin{aligned}
\eta(\texttt{L}_l) &= \eta(\texttt{K}_l) \frac{1-\kappa_l}{1-\kappa_l \eta(\texttt{M}_l)} \\
\eta(\texttt{K}_l) &= 1 - \beta_l + \beta_l \eta(\texttt{N}_l) \\
\eta(\texttt{M}_l) &= \alpha_l \eta(\texttt{S}_l) + (1-\alpha_l)\eta(\texttt{E}_l) \\
\eta(\texttt{S}_l) &= \eta(\texttt{A}_l)\left(\beta_l \eta(\texttt{N}_l) + (1-\beta_l)\right) \\
\eta(\texttt{E}_l) &= \eta(\texttt{D}_l)\left(\gamma_l \eta(\texttt{N}_l) + (1-\gamma_l)\right) \\
\eta(\texttt{N}_l) &= \eta(\texttt{I}_l) \frac{1-\beta_l}{1-\beta_l \eta(\texttt{I}_l)} \\
\eta(\texttt{A}_l) &= (1-s_l) \sum_m v_{lm} \eta(\texttt{L}_m) \\
\eta(\texttt{I}_l) &= (1-s_l) \sum_m v_{lm} \eta(\texttt{C}_m) \\
\eta(\texttt{D}_l) &= \eta(\texttt{I}_l) \\
\eta(\texttt{C}_l) &= \frac{1-\kappa_l}{1-\kappa_l \eta(\texttt{I}_l)} \\
\eta(\texttt{P}_l) &= \eta(\texttt{C}_l)
\end{aligned}
$$

A general procedure is (i) iterate to solve for the $\eta(\texttt{C}_l)$ (substituting in the downstream definition of $\eta(\texttt{I}_l)$ so the formula becomes self-referential); (ii) this directly yields $\eta(\texttt{P}_l), \eta(\texttt{I}_l), \eta(\texttt{D}_l), \eta(\texttt{N}_l), \eta(\texttt{E}_l), \eta(\texttt{K}_l)$; (iii) iterate to solve for $\eta(\texttt{L}_l)$ (again, first substituting to make it self-referential); (iv) this directly yields the remaining $\eta(\texttt{M}_l), \eta(\texttt{S}_l), \eta(\texttt{A}_l)$.

In detail: initialize $x_l^{(0)} \leftarrow 0$ for $l \in \mathcal{N}$. Iterate to convergence

$$
x_l^{(k+1)} \leftarrow \frac{1-\kappa_l}{1-\kappa_l(1-s_l)\sum_m v_{lm} x_m^{(k)}}
$$

We then set $\eta(\texttt{C}_l) \leftarrow \lim_{k\to\infty} x_l^{(k)}$ and set $\eta(\texttt{P}_l), \eta(\texttt{I}_l), \eta(\texttt{D}_l), \eta(\texttt{N}_l), \eta(\texttt{E}_l), \eta(\texttt{K}_l)$ using the above equations. Now set $y_l^{(0)} \leftarrow 0$ for $l \in \mathcal{N}$ and again iterate

$$
y_l^{(k+1)} \leftarrow \frac{(1-\kappa_l)\eta(\texttt{K}_l)}{1-\kappa_l(1-\alpha_l)\eta(\texttt{E}_l) - \kappa_l \alpha_l \left(1-\beta_l(1-\eta(\texttt{N}_l))\right)(1-s_l)\sum_m v_{lm} y_m^{(k)}}
$$

Then set $\eta(\texttt{L}_l) \leftarrow \lim_{k\to\infty} y_l^{(k)}$ and set the remaining $\eta(\texttt{M}_l), \eta(\texttt{S}_l), \eta(\texttt{A}_l)$ using the above equations.

| lhs | $\to$ | rhs | $P(b)$ |
|---|---|---|---|
| $L_b$ | $\to$ | $w\ L_b$ | $\kappa_l \pi_l(w)$ |
| | $\mid$ | $S_b\ L_b$ | $\kappa_l \pi_l(S)$ |
| | $\mid$ | $\epsilon$ | $1 - \kappa_l$ |
| $S_b$ | $\to$ | $w\ S_b\ x$ | $\kappa_s \pi_s(wx)$ |
| | $\mid$ | $L_b$ | $1 - \kappa_s$ |

Table C.3: TKF Structure Tree. Singlet rule-set for $b$.

**The Stationary Grammar**

$$\mathsf{SL} \to \mathsf{STEM} \qquad \text{weight } 1 \tag{C.198}$$

**Stem (TKF91 sequence of stem-links, from outer to inner):**

$$\mathsf{STEM} \to c_L\ \mathsf{STEM}\ c_R \qquad \kappa_S\, p_{\text{bp}}\, \pi_{\text{bp}}(c_L, c_R) \qquad \text{[single basepair, LR]} \tag{C.199}$$

$$\to c_L^1\, c_L^2\ \mathsf{STEM}\ c_R^2\, c_R^1 \qquad \kappa_S\, p_{\text{st}}\, (1 - r_K)\, \pi_K(\cdot) \qquad \text{[terminal stacked pair, LLRR]} \tag{C.200}$$

$$\to c_L^1\, c_L^2\ \mathsf{STACK}\ c_R^2\, c_R^1 \qquad \kappa_S\, p_{\text{st}}\, r_K\, \pi_K(\cdot) \qquad \text{[extended stacked pair, LLRR]} \tag{C.201}$$

$$\to \mathsf{LDECO}\ \mathsf{STEM}\ \mathsf{RDECO} \qquad \kappa_S\, p_{\text{bu}} \qquad \text{[bulge]} \tag{C.202}$$

$$\to \mathsf{CLOSE}\ \mathsf{LOOP} \qquad 1 - \kappa_S \qquad \text{[end stem]} \tag{C.203}$$

A bulge link (C.202) represents an internal loop, a single-sided bulge, or a multihelix junction branch between basepairs. All non-basepair content between consecutive basepairs is consolidated into a single TKF link type whose internal structure is governed by $\mathsf{LDECO}$ and $\mathsf{RDECO}$.

**Bulge decorations (L-side and R-side content):**

$$\mathsf{LDECO} \to \mathsf{LFRAG}\ \mathsf{LDECO} \qquad \text{[L-fragment, then more L-content]} \tag{C.204}$$

$$\to \mathsf{SL}\ \mathsf{LDECO} \qquad \text{[left branch: nested stem-loop]} \tag{C.205}$$

$$\to \epsilon \qquad \text{[end L-decorations]} \tag{C.206}$$

$$\mathsf{RDECO} \to \mathsf{RDECO}\ \mathsf{RFRAG} \qquad \text{[R-fragment, more R-content]} \tag{C.207}$$

$$\to \mathsf{RDECO}\ \mathsf{SL} \qquad \text{[right branch: nested stem-loop]} \tag{C.208}$$

$$\to \epsilon \qquad \text{[end R-decorations]} \tag{C.209}$$

$\mathsf{LDECO}$ generates all content on the $5'$ side (between the outer basepair's L-half and the continuation), while $\mathsf{RDECO}$ generates all content on the $3'$ side (between the continuation and the outer basepair's R-half). An internal loop has both $\mathsf{LDECO} \neq \epsilon$ and $\mathsf{RDECO} \neq \epsilon$; a single-sided bulge has content on only one side; a multihelix junction branch has $\mathsf{SL}$ nested within $\mathsf{LDECO}$ or $\mathsf{RDECO}$.

**Stacked-pair fragment continuation (LLRR emission):**

$$\mathsf{STACK} \to c_L^1\, c_L^2\ \mathsf{STEM}\ c_R^2\, c_R^1 \qquad (1 - r_K)\, \pi_K(\cdot) \qquad \text{[terminal]} \tag{C.210}$$

$$\to c_L^1\, c_L^2\ \mathsf{STACK}\ c_R^2\, c_R^1 \qquad r_K\, \pi_K(\cdot) \qquad \text{[extend]} \tag{C.211}$$

**Closing basepair (LR emission):**

$$\mathsf{CLOSE} \to c_L\ \mathsf{CLOSE}'\ c_R \qquad \pi_{\text{cl}}(c_L, c_R) \tag{C.212}$$

where $\mathsf{CLOSE}'$ is a unit nonterminal ($\mathsf{CLOSE}' \to \epsilon$, weight 1) that serves as a placeholder for the inside of the closing basepair.

**Loop (TKF91 sequence of loop-links):**

$$\mathsf{LOOP} \to \mathsf{LOOPLINK}\ \mathsf{LOOP} \qquad \kappa_L \qquad \text{[add loop link]} \tag{C.213}$$

$$\to \epsilon \qquad 1 - \kappa_L \qquad \text{[end loop]} \tag{C.214}$$

**Loop link types:**

$$\mathsf{LOOPLINK} \to \mathsf{LFRAG} \quad p_{\text{lf}} \quad \text{[L-fragment]} \tag{C.215}$$

$$\to \mathsf{RFRAG} \quad p_{\text{rf}} \quad \text{[R-fragment]} \tag{C.216}$$

$$\to \mathsf{SL} \quad p_{\text{sl}} \quad \text{[nested stem-loop (multiloop)]} \tag{C.217}$$

**Unpaired nucleotide fragments in loops (geometric length $\geq 1$):**

$$\mathsf{LFRAG} \to c\ \mathsf{LFRAG} \quad r_L\,\pi(c) \quad \text{[extend, L-emission]} \tag{C.218}$$

$$\to c \quad (1-r_L)\,\pi(c) \quad \text{[terminal, L-emission]} \tag{C.219}$$

$$\mathsf{RFRAG} \to \mathsf{RFRAG}\ c \quad r_R\,\pi(c) \quad \text{[extend, R-emission]} \tag{C.220}$$

$$\to c \quad (1-r_R)\,\pi(c) \quad \text{[terminal, R-emission]} \tag{C.221}$$

**Stem Link Elaboration.** The stem is a TKF91 sequence of links with rates $(\lambda_S, \mu_S)$. Each STEM nonterminal becomes $\mathsf{STEM}_{\mathtt{M}}$ (post-match/insert) and $\mathsf{STEM}_{\mathtt{D}}$ (post-delete):

$$\mathsf{STEM}_{\mathtt{M}} \to c_L^x\, c_L^y\, \mathsf{STEM}_{\mathtt{M}}\, c_R^y\, c_R^x \quad (1-\beta_S)\kappa_S\, p_{\mathrm{bp}}\, \alpha_S\, \pi_{\mathrm{bp}}(c_L^x, c_R^x)\, P_{\mathrm{bp}}(c_L^y, c_R^y | c_L^x, c_R^x) \tag{C.222}$$

$$\to c_L^y\, \mathsf{STEM}_{\mathtt{M}}\, c_R^y \quad \beta_S\, \kappa_S\, p_{\mathrm{bp}}\, \pi_{\mathrm{bp}}(c_L^y, c_R^y) \quad \text{[BP insert]} \tag{C.223}$$

$$\to c_L^x\, \mathsf{STEM}_{\mathtt{D}}\, c_R^x \quad (1-\beta_S)\kappa_S\, p_{\mathrm{bp}}\, (1-\alpha_S)\, \pi_{\mathrm{bp}}(c_L^x, c_R^x) \quad \text{[BP delete]} \tag{C.224}$$

with analogous rules for $\mathsf{STEM}_{\mathtt{D}}$ using $\gamma_S$ in place of $\beta_S$.

For matched single basepairs (C.222), the nesting is $c_L^x\, c_L^y\ \cdots\ c_R^y\, c_R^x$: ancestor terminals on the outside, descendant terminals on the inside, preserving palindromic structure. For inserted basepairs (C.223), only descendant terminals appear (LR emission). For deleted basepairs (C.224), only ancestor terminals appear (LR emission).

**Stacked-Pair Elaboration.** Matched stacked pairs have LLRR emission on both ancestor and descendant, yielding an $L^4R^4$ nesting pattern in the pair grammar:

$$\begin{aligned} \mathsf{STEM}_{\mathtt{M}} \to\ & c_{L1}^x\, c_{L2}^x\, c_{L1}^y\, c_{L2}^y\, \mathsf{STEM}_{\mathtt{M}}\, c_{R2}^y\, c_{R1}^y\, c_{R2}^x\, c_{R1}^x \\ & (1-\beta_S)\kappa_S\, p_{\mathrm{st}}\, (1-r_K)\, \alpha_S\, \pi_K(\cdot)\, P_K(\cdot|\cdot) \end{aligned} \tag{C.225}$$

with inserted stacked pairs emitting only descendant LLRR, and deleted stacked pairs emitting only ancestor LLRR.

**Bulge Elaboration.** A bulge link (C.202) gains M/I/D versions like any other stem link. A matched bulge has the form $\mathsf{LDECO}_{\mathtt{M}}\ \mathsf{STEM}_{\mathtt{M}}\ \mathsf{RDECO}_{\mathtt{M}}$, with each side elaborated independently.

Within LDECO, each sub-element (LFRAG, nested SL) gains M/I/D versions with L-emission direction preserved. For example, a matched left-branch within a bulge produces:

$$\mathsf{LDECO}_{\mathtt{M}} \to \mathsf{SL}_{\mathtt{M}}\ \mathsf{LDECO}_{\mathtt{M}} \quad \alpha_{\mathrm{deco}}\, P(\mathrm{branch}) \quad \text{[matched left branch]} \tag{C.226}$$

$$\to \mathsf{SL}_{\mathtt{I}}\ \mathsf{LDECO}_{\mathtt{M}} \quad \beta_{\mathrm{deco}} \quad \text{[inserted left branch]} \tag{C.227}$$

$$\to \mathsf{SL}_{\mathtt{D}}\ \mathsf{LDECO}_{\mathtt{D}} \quad (1-\alpha_{\mathrm{deco}})\, P(\mathrm{branch}) \quad \text{[deleted left branch]} \tag{C.228}$$

Here $\mathsf{SL}_{\mathtt{M}}$ generates aligned ancestor–descendant sub-structures, $\mathsf{SL}_{\mathtt{I}}$ generates descendant-only sub-structures, and $\mathsf{SL}_{\mathtt{D}}$ generates ancestor-only sub-structures.

RDECO is elaborated symmetrically, with R-emission direction preserved for RFRAG elements and right-side branches.

**Loop Elaboration.** The loop is a TKF91 sequence of loop-links with rates $(\lambda_L, \mu_L)$. Elaboration proceeds identically to the stem: each LOOP becomes $\mathsf{LOOP}_\mathtt{M}$/$\mathsf{LOOP}_\mathtt{D}$, and each loop-link type (LFRAG, RFRAG, nested SL) gains `M`/`I`/`D` versions.

Nested stem-loops within the loop (multiloop junctions) are handled recursively: $\mathsf{SL}_\mathtt{M}$ aligns both ancestor and descendant sub-structures, allowing multiloop branches to be independently inserted, deleted, or matched.

##### C.13.4 Example Four: The TKF Genome

**Parameters** Let $\overline{xyz}$ denote reverse complement e.g. $\overline{AAG} = CTT$.

Parameters: insertion and deletion rates $\lambda_F < \mu_F$, fragment extension probability $r_F$, substitution rate matrix $Q_F$, equilibrium probability vector $q_F$ (so $q_x Q_F = 0$) for $F \in \{R, N, G, S, L, C\}$. Splice donor/acceptor site distribution $q_{D1}$, $q_{D2}$, $q_{A1}$, $q_{A2}$. Region-type probabilities $p_G + p_N + p_S + p_C = 1$. Intron probability $p_I$.

The $Q_N$, $Q_S$, $Q_L$ and $Q_C$ models should be strand-invariant, so e.g. $Q_N(x_1, x_2) = Q_N(\overline{x_1}, \overline{x_2})$.

**Functions** For $F \in \{R, N, G, S, L, C\}$:

$$\kappa_F \quad = \quad \left(1 - \frac{\lambda_F}{\mu_F}\right)(1 - r_F)$$

**The Stationary Grammar**

| LHS | $\rightarrow$ | RHS | Transition | Emission |
|---|---|---|---|---|
| F | $\rightarrow$ | L F R | $1-\kappa_F$ | $q_F(LR)$ |
| | $\mid$ | E | $\kappa_F$ | |

...that is, for GENOME, INTER, FWDCDS, REVCDS, STRUCT, LOOP, and CONS (with $F \in \{R, N, G, S, L, C\}$), replace these rules with

| LHS | $\to$ | RHS | Transition | Emission |
|---|---|---|---|---|
| $\mathtt{F}_M$ | $\to$ | $\mathtt{L}_M$ $\mathtt{F}_M$ $\mathtt{R}_M$ | $(1-\beta_{M,F})(1-\kappa_F)\alpha_F$ | $q_F(LR_x)\exp(Q_F t)(LR_x, LR_y)$ |
| | $\vert$ | $\mathtt{L}_Y$ $\mathtt{F}_M$ $\mathtt{R}_Y$ | $\beta_{M,F}$ | $q_F(LR_y)$ |
| | $\vert$ | $\mathtt{L}_X$ $\mathtt{F}_D$ $\mathtt{R}_X$ | $(1-\beta_{M,F})(1-\kappa_F)(1-\alpha_F)$ | $q_F(LR_x)$ |
| | $\vert$ | $\mathtt{E}_M$ | $(1-\beta_{M,F})\kappa_F$ | |
| $\mathtt{F}_D$ | $\to$ | $\mathtt{L}_M$ $\mathtt{F}_M$ $\mathtt{R}_M$ | $(1-\beta_{D,F})(1-\kappa_F)\alpha_F$ | $q_F(LR_x)\exp(Q_F t)(LR_x, LR_y)$ |
| | $\vert$ | $\mathtt{L}_Y$ $\mathtt{F}_M$ $\mathtt{R}_Y$ | $\beta_{D,F}$ | $q_F(LR_y)$ |
| | $\vert$ | $\mathtt{L}_X$ $\mathtt{F}_D$ $\mathtt{R}_X$ | $(1-\beta_{D,F})(1-\kappa_F)(1-\alpha_F)$ | $q_F(LR_x)$ |
| | $\vert$ | $\mathtt{E}_M$ | $(1-\beta_{D,F})\kappa_F$ | |

...that is, two versions $\mathtt{F}_M$ and $\mathtt{F}_D$, with different $\beta$'s for each type.

For every other nonterminal N, there need to be three versions $\mathtt{N}_M$, $\mathtt{N}_X$ and $\mathtt{N}_Y$, with outgoing rules for each type going to other nonterminals of the same type.

##### Top level

| LHS | $\to$ | RHS | Transition | Emission |
|---|---|---|---|---|
| $\mathtt{GENOME}_k$ | $\to$ | $\mathtt{REGION}_M$ $\mathtt{GENOME}_M$ | $(1-\beta_{k,R})(1-\kappa_R)\alpha_R$ | |
| | $\vert$ | $\mathtt{REGION}_Y$ $\mathtt{GENOME}_M$ | $\beta_{k,R}$ | |
| | $\vert$ | $\mathtt{REGION}_X$ $\mathtt{GENOME}_D$ | $(1-\beta_{k,R})(1-\kappa_R)(1-\alpha_R)$ | |
| | $\vert$ | $\epsilon$ | $(1-\beta_{k,R})\kappa_R$ | |
| $\mathtt{REGION}_j$ | $\to$ | $\mathtt{INTER}_j$ | $p_N$ | |
| | $\vert$ | $\mathtt{FWDCDS}_j$ | $p_G/2$ | |
| | $\vert$ | $\mathtt{REVCDS}_j$ | $p_G/2$ | |
| | $\vert$ | $\mathtt{STRUCT}_j$ | $p_S$ | |
| | $\vert$ | $\mathtt{CONS}_j$ | $p_C$ | |
| INTER | $\to$ | $x_1$ INTER | $1-\kappa_N$ | $q_N(x_1)$ |
| | $\vert$ | $\epsilon$ | $\kappa_N$ | |

##### Coding sequences

| LHS | $\to$ | RHS | Transition | Emission |
|---|---|---|---|---|
| FWDCDS | $\to$ | FWDCOD FWDCDS | $1-\kappa_G$ | |
| | $\vert$ | $\epsilon$ | $\kappa_G$ | |
| FWDCOD | $\to$ | $x_1\ x_2\ x_3$ | $1-p_I$ | $q_G(xyz)$ |
| | $\vert$ | $x_1\ x_2\ x_3$ FWDINT | $p_I/3$ | $q_G(xyz)$ |
| | $\vert$ | $x_1\ x_2$ FWDINT $x_3$ | $p_I/3$ | $q_G(xyz)$ |
| | $\vert$ | $x_1$ FWDINT $x_2\ x_3$ | $p_I/3$ | $q_G(xyz)$ |
| FWDINT | $\to$ | $x_1\ x_2$ GENOME $x_3\ x_4$ | 1 | $q_{D1}(x_1)q_{D2}(x_2)q_{A1}(x_3)q_{A2}(x_4)$ |

| LHS | $\to$ | RHS | Transition | Emission |
|---|---|---|---|---|
| REVCDS | $\to$ | REVCDS REVCOD | $1-\kappa_G$ | |
| | $\vert$ | $\epsilon$ | $\kappa_G$ | |
| REVCOD | $\to$ | $x_1\ x_2\ x_3$ | $1-p_I$ | $q_G(\overline{xyz})$ |
| | $\vert$ | $x_1\ x_2\ x_3$ REVINT | $p_I/3$ | $q_G(\overline{xyz})$ |
| | $\vert$ | $x_1\ x_2$ REVINT $x_3$ | $p_I/3$ | $q_G(\overline{xyz})$ |
| | $\vert$ | $x_1$ REVINT $x_2\ x_3$ | $p_I/3$ | $q_G(\overline{xyz})$ |
| REVINT | $\to$ | $x_1\ x_2$ GENOME $x_3\ x_4$ | 1 | $q_{D1}(\overline{x_4})q_{D2}(\overline{x_3})q_{A1}(\overline{x_2})q_{A2}(\overline{x_1})$ |

$$\begin{aligned}
L(t) &\to A \cdot D(t,\, t) && \text{at weight } e^{-\mu t}, \\
L(t) &\to G_\tau \cdot D(t,\, \tau) && \text{at density } \mu e^{-\mu\tau}\, d\tau, \quad \tau \in (0,t), \\
D(t,\, T) &\to L(t-u) \cdot D(t,\, T) && \text{at density } \lambda\, du, \quad u \in (0,T), \\
D(t,\, T) &\to \varepsilon && \text{at weight } e^{-\lambda T}.
\end{aligned}$$

$$P(h_{cc'} = h \mid \mathrm{rest}) \;\propto\; \rho_h \cdot \mathcal{L}_{\mathrm{pair}}(\{c,c'\};\, H_h),$$

   where $\mathcal{L}_{\mathrm{pair}}$ is the path-DCA contribution from co-occurrences of classes $c$ and $c'$ across the corpus, evaluated at the current materialized atom $H_h$.

$$\tilde{T}_s \;=\; \sum_b \sum_x T_x^{(b,s)} \cdot \sum_{y \neq x} S_{xy}\, \pi^{(c_s)}(y)$$

is the $\pi$-weighted dwell-time integral. Under the Gamma$(a_\eta, b_\eta)$ prior the conjugate posterior is closed form

$$\eta_s \mid \text{path}, \pi^{(c_s)} \;\sim\; \text{Gamma}\big(a_\eta + N_s^{\text{acc}},\; b_\eta + \tilde{T}_s\big),$$

with $N_s^{\text{acc}} = \sum_b \sum_{x \neq y} N_{xy}^{(b,s)}$ the total observed substitution count, and the per-site marginal likelihood is closed-form Negative-Binomial.

**Laplace for $H_h$.** Each Potts atom is real-valued and the path-DCA likelihood is locally Gaussian in $H_h$ to second order. We approximate

$$\log p_{\mathrm{path-DCA}}(\mathrm{data} \mid H_h) \;\approx\; \log p_{\mathrm{path-DCA}}(\mathrm{data} \mid \hat{H}_h) \;-\; \tfrac{1}{2}(H_h - \hat{H}_h)^\top \Lambda_h (H_h - \hat{H}_h),$$

where $\hat{H}_h$ is the MAP under the current iterate, found by a few Newton or natural-gradient steps starting from a chosen seed, and $\Lambda_h = -\nabla^2_{H_h} \log p_{\mathrm{path-DCA}}$ is the Hessian at $\hat{H}_h$. Combined with the $\mathcal{N}(\mu_{kl}, \tau_{kl}^{-1})$ prior on each entry, the posterior is the closed-form Gaussian

$$H_h \mid \mathrm{data} \;\sim\; \mathcal{N}\big(\hat{H}_h^{\mathrm{post}},\, (\mathrm{diag}(\tau_{kl}) + \Lambda_h)^{-1}\big).$$

The Laplace estimate of the new-atom integral is the standard $\log p_{\mathrm{path-DCA}}(\hat{H}_h) + \log G_0^H(\hat{H}_h) + \frac{d}{2}\log(2\pi) - \frac{1}{2}\log\det(\mathrm{diag}(\tau_{kl}) + \Lambda_h)$ with $d = A(A+1)/2$ the symmetric-slice dimension.

###### The unified joint-marginal target

Let $\theta_{\text{indel}} = (\lambda, \mu, r)$ be the TKF92 indel parameters and $\alpha_z$ the Ewens partition concentration. Write Match$(A)$ for the Match cells of an alignment $A$, and let $E$ be a set of unordered Match-cell edges (a partition of Match$(A)$ into singletons of size 1 and doublets of size 2; we let $V(E) \subset$ Match$(A)$ denote the doublet endpoints). The TKF-DP joint marginal at branch length $t$ is

$$P(X,Y;t) \;=\; \sum_A \pi_{\text{TKF92}}(A \mid t, \theta_{\text{indel}}) \sum_E \pi_{\text{Ewens}}(E \mid \alpha_z, |\text{Match}(A)|) \prod_{(i,j)\in\text{Match}(A)\setminus V(E)} P_{\text{singlet}}(x_i, y_j; t) \prod_{e=((i,j),(k,l))\in E} P_{\text{do}}\text{[illegible]} \tag{D.3}$$

with the per-block primitives

$$P_{\text{singlet}}(a,b;t) \;=\; \sum_c \pi_c\, \pi^{(c)}(a)\; P_c(a\to b;\, t\cdot\eta), \tag{D.4}$$

$$P_{\text{doublet}}\big((a,c),(b,d);t\big) \;=\; \sum_{c_1,c_2} \pi_c(c_1)\,\pi_c(c_2)\; \pi_{\text{joint}}^{c_1,c_2}(a,c)\; P_{\text{joint}}^{c_1,c_2}\big((a,c)\to(b,d);\, t,\eta_1,\eta_2\big). \tag{D.5}$$

The doublet endpoint convention (matching the boost API of the released code) is that $(a,b)$ are the amino acids at the left endpoint $(i,j)$ and $(c,d)$ at the right endpoint $(k,l)$: $a=x_i$, $b=y_j$, $c=x_k$, $d=y_l$. Indices $c_1,c_2$ run over site classes and should not be confused with the endpoint-amino-acid index $c$.

The empirical class prior $\pi_c$ is the per-class column count from training (*not* uniform $1/K_c$); the per-class profile $\pi^{(c)}(\cdot)$ is the trained `pi_class`; the class-conditional transition is

$$P_c(a\to b;t) \;=\; [\exp(Q_c\cdot t)]_{a,b}, \qquad Q_c \;=\; (S_{\text{LG08}} - \operatorname{diag} S_{\text{LG08}})\cdot \pi^{(c)}[\,\cdot\,]^\top, \tag{D.6}$$

the standard F81-form GTR generator with exchangeability $S_{\text{LG08}}$ (32) and class-specific stationary $\pi^{(c)}$. The joint stationary $\pi_{\text{joint}}^{c_1,c_2}(a,c)$ at a coupled cell-pair is the joint stationary of a Potts atom $H = \texttt{atoms}[\texttt{assignments}[c_1,c_2]]$ at the trained checkpoint (the trained `potts_dp.atoms` indexed by `potts_dp.assignments`), and $P_{\text{joint}}^{c_1,c_2}$ is its time-evolved transition $\exp(Q_{\text{joint}}^{c_1,c_2}(H)\cdot t)$ built by $\texttt{build\_joint\_Q\_pair}(H,\pi_a,\pi_b,S,\eta_{\text{pair}}=(\eta_1,\eta_2))$ as described in §D.3.

The full conceptual ladder is therefore:

mean-field pre-correction (deprecated; rem. D.13) $\to$ bounded-edge aug-PHMM (§E.2)
$\to$ infinite Pair HMM (§E.3) $\to$ infinite gravestone-augmented pair SCFG (§E.3 + §D.5)
$\to$ infinite gravestone-augmented *phylogenetic* SCFG (§E.4),

each rung exact in the limit of the next, the entire ladder falling out of the simple stick-breaking-TKF92-with-Potts construction of Appendices D–E with no further mathematical machinery.
